# Neonatal Thyroxine Activation Modifies Epigenetic Programming of The Liver

**DOI:** 10.1101/2020.12.07.414938

**Authors:** Tatiana L. Fonseca, Tzintzuni Garcia, Gustavo W. Fernandes, T. Murlidharan Nair, Antonio C. Bianco

## Abstract

In the neonatal liver, a peak of type 2 deiodinase (D2) activity accelerates local T3 production and the expression of thyroid hormone (TH)-responsive genes. Here we show that this acute increase in T3 signaling permanently modifies hepatic gene expression. Liver-specific Dio2 inactivation (Alb-D2KO) transiently increased H3K9me3 levels during post-natal days 1-5 (P1-P5) in discrete chromatin areas, and methylation of 1,508 DNA sites (H-sites) that remained in the adult mouse liver. These sites were associated with 1,551 areas of reduced chromatin accessibility (RCA; Atac-seq) within core promoters and 2,426 within intergenic regions, with reduction in the expression of 1,525 genes (RNA-seq). There was strong correlation between H-sites and RCA sites (r=0.85; p<0.0002), suggesting a cause-effect relationship. The analysis of chromosome conformation capture (Hi-C) data revealed a set of 57 repressed genes that have a promoter RCA in close contact with an intergenic RCA ~300 Kbp apart, including Foxa2 that plays an important role during development. Thus, the post-natal surge in hepatic D2 activity and TH-signaling prevents discrete DNA methylation and modifies the transcriptome of the adult mouse. This explains how the systemic T3 hormone acts locally during development to define future chromatin accessibility and expression of critically relevant hepatic genes.

## Introduction

Thyroid hormone (TH) regulation of gene expression involves binding to nuclear receptors (TR) that are attached or may be directed to TH responsive elements (TREs). This is followed by recruitment of transcriptional coregulators that modify histones and chromatin accessibility to transcriptional enzymes (1); noncanonical pathways that do not require binding to TREs might also be involved (2). The genomic actions of TH signal promote localized transition of the chromatin from a tightly folded and less transcriptionally active structure known as heterochromatin to a more loosely folded and active structure known as euchromatin (3–6). As these processes involve histone modifications, regulation of gene expression by TH is largely fluid, with the rate of transcription of TH-regulated genes fluctuating rapidly according to the levels of the biologically active T3 in the cell nucleus.

In the absence of T3, most unoccupied TRs (uTR) remain attached to TREs, where they form complexes with transcriptional co-repressors to pack the DNA. Examples of uTR-recruited corepressors include nuclear receptor corepressor 1 (NCoR1) and the silencing mediator of retinoid and thyroid hormone receptors (SMRT; NCoR2), which recruit deacetylases (e.g. HDAC3), methyl transferases (e.g. HMT) and kinases (e.g.) to activate facultative heterochromatin formation and inhibit gene transcription (7, 8). Methylation of H3K9, reduction of H3R17 methylation and of phosphorylation/acetylation of H3S10/K14 are all examples of specific histone modifications triggered by uTRs that inhibit transcription (9).

uTRs are known to remain bound to TREs and have repressive effects on gene transcription. Nonetheless, exposure of cells to T3 directs additional TR units to TREs, and shifts TRs association from co-repressors to co-activators; these include steroid receptor coactivator (SRC) family, CBP/p300 or CARM1/SNF5, promoting transcriptional activity. In general, the chromatin modifications induced by T3-TR are the opposite of those caused by uTR, e.g. demethylation of H3K9, methylation of H3R17 and phosphorylation/acetylation of H3S10/K14 (9, 10). As a result, the T3-TR not only de-represses but also trans-activates transcription of target genes (1). T3-TR can also activate gene transcription via long-distance enhancer hyperacetylation, a mechanism that depends on the chromatin context (11).

Less is known about TH regulation of gene expression via DNA methylation, a process that can also result from histone methylation (12, 13), and can affect the expression of gene clusters as opposed to a single target gene (14). In general, the effects of DNA methylation on gene expression last longer when compared to histone modifications, and are involved in developmental transitions such as amphibian metamorphosis and mammalian organogenesis (15), both processes typically sensitive to TH (16, 17).

To regulate the timing and intensity of TH signaling on an organ/tissue-specific fashion, developing cells express TH deiodinases, that can both activate (D2) or inactivate (D3) TH. In the embryo, circulating T3 is kept at relatively low levels and tissues predominantly express D3. However, D3 activity diminishes towards birth at the same time that, in some tissues, D2 activity is selectively boosted. A unique blend of D3 and D2 activities seen during this transition independently controls the levels of nuclear T3 in each tissue, hence the timing and intensity of the TH signaling. In the mouse, this process may extend well into the post-natal period given that circulating T3 levels remain low (P1<<P10) through post-natal day 10 (P10) (18, 19). Indeed, peaks of D2-T3 that enhance TH signaling are seen through this period on a tissue-specific basis (3), e.g. embryonic day 17 (E17)-P1 in brown adipose tissue (BAT) (20), or P15 in the cochlea (21).

In the developing liver there is a D2-T3 peak (P1-P2) that briefly doubles T3 content (22) that coincides with the time of C/EBPa-induced maturation of the bi-potential hepatoblasts into hepatocytes (23); Dio2 expression in liver is silenced thereafter (22). This spike in TH signaling is critical for post-natal liver maturation given that liver-specific Dio2 inactivation (Alb-D2KO) was associated with the formation of 1,508 CpG sites of DNA hypermethylation (H-sites) of the adult Alb-D2KO liver genome (22). As a result, the adult Alb-D2KO liver response to high fat diet (HFD) is affected (22); the Alb-D2KO mice exhibit reduced susceptibility to obesity, liver steatosis, hyperlipidemia and alcoholic liver disease (22, 24).

Hepatocytes normally undergo postnatal epigenetic reprogramming, including changes in DNA methylation, which are the result of terminal differentiation of hepatocyte precursors (25, 26). For example, between E18.5 until adulthood, ~200,000 CpGs changed DNA methylation by more than 5%, and ~20,000 CpGs by more than 30%. These changes in DNA methylation occurred primarily in intergenic enhancer regions and coincided with the terminal differentiation of hepatoblasts into hepatocytes (25, 26). Here we investigated how D2-mediated TH signaling affect hepatocyte DNA methylation, DNA accessibility and gene expression, explaining the development effects of TH.

## Results

Two mechanisms could explain the formation of the 1,508 H-sites found in the adult Alb-D2KO liver: (i) defective neonatal DNA demethylation and/or (ii) *de novo* neonatal DNA methylation, around P1-P5 but before P10, when circulating T3 reached adult levels. To resolve this, we studied the coordinates of the H-sites (Tables S1) *vs* those of thousands of hepatic DNA sites that are normally demethylated during the P1-P21 time frame (25). Only 25 H-sites were among those that are normally demethylated (Fig. S1A), indicating that almost all H-sites in the Alb-D2KO liver resulted from *de novo* DNA methylation.

### How were the H-sites formed in the Alb-D2KO liver?

Liver-specific Dio2 inactivation reduces T3 signaling in the P1-P5 liver (22). A predominance of unoccupied thyroid hormone receptors (uTR) has been shown to foster a histone environment that has been linked to *de novo* DNA methylation in a number of developmental settings by recruiting co-repressors that triple-methylate H3K9 (9, 12, 13, 16). Thus, we tested if this could explain the H-sites in the liver of Alb-D2KO by performing a liver ChIP-seq of H3K9me3 in P1, P5 and adult Alb-D2KO mice. Indeed, we found at least 86 H-sites in the Alb-D2KO liver transiently imbedded in H3K9me3-enriched areas that were only present in the Alb-D2KO chromatin (Table S1); these sites are hereinafter referred to as HH-sites (Fig. 1A). Sixty-six HH-sites were identified in P1 and 51 in P5; in some cases, H-sites remained surrounded by areas enriched with H3K9me3 throughout the P1 and P5 period. In the adult mice, the differences in H3K9me3 enrichment level between control and Alb-D2KO mostly dissipated, remaining only around 16 of the 86 HH-sites. This dynamic is illustrated in the 20 HH-sites identified on the qA1.2 segment of Alb-D2KO chromosome 12 (Fig. 1A). Within this segment, there was an ~80 Kbp island of intense enrichment of H3k9me3 on P1 and P5; the differences in H3K9me3 dissipated in the adult mice; a smaller 14Kbp island with 5 HH-sites that behave similarly was observed 2 Mbp upstream (Table S1).

**Fig. 1.**
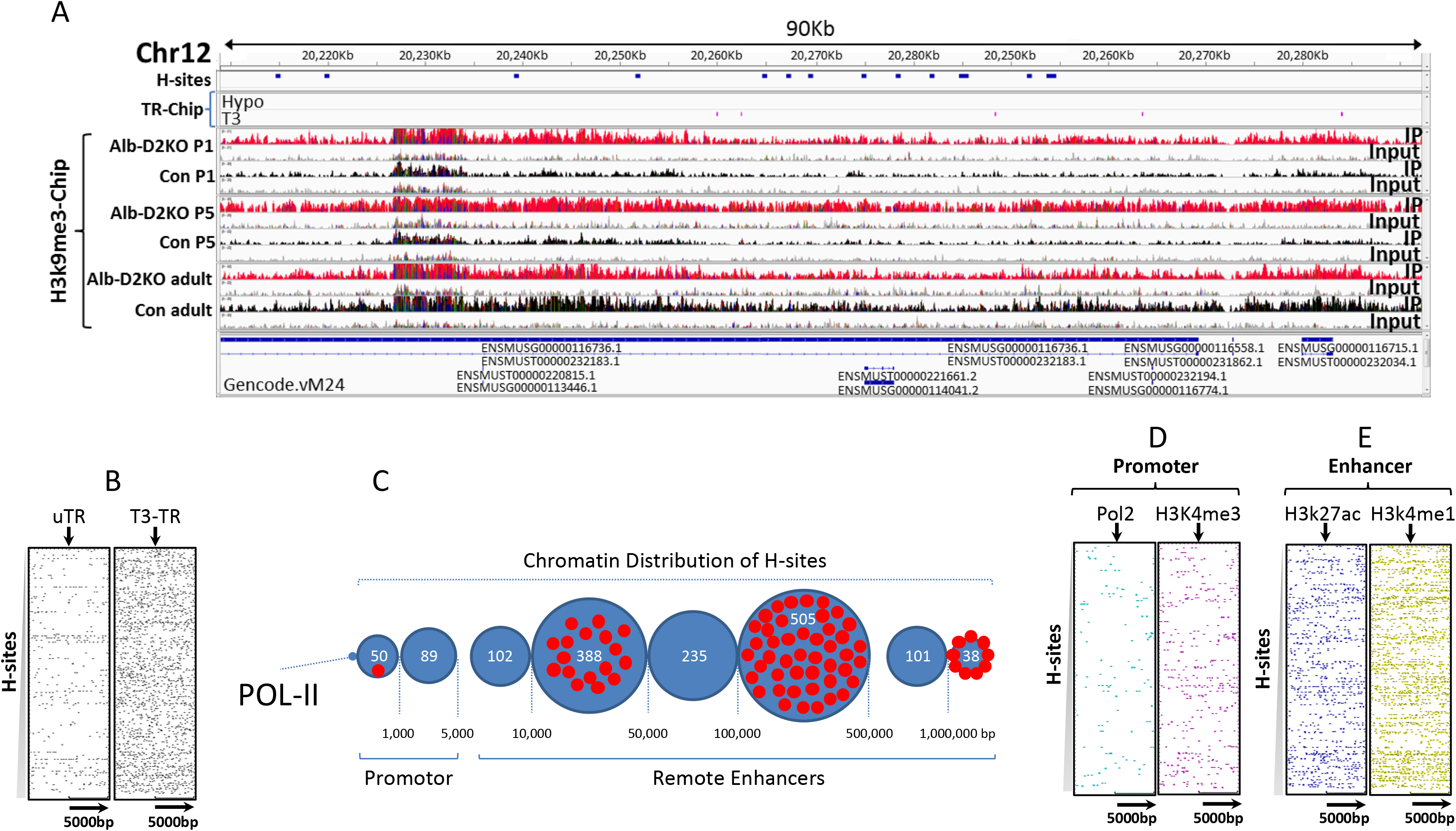
DNA hyper-methylation sites (H-sites) in the ALB-D2KO mouse liver chromatin. **(A)** browser IGV image of the indicated portion of the mouse chromosome 12 containing 15 H-sites; the following tracks contain ChIP signals for: (top to bottom) H-sites (blue), TR ChIP-seq peaks (hypo and T3-treated - pink), H3K9me3 ChIP-seq peaks in Alb-D2KO (red) and control (black) P1, P5 and adult mice; the input track is shown in gray; **(B)** heat maps of H-sites vs TR ChIP-seq peaks in liver; the X axis contains the TR peak coordinates at the center +/− 5 Kbp and the Y axis contains the H-site coordinates; the H-site coordinates start at the top and progresses downwards in the following order: Chr 1 through Chr 9, Chr X, Chr 10 through Chr 19; each dot in the heat map indicates an H-site and its relative distance to a TR peak (center); Hypo indicates TR ChIP peak coordinates obtained from hypothyroid mice, and +T3 indicates TR coordinates obtained from T3-treated mice; **(C)** distribution of H-sites as a function of polymerase II (pol-II) ChIP-seq peak locations; the numbers at the bottom indicate the distance in base pairs (bp); H-sites are within the *promoter* if up to 1 Kbp; the H-sites are grouped in blue circles according to the distance bracket; the numbers of H-sites in each bracket is indicated inside the blue circles, in white; red dots are H-sites imbedded in areas enriched with H3K9me3 during P1-P5; **(D)** same as in **B**, except that Pol-II and H3K4me3 (core promoter makers) ChIP-seq peaks were placed at the center of each respective heat map; **(E)** same as in **B**, except that H3K27ac and H3K4me1 (enhancer makers) ChIP-seq peaks were placed at the center of each respective heat map.

These results support the idea that during the first few days of life, discrete areas of the Alb-D2KO liver chromatin are enriched with H3K9me3, reflecting an environment that favors the methylation of neighboring DNA sites (H-sites). Here, we captured two snap shots, i.e. P1 and P5, of a process that could last until circulating T3 reached adult levels, i.e. around P10 (18, 19). The transient nature of the H3K9me3 pockets and the durability of the H-sites could explain why we only observed their association in 86 cases, given that we only looked at two neonatal time-points.

That liver D2 is only expressed during the first few days of life and the H-sites are only present in the Alb-D2KO livers constitute evidence that formation of H-sites is inherently connected with reduced T3-signaling (22). Thus, we next wished to define the relationship between H-sites and TRs in the liver. We reanalyzed two sets of published liver ChIP-seq data obtained with αTR – one from hypothyroid mice, in which uTRs predominate over T3-TRs, and the other from T3-treated mice, in which T3-TRs predominate over uTRs, (27). We used these data sets to assess the density of uTRs or T3-TRs in the proximity of H-site coordinates. The resulting heatmaps revealed that while only ~10% of the H-sites were flanked by uTRs in the hypothyroid liver (Fig. 1B), >90% of the H-sites were flanked by T3-TR; T3 has been shown to recruit a larger number of TR units to the TH responsive elements (TRE) (Fig. 1B).

Further assessment of the uTRs and T3-TRs revealed that the sample of 86 HH-sites we identified on P1 and P5 was only flanked by 7 uTRs (within 10 Kbp). In contrast, the same sample of HH-sites was flanked by T3-TRs in 69 occasions (Table S1). For example, no uTRs were identified on the qA1.2 segment of Alb-D2KO chromosome 12, or the smaller upstream chromatin area. In contrast, both areas combined presented 9 T3-TRs (Table S1). It is thus conceivable that uTRs might not play a decisive role in triggering H-site formation in this setting. Instead, it is more likely that the neonatal D2-mediated enrichment of the liver chromatin with T3-TR sustains recruitment of co-activators and preserves an environment unsuitable for the discrete methylation of H-sites, illustrated by the lower levels of H3K9me3 (Fig. 1A).

### Location of the H-sites in the Alb-D2KO liver chromatin

To refine the localization of H-sites in liver chromatin, we first annotated the H-sites coordinates and found widespread distribution throughout the genome (Table S2). We next reanalyzed published hepatocyte chromatin immunoprecipitation-sequencing (ChIP-seq) data of multiple chromatin markers (28) and contrasted with the H-sites coordinates. Proximity analysis revealed that only few H-sites were located within the core promoter marker Pol-II ChIP peaks; the majority of the H-sites was found between 5-500Kbp of the Pol-II peaks (Fig. 1C). This distant relationship with core promoters is illustrated in heatmaps in which the coordinates for Pol-II ChIP peaks and an additional core promoter marker H3K4me3 were plotted against the H-sites coordinates (Fig.1D)(25, 29). The low H-site density in the vicinity of these chromatin markers supports that most are not located in core promoter regions. That most H-sites are likely to be located in remote enhancer regions is suggested by their high density and proximity around the known primed and active enhancer markers, respectively H3K4me1 and H3K27ac (Fig. 1E) (25, 29).

### Reduced Chromatin Accessibility in the adult Alb-D2KO liver chromatin

Tissue- or cell type specific DNA hypermethylation is frequently associated with areas of stable heterochromatin, i.e. reduced chromatin accessibility, and has been implicated in the control of gene expression during development (30). To test if this was case in the Alb-D2KO liver, we used the transposase-accessible chromatin assay followed by high-throughput sequencing (ATAC-seq) of both adult control and Alb-D2KO mouse liver nuclei (Fig. 2A). To generate the ATAC-seq libraries we used high-quality nuclei extracted from frozen adult liver that were quality checked by light microscopy (Fig. 2A). The ATAC-seq libraries yielded the expected fragment length distribution, with the majority of fragments representing areas of inter-nucleosome open chromatin, with progressively fewer large sized fragments representing mono and oligo-nucleosomes (Fig. 2A).

**Fig. 2.**
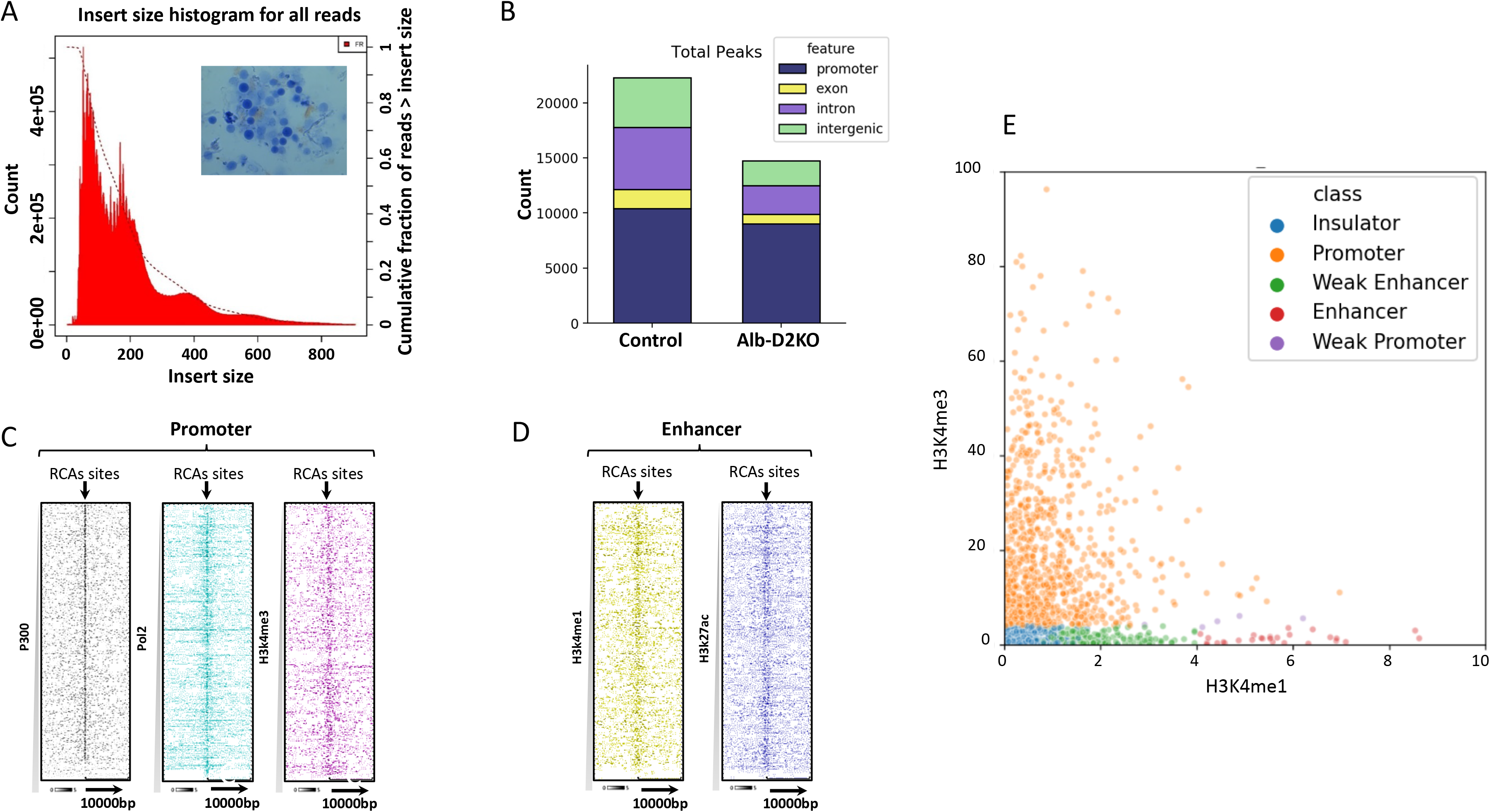
Reduced chromatin accessibility (RCA) in the ALB-D2KO mouse liver nuclei. **(A)** the inset contains a representative image of isolated liver nuclei (40X) before adding the transposase enzyme; the graph indicates the lengths of the fragments sequenced within a representative ATAC-seq library; insert size in bp; the first sharp peak (~100 bp) reflects areas of open chromatin, which is followed by 4 peaks of mono (~200 bp) or multinucleosomes of compact chromatin; **(B)** genomic distribution of peaks obtained in ATAC-Seq data; **(C)** heat maps prepared as in Fig. 1B, except that the X axis contains the RCA peak coordinates at the center +/− 10,000 bp and the Y axis contains the coordinates for P300, Pol2 and H3k4me3; **(D)** same as **C**, except that the Y axis contains the coordinates for H3k4me1 and H3k27ac; **(E)** classification of RCAs according to the relative intensities of H3K4me1 and H3K4me3 ChIP-seq peaks: promoter-RCAs, weak promoter-RCAs, enhancer-RCAs, weak enhancer-RCAs, and insulator-RCAs; each dot represents one RCA area.

A total of 22,374 areas of open chromatin were identified in the liver of control animals; after gene annotation, almost half of these open areas were placed in promoter regions up to 200 bp of transcription start site (TSS; Fig. 2B). The Alb-D2KO liver exhibited a smaller number of open chromatin areas, only 14,343, indicating the existence of 8,031 regions with reduced chromatin accessibility (RCAs). Of these RCA areas, 1,551 were annotated to core promoters and 2,426 to intergenic regions (Fig. 2B). Plotting the RCA coordinates against Pol-II, P300 and H3K4me3 ChIP-seq peaks, revealed high density with variable degrees of superimposition, confirming their proximity to promoter regions (Fig. 2C). At the same time, high density and variable degrees of superimposition features were also observed when RCA areas were plotted against H3k4me1 and H3k27ac ChIP-seq peak coordinates, confirming that they were also present in remote enhancer areas (Fig. 2D).

We next focused on the promoter RCA (p-RCA) areas, and rank them according to their ATAC-seq signal intensity and signal ratio to two typical chromatin markers H3K4me1 (enhancer) and H3K4me3 (promoter) (Fig. 2E) (29, 31). This approach revealed that 65% of these areas did function as typical core promoters, while 22% were likely to be insulators and 13% local enhancers (Fig. 2E). Of course, these are all functions that can be affected by chromatin accessibility, which is strongly coupled to binding of transcription factors; in most cases binding only occurs when the chromatin is accessible, unpacked (32). Thus, to identify the transcription factors that were affected by the p-RCA areas in the Alb-D2KO chromatin, we performed *de novo* motif profiling of these areas using the MEME suite. Motifs belonging to major families of transcription factors were identified, including Sp1-3, E2f3, Klf1, Nrf1, Elk1, and Egr2, many of which are known for playing critical roles in liver development and function (Fig. 3) (33–38). In addition, the second and the third most enriched motifs were for ZNF143 and CCCTC-binding factor (CTCF) (Fig. 3). Both are transcription factors that cooperate with the multiprotein complex cohesin to promote chromatin folding, allowing remote enhancers to interfere with transcriptional activity (39).

**Fig. 3.**
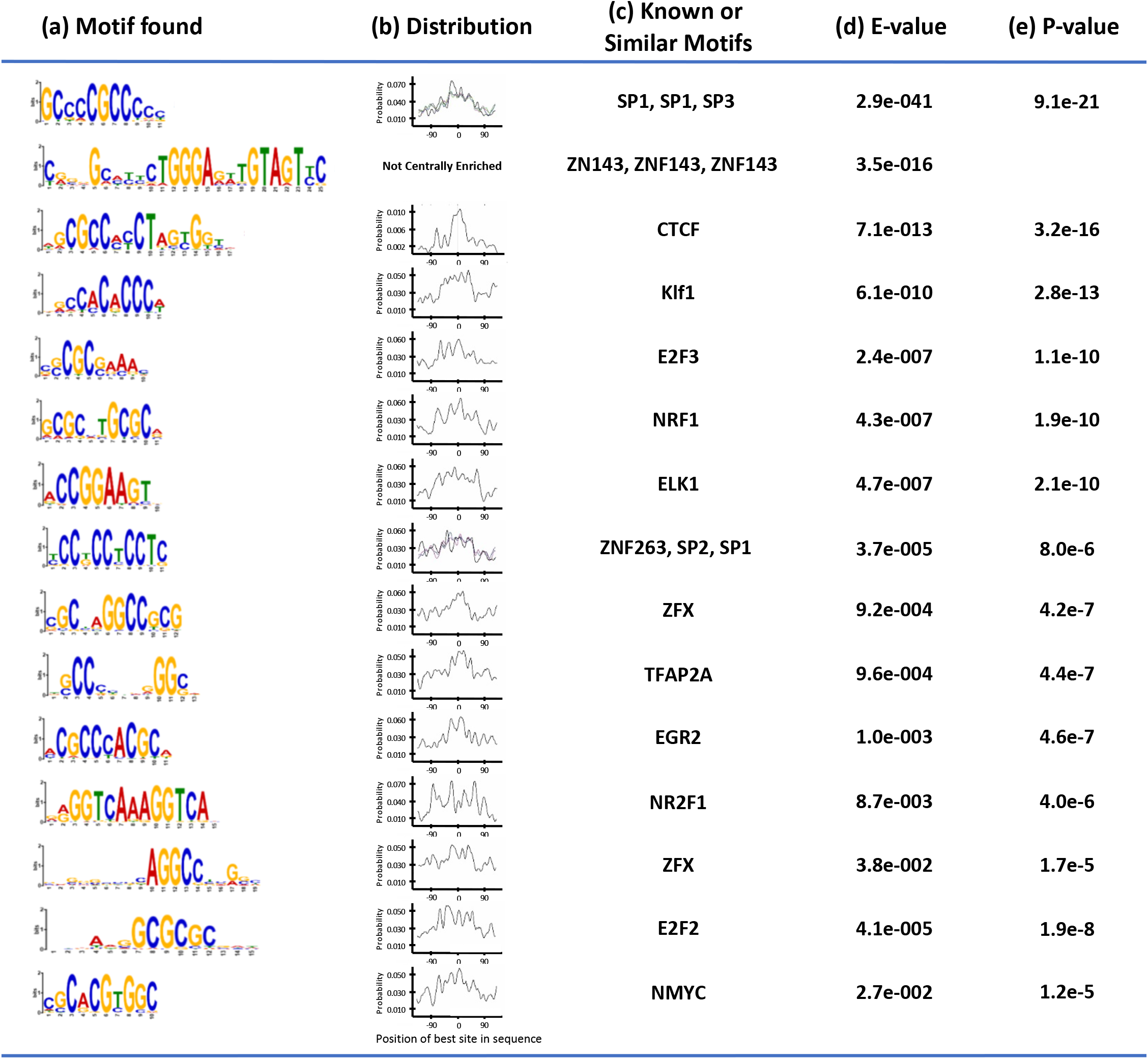
Transcription factors binding motifs impoverished in the RCA areas in Alb-D2KO liver chromatin as identified through the MEME suite: **(A)** the logo of each motif; the relative size of the letters indicates their frequency in the sequence; the total height of the letters depicts the information content of the position in bits; **(B)** the distribution of the best matches to the motif in the sequences as found by a CentriMo analysis; **(C)** the most similar motifs reported by the motif discovery programs (MEME or CentriMo) compared with known motifs in a motif database; **(D)** E-value is the adjusted p-value multiplied by the number of motifs in the input files(s) and **(E)** p-value that is the statistical significance of the motif enrichment adjusted for multiple tests; the enrichment p-value of a motif is calculated by using the one-tailed binomial test on the number of sequences with a match to the motif (“Sequence Matches”) that have their best match in the reported region (“Region Matches”), corrected for the number of regions and score thresholds tested (“Multiple Tests”). The test assumes that the probability that the best match in a sequence falls in the region is the region width divided by the number of places a motif can align in the sequence (sequence length minus motif width plus 1).

### Chromatin Accessibility and Gene expression

To study gene expression in the Alb-D2KO liver and test whether it was affected by p-RCA areas, we next performed an RNA-seq analysis of adult mouse liver (Fig. S1B) and identified 1,525 genes differentially expressed in Alb-D2KO mouse (>1.2-fold; p<0.05; Fig. 4A, S1C; Table S3). Notably, most of these genes (1,363) were downregulated in the Alb-D2KO liver, reflecting the overall reduction in chromatin accessibility. The gene set enrichment analysis (GSEA, Partek Flow) of these differentially expressed genes revealed that the top two genesets impoverished in the Alb-D2KO liver were (i) “fatty acid” and (ii) “lipid metabolic process” (ES>31 and p<2.9E-14) (Table S4). In addition, a pathway enrichment analysis (PEA, Partek Flow) identified metabolic pathways related to drugs and other xenobiotic compounds, as well as five lipid-related pathways among the top 15 pathways impoverished in the Alb-D2KO liver (ES >5.2 and p<0.01) (Table S5).

**Fig. 4.**
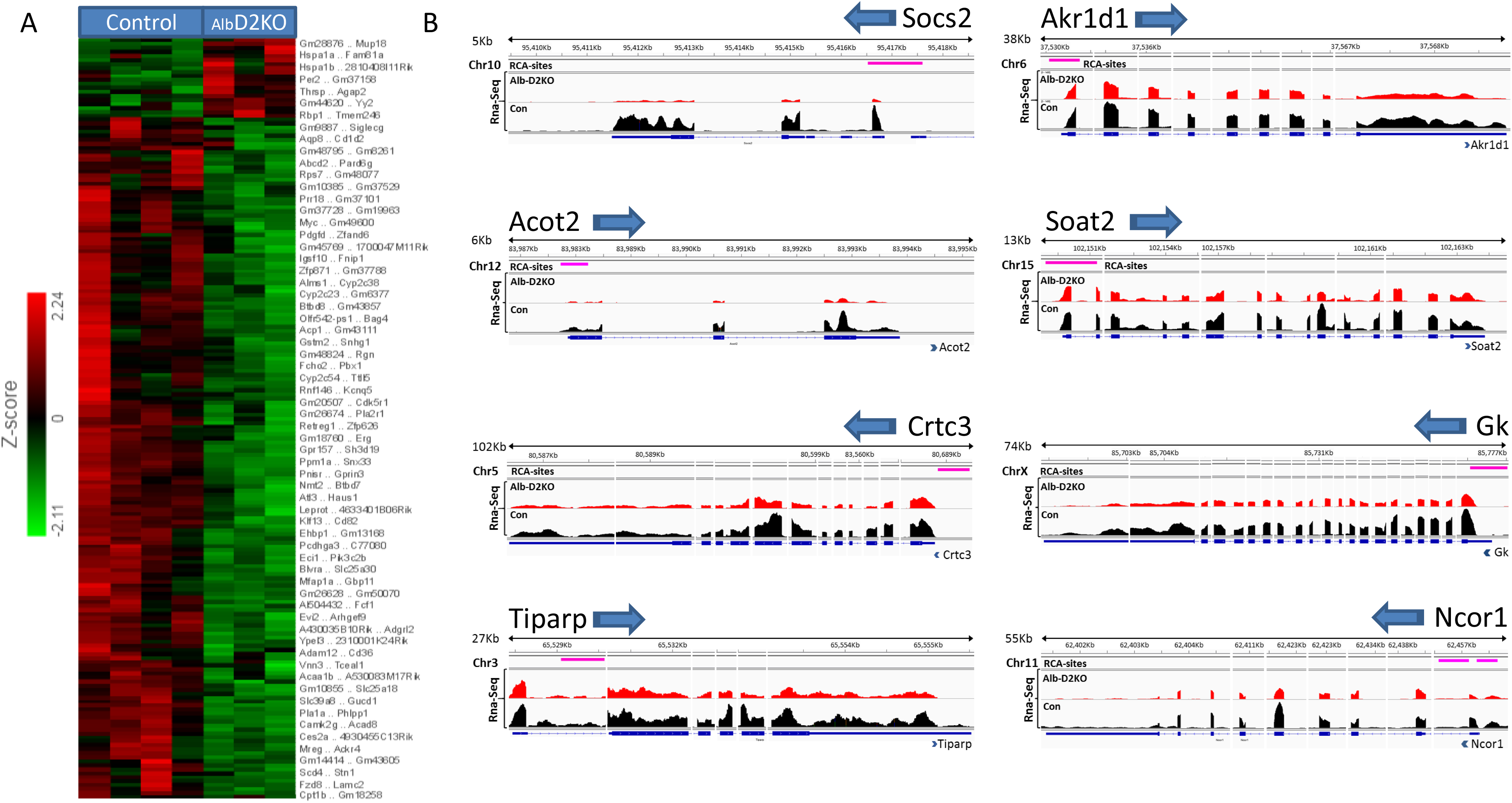
Gene expression in the ALB-D2KO mouse liver. **(A)** heatmap of liver RNA-seq data showing clustering of genes affected in the ALB-D2KO mouse; green indicates relative down-regulation; red indicates relative upregulation; fold change is shown on the left; most gene names are indicated on the right; **(B)** browser IGV display of transcript areas of selected genes (8 representative genes out of a total of 146 genes) that are downregulated in the Alb-D2KO liver <u>AND</u> exhibit a RCA area within its core promoter region; the distance between the negative RNA-seq peak and the RCA area in all cases is <5 Kbp; the gene names are indicated on the top of each panel along with the arrow indicating direction of transcription; the red tracks reflects the coverage for the Alb-D2KO RNA-seq data, and the black tracks reflects the controls; the RCA areas are indicated by a pink line.

In order to identify genes in which there was a high likelihood that a p-RCA was affecting its expression directly, we next analyzed the proximity of the 1,551 p-RCAs and the 1,363 genes with at least one negative RNA-seq peak (n-RNA-seq), selecting those that were within 5 Kbp of each other (*p-RCA:n-RNA-seq*), a configuration that strongly supports a functional relationship between them. This strict criterion led us to 154 p-RCAs (146 genes; Table S6), with key roles in liver development and function, including lipid metabolism, mitochondria, redox control, drug metabolism, TGFB-signaling and fibrosis, and cell signaling. Typical examples of these *p-RCA:n-RNA-seq* areas and the corresponding gene coverages are shown in Fig. 4B.

An analysis of these 154 p-RCA areas revealed that about ¾ contained footprints of missing transcription factors, including SP1-3 (n=63), E2F3 (n=32), Klf1 (n=37) and NRF1 (n=34) (Table S6). These footprints were rarely alone; most of the time they were present in groups of 3-4 on the same p-RCA. At the same time, about ¼ of the p-RCAs was found in insulators. The presence of both cohesin and CTCF (CAC-sites) is typical of insulators that modulate the 3D chromatin structure and gene expression by affecting the connection with remote enhancers. In contrast, insulators that contain cohesin only (CNC-sites) have a local effect, isolating functional promoters and enhancers from spreading neighboring heterochromatin (40). Using published mouse liver ChIP-seq data (31) for cohesin (Rad21) and CTCF, we looked for overlaps between the 34 RCA/insulator areas and the cohesin/CTCF peaks. There were 30-CAC sites and 4 CNC-sites, highlighting the potential importance of remote regulation in our model (Table S7).

### Functional interplay between H-sites, RCA areas and gene expression

There are multiple pathways through which H-sites could affect gene expression, mainly silencing active promoters or enhancers via creating a p-RCA, or controlling higher-order chromatin structure and the action of long-distance enhancers (30). Indeed, the analysis of the density of 1,508 H-sites and of all 8,052 RCA areas in individual chromosomes revealed a strong positive correlation (r=0.85; p<0.0002). The correlation remained strong when only the densities of H-sites *vs* p-RCA areas (r=0.69; p<0.0009; Fig. 5A) or intergenic RCA (i-RCA) areas (r=0.75; p<0.0002; Fig. 5B) were considered (Table S8). These findings support our hypothesis of a close, possibly cause-effect relationship, between H-sites and RCA areas.

**Fig. 5.**
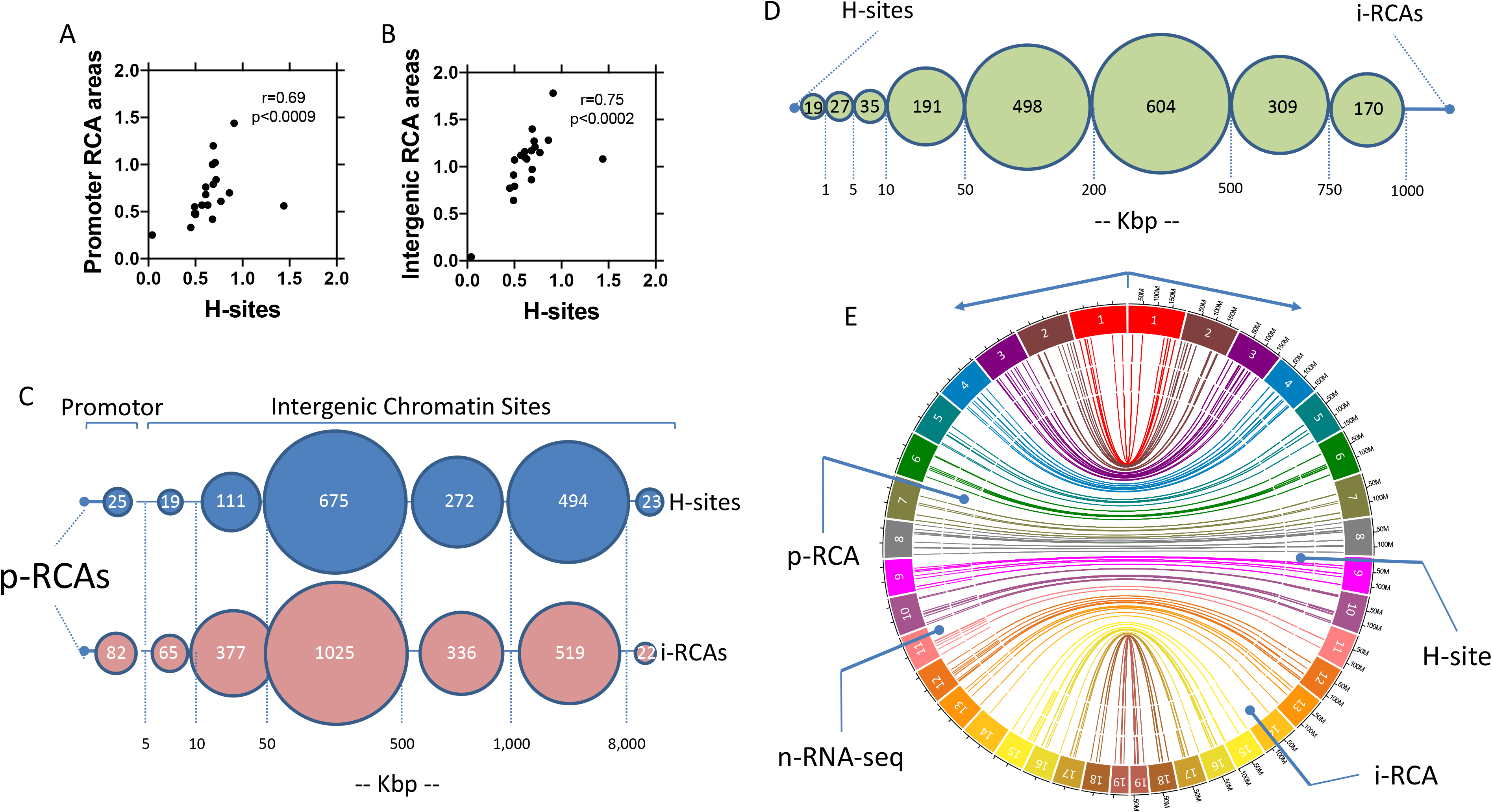
Interrelationship between H-sites and RCA areas in the ALB-D2KO mouse liver. **(A)** correlation between the density (number of H-sites/size of the chromosome) of H-sites and promoter RCAs areas in each chromosome; the Spearman r and the statistical significance are indicated; **(B)** same as **A**, except that i-RCA areas were used; **(C)** distance between H-sites and i-RCA areas vs. p-RCA areas; the number of H-sites and i-RCA areas at each distance bracket is indicated inside the circles; distances are indicated in Kbp; **(D)** same as **C** except that distances were calculated between intergenic RCA areas (i-RCA) vs. H-sites; there are 572 i-RCA located at a distance >1,000 Kbp that are not indicated; **(E)** Circa representation of the 121 n-RNA-seq areas and p-RCA areas within 1,000 bp of *area-1 vs* the closest H-sites and i-RCA to *area-2* areas across the Alb-D2KO liver genome; the outer most ring represents chromosome ideograms; on the right, the size of each chromosome is indicated in Mbp; the inner connecting lines indicate the Hi-C interacting points obtained from (42).

A more granular analysis of the H-sites coordinates revealed that of all H-sites, only 25 were within 5 Kbp of a p-RCA area (Fig. 5C). Notably, we found that 12 of the 154 *p-RCA:n-RNA-seq* areas did contain at least one H-site within 5 Kbp (10 within 1 Kbp), setting the stage for a possible local negative effect on the promoter of the following genes: 1700047M11Rik, Gstm6, 3110082I17Rik, Mtus1, Dixdc1, Eif5, Dsp, Pde4d, Sco2, Tymp, Odf3b, Cdo1 (Table S6). In these locations, the p-RCA areas included mostly core promoters, but also 2 enhancers and 1 insulator (Table S6).

Most H-sites were much further away from the p-RCA areas. About half of the H-sites was found within 500 Kbp of p-RCAs and 2/3 up to 1 Mbp away (Fig. 5C). In these cases, any effect on gene expression would need to be remote, probably involving intra-topologically associating domain (TAD) chromatin interaction, which is in keeping with the CAC sites (Table S7) and CTCF/ZNF143 footprints detected in p-RCA areas (Fig. 3). In the specific case of the remaining 134 *p-RCA:n-RNA-seq* areas, the nearest H-site was found within 50 Kbp of 18 p-RCA sites, between 50 and 500 Kbp of 62 p-RCA sites, and between 500 Kbp and 1 Mbp of 19 p-RCA sites; the remaining 35 p-RCAs were much further away from an H-site (Fig. 5C).

Whereas it is logical to assume that the distant H-sites *per se* could affect chromatin structure and folding (30), it is also feasible that the disruption of the intergenic enhancers resulted from the combined effects of both H-sites and i-RCAs. Furthermore, H-sites could have played a role in the formation and maintenance of the i-RCA areas, disrupting proper recruitment of proteins to, and the folding of, chromatin (40). Indeed, an analysis of the distance between p-RCAs and H-sites or p-RCAs and i-RCAs shows a remarkably similar profile (Fig. 5C), strengthening the latter possibility. Few H-sites and i-RCAs were near p-RCA areas; most were found between 50-1,000 Kbp (Fig. 5C). Despite very similar chromatin distribution, there is distance between H-sites and i-RCA areas (Fig. 5D), that nonetheless is compatible with them being within the same TAD (0.2-1 Mbp); DNA sequences within a TAD are known to physically interact with each other more frequently than with sequences outside the TAD (41). This was not unexpected, considering the dimension of the intergenic environment and the high-order chromatin structure (40).

We next used chromosome conformation data (Hi-C) obtained from primary liver cell cultures (42) to identify potential chromatin contacting areas that could explain how disruption of a distant enhancer would have affected the 154 *p-RCA:n-RNA-seq* sites and their genes. This was done by first building a contact matrix of all statistically significant interacting pairs separated by a minimum distance of 1 Kbp and a maximum distance of 10 Mbp (*area-1* x *area-2*) for each chromosome. This matrix was analyzed by first mining for coordinates that were within 1 Kbp of the 154 p-RCA sites. We found 528 coordinates and 121 p-RCAs that met this strict criterion; these matrix coordinates were arbitrarily named *areas-1*. The remainder p-RCAs included 4 genes that contained an H-site near the core promoter, i.e. Sco2, Tymp, Odf3b, Gstm6. Second, we further mined the matrix and verified that these 528 *areas-1* were in contact with 572 distant areas, given that a few *areas-1* exhibited more than one point of contact; we arbitrarily named these distant points of contact *areas-2*. The average distance between *areas-1* and *areas-2* was ~316 Kbp (1 Kbp – 2.4 Mbp), well within the constraints of a TAD (Fig. 6).

**Fig. 6.**
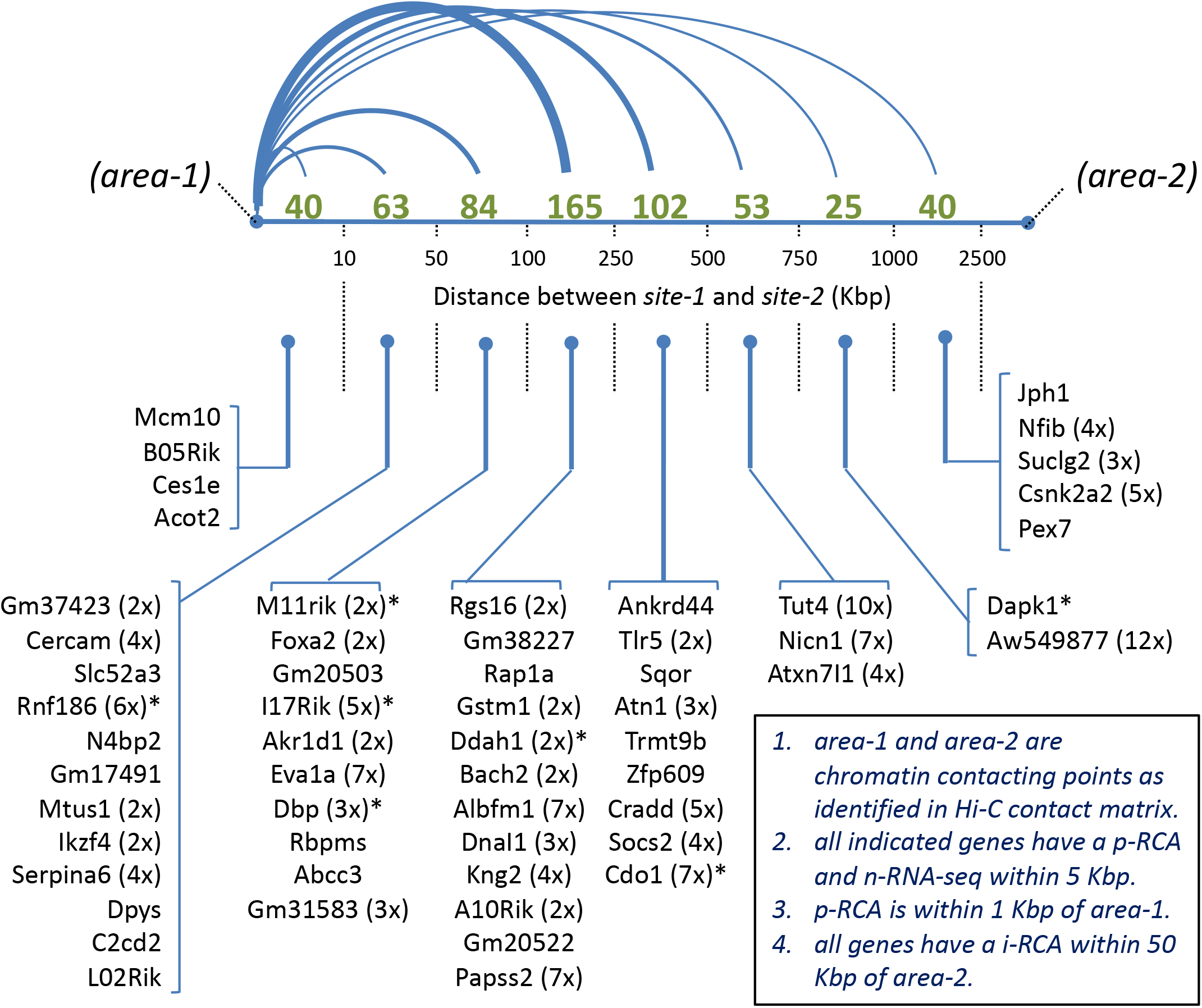
Distribution of Hi-C interacting points (*area-1* and *area-2*) located nearby p-RCA areas (*area-1*) and i-RCA areas (*area-2*); only area-1 within 1 Kbp of a p-RCA are shown; the number of *area-2* contacting points at each distance bracket is indicated in green and by the relative thickness of the connecting lines; the 57 gene names shown reflect those genes that satisfy the following criteria: (a) having a n-RNA-seq peak within 5 Kbp of a p-RCA; (b) having a p-RCA area within 1 Kbp of an *area-1*; (c) having an i-RCA area within 50 Kbp of an *area-2*; in parenthesis after each gene name is the number of interacting *areas-2*; *indicates the presence of a H-site within 50 Kbp of *area 2*. Note that the following genes that were originally included among the 146 are not shown here because they did not meetthe strict criterion of having a p-RCA within 1,000 bp of a Hi-C *area-1*: Gstt3, Gstt1, Ncor-1, Rmf167, Cdk5R1, Afmid, Sav1, Zbtb1, Gm34667, Ighm, Prkaa1, Gm49413, M09Rik, Sco2, Tymp, Odf3b, Soat2, Dnmdp, Echdc3, Gstm6, Adgrl2, Lurap1l, Gbp11, Smurf1, J07Rik, Bc024386, Mtus1, O19Rik, Pixbc1, Traip and Acaa1b.

To test whether p-RCAs could be directly affected by H-sites via high-order chromatin structure, we circa plotted for each chromosome the 1,551 p-RCA vs the 1,508 H-sites, and overlaid the 572 points of contact provided by the Hi-C analysis (Fig. S2-S6). While there was always one or more H-sites nearby an *area-2*, the smallest distance between them averaged ~1 Mbp (Table S9). Subsequently, we applied the same rationale for the i-RCAs and verified that they were in general much closer to the 572 *areas-2*, with the smallest distance between them averaging ~0.55 Mbp (Table S9). In addition, in many cases an *area-2* was in the vicinity of more than one i-RCA. For example, an analysis of a sample of all *areas-2* within <50 Kbp of an i-RCA, identified 164 instances in which an *area-2* was surrounded by up to 12 i-RCA areas, at an average distance of ~25 Kbp (Table S9); as a comparison, there were 65 H-sites located at <100 Kbp of an *area-2* (Table S9). This remarkable relative proximity between *areas-1* and *p-RCAs:n-RNA-seq* vs. *areas-2* and i-RCAs & H-sites, as revealed by the Hi-C contact matrix, is illustrated by plotting these elements in a single circa-plot with all chromosomes (Fig. 5E). Of the original 146 genes with a *p-RCA-n-RNA-seq* area, 116 genes were found within 1 Kbp of *area-1*, of which 57 exhibited one or more i-RCAs within 50 Kbp of an *area-2*, 8 of which also contained an H-site within 50 Kbp (Fig. 6).

## Discussion

The investigation of a dramatic phenotype seen in the Alb-D2KO mouse (22, 24) led to the discovery that neonatal inactivation of liver Dio2 reduced local TH-signaling and set in motion a series of events that reduced chromatin accessibility and future expression of key hepatic genes. The present data suggest that a developmental hepatic peak of D2 locally mitigates the typically low post-natal circulating T3 levels of this period of life, which then prevents creation of a nuclear environment - enrichment of discrete chromatin areas with H3K9me3 - that favors the formation of ~1,500 sites of *de novo* DNA hypermethylation (H-sites). The H-sites were mostly distant from core promoters, and distributed similarly to the ~1,550 p-RCA and ~2,400 i-RCA areas, suggesting they occupy the same or neighboring TADs. These elements are likely to have disrupted the function of long-distance enhancers, which then failed to interact and activate the core promoter areas, lowering the expression of ~1,500 genes involved in different aspects of liver development and function.

The present findings indicate that localized D2-generated T3 plays a role in the terminal maturation of the hepatocytes. This is reminiscent of the role played by D2 in the post-natal development of brown adipocytes and cochlea (20, 43). The timing and intensity of T3 signaling in developing cells is regulated via expression of deiodinases. In the embryo, circulating T3 is kept at relatively low levels and tissues predominantly express the inactivating deiodinase, i.e. D3, limiting exposure to T3. However, D3 activity diminishes towards birth at the same time that, in some tissues, D2 activity is selectively boosted. A unique blend of D3 and D2 activities seen during this transition independently controls the levels of nuclear T3 in each tissue, hence the timing and intensity of the thyroid hormone signaling.

In the case of hepatocytes, the short-lived, localized D2-T3 production, enriches discrete areas of the chromatin with T3-TRs, preventing accumulation of H3K9me3 and the formation of the H-sites. Our H3K9me3 ChIP-seq studies in P1 and P5 Alb-D2KO livers caught the moments during which the chromatin environment favored formation of H-sites, illustrated by the islands of H3K9me3 surrounding H-site coordinates; different timings could have been involved in the formation of the other H-sites, up until P10, when circulating T3 reached adult levels. Insulin signaling has been shown to play a similar role in the post-natal epigenetic programing of the liver, including differential DNA methylation. For example, changes in the DNA demethylation during the neonatal period were found to be essential for the ligand-activated PPARα-dependent gene regulates the hepatic fatty acid β-oxidation (44). At the same time, changes in the DNA methylation coincide with the hepatocyte terminal differentiation and occurs after hematopoietic stem cell migration (26).

uTRs have been shown to increase H3K9 methylation through recruitment of SUV39H1 (9) and HP1 proteins, a family of heterochromatic adaptor molecules implicated in both gene silencing and supra-nucleosomal chromatin structure (45). Indeed, the SUV39H1 was also found to cooperate with DNMTs to establish sites of *de novo* methylation (46), which could explain the formation of the H-sites found in ALB-D2KO liver. Nonetheless, our finding that few uTRs are nearby H-sites suggests that DNA hypermethylation in these areas is probably not caused by uTR-recruitment of co-repressors. Formation of the H-sites is rather the result of the hepatocyte differentiation program in the absence of a neonatal surge in T3 signaling, which is prevented by the peak of D2 activity and increased levels of T3-TR.

The reduction in gene expression found in the Alb-D2KO liver is dramatic. There are probably multiple mechanisms that explain such changes, but the ones we found evidence for in the present investigation include (i) disruption of few p-RCA function by local H-sites, (ii) disruption of long-distant chromatin interactions, looping enhancers and promoters, (iii) or a combination of both. The relative proximity and similar distribution of H-sites, i-RCAs and p-RCAs within individual chromosomes is remarkable; it is an indication that most of these elements exist inside the same or neighboring TAD unit, and probably exhibit a functional relationship. These concepts were illustrated in the finding of 57 genes that have a *p-RCA:n-RNA-seq* area along with likely long-distance i-RCA/H-site interacting chromatin areas. It is conceivable that H-sites initiate these processes by creating/maintaining i-RCAs and disrupting the function of the distant enhancers, subsequently depriving core promoters of transcriptional factors that activate gene transcription. Sp1-3, E2F3, Klf1, and Nrf1 were among the key transcription factors, which were missing in the promoter p-RCAs of the downregulated hepatic genes. It is notable that TR is not one of the transcription factors identified in p-RCAs, confirming that the T3 effects observed on gene transcription are mediated in remote locations.

The analysis of the p-RCAs also revealed negative Nrf1 and Elk1 footprints in the Ncor1 promoter, one of the 146 key downregulated genes with a p-RCA in the adult Alb-D2KO liver. Ncor1 is a TR corepressor (7, 8), which in the liver plays an essential role down-regulating T3-signaling. It is fascinating that the liver responds to a reduction in neonatal T3-signalig (due to Dio2 inactivation) and preserves T3-signaling homeostasis by inhibiting the adult expression of a TR corepressor.

In conclusion, the present studies revealed that during the hepatoblast-hepatocyte transition in mice, a short-lived surge in D2 expression and T3-signaling substantially modifies the hepatic transcriptome in adult animals. By acting through intergenic TRs, T3-signaling prevents discrete methylation of specific DNA sites, which would otherwise disrupt the function of areas that operate as remote enhancers. In the presence of the normal neonatal D2-T3 peak, these distant areas interact remotely with dozens of gene promoters, increasing chromatin accessibility and expression of genes involved in multiple hepatic functions. This constitutes a novel mechanism through which TH regulates gene expression, and explains the critical role played by deiodinases in vertebrate development.

## Material and Methods

### Animals

Experiments were approved by the local Institutional Animal Care and Use Committee. A mouse with hepatocyte-specific Dio2 inactivation (ALB-D2KO mice) was obtained and maintained as previously described (22).

### Analysis of DNA hypermethylation sites

Liver hypermethylation sites (H-sites) in the Alb-D2KO mice were obtained from a previous methylome analysis that used two methods to identify H-sites: (i) Methylation Dependent ImmunoPrecipitation followed by sequencing (MeDIP-seq) and (ii) Methylation-sensitive Restriction Enzyme digestion followed by sequencing (MRE-seq) (22). Only the H-sites present in both methods were considered. H-sites were annotated to gencode.vM24. using the Partek Genomic suite software 7.0.

### ChIP-sequencing (seq) and analysis

Liver ChIP-seq for H3K9me3 was performed in P1, P5 and adult control and Alb-D2KO mice, using the SimpleChIP® Plus Enzymatic Chromatin IP Kit (Magnetic Beads; Cell Signaling). Liver from pups and adult control and Alb-D2KO mice were obtained as described (22), snap frozen and stored at −80C. A pool of 3 livers was processed for P1 or P5; 3 adult livers were processed individually. Chromatin lysates were prepared, pre-cleared with Protein G magnetic beads, and immunoprecipitated with antibodies against the H3K9me3 (Cell Signaling). Beads were extensively washed before reverse crosslinking. Chip-enriched DNA was submitted to the Genome facility at University of Chicago for library preparation and sequence using Illumina NovaSEQ6000. FASTQ files obtained from sequencing were aligned to the Mouse mm10 genome in Partek-flow platform using the BWA. The peaks were identified using MACS2 tool, broad region and p<0.05, compared with respective input samples and annotated using the gencodev.M24. This led us to P1: 280,580, P5: 265,920 and adult: 292,552 peaks in control livers vs P1: 273,767, P5: 268,693 and adult: 286,124 peaks in Alb-D2KO livers. Areas of H3K9me3 enrichment unique to ALB-D2KO livers were also identified by comparing Alb-D2KO vs controls samples, which led us to P1: 301,105, P5: 301,098 and adult: 287,032 peaks, which were filtered down to P1: 11,303, P5: 10,005 and adult: 16,730 using −log10(pvalue)>2.

The following published ChIP-seq datasets from adult mouse liver were reanalyzed and also used in the present studies: histone markers(28) - GSM722760 (H3K4me1); GSM722761 (H3K4me3); GSM851275 (H3K27ac); GSM722762 (P300); GSM722763 (Pol2); GSM722764 (Input); chromatin folding markers(31): Rad21 and CTCF (SRP116021) and TR(27) (SRP055020). The FASTQ files were aligned to transcriptome with BWA using the Partek flow platform (Partek Inc.). The peaks were identified using MACS2 tool (p<0.05), compared to respective input samples and annotated using the gencode.vM24. In all instances, pre- and post-alignment QA/QC was done using Partek Flow (Partek Inc).

### Atac-seq and analysis

Frozen livers were processed for nuclei isolation using a detergent-free kit (Invent; cat # NI-024). Nuclei integrity was verified under light microscopy (Fig. 2A). 50,000 nuclei were used for the transposase reaction as described (47). The final libraries were purified using Agencourt AMPure XP beads (Beckman Coulter), and quality checked using a Bioanalyzer High Sensitivity DNA Analysis kit (Agilent); concentration was measured through a qPCR-based method (KAPA Library Quantification Kit for Illumina Sequencing Platforms). Samples were pair-end sequenced with the Illumina HiSeq 4000 platform at the University of Chicago Genomics Facility. The remaining adapter sequences were removed using NGmerge v 0.3. Sequencing reads were aligned to the Mouse mm10 genome build using BWA v.0.7.17 and annotated using the gencode.vM24. Sequence quality was assessed using FastQC and tools from the Picard suite including CollectInsertSizeMetrics, which showed an enrichment in short fragments as expected (Fig. 2A). Aligned reads were further filtered to remove chimeric, duplicate, supplementary, and low-quality (MAPQ < 30) alignments. We determined open-chromatin regions (peaks) using Genrich v0.6 (q-value threshold☐=☐0.05). Run by individual sample, genrich would find only a handful of peaks which very obviously were not due to ATAC enrichment. These regions were merged across samples into a blacklist of regions and excluded from consideration. Ultimately genrich was run using all samples grouped together while ignoring the blacklisted regions. Differential peaks were identified by overlap subtraction to find open chromatin regions unique to each condition.

Mouse liver RCAs were classified based on the relative strengths of H3K4me1, H3K4me3, and ATAC signals (31). Signals were summarized as transcripts per kilobase millions (TPM) values in each RCA region. Those RCA regions with a TPM >4 for either H3K4me3 or H3K4me1 were segregated into one of three categories, depending on the value of the ratio of H3K4me3 to H3K4me1. High ratios of >1.5 were classified as a ‘promoter’; ratios <0.67 were classified as ‘enhancer’, while intermediate values were classified as ‘weak promoter’. Conversely RCA regions with TPM <=4 in both H3K4me3 and H3K4me1 were segregated into one of two categories depending on the relative values of ATAC-seq to H3K4me1 ratio. If the ATAC-seq signal > H3K4me1 signal, then the RCA was categorized as ‘insulator’. If the ATAC-seq signal < H3K4me1 signal, then the RCA was categorized as ‘weak insulator’.

For the motif analysis, the nucleic acid sequence of promoter ATAC-seq peak regions were extracted and masked using RepeatMasker (v 4.0.7, using pre-defined settings for mouse). Motif analysis was carried out using the meme-chip command from MEME Suite (v 5.0.5) using the MOUSE/HOCOMOCOv10, MOUSE/uniprobe, MOUSE/chen2008, EUKARYOTE/jolma2013, and JASPAR/JASPAR_CORE_2016_vertebrates motif libraries. This multiple analysis tool identified known motifs, *de novo* motifs, characteristic distances and co-occurrences, and collapsed hits by motif similarity.

### RNA Sequencing and Analysis

RNA was isolated from mouse liver using the RNeasy Kit (Qiagen). RNA degradation was monitored using a BioAnalyzer (Agilent). Samples of total RNA with RIN >7.5 were sent to Genome Technology Access Center at Washington University in St. Louis for library preparation and sequence. Libraries were pair-end sequenced with NovaSeqS4 (Illumina). Base-calls and demultiplexing were performed with Illumina’s bcl2fastq software and a custom python demultiplexing program with a maximum of one mismatch in the indexing read. The FASTQ files were aligned to gencode.vM24 transcriptome with STAR using the Partek flow platform (Partek Inc.). All pre- and post-alignment QA/QC was performed in Partek Flow (Partek Inc.) (Fig. S1B). Aligned reads were quantified to annotation model (Partek E/M) and normalized (absolute value). Following the differential analysis (GSA), the biological significance of the changes was interpreted using gene set enrichment analysis (GSEA; Table S4) and pathway enrichment analyses (Table S5).

### Hi-C data analysis

Publicly available liver Hi-C datasets (GSE65126) were analyzed using the HICUP (48) pipeline. Briefly, the HICUP pipeline take as input FASTQ data from a Hi-C experiment and produces a filtered set of interaction pairs mapped to the reference genome (mm10). The post-pipeline analysis of the output from HICUP was done using HOMER (49) to create Hi-C interaction and contact matrices. Matrices were then analyzed for long-range interactions involving p-RCAs, i-RCAs as obtained from the analysis of the ATAC-seq, and H-sites.

### Bioinformatics tools

R-script using the Bioconductor IRanges package (50) (proximity analysis) was used to obtain the intersection of the H-sites and different chromatin markers on the genomic coordinates for the desired distance. The heat maps used to analyze proximity between different chromosomes coordinates (Fig. 1B; 1D-E; Fig. 2C-D; Fig. S1A), were generated using the interactive platform for analysis Easeq. BED files containing the start/end chromosomal coordinates of each peak were used to generate the images. The circos plots (Fig. 5E; Fig.S2A-J) were created with Circa software tool (http://omgenomics.com/circa). The Integrative Genomics Viewer (IGV) 2.7.2 version was use to visualize and integrated all the genomic data analysis (http://www.broadinstitute.org/igv).

## Supporting information

all supplemental files

## Authors contribution

T. L. F. conducted all experiments, data analyzes, prepared the manuscript figures and Tables; T.G. performed atac-seq bioinformatics analysis; T.M.N. performed HiC-seq data analysis; G.W.F. preparation of heat maps; A.C.B. planned and directed all studies and manuscript write up.

## Acknowledgement

This work was supported by National Institute of Diabetes and Digestive and Kidney Diseases grants DK58538 and DK65055. We thank Dr. Antonio Lerario for the initial Atac-seq data analysis.

## References

1. Brent GA. Mechanisms of thyroid hormone action. The Journal of clinical investigation. 2012;122(9):3035–43.

2. Hones GS, Rakov H, Logan J, Liao XH, Werbenko E, Pollard AS, et al. Noncanonical thyroid hormone signaling mediates cardiometabolic effects in vivo. Proceedings of the National Academy of Sciences of the United States of America. 2017;114(52):E11323–E32.

3. Javaid N, and Choi S. Acetylation- and Methylation-Related Epigenetic Proteins in the Context of Their Targets. Genes (Basel). 2017;8(8).

4. Hyun K, Jeon J, Park K, and Kim J. Writing, erasing and reading histone lysine methylations. Exp Mol Med. 2017;49(4):e324.

5. Briggs SD, Xiao T, Sun ZW, Caldwell JA, Shabanowitz J, Hunt DF, et al. Gene silencing: trans-histone regulatory pathway in chromatin. Nature. 2002;418(6897):498.

6. Briggs SD, and Strahl BD. Unraveling heterochromatin. Nat Genet. 2002;30(3):241–2.

7. Ritter MJ, Amano I, and Hollenberg AN. Thyroid Hormone Signaling and the Liver. Hepatology. 2020.

8. Strahl BD, Grant PA, Briggs SD, Sun ZW, Bone JR, Caldwell JA, et al. Set2 is a nucleosomal histone H3-selective methyltransferase that mediates transcriptional repression. Molecular and cellular biology. 2002;22(5):1298–306.

9. Li J, Lin Q, Yoon HG, Huang ZQ, Strahl BD, Allis CD, et al. Involvement of histone methylation and phosphorylation in regulation of transcription by thyroid hormone receptor. Mol Cell Biol. 2002;22(16):5688–97.

10. Choi HK, Choi KC, Oh SY, Kang HB, Lee YH, Haam S, et al. The functional role of the CARM1-SNF5 complex and its associated HMT activity in transcriptional activation by thyroid hormone receptor. Exp Mol Med. 2007;39(4):544–55.

11. Praestholm SM, Siersbaek MS, Nielsen R, Zhu X, Hollenberg AN, Cheng SY, et al. Multiple mechanisms regulate H3 acetylation of enhancers in response to thyroid hormone. PLoS Genet. 2020;16(5):e1008770.

12. Fuks F, Hurd PJ, Wolf D, Nan X, Bird AP, and Kouzarides T. The methyl-CpG-binding protein MeCP2 links DNA methylation to histone methylation. The Journal of biological chemistry. 2003;278(6):4035–40.

13. Fuks F, Hurd PJ, Deplus R, and Kouzarides T. The DNA methyltransferases associate with HP1 and the SUV39H1 histone methyltransferase. Nucleic acids research. 2003;31(9):2305–12.

14. Demeneix BA. Evidence for Prenatal Exposure to Thyroid Disruptors and Adverse Effects on Brain Development. Eur Thyroid J. 2019;8(6):283–92.

15. Sadakierska-Chudy A, and Filip M. A comprehensive view of the epigenetic landscape. Part II: Histone post-translational modification, nucleosome level, and chromatin regulation by ncRNAs. Neurotox Res. 2015;27(2):172–97.

16. Fu L, Wen L, and Shi YB. Role of Thyroid Hormone Receptor in Amphibian Development. Methods Mol Biol. 2018;1801:247–63.

17. Galton VA. The roles of the iodothyronine deiodinases in mammalian development. Thyroid: official journal of the American Thyroid Association. 2005;15(8):823–34.

18. Dussault JH, and Labrie F. Development of the hypothalamic-pituitary-thyroid axis in the neonatal rat. Endocrinology. 1975;97(5):1321–4.

19. Fukuda H, and Greer MA. Postnatal development of pituitary-thyroid function in male and female rats: comparison of plasma and thyroid T3 and T4 concentration. J Endocrinol Invest. 1978;1(4):311–4.

20. Hall JA, Ribich S, Christoffolete MA, Simovic G, Correa-Medina M, Patti ME, et al. Absence of thyroid hormone activation during development underlies a permanent defect in adaptive thermogenesis. Endocrinology. 2010;151(9):4573–82.

21. Campos-Barros A, Amma LL, Faris JS, Shailam R, Kelley MW, and Forrest D. Type 2 iodothyronine deiodinase expression in the cochlea before the onset of hearing. Proceedings of the National Academy of Science. 2000;97(3):1287–92.

22. Fonseca TL, Fernandes GW, McAninch EA, Bocco BM, Abdalla SM, Ribeiro MO, et al. Perinatal deiodinase 2 expression in hepatocytes defines epigenetic susceptibility to liver steatosis and obesity. Proceedings of the National Academy of Sciences of the United States of America. 2015;112(45):14018–23.

23. Darlington GJ, Wang N, and Hanson RW. C/EBP alpha: a critical regulator of genes governing integrative metabolic processes. Current opinion in genetics & development. 1995;5(5):565–70.

24. Fonseca TL, Fernandes GW, Bocco B, Keshavarzian A, Jakate S, Donohue TM, Jr., et al. Hepatic Inactivation of the Type 2 Deiodinase Confers Resistance to Alcoholic Liver Steatosis. Alcohol Clin Exp Res. 2019.

25. Reizel Y, Sabag O, Skversky Y, Spiro A, Steinberg B, Bernstein D, et al. Postnatal DNA demethylation and its role in tissue maturation. Nature communications. 2018;9(1):2040.

26. Cannon MV, Pilarowski G, Liu X, and Serre D. Extensive Epigenetic Changes Accompany Terminal Differentiation of Mouse Hepatocytes After Birth. G3 (Bethesda). 2016;6(11):3701–9.

27. Grontved L, Waterfall JJ, Kim DW, Baek S, Sung MH, Zhao L, et al. Transcriptional activation by the thyroid hormone receptor through ligand-dependent receptor recruitment and chromatin remodelling. Nature communications. 2015;6:7048.

28. Shen Y, Yue F, McCleary DF, Ye Z, Edsall L, Kuan S, et al. A map of the cis-regulatory sequences in the mouse genome. Nature. 2012;488(7409):116–20.

29. Hay D, Hughes JR, Babbs C, Davies JOJ, Graham BJ, Hanssen L, et al. Genetic dissection of the alpha-globin super-enhancer in vivo. Nat Genet. 2016;48(8):895–903.

30. Ehrlich M. DNA hypermethylation in disease: mechanisms and clinical relevance. Epigenetics. 2019;14(12):1141–63.

31. Matthews BJ, and Waxman DJ. Computational prediction of CTCF/cohesin-based intra-TAD loops that insulate chromatin contacts and gene expression in mouse liver. Elife. 2018;7.

32. Lamparter D, Marbach D, Rueedi R, Bergmann S, and Kutalik Z. Genome-Wide Association between Transcription Factor Expression and Chromatin Accessibility Reveals Regulators of Chromatin Accessibility. PLoS computational biology. 2017;13(1):e1005311.

33. Hoppe KL, and Francone OL. Binding and functional effects of transcription factors Sp1 and Sp3 on the proximal human lecithin:cholesterol acyltransferase promoter. J Lipid Res. 1998;39(5):969–77.

34. Fajas L, Landsberg RL, Huss-Garcia Y, Sardet C, Lees JA, and Auwerx J. E2Fs regulate adipocyte differentiation. Dev Cell. 2002;3(1):39–49.

35. Widenmaier SB, Snyder NA, Nguyen TB, Arduini A, Lee GY, Arruda AP, et al. NRF1 Is an ER Membrane Sensor that Is Central to Cholesterol Homeostasis. Cell. 2017;171(5):1094–109 e15.

36. Katsuoka F, Yamazaki H, and Yamamoto M. Small Maf deficiency recapitulates the liver phenotypes of Nrf1- and Nrf2-deficient mice. Genes Cells. 2016;21(12):1309–19.

37. Khan R, Raza SHA, Junjvlieke Z, Xiaoyu W, Garcia M, Elnour IE, et al. Function and Transcriptional Regulation of Bovine TORC2 Gene in Adipocytes: Roles of C/EBP, XBP1, INSM1 and ZNF263. Int J Mol Sci. 2019;20(18).

38. Lu L, Ye X, Yao Q, Lu A, Zhao Z, Ding Y, et al. Egr2 enhances insulin resistance via JAK2/STAT3/SOCS-1 pathway in HepG2 cells treated with palmitate. Gen Comp Endocrinol. 2018;260:25–31.

39. Ye BY, Shen WL, Wang D, Li P, Zhang Z, Shi ML, et al. [ZNF143 is involved in CTCF-mediated chromatin interactions by cooperation with cohesin and other partners]. Mol Biol (Mosk). 2016;50(3):496–503.

40. Allshire RC, and Madhani HD. Ten principles of heterochromatin formation and function. Nat Rev Mol Cell Biol. 2018;19(4):229–44.

41. Holwerda S, and de Laat W. Chromatin loops, gene positioning, and gene expression. Front Genet. 2012;3:217.

42. Vietri Rudan M, Barrington C, Henderson S, Ernst C, Odom DT, Tanay A, et al. Comparative Hi-C reveals that CTCF underlies evolution of chromosomal domain architecture. Cell Rep. 2015;10(8):1297–309.

43. Ng L, Goodyear RJ, Woods CA, Schneider MJ, Diamond E, Richardson GP, et al. Hearing loss and retarded cochlear development in mice lacking type 2 iodothyronine deiodinase. Proceedings of the National Academy of Sciences of the United States of America. 2004;101(10):3474–9.

44. Ehara T, Kamei Y, Yuan X, Takahashi M, Kanai S, Tamura E, et al. Ligand-activated PPARalpha-dependent DNA demethylation regulates the fatty acid beta-oxidation genes in the postnatal liver. Diabetes. 2015;64(3):775–84.

45. Lachner M, O’Carroll D, Rea S, Mechtler K, and Jenuwein T. Methylation of histone H3 lysine 9 creates a binding site for HP1 proteins. Nature. 2001;410(6824):116–20.

46. Fritsch L, Robin P, Mathieu JR, Souidi M, Hinaux H, Rougeulle C, et al. A subset of the histone H3 lysine 9 methyltransferases Suv39h1, G9a, GLP, and SETDB1 participate in a multimeric complex. Mol Cell. 2010;37(1):46–56.

47. Buenrostro JD, Giresi PG, Zaba LC, Chang HY, and Greenleaf WJ. Transposition of native chromatin for fast and sensitive epigenomic profiling of open chromatin, DNA-binding proteins and nucleosome position. Nat Methods. 2013;10(12):1213–8.

48. Wingett S, Ewels P, Furlan-Magaril M, Nagano T, Schoenfelder S, Fraser P, et al. HiCUP: pipeline for mapping and processing Hi-C data. F1000Res. 2015;4:1310.

49. Heinz S, Benner C, Spann N, Bertolino E, Lin YC, Laslo P, et al. Simple combinations of lineage-determining transcription factors prime cis-regulatory elements required for macrophage and B cell identities. Mol Cell. 2010;38(4):576–89.

50. Lawrence M, Huber W, Pages H, Aboyoun P, Carlson M, Gentleman R, et al. Software for computing and annotating genomic ranges. PLoS Comput Biol. 2013;9(8):e1003118.

