## Supplementary material for "Neonatal Thyroxine Activation Modifies Epigenetic Programming of The Liver": all supplemental files

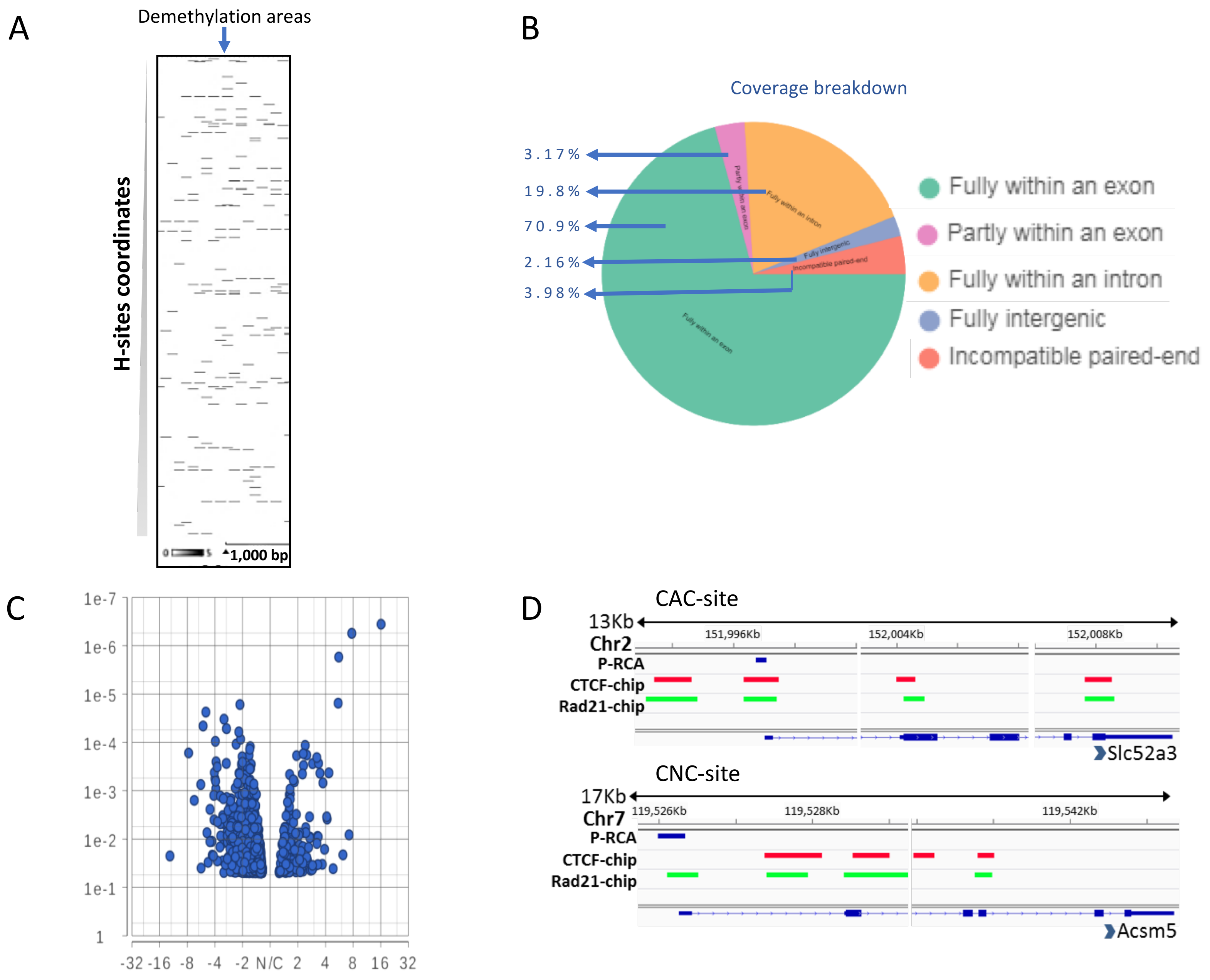

Suppl. 2- (A) Heat map of H-sites coordinates vs. demethylation sites coordinates; the bottom arrow indicates 1,000 bp; the Y axis contains the H-sites coordinates, and the X axis contains the coordinates of the sites that undergo post-natal demethylation; the coordinates in the Y axis follows the chromosomal sequence: Chr 1 to Chr 9, Chr X, Chr 10 to Chr 19); (C) RNA-seq coverage breakdown; distribution reflects the average of all samples; (B) differential gene expression (Volcano plot) in the Alb-D2KO vs control liver as detected by RNA-seq; each point represents the average value of one transcript in 3-4 animals. The expression difference is considered significant if $p<0.05$; (D) browser IGV display of two genes (Slc52a3 and Acsm5 ) that contain a p-RCA site (blue track); CTCF-ChIP-seq peak (red track) and Rad21-ChIP-seq peak “cohesin” (green track). Slc52a3 contains a cohesin and CTCF configuration typical of insulator (CAC-sites); Acsm5 contains one isolated cohesin peak (CNC-sites).

Suppl. 1

Suppl. 2

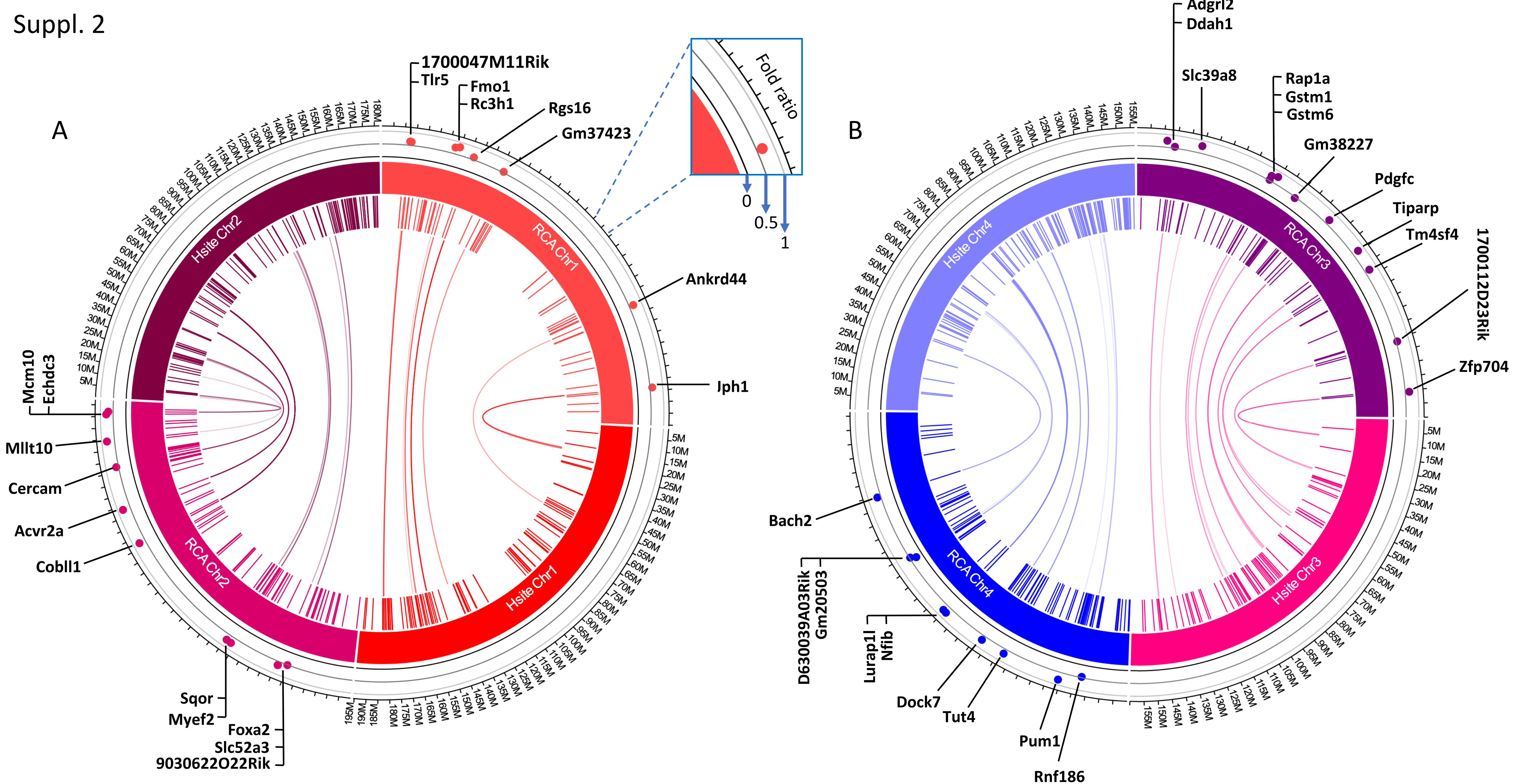

(A-J) Circa diagrams of the indicated chromosomes depicting H-sites and p-RCA areas of indicated 146 genes in which the p-RCA areas are within 5Kbp of the negative RNA-seq peaks. The outer ring (shown in the inset) is the ratio Alb-D2KO / Control gene expression: the inner black line is “0”; the dark gray line “0.5” and light gray line “1” fold-change; the location of the p-RCAs and the H-sites are indicated in the inner ring for each chromosome; the connecting lines indicate positions of areas-1 and areas-2 from Hi-C data.

Suppl. 3

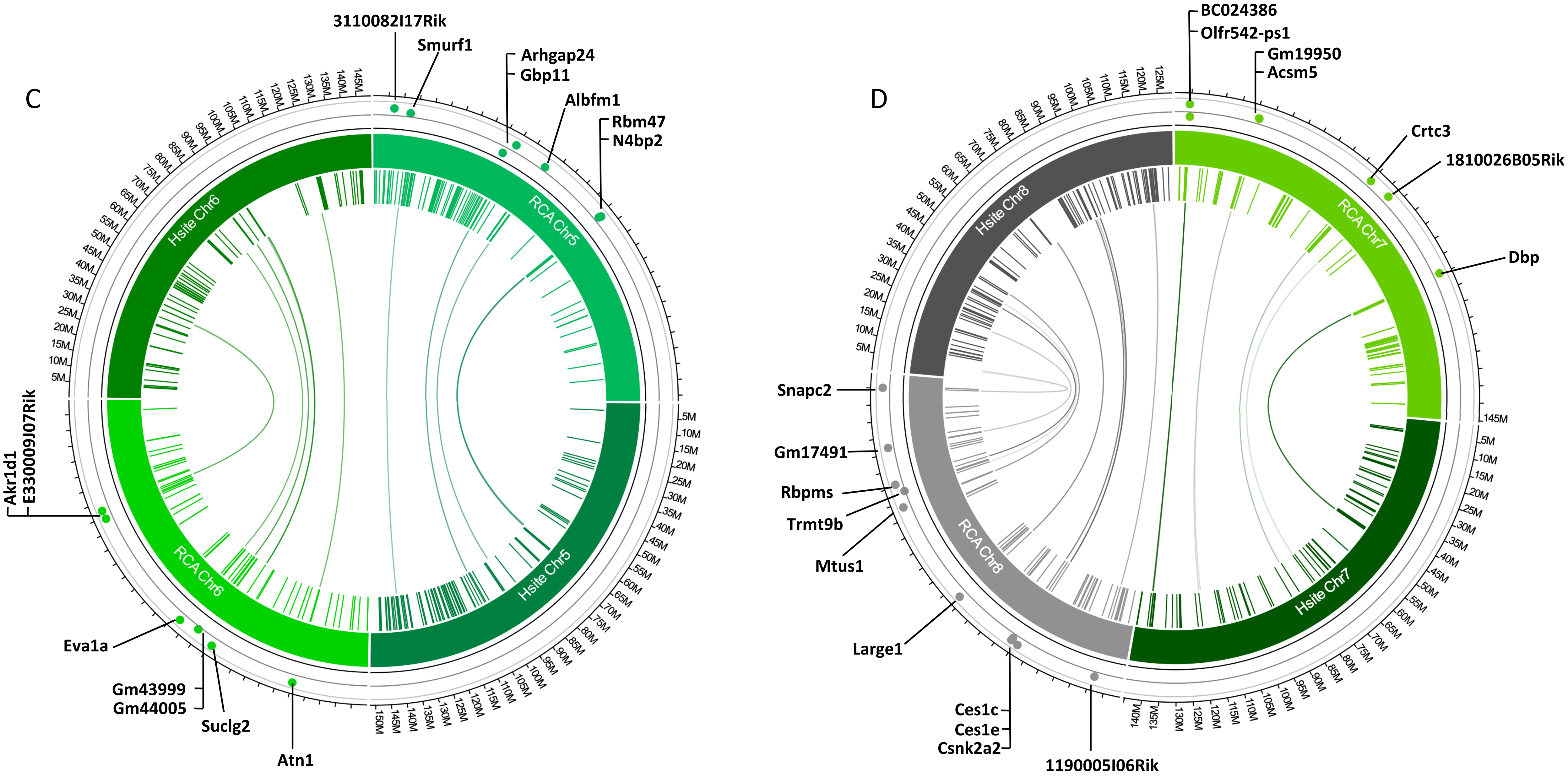

C
3110082I17Rik
Smurf1
Arhgap24
Gbp11
Albfm1
Rbm47
N4bp2
Hsite Chr6
RCA Chr5
RCA Chr6
Hsite Chr5
Akr1d1
E330009J07Rik
Eva1a
Gm43999
Gm44005
Suclg2
Atn1
D
BC024386
Olfr542-ps1
Gm19950
Acsm5
Crtc3
1810026B05Rik
Dbp
Hsite Chr8
RCA Chr7
RCA Chr8
Hsite Chr7
Snapc2
Gm17491
Rbpms
Trmt9b
Mtus1
Large1
Ces1c
Ces1e
Csnk2a2
1190005I06Rik

Suppl. 4

E

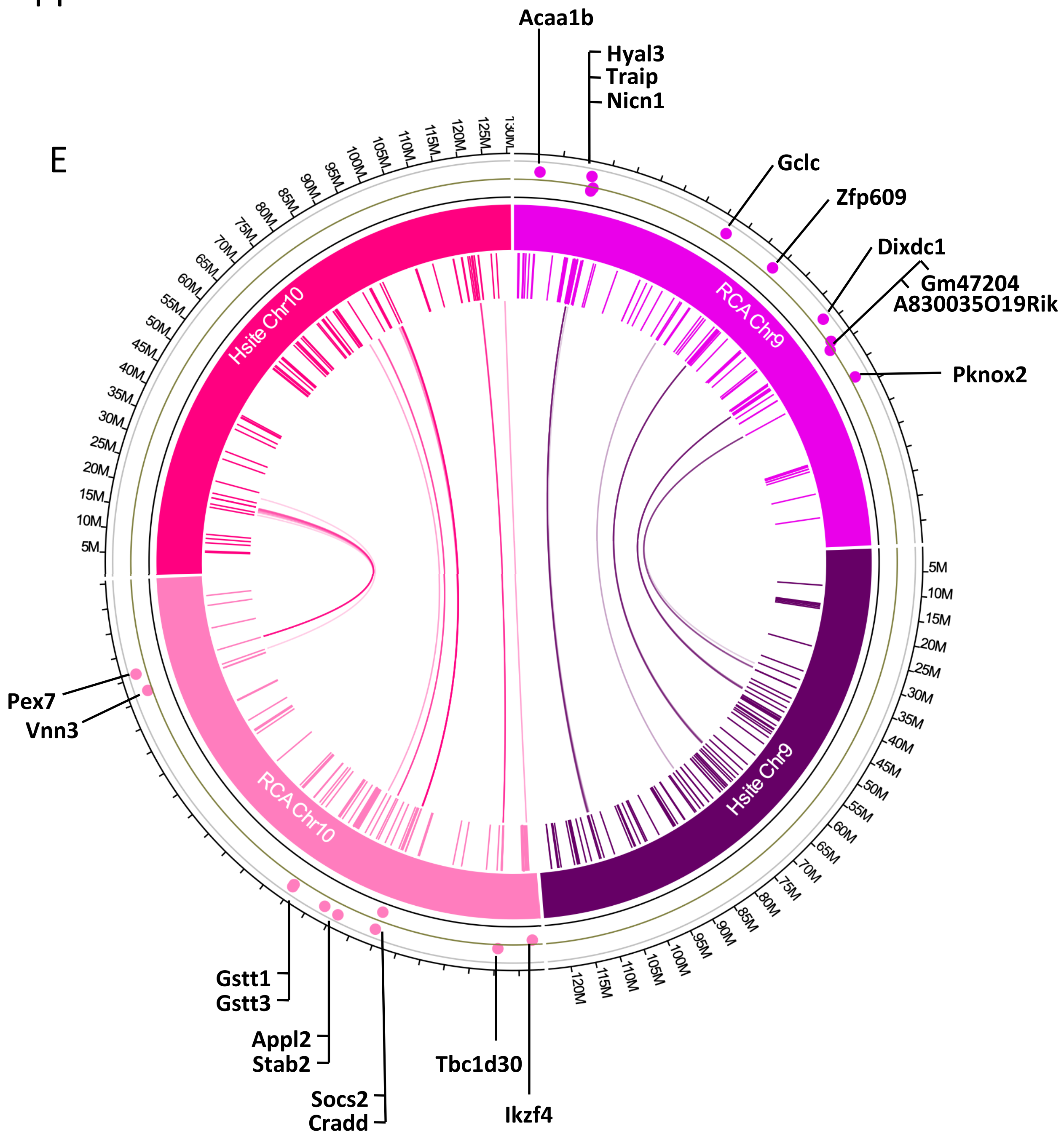
Acaa1b
Hyal3
Traip
Nicn1
Gclc
Zfp609
Dixdc1
Gm47204
A830035O19Rik
Pknox2
Hsite Chr10
RCA Chr9
Hsite Chr9
RCA Chr10
Pex7
Vnn3
Gstt1
Gstt3
Appl2
Stab2
Socs2
Cradd
Tbc1d30
Ikzf4

F

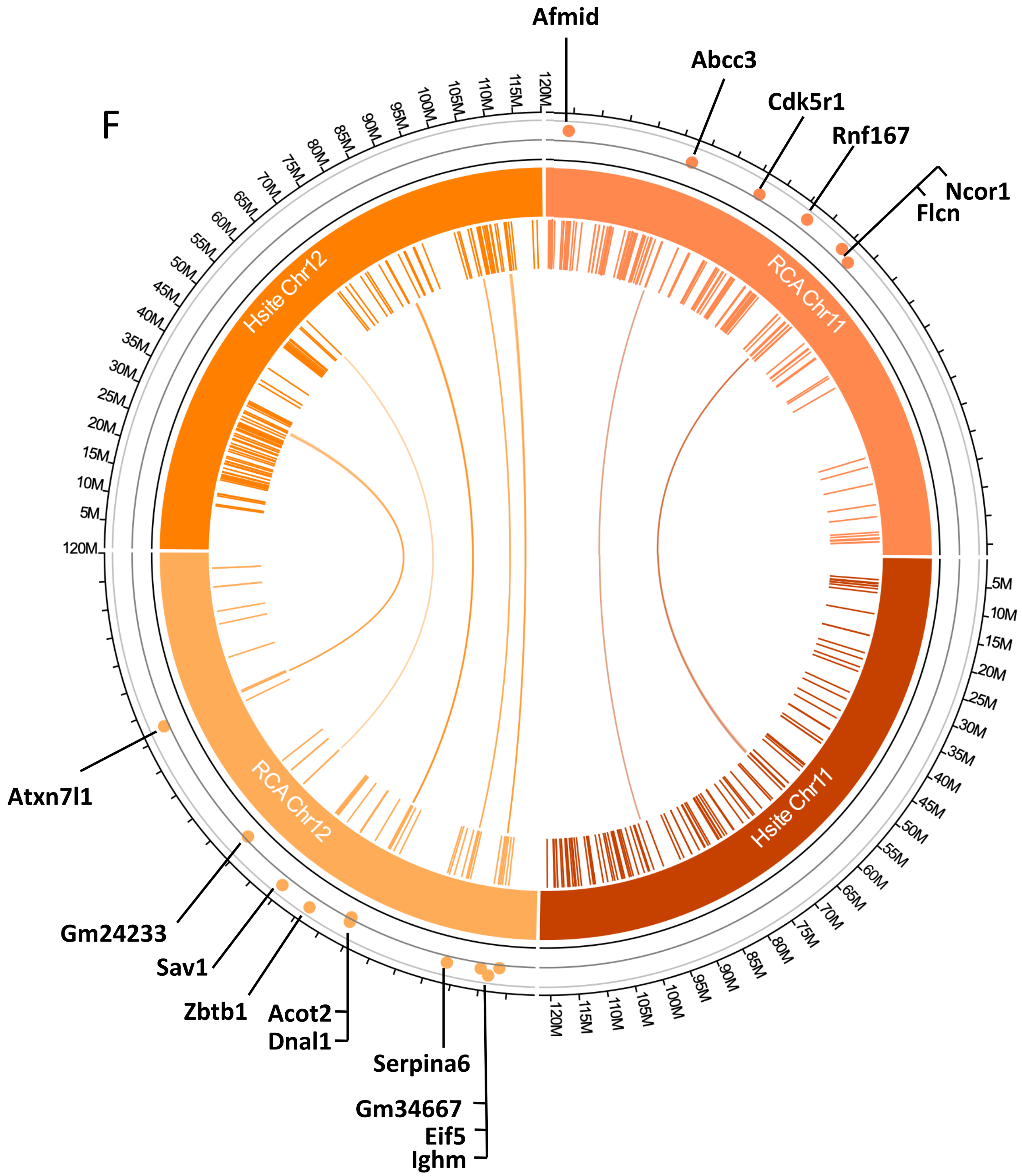
Afmid
Abcc3
Cdk5r1
Rnf167
Ncor1
Flcn
Hsite Chr12
RCA Chr11
RCA Chr12
Hsite Chr11
Atxn7l1
Gm24233
Sav1
Zbtb1
Acot2
Dnal1
Serpina6
Gm34667
Eif5
Ighm

Suppl. 5

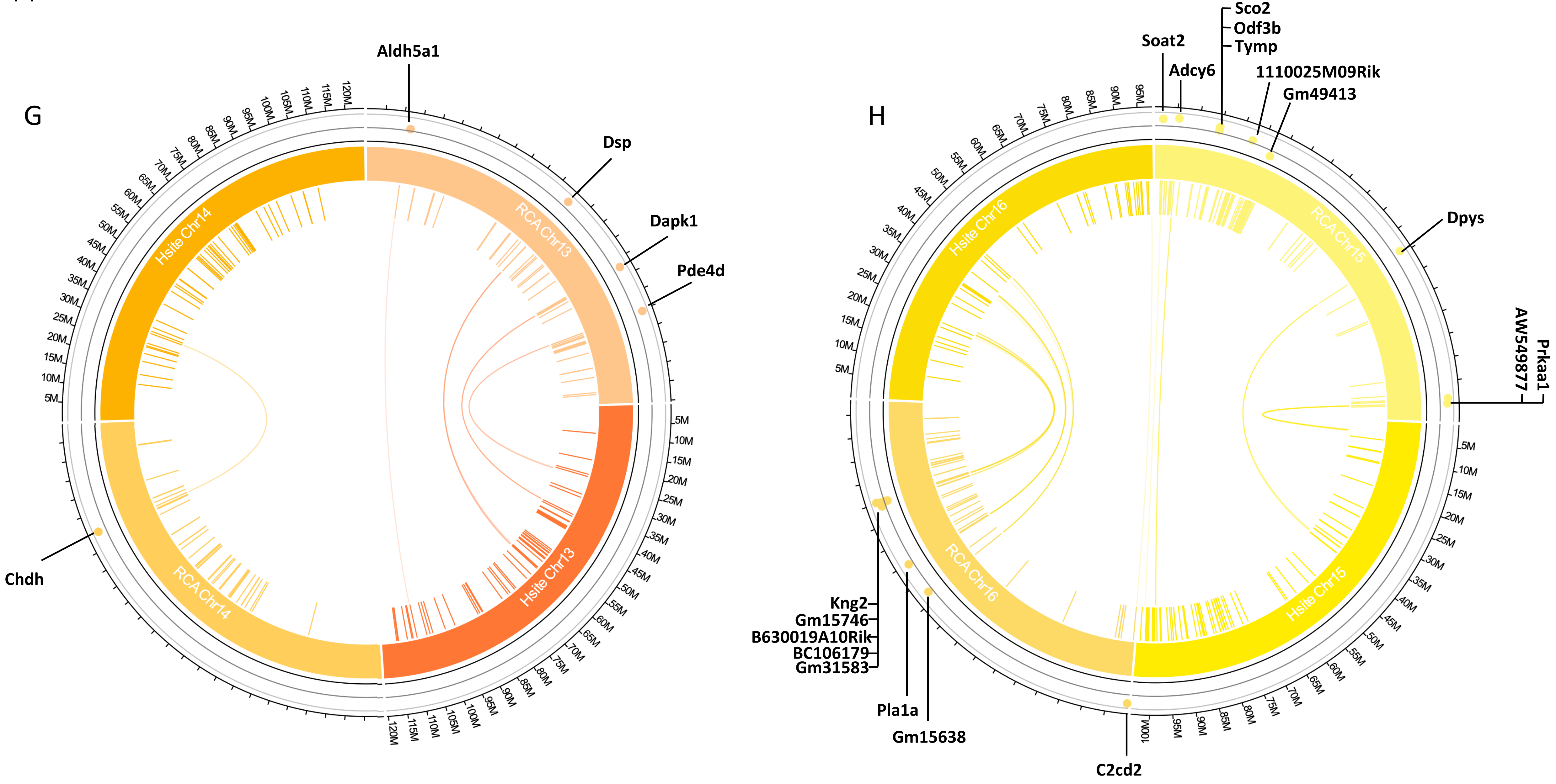
G
Aldh5a1
Dsp
Dapk1
Pde4d
Chdh
Hsite Chr14
RCA Chr13
Hsite Chr13
RCA Chr14
H
Soat2
Adcy6
Sco2
Odf3b
Tymp
1110025M09Rik
Gm49413
Dpys
Prkaa1
AW549877
Kng2
Gm15746
B630019A10Rik
BC106179
Gm31583
Pla1a
Gm15638
C2cd2
Hsite Chr16
RCA Chr15
Hsite Chr15
RCA Chr16

Suppl. 6

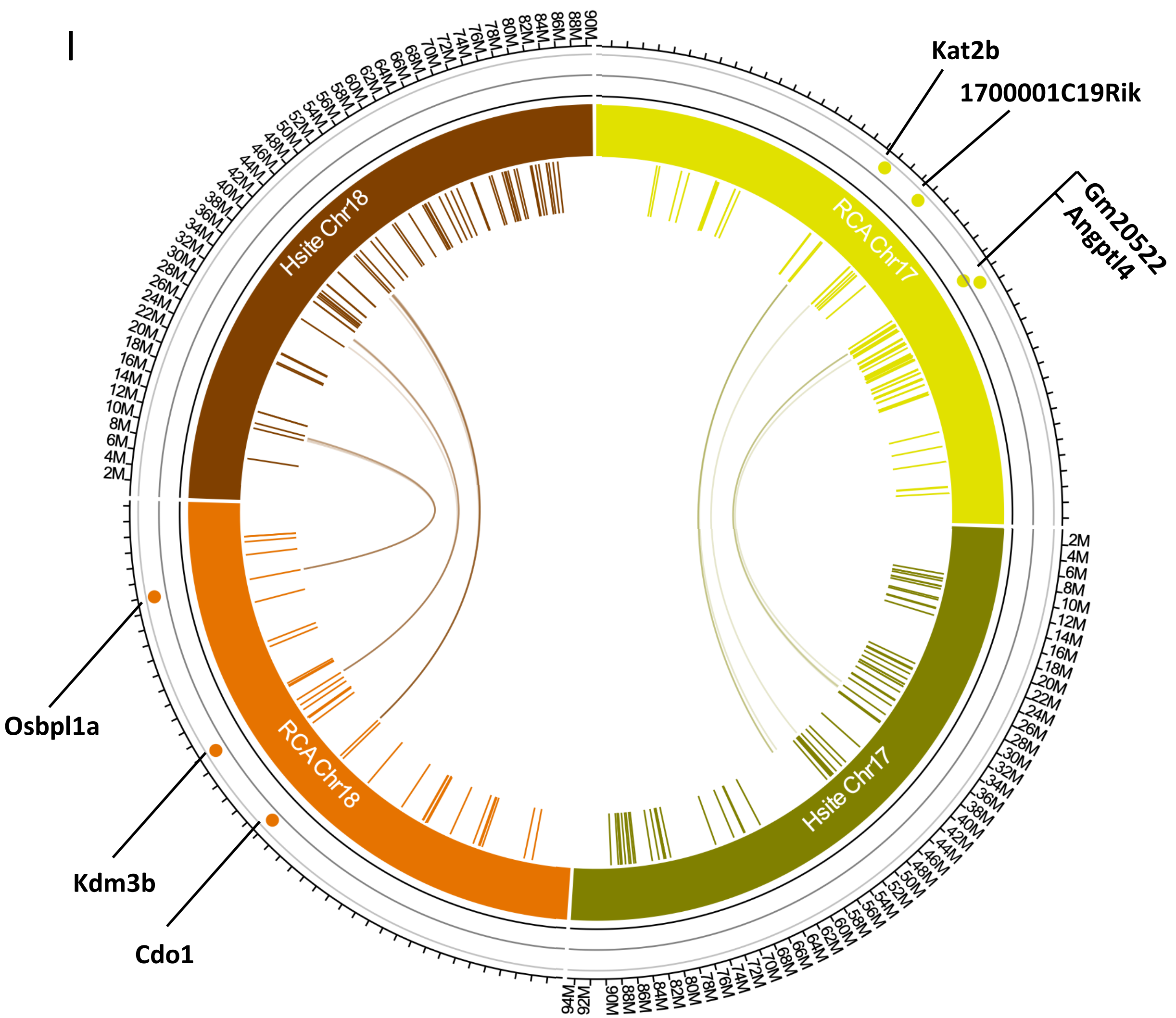
I
Kat2b
1700001C19Rik
Gm20522
Angptl4
Osbpl1a
Kdm3b
Cdo1
Hsite Chr18
RCA Chr17
Hsite Chr17
RCA Chr18

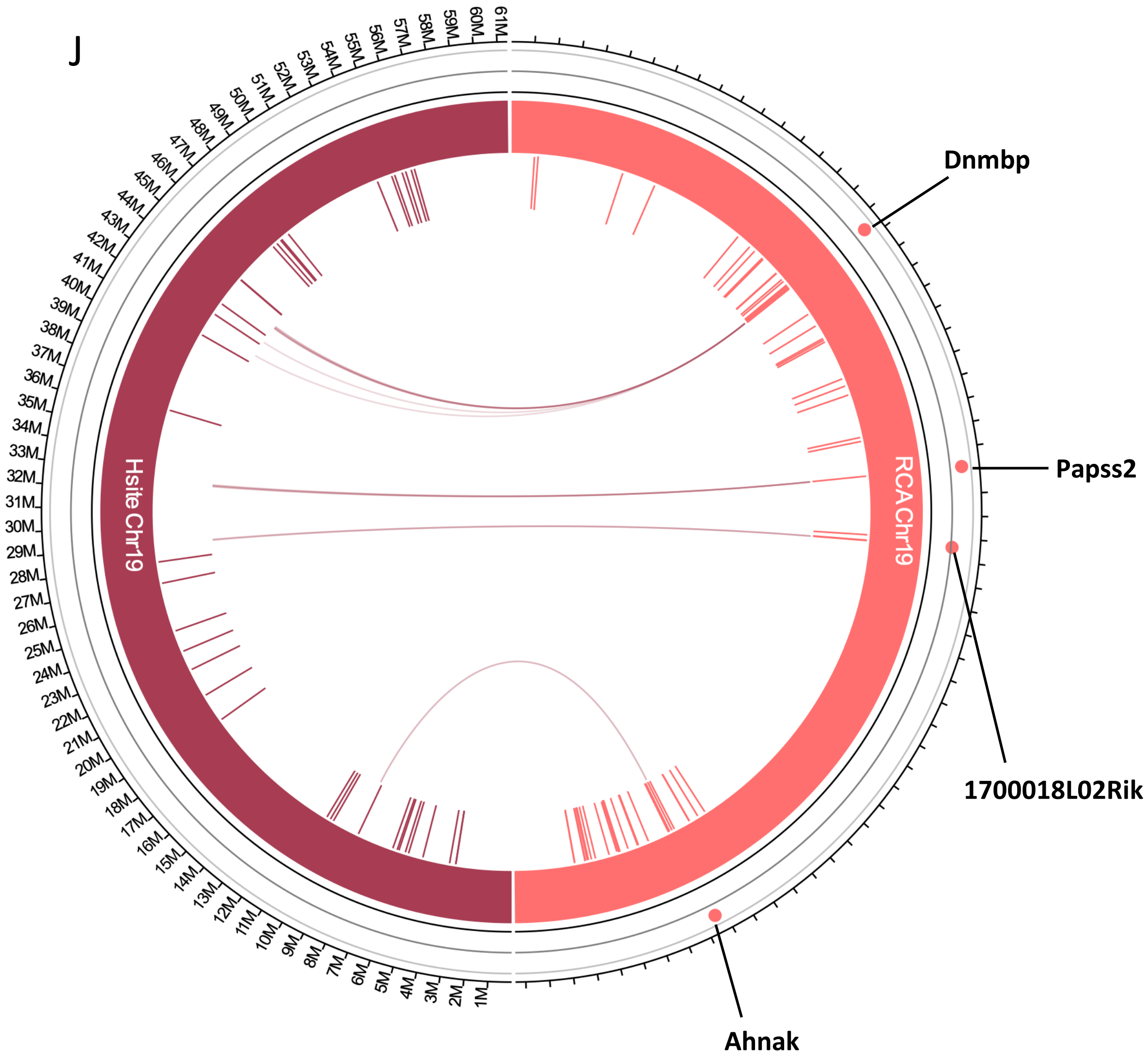
J
Dnmbp
Papss2
1700018L02Rik
Ahnak
Hsite Chr19
RCA Chr19

**Table S1**- H-sites coordinates in Alb-D2KO mouse liver that are imbedded in areas enriched with H3K9me3. Also shown is the presence of a TR ChIP-seq peak within 10,000 bp.

| **H-sites** | | | **H-sites imbedded in H3K9me3 area** | | | | **H-sites flanked by TR-ChIP peaks** | | |
|---|---|---|---|---|---|---|---|---|---|
| **Chromosome** | **start** | **end** | **P1** | **P5** | **P1+P5** | **Adult** | **H-site** | **uTRs** | **T3-TRs** |
| chr1 | 50386656 | 50387156 | 1 | 1 | 1 | 0 | 1 | 0 | 1 |
| chr1 | 25432155 | 25432655 | 1 | 1 | 1 | 0 | 1 | 0 | 0 |
| chr1 | 188902621 | 188903121 | 1 | 1 | 1 | 0 | 1 | 0 | 1 |
| chr1 | 64288926 | 64289426 | 1 | 1 | 1 | 0 | 1 | 0 | 1 |
| chr1 | 157803869 | 157804369 | 1 | 1 | 1 | 1 | 1 | 0 | 1 |
| chr1 | 55362156 | 55362656 | 0 | 1 | 0 | 1 | 1 | 0 | 1 |
| chr1 | 88222425 | 88222925 | 1 | 1 | 1 | 0 | 1 | 1 | 1 |
| chr1 | 154688870 | 154689370 | 0 | 1 | 0 | 0 | 1 | 0 | 1 |
| chr1 | 67097926 | 67098426 | 1 | 1 | 1 | 0 | 1 | 1 | 1 |
| chr1 | 146106370 | 146106870 | 1 | 1 | 1 | 0 | 1 | 0 | 0 |
| chr1 | 51747656 | 51748156 | 0 | 1 | 0 | 0 | 1 | 1 | 1 |
| chr1 | 95627423 | 95627923 | 1 | 1 | 1 | 0 | 1 | 0 | 1 |
| chr1 | 168879369 | 168879869 | 0 | 1 | 0 | 0 | 1 | 0 | 0 |
| chr2 | 138496764 | 138497264 | 1 | 0 | 0 | 1 | 1 | 0 | 1 |
| chr2 | 162142264 | 162142764 | 1 | 0 | 0 | 0 | 1 | 0 | 0 |
| chr2 | 89297843 | 89298343 | 0 | 1 | 0 | 0 | 1 | 0 | 1 |
| chr2 | 172058000 | 172058500 | 1 | 0 | 0 | 1 | 1 | 0 | 0 |
| chr3 | 93779578 | 93780078 | 1 | 1 | 1 | 0 | 1 | 0 | 1 |
| chr3 | 52415078 | 52415578 | 1 | 0 | 0 | 0 | 1 | 0 | 1 |
| chr4 | 146486538 | 146487038 | 1 | 1 | 1 | 0 | 1 | 1 | 1 |
| chr4 | 77040096 | 77040596 | 1 | 0 | 0 | 0 | 1 | 0 | 0 |
| chr4 | 61698966 | 61699466 | 1 | 0 | 0 | 0 | 1 | 0 | 1 |
| chr4 | 145449097 | 145449597 | 1 | 0 | 0 | 0 | 1 | 1 | 1 |
| chr4 | 61701966 | 61702466 | 1 | 0 | 0 | 0 | 1 | 1 | 1 |
| chr4 | 61715966 | 61716466 | 1 | 0 | 0 | 0 | 1 | 0 | 1 |
| chr7 | 14596651 | 14597151 | 1 | 0 | 0 | 1 | 1 | 0 | 1 |
| chr8 | 96151100 | 96151600 | 1 | 0 | 0 | 0 | 1 | 0 | 1 |
| chr8 | 40272646 | 40273146 | 1 | 0 | 0 | 1 | 1 | 0 | 1 |
| chr9 | 86902665 | 86903165 | 1 | 0 | 0 | 0 | 1 | 0 | 1 |
| chr10 | 126770944 | 126771444 | 0 | 1 | 0 | 0 | 1 | 0 | 0 |
| chr10 | 104933944 | 104934444 | 1 | 1 | 1 | 0 | 1 | 0 | 1 |
| chr11 | 18181997 | 18182497 | 1 | 1 | 1 | 0 | 1 | 0 | 1 |
| chr11 | 18182497 | 18182997 | 1 | 1 | 1 | 0 | 1 | 0 | 1 |
| chr11 | 38913498 | 38913998 | 1 | 0 | 0 | 0 | 1 | 0 | 1 |

| | | | | | | | | | |
|---|---|---|---|---|---|---|---|---|---|
| chr11 | 25856500 | 25857000 | 1 | 0 | 0 | 0 | 1 | 0 | 1 |
| chr12 | 20274645 | 20275145 | 0 | 1 | 0 | 0 | 1 | 0 | 1 |
| chr12 | 20267145 | 20267645 | 0 | 1 | 0 | 0 | 1 | 0 | 1 |
| chr12 | 20278145 | 20278645 | 0 | 1 | 0 | 0 | 1 | 0 | 1 |
| chr12 | 53220513 | 53221013 | 1 | 0 | 0 | 1 | 1 | 0 | 1 |
| chr12 | 20920139 | 20920639 | 1 | 1 | 1 | 0 | 1 | 0 | 1 |
| chr12 | 27136135 | 27136635 | 1 | 0 | 0 | 0 | 1 | 0 | 0 |
| chr12 | 20239145 | 20239645 | 1 | 1 | 1 | 0 | 1 | 0 | 1 |
| chr12 | 20264645 | 20265145 | 1 | 1 | 1 | 0 | 1 | 0 | 1 |
| chr12 | 20294145 | 20294645 | 0 | 1 | 0 | 0 | 1 | 0 | 1 |
| chr12 | 51492513 | 51493013 | 0 | 1 | 0 | 0 | 1 | 0 | 1 |
| chr12 | 18237694 | 18238194 | 1 | 1 | 1 | 0 | 1 | 0 | 1 |
| chr12 | 18328694 | 18329194 | 1 | 0 | 0 | 0 | 1 | 0 | 1 |
| chr12 | 18229694 | 18230194 | 1 | 1 | 1 | 0 | 1 | 0 | 1 |
| chr12 | 20281645 | 20282145 | 0 | 1 | 0 | 0 | 1 | 0 | 1 |
| chr12 | 20251645 | 20252145 | 1 | 1 | 1 | 0 | 1 | 0 | 1 |
| chr12 | 20845139 | 20845639 | 1 | 0 | 0 | 1 | 1 | 0 | 0 |
| chr12 | 16343694 | 16344194 | 0 | 1 | 0 | 1 | 1 | 0 | 1 |
| chr12 | 23267439 | 23267939 | 1 | 1 | 1 | 0 | 1 | 0 | 1 |
| chr12 | 24425135 | 24425635 | 1 | 0 | 0 | 0 | 1 | 0 | 1 |
| chr12 | 19199145 | 19199645 | 1 | 1 | 1 | 1 | 1 | 0 | 1 |
| chr12 | 68576513 | 68577013 | 1 | 0 | 0 | 0 | 1 | 0 | 1 |
| chr12 | 18243694 | 18244194 | 1 | 0 | 0 | 0 | 1 | 0 | 1 |
| chr12 | 23775135 | 23775635 | 1 | 1 | 1 | 0 | 1 | 0 | 1 |
| chr12 | 53221013 | 53221513 | 1 | 0 | 0 | 1 | 1 | 0 | 1 |
| chr12 | 20269145 | 20269645 | 1 | 1 | 1 | 0 | 1 | 0 | 1 |
| chr12 | 20284645 | 20285145 | 0 | 1 | 0 | 0 | 1 | 0 | 1 |
| chr12 | 20285145 | 20285645 | 0 | 1 | 0 | 0 | 1 | 0 | 1 |
| chr12 | 20291645 | 20292145 | 0 | 1 | 0 | 0 | 1 | 0 | 1 |
| chr12 | 20214645 | 20215145 | 0 | 1 | 0 | 1 | 1 | 0 | 0 |
| chr12 | 20293645 | 20294145 | 0 | 1 | 0 | 0 | 1 | 0 | 1 |
| chr12 | 51492013 | 51492513 | 1 | 1 | 1 | 0 | 1 | 0 | 1 |
| chr12 | 20219645 | 20220145 | 0 | 1 | 0 | 1 | 1 | 0 | 0 |
| chr13 | 111665292 | 111665792 | 1 | 1 | 1 | 1 | 1 | 0 | 1 |
| chr13 | 65311692 | 65312192 | 1 | 0 | 0 | 0 | 1 | 0 | 1 |
| chr14 | 86215693 | 86216193 | 1 | 0 | 0 | 0 | 1 | 0 | 1 |
| chr14 | 90987213 | 90987713 | 1 | 0 | 0 | 0 | 1 | 1 | 0 |
| chr14 | 78282693 | 78283193 | 1 | 0 | 0 | 0 | 1 | 0 | 1 |
| chr14 | 93899281 | 93899781 | 1 | 0 | 0 | 0 | 1 | 0 | 0 |
| chr15 | 18434245 | 18434745 | 1 | 1 | 1 | 0 | 1 | 0 | 1 |
| chr15 | 92696569 | 92697069 | 0 | 1 | 0 | 0 | 1 | 0 | 0 |

| | | | | | | | | | |
|---|---|---|---|---|---|---|---|---|---|
| chr15 | 71782070 | 71782570 | 1 | 0 | 0 | 0 | 1 | 0 | 1 |
| chr15 | 92750069 | 92750569 | 1 | 0 | 0 | 0 | 1 | 0 | 1 |
| chr16 | 84485755 | 84486255 | 1 | 0 | 0 | 1 | 1 | 0 | 0 |
| chr16 | 59284174 | 59284674 | 1 | 0 | 0 | 0 | 1 | 0 | 1 |
| chr16 | 95026393 | 95026893 | 1 | 1 | 1 | 0 | 1 | 0 | 1 |
| chr17 | 41079051 | 41079551 | 1 | 1 | 1 | 0 | 1 | 0 | 0 |
| chr17 | 14094993 | 14095493 | 1 | 0 | 0 | 0 | 1 | 0 | 1 |
| chr18 | 37341346 | 37341846 | 1 | 0 | 0 | 0 | 1 | 0 | 1 |
| chr18 | 72929846 | 72930346 | 1 | 0 | 0 | 0 | 1 | 0 | 0 |
| chr18 | 47646346 | 47646846 | 1 | 1 | 1 | 1 | 1 | 0 | 1 |
| chr18 | 37692346 | 37692846 | 1 | 1 | 1 | 0 | 1 | 0 | 1 |
| | | **Total** | **66** | **51** | **31** | **16** | **86** | **7** | **69** |

Chr is chromosome; TR is thyroid hormone receptor; uTRs is unoccupied TR, as determined in liver ChIP-seq data obtained in hypothyroid mice, present within 10,000 bp of H-site; T3-TRs is TR occupied with T3, as determined in liver ChIP-seq data obtained in T3-treated mice, present within 10,000 bp of H-site. The presence or absence of the uTR or TR ChIP-seq peak was determined by visual confirmation on the IGV browser, upstream or downstream of each H-site location.

**Table S2** - H-sites chromosome coordinates and gene annotation.

| Chr | start | stop | Gene Symbol | Gene Section | Dist. to TSS (bp) |
|---|---|---|---|---|---|
| 1 | 4766919 | 4767419 | Gm6123 | Promoter | -3712 |
| 1 | 21217919 | 21218419 | Tmem14a | Promoter | -156 |
| 1 | 21217919 | 21218419 | Tmem14a | Promoter | -243 |
| 1 | 21271419 | 21271919 | Gm28836 | Promoter | 0 |
| 1 | 21271419 | 21271919 | Gm7094 | Promoter | -484 |
| 1 | 24746155 | 24746655 | Lmbrd1 | Intron 12 | 67525 |
| 1 | 24746155 | 24746655 | Lmbrd1 | Intron 12 | 67229 |
| 1 | 24746155 | 24746655 | Lmbrd1 | Promoter | 1282 |
| 1 | 24746155 | 24746655 | Lmbrd1 | Intron 1 | 1282 |
| 1 | 25432155 | 25432655 | Adgrb3 | Intron 14 | 396857 |
| 1 | 25432155 | 25432655 | Adgrb3 | CDS 14 | 396857 |
| 1 | 25432155 | 25432655 | Adgrb3 | Intron 13 | 396857 |
| 1 | 25432155 | 25432655 | Adgrb3 | Intron 12 | 393485 |
| 1 | 25432155 | 25432655 | Adgrb3 | CDS 12 | 393485 |
| 1 | 25432155 | 25432655 | Adgrb3 | Intron 11 | 393485 |
| 1 | 25432155 | 25432655 | Adgrb3 | Intron 13 | 397053 |
| 1 | 25432155 | 25432655 | Adgrb3 | CDS 13 | 397053 |
| 1 | 25432155 | 25432655 | Adgrb3 | Intron 12 | 397053 |
| 1 | 25432155 | 25432655 | Adgrb3 | Intron 12 | 394106 |
| 1 | 25432155 | 25432655 | Adgrb3 | CDS 12 | 394106 |
| 1 | 25432155 | 25432655 | Adgrb3 | Intron 11 | 394106 |
| 1 | 40117655 | 40118155 | Il1r2 | Intron 5 | 32957 |
| 1 | 40117655 | 40118155 | Il1r2 | CDS 6 | 32957 |
| 1 | 40117655 | 40118155 | Il1r2 | Intron 6 | 32957 |
| 1 | 40117655 | 40118155 | Il1r2 | 3' UTR | 32863 |
| 1 | 51747656 | 51748156 | Gm28055 | Promoter | -712 |
| 1 | 52632656 | 52633156 | Nemp2 | Promoter | 1953 |
| 1 | 52632656 | 52633156 | Nemp2 | Intron 1 | 1953 |
| 1 | 52632656 | 52633156 | Nemp2 | Promoter | 1919 |
| 1 | 52632656 | 52633156 | Nemp2 | Intron 1 | 1919 |
| 1 | 52632656 | 52633156 | Nemp2 | Promoter | 1909 |
| 1 | 52632656 | 52633156 | Nemp2 | Intron 1 | 1909 |
| 1 | 52632656 | 52633156 | Nemp2 | Promoter | 1904 |
| 1 | 52632656 | 52633156 | Nemp2 | Promoter | 1859 |
| 1 | 52632656 | 52633156 | Nemp2 | Intron 1 | 1859 |
| 1 | 53977156 | 53977656 | Hecw2 | Intron 2 | 216393 |
| 1 | 53977156 | 53977656 | Hecw2 | Intron 2 | 217379 |
| 1 | 53977156 | 53977656 | Hecw2 | Intron 2 | 217379 |
| 1 | 53977156 | 53977656 | Hecw2 | Non-coding | 217513 |
| 1 | 55362156 | 55362656 | Boll | Promoter | -1414 |
| 1 | 55362156 | 55362656 | Boll | Promoter | 664 |
| 1 | 55362156 | 55362656 | Boll | Intron 1 | 664 |

| | | | | | |
|---|---|---|---|---|---|
| 1 | 55362156 | 55362656 | Boll | Promoter | 71 |
| 1 | 55362156 | 55362656 | Boll | 5' UTR | 71 |
| 1 | 55362156 | 55362656 | Boll | Promoter | 664 |
| 1 | 55362156 | 55362656 | Boll | Intron 1 | 664 |
| 1 | 55362156 | 55362656 | Boll | Promoter | 814 |
| 1 | 55362156 | 55362656 | Boll | Intron 1 | 814 |
| 1 | 55362156 | 55362656 | Boll | Promoter | -1414 |
| 1 | 64288926 | 64289426 | Gm28981 | Non-coding | 13618 |
| 1 | 74542426 | 74542926 | Usp37 | Promoter | 1359 |
| 1 | 74542426 | 74542926 | Usp37 | 5' UTR | 1359 |
| 1 | 74542426 | 74542926 | Plcd4 | Promoter | 0 |
| 1 | 74542426 | 74542926 | Plcd4 | 5' UTR | 0 |
| 1 | 74542426 | 74542926 | Plcd4 | Promoter | 0 |
| 1 | 74542426 | 74542926 | Plcd4 | 5' UTR | 0 |
| 1 | 74542426 | 74542926 | Plcd4 | Promoter | 0 |
| 1 | 74542426 | 74542926 | Plcd4 | 5' UTR | 0 |
| 1 | 74542426 | 74542926 | Plcd4 | Promoter | 0 |
| 1 | 74542426 | 74542926 | Plcd4 | Promoter | 0 |
| 1 | 74542426 | 74542926 | Plcd4 | 5' UTR | 0 |
| 1 | 74542426 | 74542926 | Plcd4 | Promoter | 0 |
| 1 | 74542426 | 74542926 | Plcd4 | 5' UTR | 0 |
| 1 | 74542426 | 74542926 | Plcd4 | Promoter | -1434 |
| 1 | 74542426 | 74542926 | Plcd4 | Promoter | -1438 |
| 1 | 74542426 | 74542926 | Plcd4 | Promoter | -1438 |
| 1 | 77565925 | 77566425 | Gm28385 | Promoter | -4247 |
| 1 | 83722425 | 83722925 | 4933436I20Rik | Non-coding | 188647 |
| 1 | 88222425 | 88222925 | Gm20528 | Promoter | -3840 |
| 1 | 88222425 | 88222925 | Gm20528 | Promoter | -3870 |
| 1 | 88222425 | 88222925 | Mroh2a | Promoter | -4061 |
| 1 | 88222425 | 88222925 | Mroh2a | Promoter | -4095 |
| 1 | 88222425 | 88222925 | Mroh2a | Promoter | -4106 |
| 1 | 88447425 | 88447925 | Gm29538 | Non-coding | 14684 |
| 1 | 91425923 | 91426423 | Per2 | Intron 18 | 32902 |
| 1 | 92478423 | 92478923 | Ndufa10 | Promoter | -4562 |
| 1 | 92478423 | 92478923 | Ndufa10 | Promoter | -4665 |
| 1 | 92478423 | 92478923 | Ndufa10 | Promoter | -4675 |
| 1 | 92478423 | 92478923 | Ndufa10 | Promoter | -4655 |
| 1 | 92478423 | 92478923 | Olfr1416 | 3' UTR | 6191 |
| 1 | 92478423 | 92478923 | Olfr1416 | Promoter | 1697 |
| 1 | 93832423 | 93832923 | D2hgdh | Intron 4 | 7183 |
| 1 | 93832423 | 93832923 | D2hgdh | CDS 5 | 7183 |
| 1 | 93832423 | 93832923 | D2hgdh | Intron 5 | 7183 |
| 1 | 93832423 | 93832923 | D2hgdh | Intron 4 | 7149 |
| 1 | 93832423 | 93832923 | D2hgdh | CDS 5 | 7149 |
| 1 | 93832423 | 93832923 | D2hgdh | Intron 5 | 7149 |
| 1 | 93832423 | 93832923 | D2hgdh | Intron 3 | 6138 |

| | | | | | |
|---|---|---|---|---|---|
| 1 | 93832423 | 93832923 | D2hgdh | Promoter | 2627 |
| 1 | 93832423 | 93832923 | D2hgdh | Intron 1 | 2627 |
| 1 | 93832423 | 93832923 | D2hgdh | Promoter | 2626 |
| 1 | 93832423 | 93832923 | D2hgdh | Intron 1 | 2626 |
| 1 | 95627423 | 95627923 | St8sia4 | Intron 4 | 39649 |
| 1 | 95627423 | 95627923 | St8sia4 | CDS 4 | 39649 |
| 1 | 95627423 | 95627923 | St8sia4 | Intron 3 | 39649 |
| 1 | 95627423 | 95627923 | St8sia4 | 3' UTR | 39613 |
| 1 | 95627423 | 95627923 | St8sia4 | CDS 4 | 39613 |
| 1 | 95627423 | 95627923 | St8sia4 | Intron 3 | 39613 |
| 1 | 118551423 | 118551923 | Clasp1 | Intron 29 | 162365 |
| 1 | 118551423 | 118551923 | Clasp1 | Intron 31 | 162365 |
| 1 | 118551423 | 118551923 | Clasp1 | Intron 27 | 162365 |
| 1 | 118551423 | 118551923 | Clasp1 | Intron 31 | 162365 |
| 1 | 118551423 | 118551923 | Clasp1 | 3' UTR | 161977 |
| 1 | 118551423 | 118551923 | Clasp1 | Intron 27 | 161968 |
| 1 | 118551423 | 118551923 | Clasp1 | Intron 29 | 161968 |
| 1 | 118551423 | 118551923 | Clasp1 | Intron 30 | 161968 |
| 1 | 118551423 | 118551923 | Clasp1 | Intron 28 | 161967 |
| 1 | 118551423 | 118551923 | Clasp1 | Intron 29 | 161950 |
| 1 | 118551423 | 118551923 | Clasp1 | Intron 28 | 161906 |
| 1 | 118551423 | 118551923 | Clasp1 | Intron 28 | 131863 |
| 1 | 118551423 | 118551923 | Clasp1 | Intron 29 | 131699 |
| 1 | 118551423 | 118551923 | Clasp1 | Intron 30 | 131699 |
| 1 | 118551423 | 118551923 | Clasp1 | Non-coding | 81652 |
| 1 | 118551423 | 118551923 | Clasp1 | Intron 23 | 69384 |
| 1 | 118551423 | 118551923 | Clasp1 | 3' UTR | 19344 |
| 1 | 118551423 | 118551923 | Clasp1 | Intron 2 | 3446 |
| 1 | 118551423 | 118551923 | Clasp1 | Promoter | -2311 |
| 1 | 121320423 | 121320923 | Insig2 | 5' UTR | 7102 |
| 1 | 121325923 | 121326423 | Insig2 | Promoter | 1602 |
| 1 | 121325923 | 121326423 | Insig2 | 5' UTR | 1602 |
| 1 | 121320423 | 121320923 | Insig2 | 5' UTR | 11667 |
| 1 | 121325923 | 121326423 | Insig2 | 5' UTR | 6167 |
| 1 | 121320423 | 121320923 | Insig2 | 5' UTR | 6824 |
| 1 | 121325923 | 121326423 | Insig2 | Promoter | 1324 |
| 1 | 121325923 | 121326423 | Insig2 | 5' UTR | 1324 |
| 1 | 121320423 | 121320923 | Insig2 | 5' UTR | 6756 |
| 1 | 121325923 | 121326423 | Insig2 | Promoter | 1256 |
| 1 | 121325923 | 121326423 | Insig2 | 5' UTR | 1256 |
| 1 | 121320423 | 121320923 | Insig2 | 5' UTR | 6797 |
| 1 | 121325923 | 121326423 | Insig2 | Promoter | 1297 |
| 1 | 121325923 | 121326423 | Insig2 | 5' UTR | 1297 |
| 1 | 121320423 | 121320923 | Insig2 | 5' UTR | 6871 |
| 1 | 121325923 | 121326423 | Insig2 | Promoter | 1371 |
| 1 | 121325923 | 121326423 | Insig2 | 5' UTR | 1371 |

| | | | | | |
|---|---|---|---|---|---|
| 1 | 121320423 | 121320923 | Insig2 | 5' UTR | 6843 |
| 1 | 121325923 | 121326423 | Insig2 | Promoter | 1343 |
| 1 | 121325923 | 121326423 | Insig2 | 5' UTR | 1343 |
| 1 | 121320423 | 121320923 | Insig2 | Promoter | 2269 |
| 1 | 121320423 | 121320923 | Insig2 | 5' UTR | 2269 |
| 1 | 121325923 | 121326423 | Insig2 | Promoter | -2731 |
| 1 | 121320423 | 121320923 | Insig2 | Promoter | -694 |
| 1 | 121320423 | 121320923 | Insig2 | 5' UTR | 6760 |
| 1 | 121325923 | 121326423 | Insig2 | Promoter | 1260 |
| 1 | 121325923 | 121326423 | Insig2 | 5' UTR | 1260 |
| 1 | 121320423 | 121320923 | Insig2 | 5' UTR | 6766 |
| 1 | 121325923 | 121326423 | Insig2 | Promoter | 1266 |
| 1 | 121325923 | 121326423 | Insig2 | 5' UTR | 1266 |
| 1 | 121320423 | 121320923 | Insig2 | 5' UTR | 11628 |
| 1 | 121325923 | 121326423 | Insig2 | 5' UTR | 6128 |
| 1 | 121325923 | 121326423 | Gm37174 | Promoter | -1669 |
| 1 | 133237923 | 133238423 | Plekha6 | Intron 3 | 73713 |
| 1 | 133237923 | 133238423 | Plekha6 | 5' UTR | 56602 |
| 1 | 133914923 | 133915423 | Prelp | 5' UTR | 5992 |
| 1 | 133914923 | 133915423 | Prelp | CDS 2 | 5992 |
| 1 | 133914923 | 133915423 | Prelp | Intron 1 | 5992 |
| 1 | 133914923 | 133915423 | Prelp | Non-coding | 5989 |
| 1 | 135585923 | 135586423 | Nav1 | Promoter | -567 |
| 1 | 135585923 | 135586423 | Nav1 | Promoter | -217 |
| 1 | 135674923 | 135675423 | Nav1 | 5' UTR | 12683 |
| 1 | 135585923 | 135586423 | Nav1 | Intron 3 | 101683 |
| 1 | 135585923 | 135586423 | Gm4793 | Promoter | 1150 |
| 1 | 135674923 | 135675423 | Nav1 | Non-coding | 12683 |
| 1 | 135874423 | 135874923 | Pkp1 | 3' UTR | 44285 |
| 1 | 135874423 | 135874923 | Pkp1 | 3' UTR | 44198 |
| 1 | 135874423 | 135874923 | Pkp1 | 3' UTR | 44220 |
| 1 | 150145870 | 150146370 | Gm9687 | Promoter | 0 |
| 1 | 151861870 | 151862370 | 1700025G04Rik | 3' UTR | 227601 |
| 1 | 153904870 | 153905370 | Teddm2 | Promoter | -4661 |
| 1 | 153904870 | 153905370 | Teddm2 | Promoter | -4641 |
| 1 | 153904870 | 153905370 | Glul | Intron 3 | 4926 |
| 1 | 153904870 | 153905370 | Glul | Intron 3 | 4285 |
| 1 | 153904870 | 153905370 | Glul | Intron 3 | 4264 |
| 1 | 153904870 | 153905370 | Glul | Promoter | 1072 |
| 1 | 153904870 | 153905370 | Glul | Promoter | -673 |
| 1 | 154688870 | 154689370 | Cacna1e | Intron 3 | 36987 |
| 1 | 154688870 | 154689370 | Cacna1e | 5' UTR | 36551 |
| 1 | 154688870 | 154689370 | Cacna1e | Intron 2 | 174698 |
| 1 | 154688870 | 154689370 | Cacna1e | Non-coding | 127572 |
| 1 | 154688870 | 154689370 | Cacna1e | Intron 5 | 127572 |

| | | | | | |
|---|---|---|---|---|---|
| 1 | 155019370 | 155019870 | Gm29282 | Non-coding | 37310 |
| 1 | 156642369 | 156642869 | Abl2 | 3' UTR | 83382 |
| 1 | 156642369 | 156642869 | Abl2 | CDS 13 | 83382 |
| 1 | 156642369 | 156642869 | Abl2 | 3' UTR | 83320 |
| 1 | 156642369 | 156642869 | Abl2 | CDS 12 | 83320 |
| 1 | 158509869 | 158510369 | Astn1 | CDS 6 | 147596 |
| 1 | 158509869 | 158510369 | Astn1 | Intron 6 | 147596 |
| 1 | 158509869 | 158510369 | Astn1 | CDS 6 | 147565 |
| 1 | 158509869 | 158510369 | Astn1 | Intron 6 | 147565 |
| 1 | 158509869 | 158510369 | Astn1 | CDS 6 | 147529 |
| 1 | 158509869 | 158510369 | Astn1 | Intron 6 | 147529 |
| 1 | 158509869 | 158510369 | Astn1 | CDS 6 | 147332 |
| 1 | 158509869 | 158510369 | Astn1 | Intron 6 | 147332 |
| 1 | 158509869 | 158510369 | Astn1 | CDS 6 | 147319 |
| 1 | 158509869 | 158510369 | Astn1 | Intron 6 | 147319 |
| 1 | 158509869 | 158510369 | Gm37679 | Non-coding | 43931 |
| 1 | 158509869 | 158510369 | Astn1 | Promoter | 0 |
| 1 | 160041369 | 160041869 | Gm37294 | Promoter | 2463 |
| 1 | 160041369 | 160041869 | 4930523C07Rik | Promoter | -2513 |
| 1 | 160041369 | 160041869 | 4930523C07Rik | Promoter | -2515 |
| 1 | 160041369 | 160041869 | 4930523C07Rik | Promoter | -2515 |
| 1 | 160041369 | 160041869 | 4930523C07Rik | Promoter | -2536 |
| 1 | 160041369 | 160041869 | 4930523C07Rik | Promoter | -2578 |
| 1 | 160988869 | 160989369 | Serpinc1 | 5' UTR | 10263 |
| 1 | 160988869 | 160989369 | Serpinc1 | Intron 2 | 10223 |
| 1 | 160988869 | 160989369 | Serpinc1 | 5' UTR | 10217 |
| 1 | 160988869 | 160989369 | Serpinc1 | Intron 2 | 10181 |
| 1 | 160988869 | 160989369 | Serpinc1 | Intron 2 | 4101 |
| 1 | 160988869 | 160989369 | Serpinc1 | Promoter | 1782 |
| 1 | 160988869 | 160989369 | Serpinc1 | Promoter | -8 |
| 1 | 164065869 | 164066369 | Sell | Intron 3 | 3887 |
| 1 | 164065869 | 164066369 | Sell | Intron 3 | 3784 |
| 1 | 164065869 | 164066369 | Sell | CDS 4 | 3784 |
| 1 | 164065869 | 164066369 | Sell | Intron 4 | 3784 |
| 1 | 164065869 | 164066369 | Sell | Intron 3 | 3680 |
| 1 | 164065869 | 164066369 | Sell | CDS 4 | 3680 |
| 1 | 164065869 | 164066369 | Sell | Intron 4 | 3680 |
| 1 | 164065869 | 164066369 | Sell | Promoter | -2593 |
| 1 | 164259369 | 164259869 | Slc19a2 | Intron 2 | 10323 |
| 1 | 164259369 | 164259869 | Slc19a2 | Intron 2 | 10314 |
| 1 | 164259369 | 164259869 | Slc19a2 | Intron 1 | 10116 |
| 1 | 164259369 | 164259869 | Slc19a2 | Promoter | -2484 |
| 1 | 165237869 | 165238369 | Tiprl | Promoter | -925 |
| 1 | 165237869 | 165238369 | Tiprl | Promoter | -872 |
| 1 | 165237869 | 165238369 | Tiprl | Promoter | -946 |
| 1 | 165237869 | 165238369 | Tiprl | Promoter | -1355 |

| | | | | | |
|---|---|---|---|---|---|
| 1 | 165237869 | 165238369 | Tiprl | Promoter | -936 |
| 1 | 165237869 | 165238369 | Gm38190 | Promoter | -2743 |
| 1 | 166286369 | 166286869 | Ildr2 | Intron 3 | 32230 |
| 1 | 166296369 | 166296869 | Ildr2 | Intron 6 | 42230 |
| 1 | 166286369 | 166286869 | Ildr2 | Intron 3 | 32176 |
| 1 | 166296369 | 166296869 | Ildr2 | Intron 3 | 42176 |
| 1 | 166286369 | 166286869 | Ildr2 | Intron 3 | 32176 |
| 1 | 166296369 | 166296869 | Ildr2 | Intron 5 | 42176 |
| 1 | 166286369 | 166286869 | Ildr2 | Intron 3 | 32176 |
| 1 | 166296369 | 166296869 | Ildr2 | Intron 3 | 42176 |
| 1 | 166286369 | 166286869 | Ildr2 | Intron 3 | 32176 |
| 1 | 166296369 | 166296869 | Ildr2 | Intron 4 | 42176 |
| 1 | 166286369 | 166286869 | Ildr2 | Intron 3 | 32176 |
| 1 | 166296369 | 166296869 | Ildr2 | Intron 5 | 42176 |
| 1 | 166286369 | 166286869 | Ildr2 | Intron 3 | 32176 |
| 1 | 166296369 | 166296869 | Ildr2 | Intron 6 | 42176 |
| 1 | 167079369 | 167079869 | Fam78b | 3' UTR | 77952 |
| 1 | 167079369 | 167079869 | Fam78b | 3' UTR | 77952 |
| 1 | 167079369 | 167079869 | Fam78b | 3' UTR | 77937 |
| 1 | 167079369 | 167079869 | Fam78b | 3' UTR | 77903 |
| 1 | 167079369 | 167079869 | Fam78b | 3' UTR | 77789 |
| 1 | 167079369 | 167079869 | Fam78b | 3' UTR | 77780 |
| 1 | 171238869 | 171239369 | Fcer1g | Promoter | -4503 |
| 1 | 171238869 | 171239369 | Fcer1g | Promoter | -4561 |
| 1 | 171238869 | 171239369 | Fcer1g | Promoter | -4529 |
| 1 | 171238869 | 171239369 | Ndufs2 | CDS 5 | 7754 |
| 1 | 171238869 | 171239369 | Ndufs2 | Intron 4 | 7754 |
| 1 | 171238869 | 171239369 | Ndufs2 | CDS 4 | 7754 |
| 1 | 171238869 | 171239369 | Ndufs2 | CDS 5 | 7754 |
| 1 | 171238869 | 171239369 | Ndufs2 | Intron 4 | 7754 |
| 1 | 171238869 | 171239369 | Ndufs2 | CDS 4 | 7754 |
| 1 | 171238869 | 171239369 | Ndufs2 | Non-coding | 7754 |
| 1 | 171238869 | 171239369 | Ndufs2 | Promoter | -3328 |
| 1 | 171238869 | 171239369 | Ndufs2 | Promoter | -1340 |
| 1 | 171238869 | 171239369 | Ndufs2 | Promoter | 687 |
| 1 | 171238869 | 171239369 | Ndufs2 | CDS 5 | 11883 |
| 1 | 171238869 | 171239369 | Ndufs2 | Intron 4 | 11883 |
| 1 | 171238869 | 171239369 | Ndufs2 | CDS 4 | 11883 |
| 1 | 171238869 | 171239369 | Ndufs2 | CDS 4 | 12020 |
| 1 | 171238869 | 171239369 | Ndufs2 | Intron 3 | 12020 |
| 1 | 171238869 | 171239369 | Ndufs2 | CDS 3 | 12020 |
| 1 | 177446869 | 177447369 | Gm26801 | Promoter | -3598 |
| 1 | 177446869 | 177447369 | Zbtb18 | 5' UTR | 4518 |
| 1 | 177446869 | 177447369 | Zbtb18 | Intron 1 | 4518 |
| 1 | 177446869 | 177447369 | Zbtb18 | CDS 2 | 4518 |
| 1 | 177446869 | 177447369 | Zbtb18 | 5' UTR | 4403 |

| | | | | | |
|---|---|---|---|---|---|
| 1 | 177446869 | 177447369 | Zbtb18 | Intron 2 | 4403 |
| 1 | 177446869 | 177447369 | Zbtb18 | CDS 3 | 4403 |
| 1 | 177446869 | 177447369 | Zbtb18 | 5' UTR | 4325 |
| 1 | 177446869 | 177447369 | Zbtb18 | Intron 3 | 4325 |
| 1 | 177446869 | 177447369 | Zbtb18 | CDS 4 | 4325 |
| 1 | 177446869 | 177447369 | Zbtb18 | Intron 2 | 3653 |
| 1 | 177446869 | 177447369 | Zbtb18 | CDS 3 | 3653 |
| 1 | 177446869 | 177447369 | Zbtb18 | Promoter | 2574 |
| 1 | 177446869 | 177447369 | Zbtb18 | Intron 1 | 2574 |
| 1 | 177446869 | 177447369 | Zbtb18 | CDS 2 | 2574 |
| 1 | 177446869 | 177447369 | Zbtb18 | Promoter | 2220 |
| 1 | 177446869 | 177447369 | Zbtb18 | 5' UTR | 2220 |
| 1 | 177446869 | 177447369 | Zbtb18 | Intron 1 | 2220 |
| 1 | 177446869 | 177447369 | Zbtb18 | CDS 2 | 2220 |
| 1 | 177446869 | 177447369 | Zbtb18 | Promoter | 2216 |
| 1 | 177446869 | 177447369 | Zbtb18 | Intron 1 | 2216 |
| 1 | 177446869 | 177447369 | Zbtb18 | CDS 2 | 2216 |
| 1 | 177446869 | 177447369 | Zbtb18 | Promoter | 2208 |
| 1 | 177446869 | 177447369 | Zbtb18 | 5' UTR | 2208 |
| 1 | 177446869 | 177447369 | Zbtb18 | Intron 1 | 2208 |
| 1 | 177446869 | 177447369 | Zbtb18 | CDS 2 | 2208 |
| 1 | 177446869 | 177447369 | Zbtb18 | Promoter | 2132 |
| 1 | 177446869 | 177447369 | Zbtb18 | 5' UTR | 2132 |
| 1 | 177446869 | 177447369 | Zbtb18 | Intron 2 | 2132 |
| 1 | 177446869 | 177447369 | Zbtb18 | CDS 3 | 2132 |
| 1 | 177446869 | 177447369 | Zbtb18 | Promoter | 1931 |
| 1 | 177446869 | 177447369 | Zbtb18 | 5' UTR | 1931 |
| 1 | 177446869 | 177447369 | Zbtb18 | Intron 1 | 1931 |
| 1 | 177446869 | 177447369 | Zbtb18 | CDS 2 | 1931 |
| 1 | 177446869 | 177447369 | Zbtb18 | Promoter | 1238 |
| 1 | 177446869 | 177447369 | Zbtb18 | Promoter | 1048 |
| 1 | 177446869 | 177447369 | Zbtb18 | 5' UTR | 1048 |
| 1 | 177446869 | 177447369 | Zbtb18 | CDS 1 | 1048 |
| 1 | 178419869 | 178420369 | Efcab2 | Non-coding | 13784 |
| 1 | 178419869 | 178420369 | Efcab2 | Intron 1 | 13766 |
| 1 | 179157369 | 179157869 | Smyd3 | Intron 5 | 360166 |
| 1 | 179157369 | 179157869 | Smyd3 | Non-coding | 90972 |
| 1 | 179157369 | 179157869 | Smyd3 | Non-coding | 216235 |
| 1 | 179712369 | 179712869 | Kif28 | Intron 11 | 32403 |
| 1 | 179712369 | 179712869 | Kif28 | CDS 11 | 32403 |
| 1 | 179712369 | 179712869 | Kif28 | Intron 10 | 32403 |
| 1 | 179712369 | 179712869 | Kif28 | CDS 10 | 32403 |
| 1 | 179733369 | 179733869 | Kif28 | Intron 4 | 11403 |
| 1 | 179712369 | 179712869 | Kif28 | Intron 11 | 32403 |
| 1 | 179712369 | 179712869 | Kif28 | CDS 11 | 32403 |
| 1 | 179712369 | 179712869 | Kif28 | Intron 10 | 32403 |

| | | | | | |
|---|---|---|---|---|---|
| 1 | 179712369 | 179712869 | Kif28 | CDS 10 | 32403 |
| 1 | 179733369 | 179733869 | Kif28 | Intron 4 | 11403 |
| 1 | 179712369 | 179712869 | Kif28 | Intron 9 | 32332 |
| 1 | 179733369 | 179733869 | Kif28 | Intron 4 | 11332 |
| 1 | 180829869 | 180830369 | Gm24836 | Promoter | -2400 |
| 1 | 181510369 | 181510869 | Ccdc121 | Promoter | 583 |
| 1 | 181510369 | 181510869 | Ccdc121 | 3' UTR | 583 |
| 1 | 181510369 | 181510869 | Ccdc121 | CDS 1 | 583 |
| 1 | 182283369 | 182283869 | Degs1 | Promoter | -564 |
| 1 | 182283369 | 182283869 | Degs1 | Promoter | -564 |
| 1 | 182283369 | 182283869 | Degs1 | Promoter | -1144 |
| 1 | 182283369 | 182283869 | 1700047M11Rik | Promoter | -4121 |
| 1 | 182883369 | 182883869 | Susd4 | Intron 5 | 119357 |
| 1 | 182883369 | 182883869 | Susd4 | Intron 4 | 118474 |
| 1 | 188902621 | 188903121 | Ush2a | Intron 60 | 639587 |
| 1 | 192846806 | 192847306 | Sertad4 | CDS 4 | 8451 |
| 1 | 192846806 | 192847306 | Sertad4 | Intron 3 | 8451 |
| 1 | 192846806 | 192847306 | Sertad4 | CDS 4 | 4442 |
| 1 | 192846806 | 192847306 | Sertad4 | Intron 3 | 4442 |
| 1 | 192846806 | 192847306 | Sertad4 | 3' UTR | 4442 |
| 1 | 192846806 | 192847306 | Sertad4 | CDS 4 | 4442 |
| 1 | 192846806 | 192847306 | Sertad4 | Intron 3 | 4442 |
| 1 | 195017806 | 195018306 | A330023F24Rik | Non-coding | 41431 |
| 1 | 195017806 | 195018306 | A330023F24Rik | Non-coding | 41421 |
| 1 | 195017806 | 195018306 | A330023F24Rik | Promoter | 408 |
| 1 | 195017806 | 195018306 | Gm37027 | Promoter | 0 |
| 10 | 5599540 | 5600040 | Myct1 | Intron 1 | 5765 |
| 10 | 5920540 | 5921040 | Rgs17 | Promoter | 1361 |
| 10 | 5920540 | 5921040 | Rgs17 | 5' UTR | 1361 |
| 10 | 5920540 | 5921040 | Rgs17 | Promoter | 1347 |
| 10 | 5920540 | 5921040 | Rgs17 | 5' UTR | 1347 |
| 10 | 5920540 | 5921040 | Rgs17 | Promoter | 1327 |
| 10 | 5920540 | 5921040 | Rgs17 | 5' UTR | 1327 |
| 10 | 5920540 | 5921040 | Rgs17 | Promoter | 1327 |
| 10 | 5920540 | 5921040 | Rgs17 | 5' UTR | 1327 |
| 10 | 10395202 | 10395702 | Adgb | Intron 22 | 76613 |
| 10 | 10395202 | 10395702 | Adgb | CDS 22 | 76613 |
| 10 | 10395202 | 10395702 | Adgb | Intron 21 | 76613 |
| 10 | 10395202 | 10395702 | Adgb | 3' UTR | 16043 |
| 10 | 10395202 | 10395702 | Adgb | Intron 21 | 76625 |
| 10 | 10395202 | 10395702 | Adgb | CDS 21 | 76625 |
| 10 | 10395202 | 10395702 | Adgb | Intron 20 | 76625 |
| 10 | 10395202 | 10395702 | Adgb | Intron 20 | 76613 |
| 10 | 10395202 | 10395702 | Adgb | CDS 20 | 76613 |
| 10 | 10395202 | 10395702 | Adgb | Intron 19 | 76613 |

| | | | | | |
|---|---|---|---|---|---|
| 10 | 17632194 | 17632694 | Gm47771 | Non-coding | 25820 |
| 10 | 18526194 | 18526694 | Nhsl1 | CDS 5 | 207209 |
| 10 | 18526194 | 18526694 | Nhsl1 | CDS 5 | 118519 |
| 10 | 18526194 | 18526694 | Nhsl1 | CDS 6 | 56213 |
| 10 | 20147694 | 20148194 | Map7 | Promoter | -277 |
| 10 | 20195194 | 20195694 | Map7 | Intron 1 | 46723 |
| 10 | 20147694 | 20148194 | Map7 | Promoter | -790 |
| 10 | 20195194 | 20195694 | Map7 | Intron 1 | 46210 |
| 10 | 20147694 | 20148194 | Map7 | Promoter | -801 |
| 10 | 20195194 | 20195694 | Map7 | Intron 1 | 46199 |
| 10 | 20147694 | 20148194 | Map7 | Promoter | -804 |
| 10 | 20195194 | 20195694 | Map7 | Non-coding | 46196 |
| 10 | 20195194 | 20195694 | Map7 | Non-coding | 24206 |
| 10 | 20195194 | 20195694 | Map7 | Intron 1 | 24172 |
| 10 | 20195194 | 20195694 | Map7 | Non-coding | 23488 |
| 10 | 20195194 | 20195694 | Map7 | Non-coding | 23187 |
| 10 | 21442194 | 21442694 | Gm48386 | Promoter | 2943 |
| 10 | 30845694 | 30846194 | Hey2 | Promoter | -2892 |
| 10 | 30845694 | 30846194 | Hey2 | Promoter | -3678 |
| 10 | 39386194 | 39386694 | Fyn | 5' UTR | 17339 |
| 10 | 39386194 | 39386694 | Fyn | 5' UTR | 16430 |
| 10 | 39386194 | 39386694 | Fyn | 5' UTR | 16396 |
| 10 | 39386194 | 39386694 | Fyn | 5' UTR | 16391 |
| 10 | 40810194 | 40810694 | Mettl24 | Intron 4 | 126912 |
| 10 | 40810194 | 40810694 | Mettl24 | CDS 5 | 126912 |
| 10 | 42760194 | 42760694 | Gm15199 | Promoter | -1533 |
| 10 | 42760194 | 42760694 | Sec63 | Promoter | -802 |
| 10 | 42760194 | 42760694 | Sec63 | Promoter | -921 |
| 10 | 42760194 | 42760694 | Sec63 | Promoter | -1112 |
| 10 | 43444194 | 43444694 | Pdss2 | Intron 7 | 222594 |
| 10 | 58254252 | 58254752 | Gcc2 | Promoter | -745 |
| 10 | 58254252 | 58254752 | Gcc2 | Promoter | -756 |
| 10 | 58254252 | 58254752 | Gcc2 | Promoter | -778 |
| 10 | 58254252 | 58254752 | Gcc2 | Promoter | -787 |
| 10 | 58254252 | 58254752 | Gcc2 | Promoter | -961 |
| 10 | 58254252 | 58254752 | Gcc2 | Promoter | -1139 |
| 10 | 58532752 | 58533252 | Ccdc138 | Intron 10 | 34804 |
| 10 | 59089752 | 59090252 | Sh3rf3 | Intron 7 | 276294 |
| 10 | 59089752 | 59090252 | Sh3rf3 | Non-coding | 24101 |
| 10 | 59089752 | 59090252 | Sh3rf3 | Non-coding | 9718 |
| 10 | 59447252 | 59447752 | Mcu | 3' UTR | 168941 |
| 10 | 60814752 | 60815252 | Unc5b | Intron 1 | 16330 |
| 10 | 60814752 | 60815252 | Unc5b | Non-coding | 5390 |
| 10 | 60814752 | 60815252 | Unc5b | Intron 1 | 16131 |
| 10 | 60814752 | 60815252 | Gm19972 | Promoter | -428 |
| 10 | 62126752 | 62127252 | Fam241b | Intron 2 | 12619 |

| | | | | | |
|---|---|---|---|---|---|
| 10 | 62126752 | 62127252 | Fam241b | 5' UTR | 7455 |
| 10 | 62126752 | 62127252 | Fam241b | Intron 2 | 16655 |
| 10 | 62126752 | 62127252 | Fam241b | Intron 3 | 16659 |
| 10 | 62295752 | 62296252 | Hk1 | CDS 7 | 44291 |
| 10 | 62295752 | 62296252 | Hk1 | Intron 6 | 44291 |
| 10 | 62295752 | 62296252 | Hk1 | CDS 6 | 44291 |
| 10 | 62295752 | 62296252 | Hk1 | Intron 5 | 44291 |
| 10 | 62295752 | 62296252 | Hk1 | CDS 10 | 83657 |
| 10 | 62295752 | 62296252 | Hk1 | Intron 9 | 83657 |
| 10 | 62295752 | 62296252 | Hk1 | CDS 9 | 83657 |
| 10 | 62295752 | 62296252 | Hk1 | Intron 8 | 83657 |
| 10 | 62295752 | 62296252 | Hk1 | CDS 8 | 62111 |
| 10 | 62295752 | 62296252 | Hk1 | Intron 7 | 62111 |
| 10 | 62295752 | 62296252 | Hk1 | CDS 7 | 62111 |
| 10 | 62295752 | 62296252 | Hk1 | Intron 6 | 62111 |
| 10 | 62295752 | 62296252 | Hk1 | CDS 7 | 46471 |
| 10 | 62295752 | 62296252 | Hk1 | Intron 6 | 46471 |
| 10 | 62295752 | 62296252 | Hk1 | CDS 6 | 46471 |
| 10 | 62295752 | 62296252 | Hk1 | Intron 5 | 46471 |
| 10 | 62295752 | 62296252 | Hk1 | 3' UTR | 22142 |
| 10 | 62295752 | 62296252 | Hk1 | CDS 7 | 22142 |
| 10 | 62295752 | 62296252 | Hk1 | Intron 6 | 22142 |
| 10 | 62295752 | 62296252 | Hk1 | CDS 6 | 22142 |
| 10 | 62295752 | 62296252 | Hk1 | Intron 5 | 22142 |
| 10 | 62295752 | 62296252 | Hk1 | 3' UTR | 31516 |
| 10 | 62295752 | 62296252 | Hk1 | CDS 6 | 31516 |
| 10 | 62295752 | 62296252 | Hk1 | Intron 5 | 31516 |
| 10 | 67377752 | 67378252 | Gm49454 | Non-coding | 80346 |
| 10 | 68008252 | 68008752 | Rtkn2 | Intron 6 | 28682 |
| 10 | 68008252 | 68008752 | Rtkn2 | Intron 6 | 28654 |
| 10 | 68008252 | 68008752 | Rtkn2 | Intron 6 | 28647 |
| 10 | 68008252 | 68008752 | Rtkn2 | Intron 6 | 28554 |
| 10 | 68008252 | 68008752 | Rtkn2 | Intron 6 | 28537 |
| 10 | 69929752 | 69930252 | Ank3 | Intron 25 | 530756 |
| 10 | 69890752 | 69891252 | Ank3 | Intron 17 | 491756 |
| 10 | 69929752 | 69930252 | Ank3 | Intron 25 | 395980 |
| 10 | 69890752 | 69891252 | Ank3 | Intron 16 | 356980 |
| 10 | 69929752 | 69930252 | Ank3 | Intron 23 | 395909 |
| 10 | 69890752 | 69891252 | Ank3 | Intron 16 | 356909 |
| 10 | 69929752 | 69930252 | Ank3 | Intron 24 | 395905 |
| 10 | 69890752 | 69891252 | Ank3 | Intron 16 | 356905 |
| 10 | 69929752 | 69930252 | Ank3 | Intron 24 | 395905 |
| 10 | 69890752 | 69891252 | Ank3 | Intron 16 | 356905 |
| 10 | 69929752 | 69930252 | Ank3 | Intron 1 | 395544 |
| 10 | 69890752 | 69891252 | Ank3 | Intron 1 | 356544 |
| 10 | 69929752 | 69930252 | Ank3 | Intron 1 | 395530 |

| | | | | | |
|---|---|---|---|---|---|
| 10 | 69890752 | 69891252 | Ank3 | Intron 1 | 356530 |
| 10 | 69929752 | 69930252 | Ank3 | Intron 23 | 395530 |
| 10 | 69890752 | 69891252 | Ank3 | Intron 16 | 356530 |
| 10 | 69929752 | 69930252 | Ank3 | Intron 24 | 395530 |
| 10 | 69890752 | 69891252 | Ank3 | Intron 16 | 356530 |
| 10 | 69929752 | 69930252 | Ank3 | Intron 24 | 395530 |
| 10 | 69890752 | 69891252 | Ank3 | Intron 16 | 356530 |
| 10 | 69929752 | 69930252 | Ank3 | Intron 25 | 395530 |
| 10 | 69890752 | 69891252 | Ank3 | Intron 16 | 356530 |
| 10 | 69929752 | 69930252 | Ank3 | Intron 25 | 223382 |
| 10 | 69890752 | 69891252 | Ank3 | Intron 17 | 184382 |
| 10 | 69929752 | 69930252 | Ank3 | Non-coding | 223278 |
| 10 | 69890752 | 69891252 | Ank3 | Non-coding | 184278 |
| 10 | 69929752 | 69930252 | Ank3 | Intron 5 | 31621 |
| 10 | 69929752 | 69930252 | Ank3 | Intron 3 | 30683 |
| 10 | 69929752 | 69930252 | Ank3 | Non-coding | 14887 |
| 10 | 69929752 | 69930252 | Ank3 | Intron 2 | 4257 |
| 10 | 69929752 | 69930252 | Ank3 | Intron 2 | 4220 |
| 10 | 69929752 | 69930252 | Ank3 | Intron 3 | 3979 |
| 10 | 69929752 | 69930252 | Ank3 | Intron 3 | 3979 |
| 10 | 69929752 | 69930252 | Ank3 | Intron 3 | 3979 |
| 10 | 69929752 | 69930252 | Ank3 | Intron 2 | 3947 |
| 10 | 69929752 | 69930252 | Ank3 | Non-coding | 3933 |
| 10 | 69929752 | 69930252 | Ank3 | Intron 2 | 3798 |
| 10 | 69929752 | 69930252 | Ank3 | Intron 2 | 3672 |
| 10 | 69929752 | 69930252 | Ank3 | Intron 3 | 3619 |
| 10 | 75353755 | 75354255 | Gucd1 | 3' UTR | 163019 |
| 10 | 75353755 | 75354255 | Gm5779 | Promoter | 1700 |
| 10 | 76625255 | 76625755 | Col6a2 | Promoter | -1624 |
| 10 | 76625255 | 76625755 | Col6a2 | Promoter | -1929 |
| 10 | 76625255 | 76625755 | Col6a2 | Promoter | -1929 |
| 10 | 76718255 | 76718755 | Col6a1 | Intron 13 | 7414 |
| 10 | 76718255 | 76718755 | Col6a1 | CDS 13 | 7414 |
| 10 | 76718255 | 76718755 | Col6a1 | Intron 12 | 7414 |
| 10 | 76718255 | 76718755 | Col6a1 | CDS 12 | 7414 |
| 10 | 76718255 | 76718755 | Col6a1 | Intron 11 | 7414 |
| 10 | 76718255 | 76718755 | Col6a1 | Promoter | 482 |
| 10 | 76906255 | 76906755 | Pcbp3 | 5' UTR | 55034 |
| 10 | 76906255 | 76906755 | Pcbp3 | 5' UTR | 55133 |
| 10 | 76906255 | 76906755 | Pcbp3 | Promoter | -392 |
| 10 | 77096255 | 77096755 | Col18a1 | CDS 3 | 69794 |
| 10 | 77096255 | 77096755 | Col18a1 | Intron 2 | 69794 |
| 10 | 77096255 | 77096755 | Col18a1 | CDS 2 | 16951 |
| 10 | 77096255 | 77096755 | Col18a1 | Intron 1 | 16951 |
| 10 | 77096255 | 77096755 | Col18a1 | CDS 2 | 17192 |
| 10 | 77096255 | 77096755 | Col18a1 | Intron 1 | 17192 |

| | | | | | |
|---|---|---|---|---|---|
| 10 | 78012255 | 78012755 | Pfkl | Promoter | -2447 |
| 10 | 78012255 | 78012755 | Pfkl | Promoter | -2527 |
| 10 | 78012255 | 78012755 | Pfkl | Promoter | -2171 |
| 10 | 78012255 | 78012755 | Pfkl | Promoter | -2518 |
| 10 | 80314255 | 80314755 | Apc2 | CDS 15 | 18278 |
| 10 | 80314255 | 80314755 | Apc2 | CDS 14 | 12435 |
| 10 | 80424255 | 80424755 | Tcf3 | Intron 3 | 8883 |
| 10 | 80424255 | 80424755 | Tcf3 | Intron 3 | 8893 |
| 10 | 80424255 | 80424755 | Tcf3 | Intron 3 | 8889 |
| 10 | 80424255 | 80424755 | Tcf3 | Intron 3 | 8893 |
| 10 | 80424255 | 80424755 | Tcf3 | Intron 3 | 8867 |
| 10 | 80424255 | 80424755 | Tcf3 | Intron 3 | 8867 |
| 10 | 80424255 | 80424755 | Tcf3 | Intron 2 | 6258 |
| 10 | 80424255 | 80424755 | Tcf3 | Intron 3 | 8881 |
| 10 | 80424255 | 80424755 | Tcf3 | Intron 3 | 8881 |
| 10 | 80424255 | 80424755 | Tcf3 | Intron 2 | 6328 |
| 10 | 80424255 | 80424755 | Tcf3 | Intron 2 | 6231 |
| 10 | 80424255 | 80424755 | Tcf3 | Promoter | -2394 |
| 10 | 80950255 | 80950755 | Gng7 | 3' UTR | 64171 |
| 10 | 80950255 | 80950755 | Gng7 | 3' UTR | 50611 |
| 10 | 80950255 | 80950755 | Gng7 | 3' UTR | 64191 |
| 10 | 80950255 | 80950755 | Gng7 | 3' UTR | 50624 |
| 10 | 81080755 | 81081255 | Thop1 | Intron 9 | 10720 |
| 10 | 81080755 | 81081255 | Thop1 | CDS 10 | 10720 |
| 10 | 81080755 | 81081255 | Thop1 | Intron 10 | 10720 |
| 10 | 81080755 | 81081255 | Thop1 | Promoter | 1887 |
| 10 | 81295755 | 81296255 | Tjp3 | Promoter | -4487 |
| 10 | 81295755 | 81296255 | Tjp3 | Promoter | -4173 |
| 10 | 81295755 | 81296255 | Tjp3 | Promoter | -4539 |
| 10 | 81295755 | 81296255 | Pip5k1c | Promoter | 2792 |
| 10 | 81295755 | 81296255 | Pip5k1c | Intron 1 | 2792 |
| 10 | 81295755 | 81296255 | Pip5k1c | Promoter | 2775 |
| 10 | 81295755 | 81296255 | Pip5k1c | Intron 1 | 2775 |
| 10 | 81295755 | 81296255 | Pip5k1c | Promoter | 2767 |
| 10 | 81295755 | 81296255 | Pip5k1c | Intron 1 | 2767 |
| 10 | 81295755 | 81296255 | Pip5k1c | Promoter | 2751 |
| 10 | 81295755 | 81296255 | Pip5k1c | Intron 1 | 2751 |
| 10 | 81295755 | 81296255 | Pip5k1c | Promoter | 2750 |
| 10 | 81295755 | 81296255 | Pip5k1c | Intron 1 | 2750 |
| 10 | 81364255 | 81364755 | Mfsd12 | 3' UTR | 6764 |
| 10 | 81365255 | 81365755 | Mfsd12 | 3' UTR | 7764 |
| 10 | 81364255 | 81364755 | 4930404N11Rik | Promoter | 100 |
| 10 | 81364255 | 81364755 | 4930404N11Rik | Intron 2 | 100 |

| | | | | | |
|---|---|---|---|---|---|
| 10 | 81364255 | 81364755 | 4930404N11Rik | CDS 2 | 100 |
| 10 | 81364255 | 81364755 | 4930404N11Rik | Intron 1 | 100 |
| 10 | 81364255 | 81364755 | 4930404N11Rik | CDS 1 | 100 |
| 10 | 81365255 | 81365755 | 4930404N11Rik | Promoter | -400 |
| 10 | 81364255 | 81364755 | 4930404N11Rik | Promoter | 122 |
| 10 | 81365255 | 81365755 | 4930404N11Rik | Promoter | -378 |
| 10 | 81364255 | 81364755 | 4930404N11Rik | Promoter | 1062 |
| 10 | 81364255 | 81364755 | 4930404N11Rik | Intron 3 | 1062 |
| 10 | 81364255 | 81364755 | 4930404N11Rik | CDS 3 | 1062 |
| 10 | 81364255 | 81364755 | 4930404N11Rik | Intron 2 | 1062 |
| 10 | 81364255 | 81364755 | 4930404N11Rik | CDS 2 | 1062 |
| 10 | 81365255 | 81365755 | 4930404N11Rik | Promoter | 62 |
| 10 | 81365255 | 81365755 | 4930404N11Rik | Intron 1 | 62 |
| 10 | 81365255 | 81365755 | 4930404N11Rik | CDS 1 | 62 |
| 10 | 81364255 | 81364755 | 4930404N11Rik | Promoter | 1036 |
| 10 | 81365255 | 81365755 | 4930404N11Rik | Promoter | 36 |
| 10 | 81494255 | 81494755 | Ncln | Promoter | 1638 |
| 10 | 81494255 | 81494755 | Ncln | Intron 1 | 1638 |
| 10 | 81494255 | 81494755 | Ncln | Promoter | 1572 |
| 10 | 81494255 | 81494755 | Ncln | Intron 1 | 1572 |
| 10 | 81494255 | 81494755 | Ncln | Promoter | -3986 |
| 10 | 81494255 | 81494755 | Ncln | Promoter | -2199 |
| 10 | 81494255 | 81494755 | Ncln | Promoter | 1588 |
| 10 | 89646755 | 89647255 | Scyl2 | Intron 14 | 39031 |
| 10 | 89646755 | 89647255 | Scyl2 | CDS 14 | 39031 |
| 10 | 89646755 | 89647255 | Scyl2 | Intron 13 | 39031 |
| 10 | 89646755 | 89647255 | Scyl2 | Non-coding | 7217 |
| 10 | 89646755 | 89647255 | Scyl2 | Intron 14 | 39014 |
| 10 | 89646755 | 89647255 | Scyl2 | CDS 14 | 39014 |
| 10 | 89646755 | 89647255 | Scyl2 | Intron 13 | 39014 |
| 10 | 91907755 | 91908255 | Gm31592 | Non-coding | 18926 |

| | | | | | |
|---|---|---|---|---|---|
| 10 | 92154755 | 92155255 | Rmst | Non-coding | 9916 |
| 10 | 92154755 | 92155255 | Rmst | Non-coding | 9494 |
| 10 | 92154755 | 92155255 | Rmst | Non-coding | 9412 |
| 10 | 92154755 | 92155255 | Rmst | Non-coding | 9913 |
| 10 | 92154755 | 92155255 | Rmst | Non-coding | 7886 |
| 10 | 92154755 | 92155255 | Rmst | Non-coding | 9717 |
| 10 | 94800755 | 94801255 | Plxnc1 | Intron 24 | 143581 |
| 10 | 95605366 | 95605866 | Gm23776 | Promoter | 1574 |
| 10 | 105237944 | 105238444 | Tmtc2 | Intron 11 | 335992 |
| 10 | 105237944 | 105238444 | Tmtc2 | Promoter | -2994 |
| 10 | 105237944 | 105238444 | Tmtc2 | Promoter | -3009 |
| 10 | 109683944 | 109684444 | Nav3 | 3' UTR | 315776 |
| 10 | 109683944 | 109684444 | Nav3 | CDS 39 | 315776 |
| 10 | 109683944 | 109684444 | Nav3 | Intron 38 | 315776 |
| 10 | 109683944 | 109684444 | Nav3 | 3' UTR | 771761 |
| 10 | 109683944 | 109684444 | Nav3 | CDS 41 | 771761 |
| 10 | 109683944 | 109684444 | Nav3 | Intron 40 | 771761 |
| 10 | 109683944 | 109684444 | Nav3 | Non-coding | 139142 |
| 10 | 109683944 | 109684444 | Nav3 | 3' UTR | 148778 |
| 10 | 109683944 | 109684444 | Nav3 | CDS 29 | 148778 |
| 10 | 109683944 | 109684444 | Nav3 | Intron 28 | 148778 |
| 10 | 115228944 | 115229444 | Tbc1d15 | Intron 5 | 22024 |
| 10 | 115228944 | 115229444 | Tbc1d15 | CDS 5 | 22024 |
| 10 | 115228944 | 115229444 | Tbc1d15 | Intron 4 | 22024 |
| 10 | 115228944 | 115229444 | Tbc1d15 | Non-coding | 21810 |
| 10 | 115348444 | 115348944 | Tmem19 | 5' UTR | 13288 |
| 10 | 115348444 | 115348944 | Tmem19 | Intron 3 | 13324 |
| 10 | 115348444 | 115348944 | Tmem19 | Intron 3 | 13558 |
| 10 | 115348444 | 115348944 | Tmem19 | Intron 2 | 13304 |
| 10 | 115348444 | 115348944 | Tmem19 | Intron 3 | 13281 |
| 10 | 115348444 | 115348944 | Tmem19 | Promoter | 121 |
| 10 | 115348444 | 115348944 | Tmem19 | 5' UTR | 13019 |
| 10 | 117641944 | 117642444 | Cpm | Intron 2 | 12444 |
| 10 | 117641944 | 117642444 | Cpm | Non-coding | 12444 |
| 10 | 119168944 | 119169444 | Gm34045 | Promoter | -2558 |
| 10 | 119637944 | 119638444 | Grip1 | Intron 1 | 184114 |
| 10 | 119637944 | 119638444 | Grip1 | Intron 1 | 184114 |
| 10 | 119637944 | 119638444 | Grip1 | Intron 1 | 184114 |
| 10 | 120173944 | 120174444 | Irak3 | Intron 5 | 27216 |
| 10 | 120173944 | 120174444 | Irak3 | Intron 5 | 27687 |
| 10 | 120173944 | 120174444 | Irak3 | Intron 5 | 27055 |
| 10 | 120173944 | 120174444 | Irak3 | Promoter | -3565 |
| 10 | 120173944 | 120174444 | Irak3 | Non-coding | 27144 |
| 10 | 120407444 | 120407944 | Hmga2 | Intron 3 | 68526 |
| 10 | 120407444 | 120407944 | Hmga2 | Intron 3 | 68526 |
| 10 | 120876944 | 120877444 | Msrb3 | 5' UTR | 21658 |

| | | | | | |
|---|---|---|---|---|---|
| 10 | 121918944 | 121919444 | Srgap1 | Intron 3 | 127865 |
| 10 | 122008944 | 122009444 | Srgap1 | Intron 1 | 37865 |
| 10 | 121918944 | 121919444 | Srgap1 | Intron 3 | 127872 |
| 10 | 122008944 | 122009444 | Srgap1 | Intron 1 | 37872 |
| 10 | 121918944 | 121919444 | Srgap1 | Non-coding | 127872 |
| 10 | 122008944 | 122009444 | Srgap1 | Non-coding | 37872 |
| 10 | 125372944 | 125373444 | Slc16a7 | 5' UTR | 16022 |
| 10 | 125372944 | 125373444 | Slc16a7 | Non-coding | 15925 |
| 11 | 5056997 | 5057497 | Gas2l1 | Non-coding | 7798 |
| 11 | 5056997 | 5057497 | Gas2l1 | Intron 6 | 7804 |
| 11 | 5056997 | 5057497 | Rasl10a | Promoter | -631 |
| 11 | 5056997 | 5057497 | Rasl10a | Promoter | -1469 |
| 11 | 5828997 | 5829497 | Polm | CDS 11 | 8293 |
| 11 | 5828997 | 5829497 | Polm | Intron 10 | 8293 |
| 11 | 5828997 | 5829497 | Polm | CDS 10 | 8293 |
| 11 | 5828997 | 5829497 | Polm | Intron 9 | 8293 |
| 11 | 5828997 | 5829497 | Polm | CDS 9 | 8293 |
| 11 | 5828997 | 5829497 | Polm | 3' UTR | 8520 |
| 11 | 5828997 | 5829497 | Polm | CDS 11 | 8520 |
| 11 | 5828997 | 5829497 | Polm | Intron 10 | 8520 |
| 11 | 5828997 | 5829497 | Polm | CDS 10 | 8520 |
| 11 | 5828997 | 5829497 | Polm | Intron 9 | 8520 |
| 11 | 5828997 | 5829497 | Polm | CDS 9 | 8520 |
| 11 | 5828997 | 5829497 | Polm | Promoter | 704 |
| 11 | 5972497 | 5972997 | Camk2b | Intron 19 | 92752 |
| 11 | 5972497 | 5972997 | Camk2b | CDS 19 | 92752 |
| 11 | 5972497 | 5972997 | Camk2b | Intron 18 | 92752 |
| 11 | 5972497 | 5972997 | Camk2b | CDS 18 | 92752 |
| 11 | 5972497 | 5972997 | Camk2b | Intron 17 | 92752 |
| 11 | 6015497 | 6015997 | Camk2b | CDS 3 | 49752 |
| 11 | 6015497 | 6015997 | Camk2b | Intron 2 | 49752 |
| 11 | 5984497 | 5984997 | Camk2b | Intron 13 | 80752 |
| 11 | 5971997 | 5972497 | Camk2b | 3' UTR | 93252 |
| 11 | 5971997 | 5972497 | Camk2b | Intron 20 | 93252 |
| 11 | 5971997 | 5972497 | Camk2b | CDS 20 | 93252 |
| 11 | 5971997 | 5972497 | Camk2b | Intron 19 | 93252 |
| 11 | 5972497 | 5972997 | Camk2b | Non-coding | 11142 |
| 11 | 5984497 | 5984997 | Camk2b | Promoter | -358 |
| 11 | 5971997 | 5972497 | Camk2b | Non-coding | 11642 |
| 11 | 5972497 | 5972997 | Camk2b | Intron 18 | 92752 |
| 11 | 5972497 | 5972997 | Camk2b | CDS 18 | 92752 |
| 11 | 5972497 | 5972997 | Camk2b | Intron 17 | 92752 |
| 11 | 5972497 | 5972997 | Camk2b | CDS 17 | 92752 |
| 11 | 5972497 | 5972997 | Camk2b | Intron 16 | 92752 |
| 11 | 6015497 | 6015997 | Camk2b | CDS 3 | 49752 |
| 11 | 6015497 | 6015997 | Camk2b | Intron 2 | 49752 |

| | | | | | |
|---|---|---|---|---|---|
| 11 | 5984497 | 5984997 | Camk2b | Intron 12 | 80752 |
| 11 | 5971997 | 5972497 | Camk2b | 3' UTR | 93252 |
| 11 | 5971997 | 5972497 | Camk2b | Intron 19 | 93252 |
| 11 | 5971997 | 5972497 | Camk2b | CDS 19 | 93252 |
| 11 | 5971997 | 5972497 | Camk2b | Intron 18 | 93252 |
| 11 | 5972497 | 5972997 | Camk2b | Intron 19 | 92567 |
| 11 | 5972497 | 5972997 | Camk2b | CDS 19 | 92567 |
| 11 | 5972497 | 5972997 | Camk2b | Intron 18 | 92567 |
| 11 | 5972497 | 5972997 | Camk2b | CDS 18 | 92567 |
| 11 | 5972497 | 5972997 | Camk2b | Intron 17 | 92567 |
| 11 | 6015497 | 6015997 | Camk2b | CDS 3 | 49567 |
| 11 | 6015497 | 6015997 | Camk2b | Intron 2 | 49567 |
| 11 | 5984497 | 5984997 | Camk2b | Intron 13 | 80567 |
| 11 | 5971997 | 5972497 | Camk2b | 3' UTR | 93067 |
| 11 | 5971997 | 5972497 | Camk2b | Intron 20 | 93067 |
| 11 | 5971997 | 5972497 | Camk2b | CDS 20 | 93067 |
| 11 | 5971997 | 5972497 | Camk2b | Intron 19 | 93067 |
| 11 | 5972497 | 5972997 | Camk2b | Intron 19 | 92567 |
| 11 | 5972497 | 5972997 | Camk2b | CDS 19 | 92567 |
| 11 | 5972497 | 5972997 | Camk2b | Intron 18 | 92567 |
| 11 | 5972497 | 5972997 | Camk2b | CDS 18 | 92567 |
| 11 | 5972497 | 5972997 | Camk2b | Intron 17 | 92567 |
| 11 | 6015497 | 6015997 | Camk2b | CDS 3 | 49567 |
| 11 | 6015497 | 6015997 | Camk2b | Intron 2 | 49567 |
| 11 | 5984497 | 5984997 | Camk2b | Intron 13 | 80567 |
| 11 | 5971997 | 5972497 | Camk2b | 3' UTR | 93067 |
| 11 | 5971997 | 5972497 | Camk2b | Intron 20 | 93067 |
| 11 | 5971997 | 5972497 | Camk2b | CDS 20 | 93067 |
| 11 | 5971997 | 5972497 | Camk2b | Intron 19 | 93067 |
| 11 | 5972497 | 5972997 | Camk2b | Intron 19 | 92586 |
| 11 | 5972497 | 5972997 | Camk2b | CDS 19 | 92586 |
| 11 | 5972497 | 5972997 | Camk2b | Intron 18 | 92586 |
| 11 | 5972497 | 5972997 | Camk2b | CDS 18 | 92586 |
| 11 | 5972497 | 5972997 | Camk2b | Intron 17 | 92586 |
| 11 | 6015497 | 6015997 | Camk2b | CDS 3 | 49586 |
| 11 | 6015497 | 6015997 | Camk2b | Intron 2 | 49586 |
| 11 | 5984497 | 5984997 | Camk2b | Intron 12 | 80586 |
| 11 | 5971997 | 5972497 | Camk2b | 3' UTR | 93086 |
| 11 | 5971997 | 5972497 | Camk2b | Intron 20 | 93086 |
| 11 | 5971997 | 5972497 | Camk2b | CDS 20 | 93086 |
| 11 | 5971997 | 5972497 | Camk2b | Intron 19 | 93086 |
| 11 | 5972497 | 5972997 | Camk2b | Intron 17 | 92746 |
| 11 | 5972497 | 5972997 | Camk2b | CDS 17 | 92746 |
| 11 | 5972497 | 5972997 | Camk2b | Intron 16 | 92746 |
| 11 | 5972497 | 5972997 | Camk2b | CDS 16 | 92746 |
| 11 | 5972497 | 5972997 | Camk2b | Intron 15 | 92746 |

| | | | | | |
|---|---|---|---|---|---|
| 11 | 6015497 | 6015997 | Camk2b | CDS 3 | 49746 |
| 11 | 6015497 | 6015997 | Camk2b | Intron 2 | 49746 |
| 11 | 5984497 | 5984997 | Camk2b | Intron 12 | 80746 |
| 11 | 5971997 | 5972497 | Camk2b | 3' UTR | 93246 |
| 11 | 5971997 | 5972497 | Camk2b | Intron 18 | 93246 |
| 11 | 5971997 | 5972497 | Camk2b | CDS 18 | 93246 |
| 11 | 5971997 | 5972497 | Camk2b | Intron 17 | 93246 |
| 11 | 5972497 | 5972997 | Camk2b | Intron 16 | 92549 |
| 11 | 5972497 | 5972997 | Camk2b | CDS 16 | 92549 |
| 11 | 5972497 | 5972997 | Camk2b | Intron 15 | 92549 |
| 11 | 5972497 | 5972997 | Camk2b | CDS 15 | 92549 |
| 11 | 5972497 | 5972997 | Camk2b | Intron 14 | 92549 |
| 11 | 6015497 | 6015997 | Camk2b | CDS 3 | 49549 |
| 11 | 6015497 | 6015997 | Camk2b | Intron 2 | 49549 |
| 11 | 5984497 | 5984997 | Camk2b | Intron 12 | 80549 |
| 11 | 5971997 | 5972497 | Camk2b | 3' UTR | 93049 |
| 11 | 5971997 | 5972497 | Camk2b | Intron 17 | 93049 |
| 11 | 5971997 | 5972497 | Camk2b | CDS 17 | 93049 |
| 11 | 5971997 | 5972497 | Camk2b | Intron 16 | 93049 |
| 11 | 5972497 | 5972997 | Camk2b | Intron 22 | 92616 |
| 11 | 5972497 | 5972997 | Camk2b | CDS 22 | 92616 |
| 11 | 5972497 | 5972997 | Camk2b | Intron 21 | 92616 |
| 11 | 5972497 | 5972997 | Camk2b | CDS 21 | 92616 |
| 11 | 5972497 | 5972997 | Camk2b | Intron 20 | 92616 |
| 11 | 6015497 | 6015997 | Camk2b | CDS 3 | 49616 |
| 11 | 6015497 | 6015997 | Camk2b | Intron 2 | 49616 |
| 11 | 5984497 | 5984997 | Camk2b | Intron 13 | 80616 |
| 11 | 5971997 | 5972497 | Camk2b | 3' UTR | 93116 |
| 11 | 5971997 | 5972497 | Camk2b | Intron 23 | 93116 |
| 11 | 5971997 | 5972497 | Camk2b | CDS 23 | 93116 |
| 11 | 5971997 | 5972497 | Camk2b | Intron 22 | 93116 |
| 11 | 5972497 | 5972997 | Camk2b | Intron 18 | 92542 |
| 11 | 5972497 | 5972997 | Camk2b | CDS 18 | 92542 |
| 11 | 5972497 | 5972997 | Camk2b | Intron 17 | 92542 |
| 11 | 5972497 | 5972997 | Camk2b | CDS 17 | 92542 |
| 11 | 5972497 | 5972997 | Camk2b | Intron 16 | 92542 |
| 11 | 6015497 | 6015997 | Camk2b | CDS 3 | 49542 |
| 11 | 6015497 | 6015997 | Camk2b | Intron 2 | 49542 |
| 11 | 5984497 | 5984997 | Camk2b | Intron 13 | 80542 |
| 11 | 5971997 | 5972497 | Camk2b | 3' UTR | 93042 |
| 11 | 5971997 | 5972497 | Camk2b | Intron 19 | 93042 |
| 11 | 5971997 | 5972497 | Camk2b | CDS 19 | 93042 |
| 11 | 5971997 | 5972497 | Camk2b | Intron 18 | 93042 |
| 11 | 5972497 | 5972997 | Camk2b | Intron 18 | 92574 |
| 11 | 5972497 | 5972997 | Camk2b | CDS 18 | 92574 |
| 11 | 5972497 | 5972997 | Camk2b | Intron 17 | 92574 |

| | | | | | |
|---|---|---|---|---|---|
| 11 | 5972497 | 5972997 | Camk2b | CDS 17 | 92574 |
| 11 | 5972497 | 5972997 | Camk2b | Intron 16 | 92574 |
| 11 | 6015497 | 6015997 | Camk2b | CDS 3 | 49574 |
| 11 | 6015497 | 6015997 | Camk2b | Intron 2 | 49574 |
| 11 | 5984497 | 5984997 | Camk2b | Intron 12 | 80574 |
| 11 | 5971997 | 5972497 | Camk2b | 3' UTR | 93074 |
| 11 | 5971997 | 5972497 | Camk2b | CDS 19 | 93074 |
| 11 | 5971997 | 5972497 | Camk2b | Intron 18 | 93074 |
| 11 | 6015497 | 6015997 | Camk2b | Non-coding | 49542 |
| 11 | 6015497 | 6015997 | Camk2b | Non-coding | 50366 |
| 11 | 6015497 | 6015997 | Camk2b | Non-coding | 49662 |
| 11 | 6601497 | 6601997 | Nacad | CDS 2 | 4057 |
| 11 | 6601497 | 6601997 | Nacad | Promoter | -2104 |
| 11 | 6601497 | 6601997 | Nacad | Promoter | -1558 |
| 11 | 7191997 | 7192497 | Ccdc201 | Promoter | -3427 |
| 11 | 23897000 | 23897500 | Papolg | Promoter | -1746 |
| 11 | 23897000 | 23897500 | Papolg | Promoter | -1763 |
| 11 | 23897000 | 23897500 | Gm12061 | Promoter | 2440 |
| 11 | 25856500 | 25857000 | 5730522E02Rik | Non-coding | 353570 |
| 11 | 25856500 | 25857000 | 5730522E02Rik | 5' UTR | 353573 |
| 11 | 25856500 | 25857000 | 5730522E02Rik | Non-coding | 224601 |
| 11 | 32190500 | 32191000 | Il9r | 3' UTR | 9223 |
| 11 | 32190500 | 32191000 | Il9r | CDS 9 | 9223 |
| 11 | 32190500 | 32191000 | Il9r | 3' UTR | 9280 |
| 11 | 32190500 | 32191000 | Il9r | 3' UTR | 9237 |
| 11 | 32190500 | 32191000 | Il9r | CDS 9 | 9237 |
| 11 | 32190500 | 32191000 | Il9r | 3' UTR | 9223 |
| 11 | 32190500 | 32191000 | Il9r | CDS 9 | 9223 |
| 11 | 33947000 | 33947500 | Kcnip1 | Intron 1 | 45489 |
| 11 | 33947000 | 33947500 | Kcnip1 | Non-coding | 45653 |
| 11 | 33947000 | 33947500 | Kcnip1 | Non-coding | 45489 |
| 11 | 36943998 | 36944498 | Tenm2 | 5' UTR | 291385 |
| 11 | 36943998 | 36944498 | Tenm2 | CDS 3 | 291385 |
| 11 | 36943998 | 36944498 | Tenm2 | Intron 2 | 291385 |
| 11 | 36943998 | 36944498 | Tenm2 | Promoter | 0 |
| 11 | 36943998 | 36944498 | Tenm2 | CDS 1 | 0 |
| 11 | 36943998 | 36944498 | Tenm2 | Promoter | 0 |
| 11 | 36943998 | 36944498 | Tenm2 | CDS 1 | 0 |
| 11 | 36943998 | 36944498 | Tenm2 | Promoter | 0 |
| 11 | 36943998 | 36944498 | Tenm2 | Non-coding | 291467 |
| 11 | 41867498 | 41867998 | A230004M16Rik | Non-coding | 157156 |
| 11 | 41976498 | 41976998 | Gabrg2 | Intron 2 | 23860 |
| 11 | 41976498 | 41976998 | Gabrg2 | CDS 2 | 23860 |
| 11 | 41976498 | 41976998 | Gabrg2 | Intron 1 | 23860 |
| 11 | 41976498 | 41976998 | Gabrg2 | Intron 2 | 23535 |

| | | | | | |
|---|---|---|---|---|---|
| 11 | 41976498 | 41976998 | Gabrg2 | CDS 2 | 23535 |
| 11 | 41976498 | 41976998 | Gabrg2 | Intron 1 | 23535 |
| 11 | 41976498 | 41976998 | Gabrg2 | Intron 2 | 23359 |
| 11 | 41976498 | 41976998 | Gabrg2 | CDS 2 | 23359 |
| 11 | 41976498 | 41976998 | Gabrg2 | Intron 1 | 23359 |
| 11 | 44916998 | 44917498 | Ebf1 | Intron 9 | 299681 |
| 11 | 44916998 | 44917498 | Ebf1 | Intron 9 | 299681 |
| 11 | 44916998 | 44917498 | Ebf1 | Intron 9 | 299681 |
| 11 | 44916998 | 44917498 | Ebf1 | Non-coding | 4860 |
| 11 | 52187498 | 52187998 | Gm12206 | Promoter | -955 |
| 11 | 53851998 | 53852498 | Gm12216 | Non-coding | 6647 |
| 11 | 53851998 | 53852498 | Gm12216 | 5' UTR | 6759 |
| 11 | 53851998 | 53852498 | Gm12216 | Promoter | -1949 |
| 11 | 53851998 | 53852498 | Gm12216 | Non-coding | 6732 |
| 11 | 53851998 | 53852498 | Gm12216 | 5' UTR | 6759 |
| 11 | 53851998 | 53852498 | Gm12216 | 5' UTR | 6759 |
| 11 | 57685498 | 57685998 | Galnt10 | Intron 1 | 40056 |
| 11 | 57708498 | 57708998 | Galnt10 | Intron 1 | 63056 |
| 11 | 57685498 | 57685998 | Galnt10 | Intron 1 | 40033 |
| 11 | 57708498 | 57708998 | Galnt10 | Intron 1 | 63033 |
| 11 | 58877998 | 58878498 | 2810021J22Rik | Intron 3 | 10782 |
| 11 | 58877998 | 58878498 | 2810021J22Rik | CDS 4 | 10731 |
| 11 | 58995498 | 58995998 | Obscn | CDS 101 | 140378 |
| 11 | 58995498 | 58995998 | Obscn | Intron 100 | 140378 |
| 11 | 58995498 | 58995998 | Obscn | CDS 100 | 140378 |
| 11 | 59067498 | 59067998 | Obscn | Intron 38 | 68378 |
| 11 | 59067498 | 59067998 | Obscn | CDS 38 | 68378 |
| 11 | 59067498 | 59067998 | Obscn | Intron 37 | 68378 |
| 11 | 59067498 | 59067998 | Obscn | CDS 37 | 68378 |
| 11 | 58995498 | 58995998 | Obscn | CDS 9 | 4362 |
| 11 | 58995498 | 58995998 | Obscn | Intron 8 | 4362 |
| 11 | 58995498 | 58995998 | Obscn | CDS 8 | 4362 |
| 11 | 58995498 | 58995998 | Obscn | CDS 112 | 140378 |
| 11 | 58995498 | 58995998 | Obscn | Intron 111 | 140378 |
| 11 | 58995498 | 58995998 | Obscn | CDS 111 | 140378 |
| 11 | 59067498 | 59067998 | Obscn | Intron 41 | 68378 |
| 11 | 59067498 | 59067998 | Obscn | CDS 41 | 68378 |
| 11 | 59067498 | 59067998 | Obscn | Intron 40 | 68378 |
| 11 | 59067498 | 59067998 | Obscn | CDS 40 | 68378 |
| 11 | 59067498 | 59067998 | Obscn | Intron 40 | 68399 |
| 11 | 59067498 | 59067998 | Obscn | CDS 40 | 68399 |
| 11 | 59067498 | 59067998 | Obscn | Intron 39 | 68399 |
| 11 | 59067498 | 59067998 | Obscn | CDS 39 | 68399 |
| 11 | 59067498 | 59067998 | Obscn | Intron 41 | 68378 |
| 11 | 59067498 | 59067998 | Obscn | CDS 41 | 68378 |
| 11 | 59067498 | 59067998 | Obscn | Intron 40 | 68378 |

| | | | | | |
|---|---|---|---|---|---|
| 11 | 59067498 | 59067998 | Obscn | CDS 40 | 68378 |
| 11 | 59067498 | 59067998 | Obscn | Intron 37 | 71173 |
| 11 | 59067498 | 59067998 | Obscn | CDS 37 | 71173 |
| 11 | 59067498 | 59067998 | Obscn | Intron 36 | 71173 |
| 11 | 59067498 | 59067998 | Obscn | CDS 36 | 71173 |
| 11 | 62921498 | 62921998 | Tvp23bos | Promoter | 0 |
| 11 | 66278498 | 66278998 | Gm12296 | Non-coding | 110463 |
| 11 | 66278498 | 66278998 | Shisa6 | Intron 3 | 246798 |
| 11 | 66278498 | 66278998 | Shisa6 | Intron 2 | 246967 |
| 11 | 68476498 | 68476998 | Pik3r5 | Intron 4 | 44377 |
| 11 | 68476498 | 68476998 | Pik3r5 | Non-coding | 44373 |
| 11 | 68476498 | 68476998 | Pik3r5 | Promoter | 2703 |
| 11 | 68476498 | 68476998 | Pik3r5 | Promoter | 1017 |
| 11 | 74590998 | 74591498 | Rap1gap2 | Promoter | -811 |
| 11 | 74413998 | 74414498 | Rap1gap2 | Intron 13 | 175689 |
| 11 | 74413998 | 74414498 | Rap1gap2 | CDS 13 | 175689 |
| 11 | 74590998 | 74591498 | Rap1gap2 | Intron 2 | 19418 |
| 11 | 74413998 | 74414498 | Rap1gap2 | Intron 14 | 196418 |
| 11 | 74413998 | 74414498 | Rap1gap2 | CDS 14 | 196418 |
| 11 | 74590998 | 74591498 | Rap1gap2 | Intron 2 | 19418 |
| 11 | 74413998 | 74414498 | Rap1gap2 | Intron 14 | 196418 |
| 11 | 74413998 | 74414498 | Rap1gap2 | CDS 14 | 196418 |
| 11 | 74590998 | 74591498 | Rap1gap2 | Promoter | -873 |
| 11 | 76387998 | 76388498 | Nxn | Intron 1 | 10643 |
| 11 | 76387998 | 76388498 | Nxn | Non-coding | 10571 |
| 11 | 77021498 | 77021998 | Slc6a4 | Intron 12 | 22895 |
| 11 | 77021498 | 77021998 | Slc6a4 | Intron 11 | 22882 |
| 11 | 77021498 | 77021998 | Slc6a4 | Non-coding | 3321 |
| 11 | 77769998 | 77770498 | Myo18a | 5' UTR | 6752 |
| 11 | 77769998 | 77770498 | Myo18a | Non-coding | 6697 |
| 11 | 77769998 | 77770498 | Myo18a | 5' UTR | 4395 |
| 11 | 77769998 | 77770498 | Myo18a | 5' UTR | 3262 |
| 11 | 77769998 | 77770498 | Myo18a | Promoter | -1653 |
| 11 | 77916998 | 77917498 | Gm11190 | Non-coding | 11791 |
| 11 | 78480498 | 78480998 | Sarm1 | Intron 7 | 16461 |
| 11 | 78480498 | 78480998 | Sarm1 | Promoter | 2801 |
| 11 | 78480498 | 78480998 | Sarm1 | Intron 7 | 16757 |
| 11 | 78750998 | 78751498 | Ccnq | Promoter | 232 |
| 11 | 78750998 | 78751498 | Ccnq | CDS 1 | 232 |
| 11 | 80949498 | 80949998 | Asic2 | Intron 3 | 203324 |
| 11 | 80949498 | 80949998 | Asic2 | Intron 3 | 1018460 |
| 11 | 81426498 | 81426998 | Asic2 | Intron 1 | 541460 |
| 11 | 82218998 | 82219498 | Ccl1 | Non-coding | 86193 |
| 11 | 85733998 | 85734498 | Bcas3 | Intron 24 | 380831 |
| 11 | 85719498 | 85719998 | Bcas3 | Intron 24 | 366331 |
| 11 | 85733998 | 85734498 | Bcas3 | Intron 23 | 380831 |

| | | | | | |
|---|---|---|---|---|---|
| 11 | 85719498 | 85719998 | Bcas3 | Intron 23 | 366331 |
| 11 | 85733998 | 85734498 | Bcas3 | Intron 23 | 380786 |
| 11 | 85719498 | 85719998 | Bcas3 | Intron 23 | 366286 |
| 11 | 85733998 | 85734498 | Bcas3 | Intron 22 | 380786 |
| 11 | 85719498 | 85719998 | Bcas3 | Intron 22 | 366286 |
| 11 | 85733998 | 85734498 | Bcas3 | Intron 13 | 267839 |
| 11 | 85719498 | 85719998 | Bcas3 | Intron 13 | 253339 |
| 11 | 85733998 | 85734498 | Bcas3 | Intron 9 | 224371 |
| 11 | 85719498 | 85719998 | Bcas3 | Intron 9 | 209871 |
| 11 | 85733998 | 85734498 | Bcas3 | Intron 8 | 224360 |
| 11 | 85719498 | 85719998 | Bcas3 | Intron 8 | 209860 |
| 11 | 85733998 | 85734498 | Bcas3 | Non-coding | 93294 |
| 11 | 85719498 | 85719998 | Bcas3 | Non-coding | 78794 |
| 11 | 85719498 | 85719998 | Bcas3os1 | Promoter | 0 |
| 11 | 85719498 | 85719998 | Bcas3os1 | Promoter | 0 |
| 11 | 88665686 | 88666186 | Msi2 | Intron 5 | 52044 |
| 11 | 88665686 | 88666186 | Msi2 | Intron 5 | 51916 |
| 11 | 88665686 | 88666186 | Msi2 | Non-coding | 51925 |
| 11 | 88665686 | 88666186 | Msi2 | Intron 5 | 52328 |
| 11 | 95257686 | 95258186 | Tac4 | Promoter | -3343 |
| 11 | 96308686 | 96309186 | Hoxb5os | Promoter | -1775 |
| 11 | 96308686 | 96309186 | Hoxb5os | Promoter | -3604 |
| 11 | 96613186 | 96613686 | Skap1 | Intron 4 | 148593 |
| 11 | 96613186 | 96613686 | Skap1 | Intron 4 | 148593 |
| 11 | 96613186 | 96613686 | Skap1 | Non-coding | 148562 |
| 11 | 96613186 | 96613686 | Skap1 | Intron 4 | 148559 |
| 11 | 96613186 | 96613686 | Skap1 | Intron 4 | 148505 |
| 11 | 96613186 | 96613686 | Skap1 | Non-coding | 123325 |
| 11 | 98968686 | 98969186 | Rara | Intron 4 | 40868 |
| 11 | 98968686 | 98969186 | Rara | Intron 4 | 30988 |
| 11 | 98968686 | 98969186 | Rara | Intron 5 | 30968 |
| 11 | 98968686 | 98969186 | Rara | Intron 4 | 28975 |
| 11 | 98968686 | 98969186 | Rara | Intron 3 | 8274 |
| 11 | 100745686 | 100746186 | Kcnh4 | CDS 13 | 13555 |
| 11 | 100745686 | 100746186 | Kcnh4 | Intron 12 | 13555 |
| 11 | 100745686 | 100746186 | Kcnh4 | CDS 12 | 13555 |
| 11 | 100745686 | 100746186 | Kcnh4 | CDS 13 | 13757 |
| 11 | 100745686 | 100746186 | Kcnh4 | Intron 12 | 13757 |
| 11 | 100745686 | 100746186 | Kcnh4 | CDS 12 | 13757 |
| 11 | 101178186 | 101178686 | Cntnap1 | Non-coding | 7663 |
| 11 | 101178686 | 101179186 | Ccr10 | Promoter | -3242 |
| 11 | 101179186 | 101179686 | Ccr10 | Promoter | -3742 |
| 11 | 101178186 | 101178686 | Ccr10 | Promoter | -2742 |
| 11 | 101178686 | 101179186 | Cntnap1 | Promoter | 2645 |
| 11 | 101178686 | 101179186 | Cntnap1 | Intron 5 | 2645 |
| 11 | 101178686 | 101179186 | Cntnap1 | CDS 6 | 2645 |

| | | | | | |
|---|---|---|---|---|---|
| 11 | 101178686 | 101179186 | Cntnap1 | Intron 6 | 2645 |
| 11 | 101179186 | 101179686 | Cntnap1 | Intron 6 | 3145 |
| 11 | 101179186 | 101179686 | Cntnap1 | CDS 7 | 3145 |
| 11 | 101179186 | 101179686 | Cntnap1 | Intron 7 | 3145 |
| 11 | 101178186 | 101178686 | Cntnap1 | Promoter | 2145 |
| 11 | 101178186 | 101178686 | Cntnap1 | Intron 4 | 2145 |
| 11 | 101178186 | 101178686 | Cntnap1 | CDS 5 | 2145 |
| 11 | 101178186 | 101178686 | Cntnap1 | Intron 5 | 2145 |
| 11 | 102779686 | 102780186 | Adam11 | 3' UTR | 18247 |
| 11 | 102779686 | 102780186 | Adam11 | 3' UTR | 18247 |
| 11 | 103983686 | 103984186 | Arf2 | 3' UTR | 16947 |
| 11 | 103983686 | 103984186 | Arf2 | CDS 5 | 16947 |
| 11 | 103983686 | 103984186 | Arf2 | 3' UTR | 16815 |
| 11 | 103983686 | 103984186 | Arf2 | CDS 5 | 16815 |
| 11 | 104688686 | 104689186 | Gm11639 | Promoter | 2979 |
| 11 | 104688686 | 104689186 | Gm11639 | Intron 2 | 2979 |
| 11 | 104688686 | 104689186 | Gm11639 | Promoter | 2979 |
| 11 | 104688686 | 104689186 | Gm11639 | Intron 2 | 2979 |
| 11 | 106345686 | 106346186 | Scn4a | CDS 6 | 3204 |
| 11 | 106345686 | 106346186 | Scn4a | Intron 5 | 3204 |
| 11 | 107393686 | 107394186 | Pitpnc1 | Intron 1 | 76514 |
| 11 | 109297186 | 109297686 | Rgs9 | Promoter | 444 |
| 11 | 109297186 | 109297686 | Rgs9 | Intron 1 | 444 |
| 11 | 109297186 | 109297686 | Rgs9 | Promoter | 381 |
| 11 | 109297186 | 109297686 | Rgs9 | Intron 1 | 381 |
| 11 | 109297186 | 109297686 | Rgs9 | Promoter | 421 |
| 11 | 109297186 | 109297686 | Rgs9 | Intron 1 | 421 |
| 11 | 109559186 | 109559686 | Arsg | Intron 12 | 85812 |
| 11 | 109559186 | 109559686 | Arsg | Intron 10 | 73580 |
| 11 | 109559186 | 109559686 | Arsg | Intron 3 | 15432 |
| 11 | 110450186 | 110450686 | Map2k6 | Intron 1 | 51064 |
| 11 | 110450186 | 110450686 | Map2k6 | Intron 1 | 51064 |
| 11 | 110450186 | 110450686 | Map2k6 | Non-coding | 50927 |
| 11 | 113172186 | 113172686 | 2610035D17Rik | Non-coding | 29153 |
| 11 | 113172186 | 113172686 | 2610035D17Rik | Promoter | 392 |
| 11 | 113172186 | 113172686 | 2610035D17Rik | Non-coding | 29137 |
| 11 | 113172186 | 113172686 | 2610035D17Rik | Promoter | 392 |
| 11 | 113805686 | 113806186 | Sdk2 | Intron 38 | 260033 |
| 11 | 114067686 | 114068186 | Sdk2 | Promoter | -1467 |
| 11 | 113806186 | 113806686 | Sdk2 | Intron 38 | 259533 |
| 11 | 113806186 | 113806686 | Sdk2 | CDS 38 | 259533 |
| 11 | 113806186 | 113806686 | Sdk2 | Intron 37 | 259533 |

| | | | | | |
|---|---|---|---|---|---|
| 11 | 113805686 | 113806186 | Sdk2 | Non-coding | 4238 |
| 11 | 113806186 | 113806686 | Sdk2 | Non-coding | 3738 |
| 11 | 113806186 | 113806686 | Sdk2 | Non-coding | 19534 |
| 11 | 114067686 | 114068186 | Sdk2 | Promoter | -1710 |
| 11 | 114067686 | 114068186 | Sdk2 | Promoter | -639 |
| 11 | 114067686 | 114068186 | Sdk2 | Promoter | -656 |
| 11 | 115237686 | 115238186 | Tmem104 | Intron 9 | 50199 |
| 11 | 115237686 | 115238186 | Tmem104 | Intron 9 | 50199 |
| 11 | 115237686 | 115238186 | Tmem104 | Non-coding | 50135 |
| 11 | 115317686 | 115318186 | Otop2 | 5' UTR | 10523 |
| 11 | 115317686 | 115318186 | Ush1g | Intron 2 | 3856 |
| 11 | 115317686 | 115318186 | Ush1g | CDS 2 | 3856 |
| 11 | 115317686 | 115318186 | Otop2 | Promoter | -4344 |
| 11 | 115317686 | 115318186 | Otop2 | Promoter | -4575 |
| 11 | 115317686 | 115318186 | Otop2 | Promoter | -4739 |
| 11 | 115382686 | 115383186 | Cdr2l | Promoter | 770 |
| 11 | 115382686 | 115383186 | Cdr2l | Intron 1 | 770 |
| 11 | 115446686 | 115447186 | Trim80 | Intron 6 | 6141 |
| 11 | 115446686 | 115447186 | Trim80 | CDS 7 | 6141 |
| 11 | 115446686 | 115447186 | Trim80 | Intron 7 | 6141 |
| 11 | 115446686 | 115447186 | Mir3968 | Promoter | 875 |
| 11 | 115699686 | 115700186 | Grb2 | 5' UTR | 8412 |
| 11 | 115699686 | 115700186 | Grb2 | Promoter | -151 |
| 11 | 115699686 | 115700186 | Grb2 | Promoter | -378 |
| 11 | 115699686 | 115700186 | Grb2 | 5' UTR | 8391 |
| 11 | 115699686 | 115700186 | Grb2 | Promoter | 0 |
| 11 | 115699686 | 115700186 | Grb2 | 5' UTR | 0 |
| 11 | 116647186 | 116647686 | Cygb | Intron 3 | 6628 |
| 11 | 117038686 | 117039186 | Gm11728 | Promoter | 1008 |
| 11 | 117038686 | 117039186 | Gm11728 | Promoter | 1003 |
| 11 | 118100686 | 118101186 | Dnah17 | Intron 23 | 28034 |
| 11 | 118100686 | 118101186 | Dnah17 | CDS 23 | 28034 |
| 11 | 118100686 | 118101186 | Dnah17 | Intron 22 | 28034 |
| 11 | 118059686 | 118060186 | Dnah17 | Intron 55 | 69034 |
| 11 | 118059686 | 118060186 | Dnah17 | CDS 55 | 69034 |
| 11 | 118100686 | 118101186 | Dnah17 | Intron 23 | 28034 |
| 11 | 118100686 | 118101186 | Dnah17 | CDS 23 | 28034 |
| 11 | 118100686 | 118101186 | Dnah17 | Intron 22 | 28034 |
| 11 | 118059686 | 118060186 | Dnah17 | Intron 54 | 69034 |
| 11 | 118059686 | 118060186 | Dnah17 | CDS 54 | 69034 |
| 11 | 118100686 | 118101186 | Dnah17 | Intron 24 | 29449 |
| 11 | 118100686 | 118101186 | Dnah17 | CDS 24 | 29449 |
| 11 | 118100686 | 118101186 | Dnah17 | Intron 23 | 29449 |
| 11 | 118059686 | 118060186 | Dnah17 | Intron 55 | 70449 |
| 11 | 118059686 | 118060186 | Dnah17 | CDS 55 | 70449 |
| 11 | 118668686 | 118669186 | Rbfox3 | 5' UTR | 240365 |

| | | | | | |
|---|---|---|---|---|---|
| 11 | 118877186 | 118877686 | Rbfox3 | 5' UTR | 31865 |
| 11 | 118668686 | 118669186 | Rbfox3 | 5' UTR | 91856 |
| 11 | 118668686 | 118669186 | Rbfox3 | 5' UTR | 240387 |
| 11 | 118877186 | 118877686 | Rbfox3 | 5' UTR | 31887 |
| 11 | 118668686 | 118669186 | Rbfox3 | 5' UTR | 242412 |
| 11 | 118877186 | 118877686 | Rbfox3 | 5' UTR | 33912 |
| 11 | 118668686 | 118669186 | Rbfox3 | 5' UTR | 240365 |
| 11 | 118877186 | 118877686 | Rbfox3 | 5' UTR | 31865 |
| 11 | 118668686 | 118669186 | Rbfox3 | 5' UTR | 240367 |
| 11 | 118877186 | 118877686 | Rbfox3 | 5' UTR | 31867 |
| 11 | 118877186 | 118877686 | Rbfox3 | Non-coding | 11004 |
| 11 | 118877186 | 118877686 | Rbfox3 | Non-coding | 30320 |
| 11 | 120498186 | 120498686 | Slc25a10 | 3' UTR | 6346 |
| 11 | 120498186 | 120498686 | Slc25a10 | Non-coding | 6298 |
| 11 | 120498186 | 120498686 | Slc25a10 | Promoter | 2087 |
| 11 | 120498186 | 120498686 | Slc25a10 | Promoter | 461 |
| 12 | 10212194 | 10212694 | Gm48791 | Non-coding | 35051 |
| 12 | 13126194 | 13126694 | Gm48213 | Promoter | 418 |
| 12 | 16765694 | 16766194 | Greb1 | 5' UTR | 34491 |
| 12 | 16765694 | 16766194 | Greb1 | 5' UTR | 34693 |
| 12 | 16765694 | 16766194 | Gm44305 | Promoter | 1030 |
| 12 | 16853194 | 16853694 | Gm36495 | Promoter | -2312 |
| 12 | 17330694 | 17331194 | Atp6v1c2 | Promoter | -1345 |
| 12 | 17330694 | 17331194 | Atp6v1c2 | Promoter | -1334 |
| 12 | 18243694 | 18244194 | Gm19368 | Promoter | -4619 |
| 12 | 18229694 | 18230194 | Gm19368 | Non-coding | 8881 |
| 12 | 18237694 | 18238194 | Gm19368 | Promoter | 881 |
| 12 | 18328694 | 18329194 | Gm7093 | Promoter | -3679 |
| 12 | 20214645 | 20215145 | Gm49692 | Non-coding | 22745 |
| 12 | 20284645 | 20285145 | Gm49692 | Non-coding | 92745 |
| 12 | 20293645 | 20294145 | Gm49692 | Non-coding | 101745 |
| 12 | 20294145 | 20294645 | Gm49692 | Non-coding | 102245 |
| 12 | 20291645 | 20292145 | Gm49692 | Non-coding | 99745 |
| 12 | 20285145 | 20285645 | Gm49692 | Non-coding | 93245 |
| 12 | 20267145 | 20267645 | Gm49692 | Non-coding | 75245 |
| 12 | 20278145 | 20278645 | Gm49692 | Non-coding | 86245 |
| 12 | 20281645 | 20282145 | Gm49692 | Non-coding | 89745 |
| 12 | 20274645 | 20275145 | Gm49692 | Non-coding | 82745 |
| 12 | 20269145 | 20269645 | Gm49692 | Non-coding | 77245 |
| 12 | 20251645 | 20252145 | Gm49692 | Non-coding | 59745 |
| 12 | 20264645 | 20265145 | Gm49692 | Non-coding | 72745 |
| 12 | 20239145 | 20239645 | Gm49692 | Non-coding | 47245 |
| 12 | 20219645 | 20220145 | Gm49692 | Non-coding | 27745 |
| 12 | 20239145 | 20239645 | Gm48668 | Promoter | -3026 |
| 12 | 20274645 | 20275145 | Gm9269 | Promoter | 0 |
| 12 | 20814639 | 20815139 | 1700030C10Rik | Promoter | 641 |

| | | | | | |
|---|---|---|---|---|---|
| 12 | 20814639 | 20815139 | 1700030C10Rik | Promoter | 589 |
| 12 | 20814639 | 20815139 | 1700030C10Rik | Promoter | 333 |
| 12 | 20920139 | 20920639 | 3110053B16Rik | Promoter | 624 |
| 12 | 20920139 | 20920639 | 3110053B16Rik | Promoter | 415 |
| 12 | 20920139 | 20920639 | 3110053B16Rik | Promoter | 590 |
| 12 | 20920139 | 20920639 | Zfp125 | Promoter | 624 |
| 12 | 20920139 | 20920639 | Zfp125 | Promoter | 624 |
| 12 | 23267439 | 23267939 | Gm20472 | Non-coding | 47420 |
| 12 | 24493135 | 24493635 | Gm16372 | Promoter | 22 |
| 12 | 24493135 | 24493635 | Taf1b | Promoter | -4724 |
| 12 | 24493135 | 24493635 | Taf1b | Promoter | -4976 |
| 12 | 24493135 | 24493635 | Taf1b | Promoter | -4994 |
| 12 | 24493135 | 24493635 | Taf1b | Promoter | -4995 |
| 12 | 24654635 | 24655135 | Klf11 | Intron 2 | 3361 |
| 12 | 24654635 | 24655135 | Klf11 | CDS 3 | 3361 |
| 12 | 24654635 | 24655135 | Klf11 | Promoter | 2739 |
| 12 | 24654635 | 24655135 | Klf11 | Promoter | 2739 |
| 12 | 24654635 | 24655135 | Klf11 | Intron 2 | 2739 |
| 12 | 24654635 | 24655135 | Klf11 | CDS 3 | 2739 |
| 12 | 24654635 | 24655135 | Klf11 | Promoter | 2481 |
| 12 | 24654635 | 24655135 | Klf11 | Intron 2 | 2481 |
| 12 | 24654635 | 24655135 | Klf11 | CDS 3 | 2481 |
| 12 | 26180635 | 26181135 | Gm29687 | Non-coding | 29371 |
| 12 | 26387135 | 26387635 | Rnf144a | 5' UTR | 27620 |
| 12 | 26405135 | 26405635 | Rnf144a | 5' UTR | 9620 |
| 12 | 26387135 | 26387635 | Rnf144a | 5' UTR | 27582 |
| 12 | 26405135 | 26405635 | Rnf144a | 5' UTR | 9582 |
| 12 | 26387135 | 26387635 | Rnf144a | 5' UTR | 27497 |
| 12 | 26405135 | 26405635 | Rnf144a | 5' UTR | 9497 |
| 12 | 26405135 | 26405635 | Rnf144a | Non-coding | 8370 |
| 12 | 26405135 | 26405635 | Rnf144a | Promoter | 813 |
| 12 | 27136135 | 27136635 | Gm9866 | Non-coding | 23869 |
| 12 | 27136135 | 27136635 | Gm9866 | Non-coding | 23999 |
| 12 | 27136135 | 27136635 | Gm9866 | Non-coding | 23999 |
| 12 | 28800635 | 28801135 | Eipr1 | Intron 3 | 48832 |
| 12 | 28800635 | 28801135 | Eipr1 | Intron 4 | 48780 |
| 12 | 28800635 | 28801135 | Eipr1 | Intron 3 | 48774 |
| 12 | 28905135 | 28905635 | Gm31508 | Promoter | -4594 |
| 12 | 28949135 | 28949635 | Gm31508 | Non-coding | 38906 |
| 12 | 29894635 | 29895135 | Myt1l | Intron 20 | 366251 |
| 12 | 29894635 | 29895135 | Myt1l | CDS 21 | 366251 |
| 12 | 29616635 | 29617135 | Myt1l | 5' UTR | 88251 |
| 12 | 29805635 | 29806135 | Myt1l | Intron 8 | 277251 |
| 12 | 29829635 | 29830135 | Myt1l | Intron 10 | 301251 |
| 12 | 29627635 | 29628135 | Myt1l | 5' UTR | 99251 |
| 12 | 29618635 | 29619135 | Myt1l | 5' UTR | 90251 |

| | | | | | |
|---|---|---|---|---|---|
| 12 | 29616135 | 29616635 | Myt1l | 5' UTR | 87751 |
| 12 | 29894635 | 29895135 | Myt1l | Intron 20 | 366251 |
| 12 | 29894635 | 29895135 | Myt1l | CDS 21 | 366251 |
| 12 | 29616635 | 29617135 | Myt1l | 5' UTR | 88251 |
| 12 | 29805635 | 29806135 | Myt1l | Intron 8 | 277251 |
| 12 | 29829635 | 29830135 | Myt1l | Intron 10 | 301251 |
| 12 | 29627635 | 29628135 | Myt1l | 5' UTR | 99251 |
| 12 | 29618635 | 29619135 | Myt1l | 5' UTR | 90251 |
| 12 | 29616135 | 29616635 | Myt1l | 5' UTR | 87751 |
| 12 | 29894635 | 29895135 | Myt1l | Intron 20 | 360020 |
| 12 | 29894635 | 29895135 | Myt1l | CDS 21 | 360020 |
| 12 | 29616635 | 29617135 | Myt1l | 5' UTR | 82020 |
| 12 | 29805635 | 29806135 | Myt1l | Intron 8 | 271020 |
| 12 | 29829635 | 29830135 | Myt1l | Intron 10 | 295020 |
| 12 | 29627635 | 29628135 | Myt1l | 5' UTR | 93020 |
| 12 | 29618635 | 29619135 | Myt1l | 5' UTR | 84020 |
| 12 | 29616135 | 29616635 | Myt1l | 5' UTR | 81520 |
| 12 | 29616635 | 29617135 | Myt1l | Non-coding | 81413 |
| 12 | 29627635 | 29628135 | Myt1l | Non-coding | 92413 |
| 12 | 29618635 | 29619135 | Myt1l | Non-coding | 83413 |
| 12 | 29616135 | 29616635 | Myt1l | Non-coding | 80913 |
| 12 | 29616635 | 29617135 | Myt1l | Non-coding | 49935 |
| 12 | 29627635 | 29628135 | Myt1l | Non-coding | 60935 |
| 12 | 29618635 | 29619135 | Myt1l | Non-coding | 51935 |
| 12 | 29616135 | 29616635 | Myt1l | Non-coding | 49435 |
| 12 | 29894635 | 29895135 | Myt1l | Promoter | 0 |
| 12 | 29894635 | 29895135 | Myt1l | 5' UTR | 0 |
| 12 | 29894635 | 29895135 | Myt1l | CDS 1 | 0 |
| 12 | 30103135 | 30103635 | Tpo | CDS 8 | 28990 |
| 12 | 30103135 | 30103635 | Tpo | Intron 7 | 28990 |
| 12 | 30416635 | 30417135 | 2310016D03Rik | Non-coding | 50224 |
| 12 | 31295635 | 31296135 | Lamb1 | Intron 13 | 30401 |
| 12 | 31295635 | 31296135 | Lamb1 | Intron 12 | 30342 |
| 12 | 31973635 | 31974135 | Prkar2b | Intron 5 | 87101 |
| 12 | 31973635 | 31974135 | Prkar2b | Intron 5 | 87162 |
| 12 | 31973635 | 31974135 | Prkar2b | Intron 5 | 86455 |
| 12 | 32929635 | 32930135 | 4933406C10Rik | Non-coding | 23655 |
| 12 | 32930135 | 32930635 | 4933406C10Rik | Non-coding | 23155 |
| 12 | 32929635 | 32930135 | 4933406C10Rik | Non-coding | 23649 |
| 12 | 32930135 | 32930635 | 4933406C10Rik | Non-coding | 23149 |
| 12 | 32929635 | 32930135 | Gm19056 | Promoter | 0 |
| 12 | 32930135 | 32930635 | Gm19056 | Promoter | -187 |
| 12 | 32929635 | 32930135 | Gm18622 | Promoter | 0 |
| 12 | 32930135 | 32930635 | Gm18622 | Promoter | 11 |
| 12 | 32929635 | 32930135 | Gm47961 | Promoter | 1598 |

| | | | | | |
|---|---|---|---|---|---|
| 12 | 32930135 | 32930635 | Gm47961 | Promoter | 1098 |
| 12 | 33382635 | 33383135 | Atxn7l1os2 | Non-coding | 11626 |
| 12 | 34693135 | 34693635 | Hdac9 | Intron 3 | 223461 |
| 12 | 34693135 | 34693635 | Hdac9 | Intron 2 | 213423 |
| 12 | 34693135 | 34693635 | Hdac9 | Non-coding | 213345 |
| 12 | 35529328 | 35529828 | Ahr | Intron 1 | 5211 |
| 12 | 35529328 | 35529828 | Ahr | Intron 1 | 4796 |
| 12 | 35529328 | 35529828 | Ahr | Promoter | -2548 |
| 12 | 35529328 | 35529828 | Gm48236 | Promoter | -4593 |
| 12 | 40472413 | 40472913 | Dock4 | Intron 1 | 26461 |
| 12 | 40472413 | 40472913 | Dock4 | Non-coding | 26377 |
| 12 | 40472413 | 40472913 | Dock4 | Non-coding | 26374 |
| 12 | 40472413 | 40472913 | Dock4 | Non-coding | 26360 |
| 12 | 40472413 | 40472913 | Dock4 | Non-coding | 26325 |
| 12 | 40472413 | 40472913 | Dock4 | Intron 1 | 26077 |
| 12 | 41483413 | 41483913 | Immp2l | Intron 4 | 459323 |
| 12 | 41483413 | 41483913 | Immp2l | Intron 3 | 459084 |
| 12 | 41483413 | 41483913 | Lrrn3 | Promoter | 2519 |
| 12 | 41483413 | 41483913 | Lrrn3 | 5' UTR | 2519 |
| 12 | 50770013 | 50770513 | Gm48779 | Non-coding | 16916 |
| 12 | 50997513 | 50998013 | Gm40421 | Non-coding | 6861 |
| 12 | 51769013 | 51769513 | Hectd1 | CDS 25 | 60024 |
| 12 | 51769013 | 51769513 | Hectd1 | Intron 24 | 60024 |
| 12 | 51769013 | 51769513 | Hectd1 | CDS 25 | 60024 |
| 12 | 51769013 | 51769513 | Hectd1 | Intron 24 | 60024 |
| 12 | 51769013 | 51769513 | Hectd1 | Promoter | -3781 |
| 12 | 51843513 | 51844013 | Gm19309 | Promoter | -2891 |
| 12 | 52447513 | 52448013 | Gm47431 | Promoter | 113 |
| 12 | 52963513 | 52964013 | Akap6 | Intron 7 | 264130 |
| 12 | 52944513 | 52945013 | Akap6 | Intron 7 | 245130 |
| 12 | 52963513 | 52964013 | Akap6 | Intron 6 | 167677 |
| 12 | 52944513 | 52945013 | Akap6 | Intron 6 | 148677 |
| 12 | 56545513 | 56546013 | Gm26973 | Non-coding | 10170 |
| 12 | 56545513 | 56546013 | Gm26973 | Non-coding | 8148 |
| 12 | 56545513 | 56546013 | Gm26973 | Non-coding | 8097 |
| 12 | 56958013 | 56958513 | Slc25a21 | Intron 1 | 238960 |
| 12 | 56958013 | 56958513 | Slc25a21 | Intron 2 | 202131 |
| 12 | 56958013 | 56958513 | Slc25a21 | Intron 1 | 200732 |
| 12 | 56958013 | 56958513 | Slc25a21 | Non-coding | 21881 |
| 12 | 58925013 | 58925513 | Gm47647 | Promoter | -553 |
| 12 | 58925013 | 58925513 | Gm47647 | Promoter | -718 |
| 12 | 69157513 | 69158013 | Rps29 | Promoter | 1174 |
| 12 | 69157513 | 69158013 | Rps29 | 3' UTR | 1174 |
| 12 | 69157513 | 69158013 | Rps29 | CDS 3 | 1174 |
| 12 | 69157513 | 69158013 | Rps29 | Intron 2 | 1174 |
| 12 | 69157513 | 69158013 | Rps29 | Promoter | 1042 |

| | | | | | |
|---|---|---|---|---|---|
| 12 | 69157513 | 69158013 | Rn7s1 | Promoter | -1282 |
| 12 | 69157513 | 69158013 | Gm47287 | Promoter | -1424 |
| 12 | 69157513 | 69158013 | Gm47295 | Promoter | -1505 |
| 12 | 70974013 | 70974513 | Psma3 | Promoter | -108 |
| 12 | 70974013 | 70974513 | Psma3 | Promoter | -158 |
| 12 | 70974013 | 70974513 | Psma3 | Promoter | -158 |
| 12 | 70974013 | 70974513 | Psma3 | Promoter | -170 |
| 12 | 70974013 | 70974513 | Psma3 | Promoter | -176 |
| 12 | 70974013 | 70974513 | Psma3 | Promoter | -195 |
| 12 | 70974013 | 70974513 | Psma3 | Promoter | -202 |
| 12 | 70974013 | 70974513 | Psma3 | Promoter | -206 |
| 12 | 72116013 | 72116513 | Ccdc175 | Intron 14 | 68517 |
| 12 | 75175513 | 75176013 | Kcnh5 | Promoter | 1320 |
| 12 | 75175513 | 75176013 | Kcnh5 | Intron 1 | 1320 |
| 12 | 80223513 | 80224013 | Actn1 | Intron 1 | 36359 |
| 12 | 80223513 | 80224013 | Actn1 | Intron 1 | 36079 |
| 12 | 80223513 | 80224013 | Actn1 | Non-coding | 35878 |
| 12 | 80223513 | 80224013 | Actn1 | Non-coding | 34772 |
| 12 | 81975013 | 81975513 | Pcnx | Intron 22 | 114990 |
| 12 | 81975013 | 81975513 | Pcnx | CDS 23 | 114990 |
| 12 | 81975013 | 81975513 | Pcnx | Intron 23 | 114990 |
| 12 | 81975013 | 81975513 | Pcnx | Intron 22 | 114983 |
| 12 | 81975013 | 81975513 | Pcnx | CDS 23 | 114983 |
| 12 | 81975013 | 81975513 | Pcnx | Intron 23 | 114983 |
| 12 | 81975013 | 81975513 | Pcnx | 3' UTR | 114672 |
| 12 | 81975013 | 81975513 | Pcnx | Intron 15 | 56370 |
| 12 | 81975013 | 81975513 | Pcnx | CDS 16 | 56370 |
| 12 | 81975013 | 81975513 | Pcnx | Intron 16 | 56370 |
| 12 | 81975013 | 81975513 | Pcnx | Promoter | 1826 |
| 12 | 82171013 | 82171513 | Gm48242 | Promoter | -1907 |
| 12 | 82179013 | 82179513 | Sipa1l1 | Non-coding | 9693 |
| 12 | 82171013 | 82171513 | Sipa1l1 | Promoter | 1693 |
| 12 | 82179013 | 82179513 | Sipa1l1 | 5' UTR | 9593 |
| 12 | 82171013 | 82171513 | Sipa1l1 | Promoter | 1593 |
| 12 | 82171013 | 82171513 | Sipa1l1 | 5' UTR | 1593 |
| 12 | 82179013 | 82179513 | Sipa1l1 | 5' UTR | 9567 |
| 12 | 82171013 | 82171513 | Sipa1l1 | Promoter | 1567 |
| 12 | 82171013 | 82171513 | Sipa1l1 | 5' UTR | 1567 |
| 12 | 82179013 | 82179513 | Sipa1l1 | Non-coding | 9538 |
| 12 | 82171013 | 82171513 | Sipa1l1 | Promoter | 1538 |
| 12 | 82179013 | 82179513 | Sipa1l1 | Non-coding | 9537 |
| 12 | 82171013 | 82171513 | Sipa1l1 | Promoter | 1537 |
| 12 | 82179013 | 82179513 | Sipa1l1 | 5' UTR | 8987 |
| 12 | 82171013 | 82171513 | Sipa1l1 | Promoter | 987 |
| 12 | 82171013 | 82171513 | Sipa1l1 | 5' UTR | 987 |
| 12 | 85336050 | 85336550 | Nek9 | Promoter | 2813 |

| | | | | | |
|---|---|---|---|---|---|
| 12 | 85336050 | 85336550 | Nek9 | Intron 2 | 2813 |
| 12 | 85336050 | 85336550 | Nek9 | CDS 2 | 2813 |
| 12 | 85336050 | 85336550 | Nek9 | Intron 1 | 2813 |
| 12 | 86761050 | 86761550 | Lrrc74a | Intron 11 | 26681 |
| 12 | 86761050 | 86761550 | Lrrc74a | Intron 11 | 26667 |
| 12 | 86761050 | 86761550 | Lrrc74a | 3' UTR | 26622 |
| 12 | 87052550 | 87053050 | Tmem63c | 5' UTR | 31210 |
| 12 | 87052550 | 87053050 | Tmem63c | 5' UTR | 26258 |
| 12 | 87052550 | 87053050 | Tmem63c | 5' UTR | 25986 |
| 12 | 87052550 | 87053050 | Tmem63c | 5' UTR | 25934 |
| 12 | 87052550 | 87053050 | Tmem63c | 5' UTR | 14111 |
| 12 | 89135056 | 89135556 | Nrxn3 | Non-coding | 412180 |
| 12 | 89135056 | 89135556 | Nrxn3 | 5' UTR | 412070 |
| 12 | 89406556 | 89407056 | Nrxn3 | Intron 7 | 683570 |
| 12 | 89368056 | 89368556 | Nrxn3 | Intron 7 | 645070 |
| 12 | 89135056 | 89135556 | Nrxn3 | Intron 4 | 411947 |
| 12 | 89406556 | 89407056 | Nrxn3 | Intron 9 | 683447 |
| 12 | 89368056 | 89368556 | Nrxn3 | Intron 9 | 644947 |
| 12 | 89135056 | 89135556 | Nrxn3 | Intron 3 | 340552 |
| 12 | 89406556 | 89407056 | Nrxn3 | Intron 9 | 612052 |
| 12 | 89368056 | 89368556 | Nrxn3 | Intron 9 | 573552 |
| 12 | 89135056 | 89135556 | Nrxn3 | Non-coding | 181666 |
| 12 | 89135056 | 89135556 | Nrxn3 | 5' UTR | 181657 |
| 12 | 89406556 | 89407056 | Nrxn3 | Intron 8 | 453157 |
| 12 | 89368056 | 89368556 | Nrxn3 | Intron 8 | 414657 |
| 12 | 89406556 | 89407056 | Nrxn3 | Intron 5 | 213450 |
| 12 | 89368056 | 89368556 | Nrxn3 | Intron 5 | 174950 |
| 12 | 91400060 | 91400560 | Tshr | 5' UTR | 15494 |
| 12 | 91400060 | 91400560 | Tshr | Promoter | -434 |
| 12 | 91400060 | 91400560 | Tshr | Promoter | -440 |
| 12 | 99282290 | 99282790 | Foxn3 | Intron 4 | 111008 |
| 12 | 99282290 | 99282790 | Foxn3 | Intron 4 | 167285 |
| 12 | 99282290 | 99282790 | Foxn3 | Intron 4 | 110283 |
| 12 | 99282290 | 99282790 | Foxn3 | Intron 1 | 8489 |
| 12 | 99282290 | 99282790 | Foxn3 | Non-coding | 47274 |
| 12 | 99282290 | 99282790 | Foxn3 | Non-coding | 8600 |
| 12 | 99282290 | 99282790 | Foxn3 | Non-coding | 8488 |
| 12 | 100128790 | 100129290 | Nrde2 | Intron 12 | 30359 |
| 12 | 100128790 | 100129290 | Nrde2 | Promoter | 483 |
| 12 | 102559290 | 102559790 | Chga | CDS 4 | 4321 |
| 12 | 102559290 | 102559790 | Chga | Intron 4 | 4321 |
| 12 | 102559290 | 102559790 | Chga | Promoter | 1222 |
| 12 | 103236290 | 103236790 | Prima1 | Intron 2 | 5361 |
| 12 | 103236290 | 103236790 | Prima1 | Intron 2 | 5357 |
| 12 | 103236290 | 103236790 | Prima1 | Intron 2 | 5339 |
| 12 | 103236290 | 103236790 | Prima1 | Promoter | -265 |

| | | | | | |
|---|---|---|---|---|---|
| 12 | 103316290 | 103316790 | Fam181a | 3' UTR | 5315 |
| 12 | 103316290 | 103316790 | Fam181a | CDS 2 | 5315 |
| 12 | 103316290 | 103316790 | Gm29508 | Promoter | -2284 |
| 12 | 103316290 | 103316790 | Gm29508 | Promoter | -1520 |
| 12 | 103316290 | 103316790 | Fam181a | Promoter | 1337 |
| 12 | 103316290 | 103316790 | Fam181a | CDS 2 | 1337 |
| 12 | 103316290 | 103316790 | Fam181a | Promoter | 1331 |
| 12 | 103316290 | 103316790 | Fam181a | 3' UTR | 1331 |
| 12 | 103316290 | 103316790 | Fam181a | CDS 2 | 1331 |
| 12 | 104986790 | 104987290 | Syne3 | 5' UTR | 22520 |
| 12 | 104986790 | 104987290 | Syne3 | 5' UTR | 11388 |
| 12 | 104986790 | 104987290 | Syne3 | 5' UTR | 11388 |
| 12 | 106098790 | 106099290 | Gm32635 | Promoter | 0 |
| 12 | 107729290 | 107729790 | 4930465M20Rik | Non-coding | 5699 |
| 12 | 107729290 | 107729790 | 4930465M20Rik | Non-coding | 5707 |
| 12 | 107924290 | 107924790 | Bcl11b | Intron 3 | 78813 |
| 12 | 107924290 | 107924790 | Bcl11b | Intron 1 | 78625 |
| 12 | 107924290 | 107924790 | Bcl11b | Intron 2 | 78625 |
| 12 | 108959290 | 108959790 | Wdr25 | Intron 2 | 65018 |
| 12 | 108959290 | 108959790 | Wdr25 | Intron 2 | 64997 |
| 12 | 108959290 | 108959790 | Wdr25 | Intron 3 | 64973 |
| 12 | 108959290 | 108959790 | Wdr25 | Non-coding | 46558 |
| 12 | 108959290 | 108959790 | Gm47980 | Promoter | -1122 |
| 12 | 109459790 | 109460290 | Dlk1 | CDS 6 | 6941 |
| 12 | 109459790 | 109460290 | Dlk1 | CDS 5 | 6335 |
| 12 | 109459790 | 109460290 | Dlk1 | CDS 5 | 6299 |
| 12 | 109459790 | 109460290 | Dlk1 | Intron 5 | 6299 |
| 12 | 109459790 | 109460290 | Dlk1 | CDS 6 | 6299 |
| 12 | 109459790 | 109460290 | Dlk1 | CDS 5 | 6299 |
| 12 | 109459790 | 109460290 | Dlk1 | Intron 5 | 6299 |
| 12 | 109459790 | 109460290 | Dlk1 | CDS 6 | 6299 |
| 12 | 109459790 | 109460290 | Dlk1 | CDS 5 | 6255 |
| 12 | 109459790 | 109460290 | Dlk1 | Intron 5 | 6255 |
| 12 | 109459790 | 109460290 | Dlk1 | CDS 6 | 6255 |
| 12 | 109459790 | 109460290 | Dlk1 | CDS 5 | 6135 |
| 12 | 109459790 | 109460290 | Dlk1 | Intron 5 | 6135 |
| 12 | 109459790 | 109460290 | Dlk1 | CDS 6 | 6135 |
| 12 | 109459790 | 109460290 | Dlk1 | CDS 5 | 6130 |
| 12 | 109459790 | 109460290 | Dlk1 | Intron 5 | 6130 |
| 12 | 109459790 | 109460290 | Dlk1 | CDS 6 | 6130 |
| 12 | 109459790 | 109460290 | Dlk1 | Promoter | 0 |
| 12 | 109459790 | 109460290 | Dlk1 | 5' UTR | 0 |
| 12 | 109459790 | 109460290 | Dlk1 | CDS 1 | 0 |
| 12 | 109459790 | 109460290 | Dlk1 | Intron 1 | 0 |

| | | | | | |
|---|---|---|---|---|---|
| 12 | 109459790 | 109460290 | Dlk1 | CDS 2 | 0 |
| 12 | 111161289 | 111161789 | 4930595D18Rik | Intron 3 | 22703 |
| 12 | 111161289 | 111161789 | 4930595D18Rik | CDS 3 | 22703 |
| 12 | 111161289 | 111161789 | Traf3 | Promoter | -4581 |
| 12 | 111161289 | 111161789 | Traf3 | Promoter | -4674 |
| 12 | 111161289 | 111161789 | Traf3 | Promoter | -4697 |
| 12 | 111161289 | 111161789 | Traf3 | Promoter | -4759 |
| 12 | 111357289 | 111357789 | Cdc42bpb | Intron 1 | 19930 |
| 12 | 111357289 | 111357789 | Cdc42bpb | Intron 2 | 19831 |
| 12 | 111539289 | 111539789 | Eif5 | Promoter | 1273 |
| 12 | 111539289 | 111539789 | Eif5 | 5' UTR | 1273 |
| 12 | 111539289 | 111539789 | Eif5 | Intron 2 | 1273 |
| 12 | 111539289 | 111539789 | Eif5 | CDS 3 | 1273 |
| 12 | 111539289 | 111539789 | Eif5 | Intron 3 | 1273 |
| 12 | 111539789 | 111540289 | Eif5 | Promoter | 1773 |
| 12 | 111539789 | 111540289 | Eif5 | Intron 3 | 1773 |
| 12 | 111539789 | 111540289 | Eif5 | CDS 4 | 1773 |
| 12 | 111539789 | 111540289 | Eif5 | Intron 4 | 1773 |
| 12 | 111539789 | 111540289 | Eif5 | CDS 5 | 1773 |
| 12 | 111539289 | 111539789 | Eif5 | Promoter | 1092 |
| 12 | 111539289 | 111539789 | Eif5 | 5' UTR | 1092 |
| 12 | 111539289 | 111539789 | Eif5 | Intron 1 | 1092 |
| 12 | 111539289 | 111539789 | Eif5 | CDS 2 | 1092 |
| 12 | 111539289 | 111539789 | Eif5 | Intron 2 | 1092 |
| 12 | 111539789 | 111540289 | Eif5 | Promoter | 1592 |
| 12 | 111539789 | 111540289 | Eif5 | Intron 2 | 1592 |
| 12 | 111539789 | 111540289 | Eif5 | CDS 3 | 1592 |
| 12 | 111539789 | 111540289 | Eif5 | Intron 3 | 1592 |
| 12 | 111539789 | 111540289 | Eif5 | CDS 4 | 1592 |
| 12 | 111539289 | 111539789 | Eif5 | Promoter | 1073 |
| 12 | 111539789 | 111540289 | Eif5 | Promoter | 1573 |
| 12 | 111539289 | 111539789 | Eif5 | Promoter | 1073 |
| 12 | 111539789 | 111540289 | Eif5 | Promoter | 1573 |
| 12 | 111539289 | 111539789 | Eif5 | Promoter | 1073 |
| 12 | 111539289 | 111539789 | Eif5 | 5' UTR | 1073 |
| 12 | 111539289 | 111539789 | Eif5 | Intron 2 | 1073 |
| 12 | 111539289 | 111539789 | Eif5 | CDS 3 | 1073 |
| 12 | 111539289 | 111539789 | Eif5 | Intron 3 | 1073 |
| 12 | 111539789 | 111540289 | Eif5 | Promoter | 1573 |
| 12 | 111539789 | 111540289 | Eif5 | Intron 3 | 1573 |
| 12 | 111539789 | 111540289 | Eif5 | CDS 4 | 1573 |
| 12 | 111539789 | 111540289 | Eif5 | Intron 4 | 1573 |
| 12 | 111539789 | 111540289 | Eif5 | CDS 5 | 1573 |
| 12 | 111539289 | 111539789 | Eif5 | Promoter | 1073 |

| | | | | | |
|---|---|---|---|---|---|
| 12 | 111539289 | 111539789 | Eif5 | 5' UTR | 1073 |
| 12 | 111539289 | 111539789 | Eif5 | Intron 2 | 1073 |
| 12 | 111539289 | 111539789 | Eif5 | CDS 3 | 1073 |
| 12 | 111539289 | 111539789 | Eif5 | Intron 3 | 1073 |
| 12 | 111539789 | 111540289 | Eif5 | Promoter | 1573 |
| 12 | 111539789 | 111540289 | Eif5 | Intron 3 | 1573 |
| 12 | 111539789 | 111540289 | Eif5 | CDS 4 | 1573 |
| 12 | 111539789 | 111540289 | Eif5 | Intron 4 | 1573 |
| 12 | 111539789 | 111540289 | Eif5 | CDS 5 | 1573 |
| 12 | 111539289 | 111539789 | Eif5 | Promoter | 936 |
| 12 | 111539289 | 111539789 | Eif5 | 5' UTR | 936 |
| 12 | 111539289 | 111539789 | Eif5 | Intron 1 | 936 |
| 12 | 111539289 | 111539789 | Eif5 | CDS 2 | 936 |
| 12 | 111539289 | 111539789 | Eif5 | Intron 2 | 936 |
| 12 | 111539789 | 111540289 | Eif5 | Promoter | 1436 |
| 12 | 111539789 | 111540289 | Eif5 | Intron 2 | 1436 |
| 12 | 111539789 | 111540289 | Eif5 | CDS 3 | 1436 |
| 12 | 111539289 | 111539789 | Eif5 | Promoter | 934 |
| 12 | 111539789 | 111540289 | Eif5 | Promoter | 1434 |
| 12 | 111539289 | 111539789 | Eif5 | Promoter | 507 |
| 12 | 111539289 | 111539789 | Eif5 | 5' UTR | 507 |
| 12 | 111539289 | 111539789 | Eif5 | Intron 1 | 507 |
| 12 | 111539289 | 111539789 | Eif5 | CDS 2 | 507 |
| 12 | 111539289 | 111539789 | Eif5 | Intron 2 | 507 |
| 12 | 111539789 | 111540289 | Eif5 | Promoter | 1007 |
| 12 | 111539789 | 111540289 | Eif5 | Intron 2 | 1007 |
| 12 | 111539789 | 111540289 | Eif5 | CDS 3 | 1007 |
| 12 | 111539789 | 111540289 | Eif5 | Intron 3 | 1007 |
| 12 | 111539789 | 111540289 | Eif5 | CDS 4 | 1007 |
| 12 | 111539289 | 111539789 | Eif5 | Promoter | 458 |
| 12 | 111539289 | 111539789 | Eif5 | 5' UTR | 458 |
| 12 | 111539289 | 111539789 | Eif5 | Intron 1 | 458 |
| 12 | 111539289 | 111539789 | Eif5 | CDS 2 | 458 |
| 12 | 111539289 | 111539789 | Eif5 | Intron 2 | 458 |
| 12 | 111539789 | 111540289 | Eif5 | Promoter | 958 |
| 12 | 111539789 | 111540289 | Eif5 | Intron 2 | 958 |
| 12 | 111539789 | 111540289 | Eif5 | CDS 3 | 958 |
| 12 | 111539289 | 111539789 | Eif5 | Promoter | 258 |
| 12 | 111539789 | 111540289 | Eif5 | Promoter | 758 |
| 12 | 111539289 | 111539789 | Eif5 | Promoter | 218 |
| 12 | 111539289 | 111539789 | Eif5 | 5' UTR | 218 |
| 12 | 111539289 | 111539789 | Eif5 | CDS 1 | 218 |
| 12 | 111539289 | 111539789 | Eif5 | Intron 1 | 218 |
| 12 | 111539789 | 111540289 | Eif5 | Promoter | 718 |
| 12 | 111539789 | 111540289 | Eif5 | Intron 1 | 718 |
| 12 | 111539789 | 111540289 | Eif5 | CDS 2 | 718 |

| | | | | | |
|---|---|---|---|---|---|
| 12 | 111539789 | 111540289 | Eif5 | Intron 2 | 718 |
| 12 | 111539789 | 111540289 | Eif5 | CDS 3 | 718 |
| 12 | 111539289 | 111539789 | Eif5 | Promoter | 0 |
| 12 | 111539289 | 111539789 | Eif5 | 5' UTR | 0 |
| 12 | 111539289 | 111539789 | Eif5 | CDS 1 | 0 |
| 12 | 111539289 | 111539789 | Eif5 | Intron 1 | 0 |
| 12 | 111539789 | 111540289 | Eif5 | Promoter | 351 |
| 12 | 111539789 | 111540289 | Eif5 | 3' UTR | 351 |
| 12 | 111539789 | 111540289 | Eif5 | Intron 1 | 351 |
| 12 | 111539789 | 111540289 | Eif5 | CDS 2 | 351 |
| 12 | 111539789 | 111540289 | Eif5 | Intron 2 | 351 |
| 12 | 111539789 | 111540289 | Eif5 | CDS 3 | 351 |
| 12 | 111539289 | 111539789 | Eif5 | Promoter | 0 |
| 12 | 111539789 | 111540289 | Eif5 | Promoter | 210 |
| 12 | 111539289 | 111539789 | Eif5 | Promoter | 0 |
| 12 | 111539789 | 111540289 | Eif5 | Promoter | 146 |
| 12 | 111539289 | 111539789 | Snora28 | Promoter | -1152 |
| 12 | 111539789 | 111540289 | Snora28 | Promoter | -652 |
| 12 | 111539289 | 111539789 | Eif5 | Promoter | -2436 |
| 12 | 111539789 | 111540289 | Eif5 | Promoter | -1936 |
| 12 | 111539289 | 111539789 | Eif5 | Promoter | -3042 |
| 12 | 111539789 | 111540289 | Eif5 | Promoter | -2542 |
| 12 | 112131289 | 112131789 | Mir203 | Promoter | 409 |
| 12 | 112793789 | 112794289 | Ahnak2 | Intron 1 | 8369 |
| 12 | 117743527 | 117744027 | Rapgef5 | Intron 24 | 227048 |
| 12 | 117743527 | 117744027 | Rapgef5 | Intron 14 | 85493 |
| 12 | 117743527 | 117744027 | Rapgef5 | Intron 12 | 54691 |
| 12 | 117743527 | 117744027 | Rapgef5 | Promoter | 2932 |
| 13 | 24296131 | 24296631 | Gm11340 | Promoter | -4325 |
| 13 | 31797631 | 31798131 | Gm11379 | Promoter | 92 |
| 13 | 38185131 | 38185631 | Dsp | CDS 16 | 33837 |
| 13 | 38185131 | 38185631 | Dsp | Intron 16 | 33837 |
| 13 | 38185131 | 38185631 | Dsp | CDS 17 | 33837 |
| 13 | 38185131 | 38185631 | Dsp | CDS 16 | 33803 |
| 13 | 38185131 | 38185631 | Dsp | Intron 16 | 33803 |
| 13 | 38185131 | 38185631 | Dsp | CDS 17 | 33803 |
| 13 | 38937131 | 38937631 | Slc35b3 | Intron 8 | 23245 |
| 13 | 38937131 | 38937631 | Slc35b3 | CDS 8 | 23245 |
| 13 | 38937131 | 38937631 | Slc35b3 | Intron 7 | 23245 |
| 13 | 38937131 | 38937631 | Slc35b3 | Intron 8 | 23194 |
| 13 | 38937131 | 38937631 | Slc35b3 | CDS 8 | 23194 |
| 13 | 38937131 | 38937631 | Slc35b3 | Intron 7 | 23194 |
| 13 | 38937131 | 38937631 | Slc35b3 | Intron 8 | 22899 |
| 13 | 38937131 | 38937631 | Slc35b3 | CDS 8 | 22899 |
| 13 | 38937131 | 38937631 | Slc35b3 | Intron 7 | 22899 |
| 13 | 38937131 | 38937631 | Slc35b3 | 3' UTR | 22866 |

| | | | | | |
|---|---|---|---|---|---|
| 13 | 38937131 | 38937631 | Slc35b3 | Intron 4 | 5522 |
| 13 | 38937131 | 38937631 | Slc35b3 | CDS 4 | 5522 |
| 13 | 38937131 | 38937631 | Slc35b3 | Intron 3 | 5522 |
| 13 | 38937131 | 38937631 | Slc35b3 | Intron 7 | 22907 |
| 13 | 38937131 | 38937631 | Slc35b3 | CDS 7 | 22907 |
| 13 | 38937131 | 38937631 | Slc35b3 | Intron 6 | 22907 |
| 13 | 38937131 | 38937631 | Slc35b3 | Intron 7 | 22877 |
| 13 | 38937131 | 38937631 | Slc35b3 | CDS 7 | 22877 |
| 13 | 38937131 | 38937631 | Slc35b3 | Intron 6 | 22877 |
| 13 | 38937131 | 38937631 | Slc35b3 | 3' UTR | 22855 |
| 13 | 38937131 | 38937631 | Slc35b3 | 3' UTR | 22883 |
| 13 | 40887131 | 40887631 | Gcnt2 | Intron 1 | 27377 |
| 13 | 40887131 | 40887631 | Gcnt2 | Promoter | 0 |
| 13 | 40887131 | 40887631 | Gcnt2 | 5' UTR | 0 |
| 13 | 40887131 | 40887631 | Gcnt2 | CDS 1 | 0 |
| 13 | 41042131 | 41042631 | Mak | CDS 9 | 27839 |
| 13 | 41042131 | 41042631 | Mak | Intron 8 | 27839 |
| 13 | 41042131 | 41042631 | Mak | CDS 9 | 27835 |
| 13 | 41042131 | 41042631 | Mak | Intron 8 | 27835 |
| 13 | 41077631 | 41078131 | Mak | Promoter | 1576 |
| 13 | 41077631 | 41078131 | Mak | 5' UTR | 1576 |
| 13 | 41042131 | 41042631 | Mak | CDS 10 | 37076 |
| 13 | 41042131 | 41042631 | Mak | Intron 9 | 37076 |
| 13 | 41077631 | 41078131 | Mak | Promoter | 1512 |
| 13 | 41042131 | 41042631 | Mak | Non-coding | 37012 |
| 13 | 41077631 | 41078131 | Mak | Promoter | 1574 |
| 13 | 41077631 | 41078131 | Mak | 5' UTR | 1574 |
| 13 | 41042131 | 41042631 | Mak | CDS 9 | 37074 |
| 13 | 41042131 | 41042631 | Mak | Intron 8 | 37074 |
| 13 | 41042131 | 41042631 | Mak | CDS 9 | 27829 |
| 13 | 41042131 | 41042631 | Mak | Intron 8 | 27829 |
| 13 | 41042131 | 41042631 | Mak | Promoter | -2359 |
| 13 | 41077631 | 41078131 | Mak | Promoter | 1408 |
| 13 | 41077631 | 41078131 | Mak | 5' UTR | 1408 |
| 13 | 42952131 | 42952631 | Phactr1 | Intron 3 | 271508 |
| 13 | 42952131 | 42952631 | Phactr1 | Non-coding | 271332 |
| 13 | 42952131 | 42952631 | Phactr1 | Intron 3 | 270618 |
| 13 | 42952131 | 42952631 | Phactr1 | Intron 2 | 269433 |
| 13 | 42952131 | 42952631 | Phactr1 | Intron 1 | 242588 |
| 13 | 42952131 | 42952631 | Phactr1 | Intron 1 | 242550 |
| 13 | 42952131 | 42952631 | Phactr1 | 5' UTR | 85884 |
| 13 | 42952131 | 42952631 | Phactr1 | Non-coding | 85297 |
| 13 | 42952131 | 42952631 | Phactr1 | Intron 1 | 35480 |
| 13 | 43300631 | 43301131 | Gfod1 | Intron 1 | 3042 |
| 13 | 43300631 | 43301131 | Gfod1 | Promoter | 2275 |
| 13 | 43300631 | 43301131 | Gfod1 | Intron 1 | 2275 |

| | | | | | |
|---|---|---|---|---|---|
| 13 | 43797131 | 43797631 | Cd83 | Intron 2 | 12356 |
| 13 | 43797131 | 43797631 | Cd83 | CDS 3 | 12356 |
| 13 | 43797131 | 43797631 | Cd83 | Non-coding | 11946 |
| 13 | 52903131 | 52903631 | Auh | Intron 3 | 26031 |
| 13 | 52903131 | 52903631 | Auh | Intron 2 | 15602 |
| 13 | 52903131 | 52903631 | Auh | Intron 3 | 25828 |
| 13 | 52903131 | 52903631 | Auh | Non-coding | 26018 |
| 13 | 52903131 | 52903631 | Auh | Intron 3 | 26022 |
| 13 | 52903131 | 52903631 | Auh | Non-coding | 20235 |
| 13 | 54564139 | 54564639 | 4833439L19Rik | Promoter | 797 |
| 13 | 54564139 | 54564639 | 4833439L19Rik | 5' UTR | 797 |
| 13 | 54564139 | 54564639 | 4833439L19Rik | Intron 3 | 797 |
| 13 | 54564139 | 54564639 | 4833439L19Rik | CDS 3 | 797 |
| 13 | 54564139 | 54564639 | 4833439L19Rik | Intron 2 | 797 |
| 13 | 54564139 | 54564639 | 4833439L19Rik | CDS 2 | 797 |
| 13 | 54564139 | 54564639 | 4833439L19Rik | Intron 1 | 797 |
| 13 | 54564139 | 54564639 | 4833439L19Rik | Promoter | 665 |
| 13 | 54564139 | 54564639 | 4833439L19Rik | 5' UTR | 665 |
| 13 | 54564139 | 54564639 | 4833439L19Rik | Promoter | 695 |
| 13 | 54564139 | 54564639 | 4833439L19Rik | Promoter | 653 |
| 13 | 54564139 | 54564639 | 4833439L19Rik | Promoter | 700 |
| 13 | 54564139 | 54564639 | 4833439L19Rik | Promoter | 692 |
| 13 | 54564139 | 54564639 | 4833439L19Rik | 5' UTR | 692 |
| 13 | 54564139 | 54564639 | 4833439L19Rik | Intron 3 | 692 |
| 13 | 54564139 | 54564639 | 4833439L19Rik | CDS 3 | 692 |
| 13 | 54564139 | 54564639 | 4833439L19Rik | Intron 2 | 692 |
| 13 | 54564139 | 54564639 | 4833439L19Rik | CDS 2 | 692 |
| 13 | 54564139 | 54564639 | 4833439L19Rik | Intron 1 | 692 |
| 13 | 54564139 | 54564639 | 4833439L19Rik | Promoter | 706 |
| 13 | 54564139 | 54564639 | 4833439L19Rik | Promoter | 718 |
| 13 | 54564139 | 54564639 | 4833439L19Rik | 5' UTR | 718 |
| 13 | 54564139 | 54564639 | 4833439L19Rik | Intron 2 | 718 |
| 13 | 54564139 | 54564639 | 4833439L19Rik | CDS 2 | 718 |
| 13 | 54564139 | 54564639 | 4833439L19Rik | Intron 1 | 718 |
| 13 | 54564139 | 54564639 | 4833439L19Rik | Promoter | 687 |
| 13 | 54564139 | 54564639 | 4833439L19Rik | 5' UTR | 687 |
| 13 | 55962139 | 55962639 | Gm47071 | Non-coding | 135889 |
| 13 | 57450139 | 57450639 | Spock1 | Intron 6 | 457676 |
| 13 | 57450139 | 57450639 | Spock1 | Intron 7 | 457694 |
| 13 | 57450139 | 57450639 | Spock1 | Intron 7 | 457637 |
| 13 | 57450139 | 57450639 | Spock1 | Intron 6 | 457653 |
| 13 | 57450139 | 57450639 | Spock1 | Intron 5 | 456949 |
| 13 | 57450139 | 57450639 | Spock1 | Intron 6 | 457661 |
| 13 | 60056139 | 60056639 | A530065N20Rik | Non-coding | 26428 |
| 13 | 60056139 | 60056639 | Gm48396 | Promoter | 987 |

| | | | | | |
|---|---|---|---|---|---|
| 13 | 60427639 | 60428139 | Gm35333 | Non-coding | 6593 |
| 13 | 60427639 | 60428139 | Gm35333 | Promoter | -440 |
| 13 | 63247691 | 63248191 | Aopep | Intron 12 | 232757 |
| 13 | 63247691 | 63248191 | Aopep | Intron 12 | 232524 |
| 13 | 63247691 | 63248191 | Aopep | Intron 12 | 232198 |
| 13 | 63247691 | 63248191 | Aopep | Intron 3 | 56579 |
| 13 | 63247691 | 63248191 | Aopep | Intron 2 | 7662 |
| 13 | 63247691 | 63248191 | Aopep | Non-coding | 7646 |
| 13 | 63247691 | 63248191 | Aopep | Intron 2 | 7617 |
| 13 | 63247691 | 63248191 | Aopep | Non-coding | 7617 |
| 13 | 63247691 | 63248191 | Aopep | Intron 2 | 7597 |
| 13 | 63247691 | 63248191 | Aopep | Intron 2 | 7583 |
| 13 | 65311692 | 65312192 | Gm3800 | Promoter | -1566 |
| 13 | 70840622 | 70841122 | Adamts16 | Promoter | 690 |
| 13 | 70840622 | 70841122 | Adamts16 | Intron 2 | 690 |
| 13 | 70840622 | 70841122 | Adamts16 | Promoter | 654 |
| 13 | 70840622 | 70841122 | Adamts16 | Intron 2 | 654 |
| 13 | 70840622 | 70841122 | Adamts16 | Promoter | 690 |
| 13 | 70840622 | 70841122 | Adamts16 | Intron 2 | 690 |
| 13 | 70840622 | 70841122 | Adamts16 | Promoter | 687 |
| 13 | 70840622 | 70841122 | Gm36529 | Promoter | 171 |
| 13 | 72785552 | 72786052 | Gm47902 | Non-coding | 16121 |
| 13 | 73627552 | 73628052 | Tert | Promoter | 641 |
| 13 | 73627552 | 73628052 | Tert | Promoter | 636 |
| 13 | 73627552 | 73628052 | Tert | CDS 2 | 636 |
| 13 | 73627552 | 73628052 | Tert | Promoter | 562 |
| 13 | 73627552 | 73628052 | Tert | CDS 2 | 562 |
| 13 | 73627552 | 73628052 | Tert | Intron 2 | 562 |
| 13 | 73627552 | 73628052 | Tert | Promoter | 541 |
| 13 | 73627552 | 73628052 | Tert | CDS 2 | 541 |
| 13 | 73627552 | 73628052 | Tert | Intron 2 | 541 |
| 13 | 76210552 | 76211052 | Fam81b | Intron 10 | 94021 |
| 13 | 84062395 | 84062895 | Gm17750 | Promoter | 1878 |
| 13 | 85224395 | 85224895 | Rasa1 | Intron 18 | 64135 |
| 13 | 85224395 | 85224895 | Rasa1 | Intron 8 | 64236 |
| 13 | 85224395 | 85224895 | Rasa1 | Promoter | -2693 |
| 13 | 85224395 | 85224895 | Rasa1 | Promoter | 1801 |
| 13 | 85224395 | 85224895 | Rasa1 | Promoter | 1111 |
| 13 | 103695920 | 103696420 | Gm30551 | Non-coding | 11214 |
| 13 | 103695920 | 103696420 | Gm30551 | Non-coding | 11200 |
| 13 | 104798920 | 104799420 | Cwc27 | Intron 7 | 17534 |
| 13 | 104798920 | 104799420 | Srek1ip1 | Non-coding | 6436 |
| 13 | 108271806 | 108272306 | Elovl7 | CDS 5 | 57402 |
| 13 | 108271806 | 108272306 | Elovl7 | Intron 5 | 57402 |
| 13 | 108271806 | 108272306 | Elovl7 | 3' UTR | 57396 |
| 13 | 109657303 | 109657803 | Pde4d | Intron 3 | 1003126 |

| | | | | | |
|---|---|---|---|---|---|
| 13 | 109657303 | 109657803 | Pde4d | Intron 2 | 797137 |
| 13 | 109657303 | 109657803 | Pde4d | Non-coding | 540944 |
| 13 | 109657303 | 109657803 | Pde4d | Intron 1 | 540662 |
| 13 | 109657303 | 109657803 | Pde4d | Intron 1 | 396649 |
| 13 | 109657303 | 109657803 | Pde4d | Intron 1 | 215120 |
| 13 | 109657303 | 109657803 | Pde4d | Intron 1 | 75898 |
| 13 | 109657303 | 109657803 | Pde4d | Intron 1 | 24523 |
| 13 | 114885292 | 114885792 | Itga2 | Intron 3 | 46309 |
| 13 | 114914792 | 114915292 | Itga2 | Intron 1 | 16809 |
| 13 | 114885292 | 114885792 | Itga2 | Intron 3 | 16152 |
| 14 | 11994986 | 11995486 | Ptprg | Intron 4 | 441454 |
| 14 | 11994986 | 11995486 | Ptprg | Intron 4 | 441433 |
| 14 | 15384486 | 15384986 | Lrrc3b | 5' UTR | 51520 |
| 14 | 15384486 | 15384986 | Lrrc3b | 5' UTR | 54002 |
| 14 | 15384486 | 15384986 | Lrrc3b | 5' UTR | 53161 |
| 14 | 19981278 | 19981778 | Gng2 | Promoter | -4028 |
| 14 | 19981278 | 19981778 | Gng2 | Promoter | -4105 |
| 14 | 19981278 | 19981778 | Gng2 | Promoter | -3650 |
| 14 | 19981278 | 19981778 | Gng2 | Promoter | -4218 |
| 14 | 19981278 | 19981778 | Gng2 | Promoter | -4093 |
| 14 | 19981278 | 19981778 | Gng2 | Promoter | -4584 |
| 14 | 22735278 | 22735778 | Lrmda | Intron 5 | 715566 |
| 14 | 22735278 | 22735778 | Lrmda | Intron 3 | 715457 |
| 14 | 22735278 | 22735778 | Lrmda | 3' UTR | 715430 |
| 14 | 25480514 | 25481014 | Zmiz1 | Non-coding | 21329 |
| 14 | 25480514 | 25481014 | Zmiz1 | 5' UTR | 21290 |
| 14 | 25480514 | 25481014 | Zmiz1 | 5' UTR | 21247 |
| 14 | 25480514 | 25481014 | Zmiz1 | Non-coding | 21222 |
| 14 | 25480514 | 25481014 | Gm47921 | Promoter | 460 |
| 14 | 26357514 | 26358014 | 4933413J09Rik | Promoter | 940 |
| 14 | 26357514 | 26358014 | 4933413J09Rik | Promoter | 0 |
| 14 | 26357514 | 26358014 | 4933413J09Rik | Promoter | 0 |
| 14 | 26357514 | 26358014 | 4933413J09Rik | Promoter | -4617 |
| 14 | 32093814 | 32094314 | Oxnad1 | Intron 4 | 8438 |
| 14 | 32093814 | 32094314 | Oxnad1 | Intron 3 | 8401 |
| 14 | 32093814 | 32094314 | Oxnad1 | 3' UTR | 8029 |
| 14 | 32093814 | 32094314 | Oxnad1 | Intron 3 | 8027 |
| 14 | 32093814 | 32094314 | Oxnad1 | Intron 3 | 8003 |
| 14 | 32093814 | 32094314 | Oxnad1 | Intron 3 | 7988 |
| 14 | 32093814 | 32094314 | Oxnad1 | Intron 3 | 7980 |
| 14 | 32093814 | 32094314 | Oxnad1 | Non-coding | 7980 |
| 14 | 32093814 | 32094314 | Oxnad1 | Intron 2 | 7937 |
| 14 | 32093814 | 32094314 | Oxnad1 | Promoter | -1085 |
| 14 | 32093814 | 32094314 | Oxnad1 | Promoter | -1148 |
| 14 | 34822314 | 34822814 | Grid1 | Promoter | 2206 |
| 14 | 34822314 | 34822814 | Grid1 | Intron 1 | 2206 |

| | | | | | |
|---|---|---|---|---|---|
| 14 | 34822314 | 34822814 | Grid1 | CDS 2 | 2206 |
| 14 | 34822314 | 34822814 | Grid1 | Intron 2 | 2206 |
| 14 | 40918711 | 40919211 | Tspan14 | Intron 2 | 47597 |
| 14 | 40918711 | 40919211 | Tspan14 | Intron 1 | 15111 |
| 14 | 40918711 | 40919211 | Tspan14 | Non-coding | 15106 |
| 14 | 45982825 | 45983325 | Gm15218 | Non-coding | 228136 |
| 14 | 45982825 | 45983325 | Gm49192 | Non-coding | 7713 |
| 14 | 48138825 | 48139325 | Peli2 | Intron 1 | 18002 |
| 14 | 48138825 | 48139325 | Peli2 | Intron 1 | 17956 |
| 14 | 48138825 | 48139325 | Peli2 | Intron 1 | 17903 |
| 14 | 48138825 | 48139325 | Peli2 | Non-coding | 17888 |
| 14 | 48138825 | 48139325 | Peli2 | Non-coding | 16796 |
| 14 | 50892825 | 50893325 | Klhl33 | CDS 2 | 4183 |
| 14 | 50892825 | 50893325 | Klhl33 | Intron 1 | 4183 |
| 14 | 50892825 | 50893325 | Klhl33 | Promoter | 0 |
| 14 | 50892825 | 50893325 | Klhl33 | CDS 1 | 0 |
| 14 | 51071325 | 51071825 | Olfr750 | Promoter | 2611 |
| 14 | 51071325 | 51071825 | Olfr750 | 5' UTR | 2611 |
| 14 | 51071325 | 51071825 | Olfr750 | CDS 2 | 2611 |
| 14 | 51071325 | 51071825 | Olfr750 | Intron 1 | 2611 |
| 14 | 51071325 | 51071825 | Olfr750 | Promoter | 0 |
| 14 | 51071325 | 51071825 | Olfr750 | 5' UTR | 0 |
| 14 | 51071325 | 51071825 | Olfr750 | CDS 1 | 0 |
| 14 | 55618163 | 55618663 | Rec8 | Promoter | 126 |
| 14 | 55618163 | 55618663 | Rec8 | 5' UTR | 126 |
| 14 | 55618163 | 55618663 | Rec8 | CDS 1 | 126 |
| 14 | 55618163 | 55618663 | Rec8 | Intron 1 | 126 |
| 14 | 55618163 | 55618663 | Rec8 | CDS 2 | 126 |
| 14 | 55618163 | 55618663 | Rec8 | Intron 2 | 126 |
| 14 | 55617663 | 55618163 | Rec8 | Promoter | 0 |
| 14 | 55617663 | 55618163 | Rec8 | 5' UTR | 0 |
| 14 | 55618163 | 55618663 | Rec8 | Promoter | -4835 |
| 14 | 55618163 | 55618663 | Rec8 | Promoter | -4906 |
| 14 | 60152163 | 60152663 | Atp8a2 | Intron 1 | 44517 |
| 14 | 60152163 | 60152663 | Atp8a2 | Intron 1 | 24877 |
| 14 | 60152163 | 60152663 | Gm16261 | Promoter | -1488 |
| 14 | 60670663 | 60671163 | Spata13 | 5' UTR | 36662 |
| 14 | 60670663 | 60671163 | Spata13 | 5' UTR | 36420 |
| 14 | 60670663 | 60671163 | Spata13 | 5' UTR | 35934 |
| 14 | 60670663 | 60671163 | Spata13 | 5' UTR | 35929 |
| 14 | 60670663 | 60671163 | Spata13 | Non-coding | 35908 |
| 14 | 61143663 | 61144163 | Sacs | 5' UTR | 5206 |
| 14 | 61143663 | 61144163 | Sacs | 5' UTR | 5172 |
| 14 | 61143663 | 61144163 | Sacs | Promoter | -3558 |
| 14 | 61143663 | 61144163 | Sacs | Promoter | -3889 |
| 14 | 64011663 | 64012163 | Rp1l1 | 5' UTR | 19157 |

| | | | | | |
|---|---|---|---|---|---|
| 14 | 65023663 | 65024163 | Ints9 | Intron 10 | 73618 |
| 14 | 65023663 | 65024163 | Ints9 | Non-coding | 73350 |
| 14 | 65865163 | 65865663 | Ccdc25 | 3' UTR | 27861 |
| 14 | 65865163 | 65865663 | Ccdc25 | Non-coding | 27806 |
| 14 | 66263663 | 66264163 | Ptk2b | 5' UTR | 16890 |
| 14 | 66263663 | 66264163 | Ptk2b | 5' UTR | 16890 |
| 14 | 66263663 | 66264163 | Ptk2b | 5' UTR | 16820 |
| 14 | 66263663 | 66264163 | Ptk2b | Non-coding | 16830 |
| 14 | 68122443 | 68122943 | Nefm | Promoter | 1904 |
| 14 | 68122443 | 68122943 | Nefm | Intron 2 | 1904 |
| 14 | 68122443 | 68122943 | Nefm | CDS 2 | 1904 |
| 14 | 68122443 | 68122943 | Nefm | Promoter | 1904 |
| 14 | 68122443 | 68122943 | Nefm | Intron 2 | 1904 |
| 14 | 68122443 | 68122943 | Nefm | CDS 2 | 1904 |
| 14 | 70562693 | 70563193 | Hr | Intron 6 | 9006 |
| 14 | 70562693 | 70563193 | Hr | Intron 7 | 8637 |
| 14 | 70562693 | 70563193 | Hr | Intron 5 | 6793 |
| 14 | 75249693 | 75250193 | Cpb2 | Intron 1 | 7406 |
| 14 | 77706693 | 77707193 | Enox1 | Intron 17 | 549930 |
| 14 | 77176693 | 77177193 | Enox1 | 5' UTR | 19930 |
| 14 | 77176693 | 77177193 | Enox1 | Non-coding | 19913 |
| 14 | 77706693 | 77707193 | Enox1 | 3' UTR | 326114 |
| 14 | 77706693 | 77707193 | Enox1 | Intron 15 | 326047 |
| 14 | 78038693 | 78039193 | Gm48954 | Non-coding | 18965 |
| 14 | 78038193 | 78038693 | Gm48954 | Non-coding | 18465 |
| 14 | 78282693 | 78283193 | Tnfsf11 | Intron 4 | 24851 |
| 14 | 78526693 | 78527193 | Akap11 | 5' UTR | 9616 |
| 14 | 78526693 | 78527193 | Akap11 | Promoter | -4042 |
| 14 | 86215693 | 86216193 | Gm49061 | Promoter | 238 |
| 14 | 103586783 | 103587283 | Scel | Intron 20 | 73441 |
| 14 | 103586783 | 103587283 | Scel | Intron 21 | 73362 |
| 14 | 103681783 | 103682283 | Slain1 | Intron 2 | 31555 |
| 14 | 103646783 | 103647283 | Slain1 | Promoter | -2945 |
| 14 | 103681783 | 103682283 | Slain1 | Intron 2 | 31137 |
| 14 | 103646783 | 103647283 | Slain1 | Promoter | -3363 |
| 14 | 103681783 | 103682283 | Slain1 | Intron 2 | 26012 |
| 14 | 103681783 | 103682283 | Slain1 | Intron 1 | 17285 |
| 14 | 119106278 | 119106778 | Gm16835 | Non-coding | 7313 |
| 14 | 120368778 | 120369278 | Mbnl2 | Intron 2 | 93109 |
| 14 | 120368778 | 120369278 | Mbnl2 | Intron 2 | 93072 |
| 14 | 120368778 | 120369278 | Mbnl2 | Non-coding | 93071 |
| 14 | 120368778 | 120369278 | Mbnl2 | Intron 2 | 93057 |
| 14 | 120368778 | 120369278 | Mbnl2 | Intron 2 | 71104 |
| 14 | 120368778 | 120369278 | Mbnl2 | Intron 2 | 62326 |
| 14 | 120368778 | 120369278 | Mbnl2 | Non-coding | 62322 |
| 14 | 121796278 | 121796778 | Dock9 | Promoter | 1060 |

| | | | | | |
|---|---|---|---|---|---|
| 14 | 121796278 | 121796778 | Dock9 | Intron 1 | 1060 |
| 14 | 121961278 | 121961778 | Ubac2 | Intron 4 | 82658 |
| 14 | 121961278 | 121961778 | Ubac2 | Intron 3 | 82616 |
| 14 | 121961278 | 121961778 | Ubac2 | Non-coding | 45075 |
| 14 | 121961278 | 121961778 | Ubac2 | Non-coding | 14964 |
| 14 | 121961278 | 121961778 | Gpr183 | 5' UTR | 3418 |
| 14 | 121961278 | 121961778 | Gpr183 | 5' UTR | 3328 |
| 15 | 11626745 | 11627245 | Gm49106 | Non-coding | 3496 |
| 15 | 11626745 | 11627245 | Gm49106 | Non-coding | 3502 |
| 15 | 11626745 | 11627245 | Gm49107 | Promoter | -3484 |
| 15 | 27981745 | 27982245 | Trio | Intron 1 | 43604 |
| 15 | 27981745 | 27982245 | Trio | 5' UTR | 42710 |
| 15 | 27981745 | 27982245 | Trio | Non-coding | 13119 |
| 15 | 32296245 | 32296745 | Sema5a | Non-coding | 51435 |
| 15 | 32296245 | 32296745 | Sema5a | 5' UTR | 51432 |
| 15 | 32296245 | 32296745 | Sema5a | Non-coding | 51423 |
| 15 | 32296245 | 32296745 | Sema5a | Non-coding | 51420 |
| 15 | 32296245 | 32296745 | Sema5a | Non-coding | 51414 |
| 15 | 32296245 | 32296745 | Sema5a | Non-coding | 51207 |
| 15 | 32296245 | 32296745 | Sema5a | Non-coding | 51142 |
| 15 | 34904745 | 34905245 | Stk3 | Intron 10 | 250562 |
| 15 | 34904745 | 34905245 | Stk3 | Intron 10 | 273677 |
| 15 | 34904745 | 34905245 | Stk3 | Intron 8 | 250506 |
| 15 | 38273245 | 38273745 | Gm49314 | Promoter | 2697 |
| 15 | 39937454 | 39937954 | Lrp12 | Intron 1 | 5835 |
| 15 | 39937454 | 39937954 | Lrp12 | Intron 1 | 6041 |
| 15 | 39937454 | 39937954 | Lrp12 | Intron 1 | 5737 |
| 15 | 39937454 | 39937954 | Lrp12 | Non-coding | 5753 |
| 15 | 40065954 | 40066454 | Gm33251 | Non-coding | 20763 |
| 15 | 59057945 | 59058445 | Mtss1 | Intron 2 | 23545 |
| 15 | 59057945 | 59058445 | Mtss1 | CDS 2 | 23545 |
| 15 | 59057945 | 59058445 | Mtss1 | Intron 1 | 23545 |
| 15 | 59057945 | 59058445 | Mtss1 | Non-coding | 23561 |
| 15 | 59057945 | 59058445 | Mtss1 | Promoter | -2480 |
| 15 | 59592445 | 59592945 | Nsmce2 | Intron 7 | 218160 |
| 15 | 59592445 | 59592945 | Nsmce2 | Intron 6 | 218158 |
| 15 | 75996570 | 75997070 | Mapk15 | Promoter | 2801 |
| 15 | 75996570 | 75997070 | Mapk15 | Intron 7 | 2801 |
| 15 | 75996570 | 75997070 | Mapk15 | Promoter | 2801 |
| 15 | 75996570 | 75997070 | Mapk15 | Promoter | 631 |
| 15 | 75996570 | 75997070 | Mapk15 | Promoter | 0 |
| 15 | 75996570 | 75997070 | Mapk15 | Promoter | -1301 |
| 15 | 76669070 | 76669570 | Foxh1 | Promoter | 379 |
| 15 | 76669070 | 76669570 | Foxh1 | CDS 3 | 379 |
| 15 | 76669070 | 76669570 | Foxh1 | Intron 2 | 379 |
| 15 | 76669070 | 76669570 | Foxh1 | CDS 2 | 379 |

| | | | | | |
|---|---|---|---|---|---|
| 15 | 76669070 | 76669570 | Foxh1 | Intron 1 | 379 |
| 15 | 76669070 | 76669570 | Foxh1 | Promoter | 233 |
| 15 | 76669070 | 76669570 | Ppp1r16a | Promoter | -2045 |
| 15 | 76669070 | 76669570 | Ppp1r16a | Promoter | -2077 |
| 15 | 77765570 | 77766070 | Myh9 | Intron 35 | 76106 |
| 15 | 77765570 | 77766070 | Myh9 | CDS 35 | 76106 |
| 15 | 77765570 | 77766070 | Myh9 | Intron 34 | 76106 |
| 15 | 77765570 | 77766070 | Myh9 | Intron 3 | 3710 |
| 15 | 77765570 | 77766070 | Myh9 | Promoter | 1121 |
| 15 | 78387570 | 78388070 | Tex33 | Intron 2 | 7843 |
| 15 | 78387570 | 78388070 | Tex33 | Intron 3 | 7843 |
| 15 | 78387570 | 78388070 | Tex33 | Promoter | 700 |
| 15 | 78387570 | 78388070 | Tex33 | 5' UTR | 700 |
| 15 | 78387570 | 78388070 | Tex33 | Intron 2 | 7833 |
| 15 | 79098070 | 79098570 | Gm25072 | Promoter | -3012 |
| 15 | 79201570 | 79202070 | Gm10863 | Non-coding | 35529 |
| 15 | 79201570 | 79202070 | Gm10863 | Non-coding | 35512 |
| 15 | 79201570 | 79202070 | Gm10863 | Non-coding | 35504 |
| 15 | 79201570 | 79202070 | Gm10863 | Non-coding | 35504 |
| 15 | 79201570 | 79202070 | Gm10863 | Non-coding | 28850 |
| 15 | 79950070 | 79950570 | Cbx7 | 5' UTR | 20550 |
| 15 | 79950070 | 79950570 | Cbx7 | 5' UTR | 20549 |
| 15 | 80078070 | 80078570 | Rpl3 | CDS 9 | 4837 |
| 15 | 80078070 | 80078570 | Rpl3 | Intron 8 | 4837 |
| 15 | 80078070 | 80078570 | Rpl3 | CDS 8 | 4837 |
| 15 | 80078070 | 80078570 | Rpl3 | Intron 7 | 4837 |
| 15 | 80078070 | 80078570 | Rpl3 | Promoter | 1912 |
| 15 | 80078070 | 80078570 | Rpl3 | Intron 2 | 1912 |
| 15 | 80078070 | 80078570 | Rpl3 | Non-coding | 4014 |
| 15 | 80078070 | 80078570 | Rpl3 | Non-coding | 4784 |
| 15 | 80078070 | 80078570 | Snord83b | Promoter | 22 |
| 15 | 80078070 | 80078570 | Gm22077 | Promoter | 1774 |
| 15 | 81290570 | 81291070 | 8430426J06Rik | Non-coding | 21797 |
| 15 | 81948570 | 81949070 | Csdc2 | Intron 2 | 11811 |
| 15 | 81948570 | 81949070 | Csdc2 | CDS 3 | 11811 |
| 15 | 81948570 | 81949070 | Csdc2 | Intron 3 | 11811 |
| 15 | 81948570 | 81949070 | Csdc2 | Intron 2 | 11778 |
| 15 | 81948570 | 81949070 | Csdc2 | Promoter | 2839 |
| 15 | 81948570 | 81949070 | Csdc2 | Promoter | 1609 |
| 15 | 84086070 | 84086570 | Sult4a1 | Intron 5 | 19185 |
| 15 | 84086070 | 84086570 | Sult4a1 | Intron 5 | 19040 |
| 15 | 84809570 | 84810070 | Phf21b | Intron 2 | 45041 |
| 15 | 84809570 | 84810070 | Phf21b | Intron 2 | 45980 |
| 15 | 84809570 | 84810070 | Phf21b | Intron 2 | 45682 |
| 15 | 89376069 | 89376569 | Sco2 | Promoter | -2250 |
| 15 | 89376069 | 89376569 | Sco2 | Promoter | -2222 |

| | | | | | |
|---|---|---|---|---|---|
| 15 | 89376069 | 89376569 | Tymp | Promoter | 471 |
| 15 | 89376069 | 89376569 | Tymp | Intron 3 | 471 |
| 15 | 89376069 | 89376569 | Tymp | CDS 3 | 471 |
| 15 | 89376069 | 89376569 | Tymp | Intron 2 | 471 |
| 15 | 89376069 | 89376569 | Odf3b | Promoter | 2686 |
| 15 | 89376069 | 89376569 | Odf3b | Promoter | 1658 |
| 15 | 89376069 | 89376569 | Odf3b | Promoter | 2357 |
| 15 | 89376069 | 89376569 | Odf3b | Promoter | 2421 |
| 15 | 89376069 | 89376569 | Odf3b | Promoter | 2294 |
| 15 | 89376069 | 89376569 | Odf3b | Promoter | 2435 |
| 15 | 89376069 | 89376569 | Odf3b | Promoter | 2714 |
| 15 | 89424069 | 89424569 | Cpt1b | Promoter | 1295 |
| 15 | 89424069 | 89424569 | Cpt1b | Intron 4 | 1295 |
| 15 | 89424069 | 89424569 | Cpt1b | CDS 4 | 1295 |
| 15 | 89424069 | 89424569 | Cpt1b | Intron 3 | 1295 |
| 15 | 89424069 | 89424569 | Gm44502 | 3' UTR | 5128 |
| 15 | 89424069 | 89424569 | Cpt1b | Promoter | -2667 |
| 15 | 89424069 | 89424569 | Cpt1b | Promoter | 0 |
| 15 | 89424069 | 89424569 | Cpt1b | Promoter | 1227 |
| 15 | 89424069 | 89424569 | Chkb | Promoter | 2593 |
| 15 | 89424069 | 89424569 | Chkb | Promoter | 2591 |
| 15 | 92750069 | 92750569 | Pdzrn4 | Intron 7 | 353188 |
| 15 | 92696569 | 92697069 | Pdzrn4 | Intron 5 | 299688 |
| 15 | 92750069 | 92750569 | Pdzrn4 | Intron 5 | 152943 |
| 15 | 92696569 | 92697069 | Pdzrn4 | Intron 3 | 99443 |
| 15 | 93513069 | 93513569 | Prickle1 | 5' UTR | 82323 |
| 15 | 93513069 | 93513569 | Prickle1 | 5' UTR | 5955 |
| 15 | 94984069 | 94984569 | Tmem117 | Intron 5 | 354819 |
| 15 | 94984069 | 94984569 | Gm23129 | Promoter | 2570 |
| 15 | 97002569 | 97003069 | Slc38a4 | Intron 12 | 16883 |
| 15 | 97002569 | 97003069 | Slc38a4 | Intron 13 | 52888 |
| 15 | 97052569 | 97053069 | Slc38a4 | Promoter | 2888 |
| 15 | 97052569 | 97053069 | Slc38a4 | 5' UTR | 2888 |
| 15 | 97002569 | 97003069 | Slc38a4 | Intron 13 | 17254 |
| 15 | 97002569 | 97003069 | Slc38a4 | Intron 13 | 28143 |
| 15 | 97052569 | 97053069 | Slc38a4 | Promoter | -2481 |
| 15 | 98226069 | 98226569 | Olfr286 | 3' UTR | 7882 |
| 15 | 98226069 | 98226569 | Olfr286 | Promoter | 1075 |
| 15 | 98984569 | 98985069 | 4930578M01Rik | Promoter | 1221 |
| 15 | 98984569 | 98985069 | 4930578M01Rik | Promoter | 942 |
| 15 | 98984569 | 98985069 | 4930578M01Rik | Promoter | -825 |
| 15 | 99315069 | 99315569 | Prpf40b | Non-coding | 19830 |
| 15 | 99315069 | 99315569 | Prpf40b | Non-coding | 19828 |

| | | | | | |
|---|---|---|---|---|---|
| 15 | 99315069 | 99315569 | Prpf40b | Non-coding | 19828 |
| 15 | 99315069 | 99315569 | Prpf40b | Intron 20 | 19798 |
| 15 | 99315069 | 99315569 | Prpf40b | CDS 21 | 19798 |
| 15 | 99315069 | 99315569 | Prpf40b | Intron 21 | 19798 |
| 15 | 99315069 | 99315569 | Prpf40b | CDS 22 | 19798 |
| 15 | 99315069 | 99315569 | Prpf40b | Intron 19 | 19174 |
| 15 | 99315069 | 99315569 | Prpf40b | CDS 20 | 19174 |
| 15 | 99315069 | 99315569 | Prpf40b | Intron 20 | 19174 |
| 15 | 99315069 | 99315569 | Prpf40b | CDS 21 | 19174 |
| 15 | 99315069 | 99315569 | Prpf40b | Intron 19 | 11786 |
| 15 | 99315069 | 99315569 | Prpf40b | CDS 20 | 11786 |
| 15 | 99315069 | 99315569 | Prpf40b | Intron 20 | 11786 |
| 15 | 99315069 | 99315569 | Prpf40b | CDS 21 | 11786 |
| 15 | 99315069 | 99315569 | Prpf40b | Intron 19 | 11785 |
| 15 | 99315069 | 99315569 | Prpf40b | CDS 20 | 11785 |
| 15 | 99315069 | 99315569 | Prpf40b | Intron 20 | 11785 |
| 15 | 99315069 | 99315569 | Prpf40b | CDS 21 | 11785 |
| 15 | 99315069 | 99315569 | Prpf40b | Non-coding | 7184 |
| 15 | 99315069 | 99315569 | Prpf40b | Non-coding | 5166 |
| 15 | 99315069 | 99315569 | Prpf40b | Promoter | 201 |
| 15 | 99315069 | 99315569 | Prpf40b | Promoter | 192 |
| 15 | 99315069 | 99315569 | Prpf40b | Intron 1 | 192 |
| 15 | 99315069 | 99315569 | Prpf40b | CDS 2 | 192 |
| 15 | 99315069 | 99315569 | Prpf40b | Intron 2 | 192 |
| 15 | 99315069 | 99315569 | Prpf40b | CDS 3 | 192 |
| 15 | 99315069 | 99315569 | Prpf40b | Promoter | 163 |
| 15 | 99315069 | 99315569 | Prpf40b | Intron 1 | 163 |
| 15 | 99315069 | 99315569 | Prpf40b | CDS 2 | 163 |
| 15 | 99315069 | 99315569 | Prpf40b | Intron 2 | 163 |
| 15 | 99315069 | 99315569 | Prpf40b | CDS 3 | 163 |
| 15 | 99315069 | 99315569 | Prpf40b | Promoter | 0 |
| 15 | 99315069 | 99315569 | Prpf40b | 5' UTR | 0 |
| 15 | 99315069 | 99315569 | Prpf40b | CDS 1 | 0 |
| 15 | 99601569 | 99602069 | Aqp6 | Promoter | 169 |
| 15 | 99601569 | 99602069 | Aqp6 | CDS 1 | 169 |
| 15 | 99601569 | 99602069 | Aqp6 | Intron 1 | 169 |
| 15 | 99601569 | 99602069 | Aqp6 | Promoter | 169 |
| 15 | 99601569 | 99602069 | Aqp6 | CDS 1 | 169 |
| 15 | 99601569 | 99602069 | Aqp6 | Intron 1 | 169 |
| 15 | 100645069 | 100645569 | Bin2 | CDS 10 | 23985 |
| 15 | 100645069 | 100645569 | Bin2 | Intron 9 | 23985 |
| 15 | 100645069 | 100645569 | Bin2 | CDS 9 | 23959 |
| 15 | 100645069 | 100645569 | Bin2 | Intron 8 | 23959 |
| 15 | 100645069 | 100645569 | Bin2 | CDS 9 | 23937 |
| 15 | 100645069 | 100645569 | Bin2 | Intron 8 | 23937 |
| 15 | 100645069 | 100645569 | Bin2 | CDS 10 | 23937 |

| | | | | | |
|---|---|---|---|---|---|
| 15 | 100645069 | 100645569 | Bin2 | Intron 9 | 23937 |
| 15 | 100645069 | 100645569 | Bin2 | 3' UTR | 23937 |
| 15 | 100645069 | 100645569 | Bin2 | CDS 4 | 23937 |
| 15 | 100645069 | 100645569 | Bin2 | Intron 3 | 23937 |
| 15 | 100645069 | 100645569 | Bin2 | 3' UTR | 23957 |
| 15 | 100645069 | 100645569 | Bin2 | CDS 7 | 23957 |
| 15 | 100645069 | 100645569 | Bin2 | Intron 6 | 23957 |
| 15 | 100645069 | 100645569 | Bin2 | Non-coding | 6092 |
| 15 | 100685069 | 100685569 | Cela1 | Promoter | 2353 |
| 15 | 100685069 | 100685569 | Cela1 | Intron 3 | 2353 |
| 15 | 100685069 | 100685569 | Cela1 | CDS 3 | 2353 |
| 15 | 100685069 | 100685569 | Cela1 | Intron 2 | 2353 |
| 15 | 100685069 | 100685569 | Cela1 | Promoter | -3106 |
| 15 | 100685069 | 100685569 | Cela1 | Promoter | -3485 |
| 15 | 100685069 | 100685569 | Cela1 | Promoter | -1949 |
| 15 | 100685069 | 100685569 | Cela1 | Promoter | 0 |
| 15 | 100685069 | 100685569 | Cela1 | Intron 1 | 0 |
| 15 | 100685069 | 100685569 | Cela1 | CDS 1 | 0 |
| 15 | 100685069 | 100685569 | Cela1 | Promoter | 0 |
| 15 | 100685069 | 100685569 | Cela1 | Promoter | 2347 |
| 15 | 101030069 | 101030569 | Scn8a | Intron 22 | 159386 |
| 15 | 101030069 | 101030569 | Scn8a | Intron 22 | 159165 |
| 15 | 101030069 | 101030569 | Scn8a | Intron 22 | 159124 |
| 15 | 101030069 | 101030569 | Scn8a | Intron 22 | 159117 |
| 15 | 101030069 | 101030569 | Scn8a | Intron 21 | 93798 |
| 15 | 101030069 | 101030569 | Scn8a | Intron 20 | 93744 |
| 15 | 101254569 | 101255069 | Nr4a1 | Promoter | 300 |
| 15 | 101254569 | 101255069 | Nr4a1 | 5' UTR | 300 |
| 15 | 101548569 | 101549069 | Krt82 | Promoter | 1599 |
| 15 | 101548569 | 101549069 | Krt82 | Intron 1 | 1599 |
| 15 | 102985069 | 102985569 | Hoxc5 | Non-coding | 17626 |
| 15 | 102985069 | 102985569 | Hoxc9 | 3' UTR | 8037 |
| 16 | 6876907 | 6877407 | Rbfox1 | Intron 4 | 991552 |
| 16 | 6876907 | 6877407 | Rbfox1 | Intron 3 | 527769 |
| 16 | 6876907 | 6877407 | Rbfox1 | Intron 1 | 67685 |
| 16 | 11648407 | 11648907 | Snx29 | Intron 16 | 325499 |
| 16 | 11648407 | 11648907 | Snx29 | Intron 9 | 242759 |
| 16 | 11648407 | 11648907 | Snx29 | Intron 9 | 241750 |
| 16 | 11648407 | 11648907 | Snx29 | Intron 6 | 227699 |
| 16 | 11648407 | 11648907 | Snx29 | Non-coding | 201099 |
| 16 | 15861907 | 15862407 | Mzt2 | Promoter | 941 |
| 16 | 15861907 | 15862407 | Mzt2 | Intron 2 | 941 |
| 16 | 15861907 | 15862407 | Mzt2 | CDS 2 | 941 |
| 16 | 15861907 | 15862407 | Mzt2 | Promoter | 316 |
| 16 | 15861907 | 15862407 | Mzt2 | Promoter | 904 |
| 16 | 15861907 | 15862407 | Mzt2 | Intron 2 | 904 |

| | | | | | |
|---|---|---|---|---|---|
| 16 | 15861907 | 15862407 | Mzt2 | CDS 2 | 904 |
| 16 | 15861907 | 15862407 | Mzt2 | Promoter | 963 |
| 16 | 15861907 | 15862407 | Mzt2 | Intron 3 | 963 |
| 16 | 15861907 | 15862407 | Mzt2 | CDS 3 | 963 |
| 16 | 18505907 | 18506407 | Gnb1l | 5' UTR | 7194 |
| 16 | 18505907 | 18506407 | Gnb1l | 5' UTR | 7123 |
| 16 | 18505907 | 18506407 | Gnb1l | 5' UTR | 7084 |
| 16 | 18505907 | 18506407 | Gnb1l | Non-coding | 7029 |
| 16 | 18505907 | 18506407 | Gnb1l | 5' UTR | 7021 |
| 16 | 18505907 | 18506407 | Gnb1l | 5' UTR | 7021 |
| 16 | 18505907 | 18506407 | Gnb1l | 5' UTR | 7019 |
| 16 | 18505907 | 18506407 | Gnb1l | 5' UTR | 7019 |
| 16 | 18505907 | 18506407 | Gnb1l | Non-coding | 7016 |
| 16 | 18505907 | 18506407 | Gnb1l | 5' UTR | 7010 |
| 16 | 18505907 | 18506407 | Gnb1l | 5' UTR | 7005 |
| 16 | 18505907 | 18506407 | Gnb1l | Non-coding | 6967 |
| 16 | 21330427 | 21330927 | Magef1 | Promoter | 2430 |
| 16 | 21330427 | 21330927 | Magef1 | Promoter | 2203 |
| 16 | 22621427 | 22621927 | Dgkg | Intron 2 | 35247 |
| 16 | 22621427 | 22621927 | Dgkg | Intron 2 | 35295 |
| 16 | 22621427 | 22621927 | Dgkg | Intron 2 | 35262 |
| 16 | 22621427 | 22621927 | Dgkg | Non-coding | 35102 |
| 16 | 28750414 | 28750914 | Fgf12 | Promoter | 2155 |
| 16 | 28750414 | 28750914 | Fgf12 | Intron 1 | 2155 |
| 16 | 30873914 | 30874414 | Gm49756 | Promoter | -3822 |
| 16 | 30873914 | 30874414 | Gm49756 | Promoter | -3189 |
| 16 | 30873914 | 30874414 | Gm49757 | Non-coding | 3723 |
| 16 | 33529414 | 33529914 | Slc12a8 | Intron 2 | 12086 |
| 16 | 33529414 | 33529914 | Slc12a8 | Intron 1 | 11136 |
| 16 | 33529414 | 33529914 | Slc12a8 | Intron 1 | 11085 |
| 16 | 33529414 | 33529914 | Slc12a8 | 5' UTR | 10878 |
| 16 | 34574414 | 34574914 | Kalrn | Promoter | -881 |
| 16 | 34658414 | 34658914 | Ropn1 | 5' UTR | 8493 |
| 16 | 34658414 | 34658914 | Ropn1 | 5' UTR | 7226 |
| 16 | 34806914 | 34807414 | Mylk | Non-coding | 61704 |
| 16 | 34976914 | 34977414 | Mylk | CDS 26 | 191993 |
| 16 | 34976914 | 34977414 | Mylk | Intron 26 | 191993 |
| 16 | 34806914 | 34807414 | Mylk | Intron 1 | 21993 |
| 16 | 34976914 | 34977414 | Mylk | Non-coding | 76659 |
| 16 | 35183414 | 35183914 | Adcy5 | Intron 1 | 28537 |
| 16 | 35178914 | 35179414 | Adcy5 | Intron 1 | 24037 |
| 16 | 43385887 | 43386387 | Zbtb20 | Non-coding | 510006 |
| 16 | 43385887 | 43386387 | Zbtb20 | Non-coding | 478345 |
| 16 | 43385887 | 43386387 | Zbtb20 | 5' UTR | 478242 |
| 16 | 43385887 | 43386387 | Zbtb20 | 5' UTR | 478235 |
| 16 | 43385887 | 43386387 | Zbtb20 | Non-coding | 478235 |

| | | | | | |
|---|---|---|---|---|---|
| 16 | 43385887 | 43386387 | Zbtb20 | 5' UTR | 150007 |
| 16 | 43385887 | 43386387 | Zbtb20 | 5' UTR | 138603 |
| 16 | 43385887 | 43386387 | Zbtb20 | 5' UTR | 138603 |
| 16 | 43385887 | 43386387 | Zbtb20 | Non-coding | 22839 |
| 16 | 43385887 | 43386387 | Zbtb20 | Non-coding | 22423 |
| 16 | 43385887 | 43386387 | Zbtb20 | 5' UTR | 22014 |
| 16 | 43385887 | 43386387 | Zbtb20 | 5' UTR | 21683 |
| 16 | 44765387 | 44765887 | Cd200r1 | Promoter | 0 |
| 16 | 44765387 | 44765887 | Cd200r1 | 5' UTR | 0 |
| 16 | 44765387 | 44765887 | Cd200r1 | Promoter | 0 |
| 16 | 44765387 | 44765887 | Cd200r1 | 5' UTR | 0 |
| 16 | 44765387 | 44765887 | Cd200r1 | Promoter | 0 |
| 16 | 44765387 | 44765887 | Cd200r1 | 5' UTR | 0 |
| 16 | 45579387 | 45579887 | Slc9c1 | Intron 16 | 44078 |
| 16 | 45579387 | 45579887 | Slc9c1 | 3' UTR | 39716 |
| 16 | 45579387 | 45579887 | Slc9c1 | 3' UTR | 39716 |
| 16 | 57177887 | 57178387 | Tbc1d23 | Intron 15 | 53118 |
| 16 | 57177887 | 57178387 | Tbc1d23 | CDS 15 | 53118 |
| 16 | 57177887 | 57178387 | Tbc1d23 | Intron 14 | 53118 |
| 16 | 57177887 | 57178387 | Tbc1d23 | Intron 16 | 52980 |
| 16 | 57177887 | 57178387 | Tbc1d23 | CDS 16 | 52980 |
| 16 | 57177887 | 57178387 | Tbc1d23 | Intron 15 | 52980 |
| 16 | 57177887 | 57178387 | Tbc1d23 | Promoter | 0 |
| 16 | 57177887 | 57178387 | Tbc1d23 | Non-coding | 4476 |
| 16 | 57177887 | 57178387 | Tbc1d23 | Non-coding | 4166 |
| 16 | 59284174 | 59284674 | Olfr202 | Promoter | 2035 |
| 16 | 59284174 | 59284674 | Olfr202 | 5' UTR | 2035 |
| 16 | 59284174 | 59284674 | Olfr202 | CDS 3 | 2035 |
| 16 | 59284174 | 59284674 | Olfr202 | Intron 2 | 2035 |
| 16 | 59284174 | 59284674 | Olfr202 | Promoter | 2120 |
| 16 | 59284174 | 59284674 | Olfr202 | 5' UTR | 2120 |
| 16 | 59284174 | 59284674 | Olfr202 | CDS 3 | 2120 |
| 16 | 59284174 | 59284674 | Olfr202 | Intron 2 | 2120 |
| 16 | 59284174 | 59284674 | Olfr202 | Promoter | 0 |
| 16 | 59284174 | 59284674 | Olfr202 | CDS 1 | 0 |
| 16 | 70320255 | 70320755 | Gbe1 | Intron 1 | 6306 |
| 16 | 70320255 | 70320755 | Gbe1 | Non-coding | 6192 |
| 16 | 70320255 | 70320755 | Gbe1 | Non-coding | 6191 |
| 16 | 70320255 | 70320755 | Gbe1 | Intron 1 | 6191 |
| 16 | 70320255 | 70320755 | Gbe1 | Intron 1 | 6168 |
| 16 | 70320255 | 70320755 | Gbe1 | Intron 1 | 6121 |
| 16 | 76401255 | 76401755 | Gm9843 | Promoter | 1898 |
| 16 | 76454755 | 76455255 | Gm45030 | Non-coding | 47655 |
| 16 | 78311755 | 78312255 | Cxadr | Intron 2 | 10266 |
| 16 | 78311755 | 78312255 | Cxadr | Intron 1 | 10259 |
| 16 | 78311755 | 78312255 | Cxadr | Intron 1 | 9999 |

| | | | | | |
|---|---|---|---|---|---|
| 16 | 78311755 | 78312255 | Cxadr | Intron 1 | 9996 |
| 16 | 78954255 | 78954755 | Tmprss15 | Intron 27 | 136338 |
| 16 | 78954255 | 78954755 | Tmprss15 | Intron 26 | 136343 |
| 16 | 87493755 | 87494255 | Cct8 | Promoter | 1619 |
| 16 | 87493755 | 87494255 | Cct8 | CDS 3 | 1619 |
| 16 | 87493755 | 87494255 | Cct8 | Intron 2 | 1619 |
| 16 | 87493755 | 87494255 | Cct8 | Promoter | 1534 |
| 16 | 87493755 | 87494255 | Cct8 | CDS 3 | 1534 |
| 16 | 87493755 | 87494255 | Cct8 | Intron 2 | 1534 |
| 16 | 87493755 | 87494255 | Cct8 | Promoter | 1534 |
| 16 | 87493755 | 87494255 | Cct8 | Intron 2 | 1534 |
| 16 | 87493755 | 87494255 | Cct8 | Promoter | -4911 |
| 16 | 87493755 | 87494255 | Cct8 | Promoter | -2306 |
| 16 | 87493755 | 87494255 | Cct8 | Promoter | 1605 |
| 16 | 87493755 | 87494255 | Cct8 | 5' UTR | 1605 |
| 16 | 87493755 | 87494255 | Cct8 | CDS 2 | 1605 |
| 16 | 87493755 | 87494255 | Cct8 | Intron 1 | 1605 |
| 16 | 87493755 | 87494255 | Cct8 | Promoter | 1450 |
| 16 | 87493755 | 87494255 | Cct8 | 5' UTR | 1450 |
| 16 | 87493755 | 87494255 | Cct8 | CDS 3 | 1450 |
| 16 | 87493755 | 87494255 | Cct8 | Intron 2 | 1450 |
| 16 | 87493755 | 87494255 | B130034C11Rik | Promoter | -1744 |
| 16 | 87493755 | 87494255 | B130034C11Rik | Promoter | -1818 |
| 16 | 87906255 | 87906755 | Grik1 | Intron 16 | 383110 |
| 16 | 87906255 | 87906755 | Grik1 | Intron 15 | 383489 |
| 16 | 87906255 | 87906755 | Grik1 | Intron 15 | 383504 |
| 16 | 87906255 | 87906755 | Grik1 | Intron 16 | 382977 |
| 16 | 87906255 | 87906755 | Grik1 | Intron 16 | 382977 |
| 16 | 90359755 | 90360255 | Gm49709 | Non-coding | 10338 |
| 16 | 90359755 | 90360255 | Gm49710 | Promoter | 2863 |
| 16 | 90876255 | 90876755 | Eva1c | Non-coding | 49536 |
| 16 | 90872755 | 90873255 | Eva1c | Non-coding | 46036 |
| 16 | 90876255 | 90876755 | Eva1c | Non-coding | 49536 |
| 16 | 90872755 | 90873255 | Eva1c | Non-coding | 46036 |
| 16 | 90876255 | 90876755 | Eva1c | Intron 4 | 49509 |
| 16 | 90872755 | 90873255 | Eva1c | Intron 3 | 46009 |
| 16 | 90876255 | 90876755 | Eva1c | 3' UTR | 45447 |
| 16 | 90872755 | 90873255 | Eva1c | 3' UTR | 41947 |
| 16 | 90876255 | 90876755 | Eva1c | Intron 4 | 45396 |
| 16 | 90872755 | 90873255 | Eva1c | Intron 3 | 41896 |
| 16 | 90876255 | 90876755 | Eva1c | 3' UTR | 45388 |
| 16 | 90872755 | 90873255 | Eva1c | 3' UTR | 41888 |
| 16 | 90876255 | 90876755 | Eva1c | Intron 2 | 45267 |
| 16 | 90872755 | 90873255 | Eva1c | Intron 1 | 41767 |

| | | | | | |
|---|---|---|---|---|---|
| 16 | 90876255 | 90876755 | Eva1c | Intron 4 | 45142 |
| 16 | 90872755 | 90873255 | Eva1c | Intron 3 | 41642 |
| 16 | 90876255 | 90876755 | Eva1c | Intron 4 | 29857 |
| 16 | 90872755 | 90873255 | Eva1c | Intron 3 | 26357 |
| 16 | 90876255 | 90876755 | Eva1c | Intron 3 | 9875 |
| 16 | 90872755 | 90873255 | Eva1c | Intron 2 | 6375 |
| 16 | 91320755 | 91321255 | Gm15966 | Non-coding | 7743 |
| 16 | 91301255 | 91301755 | Gm15966 | Non-coding | 27243 |
| 16 | 91320755 | 91321255 | Gm15966 | Non-coding | 7743 |
| 16 | 91301255 | 91301755 | Gm15966 | Non-coding | 27243 |
| 16 | 91320755 | 91321255 | Gm15966 | Non-coding | 7789 |
| 16 | 91320755 | 91321255 | Gm15966 | Non-coding | 7751 |
| 16 | 91320755 | 91321255 | Gm15966 | Non-coding | 7789 |
| 16 | 91320755 | 91321255 | Gm15966 | Non-coding | 7751 |
| 16 | 91320755 | 91321255 | Gm15966 | Non-coding | 7732 |
| 16 | 91320755 | 91321255 | Gm15966 | Non-coding | 7741 |
| 16 | 91320755 | 91321255 | Gm15966 | Non-coding | 4486 |
| 16 | 92762255 | 92762755 | Runx1 | Intron 2 | 63395 |
| 16 | 92762255 | 92762755 | Runx1 | Intron 1 | 63024 |
| 16 | 93486755 | 93487255 | Gm30695 | Promoter | 0 |
| 16 | 94005755 | 94006255 | Cldn14 | Promoter | 2583 |
| 16 | 94005755 | 94006255 | Cldn14 | 5' UTR | 2583 |
| 16 | 94721393 | 94721893 | Gm41505 | Promoter | -1 |
| 16 | 94996393 | 94996893 | Kcnj6 | Promoter | 763 |
| 16 | 94996393 | 94996893 | Kcnj6 | 5' UTR | 763 |
| 16 | 94996393 | 94996893 | Kcnj6 | Promoter | 804 |
| 16 | 94996393 | 94996893 | Kcnj6 | 5' UTR | 804 |
| 16 | 94996393 | 94996893 | Kcnj6 | Promoter | 809 |
| 16 | 94996393 | 94996893 | Kcnj6 | Promoter | 691 |
| 16 | 94996393 | 94996893 | Kcnj6 | Promoter | 360 |
| 16 | 96508893 | 96509393 | Gm49948 | Intron 8 | 147099 |
| 16 | 96508893 | 96509393 | Pcp4 | Intron 2 | 41287 |
| 16 | 96508893 | 96509393 | Pcp4 | 3' UTR | 15077 |
| 16 | 96508893 | 96509393 | Pcp4 | Intron 2 | 15077 |
| 16 | 96508893 | 96509393 | Pcp4 | Non-coding | 9273 |
| 16 | 96903393 | 96903893 | Dscam | Intron 3 | 266860 |
| 17 | 10841134 | 10841634 | Pacrg | Promoter | -822 |
| 17 | 10841134 | 10841634 | Prkn | Promoter | 750 |
| 17 | 10841134 | 10841634 | Prkn | Promoter | 750 |
| 17 | 10841134 | 10841634 | Prkn | Promoter | 750 |
| 17 | 10841134 | 10841634 | Prkn | Intron 1 | 750 |
| 17 | 10841134 | 10841634 | Prkn | Promoter | 747 |
| 17 | 10841134 | 10841634 | Prkn | Intron 1 | 747 |
| 17 | 10841134 | 10841634 | Prkn | Promoter | 743 |
| 17 | 10841134 | 10841634 | Prkn | Promoter | 721 |
| 17 | 10841134 | 10841634 | Prkn | Intron 1 | 721 |

| | | | | | |
|---|---|---|---|---|---|
| 17 | 10841134 | 10841634 | Prkn | Promoter | 721 |
| 17 | 10841134 | 10841634 | Prkn | Intron 1 | 721 |
| 17 | 10841134 | 10841634 | Prkn | Promoter | 721 |
| 17 | 10841134 | 10841634 | Prkn | Intron 1 | 721 |
| 17 | 10841134 | 10841634 | Prkn | Promoter | 721 |
| 17 | 10841134 | 10841634 | Prkn | Intron 1 | 721 |
| 17 | 10841134 | 10841634 | Prkn | Promoter | 664 |
| 17 | 10841134 | 10841634 | Prkn | 5' UTR | 664 |
| 17 | 12413134 | 12413634 | Plg | Intron 16 | 34526 |
| 17 | 12734134 | 12734634 | Igf2r | Intron 5 | 35031 |
| 17 | 15413536 | 15414036 | Fam120b | Intron 4 | 17334 |
| 17 | 15413536 | 15414036 | Fam120b | Non-coding | 17288 |
| 17 | 24164533 | 24165033 | Amdhd2 | Promoter | -766 |
| 17 | 24164533 | 24165033 | Amdhd2 | Promoter | -868 |
| 17 | 24164533 | 24165033 | Amdhd2 | Promoter | -802 |
| 17 | 24164533 | 24165033 | Amdhd2 | Promoter | -868 |
| 17 | 24164533 | 24165033 | Amdhd2 | Promoter | -839 |
| 17 | 24164533 | 24165033 | Atp6v0c | CDS 3 | 4429 |
| 17 | 24164533 | 24165033 | Atp6v0c | Intron 2 | 4429 |
| 17 | 24164533 | 24165033 | Atp6v0c | CDS 2 | 4429 |
| 17 | 24164533 | 24165033 | Atp6v0c | Intron 1 | 4429 |
| 17 | 24164533 | 24165033 | Atp6v0c | Promoter | 1997 |
| 17 | 24164533 | 24165033 | Atp6v0c | CDS 4 | 4645 |
| 17 | 24164533 | 24165033 | Atp6v0c | Intron 3 | 4645 |
| 17 | 24164533 | 24165033 | Atp6v0c | CDS 3 | 4645 |
| 17 | 24164533 | 24165033 | Atp6v0c | Intron 2 | 4645 |
| 17 | 24164533 | 24165033 | Atp6v0c | CDS 4 | 4616 |
| 17 | 24164533 | 24165033 | Atp6v0c | Intron 3 | 4616 |
| 17 | 24164533 | 24165033 | Atp6v0c | CDS 3 | 4616 |
| 17 | 24164533 | 24165033 | Atp6v0c | Intron 2 | 4616 |
| 17 | 24164533 | 24165033 | Atp6v0c | Non-coding | 4034 |
| 17 | 24164533 | 24165033 | Atp6v0c | Non-coding | 4670 |
| 17 | 25077555 | 25078055 | Ift140 | Intron 18 | 61470 |
| 17 | 25077555 | 25078055 | Ift140 | Intron 18 | 61464 |
| 17 | 25077555 | 25078055 | Ift140 | Intron 18 | 61164 |
| 17 | 25077555 | 25078055 | Ift140 | Non-coding | 32038 |
| 17 | 25077555 | 25078055 | Ift140 | Non-coding | 31989 |
| 17 | 25077555 | 25078055 | Tmem204 | Intron 1 | 3127 |
| 17 | 25077555 | 25078055 | Tmem204 | Non-coding | 3136 |
| 17 | 25077555 | 25078055 | Tmem204 | Non-coding | 5537 |
| 17 | 25077555 | 25078055 | Tmem204 | Promoter | 170 |
| 17 | 25077555 | 25078055 | Tmem204 | 5' UTR | 170 |
| 17 | 25277555 | 25278055 | Ube2i | Promoter | -3244 |
| 17 | 25277555 | 25278055 | Ube2i | Promoter | -4171 |
| 17 | 25277555 | 25278055 | Ube2i | Promoter | -3244 |
| 17 | 25277555 | 25278055 | Ube2i | Promoter | -3640 |

| | | | | | |
|---|---|---|---|---|---|
| 17 | 25277555 | 25278055 | Ube2i | Promoter | -4020 |
| 17 | 25277555 | 25278055 | Ube2i | Promoter | -2932 |
| 17 | 25277555 | 25278055 | Ube2i | Promoter | -3964 |
| 17 | 25277555 | 25278055 | Ube2i | Promoter | -3687 |
| 17 | 25277555 | 25278055 | Ube2i | Promoter | -3252 |
| 17 | 25277555 | 25278055 | Ube2i | Promoter | -3954 |
| 17 | 25277555 | 25278055 | Ube2i | Promoter | -4399 |
| 17 | 25277555 | 25278055 | Ube2i | Promoter | -4295 |
| 17 | 26112555 | 26113055 | Tmem8 | Promoter | -244 |
| 17 | 26112555 | 26113055 | Tmem8 | Promoter | -261 |
| 17 | 26112555 | 26113055 | Tmem8 | Promoter | -288 |
| 17 | 26112555 | 26113055 | Tmem8 | Promoter | -3576 |
| 17 | 26112555 | 26113055 | Tmem8 | Promoter | -4774 |
| 17 | 27133055 | 27133555 | Uqcc2 | Promoter | 362 |
| 17 | 27133055 | 27133555 | Uqcc2 | Intron 1 | 362 |
| 17 | 27133055 | 27133555 | Uqcc2 | Promoter | 75 |
| 17 | 27133055 | 27133555 | Uqcc2 | 5' UTR | 75 |
| 17 | 27133055 | 27133555 | Uqcc2 | Intron 1 | 75 |
| 17 | 27133055 | 27133555 | Uqcc2 | CDS 1 | 75 |
| 17 | 27133055 | 27133555 | Uqcc2 | Promoter | 69 |
| 17 | 27133055 | 27133555 | Uqcc2 | Promoter | 348 |
| 17 | 27133055 | 27133555 | Uqcc2 | Intron 1 | 348 |
| 17 | 28058555 | 28059055 | Anks1 | Intron 23 | 149249 |
| 17 | 28058555 | 28059055 | Anks1 | Intron 22 | 149238 |
| 17 | 28058555 | 28059055 | Anks1 | CDS 23 | 149238 |
| 17 | 28058555 | 28059055 | Anks1 | Intron 23 | 149238 |
| 17 | 28058555 | 28059055 | Anks1 | Intron 22 | 149222 |
| 17 | 28058555 | 28059055 | Anks1 | Intron 23 | 149183 |
| 17 | 28058555 | 28059055 | Anks1 | CDS 24 | 149183 |
| 17 | 28058555 | 28059055 | Anks1 | Intron 24 | 149183 |
| 17 | 28058555 | 28059055 | Anks1 | Intron 12 | 51210 |
| 17 | 28058555 | 28059055 | Anks1 | Intron 10 | 15940 |
| 17 | 28058555 | 28059055 | Anks1 | Intron 4 | 4222 |
| 17 | 28058555 | 28059055 | Anks1 | CDS 5 | 4222 |
| 17 | 28058555 | 28059055 | Anks1 | Intron 5 | 4222 |
| 17 | 28058555 | 28059055 | Anks1 | Promoter | 1114 |
| 17 | 28523555 | 28524055 | Gm20109 | Promoter | -406 |
| 17 | 28523555 | 28524055 | E230001N04Rik | Promoter | 835 |
| 17 | 28523555 | 28524055 | E230001N04Rik | Promoter | 262 |
| 17 | 28523555 | 28524055 | Gm25195 | Promoter | -2277 |
| 17 | 28538555 | 28539055 | Armc12 | 3' UTR | 7749 |
| 17 | 28538555 | 28539055 | Armc12 | Intron 5 | 7749 |
| 17 | 28538555 | 28539055 | Armc12 | CDS 6 | 7749 |
| 17 | 28538555 | 28539055 | Armc12 | 3' UTR | 7698 |

| | | | | | |
|---|---|---|---|---|---|
| 17 | 28538555 | 28539055 | Armc12 | Intron 5 | 7698 |
| 17 | 28538555 | 28539055 | Armc12 | CDS 6 | 7698 |
| 17 | 28538555 | 28539055 | Armc12 | 3' UTR | 7698 |
| 17 | 28538555 | 28539055 | Armc12 | Intron 5 | 7698 |
| 17 | 28538555 | 28539055 | Armc12 | CDS 6 | 7698 |
| 17 | 28538555 | 28539055 | Armc12 | Non-coding | 7694 |
| 17 | 28538555 | 28539055 | Armc12 | 3' UTR | 7688 |
| 17 | 29627555 | 29628055 | Gm28052 | Non-coding | 77756 |
| 17 | 29627555 | 29628055 | Rnf8 | Intron 3 | 12765 |
| 17 | 29627555 | 29628055 | Rnf8 | 3' UTR | 12701 |
| 17 | 29627555 | 29628055 | Gm28043 | Intron 3 | 12679 |
| 17 | 29627555 | 29628055 | Rnf8 | 3' UTR | 12587 |
| 17 | 29627555 | 29628055 | Rnf8 | Promoter | -581 |
| 17 | 29627555 | 29628055 | Rnf8 | Promoter | -3477 |
| 17 | 31125055 | 31125555 | Tff3 | 3' UTR | 4092 |
| 17 | 31125055 | 31125555 | Tff3 | CDS 3 | 4092 |
| 17 | 31125055 | 31125555 | Tff3 | Intron 2 | 4092 |
| 17 | 31277555 | 31278055 | Rsph1 | Promoter | -171 |
| 17 | 31277555 | 31278055 | Rsph1 | Promoter | -593 |
| 17 | 31277555 | 31278055 | Rsph1 | Promoter | -216 |
| 17 | 32334055 | 32334555 | Akap8l | Intron 9 | 16053 |
| 17 | 32334055 | 32334555 | Akap8l | Non-coding | 16021 |
| 17 | 32334055 | 32334555 | Akap8l | Non-coding | 16023 |
| 17 | 32334055 | 32334555 | Akap8l | Intron 10 | 16041 |
| 17 | 32334055 | 32334555 | Akap8l | Promoter | -1155 |
| 17 | 32334055 | 32334555 | Akap8l | Promoter | -573 |
| 17 | 32334055 | 32334555 | Akap8l | Promoter | 1025 |
| 17 | 32334055 | 32334555 | Akap8l | Intron 2 | 1025 |
| 17 | 32334055 | 32334555 | Akap8l | Promoter | 0 |
| 17 | 32334055 | 32334555 | Akap8l | Promoter | -580 |
| 17 | 32334055 | 32334555 | Akap8l | Promoter | 1856 |
| 17 | 32334055 | 32334555 | Akap8l | Intron 5 | 1856 |
| 17 | 32334055 | 32334555 | Akap8l | Promoter | 1999 |
| 17 | 32334055 | 32334555 | Akap8l | 3' UTR | 1999 |
| 17 | 34130055 | 34130555 | H2-DMa | Non-coding | 10514 |
| 17 | 34130055 | 34130555 | H2-DMa | Non-coding | 10467 |
| 17 | 34130055 | 34130555 | H2-DMa | Non-coding | 7233 |
| 17 | 34130055 | 34130555 | H2-DMa | 5' UTR | 7223 |
| 17 | 34130055 | 34130555 | H2-DMa | Non-coding | 7209 |
| 17 | 34130055 | 34130555 | Gm50335 | Promoter | -644 |
| 17 | 35554055 | 35554555 | Cdsn | Promoter | 1927 |
| 17 | 35554055 | 35554555 | Cdsn | Intron 1 | 1927 |
| 17 | 35774055 | 35774555 | 4833427F10Rik | Promoter | 1605 |
| 17 | 41079051 | 41079551 | Gm49857 | Non-coding | 7928 |
| 17 | 44560051 | 44560551 | Runx2 | 3' UTR | 175626 |
| 17 | 44560051 | 44560551 | Runx2 | Intron 6 | 253676 |

| | | | | | |
|---|---|---|---|---|---|
| 17 | 44560051 | 44560551 | Runx2os3 | Promoter | -1439 |
| 17 | 45530551 | 45531051 | Tcte1 | 5' UTR | 7117 |
| 17 | 45530551 | 45531051 | Tcte1 | Promoter | -2002 |
| 17 | 46768051 | 46768551 | Ptcra | Promoter | -4338 |
| 17 | 46768051 | 46768551 | 2310039H08Rik | Promoter | -4084 |
| 17 | 47518051 | 47518551 | Ccnd3 | 5' UTR | 13000 |
| 17 | 47512551 | 47513051 | Ccnd3 | 5' UTR | 7500 |
| 17 | 47518051 | 47518551 | Ccnd3 | 5' UTR | 13000 |
| 17 | 47512551 | 47513051 | Ccnd3 | 5' UTR | 7500 |
| 17 | 47518051 | 47518551 | Ccnd3 | Intron 1 | 12972 |
| 17 | 47512551 | 47513051 | Ccnd3 | Intron 1 | 7472 |
| 17 | 47518051 | 47518551 | Ccnd3 | 5' UTR | 12932 |
| 17 | 47512551 | 47513051 | Ccnd3 | 5' UTR | 7432 |
| 17 | 47518051 | 47518551 | Ccnd3 | 5' UTR | 12923 |
| 17 | 47512551 | 47513051 | Ccnd3 | 5' UTR | 7423 |
| 17 | 47518051 | 47518551 | Ccnd3 | 5' UTR | 12902 |
| 17 | 47512551 | 47513051 | Ccnd3 | 5' UTR | 7402 |
| 17 | 47518051 | 47518551 | Ccnd3 | 5' UTR | 12895 |
| 17 | 47512551 | 47513051 | Ccnd3 | 5' UTR | 7395 |
| 17 | 47518051 | 47518551 | Ccnd3 | Non-coding | 12889 |
| 17 | 47512551 | 47513051 | Ccnd3 | Non-coding | 7389 |
| 17 | 47518051 | 47518551 | Ccnd3 | Non-coding | 12832 |
| 17 | 47512551 | 47513051 | Ccnd3 | Non-coding | 7332 |
| 17 | 47518051 | 47518551 | Ccnd3 | 5' UTR | 12820 |
| 17 | 47512551 | 47513051 | Ccnd3 | 5' UTR | 7320 |
| 17 | 47518051 | 47518551 | Ccnd3 | Non-coding | 12811 |
| 17 | 47512551 | 47513051 | Ccnd3 | Non-coding | 7311 |
| 17 | 47518051 | 47518551 | Ccnd3 | 5' UTR | 12810 |
| 17 | 47512551 | 47513051 | Ccnd3 | 5' UTR | 7310 |
| 17 | 47768551 | 47769051 | Tfeb | 5' UTR | 31521 |
| 17 | 47768551 | 47769051 | Tfeb | 5' UTR | 30951 |
| 17 | 47768551 | 47769051 | Tfeb | 5' UTR | 30623 |
| 17 | 47768551 | 47769051 | Tfeb | 5' UTR | 29648 |
| 17 | 47768551 | 47769051 | Tfeb | 5' UTR | 14225 |
| 17 | 47768551 | 47769051 | Tfeb | Intron 1 | 9376 |
| 17 | 47768551 | 47769051 | Tfeb | 5' UTR | 5381 |
| 17 | 47768551 | 47769051 | Tfeb | Promoter | -187 |
| 17 | 47768551 | 47769051 | Tfeb | Promoter | -187 |
| 17 | 47898551 | 47899051 | Foxp4 | Intron 2 | 25595 |
| 17 | 47898551 | 47899051 | Foxp4 | Intron 2 | 25385 |
| 17 | 47898551 | 47899051 | Foxp4 | Intron 2 | 25350 |
| 17 | 47898551 | 47899051 | Foxp4 | Intron 2 | 25426 |
| 17 | 47898551 | 47899051 | Foxp4 | Non-coding | 25376 |
| 17 | 48055050 | 48055550 | Gm20596 | Promoter | 598 |
| 17 | 65386160 | 65386660 | Tmem232 | Intron 10 | 130570 |

| | | | | | |
|---|---|---|---|---|---|
| 17 | 65386160 | 65386660 | Tmem232 | Intron 12 | 154123 |
| 17 | 65386160 | 65386660 | Tmem232 | Intron 10 | 123394 |
| 17 | 65386160 | 65386660 | Tmem232 | Intron 13 | 154117 |
| 17 | 65386160 | 65386660 | Tmem232 | Intron 12 | 154085 |
| 17 | 65386160 | 65386660 | Tmem232 | Non-coding | 97842 |
| 17 | 65386160 | 65386660 | Tmem232 | Non-coding | 49736 |
| 17 | 70705660 | 70706160 | Dlgap1 | Intron 8 | 736587 |
| 17 | 70705660 | 70706160 | Dlgap1 | Intron 9 | 736414 |
| 17 | 70705660 | 70706160 | Dlgap1 | Intron 8 | 736258 |
| 17 | 70705660 | 70706160 | Dlgap1 | Intron 5 | 280866 |
| 17 | 70705660 | 70706160 | Dlgap1 | Intron 4 | 189638 |
| 17 | 70705660 | 70706160 | Dlgap1 | Intron 4 | 189012 |
| 17 | 70705660 | 70706160 | Dlgap1 | Intron 4 | 183983 |
| 17 | 70705660 | 70706160 | Dlgap1 | Intron 4 | 183605 |
| 17 | 70705660 | 70706160 | Dlgap1 | Non-coding | 183524 |
| 17 | 70705660 | 70706160 | Dlgap1 | Intron 4 | 183524 |
| 17 | 70705660 | 70706160 | Dlgap1 | Intron 4 | 183498 |
| 17 | 70705660 | 70706160 | Dlgap1 | Intron 4 | 144108 |
| 17 | 70705660 | 70706160 | Dlgap1 | Non-coding | 144108 |
| 17 | 78604660 | 78605160 | Vit | Intron 9 | 96596 |
| 17 | 78604660 | 78605160 | Vit | Intron 8 | 96488 |
| 17 | 80601160 | 80601660 | Map4k3 | Intron 28 | 126434 |
| 17 | 80601160 | 80601660 | Map4k3 | Intron 28 | 126434 |
| 17 | 80601160 | 80601660 | Map4k3 | Intron 27 | 126365 |
| 17 | 80601160 | 80601660 | Map4k3 | Non-coding | 23327 |
| 17 | 84584160 | 84584660 | Plekhh2 | Intron 16 | 72265 |
| 17 | 84584160 | 84584660 | Gm49969 | Non-coding | 41736 |
| 17 | 85400160 | 85400660 | Camkmt | Intron 6 | 309608 |
| 17 | 86302160 | 86302660 | Prkce | Intron 1 | 134375 |
| 17 | 86302160 | 86302660 | Prkce | Intron 2 | 133986 |
| 17 | 87134160 | 87134660 | Socs5 | CDS 3 | 26481 |
| 17 | 87133660 | 87134160 | Socs5 | CDS 3 | 25981 |
| 17 | 87134160 | 87134660 | Socs5 | CDS 2 | 26469 |
| 17 | 87133660 | 87134160 | Socs5 | CDS 2 | 25969 |
| 17 | 88752160 | 88752660 | Lhcgr | Non-coding | 21397 |
| 17 | 88752160 | 88752660 | Lhcgr | Intron 9 | 39317 |
| 17 | 88752160 | 88752660 | Lhcgr | 3' UTR | 39331 |
| 18 | 6989502 | 6990002 | Mkx | Intron 5 | 14779 |
| 18 | 6989502 | 6990002 | Mkx | Promoter | 2987 |
| 18 | 12211991 | 12212491 | Npc1 | Intron 7 | 23910 |
| 18 | 12211991 | 12212491 | Npc1 | CDS 7 | 23910 |
| 18 | 12211991 | 12212491 | Npc1 | Intron 6 | 23910 |
| 18 | 12211991 | 12212491 | Npc1 | Promoter | 64 |
| 18 | 15076991 | 15077491 | Kctd1 | Intron 1 | 73956 |
| 18 | 15084491 | 15084991 | Kctd1 | Intron 1 | 66456 |
| 18 | 23760499 | 23760999 | Mapre2 | Non-coding | 8166 |

| | | | | | |
|---|---|---|---|---|---|
| 18 | 23760499 | 23760999 | Mapre2 | 5' UTR | 8166 |
| 18 | 23760499 | 23760999 | Mapre2 | 5' UTR | 8160 |
| 18 | 23760499 | 23760999 | Mapre2 | 5' UTR | 8106 |
| 18 | 23760499 | 23760999 | Mapre2 | Intron 1 | 6774 |
| 18 | 23760499 | 23760999 | Mapre2 | Non-coding | 6751 |
| 18 | 25547999 | 25548499 | Celf4 | Intron 2 | 205485 |
| 18 | 25727499 | 25727999 | Celf4 | Intron 1 | 25985 |
| 18 | 25547999 | 25548499 | Celf4 | Intron 2 | 205252 |
| 18 | 25727499 | 25727999 | Celf4 | Intron 1 | 25752 |
| 18 | 25547999 | 25548499 | Celf4 | Intron 2 | 205485 |
| 18 | 25727499 | 25727999 | Celf4 | Intron 1 | 25985 |
| 18 | 25547999 | 25548499 | Celf4 | Intron 2 | 205659 |
| 18 | 25727499 | 25727999 | Celf4 | Intron 1 | 26159 |
| 18 | 25547999 | 25548499 | Celf4 | Intron 2 | 205204 |
| 18 | 25727499 | 25727999 | Celf4 | Intron 1 | 25704 |
| 18 | 25547999 | 25548499 | Celf4 | Non-coding | 204194 |
| 18 | 25727499 | 25727999 | Celf4 | Non-coding | 24694 |
| 18 | 25727499 | 25727999 | Celf4 | Non-coding | 25955 |
| 18 | 32406846 | 32407346 | Bin1 | Intron 2 | 29616 |
| 18 | 32406846 | 32407346 | Bin1 | Intron 2 | 29601 |
| 18 | 32406846 | 32407346 | Bin1 | Intron 2 | 29601 |
| 18 | 32406846 | 32407346 | Bin1 | Intron 2 | 29589 |
| 18 | 32406846 | 32407346 | Bin1 | Promoter | -288 |
| 18 | 36006846 | 36007346 | Psd2 | Intron 12 | 42016 |
| 18 | 35979846 | 35980346 | Psd2 | CDS 3 | 15016 |
| 18 | 35979846 | 35980346 | Psd2 | Intron 3 | 15016 |
| 18 | 35979846 | 35980346 | Psd2 | CDS 4 | 15016 |
| 18 | 36006846 | 36007346 | Psd2 | Intron 12 | 41781 |
| 18 | 35979846 | 35980346 | Psd2 | CDS 3 | 14781 |
| 18 | 35979846 | 35980346 | Psd2 | Intron 3 | 14781 |
| 18 | 35979846 | 35980346 | Psd2 | CDS 4 | 14781 |
| 18 | 36006846 | 36007346 | Psd2 | Intron 12 | 41779 |
| 18 | 35979846 | 35980346 | Psd2 | CDS 3 | 14779 |
| 18 | 35979846 | 35980346 | Psd2 | Intron 3 | 14779 |
| 18 | 35979846 | 35980346 | Psd2 | CDS 4 | 14779 |
| 18 | 36006846 | 36007346 | Psd2 | 3' UTR | 41744 |
| 18 | 35979846 | 35980346 | Psd2 | CDS 3 | 14744 |
| 18 | 35979846 | 35980346 | Psd2 | Intron 3 | 14744 |
| 18 | 35979846 | 35980346 | Psd2 | CDS 4 | 14744 |
| 18 | 36006846 | 36007346 | Psd2 | Intron 12 | 41741 |
| 18 | 35979846 | 35980346 | Psd2 | CDS 3 | 14741 |
| 18 | 35979846 | 35980346 | Psd2 | Intron 3 | 14741 |
| 18 | 35979846 | 35980346 | Psd2 | CDS 4 | 14741 |
| 18 | 36999346 | 36999846 | Pcdha1 | Intron 1 | 69162 |
| 18 | 36999346 | 36999846 | Pcdha1 | Intron 1 | 69061 |
| 18 | 36999346 | 36999846 | Pcdha2 | Intron 1 | 60141 |

| | | | | | |
|---|---|---|---|---|---|
| 18 | 36999346 | 36999846 | Pcdha2 | Intron 1 | 60122 |
| 18 | 36999346 | 36999846 | Pcdha3 | Intron 1 | 53241 |
| 18 | 36999346 | 36999846 | Pcdha4 | Intron 1 | 46698 |
| 18 | 36999346 | 36999846 | Pcdha4 | Intron 1 | 46671 |
| 18 | 37674846 | 37675346 | Gm42416 | Intron 1 | 722105 |
| 18 | 37763846 | 37764346 | Gm42416 | Intron 1 | 811105 |
| 18 | 37692346 | 37692846 | Gm42416 | Intron 1 | 739605 |
| 18 | 37341346 | 37341846 | Gm42416 | Intron 1 | 388605 |
| 18 | 36999346 | 36999846 | Gm42416 | Intron 1 | 46605 |
| 18 | 36999346 | 36999846 | Pcdha5 | Intron 1 | 38906 |
| 18 | 36999346 | 36999846 | Pcdha6 | Intron 1 | 31715 |
| 18 | 36999346 | 36999846 | Pcdha6 | Intron 1 | 31698 |
| 18 | 36999346 | 36999846 | Pcdha6 | 3' UTR | 31696 |
| 18 | 37674846 | 37675346 | Gm37013 | Intron 1 | 704930 |
| 18 | 37763846 | 37764346 | Gm37013 | Intron 1 | 793930 |
| 18 | 37692346 | 37692846 | Gm37013 | Intron 1 | 722430 |
| 18 | 37341346 | 37341846 | Gm37013 | Intron 1 | 371430 |
| 18 | 36999346 | 36999846 | Gm37013 | Intron 1 | 29430 |
| 18 | 36999346 | 36999846 | Pcdha7 | Intron 1 | 25544 |
| 18 | 37674846 | 37675346 | Gm37388 | Intron 1 | 698905 |
| 18 | 37763846 | 37764346 | Gm37388 | Intron 1 | 787905 |
| 18 | 37692346 | 37692846 | Gm37388 | Intron 1 | 716405 |
| 18 | 37341346 | 37341846 | Gm37388 | Intron 1 | 365405 |
| 18 | 36999346 | 36999846 | Gm37388 | Intron 1 | 23405 |
| 18 | 36999346 | 36999846 | Pcdha8 | Intron 1 | 6975 |
| 18 | 36999346 | 36999846 | Pcdha9 | Promoter | 1466 |
| 18 | 36999346 | 36999846 | Pcdha9 | CDS 1 | 1466 |
| 18 | 37341346 | 37341846 | Pcdhb7 | Promoter | 0 |
| 18 | 37341346 | 37341846 | Pcdhb7 | 5' UTR | 0 |
| 18 | 37341346 | 37341846 | Pcdhb7 | CDS 1 | 0 |
| 18 | 37674846 | 37675346 | Pcdhga1 | Intron 1 | 13053 |
| 18 | 37763846 | 37764346 | Pcdhga1 | Intron 1 | 102053 |
| 18 | 37692346 | 37692846 | Pcdhga1 | Intron 1 | 30553 |
| 18 | 37674846 | 37675346 | Pcdhga2 | Intron 1 | 5893 |
| 18 | 37763846 | 37764346 | Pcdhga2 | Intron 1 | 94893 |
| 18 | 37692346 | 37692846 | Pcdhga2 | Intron 1 | 23393 |
| 18 | 37674846 | 37675346 | Pcdhga3 | Promoter | 539 |
| 18 | 37674846 | 37675346 | Pcdhga3 | CDS 1 | 539 |
| 18 | 37763846 | 37764346 | Pcdhga3 | Intron 1 | 89539 |
| 18 | 37692346 | 37692846 | Pcdhga3 | Intron 1 | 18039 |
| 18 | 37674846 | 37675346 | Pcdhgb1 | Promoter | -4887 |
| 18 | 37763846 | 37764346 | Pcdhgb1 | Intron 1 | 83613 |
| 18 | 37692346 | 37692846 | Pcdhgb1 | Intron 1 | 12113 |
| 18 | 37674846 | 37675346 | Pcdhgb1 | Promoter | -4911 |
| 18 | 37763846 | 37764346 | Pcdhga4 | Intron 1 | 78636 |
| 18 | 37692346 | 37692846 | Pcdhga4 | Intron 1 | 7136 |

| | | | | | |
|---|---|---|---|---|---|
| 18 | 37763846 | 37764346 | Pcdhgb2 | Intron 1 | 74018 |
| 18 | 37692346 | 37692846 | Pcdhgb2 | Promoter | 2518 |
| 18 | 37692346 | 37692846 | Pcdhgb2 | CDS 1 | 2518 |
| 18 | 37692346 | 37692846 | Pcdhgb2 | Intron 1 | 2518 |
| 18 | 37692346 | 37692846 | Pcdhgb2 | Promoter | 2511 |
| 18 | 37763846 | 37764346 | Pcdhga5 | Intron 1 | 69466 |
| 18 | 37692346 | 37692846 | Pcdhga5 | Promoter | -1534 |
| 18 | 37763846 | 37764346 | Pcdhga6 | Intron 1 | 56807 |
| 18 | 37763846 | 37764346 | Pcdhga7 | Intron 1 | 49082 |
| 18 | 37763846 | 37764346 | Pcdhgb4 | Intron 1 | 43477 |
| 18 | 37763846 | 37764346 | Pcdhga8 | Intron 1 | 38140 |
| 18 | 37763846 | 37764346 | Pcdhgb5 | Intron 1 | 32837 |
| 18 | 37763846 | 37764346 | Pcdhga9 | Intron 1 | 26910 |
| 18 | 37763846 | 37764346 | Pcdhgb6 | Intron 1 | 21752 |
| 18 | 37763846 | 37764346 | Pcdhga10 | Intron 1 | 16853 |
| 18 | 37763846 | 37764346 | Pcdhgb7 | Intron 1 | 12246 |
| 18 | 37763846 | 37764346 | Gm26672 | Non-coding | 3196 |
| 18 | 37763846 | 37764346 | Gm26672 | Non-coding | 3897 |
| 18 | 37763846 | 37764346 | Pcdhga11 | Intron 1 | 8115 |
| 18 | 37763846 | 37764346 | Pcdhgb8 | Promoter | 2045 |
| 18 | 37763846 | 37764346 | Pcdhgb8 | CDS 1 | 2045 |
| 18 | 37763846 | 37764346 | Pcdhgb8 | Intron 1 | 2045 |
| 18 | 37763846 | 37764346 | Pcdhgb8 | Promoter | 1967 |
| 18 | 37763846 | 37764346 | Pcdhgb8 | CDS 1 | 1967 |
| 18 | 37763846 | 37764346 | Pcdhgb8 | Intron 1 | 1967 |
| 18 | 37763846 | 37764346 | Pcdhga12 | Promoter | -1234 |
| 18 | 37763846 | 37764346 | Pcdhga12 | Promoter | -1427 |
| 18 | 38264346 | 38264846 | Dele1 | 3' UTR | 14058 |
| 18 | 39669346 | 39669846 | Gm50397 | Non-coding | 65114 |
| 18 | 42443846 | 42444346 | Gm16415 | Promoter | 2827 |
| 18 | 46273846 | 46274346 | Pggt1b | Intron 2 | 6648 |
| 18 | 46273846 | 46274346 | Pggt1b | Intron 1 | 6485 |
| 18 | 46273846 | 46274346 | Pggt1b | Intron 2 | 6482 |
| 18 | 46273846 | 46274346 | Pggt1b | Intron 2 | 6474 |
| 18 | 46273846 | 46274346 | Pggt1b | Non-coding | 6482 |
| 18 | 46273846 | 46274346 | Pggt1b | Promoter | 1584 |
| 18 | 46273846 | 46274346 | Pggt1b | Non-coding | 6601 |
| 18 | 46714846 | 46715346 | Cdo1 | Intron 4 | 13050 |
| 18 | 46714846 | 46715346 | Cdo1 | Intron 3 | 13010 |
| 18 | 46714846 | 46715346 | Cdo1 | 3' UTR | 12995 |
| 18 | 47646346 | 47646846 | Gm5095 | Non-coding | 108611 |
| 18 | 50027846 | 50028346 | Tnfaip8 | Intron 1 | 48419 |
| 18 | 50027846 | 50028346 | Tnfaip8 | Non-coding | 48355 |
| 18 | 50027846 | 50028346 | Tnfaip8 | Intron 1 | 48317 |
| 18 | 50027846 | 50028346 | Tnfaip8 | Non-coding | 48311 |
| 18 | 50027846 | 50028346 | Tnfaip8 | Intron 1 | 8279 |

| | | | | | |
|---|---|---|---|---|---|
| 18 | 50027846 | 50028346 | Tnfaip8 | Promoter | -2672 |
| 18 | 50027846 | 50028346 | Tnfaip8 | Promoter | -2682 |
| 18 | 50027846 | 50028346 | Tnfaip8 | Promoter | -2683 |
| 18 | 54631846 | 54632346 | 9330117O12Rik | Non-coding | 20102 |
| 18 | 57939346 | 57939846 | Slc12a2 | Intron 23 | 60668 |
| 18 | 60652846 | 60653346 | Synpo | Promoter | -4526 |
| 18 | 60652846 | 60653346 | Synpo | Non-coding | 6797 |
| 18 | 60652846 | 60653346 | Synpo | Promoter | -4573 |
| 18 | 60652846 | 60653346 | Synpo | Promoter | -4573 |
| 18 | 61018346 | 61018846 | Slc6a7 | Promoter | -4123 |
| 18 | 61018346 | 61018846 | Slc6a7 | Promoter | -4139 |
| 18 | 61054846 | 61055346 | Pdgfrb | 5' UTR | 9696 |
| 18 | 61054846 | 61055346 | Pdgfrb | Intron 2 | 9646 |
| 18 | 61219846 | 61220346 | Pde6a | Promoter | 0 |
| 18 | 61219846 | 61220346 | Pde6a | 5' UTR | 0 |
| 18 | 61736846 | 61737346 | Afap1l1 | CDS 16 | 49357 |
| 18 | 61736846 | 61737346 | Afap1l1 | Intron 15 | 49357 |
| 18 | 61736846 | 61737346 | Afap1l1 | 3' UTR | 49266 |
| 18 | 61736846 | 61737346 | Afap1l1 | Non-coding | 49315 |
| 18 | 61736846 | 61737346 | 1500015A07Rik | Promoter | 2489 |
| 18 | 63803346 | 63803846 | Wdr7 | Intron 19 | 94661 |
| 18 | 63803346 | 63803846 | Wdr7 | 3' UTR | 94633 |
| 18 | 63803346 | 63803846 | Wdr7 | Intron 18 | 94631 |
| 18 | 63803346 | 63803846 | Wdr7 | Intron 2 | 25374 |
| 18 | 64930846 | 64931346 | Nedd4l | Intron 1 | 43141 |
| 18 | 64930846 | 64931346 | Nedd4l | Intron 1 | 43014 |
| 18 | 64930846 | 64931346 | Nedd4l | Non-coding | 42998 |
| 18 | 64930846 | 64931346 | Nedd4l | Intron 1 | 42888 |
| 18 | 64930846 | 64931346 | Nedd4l | Intron 1 | 42798 |
| 18 | 64930846 | 64931346 | Nedd4l | Intron 1 | 41917 |
| 18 | 64930846 | 64931346 | Nedd4l | Non-coding | 41856 |
| 18 | 64930846 | 64931346 | Nedd4l | 5' UTR | 41855 |
| 18 | 64930846 | 64931346 | Nedd4l | Non-coding | 41831 |
| 18 | 64930846 | 64931346 | Nedd4l | 5' UTR | 41817 |
| 18 | 64930846 | 64931346 | Nedd4l | 5' UTR | 40353 |
| 18 | 64930846 | 64931346 | Nedd4l | Intron 1 | 9729 |
| 18 | 64930846 | 64931346 | Nedd4l | Intron 1 | 9729 |
| 18 | 64930846 | 64931346 | Nedd4l | Non-coding | 9717 |
| 18 | 66037346 | 66037846 | Gm50142 | Promoter | 519 |
| 18 | 67089346 | 67089846 | Gnal | Promoter | 1130 |
| 18 | 67089346 | 67089846 | Gnal | Intron 1 | 1130 |
| 18 | 67089346 | 67089846 | Gnal | Promoter | 1062 |
| 18 | 67089346 | 67089846 | Gnal | 3' UTR | 1062 |
| 18 | 67089346 | 67089846 | Gnal | Promoter | 1048 |

| | | | | | |
|---|---|---|---|---|---|
| 18 | 67089346 | 67089846 | Gnal | Intron 1 | 1048 |
| 18 | 67089346 | 67089846 | Gnal | Promoter | 1011 |
| 18 | 67089346 | 67089846 | Gnal | Intron 1 | 1011 |
| 18 | 69666346 | 69666846 | Tcf4 | Intron 16 | 322200 |
| 18 | 69666346 | 69666846 | Tcf4 | Intron 15 | 321857 |
| 18 | 69666346 | 69666846 | Tcf4 | Intron 14 | 321854 |
| 18 | 69666346 | 69666846 | Tcf4 | Intron 14 | 321817 |
| 18 | 69666346 | 69666846 | Tcf4 | Intron 15 | 320627 |
| 18 | 69666346 | 69666846 | Tcf4 | Intron 15 | 320625 |
| 18 | 69666346 | 69666846 | Tcf4 | Intron 15 | 320625 |
| 18 | 69666346 | 69666846 | Tcf4 | Intron 13 | 320585 |
| 18 | 69666346 | 69666846 | Tcf4 | Intron 14 | 320495 |
| 18 | 69666346 | 69666846 | Tcf4 | Intron 15 | 319986 |
| 18 | 69666346 | 69666846 | Tcf4 | Intron 12 | 319047 |
| 18 | 69666346 | 69666846 | Tcf4 | Intron 14 | 318079 |
| 18 | 69666346 | 69666846 | Tcf4 | Intron 14 | 317823 |
| 18 | 69666346 | 69666846 | Tcf4 | Intron 13 | 250449 |
| 18 | 69666346 | 69666846 | Tcf4 | Intron 11 | 165649 |
| 18 | 69666346 | 69666846 | Tcf4 | Intron 11 | 146867 |
| 18 | 69666346 | 69666846 | Tcf4 | Intron 11 | 146580 |
| 18 | 69666346 | 69666846 | Tcf4 | Intron 10 | 146570 |
| 18 | 69666346 | 69666846 | Tcf4 | Intron 10 | 145048 |
| 18 | 69666346 | 69666846 | Tcf4 | Intron 10 | 144485 |
| 18 | 69666346 | 69666846 | Tcf4 | Intron 8 | 73358 |
| 18 | 69666346 | 69666846 | Tcf4 | Intron 6 | 72933 |
| 18 | 69666346 | 69666846 | Tcf4 | Intron 6 | 72927 |
| 18 | 69666346 | 69666846 | Tcf4 | Intron 8 | 72920 |
| 18 | 69666346 | 69666846 | Tcf4 | Intron 8 | 53902 |
| 18 | 75183846 | 75184346 | Dym | Intron 13 | 165061 |
| 18 | 75183846 | 75184346 | Gm8807 | Promoter | -1031 |
| 18 | 75607846 | 75608346 | Ctif | Intron 5 | 89209 |
| 18 | 75838846 | 75839346 | Zbtb7c | 5' UTR | 18668 |
| 18 | 75838846 | 75839346 | Zbtb7c | 5' UTR | 18631 |
| 18 | 76223846 | 76224346 | Gm9028 | Promoter | 0 |
| 18 | 77281261 | 77281761 | Gm50284 | Non-coding | 16918 |
| 18 | 77281261 | 77281761 | Loxhd1 | Promoter | -197 |
| 18 | 77281261 | 77281761 | Loxhd1 | Promoter | -197 |
| 18 | 77281261 | 77281761 | Loxhd1 | Promoter | -197 |
| 18 | 80083261 | 80083761 | Pard6g | Intron 2 | 36371 |
| 18 | 80083261 | 80083761 | Gm50195 | Promoter | -3045 |
| 18 | 80305761 | 80306261 | Kcng2 | Intron 3 | 57994 |
| 18 | 80305761 | 80306261 | Gm37015 | Promoter | -2080 |
| 18 | 80974261 | 80974761 | Sall3 | CDS 2 | 12011 |
| 18 | 80974261 | 80974761 | Sall3 | Intron 1 | 12011 |
| 18 | 80974261 | 80974761 | Sall3 | CDS 2 | 11776 |
| 18 | 80974261 | 80974761 | Sall3 | Intron 1 | 11776 |

| | | | | | |
|---|---|---|---|---|---|
| 18 | 80974261 | 80974761 | Sall3 | Promoter | 271 |
| 18 | 81511261 | 81511761 | Gm50412 | Non-coding | 30904 |
| 18 | 81511261 | 81511761 | Gm50412 | Non-coding | 30904 |
| 18 | 84729608 | 84730108 | Dipk1c | Intron 1 | 9589 |
| 18 | 84729608 | 84730108 | Dipk1c | CDS 2 | 9589 |
| 18 | 84729608 | 84730108 | Dipk1c | 5' UTR | 9366 |
| 18 | 84729608 | 84730108 | Dipk1c | 3' UTR | 9363 |
| 18 | 84729608 | 84730108 | Dipk1c | 3' UTR | 9331 |
| 18 | 84729608 | 84730108 | Dipk1c | Non-coding | 8725 |
| 18 | 84729608 | 84730108 | Dipk1c | 5' UTR | 8725 |
| 19 | 3084000 | 3084500 | 1700030N03Rik | Non-coding | 113233 |
| 19 | 3083500 | 3084000 | 1700030N03Rik | Non-coding | 113733 |
| 19 | 3084000 | 3084500 | 1700030N03Rik | Non-coding | 113215 |
| 19 | 3083500 | 3084000 | 1700030N03Rik | Non-coding | 113715 |
| 19 | 3084000 | 3084500 | 1700030N03Rik | Non-coding | 113215 |
| 19 | 3083500 | 3084000 | 1700030N03Rik | Non-coding | 113715 |
| 19 | 4933000 | 4933500 | Dpp3 | Promoter | -4712 |
| 19 | 4933000 | 4933500 | Dpp3 | Promoter | -4752 |
| 19 | 4933000 | 4933500 | Dpp3 | Promoter | -4789 |
| 19 | 4933000 | 4933500 | Peli3 | Intron 6 | 9628 |
| 19 | 4933000 | 4933500 | Peli3 | Intron 5 | 9593 |
| 19 | 5750500 | 5751000 | Ltbp3 | Intron 12 | 9596 |
| 19 | 5750500 | 5751000 | Ltbp3 | CDS 13 | 9596 |
| 19 | 5750500 | 5751000 | Ltbp3 | Non-coding | 9596 |
| 19 | 5750500 | 5751000 | Ltbp3 | Non-coding | 8041 |
| 19 | 5750500 | 5751000 | Ltbp3 | Non-coding | 6597 |
| 19 | 5750500 | 5751000 | Ltbp3 | Promoter | 0 |
| 19 | 5750500 | 5751000 | Ltbp3 | Promoter | -4062 |
| 19 | 5750500 | 5751000 | Ltbp3 | Promoter | -4616 |
| 19 | 6305500 | 6306000 | Cdc42bpg | Promoter | -456 |
| 19 | 6403500 | 6404000 | Rasgrp2 | CDS 5 | 4160 |
| 19 | 6403500 | 6404000 | Rasgrp2 | Intron 5 | 4160 |
| 19 | 6403500 | 6404000 | Rasgrp2 | 3' UTR | 4044 |
| 19 | 6403500 | 6404000 | Rasgrp2 | CDS 5 | 4044 |
| 19 | 6403500 | 6404000 | Rasgrp2 | CDS 5 | 3626 |
| 19 | 6403500 | 6404000 | Rasgrp2 | Intron 5 | 3626 |
| 19 | 6403500 | 6404000 | Rasgrp2 | 3' UTR | 3387 |
| 19 | 6403500 | 6404000 | Rasgrp2 | CDS 4 | 3387 |
| 19 | 6403500 | 6404000 | Rasgrp2 | Promoter | 2973 |
| 19 | 6403500 | 6404000 | Rasgrp2 | Promoter | 2967 |

| | | | | | |
|---|---|---|---|---|---|
| 19 | 6403500 | 6404000 | Rasgrp2 | 3' UTR | 2967 |
| 19 | 6403500 | 6404000 | Rasgrp2 | Promoter | 2953 |
| 19 | 6403500 | 6404000 | Rasgrp2 | CDS 5 | 2953 |
| 19 | 6403500 | 6404000 | Rasgrp2 | Intron 5 | 2953 |
| 19 | 6403500 | 6404000 | Rasgrp2 | Promoter | 2938 |
| 19 | 6403500 | 6404000 | Rasgrp2 | CDS 5 | 2938 |
| 19 | 6403500 | 6404000 | Rasgrp2 | Intron 5 | 2938 |
| 19 | 6403500 | 6404000 | Rasgrp2 | Promoter | 2936 |
| 19 | 6403500 | 6404000 | Rasgrp2 | CDS 5 | 2936 |
| 19 | 6403500 | 6404000 | Rasgrp2 | Intron 5 | 2936 |
| 19 | 6403500 | 6404000 | Rasgrp2 | Promoter | 2917 |
| 19 | 6403500 | 6404000 | Rasgrp2 | CDS 5 | 2917 |
| 19 | 6403500 | 6404000 | Rasgrp2 | Intron 5 | 2917 |
| 19 | 6403500 | 6404000 | Rasgrp2 | Promoter | 2877 |
| 19 | 6403500 | 6404000 | Rasgrp2 | 3' UTR | 2877 |
| 19 | 6403500 | 6404000 | Rasgrp2 | Promoter | 1805 |
| 19 | 6403500 | 6404000 | Rasgrp2 | Promoter | 1798 |
| 19 | 6403500 | 6404000 | Rasgrp2 | 3' UTR | 1798 |
| 19 | 6403500 | 6404000 | Rasgrp2 | CDS 4 | 1798 |
| 19 | 6403500 | 6404000 | Rasgrp2 | Promoter | 1782 |
| 19 | 6403500 | 6404000 | Rasgrp2 | 3' UTR | 1782 |
| 19 | 6403500 | 6404000 | Rasgrp2 | Promoter | 1781 |
| 19 | 6403500 | 6404000 | Rasgrp2 | Promoter | 582 |
| 19 | 6403500 | 6404000 | Rasgrp2 | Promoter | -582 |
| 19 | 6403500 | 6404000 | Rasgrp2 | Promoter | -741 |
| 19 | 6403500 | 6404000 | Rasgrp2 | Promoter | -827 |
| 19 | 6403500 | 6404000 | Rasgrp2 | Promoter | -4405 |
| 19 | 6660000 | 6660500 | 2410152P15Rik | Non-coding | 14049 |
| 19 | 8712510 | 8713010 | Slc3a2 | Intron 3 | 10360 |
| 19 | 8712510 | 8713010 | Slc3a2 | CDS 3 | 10360 |
| 19 | 8712510 | 8713010 | Slc3a2 | Intron 2 | 10360 |
| 19 | 8712510 | 8713010 | Slc3a2 | Promoter | 0 |
| 19 | 8712510 | 8713010 | Slc3a2 | 5' UTR | 0 |
| 19 | 8712510 | 8713010 | Slc3a2 | Promoter | 870 |
| 19 | 8712510 | 8713010 | Slc3a2 | Intron 2 | 870 |
| 19 | 8712510 | 8713010 | Slc3a2 | CDS 2 | 870 |
| 19 | 8712510 | 8713010 | Slc3a2 | Intron 1 | 870 |
| 19 | 8712510 | 8713010 | Slc3a2 | Promoter | 346 |
| 19 | 8712510 | 8713010 | Slc3a2 | Intron 2 | 346 |
| 19 | 8712510 | 8713010 | Slc3a2 | CDS 2 | 346 |
| 19 | 8712510 | 8713010 | Slc3a2 | Intron 1 | 346 |
| 19 | 8712510 | 8713010 | Slc3a2 | Promoter | 347 |
| 19 | 8712510 | 8713010 | Slc3a2 | Intron 2 | 347 |
| 19 | 8712510 | 8713010 | Slc3a2 | CDS 2 | 347 |
| 19 | 8712510 | 8713010 | Slc3a2 | Intron 1 | 347 |
| 19 | 8712510 | 8713010 | Slc3a2 | Promoter | 1424 |

| | | | | | |
|---|---|---|---|---|---|
| 19 | 8712510 | 8713010 | Slc3a2 | Intron 3 | 1424 |
| 19 | 8712510 | 8713010 | Slc3a2 | CDS 3 | 1424 |
| 19 | 8712510 | 8713010 | Slc3a2 | Intron 2 | 1424 |
| 19 | 8712510 | 8713010 | Slc3a2 | Promoter | 1087 |
| 19 | 8712510 | 8713010 | Slc3a2 | Promoter | 1174 |
| 19 | 10260510 | 10261010 | Dagla | Intron 9 | 43868 |
| 19 | 10260510 | 10261010 | Dagla | CDS 9 | 43868 |
| 19 | 10260510 | 10261010 | Dagla | Intron 8 | 43868 |
| 19 | 10260510 | 10261010 | Dagla | 3' UTR | 43848 |
| 19 | 10260510 | 10261010 | Dagla | Promoter | -3638 |
| 19 | 10453010 | 10453510 | Syt7 | 3' UTR | 63579 |
| 19 | 10676510 | 10677010 | Pga5 | Promoter | 1062 |
| 19 | 10676510 | 10677010 | Pga5 | Intron 2 | 1062 |
| 19 | 10676510 | 10677010 | Pga5 | CDS 2 | 1062 |
| 19 | 10676510 | 10677010 | Pga5 | Intron 1 | 1062 |
| 19 | 10676510 | 10677010 | Pga5 | Promoter | 0 |
| 19 | 18630010 | 18630510 | Ostf1 | Promoter | 1314 |
| 19 | 18630010 | 18630510 | Ostf1 | Intron 1 | 1314 |
| 19 | 18630010 | 18630510 | Ostf1 | Promoter | 1280 |
| 19 | 18630010 | 18630510 | Ostf1 | Intron 1 | 1280 |
| 19 | 18630010 | 18630510 | Ostf1 | Promoter | 1280 |
| 19 | 18630010 | 18630510 | Ostf1 | Intron 1 | 1280 |
| 19 | 18630010 | 18630510 | Ostf1 | Promoter | 1231 |
| 19 | 18630010 | 18630510 | Ostf1 | Promoter | 1209 |
| 19 | 18630010 | 18630510 | Nmrk1 | Promoter | -1440 |
| 19 | 18630010 | 18630510 | Nmrk1 | Promoter | -1440 |
| 19 | 18630010 | 18630510 | Nmrk1 | Promoter | -1440 |
| 19 | 18630010 | 18630510 | Nmrk1 | Promoter | -1449 |
| 19 | 18630010 | 18630510 | Nmrk1 | Promoter | -1452 |
| 19 | 18630010 | 18630510 | Nmrk1 | Promoter | -1488 |
| 19 | 21833510 | 21834010 | Cemip2 | Intron 18 | 55168 |
| 19 | 21833510 | 21834010 | Cemip2 | Intron 17 | 55152 |
| 19 | 21833510 | 21834010 | Gm50130 | Promoter | 0 |
| 19 | 22938510 | 22939010 | Trpm3 | Intron 21 | 799464 |
| 19 | 22938510 | 22939010 | Trpm3 | Intron 21 | 799393 |
| 19 | 22938510 | 22939010 | Trpm3 | Intron 22 | 799393 |
| 19 | 22938510 | 22939010 | Trpm3 | Intron 20 | 799393 |
| 19 | 22938510 | 22939010 | Trpm3 | Intron 21 | 799393 |
| 19 | 22938510 | 22939010 | Trpm3 | Intron 22 | 490327 |
| 19 | 22938510 | 22939010 | Trpm3 | Intron 21 | 490327 |
| 19 | 22938510 | 22939010 | Trpm3 | Intron 22 | 490327 |
| 19 | 22938510 | 22939010 | Trpm3 | Intron 20 | 490327 |
| 19 | 22938510 | 22939010 | Trpm3 | Intron 21 | 490327 |
| 19 | 22938510 | 22939010 | Trpm3 | 3' UTR | 490327 |
| 19 | 22938510 | 22939010 | Trpm3 | Intron 22 | 490278 |
| 19 | 22938510 | 22939010 | Trpm3 | Intron 22 | 245724 |

| | | | | | |
|---|---|---|---|---|---|
| 19 | 22938510 | 22939010 | Trpm3 | Intron 17 | 171876 |
| 19 | 24135010 | 24135510 | Tjp2 | Intron 3 | 89521 |
| 19 | 24135010 | 24135510 | Tjp2 | Intron 2 | 38680 |
| 19 | 24135010 | 24135510 | Tjp2 | Intron 3 | 65141 |
| 19 | 24135010 | 24135510 | Tjp2 | Intron 2 | 89517 |
| 19 | 24135010 | 24135510 | Tjp2 | Intron 2 | 7573 |
| 19 | 24135010 | 24135510 | Tjp2 | Non-coding | 11558 |
| 19 | 24135010 | 24135510 | Tjp2 | Intron 2 | 37627 |
| 19 | 40994010 | 40994510 | Blnk | Promoter | 26 |
| 19 | 40994010 | 40994510 | Blnk | 5' UTR | 26 |
| 19 | 40994010 | 40994510 | Blnk | Intron 1 | 26 |
| 19 | 40994010 | 40994510 | Blnk | CDS 1 | 26 |
| 19 | 40994010 | 40994510 | Blnk | Promoter | 0 |
| 19 | 40994010 | 40994510 | Blnk | 5' UTR | 0 |
| 19 | 40994010 | 40994510 | Blnk | Intron 1 | 0 |
| 19 | 40994010 | 40994510 | Blnk | CDS 1 | 0 |
| 19 | 40994010 | 40994510 | Blnk | Promoter | 26 |
| 19 | 42326010 | 42326510 | Crtac1 | Intron 3 | 105276 |
| 19 | 42326010 | 42326510 | Crtac1 | Intron 3 | 105274 |
| 19 | 43016010 | 43016510 | Hpse2 | Intron 3 | 371846 |
| 19 | 43016010 | 43016510 | Hpse2 | Non-coding | 207180 |
| 19 | 44694510 | 44695010 | Gm26644 | Promoter | 451 |
| 19 | 44694510 | 44695010 | Gm26644 | Promoter | 266 |
| 19 | 47257510 | 47258010 | Neurl1a | 3' UTR | 78690 |
| 19 | 47257510 | 47258010 | Neurl1a | CDS 6 | 78690 |
| 19 | 47257510 | 47258010 | Neurl1a | 3' UTR | 28599 |
| 19 | 47257510 | 47258010 | Neurl1a | CDS 6 | 28599 |
| 19 | 47526510 | 47527010 | Stn1 | Intron 3 | 10110 |
| 19 | 47526510 | 47527010 | Stn1 | Intron 3 | 10027 |
| 19 | 47526510 | 47527010 | Stn1 | Non-coding | 9955 |
| 19 | 47526510 | 47527010 | Stn1 | Non-coding | 9887 |
| 19 | 47526510 | 47527010 | Stn1 | Non-coding | 10085 |
| 19 | 47526510 | 47527010 | Stn1 | Intron 4 | 10498 |
| 19 | 47795010 | 47795510 | Cfap43 | Intron 11 | 41873 |
| 19 | 47875010 | 47875510 | Cfap43 | Non-coding | 43778 |
| 19 | 47795010 | 47795510 | Cfap43 | Non-coding | 123778 |
| 19 | 47795010 | 47795510 | Cfap43 | Non-coding | 16637 |
| 19 | 47875010 | 47875510 | Gsto2 | CDS 6 | 9476 |
| 19 | 47875010 | 47875510 | Gsto2 | Intron 6 | 9476 |
| 19 | 47875010 | 47875510 | Gsto2 | CDS 6 | 9465 |
| 19 | 47875010 | 47875510 | Gsto2 | Intron 6 | 9465 |
| 19 | 47875010 | 47875510 | Gsto2 | CDS 6 | 9430 |
| 19 | 47875010 | 47875510 | Gsto2 | Intron 6 | 9430 |
| 19 | 47875010 | 47875510 | Gsto2 | CDS 6 | 9243 |
| 19 | 47875010 | 47875510 | Gsto2 | Intron 6 | 9243 |
| 19 | 47875010 | 47875510 | Gsto2 | Intron 5 | 9233 |

| | | | | | |
|---|---|---|---|---|---|
| 19 | 47875010 | 47875510 | Gsto2 | CDS 6 | 9184 |
| 19 | 47875010 | 47875510 | Gsto2 | Intron 6 | 9184 |
| 19 | 47875010 | 47875510 | Gsto2 | CDS 4 | 4748 |
| 19 | 47875010 | 47875510 | Gsto2 | Intron 4 | 4748 |
| 19 | 48339010 | 48339510 | Sorcs3 | Intron 1 | 132985 |
| 19 | 55492010 | 55492510 | Vti1a | Intron 5 | 175715 |
| 19 | 55492010 | 55492510 | Vti1a | Intron 6 | 175660 |
| 19 | 55492010 | 55492510 | Vti1a | Intron 5 | 175659 |
| 19 | 56013010 | 56013510 | Gm50135 | Promoter | 481 |
| 2 | 4833954 | 4834454 | Gm13176 | Promoter | 159 |
| 2 | 5454454 | 5454954 | Camk1d | Intron 2 | 259562 |
| 2 | 5576454 | 5576954 | Camk1d | Intron 1 | 137562 |
| 2 | 5454454 | 5454954 | Camk1d | Intron 2 | 221093 |
| 2 | 5576454 | 5576954 | Camk1d | Intron 1 | 99093 |
| 2 | 5454454 | 5454954 | Camk1d | Non-coding | 221447 |
| 2 | 5576454 | 5576954 | Camk1d | Non-coding | 99447 |
| 2 | 5576454 | 5576954 | Camk1d | Non-coding | 137550 |
| 2 | 5876954 | 5877454 | Sec61a2 | CDS 8 | 17979 |
| 2 | 5876954 | 5877454 | Sec61a2 | Intron 7 | 17979 |
| 2 | 5876954 | 5877454 | Sec61a2 | CDS 8 | 17881 |
| 2 | 5876954 | 5877454 | Sec61a2 | Intron 7 | 17881 |
| 2 | 5876954 | 5877454 | Sec61a2 | Promoter | -493 |
| 2 | 5876954 | 5877454 | Sec61a2 | Promoter | -2157 |
| 2 | 5876954 | 5877454 | Sec61a2 | Promoter | -2663 |
| 2 | 5876954 | 5877454 | Sec61a2 | Promoter | 0 |
| 2 | 5876954 | 5877454 | Sec61a2 | Promoter | 537 |
| 2 | 5876954 | 5877454 | Sec61a2 | Promoter | 918 |
| 2 | 11545873 | 11546373 | Pfkfb3 | Intron 1 | 7705 |
| 2 | 11545873 | 11546373 | Pfkfb3 | Intron 1 | 7550 |
| 2 | 13491373 | 13491873 | Cubn | Promoter | 0 |
| 2 | 13491373 | 13491873 | Cubn | Intron 1 | 0 |
| 2 | 13491373 | 13491873 | Cubn | CDS 1 | 0 |
| 2 | 19517873 | 19518373 | 4921504E06Rik | Intron 5 | 35542 |
| 2 | 19517873 | 19518373 | 4921504E06Rik | Non-coding | 35542 |
| 2 | 24427980 | 24428480 | Pax8 | Intron 10 | 47120 |
| 2 | 24427980 | 24428480 | Pax8 | Promoter | -4897 |
| 2 | 24427980 | 24428480 | Pax8 | Intron 9 | 46618 |
| 2 | 24427980 | 24428480 | Pax8 | Intron 8 | 47085 |
| 2 | 25288480 | 25288980 | Tmem210 | Promoter | 335 |
| 2 | 25288480 | 25288980 | Tmem210 | CDS 2 | 335 |
| 2 | 25288480 | 25288980 | Tmem210 | Intron 2 | 335 |
| 2 | 25288480 | 25288980 | Tmem210 | CDS 3 | 335 |
| 2 | 25288480 | 25288980 | Tmem210 | Intron 3 | 335 |
| 2 | 25288480 | 25288980 | Tmem210 | CDS 4 | 335 |
| 2 | 25288480 | 25288980 | Tmem210 | Promoter | 279 |
| 2 | 25288480 | 25288980 | Lrrc26 | Promoter | -935 |

| | | | | | |
|---|---|---|---|---|---|
| 2 | 25288480 | 25288980 | Grin1os | Promoter | -2240 |
| 2 | 25900980 | 25901480 | Kcnt1 | CDS 15 | 37246 |
| 2 | 25900980 | 25901480 | Kcnt1 | Intron 15 | 37246 |
| 2 | 25900980 | 25901480 | Kcnt1 | CDS 16 | 37246 |
| 2 | 25900980 | 25901480 | Kcnt1 | Intron 16 | 37246 |
| 2 | 25900980 | 25901480 | Kcnt1 | CDS 16 | 37126 |
| 2 | 25900980 | 25901480 | Kcnt1 | Intron 16 | 37126 |
| 2 | 25900980 | 25901480 | Kcnt1 | CDS 17 | 37126 |
| 2 | 25900980 | 25901480 | Kcnt1 | Intron 17 | 37126 |
| 2 | 25900980 | 25901480 | Kcnt1 | CDS 15 | 37012 |
| 2 | 25900980 | 25901480 | Kcnt1 | Intron 15 | 37012 |
| 2 | 25900980 | 25901480 | Kcnt1 | CDS 16 | 37012 |
| 2 | 25900980 | 25901480 | Kcnt1 | Intron 16 | 37012 |
| 2 | 25900980 | 25901480 | Kcnt1 | CDS 15 | 23197 |
| 2 | 25900980 | 25901480 | Kcnt1 | Intron 15 | 23197 |
| 2 | 25900980 | 25901480 | Kcnt1 | CDS 16 | 23197 |
| 2 | 25900980 | 25901480 | Kcnt1 | Intron 16 | 23197 |
| 2 | 25900980 | 25901480 | Kcnt1 | CDS 16 | 23117 |
| 2 | 25900980 | 25901480 | Kcnt1 | Intron 16 | 23117 |
| 2 | 25900980 | 25901480 | Kcnt1 | CDS 17 | 23117 |
| 2 | 25900980 | 25901480 | Kcnt1 | Intron 17 | 23117 |
| 2 | 25900980 | 25901480 | Kcnt1 | Non-coding | 23110 |
| 2 | 25900980 | 25901480 | Kcnt1 | CDS 16 | 23107 |
| 2 | 25900980 | 25901480 | Kcnt1 | Intron 16 | 23107 |
| 2 | 25900980 | 25901480 | Kcnt1 | CDS 17 | 23107 |
| 2 | 25900980 | 25901480 | Kcnt1 | Intron 17 | 23107 |
| 2 | 25900980 | 25901480 | Kcnt1 | CDS 16 | 23099 |
| 2 | 25900980 | 25901480 | Kcnt1 | Intron 16 | 23099 |
| 2 | 25900980 | 25901480 | Kcnt1 | CDS 17 | 23099 |
| 2 | 25900980 | 25901480 | Kcnt1 | Intron 17 | 23099 |
| 2 | 25900980 | 25901480 | Kcnt1 | Non-coding | 6637 |
| 2 | 25900980 | 25901480 | Kcnt1 | Promoter | 89 |
| 2 | 25900980 | 25901480 | Kcnt1 | 3' UTR | 89 |
| 2 | 25900980 | 25901480 | Kcnt1 | CDS 1 | 89 |
| 2 | 25900980 | 25901480 | Kcnt1 | Intron 1 | 89 |
| 2 | 25900980 | 25901480 | Kcnt1 | CDS 2 | 89 |
| 2 | 25900980 | 25901480 | Kcnt1 | Intron 2 | 89 |
| 2 | 25900980 | 25901480 | Kcnt1 | Promoter | -2022 |
| 2 | 26257480 | 26257980 | Ccdc187 | Intron 16 | 36459 |
| 2 | 26257480 | 26257980 | Ccdc187 | CDS 16 | 36459 |
| 2 | 26257480 | 26257980 | Ccdc187 | Intron 15 | 36459 |
| 2 | 26257480 | 26257980 | Ccdc187 | Intron 17 | 36459 |
| 2 | 26257480 | 26257980 | Ccdc187 | CDS 17 | 36459 |
| 2 | 26257480 | 26257980 | Ccdc187 | Intron 16 | 36459 |
| 2 | 27008980 | 27009480 | Adamts13 | 3' UTR | 35487 |
| 2 | 27008980 | 27009480 | Adamts13 | Intron 28 | 35487 |

| | | | | | |
|---|---|---|---|---|---|
| 2 | 27008980 | 27009480 | Adamts13 | CDS 29 | 35487 |
| 2 | 27008980 | 27009480 | Adamts13 | Promoter | 2409 |
| 2 | 27008980 | 27009480 | Cacfd1 | Promoter | -446 |
| 2 | 27008980 | 27009480 | Cacfd1 | Promoter | -446 |
| 2 | 27008980 | 27009480 | Cacfd1 | Promoter | -446 |
| 2 | 27008980 | 27009480 | Cacfd1 | Promoter | -446 |
| 2 | 27008980 | 27009480 | Cacfd1 | Promoter | -533 |
| 2 | 27008980 | 27009480 | Cacfd1 | Promoter | -602 |
| 2 | 27008980 | 27009480 | Cacfd1 | Promoter | -1113 |
| 2 | 27084480 | 27084980 | Adamtsl2 | Intron 5 | 5101 |
| 2 | 27084480 | 27084980 | Adamtsl2 | CDS 6 | 5101 |
| 2 | 27084480 | 27084980 | Adamtsl2 | Intron 6 | 5101 |
| 2 | 27449980 | 27450480 | Brd3 | Intron 10 | 24919 |
| 2 | 27449980 | 27450480 | Brd3 | CDS 10 | 24919 |
| 2 | 27449980 | 27450480 | Brd3 | Intron 10 | 25207 |
| 2 | 27449980 | 27450480 | Brd3 | CDS 10 | 25207 |
| 2 | 27449980 | 27450480 | Brd3 | Intron 11 | 25194 |
| 2 | 27449980 | 27450480 | Brd3 | CDS 11 | 25194 |
| 2 | 27449980 | 27450480 | Brd3 | Intron 9 | 13637 |
| 2 | 27449980 | 27450480 | Brd3 | CDS 9 | 13637 |
| 2 | 27449980 | 27450480 | Brd3 | Intron 11 | 25175 |
| 2 | 27449980 | 27450480 | Brd3 | CDS 11 | 25175 |
| 2 | 27449980 | 27450480 | Brd3 | Intron 11 | 57183 |
| 2 | 27449980 | 27450480 | Brd3 | CDS 11 | 57183 |
| 2 | 27449980 | 27450480 | Mir7578 | Promoter | 151 |
| 2 | 31082480 | 31082980 | Fnbp1 | Intron 6 | 58823 |
| 2 | 31082480 | 31082980 | Fnbp1 | CDS 6 | 58823 |
| 2 | 31082480 | 31082980 | Fnbp1 | Intron 4 | 13521 |
| 2 | 31082480 | 31082980 | Fnbp1 | CDS 4 | 13521 |
| 2 | 31082480 | 31082980 | Fnbp1 | Intron 6 | 59029 |
| 2 | 31082480 | 31082980 | Fnbp1 | CDS 6 | 59029 |
| 2 | 31082480 | 31082980 | Fnbp1 | Intron 6 | 59017 |
| 2 | 31082480 | 31082980 | Fnbp1 | CDS 6 | 59017 |
| 2 | 31082480 | 31082980 | Fnbp1 | Intron 6 | 58978 |
| 2 | 31082480 | 31082980 | Fnbp1 | CDS 6 | 58978 |
| 2 | 31082480 | 31082980 | Fnbp1 | Non-coding | 13521 |
| 2 | 31082480 | 31082980 | Fnbp1 | Intron 5 | 22373 |
| 2 | 31082480 | 31082980 | Fnbp1 | CDS 5 | 22373 |
| 2 | 31082480 | 31082980 | Fnbp1 | Intron 6 | 58823 |
| 2 | 31082480 | 31082980 | Fnbp1 | CDS 6 | 58823 |
| 2 | 31082480 | 31082980 | Fnbp1 | Intron 6 | 58823 |
| 2 | 31082480 | 31082980 | Fnbp1 | CDS 6 | 58823 |
| 2 | 31082480 | 31082980 | Fnbp1 | Intron 6 | 58823 |
| 2 | 31082480 | 31082980 | Fnbp1 | CDS 6 | 58823 |
| 2 | 31082480 | 31082980 | Fnbp1 | Intron 6 | 58823 |
| 2 | 31082480 | 31082980 | Fnbp1 | CDS 6 | 58823 |

| | | | | | |
|---|---|---|---|---|---|
| 2 | 31082480 | 31082980 | Fnbp1 | Intron 6 | 58960 |
| 2 | 31082480 | 31082980 | Fnbp1 | CDS 6 | 58960 |
| 2 | 31082480 | 31082980 | Fnbp1 | Intron 4 | 58976 |
| 2 | 31082480 | 31082980 | Fnbp1 | Intron 6 | 33333 |
| 2 | 31082480 | 31082980 | Fnbp1 | CDS 6 | 33333 |
| 2 | 32087980 | 32088480 | Fam78a | Promoter | -3310 |
| 2 | 32087980 | 32088480 | Fam78a | Promoter | -4196 |
| 2 | 32087980 | 32088480 | Fam78a | Promoter | -3122 |
| 2 | 33455480 | 33455980 | Zbtb43 | 5' UTR | 12553 |
| 2 | 33455480 | 33455980 | Zbtb43 | Intron 1 | 6374 |
| 2 | 33455480 | 33455980 | Zbtb43 | Intron 3 | 12526 |
| 2 | 33455480 | 33455980 | Zbtb43 | 5' UTR | 9103 |
| 2 | 33455480 | 33455980 | Zbtb43 | 5' UTR | 12580 |
| 2 | 41428980 | 41429480 | Lrp1b | Intron 18 | 1224145 |
| 2 | 42056480 | 42056980 | Lrp1b | Intron 2 | 596645 |
| 2 | 41428980 | 41429480 | Lrp1b | Intron 15 | 359599 |
| 2 | 41428980 | 41429480 | Lrp1b | Intron 3 | 1223578 |
| 2 | 42056480 | 42056980 | Lrp1b | Intron 2 | 596078 |
| 2 | 42056480 | 42056980 | Lrp1b | Non-coding | 596094 |
| 2 | 42056480 | 42056980 | Lrp1b | 3' UTR | 595947 |
| 2 | 58552089 | 58552589 | Acvr1 | 5' UTR | 14238 |
| 2 | 58552089 | 58552589 | Acvr1 | 5' UTR | 14240 |
| 2 | 58552089 | 58552589 | Acvr1 | 5' UTR | 14569 |
| 2 | 62384943 | 62385443 | Dpp4 | Intron 5 | 26789 |
| 2 | 62384943 | 62385443 | Dpp4 | CDS 5 | 26789 |
| 2 | 62384943 | 62385443 | Dpp4 | Intron 4 | 26789 |
| 2 | 62384943 | 62385443 | Gm13561 | Non-coding | 27872 |
| 2 | 62384943 | 62385443 | Dpp4 | Non-coding | 26579 |
| 2 | 66784943 | 66785443 | Scn7a | Promoter | -28 |
| 2 | 68975943 | 68976443 | Cers6 | Intron 3 | 114502 |
| 2 | 68975943 | 68976443 | Cers6 | Non-coding | 114493 |
| 2 | 68975943 | 68976443 | Cers6 | Intron 3 | 114357 |
| 2 | 68975943 | 68976443 | Cers6 | Non-coding | 28683 |
| 2 | 68975943 | 68976443 | Cers6 | Non-coding | 16032 |
| 2 | 70047443 | 70047943 | Myo3b | Intron 2 | 8317 |
| 2 | 70663443 | 70663943 | Gm26558 | Promoter | -1333 |
| 2 | 70663443 | 70663943 | Gm26558 | Promoter | -1334 |
| 2 | 70663443 | 70663943 | Gorasp2 | Promoter | 1867 |
| 2 | 70663443 | 70663943 | Gorasp2 | Intron 1 | 1867 |
| 2 | 70663443 | 70663943 | Gorasp2 | Promoter | 1867 |
| 2 | 70663443 | 70663943 | Gorasp2 | 5' UTR | 1867 |
| 2 | 70663443 | 70663943 | Gorasp2 | Promoter | 1867 |
| 2 | 70663443 | 70663943 | Gorasp2 | Intron 1 | 1867 |
| 2 | 70663443 | 70663943 | Gorasp2 | Promoter | 1853 |
| 2 | 70663443 | 70663943 | Gorasp2 | Intron 1 | 1853 |
| 2 | 70663443 | 70663943 | Gorasp2 | Promoter | 1243 |

| | | | | | |
|---|---|---|---|---|---|
| 2 | 73486443 | 73486943 | Wipf1 | 5' UTR | 42792 |
| 2 | 73495443 | 73495943 | Wipf1 | 5' UTR | 33792 |
| 2 | 73486443 | 73486943 | Wipf1 | Promoter | -679 |
| 2 | 73486443 | 73486943 | Wipf1 | Promoter | 0 |
| 2 | 73486443 | 73486943 | Wipf1 | 5' UTR | 0 |
| 2 | 73486443 | 73486943 | Wipf1 | Non-coding | 42537 |
| 2 | 73495443 | 73495943 | Wipf1 | Non-coding | 33537 |
| 2 | 73486443 | 73486943 | Wipf1 | Non-coding | 42440 |
| 2 | 73495443 | 73495943 | Wipf1 | Non-coding | 33440 |
| 2 | 78072443 | 78072943 | 4930440I19Rik | Non-coding | 21220 |
| 2 | 86365843 | 86366343 | Olfr1056 | Promoter | -3373 |
| 2 | 89297843 | 89298343 | Olfr1230 | Promoter | -574 |
| 2 | 91928343 | 91928843 | Chrm4 | Promoter | 1094 |
| 2 | 91928343 | 91928843 | Chrm4 | 3' UTR | 1094 |
| 2 | 91928343 | 91928843 | Chrm4 | CDS 1 | 1094 |
| 2 | 91928343 | 91928843 | Mdk | Promoter | 2846 |
| 2 | 91928343 | 91928843 | Mdk | Promoter | 2938 |
| 2 | 91928343 | 91928843 | Mdk | Promoter | 2857 |
| 2 | 92369843 | 92370343 | Large2 | Promoter | 715 |
| 2 | 92369843 | 92370343 | Large2 | Intron 5 | 715 |
| 2 | 92369843 | 92370343 | Large2 | CDS 5 | 715 |
| 2 | 92369843 | 92370343 | Large2 | Intron 4 | 715 |
| 2 | 92369843 | 92370343 | Large2 | CDS 4 | 715 |
| 2 | 92369843 | 92370343 | Large2 | Intron 3 | 715 |
| 2 | 92369843 | 92370343 | Large2 | Promoter | 644 |
| 2 | 92369843 | 92370343 | Large2 | Intron 5 | 644 |
| 2 | 92369843 | 92370343 | Large2 | CDS 5 | 644 |
| 2 | 92369843 | 92370343 | Large2 | Intron 4 | 644 |
| 2 | 92369843 | 92370343 | Large2 | CDS 4 | 644 |
| 2 | 92369843 | 92370343 | Large2 | Intron 3 | 644 |
| 2 | 92369843 | 92370343 | Large2 | Promoter | 694 |
| 2 | 92369843 | 92370343 | Large2 | Promoter | 676 |
| 2 | 92369843 | 92370343 | Large2 | 3' UTR | 676 |
| 2 | 92369843 | 92370343 | Large2 | Promoter | 669 |
| 2 | 92369843 | 92370343 | Large2 | 3' UTR | 669 |
| 2 | 92369843 | 92370343 | Large2 | Intron 4 | 669 |
| 2 | 92369843 | 92370343 | Large2 | CDS 4 | 669 |
| 2 | 92369843 | 92370343 | Large2 | Intron 3 | 669 |
| 2 | 92369843 | 92370343 | Large2 | Promoter | 1625 |
| 2 | 92369843 | 92370343 | Large2 | Intron 4 | 1625 |
| 2 | 92369843 | 92370343 | Large2 | CDS 4 | 1625 |
| 2 | 92369843 | 92370343 | Large2 | Intron 3 | 1625 |
| 2 | 92369843 | 92370343 | Large2 | CDS 3 | 1625 |
| 2 | 92369843 | 92370343 | Large2 | Intron 2 | 1625 |
| 2 | 92369843 | 92370343 | Large2 | Promoter | 686 |
| 2 | 92369843 | 92370343 | Large2 | Intron 4 | 686 |

| | | | | | |
|---|---|---|---|---|---|
| 2 | 92369843 | 92370343 | Large2 | CDS 4 | 686 |
| 2 | 92369843 | 92370343 | Large2 | Intron 3 | 686 |
| 2 | 92369843 | 92370343 | Large2 | CDS 3 | 686 |
| 2 | 92369843 | 92370343 | Large2 | Intron 2 | 686 |
| 2 | 92369843 | 92370343 | Large2 | Promoter | 691 |
| 2 | 92369843 | 92370343 | Large2 | Intron 5 | 691 |
| 2 | 92369843 | 92370343 | Large2 | CDS 5 | 691 |
| 2 | 92369843 | 92370343 | Large2 | Intron 4 | 691 |
| 2 | 92369843 | 92370343 | Large2 | CDS 4 | 691 |
| 2 | 92369843 | 92370343 | Large2 | Intron 3 | 691 |
| 2 | 92369843 | 92370343 | Large2 | Promoter | -3321 |
| 2 | 92369843 | 92370343 | Large2 | Promoter | -3066 |
| 2 | 92369843 | 92370343 | Large2 | Promoter | -368 |
| 2 | 92369843 | 92370343 | Large2 | Promoter | -181 |
| 2 | 92369843 | 92370343 | Large2 | Promoter | 0 |
| 2 | 92369843 | 92370343 | Large2 | Promoter | 691 |
| 2 | 92369843 | 92370343 | Large2 | Promoter | 676 |
| 2 | 92369843 | 92370343 | Large2 | 3' UTR | 4527 |
| 2 | 92369843 | 92370343 | Large2 | CDS 5 | 4527 |
| 2 | 92369843 | 92370343 | Large2 | Intron 4 | 4527 |
| 2 | 92369843 | 92370343 | Large2 | CDS 4 | 4527 |
| 2 | 92369843 | 92370343 | Large2 | Intron 3 | 4527 |
| 2 | 92369843 | 92370343 | Large2 | Promoter | 1603 |
| 2 | 92369843 | 92370343 | Pex16 | Promoter | -4333 |
| 2 | 92369843 | 92370343 | Pex16 | Promoter | -4963 |
| 2 | 92393343 | 92393843 | Mapk8ip1 | Promoter | -611 |
| 2 | 92393343 | 92393843 | Mapk8ip1 | Intron 1 | 7421 |
| 2 | 92393343 | 92393843 | Mapk8ip1 | Promoter | -610 |
| 2 | 92529843 | 92530343 | Gm13817 | Non-coding | 10638 |
| 2 | 92596843 | 92597343 | Mir7222 | Promoter | 2241 |
| 2 | 92596843 | 92597343 | Chst1 | Promoter | -2364 |
| 2 | 92987343 | 92987843 | Prdm11 | Intron 5 | 57729 |
| 2 | 92987343 | 92987843 | Prdm11 | Intron 5 | 57729 |
| 2 | 92987343 | 92987843 | Prdm11 | Intron 5 | 58302 |
| 2 | 92987343 | 92987843 | Prdm11 | Intron 5 | 58211 |
| 2 | 93211343 | 93211843 | Tspan18 | Intron 6 | 122645 |
| 2 | 93211343 | 93211843 | Tspan18 | CDS 6 | 122645 |
| 2 | 93211343 | 93211843 | Tspan18 | Intron 5 | 15944 |
| 2 | 93211343 | 93211843 | Tspan18 | CDS 5 | 15944 |
| 2 | 102153843 | 102154343 | Ldlrad3 | Intron 1 | 32043 |
| 2 | 102153843 | 102154343 | Ldlrad3 | Intron 1 | 32016 |
| 2 | 107294843 | 107295343 | Kcna4 | 5' UTR | 4204 |
| 2 | 107294843 | 107295343 | Kcna4 | CDS 2 | 4204 |
| 2 | 115579264 | 115579764 | Gm13977 | Promoter | 1792 |
| 2 | 115579264 | 115579764 | BC052040 | Promoter | -1952 |
| 2 | 115579264 | 115579764 | BC052040 | Promoter | -1969 |

| | | | | | |
|---|---|---|---|---|---|
| 2 | 115579264 | 115579764 | BC052040 | Promoter | -2037 |
| 2 | 119126264 | 119126764 | Rad51 | Intron 5 | 13457 |
| 2 | 119126264 | 119126764 | Rad51 | Non-coding | 13450 |
| 2 | 119126264 | 119126764 | Rad51 | Intron 5 | 13447 |
| 2 | 119126264 | 119126764 | Rad51 | Promoter | 2461 |
| 2 | 119126264 | 119126764 | Rad51 | Intron 1 | 2461 |
| 2 | 119126264 | 119126764 | Rad51 | Promoter | -4754 |
| 2 | 125660764 | 125661264 | Shc4 | Intron 6 | 62124 |
| 2 | 125660764 | 125661264 | Shc4 | Intron 3 | 12369 |
| 2 | 125660764 | 125661264 | Shc4 | Intron 3 | 12239 |
| 2 | 127773764 | 127774264 | Nphp1 | Intron 5 | 14591 |
| 2 | 127773764 | 127774264 | Nphp1 | CDS 5 | 14591 |
| 2 | 127773764 | 127774264 | Nphp1 | Intron 4 | 14591 |
| 2 | 127773764 | 127774264 | Nphp1 | Intron 5 | 14634 |
| 2 | 127773764 | 127774264 | Nphp1 | CDS 5 | 14634 |
| 2 | 127773764 | 127774264 | Nphp1 | Intron 4 | 14634 |
| 2 | 130586764 | 130587264 | Avp | Promoter | -4209 |
| 2 | 132027264 | 132027764 | Rassf2 | Promoter | 2238 |
| 2 | 132027264 | 132027764 | Rassf2 | 5' UTR | 2238 |
| 2 | 132027264 | 132027764 | Rassf2 | Promoter | 2439 |
| 2 | 132027264 | 132027764 | Rassf2 | 5' UTR | 2439 |
| 2 | 132027264 | 132027764 | Rassf2 | Promoter | 2495 |
| 2 | 132027264 | 132027764 | Rassf2 | 5' UTR | 2495 |
| 2 | 132027264 | 132027764 | Rassf2 | Promoter | 1403 |
| 2 | 132027264 | 132027764 | Rassf2 | 5' UTR | 1403 |
| 2 | 132140264 | 132140764 | Slc23a2 | 5' UTR | 4345 |
| 2 | 132140264 | 132140764 | Slc23a2 | Non-coding | 4345 |
| 2 | 132140264 | 132140764 | Gm14051 | Promoter | -3769 |
| 2 | 138496764 | 138497264 | Btbd3 | Non-coding | 240102 |
| 2 | 138496764 | 138497264 | Btbd3 | Non-coding | 240094 |
| 2 | 143834764 | 143835264 | Bfsp1 | Intron 5 | 27910 |
| 2 | 143834764 | 143835264 | Bfsp1 | Intron 5 | 27680 |
| 2 | 144196264 | 144196764 | Gm5535 | Non-coding | 11227 |
| 2 | 144196264 | 144196764 | Gm5535 | Non-coding | 11109 |
| 2 | 146260264 | 146260764 | Ralgapa2 | Intron 39 | 251241 |
| 2 | 146260264 | 146260764 | Ralgapa2 | CDS 39 | 251241 |
| 2 | 146260264 | 146260764 | Ralgapa2 | Intron 38 | 251241 |
| 2 | 146260264 | 146260764 | Ralgapa2 | Intron 40 | 251581 |
| 2 | 146260264 | 146260764 | Ralgapa2 | CDS 40 | 251581 |
| 2 | 146260264 | 146260764 | Ralgapa2 | Intron 39 | 251581 |
| 2 | 146260264 | 146260764 | Ralgapa2 | Intron 30 | 186529 |
| 2 | 146260264 | 146260764 | Ralgapa2 | CDS 30 | 186529 |
| 2 | 146260264 | 146260764 | Ralgapa2 | Intron 29 | 186529 |
| 2 | 146260264 | 146260764 | Ralgapa2 | Promoter | 612 |
| 2 | 146260264 | 146260764 | Ralgapa2 | Intron 38 | 251136 |
| 2 | 146260264 | 146260764 | Ralgapa2 | CDS 38 | 251136 |

| | | | | | |
|---|---|---|---|---|---|
| 2 | 146260264 | 146260764 | Ralgapa2 | Intron 37 | 251136 |
| 2 | 146260264 | 146260764 | Ralgapa2 | 3' UTR | 126920 |
| 2 | 146260264 | 146260764 | A930019D19Rik | Promoter | 978 |
| 2 | 151865764 | 151866264 | Rspo4 | Intron 1 | 22837 |
| 2 | 151865764 | 151866264 | Rspo4os | Promoter | 697 |
| 2 | 153078264 | 153078764 | Ccm2l | 3' UTR | 12309 |
| 2 | 153078264 | 153078764 | Ccm2l | Intron 7 | 12309 |
| 2 | 153078264 | 153078764 | Ccm2l | CDS 8 | 12309 |
| 2 | 153078264 | 153078764 | Ccm2l | Intron 8 | 12309 |
| 2 | 153078264 | 153078764 | Ccm2l | Intron 6 | 12262 |
| 2 | 153078264 | 153078764 | Ccm2l | CDS 7 | 12262 |
| 2 | 153078264 | 153078764 | Ccm2l | Intron 7 | 12262 |
| 2 | 153834264 | 153834764 | Efcab8 | Intron 23 | 54333 |
| 2 | 154706264 | 154706764 | Rpl5-ps2 | Promoter | 2455 |
| 2 | 155753264 | 155753764 | Procr | Promoter | 2147 |
| 2 | 155753264 | 155753764 | Procr | Intron 1 | 2147 |
| 2 | 155753264 | 155753764 | Procr | CDS 2 | 2147 |
| 2 | 155753264 | 155753764 | Procr | Intron 2 | 2147 |
| 2 | 155753264 | 155753764 | Procr | Promoter | 2141 |
| 2 | 155753264 | 155753764 | Procr | Intron 2 | 2141 |
| 2 | 155753264 | 155753764 | Procr | CDS 3 | 2141 |
| 2 | 155753264 | 155753764 | Procr | Intron 3 | 2141 |
| 2 | 155753264 | 155753764 | Procr | Promoter | 2114 |
| 2 | 155753264 | 155753764 | Procr | Promoter | 1920 |
| 2 | 156048264 | 156048764 | Fer1l4 | CDS 9 | 4184 |
| 2 | 156048264 | 156048764 | Fer1l4 | Intron 8 | 4184 |
| 2 | 156048264 | 156048764 | Fer1l4 | CDS 8 | 4184 |
| 2 | 156048264 | 156048764 | Fer1l4 | Intron 7 | 4184 |
| 2 | 156093764 | 156094264 | Cpne1 | 5' UTR | 17672 |
| 2 | 156093764 | 156094264 | Cpne1 | 5' UTR | 17676 |
| 2 | 156093764 | 156094264 | Cpne1 | Non-coding | 17625 |
| 2 | 156093764 | 156094264 | Gm28036 | Intron 1 | 3822 |
| 2 | 156093764 | 156094264 | Gm28036 | Intron 3 | 17638 |
| 2 | 156093764 | 156094264 | Cpne1 | 5' UTR | 17680 |
| 2 | 156093764 | 156094264 | Cpne1 | 5' UTR | 17638 |
| 2 | 156093764 | 156094264 | Cpne1 | 5' UTR | 17641 |
| 2 | 156093764 | 156094264 | Cpne1 | 5' UTR | 17702 |
| 2 | 156093764 | 156094264 | Cpne1 | 5' UTR | 17646 |
| 2 | 156093764 | 156094264 | Cpne1 | 5' UTR | 17696 |
| 2 | 156093764 | 156094264 | Cpne1 | 5' UTR | 17646 |
| 2 | 156093764 | 156094264 | Cpne1 | 5' UTR | 17600 |
| 2 | 156093764 | 156094264 | Cpne1 | 5' UTR | 8365 |
| 2 | 156093764 | 156094264 | Gm28036 | Intron 5 | 32829 |
| 2 | 156093764 | 156094264 | Gm28036 | 3' UTR | 32829 |
| 2 | 156093764 | 156094264 | Gm28036 | Intron 5 | 32829 |

| | | | | | |
|---|---|---|---|---|---|
| 2 | 156093764 | 156094264 | Gm28036 | Intron 3 | 17663 |
| 2 | 156093764 | 156094264 | Gm28036 | Promoter | 2665 |
| 2 | 156093764 | 156094264 | Gm28036 | Intron 1 | 2665 |
| 2 | 156093764 | 156094264 | Gm28036 | Intron 3 | 17668 |
| 2 | 156093764 | 156094264 | Rbm12 | 3' UTR | 17693 |
| 2 | 156093764 | 156094264 | Rbm12 | 3' UTR | 17693 |
| 2 | 157130764 | 157131264 | Samhd1 | Intron 1 | 4002 |
| 2 | 157130764 | 157131264 | Samhd1 | Intron 1 | 3830 |
| 2 | 157130764 | 157131264 | Samhd1 | Intron 1 | 3857 |
| 2 | 157130764 | 157131264 | Samhd1 | Intron 1 | 3959 |
| 2 | 157130764 | 157131264 | Samhd1 | Intron 1 | 3943 |
| 2 | 157130764 | 157131264 | Samhd1 | Non-coding | 3674 |
| 2 | 157130764 | 157131264 | Samhd1 | Promoter | -706 |
| 2 | 158093764 | 158094264 | Rprd1b | Intron 5 | 64965 |
| 2 | 158093764 | 158094264 | 2010009K17Rik | Promoter | -1966 |
| 2 | 160908264 | 160908764 | Emilin3 | CDS 4 | 3554 |
| 2 | 160908264 | 160908764 | Emilin3 | CDS 4 | 3565 |
| 2 | 160973264 | 160973764 | Chd6 | Intron 27 | 135259 |
| 2 | 162142264 | 162142764 | Ptprt | Intron 6 | 518384 |
| 2 | 162491264 | 162491764 | Ptprt | Intron 1 | 169384 |
| 2 | 162142264 | 162142764 | Ptprt | Intron 6 | 518384 |
| 2 | 162491264 | 162491764 | Ptprt | Intron 1 | 169384 |
| 2 | 162142264 | 162142764 | Ptprt | Intron 6 | 518331 |
| 2 | 162491264 | 162491764 | Ptprt | Intron 1 | 169331 |
| 2 | 162142264 | 162142764 | Ptprt | Intron 6 | 518384 |
| 2 | 162491264 | 162491764 | Ptprt | Intron 1 | 169384 |
| 2 | 163387264 | 163387764 | Jph2 | Intron 1 | 10186 |
| 2 | 163387264 | 163387764 | Jph2 | Intron 2 | 10230 |
| 2 | 164558500 | 164559000 | Gm14317 | Non-coding | 7970 |
| 2 | 164558500 | 164559000 | Wfdc2 | Promoter | -3413 |
| 2 | 164558500 | 164559000 | Wfdc2 | Promoter | -3579 |
| 2 | 164558500 | 164559000 | Wfdc2 | Promoter | -3699 |
| 2 | 164558500 | 164559000 | Wfdc2 | Promoter | -3714 |
| 2 | 164558500 | 164559000 | Wfdc2 | Promoter | -4019 |
| 2 | 165652500 | 165653000 | Eya2 | 5' UTR | 57468 |
| 2 | 165652500 | 165653000 | Eya2 | Promoter | -2298 |
| 2 | 165652500 | 165653000 | Eya2 | Promoter | -2318 |
| 2 | 166142500 | 166143000 | Sulf2 | 5' UTR | 12664 |
| 2 | 166142500 | 166143000 | Sulf2 | 5' UTR | 12664 |
| 2 | 166142500 | 166143000 | Sulf2 | 5' UTR | 12286 |
| 2 | 166142500 | 166143000 | Sulf2 | 5' UTR | 11807 |
| 2 | 166142500 | 166143000 | Sulf2 | 5' UTR | 11424 |
| 2 | 167144500 | 167145000 | Kcnb1 | Intron 1 | 43819 |
| 2 | 167144500 | 167145000 | Kcnb1 | Intron 2 | 45156 |
| 2 | 167328500 | 167329000 | B4galt5 | Intron 1 | 20184 |
| 2 | 167657000 | 167657500 | Gm20431 | Intron 1 | 3997 |

| | | | | | |
|---|---|---|---|---|---|
| 2 | 167657000 | 167657500 | Tmem189 | Intron 1 | 4057 |
| 2 | 167657000 | 167657500 | Tmem189 | Non-coding | 32323 |
| 2 | 167657000 | 167657500 | Tmem189 | Non-coding | 4055 |
| 2 | 167657000 | 167657500 | Gm14320 | Promoter | -4609 |
| 2 | 168268500 | 168269000 | Kcng1 | CDS 2 | 12133 |
| 2 | 168268500 | 168269000 | Kcng1 | Promoter | 332 |
| 2 | 168268500 | 168269000 | Kcng1 | CDS 1 | 332 |
| 2 | 168505000 | 168505500 | Nfatc2 | CDS 9 | 84761 |
| 2 | 168505000 | 168505500 | Nfatc2 | Intron 8 | 84761 |
| 2 | 168505000 | 168505500 | Nfatc2 | CDS 9 | 84866 |
| 2 | 168505000 | 168505500 | Nfatc2 | Intron 8 | 84866 |
| 2 | 168505000 | 168505500 | Nfatc2 | Non-coding | 46777 |
| 2 | 170727000 | 170727500 | Dok5 | Promoter | -4307 |
| 2 | 172925999 | 172926499 | Bmp7 | Intron 1 | 13823 |
| 2 | 172925999 | 172926499 | Bmp7 | Non-coding | 13594 |
| 2 | 174670499 | 174670999 | Zfp831 | Intron 3 | 26965 |
| 2 | 178264795 | 178265295 | Phactr3 | Intron 3 | 145820 |
| 2 | 178264795 | 178265295 | Phactr3 | Intron 3 | 145616 |
| 2 | 178264795 | 178265295 | Phactr3 | Intron 3 | 122862 |
| 2 | 178264795 | 178265295 | Phactr3 | Intron 3 | 89446 |
| 2 | 178264795 | 178265295 | Phactr3 | Intron 3 | 71711 |
| 2 | 179411295 | 179411795 | 4930591A17Rik | Promoter | -3141 |
| 2 | 180337295 | 180337795 | Gata5 | Promoter | -2595 |
| 2 | 180337295 | 180337795 | Gata5os | Non-coding | 4438 |
| 2 | 180642795 | 180643295 | Tcfl5 | Promoter | -86 |
| 2 | 180642795 | 180643295 | Tcfl5 | Promoter | -205 |
| 2 | 180642795 | 180643295 | Tcfl5 | Promoter | -447 |
| 3 | 5203000 | 5203500 | 2700069I18Rik | Non-coding | 17324 |
| 3 | 5203000 | 5203500 | 2700069I18Rik | Non-coding | 17342 |
| 3 | 31912578 | 31913078 | Kcnmb2 | 5' UTR | 10071 |
| 3 | 31912578 | 31913078 | Kcnmb2 | 5' UTR | 9904 |
| 3 | 31912578 | 31913078 | Kcnmb2 | 5' UTR | 9882 |
| 3 | 31912578 | 31913078 | Kcnmb2 | 5' UTR | 9875 |
| 3 | 31912578 | 31913078 | Kcnmb2 | 5' UTR | 9875 |
| 3 | 32934078 | 32934578 | Usp13 | 3' UTR | 116532 |
| 3 | 32934078 | 32934578 | Gm37459 | Promoter | -4736 |
| 3 | 33131578 | 33132078 | Pex5l | Intron 1 | 11005 |
| 3 | 33131578 | 33132078 | Pex5l | Non-coding | 11170 |
| 3 | 33131578 | 33132078 | Pex5l | Non-coding | 10999 |
| 3 | 34868578 | 34869078 | Gm21388 | Non-coding | 41962 |
| 3 | 34868578 | 34869078 | Gm21388 | Non-coding | 41946 |
| 3 | 34868578 | 34869078 | Gm21388 | Non-coding | 41909 |
| 3 | 34868578 | 34869078 | Gm21388 | Non-coding | 41893 |
| 3 | 34868578 | 34869078 | Gm21388 | Non-coding | 31428 |
| 3 | 38482578 | 38483078 | Ankrd50 | Promoter | 1767 |

| | | | | | |
|---|---|---|---|---|---|
| 3 | 38482578 | 38483078 | Ankrd50 | Intron 2 | 1767 |
| 3 | 38482578 | 38483078 | Ankrd50 | CDS 2 | 1767 |
| 3 | 38482578 | 38483078 | Ankrd50 | Promoter | 14 |
| 3 | 38482578 | 38483078 | Ankrd50 | Intron 1 | 14 |
| 3 | 38482578 | 38483078 | Ankrd50 | CDS 1 | 14 |
| 3 | 38482578 | 38483078 | Ankrd50 | Promoter | 1738 |
| 3 | 51226078 | 51226578 | Noct | Promoter | 1631 |
| 3 | 51226078 | 51226578 | Noct | Intron 1 | 1631 |
| 3 | 51226078 | 51226578 | Noct | Promoter | 1608 |
| 3 | 51226078 | 51226578 | Noct | Intron 1 | 1608 |
| 3 | 63297078 | 63297578 | Mme | 5' UTR | 55541 |
| 3 | 63297078 | 63297578 | Mme | Promoter | 1812 |
| 3 | 63297078 | 63297578 | Mme | 5' UTR | 1812 |
| 3 | 63297078 | 63297578 | Mme | Promoter | 1630 |
| 3 | 63297078 | 63297578 | Mme | 5' UTR | 1630 |
| 3 | 63297078 | 63297578 | Mme | Promoter | 1517 |
| 3 | 63297078 | 63297578 | Mme | 5' UTR | 1517 |
| 3 | 63297078 | 63297578 | Mme | Promoter | 1206 |
| 3 | 63297078 | 63297578 | Mme | 5' UTR | 1206 |
| 3 | 63297078 | 63297578 | Mme | Promoter | 983 |
| 3 | 63297078 | 63297578 | Mme | 5' UTR | 983 |
| 3 | 63297078 | 63297578 | Mme | Promoter | 26 |
| 3 | 63297078 | 63297578 | Mme | 5' UTR | 26 |
| 3 | 63297078 | 63297578 | Mme | Promoter | -2582 |
| 3 | 68561078 | 68561578 | Iqschfp | Intron 5 | 668846 |
| 3 | 68561078 | 68561578 | Schip1 | Intron 2 | 496276 |
| 3 | 68561078 | 68561578 | Schip1 | Intron 1 | 496276 |
| 3 | 68561078 | 68561578 | Schip1 | Intron 2 | 92895 |
| 3 | 68561078 | 68561578 | Schip1 | Intron 2 | 67587 |
| 3 | 68561078 | 68561578 | Schip1 | Intron 2 | 66870 |
| 3 | 69588078 | 69588578 | B3galnt1 | 5' UTR | 10383 |
| 3 | 79519578 | 79520078 | Fnip2 | Intron 1 | 47719 |
| 3 | 79519578 | 79520078 | Fnip2 | Intron 1 | 47605 |
| 3 | 79519578 | 79520078 | Fnip2 | Intron 1 | 11799 |
| 3 | 79519578 | 79520078 | Fnip2 | Promoter | -4325 |
| 3 | 79519578 | 79520078 | Fnip2 | Non-coding | 47602 |
| 3 | 82999078 | 82999578 | Gm30097 | Non-coding | 8911 |
| 3 | 82999078 | 82999578 | Gm30097 | Promoter | -3492 |
| 3 | 83799578 | 83800078 | Gm7115 | Promoter | 0 |
| 3 | 84445578 | 84446078 | Fhdc1 | CDS 12 | 34352 |
| 3 | 84445578 | 84446078 | Fhdc1 | CDS 12 | 33414 |
| 3 | 84445578 | 84446078 | Fhdc1 | Intron 5 | 34316 |
| 3 | 85197078 | 85197578 | Gm37240 | 5' UTR | 689939 |
| 3 | 85197078 | 85197578 | Gm38313 | Promoter | -630 |
| 3 | 86397578 | 86398078 | Lrba | Intron 35 | 172898 |
| 3 | 86397578 | 86398078 | Lrba | Intron 35 | 172898 |

| | | | | | |
|---|---|---|---|---|---|
| 3 | 86397578 | 86398078 | Lrba | Intron 35 | 172898 |
| 3 | 86397578 | 86398078 | Lrba | Intron 34 | 172289 |
| 3 | 87620578 | 87621078 | Arhgef11 | Non-coding | 3019 |
| 3 | 87620578 | 87621078 | Arhgef11 | Intron 1 | 3003 |
| 3 | 87620578 | 87621078 | Arhgef11 | Promoter | 2993 |
| 3 | 87620578 | 87621078 | Arhgef11 | Intron 1 | 2993 |
| 3 | 87620578 | 87621078 | Arhgef11 | Promoter | 2986 |
| 3 | 87620578 | 87621078 | Arhgef11 | Promoter | 2477 |
| 3 | 87620578 | 87621078 | Arhgef11 | Intron 1 | 2477 |
| 3 | 87620578 | 87621078 | Arhgef11 | Promoter | 1817 |
| 3 | 87620578 | 87621078 | Arhgef11 | Intron 1 | 1817 |
| 3 | 88032578 | 88033078 | Hapln2 | Promoter | -4994 |
| 3 | 88032578 | 88033078 | Gm19439 | Promoter | 1700 |
| 3 | 88730578 | 88731078 | Rit1 | 3' UTR | 13684 |
| 3 | 88730578 | 88731078 | Rit1 | Non-coding | 13680 |
| 3 | 90242578 | 90243078 | Creb3l4 | Promoter | 435 |
| 3 | 90242578 | 90243078 | Creb3l4 | 5' UTR | 435 |
| 3 | 90242578 | 90243078 | Creb3l4 | Intron 2 | 435 |
| 3 | 90242578 | 90243078 | Creb3l4 | CDS 2 | 435 |
| 3 | 90242578 | 90243078 | Creb3l4 | Intron 1 | 435 |
| 3 | 90242578 | 90243078 | Creb3l4 | Promoter | 66 |
| 3 | 90242578 | 90243078 | Creb3l4 | 5' UTR | 66 |
| 3 | 90242578 | 90243078 | Creb3l4 | Intron 2 | 66 |
| 3 | 90242578 | 90243078 | Creb3l4 | CDS 2 | 66 |
| 3 | 90242578 | 90243078 | Creb3l4 | Intron 1 | 66 |
| 3 | 90242578 | 90243078 | Creb3l4 | CDS 1 | 66 |
| 3 | 93553578 | 93554078 | S100a10 | Promoter | -1002 |
| 3 | 93553578 | 93554078 | S100a10 | Promoter | -1085 |
| 3 | 94695578 | 94696078 | Selenbp2 | Promoter | 2022 |
| 3 | 94695578 | 94696078 | Selenbp2 | Intron 1 | 2022 |
| 3 | 94697078 | 94697578 | Selenbp2 | CDS 3 | 3522 |
| 3 | 94697078 | 94697578 | Selenbp2 | Intron 3 | 3522 |
| 3 | 94697078 | 94697578 | Selenbp2 | CDS 4 | 3522 |
| 3 | 94697078 | 94697578 | Selenbp2 | Intron 4 | 3522 |
| 3 | 94695578 | 94696078 | Selenbp2 | Promoter | 2012 |
| 3 | 94697078 | 94697578 | Selenbp2 | Non-coding | 3512 |
| 3 | 94695578 | 94696078 | Selenbp2 | Promoter | 2010 |
| 3 | 94695578 | 94696078 | Selenbp2 | Intron 1 | 2010 |
| 3 | 94697078 | 94697578 | Selenbp2 | CDS 2 | 3510 |
| 3 | 94697078 | 94697578 | Selenbp2 | Intron 2 | 3510 |
| 3 | 94697078 | 94697578 | Selenbp2 | CDS 3 | 3510 |
| 3 | 94697078 | 94697578 | Selenbp2 | Intron 3 | 3510 |
| 3 | 94695578 | 94696078 | Selenbp2 | Promoter | 2009 |
| 3 | 94695578 | 94696078 | Selenbp2 | Intron 1 | 2009 |
| 3 | 94697078 | 94697578 | Selenbp2 | CDS 3 | 3509 |
| 3 | 94697078 | 94697578 | Selenbp2 | Intron 3 | 3509 |

| | | | | | |
|---|---|---|---|---|---|
| 3 | 94695578 | 94696078 | Selenbp2 | Promoter | 1931 |
| 3 | 94695578 | 94696078 | Selenbp2 | Intron 1 | 1931 |
| 3 | 94697078 | 94697578 | Selenbp2 | 3' UTR | 3431 |
| 3 | 94697078 | 94697578 | Selenbp2 | Intron 2 | 3431 |
| 3 | 94697078 | 94697578 | Selenbp2 | CDS 3 | 3431 |
| 3 | 94697078 | 94697578 | Selenbp2 | Intron 3 | 3431 |
| 3 | 94695578 | 94696078 | Selenbp2 | Promoter | 1930 |
| 3 | 94697078 | 94697578 | Selenbp2 | Non-coding | 3430 |
| 3 | 94695578 | 94696078 | Selenbp2 | Promoter | 1919 |
| 3 | 94695578 | 94696078 | Selenbp2 | Intron 1 | 1919 |
| 3 | 94697078 | 94697578 | Selenbp2 | CDS 3 | 3419 |
| 3 | 94697078 | 94697578 | Selenbp2 | Intron 3 | 3419 |
| 3 | 94697078 | 94697578 | Selenbp2 | CDS 4 | 3419 |
| 3 | 94697078 | 94697578 | Selenbp2 | Intron 4 | 3419 |
| 3 | 97204077 | 97204577 | Bcl9 | 3' UTR | 22788 |
| 3 | 101409577 | 101410077 | Igsf3 | Intron 2 | 32494 |
| 3 | 101409577 | 101410077 | Igsf3 | Intron 1 | 31239 |
| 3 | 101409577 | 101410077 | Gm42939 | Promoter | 1784 |
| 3 | 101409577 | 101410077 | Gm42938 | Promoter | -1462 |
| 3 | 101817577 | 101818077 | Mab21l3 | Intron 6 | 18147 |
| 3 | 101817577 | 101818077 | Mab21l3 | Intron 6 | 30883 |
| 3 | 101897077 | 101897577 | Slc22a15 | Intron 4 | 26817 |
| 3 | 101897077 | 101897577 | Slc22a15 | CDS 4 | 26817 |
| 3 | 101897077 | 101897577 | Slc22a15 | Intron 3 | 26817 |
| 3 | 101897077 | 101897577 | Slc22a15 | Intron 2 | 26873 |
| 3 | 101897077 | 101897577 | Slc22a15 | Promoter | -206 |
| 3 | 101897077 | 101897577 | Slc22a15 | Intron 4 | 26803 |
| 3 | 101897077 | 101897577 | Slc22a15 | CDS 4 | 26803 |
| 3 | 101897077 | 101897577 | Slc22a15 | Intron 3 | 26803 |
| 3 | 102310077 | 102310577 | A230001M10Rik | Non-coding | 47672 |
| 3 | 103734077 | 103734577 | Olfml3 | Promoter | 0 |
| 3 | 103734077 | 103734577 | Olfml3 | 3' UTR | 3425 |
| 3 | 103734077 | 103734577 | Olfml3 | Promoter | -1523 |
| 3 | 103734077 | 103734577 | Olfml3 | Non-coding | 3392 |
| 3 | 103734077 | 103734577 | Gm10964 | Promoter | -4789 |
| 3 | 104424077 | 104424577 | Lrig2 | Intron 1 | 36935 |
| 3 | 104753082 | 104753582 | Gm6485 | Promoter | 0 |
| 3 | 104753082 | 104753582 | Gm43584 | Non-coding | 19689 |
| 3 | 106042582 | 106043082 | Gm9504 | Promoter | 350 |
| 3 | 107943082 | 107943582 | Gstm6 | Promoter | 168 |
| 3 | 107943082 | 107943582 | Gstm6 | Intron 2 | 168 |
| 3 | 107943082 | 107943582 | Gstm6 | CDS 2 | 168 |
| 3 | 107943082 | 107943582 | Gstm6 | Intron 1 | 168 |
| 3 | 107943082 | 107943582 | Gstm6 | Promoter | 153 |
| 3 | 107943082 | 107943582 | Gstm6 | Intron 2 | 153 |

| | | | | | |
|---|---|---|---|---|---|
| 3 | 107943082 | 107943582 | Gstm6 | CDS 2 | 153 |
| 3 | 107943082 | 107943582 | Gstm6 | Intron 1 | 153 |
| 3 | 107943082 | 107943582 | Gstm6 | Promoter | 0 |
| 3 | 107943082 | 107943582 | Gstm6 | 5' UTR | 0 |
| 3 | 107943082 | 107943582 | Gstm6 | Intron 1 | 0 |
| 3 | 107943082 | 107943582 | Gstm6 | CDS 1 | 0 |
| 3 | 107943082 | 107943582 | Gstm6 | Promoter | 0 |
| 3 | 107943082 | 107943582 | Gstm6 | 5' UTR | 0 |
| 3 | 107943082 | 107943582 | Gstm6 | Intron 1 | 0 |
| 3 | 107943082 | 107943582 | Gstm6 | CDS 1 | 0 |
| 3 | 107943082 | 107943582 | Gstm6 | Promoter | 0 |
| 3 | 107943082 | 107943582 | Gstm6 | 5' UTR | 0 |
| 3 | 107943082 | 107943582 | Gstm6 | Intron 1 | 0 |
| 3 | 107943082 | 107943582 | Gstm6 | CDS 1 | 0 |
| 3 | 107943082 | 107943582 | Gstm6 | Promoter | 134 |
| 3 | 114066082 | 114066582 | Col11a1 | Intron 3 | 35542 |
| 3 | 114066082 | 114066582 | Col11a1 | CDS 4 | 35542 |
| 3 | 114066082 | 114066582 | Col11a1 | Intron 4 | 35542 |
| 3 | 114066082 | 114066582 | Col11a1 | Intron 3 | 35542 |
| 3 | 114066082 | 114066582 | Col11a1 | CDS 4 | 35542 |
| 3 | 114066082 | 114066582 | Col11a1 | Intron 4 | 35542 |
| 3 | 122322582 | 122323082 | Bcar3 | Non-coding | 28446 |
| 3 | 123179582 | 123180082 | Synpo2 | Intron 1 | 56015 |
| 3 | 123179582 | 123180082 | Synpo2 | Intron 1 | 56066 |
| 3 | 123179582 | 123180082 | Synpo2 | Intron 1 | 56066 |
| 3 | 123179582 | 123180082 | Synpo2 | Non-coding | 56068 |
| 3 | 127487082 | 127487582 | Ank2 | Intron 1 | 11769 |
| 3 | 127480582 | 127481082 | Ank2 | Intron 1 | 18269 |
| 3 | 127487082 | 127487582 | Ank2 | Non-coding | 11421 |
| 3 | 127480582 | 127481082 | Ank2 | Non-coding | 17921 |
| 3 | 127480582 | 127481082 | Gm43355 | Promoter | 346 |
| 3 | 127480582 | 127481082 | Gm42525 | Promoter | 109 |
| 3 | 127480582 | 127481082 | Gm44494 | Promoter | -38 |
| 3 | 127687082 | 127687582 | Alpk1 | Intron 5 | 92946 |
| 3 | 127687082 | 127687582 | Alpk1 | Intron 4 | 41990 |
| 3 | 127687082 | 127687582 | Alpk1 | Non-coding | 92946 |
| 3 | 127687082 | 127687582 | Alpk1 | Promoter | -741 |
| 3 | 129902082 | 129902582 | Casp6 | Promoter | 657 |
| 3 | 129902082 | 129902582 | Casp6 | 5' UTR | 657 |
| 3 | 129902082 | 129902582 | Casp6 | Promoter | 624 |
| 3 | 129902082 | 129902582 | Casp6 | Promoter | 581 |
| 3 | 130620082 | 130620582 | Etnppl | Promoter | 2634 |
| 3 | 130620082 | 130620582 | Etnppl | Intron 2 | 2634 |
| 3 | 130620082 | 130620582 | Etnppl | CDS 3 | 2634 |
| 3 | 130620082 | 130620582 | Etnppl | Intron 3 | 2634 |
| 3 | 130620082 | 130620582 | Etnppl | Promoter | 2513 |

| | | | | | |
|---|---|---|---|---|---|
| 3 | 130620082 | 130620582 | Etnppl | Intron 1 | 2513 |
| 3 | 130620082 | 130620582 | Etnppl | CDS 2 | 2513 |
| 3 | 130620082 | 130620582 | Etnppl | Intron 2 | 2513 |
| 3 | 130620082 | 130620582 | Etnppl | Promoter | 2510 |
| 3 | 130620082 | 130620582 | Etnppl | Promoter | 2483 |
| 3 | 130620082 | 130620582 | Etnppl | Promoter | 2389 |
| 3 | 130620082 | 130620582 | Etnppl | Intron 1 | 2389 |
| 3 | 130620082 | 130620582 | Etnppl | CDS 2 | 2389 |
| 3 | 130620082 | 130620582 | Etnppl | Intron 2 | 2389 |
| 3 | 130728582 | 130729082 | Rpl34 | Promoter | 1256 |
| 3 | 130728582 | 130729082 | Rpl34 | Intron 4 | 1256 |
| 3 | 130728582 | 130729082 | Rpl34 | CDS 4 | 1256 |
| 3 | 130728582 | 130729082 | Rpl34 | Intron 3 | 1256 |
| 3 | 130728582 | 130729082 | Rpl34 | CDS 3 | 1256 |
| 3 | 130728582 | 130729082 | Rpl34 | Promoter | -1175 |
| 3 | 130728582 | 130729082 | Rpl34 | Promoter | 1317 |
| 3 | 130728582 | 130729082 | Rpl34 | Intron 4 | 1317 |
| 3 | 130728582 | 130729082 | Rpl34 | CDS 4 | 1317 |
| 3 | 130728582 | 130729082 | Rpl34 | Intron 3 | 1317 |
| 3 | 130728582 | 130729082 | Rpl34 | CDS 3 | 1317 |
| 3 | 130728582 | 130729082 | Rpl34 | Promoter | 1261 |
| 3 | 130728582 | 130729082 | Rpl34 | Promoter | 1240 |
| 3 | 130728582 | 130729082 | Rpl34 | Intron 4 | 1240 |
| 3 | 130728582 | 130729082 | Rpl34 | CDS 4 | 1240 |
| 3 | 130728582 | 130729082 | Rpl34 | Intron 3 | 1240 |
| 3 | 130728582 | 130729082 | Rpl34 | CDS 3 | 1240 |
| 3 | 130728582 | 130729082 | Rpl34 | Promoter | 1253 |
| 3 | 130728582 | 130729082 | Rpl34 | 3' UTR | 1253 |
| 3 | 130728582 | 130729082 | Rpl34 | CDS 4 | 1253 |
| 3 | 130728582 | 130729082 | Rpl34 | Intron 3 | 1253 |
| 3 | 130728582 | 130729082 | Rpl34 | CDS 3 | 1253 |
| 3 | 130728582 | 130729082 | Rpl34 | Promoter | 1266 |
| 3 | 130728582 | 130729082 | Rpl34 | Promoter | 1265 |
| 3 | 130728582 | 130729082 | Rpl34 | 3' UTR | 1265 |
| 3 | 130728582 | 130729082 | Rpl34 | CDS 3 | 1265 |
| 3 | 130728582 | 130729082 | Rpl34 | Intron 2 | 1265 |
| 3 | 130728582 | 130729082 | Rpl34 | CDS 2 | 1265 |
| 3 | 130728582 | 130729082 | Gm42997 | Promoter | -1378 |
| 3 | 130728582 | 130729082 | Gm42997 | Promoter | -1519 |
| 3 | 135493036 | 135493536 | Manba | Intron 1 | 7425 |
| 3 | 135493036 | 135493536 | Manba | Intron 1 | 7425 |
| 3 | 135493036 | 135493536 | Manba | Non-coding | 7417 |
| 3 | 135493036 | 135493536 | Manba | Non-coding | 7411 |
| 3 | 135493036 | 135493536 | Gm16100 | Promoter | -2117 |
| 3 | 138643536 | 138644036 | Gm43723 | Promoter | 0 |
| 3 | 143034536 | 143035036 | 4930519L02Rik | Non-coding | 16876 |

| | | | | | |
|---|---|---|---|---|---|
| 3 | 143034536 | 143035036 | 4930519L02Rik | Non-coding | 3922 |
| 3 | 146038536 | 146039036 | Gm35409 | Promoter | 91 |
| 3 | 149335036 | 149335536 | Gm30382 | Non-coding | 109741 |
| 3 | 149335036 | 149335536 | Gm5149 | Non-coding | 30207 |
| 3 | 149335036 | 149335536 | Gm30382 | Non-coding | 109766 |
| 3 | 149335036 | 149335536 | Gm5149 | Promoter | 190 |
| 3 | 153536536 | 153537036 | St6galnac3 | Intron 1 | 188139 |
| 3 | 153536536 | 153537036 | St6galnac3 | Intron 1 | 188078 |
| 3 | 153536536 | 153537036 | St6galnac3 | Non-coding | 183961 |
| 3 | 154654536 | 154655036 | Gm38139 | Promoter | -1665 |
| 4 | 8728353 | 8728853 | Chd7 | Non-coding | 37947 |
| 4 | 8728353 | 8728853 | Chd7 | 5' UTR | 37883 |
| 4 | 8728353 | 8728853 | Chd7 | 5' UTR | 37432 |
| 4 | 8728353 | 8728853 | Chd7 | 5' UTR | 36988 |
| 4 | 8728353 | 8728853 | Chd7 | 5' UTR | 18013 |
| 4 | 34429751 | 34430251 | Gm23466 | Promoter | -4045 |
| 4 | 41583967 | 41584467 | Dnaic1 | Intron 1 | 14192 |
| 4 | 41758127 | 41758627 | Il11ra1 | 5' UTR | 58138 |
| 4 | 41758127 | 41758627 | Sigmar1 | Promoter | -2727 |
| 4 | 41758127 | 41758627 | Sigmar1 | Promoter | -1969 |
| 4 | 41758127 | 41758627 | Galt | Promoter | 2899 |
| 4 | 41758127 | 41758627 | Galt | 3' UTR | 2899 |
| 4 | 41758127 | 41758627 | Galt | CDS 10 | 2899 |
| 4 | 41758127 | 41758627 | Galt | Intron 10 | 2899 |
| 4 | 41758127 | 41758627 | Galt | CDS 11 | 2899 |
| 4 | 41758127 | 41758627 | Galt | Promoter | 2899 |
| 4 | 41758127 | 41758627 | Galt | 3' UTR | 2899 |
| 4 | 41758127 | 41758627 | Galt | CDS 10 | 2899 |
| 4 | 41758127 | 41758627 | Galt | Promoter | 2874 |
| 4 | 41758127 | 41758627 | Galt | Promoter | 2874 |
| 4 | 41758127 | 41758627 | Galt | Promoter | 2369 |
| 4 | 41758127 | 41758627 | Galt | Promoter | 2287 |
| 4 | 41758127 | 41758627 | Galt | Promoter | 2166 |
| 4 | 41758127 | 41758627 | Galt | Promoter | 1917 |
| 4 | 41758127 | 41758627 | Galt | Promoter | 1873 |
| 4 | 41758127 | 41758627 | Galt | Promoter | 1865 |
| 4 | 41758127 | 41758627 | Il11ra1 | Promoter | -1569 |
| 4 | 41758127 | 41758627 | Il11ra1 | Promoter | -1827 |
| 4 | 41758127 | 41758627 | Il11ra1 | Promoter | -1871 |
| 4 | 41758127 | 41758627 | Il11ra1 | Promoter | -1885 |
| 4 | 41758127 | 41758627 | Il11ra1 | Promoter | -3682 |
| 4 | 41758127 | 41758627 | Il11ra1 | Promoter | -3768 |
| 4 | 43422128 | 43422628 | Rusc2 | CDS 3 | 40149 |
| 4 | 43422128 | 43422628 | Rusc2 | Intron 3 | 40149 |
| 4 | 43422128 | 43422628 | Rusc2 | CDS 4 | 40149 |
| 4 | 43422128 | 43422628 | Rusc2 | Intron 4 | 40149 |

| | | | | | |
|---|---|---|---|---|---|
| 4 | 43422128 | 43422628 | Rusc2 | Non-coding | 20890 |
| 4 | 43422128 | 43422628 | Rusc2 | CDS 3 | 15693 |
| 4 | 43422128 | 43422628 | Rusc2 | Intron 3 | 15693 |
| 4 | 43422128 | 43422628 | Rusc2 | CDS 4 | 15693 |
| 4 | 43422128 | 43422628 | Rusc2 | Intron 4 | 15693 |
| 4 | 43422128 | 43422628 | Rusc2 | Non-coding | 15626 |
| 4 | 43422128 | 43422628 | Rusc2 | CDS 2 | 7432 |
| 4 | 43422128 | 43422628 | Rusc2 | Intron 2 | 7432 |
| 4 | 43422128 | 43422628 | Rusc2 | CDS 3 | 7432 |
| 4 | 43422128 | 43422628 | Rusc2 | Intron 3 | 7432 |
| 4 | 43422128 | 43422628 | Rusc2 | 3' UTR | 7432 |
| 4 | 43422128 | 43422628 | Rusc2 | CDS 2 | 7432 |
| 4 | 43422128 | 43422628 | Rusc2 | Intron 2 | 7432 |
| 4 | 43422128 | 43422628 | Rusc2 | CDS 3 | 7432 |
| 4 | 43422128 | 43422628 | Rusc2 | Intron 3 | 7432 |
| 4 | 43422128 | 43422628 | Rusc2 | Non-coding | 5690 |
| 4 | 43422128 | 43422628 | Rusc2 | Promoter | 0 |
| 4 | 43422128 | 43422628 | Rusc2 | Promoter | -963 |
| 4 | 44607128 | 44607628 | Pax5 | Intron 7 | 102860 |
| 4 | 44607128 | 44607628 | Pax5 | Intron 7 | 102791 |
| 4 | 44607128 | 44607628 | Pax5 | Intron 7 | 102791 |
| 4 | 44607128 | 44607628 | Pax5 | Intron 6 | 102791 |
| 4 | 44607128 | 44607628 | Pax5 | Intron 6 | 102791 |
| 4 | 44607128 | 44607628 | Pax5 | Intron 6 | 102791 |
| 4 | 44607128 | 44607628 | Pax5 | Intron 6 | 102781 |
| 4 | 44607128 | 44607628 | Pax5 | Intron 6 | 102781 |
| 4 | 44607128 | 44607628 | Pax5 | Intron 6 | 102781 |
| 4 | 44607128 | 44607628 | Pax5 | Intron 6 | 102781 |
| 4 | 44607128 | 44607628 | Pax5 | Intron 7 | 102781 |
| 4 | 44607128 | 44607628 | Pax5 | Intron 7 | 102781 |
| 4 | 44607128 | 44607628 | Pax5 | Intron 6 | 102781 |
| 4 | 44607128 | 44607628 | Pax5 | Intron 3 | 38096 |
| 4 | 44607128 | 44607628 | Pax5 | Non-coding | 76188 |
| 4 | 44607128 | 44607628 | Mir5120 | Promoter | 0 |
| 4 | 45764128 | 45764628 | Gm12409 | Promoter | 0 |
| 4 | 46533628 | 46534128 | Trim14 | Promoter | 1995 |
| 4 | 46533628 | 46534128 | Trim14 | Intron 1 | 1995 |
| 4 | 46533628 | 46534128 | Trim14 | Promoter | 287 |
| 4 | 46533628 | 46534128 | Trim14 | Promoter | 2014 |
| 4 | 46533628 | 46534128 | Trim14 | Intron 1 | 2014 |
| 4 | 46533628 | 46534128 | Trim14 | Promoter | 315 |
| 4 | 46533628 | 46534128 | Trim14 | Promoter | 1987 |
| 4 | 46533628 | 46534128 | Trim14 | Intron 1 | 1987 |
| 4 | 46533628 | 46534128 | Trim14 | Promoter | 477 |
| 4 | 46533628 | 46534128 | Gm16731 | Promoter | -1773 |
| 4 | 46809628 | 46810128 | Gabbr2 | Intron 4 | 181746 |

| | | | | | |
|---|---|---|---|---|---|
| 4 | 46809628 | 46810128 | Gabbr2 | Non-coding | 49775 |
| 4 | 47350128 | 47350628 | Tgfbr1 | Promoter | -2594 |
| 4 | 47350128 | 47350628 | Tgfbr1 | Promoter | -2679 |
| 4 | 47350128 | 47350628 | Tgfbr1 | Promoter | -2970 |
| 4 | 47350128 | 47350628 | Tgfbr1 | Promoter | -2982 |
| 4 | 48536128 | 48536628 | Msantd3 | Promoter | -3307 |
| 4 | 48536128 | 48536628 | Msantd3 | Promoter | -3451 |
| 4 | 48536128 | 48536628 | Msantd3 | Promoter | -3871 |
| 4 | 48536128 | 48536628 | Msantd3 | Promoter | -3918 |
| 4 | 48895628 | 48896128 | Gm23746 | Promoter | -371 |
| 4 | 48895628 | 48896128 | Gm12436 | Promoter | -2465 |
| 4 | 53011128 | 53011628 | Nipsnap3b | Promoter | -252 |
| 4 | 53011128 | 53011628 | Nipsnap3b | Promoter | -322 |
| 4 | 61698966 | 61699466 | Mup-ps15 | Promoter | 125 |
| 4 | 62622466 | 62622966 | Rgs3 | Intron 10 | 62619 |
| 4 | 62622466 | 62622966 | Rgs3 | Intron 8 | 38865 |
| 4 | 62622466 | 62622966 | Rgs3 | Non-coding | 18007 |
| 4 | 62622466 | 62622966 | Rgs3 | Promoter | 2946 |
| 4 | 62622466 | 62622966 | Rgs3 | Intron 1 | 2946 |
| 4 | 62622466 | 62622966 | Rgs3 | Promoter | 2884 |
| 4 | 62622466 | 62622966 | Rgs3 | Intron 1 | 2884 |
| 4 | 62622466 | 62622966 | Rgs3 | Promoter | -565 |
| 4 | 62622466 | 62622966 | Rgs3 | Promoter | -743 |
| 4 | 62973466 | 62973966 | Zfp618 | 5' UTR | 7893 |
| 4 | 62973466 | 62973966 | Zfp618 | Intron 1 | 7893 |
| 4 | 62973466 | 62973966 | Zfp618 | Intron 1 | 7892 |
| 4 | 68954966 | 68955466 | Brinp1 | Promoter | -568 |
| 4 | 77040096 | 77040596 | Ptprd | Non-coding | 1171300 |
| 4 | 77040096 | 77040596 | Ptprd | Non-coding | 1171272 |
| 4 | 77040096 | 77040596 | Ptprd | Non-coding | 1171144 |
| 4 | 83655596 | 83656096 | Ccdc171 | CDS 12 | 130051 |
| 4 | 83655596 | 83656096 | Ccdc171 | Intron 12 | 130051 |
| 4 | 83655596 | 83656096 | Ccdc171 | CDS 12 | 130051 |
| 4 | 83655596 | 83656096 | Ccdc171 | Intron 12 | 130051 |
| 4 | 83655596 | 83656096 | Ccdc171 | 3' UTR | 129957 |
| 4 | 87432596 | 87433096 | Gm23154 | Promoter | 324 |
| 4 | 91385309 | 91385809 | Elavl2 | Intron 2 | 14176 |
| 4 | 91385309 | 91385809 | Elavl2 | Non-coding | 14168 |
| 4 | 94739809 | 94740309 | Tek | Promoter | 520 |
| 4 | 94739809 | 94740309 | Tek | Intron 1 | 520 |
| 4 | 94739809 | 94740309 | Tek | Promoter | 509 |
| 4 | 94739809 | 94740309 | Tek | Intron 1 | 509 |
| 4 | 94739809 | 94740309 | Tek | Promoter | 291 |
| 4 | 94739809 | 94740309 | Tek | Intron 1 | 291 |
| 4 | 97900309 | 97900809 | Nfia | Intron 2 | 127575 |
| 4 | 97855809 | 97856309 | Nfia | Intron 2 | 83075 |

| | | | | | |
|---|---|---|---|---|---|
| 4 | 97900309 | 97900809 | Nfia | Intron 2 | 122701 |
| 4 | 97855809 | 97856309 | Nfia | Intron 2 | 78201 |
| 4 | 97900309 | 97900809 | Nfia | Intron 2 | 122683 |
| 4 | 97855809 | 97856309 | Nfia | Intron 2 | 78183 |
| 4 | 97900309 | 97900809 | Nfia | Intron 2 | 122506 |
| 4 | 97855809 | 97856309 | Nfia | Intron 2 | 78006 |
| 4 | 97900309 | 97900809 | Nfia | Intron 2 | 122490 |
| 4 | 97855809 | 97856309 | Nfia | Intron 2 | 77990 |
| 4 | 97900309 | 97900809 | Nfia | Intron 2 | 122475 |
| 4 | 97855809 | 97856309 | Nfia | Intron 2 | 77975 |
| 4 | 97900309 | 97900809 | Nfia | Intron 1 | 20190 |
| 4 | 100527895 | 100528395 | Ube2u | Intron 7 | 49046 |
| 4 | 100527895 | 100528395 | Ube2u | Intron 6 | 49028 |
| 4 | 100527895 | 100528395 | Ube2uos | Promoter | -93 |
| 4 | 101079395 | 101079895 | Raver2 | Intron 1 | 10412 |
| 4 | 101079395 | 101079895 | Raver2 | Intron 1 | 10412 |
| 4 | 101079395 | 101079895 | Raver2 | Non-coding | 9916 |
| 4 | 101196895 | 101197395 | Jak1 | 5' UTR | 67888 |
| 4 | 101196895 | 101197395 | Jak1 | Non-coding | 67887 |
| 4 | 101196895 | 101197395 | Jak1 | Non-coding | 67829 |
| 4 | 101196895 | 101197395 | Jak1 | 5' UTR | 67879 |
| 4 | 102965395 | 102965895 | Sgip1 | Intron 20 | 205260 |
| 4 | 102965395 | 102965895 | Sgip1 | Intron 19 | 205050 |
| 4 | 102965395 | 102965895 | Sgip1 | Intron 20 | 205032 |
| 4 | 102965395 | 102965895 | Sgip1 | Intron 21 | 204870 |
| 4 | 102965395 | 102965895 | Sgip1 | Intron 13 | 94831 |
| 4 | 102965395 | 102965895 | Sgip1 | Non-coding | 3039 |
| 4 | 106726895 | 106727395 | Mroh7 | Promoter | 536 |
| 4 | 106726895 | 106727395 | Mroh7 | 5' UTR | 536 |
| 4 | 106726895 | 106727395 | Mroh7 | Promoter | 582 |
| 4 | 106726895 | 106727395 | Mroh7 | 5' UTR | 582 |
| 4 | 106726895 | 106727395 | Mroh7 | 5' UTR | 3531 |
| 4 | 107442895 | 107443395 | Glis1 | Intron 2 | 8278 |
| 4 | 107442895 | 107443395 | Glis1 | Intron 2 | 8176 |
| 4 | 107442895 | 107443395 | Glis1 | 5' UTR | 3897 |
| 4 | 108890395 | 108890895 | Rab3b | CDS 2 | 11332 |
| 4 | 108890395 | 108890895 | Rab3b | Intron 2 | 11332 |
| 4 | 108890395 | 108890895 | Rab3b | CDS 2 | 11195 |
| 4 | 108890395 | 108890895 | Rab3b | Intron 2 | 11195 |
| 4 | 108890395 | 108890895 | Rab3b | Promoter | 105 |
| 4 | 108890395 | 108890895 | Rab3b | CDS 1 | 105 |
| 4 | 108890395 | 108890895 | Rab3b | Intron 1 | 105 |
| 4 | 113867395 | 113867895 | Skint5 | Intron 12 | 131609 |
| 4 | 113867395 | 113867895 | Skint5 | Intron 12 | 131609 |
| 4 | 113867395 | 113867895 | Skint5 | Intron 12 | 131609 |
| 4 | 114468395 | 114468895 | Trabd2b | Intron 2 | 61671 |

| | | | | | |
|---|---|---|---|---|---|
| 4 | 116299395 | 116299895 | Gm49337 | 3' UTR | 126011 |
| 4 | 116299395 | 116299895 | Gm49337 | Intron 9 | 126011 |
| 4 | 116299395 | 116299895 | Gm49337 | CDS 10 | 126011 |
| 4 | 116299395 | 116299895 | Pik3r3 | 3' UTR | 77734 |
| 4 | 116299395 | 116299895 | Pik3r3 | Intron 9 | 77734 |
| 4 | 116299395 | 116299895 | Pik3r3 | CDS 10 | 77734 |
| 4 | 116299395 | 116299895 | Pik3r3 | 3' UTR | 77699 |
| 4 | 116299395 | 116299895 | Pik3r3 | Intron 8 | 77699 |
| 4 | 116299395 | 116299895 | Pik3r3 | CDS 9 | 77699 |
| 4 | 116853895 | 116854395 | Gm12994 | Promoter | 1231 |
| 4 | 117977895 | 117978395 | St3gal3 | Intron 5 | 156496 |
| 4 | 117977895 | 117978395 | St3gal3 | Intron 5 | 156520 |
| 4 | 117977895 | 117978395 | St3gal3 | Intron 4 | 156488 |
| 4 | 117977895 | 117978395 | St3gal3 | Intron 2 | 53488 |
| 4 | 117977895 | 117978395 | 9530034E10Rik | Promoter | 2859 |
| 4 | 119399395 | 119399895 | Ccdc30 | Intron 3 | 15627 |
| 4 | 119399395 | 119399895 | Ccdc30 | Intron 3 | 15567 |
| 4 | 119399395 | 119399895 | Ccdc30 | Intron 3 | 15627 |
| 4 | 119399395 | 119399895 | Ccdc30 | Intron 3 | 15608 |
| 4 | 119399395 | 119399895 | Ccdc30 | Intron 3 | 15585 |
| 4 | 119399395 | 119399895 | Gm50484 | Non-coding | 5895 |
| 4 | 124904256 | 124904756 | Epha10 | Intron 5 | 18457 |
| 4 | 124899256 | 124899756 | Epha10 | Intron 3 | 13457 |
| 4 | 124899256 | 124899756 | 1700125G02Rik | Promoter | -4473 |
| 4 | 124899256 | 124899756 | Epha10 | Promoter | 0 |
| 4 | 125515756 | 125516256 | Grik3 | Intron 1 | 25056 |
| 4 | 126149256 | 126149756 | Eva1b | Promoter | 1512 |
| 4 | 126149256 | 126149756 | Eva1b | 3' UTR | 1512 |
| 4 | 126149256 | 126149756 | Eva1b | CDS 3 | 1512 |
| 4 | 126149256 | 126149756 | Eva1b | Promoter | 1253 |
| 4 | 126149256 | 126149756 | Eva1b | 3' UTR | 1253 |
| 4 | 126149256 | 126149756 | Eva1b | CDS 3 | 1253 |
| 4 | 126149256 | 126149756 | Eva1b | Promoter | 724 |
| 4 | 126149256 | 126149756 | Eva1b | 3' UTR | 724 |
| 4 | 126149256 | 126149756 | Eva1b | CDS 2 | 724 |
| 4 | 127984756 | 127985256 | Csmd2 | Promoter | -2601 |
| 4 | 128183256 | 128183756 | Csmd2 | Non-coding | 195399 |
| 4 | 127984756 | 127985256 | Csmd2 | Promoter | -2788 |
| 4 | 128183256 | 128183756 | Csmd2 | Non-coding | 195212 |
| 4 | 127984756 | 127985256 | Csmd2 | Promoter | -2788 |
| 4 | 128183256 | 128183756 | Csmd2 | Intron 3 | 195212 |
| 4 | 129630256 | 129630756 | Txlna | Intron 9 | 10310 |
| 4 | 129630256 | 129630756 | Txlna | CDS 9 | 10310 |
| 4 | 129630256 | 129630756 | Txlna | Intron 8 | 10310 |
| 4 | 129630256 | 129630756 | Txlna | Intron 8 | 10050 |

| | | | | | |
|---|---|---|---|---|---|
| 4 | 129630256 | 129630756 | Txlna | CDS 8 | 10050 |
| 4 | 129630256 | 129630756 | Txlna | Intron 7 | 10050 |
| 4 | 129630256 | 129630756 | Txlna | Non-coding | 9885 |
| 4 | 129949256 | 129949756 | Spocd1 | Intron 7 | 20007 |
| 4 | 129949256 | 129949756 | Spocd1 | Promoter | 1709 |
| 4 | 129949256 | 129949756 | Spocd1 | Intron 3 | 1709 |
| 4 | 131797585 | 131798085 | Ptpru | Intron 14 | 40204 |
| 4 | 131797585 | 131798085 | Ptpru | CDS 14 | 40204 |
| 4 | 131797585 | 131798085 | Ptpru | Intron 13 | 40204 |
| 4 | 131797585 | 131798085 | Ptpru | Intron 14 | 40147 |
| 4 | 131797585 | 131798085 | Ptpru | CDS 14 | 40147 |
| 4 | 131797585 | 131798085 | Ptpru | Intron 13 | 40147 |
| 4 | 131797585 | 131798085 | Ptpru | Non-coding | 40168 |
| 4 | 131797585 | 131798085 | Ptpru | Intron 13 | 23432 |
| 4 | 131797585 | 131798085 | Ptpru | CDS 13 | 23432 |
| 4 | 131797585 | 131798085 | Ptpru | Intron 12 | 23432 |
| 4 | 131797585 | 131798085 | Ptpru | Non-coding | 4912 |
| 4 | 131797585 | 131798085 | Ptpru | Promoter | 286 |
| 4 | 133615085 | 133615585 | Zdhhc18 | Intron 3 | 17885 |
| 4 | 133615085 | 133615585 | Zdhhc18 | CDS 3 | 17885 |
| 4 | 133615085 | 133615585 | Zdhhc18 | Intron 2 | 17885 |
| 4 | 133615085 | 133615585 | Zdhhc18 | Promoter | 0 |
| 4 | 133615085 | 133615585 | Zdhhc18 | Promoter | -2050 |
| 4 | 133615085 | 133615585 | Zdhhc18 | Promoter | 0 |
| 4 | 134060085 | 134060585 | Crybg2 | Promoter | -230 |
| 4 | 134060085 | 134060585 | Crybg2 | Promoter | -230 |
| 4 | 134166085 | 134166585 | Cep85 | Intron 3 | 20528 |
| 4 | 134166085 | 134166585 | Cep85 | Intron 3 | 20397 |
| 4 | 134166085 | 134166585 | Cep85 | Intron 3 | 20501 |
| 4 | 134166085 | 134166585 | Cep85 | Non-coding | 20501 |
| 4 | 134166085 | 134166585 | Gm27619 | Promoter | 1311 |
| 4 | 134636585 | 134637085 | Man1c1 | Intron 3 | 67206 |
| 4 | 134636585 | 134637085 | Man1c1 | Intron 2 | 67206 |
| 4 | 134636585 | 134637085 | Man1c1 | Non-coding | 66683 |
| 4 | 135265085 | 135265585 | Clic4 | Intron 1 | 7230 |
| 4 | 135265085 | 135265585 | Clic4 | Non-coding | 7230 |
| 4 | 136981585 | 136982085 | Zbtb40 | 3' UTR | 66611 |
| 4 | 137716585 | 137717085 | Rap1gap | Intron 10 | 35832 |
| 4 | 137716585 | 137717085 | Rap1gap | Intron 8 | 35832 |
| 4 | 137716585 | 137717085 | Rap1gap | Intron 10 | 34918 |
| 4 | 137716585 | 137717085 | Rap1gap | Intron 8 | 9063 |
| 4 | 137716585 | 137717085 | Rap1gap | Promoter | -1787 |
| 4 | 139217085 | 139217585 | Capzb | Intron 1 | 24186 |
| 4 | 139217085 | 139217585 | Capzb | Intron 1 | 24163 |
| 4 | 139217085 | 139217585 | Capzb | Intron 1 | 24161 |
| 4 | 139217085 | 139217585 | Capzb | Non-coding | 24146 |

| | | | | | |
|---|---|---|---|---|---|
| 4 | 139217085 | 139217585 | Capzb | Non-coding | 24116 |
| 4 | 139217085 | 139217585 | Capzb | Intron 1 | 23089 |
| 4 | 139217085 | 139217585 | Capzb | Non-coding | 22974 |
| 4 | 139217085 | 139217585 | Capzb | Intron 1 | 17818 |
| 4 | 139597085 | 139597585 | Iffo2 | 5' UTR | 66537 |
| 4 | 139597085 | 139597585 | Iffo2 | 5' UTR | 65396 |
| 4 | 139597085 | 139597585 | Iffo2 | Intron 1 | 22365 |
| 4 | 139795585 | 139796085 | Pax7 | Intron 4 | 37444 |
| 4 | 139795585 | 139796085 | Pax7 | Intron 4 | 36923 |
| 4 | 139905585 | 139906085 | Gm1667 | Promoter | -4403 |
| 4 | 140597585 | 140598085 | Arhgef10l | Intron 4 | 50666 |
| 4 | 140622585 | 140623085 | Arhgef10l | 5' UTR | 25666 |
| 4 | 140597585 | 140598085 | Arhgef10l | Intron 4 | 67928 |
| 4 | 140622585 | 140623085 | Arhgef10l | 5' UTR | 42928 |
| 4 | 140597585 | 140598085 | Arhgef10l | Intron 4 | 50672 |
| 4 | 140622585 | 140623085 | Arhgef10l | 5' UTR | 25672 |
| 4 | 140597585 | 140598085 | Arhgef10l | Intron 4 | 50666 |
| 4 | 140622585 | 140623085 | Arhgef10l | 5' UTR | 25666 |
| 4 | 140597585 | 140598085 | Arhgef10l | Intron 5 | 51276 |
| 4 | 140622585 | 140623085 | Arhgef10l | 5' UTR | 26276 |
| 4 | 140597585 | 140598085 | Arhgef10l | Intron 3 | 18981 |
| 4 | 140992085 | 140992585 | Atp13a2 | Intron 1 | 5212 |
| 4 | 140992085 | 140992585 | Atp13a2 | CDS 2 | 5212 |
| 4 | 140992085 | 140992585 | Atp13a2 | Intron 2 | 5212 |
| 4 | 140992085 | 140992585 | Atp13a2 | CDS 3 | 5212 |
| 4 | 140992085 | 140992585 | Atp13a2 | Intron 3 | 5212 |
| 4 | 140992085 | 140992585 | Atp13a2 | Intron 1 | 5212 |
| 4 | 140992085 | 140992585 | Atp13a2 | CDS 2 | 5212 |
| 4 | 140992085 | 140992585 | Atp13a2 | Intron 2 | 5212 |
| 4 | 140992085 | 140992585 | Atp13a2 | CDS 3 | 5212 |
| 4 | 140992085 | 140992585 | Atp13a2 | Intron 3 | 5212 |
| 4 | 140992085 | 140992585 | Atp13a2 | Intron 1 | 5212 |
| 4 | 140992085 | 140992585 | Atp13a2 | CDS 2 | 5212 |
| 4 | 140992085 | 140992585 | Atp13a2 | Intron 2 | 5212 |
| 4 | 140992085 | 140992585 | Atp13a2 | CDS 3 | 5212 |
| 4 | 140992085 | 140992585 | Atp13a2 | Intron 3 | 5212 |
| 4 | 140992085 | 140992585 | Atp13a2 | Non-coding | 5185 |
| 4 | 140992085 | 140992585 | Atp13a2 | Non-coding | 5095 |
| 4 | 140992085 | 140992585 | Atp13a2 | Promoter | -3017 |
| 4 | 140992085 | 140992585 | Atp13a2 | Promoter | -3627 |
| 4 | 145449097 | 145449597 | Platr16 | Non-coding | 38831 |
| 4 | 145449097 | 145449597 | Gm13230 | Promoter | 2357 |
| 4 | 148040891 | 148041391 | Clcn6 | Promoter | -2069 |
| 4 | 148040891 | 148041391 | Clcn6 | Promoter | -2077 |
| 4 | 148040891 | 148041391 | Clcn6 | Promoter | -2120 |
| 4 | 148040891 | 148041391 | Clcn6 | Promoter | -2127 |

| | | | | | |
|---|---|---|---|---|---|
| 4 | 148040891 | 148041391 | Clcn6 | Promoter | -2328 |
| 4 | 148040891 | 148041391 | Mthfr | Promoter | 1814 |
| 4 | 148040891 | 148041391 | Mthfr | 5' UTR | 1814 |
| 4 | 148040891 | 148041391 | Mthfr | Promoter | 1732 |
| 4 | 148040891 | 148041391 | Mthfr | 5' UTR | 1732 |
| 4 | 148040891 | 148041391 | Mthfr | Promoter | 81 |
| 4 | 148040891 | 148041391 | Mthfr | Promoter | 0 |
| 4 | 148040891 | 148041391 | Mthfr | 5' UTR | 0 |
| 4 | 148040891 | 148041391 | Mthfr | CDS 1 | 0 |
| 4 | 148040891 | 148041391 | Mthfr | Intron 1 | 0 |
| 4 | 148040891 | 148041391 | Mthfr | Promoter | 0 |
| 4 | 148040891 | 148041391 | Mthfr | 5' UTR | 0 |
| 4 | 148040891 | 148041391 | Mthfr | CDS 1 | 0 |
| 4 | 148040891 | 148041391 | Mthfr | Intron 1 | 0 |
| 4 | 148040891 | 148041391 | Mthfr | Promoter | -211 |
| 4 | 148661391 | 148661891 | Gm572 | Intron 5 | 18074 |
| 4 | 148844391 | 148844891 | Casz1 | 5' UTR | 39962 |
| 4 | 148844391 | 148844891 | Casz1 | 5' UTR | 39962 |
| 4 | 151103891 | 151104391 | Camta1 | Intron 11 | 757279 |
| 4 | 151560891 | 151561391 | Camta1 | Intron 4 | 300279 |
| 4 | 151670891 | 151671391 | Camta1 | Intron 3 | 190279 |
| 4 | 151170391 | 151170891 | Camta1 | Intron 6 | 690779 |
| 4 | 151103891 | 151104391 | Camta1 | Intron 11 | 757486 |
| 4 | 151560891 | 151561391 | Camta1 | Intron 4 | 300486 |
| 4 | 151670891 | 151671391 | Camta1 | Intron 3 | 190486 |
| 4 | 151170391 | 151170891 | Camta1 | Intron 6 | 690986 |
| 4 | 151103891 | 151104391 | Camta1 | Intron 11 | 757356 |
| 4 | 151560891 | 151561391 | Camta1 | Intron 4 | 300356 |
| 4 | 151670891 | 151671391 | Camta1 | Intron 3 | 190356 |
| 4 | 151170391 | 151170891 | Camta1 | Intron 6 | 690856 |
| 4 | 151103891 | 151104391 | Camta1 | 5' UTR | 25105 |
| 4 | 151103891 | 151104391 | Camta1 | 5' UTR | 4186 |
| 4 | 151103891 | 151104391 | Camta1 | Non-coding | 35194 |
| 4 | 151103891 | 151104391 | Camta1 | 5' UTR | 3913 |
| 4 | 151560891 | 151561391 | Camta1 | Non-coding | 300260 |
| 4 | 151670891 | 151671391 | Camta1 | Non-coding | 190260 |
| 4 | 152100891 | 152101391 | Plekhg5 | Intron 2 | 28393 |
| 4 | 152100891 | 152101391 | Plekhg5 | Intron 2 | 24508 |
| 4 | 152100891 | 152101391 | Plekhg5 | Intron 2 | 14033 |
| 4 | 152100891 | 152101391 | Plekhg5 | Intron 1 | 4172 |
| 4 | 152100891 | 152101391 | Plekhg5 | Intron 1 | 4172 |
| 4 | 152100891 | 152101391 | Plekhg5 | Promoter | -3109 |
| 4 | 152126391 | 152126891 | Espn | Intron 13 | 25317 |
| 4 | 152126391 | 152126891 | Espn | Intron 10 | 25481 |
| 4 | 152126391 | 152126891 | Espn | Promoter | 2035 |
| 4 | 152126391 | 152126891 | Espn | Intron 2 | 2035 |

| | | | | | |
|---|---|---|---|---|---|
| 4 | 152126391 | 152126891 | Espn | Intron 11 | 25332 |
| 4 | 152126391 | 152126891 | Espn | Intron 5 | 4975 |
| 4 | 152126391 | 152126891 | Espn | Non-coding | 5447 |
| 4 | 152126391 | 152126891 | Espn | Intron 7 | 9435 |
| 4 | 152126391 | 152126891 | Espn | Intron 7 | 9435 |
| 4 | 152126391 | 152126891 | Espn | Intron 8 | 9435 |
| 4 | 152126391 | 152126891 | Espn | Intron 6 | 9392 |
| 4 | 152126391 | 152126891 | Espn | Intron 7 | 9392 |
| 4 | 152126391 | 152126891 | Espn | Intron 7 | 9392 |
| 4 | 152126391 | 152126891 | Espn | Intron 8 | 9392 |
| 4 | 152126391 | 152126891 | Espn | Intron 6 | 9435 |
| 4 | 152126391 | 152126891 | Espn | Non-coding | 4951 |
| 4 | 152126391 | 152126891 | Espn | Non-coding | 4976 |
| 4 | 152126391 | 152126891 | Espn | Intron 10 | 22162 |
| 4 | 152126391 | 152126891 | Espn | CDS 12 | 25332 |
| 4 | 152126391 | 152126891 | Espn | Intron 11 | 25332 |
| 4 | 152311891 | 152312391 | Rnf207 | Intron 13 | 6218 |
| 4 | 152311891 | 152312391 | Rnf207 | CDS 13 | 6218 |
| 4 | 152311891 | 152312391 | Rnf207 | Intron 12 | 6218 |
| 4 | 152311891 | 152312391 | Rnf207 | CDS 12 | 6218 |
| 4 | 152311891 | 152312391 | Rnf207 | Promoter | -2857 |
| 4 | 152311891 | 152312391 | Rnf207 | Intron 12 | 6068 |
| 4 | 152311891 | 152312391 | Rnf207 | CDS 12 | 6068 |
| 4 | 152311891 | 152312391 | Rnf207 | Intron 11 | 6068 |
| 4 | 152311891 | 152312391 | Rnf207 | CDS 11 | 6068 |
| 4 | 152311891 | 152312391 | Rnf207 | Promoter | 1013 |
| 4 | 152311891 | 152312391 | Rnf207 | Promoter | 958 |
| 4 | 152311891 | 152312391 | Rnf207 | Promoter | 1010 |
| 4 | 152359391 | 152359891 | Chd5 | Intron 6 | 20740 |
| 4 | 152359391 | 152359891 | Chd5 | CDS 7 | 20740 |
| 4 | 152359391 | 152359891 | Chd5 | Intron 7 | 20740 |
| 4 | 152359391 | 152359891 | Chd5 | Intron 6 | 20740 |
| 4 | 152359391 | 152359891 | Chd5 | CDS 7 | 20740 |
| 4 | 152359391 | 152359891 | Chd5 | Intron 7 | 20740 |
| 4 | 152359391 | 152359891 | Chd5 | Intron 6 | 20504 |
| 4 | 152359391 | 152359891 | Chd5 | CDS 7 | 20504 |
| 4 | 152359391 | 152359891 | Chd5 | Intron 7 | 20504 |
| 4 | 152430391 | 152430891 | Kcnab2 | Intron 2 | 17950 |
| 4 | 152430391 | 152430891 | Kcnab2 | Intron 3 | 47020 |
| 4 | 152430391 | 152430891 | Kcnab2 | Intron 2 | 46699 |
| 4 | 152430391 | 152430891 | Kcnab2 | Intron 3 | 46549 |
| 4 | 152430391 | 152430891 | Kcnab2 | Promoter | 0 |
| 4 | 152430391 | 152430891 | Kcnab2 | 5' UTR | 0 |
| 4 | 152430391 | 152430891 | Kcnab2 | Intron 3 | 17966 |
| 4 | 152430391 | 152430891 | Kcnab2 | Promoter | 0 |
| 4 | 152430391 | 152430891 | Kcnab2 | 5' UTR | 0 |

| | | | | | |
|---|---|---|---|---|---|
| 4 | 152430391 | 152430891 | Kcnab2 | Intron 3 | 24153 |
| 4 | 152430391 | 152430891 | Kcnab2 | Promoter | 0 |
| 4 | 152430391 | 152430891 | Kcnab2 | 5' UTR | 0 |
| 4 | 152430391 | 152430891 | Kcnab2 | 5' UTR | 17851 |
| 4 | 152430391 | 152430891 | Kcnab2 | Non-coding | 46629 |
| 4 | 152430391 | 152430891 | Kcnab2 | Non-coding | 46654 |
| 4 | 152430391 | 152430891 | Kcnab2 | Non-coding | 17851 |
| 4 | 152554391 | 152554891 | Nphp4 | Non-coding | 77685 |
| 4 | 152554391 | 152554891 | Nphp4 | Intron 19 | 76249 |
| 4 | 152554391 | 152554891 | Nphp4 | CDS 20 | 76249 |
| 4 | 152554391 | 152554891 | Nphp4 | Intron 20 | 76249 |
| 4 | 152554391 | 152554891 | Nphp4 | Intron 18 | 70021 |
| 4 | 152554391 | 152554891 | Nphp4 | CDS 19 | 70021 |
| 4 | 152554391 | 152554891 | Nphp4 | Intron 19 | 70021 |
| 4 | 154041891 | 154042391 | Ccdc27 | Promoter | 287 |
| 4 | 154041891 | 154042391 | Ccdc27 | Intron 1 | 287 |
| 4 | 154041891 | 154042391 | Gm13117 | Promoter | 2033 |
| 4 | 155859891 | 155860391 | Tas1r3 | Promoter | 2972 |
| 4 | 155859891 | 155860391 | Tas1r3 | 3' UTR | 2972 |
| 4 | 155859891 | 155860391 | Tas1r3 | CDS 6 | 2972 |
| 5 | 16846682 | 16847182 | Gm28710 | Intron 16 | 54872 |
| 5 | 24693682 | 24694182 | Nub1 | Intron 3 | 8150 |
| 5 | 24693682 | 24694182 | Nub1 | Intron 3 | 7858 |
| 5 | 24693682 | 24694182 | Nub1 | Non-coding | 7695 |
| 5 | 25848182 | 25848682 | Actr3b | Intron 8 | 88185 |
| 5 | 25848182 | 25848682 | Actr3b | CDS 9 | 88185 |
| 5 | 25848182 | 25848682 | Actr3b | Intron 9 | 88185 |
| 5 | 25848182 | 25848682 | Actr3b | Intron 9 | 88156 |
| 5 | 25848182 | 25848682 | Actr3b | CDS 10 | 88156 |
| 5 | 25848182 | 25848682 | Actr3b | Intron 10 | 88156 |
| 5 | 27160960 | 27161460 | Dpp6 | Intron 1 | 343603 |
| 5 | 27160960 | 27161460 | Dpp6 | Non-coding | 343526 |
| 5 | 27160960 | 27161460 | Dpp6 | Intron 1 | 256265 |
| 5 | 27160960 | 27161460 | Dpp6 | Intron 1 | 111761 |
| 5 | 28160460 | 28160960 | En2 | Promoter | -4734 |
| 5 | 35831351 | 35831851 | Ablim2 | Intron 8 | 73471 |
| 5 | 35831351 | 35831851 | Ablim2 | Intron 8 | 73471 |
| 5 | 35831351 | 35831851 | Ablim2 | CDS 9 | 73471 |
| 5 | 35831351 | 35831851 | Ablim2 | Intron 9 | 73471 |
| 5 | 35831351 | 35831851 | Ablim2 | Intron 8 | 73427 |
| 5 | 35831351 | 35831851 | Ablim2 | Intron 8 | 73420 |
| 5 | 35831351 | 35831851 | Ablim2 | Intron 8 | 73420 |
| 5 | 35831351 | 35831851 | Ablim2 | Intron 8 | 73385 |
| 5 | 35831351 | 35831851 | Ablim2 | Intron 8 | 73300 |
| 5 | 35831351 | 35831851 | Ablim2 | CDS 9 | 73300 |
| 5 | 35831351 | 35831851 | Ablim2 | Intron 9 | 73300 |

| | | | | | |
|---|---|---|---|---|---|
| 5 | 35831351 | 35831851 | Ablim2 | Intron 3 | 16992 |
| 5 | 35831351 | 35831851 | Ablim2 | CDS 4 | 16992 |
| 5 | 35831351 | 35831851 | Ablim2 | Intron 4 | 16992 |
| 5 | 35831351 | 35831851 | Ablim2 | Intron 3 | 16925 |
| 5 | 35831351 | 35831851 | Ablim2 | 3' UTR | 16911 |
| 5 | 35831351 | 35831851 | Ablim2 | Intron 1 | 3227 |
| 5 | 35831351 | 35831851 | Ablim2 | Non-coding | 3196 |
| 5 | 35831351 | 35831851 | Ablim2 | Intron 1 | 3196 |
| 5 | 37551761 | 37552261 | Stk32b | Intron 3 | 164911 |
| 5 | 38315261 | 38315761 | Drd5 | Promoter | -3606 |
| 5 | 52360761 | 52361261 | Sod3 | Promoter | -2530 |
| 5 | 52996761 | 52997261 | 5033403H07Rik | Promoter | -2324 |
| 5 | 52996761 | 52997261 | 5033403H07Rik | Promoter | -4468 |
| 5 | 52996761 | 52997261 | 5033403H07Rik | Promoter | -4417 |
| 5 | 55722761 | 55723261 | Gm20223 | Promoter | 161 |
| 5 | 66527261 | 66527761 | Apbb2 | 5' UTR | 90998 |
| 5 | 66527261 | 66527761 | Apbb2 | 5' UTR | 91024 |
| 5 | 66527261 | 66527761 | Apbb2 | 5' UTR | 90921 |
| 5 | 66527261 | 66527761 | Apbb2 | 5' UTR | 90903 |
| 5 | 66527261 | 66527761 | Apbb2 | 5' UTR | 90911 |
| 5 | 66527261 | 66527761 | Apbb2 | 5' UTR | 90911 |
| 5 | 66527261 | 66527761 | Apbb2 | Non-coding | 90923 |
| 5 | 66527261 | 66527761 | Apbb2 | Non-coding | 90910 |
| 5 | 66527261 | 66527761 | Apbb2 | 5' UTR | 90880 |
| 5 | 66527261 | 66527761 | Apbb2 | 5' UTR | 90911 |
| 5 | 66527261 | 66527761 | Apbb2 | 5' UTR | 91014 |
| 5 | 66527261 | 66527761 | Apbb2 | 5' UTR | 19197 |
| 5 | 66527261 | 66527761 | Apbb2 | Non-coding | 90912 |
| 5 | 66527261 | 66527761 | Gm43343 | Promoter | -1148 |
| 5 | 66650761 | 66651261 | Uchl1os | Non-coding | 25238 |
| 5 | 73014261 | 73014761 | Gm15847 | Promoter | 733 |
| 5 | 73014261 | 73014761 | Gm42668 | Promoter | -438 |
| 5 | 73153761 | 73154261 | Fryl | 5' UTR | 102359 |
| 5 | 73153761 | 73154261 | Fryl | Non-coding | 14180 |
| 5 | 73153761 | 73154261 | Fryl | Non-coding | 102165 |
| 5 | 73153761 | 73154261 | Fryl | Intron 1 | 37619 |
| 5 | 73153761 | 73154261 | 1700071G01Rik | Promoter | -2753 |
| 5 | 73627761 | 73628261 | Lrrc66 | Intron 2 | 4266 |
| 5 | 73627761 | 73628261 | Lrrc66 | Promoter | 2052 |
| 5 | 77468975 | 77469475 | Gm34728 | Non-coding | 34941 |
| 5 | 86664975 | 86665475 | Tmprss11b | CDS 5 | 10888 |
| 5 | 86664975 | 86665475 | Tmprss11b | Intron 4 | 10888 |

| | | | | | |
|---|---|---|---|---|---|
| 5 | 92320974 | 92321474 | Cxcl9 | 3' UTR | 6606 |
| 5 | 92402974 | 92403474 | Art3 | Intron 5 | 71133 |
| 5 | 92402974 | 92403474 | Art3 | Intron 5 | 71133 |
| 5 | 92402974 | 92403474 | Art3 | Intron 3 | 14831 |
| 5 | 92402974 | 92403474 | Art3 | Intron 3 | 14831 |
| 5 | 92402974 | 92403474 | Art3 | Intron 3 | 14831 |
| 5 | 92402974 | 92403474 | Art3 | Intron 3 | 14831 |
| 5 | 92402974 | 92403474 | Art3 | Intron 3 | 14831 |
| 5 | 92402974 | 92403474 | Art3 | Non-coding | 14831 |
| 5 | 92402974 | 92403474 | Art3 | Intron 2 | 10389 |
| 5 | 92402974 | 92403474 | Art3 | Intron 2 | 9915 |
| 5 | 92402974 | 92403474 | Art3 | Promoter | 976 |
| 5 | 92402974 | 92403474 | Art3 | Intron 1 | 976 |
| 5 | 92930974 | 92931474 | Shroom3 | Intron 3 | 247539 |
| 5 | 92930974 | 92931474 | Shroom3 | Intron 3 | 247349 |
| 5 | 92930974 | 92931474 | Shroom3 | 5' UTR | 121596 |
| 5 | 92930974 | 92931474 | Shroom3 | Intron 2 | 74503 |
| 5 | 92930974 | 92931474 | Shroom3 | 5' UTR | 32981 |
| 5 | 92930974 | 92931474 | Shroom3 | 5' UTR | 5567 |
| 5 | 93098474 | 93098974 | 11-Sep | Intron 1 | 5037 |
| 5 | 93098474 | 93098974 | 11-Sep | Intron 1 | 5036 |
| 5 | 93098474 | 93098974 | 11-Sep | 5' UTR | 5026 |
| 5 | 93098474 | 93098974 | 11-Sep | Intron 1 | 5023 |
| 5 | 93098474 | 93098974 | 11-Sep | Intron 1 | 4948 |
| 5 | 93098474 | 93098974 | 11-Sep | 5' UTR | 4938 |
| 5 | 93098474 | 93098974 | 11-Sep | Intron 1 | 4903 |
| 5 | 100058481 | 100058981 | Enoph1 | Intron 1 | 18496 |
| 5 | 100058481 | 100058981 | Enoph1 | Intron 1 | 18437 |
| 5 | 100058481 | 100058981 | Enoph1 | Non-coding | 18413 |
| 5 | 100058481 | 100058981 | Gm26363 | Promoter | -1489 |
| 5 | 100058481 | 100058981 | Enoph1 | Promoter | -2990 |
| 5 | 103423981 | 103424481 | Ptpn13 | Promoter | -711 |
| 5 | 103423981 | 103424481 | Ptpn13 | Promoter | -759 |
| 5 | 103423981 | 103424481 | Ptpn13 | Promoter | -775 |
| 5 | 103423981 | 103424481 | Ptpn13 | Promoter | -1020 |
| 5 | 104824981 | 104825481 | Zfp951 | Intron 1 | 34590 |
| 5 | 104824981 | 104825481 | Zfp951 | Intron 1 | 3477 |
| 5 | 106580481 | 106580981 | Gm28039 | Non-coding | 115591 |
| 5 | 110387481 | 110387981 | Fbrsl1 | Intron 5 | 60523 |
| 5 | 110387481 | 110387981 | Fbrsl1 | Intron 5 | 60622 |
| 5 | 110387481 | 110387981 | Fbrsl1 | Promoter | -384 |
| 5 | 110387481 | 110387981 | Fbrsl1 | Intron 5 | 60506 |
| 5 | 110387481 | 110387981 | Fbrsl1 | Promoter | -518 |
| 5 | 111227481 | 111227981 | Ttc28 | Intron 9 | 347678 |
| 5 | 111227481 | 111227981 | Ttc28 | Intron 8 | 347678 |
| 5 | 111227481 | 111227981 | Ttc28 | Promoter | 2023 |

| | | | | | |
|---|---|---|---|---|---|
| 5 | 111442480 | 111442980 | Mn1 | Intron 1 | 25038 |
| 5 | 112542480 | 112542980 | Sez6l | Intron 1 | 33996 |
| 5 | 112542480 | 112542980 | Sez6l | Intron 1 | 33996 |
| 5 | 112542480 | 112542980 | Sez6l | Intron 1 | 34206 |
| 5 | 112542480 | 112542980 | Sez6l | Intron 1 | 34198 |
| 5 | 112542480 | 112542980 | Sez6l | Intron 1 | 33889 |
| 5 | 112950480 | 112950980 | Grk3 | Intron 9 | 64535 |
| 5 | 112950480 | 112950980 | Grk3 | 3' UTR | 64535 |
| 5 | 112950480 | 112950980 | Grk3 | Intron 9 | 64812 |
| 5 | 113075980 | 113076480 | Crybb3 | CDS 5 | 4188 |
| 5 | 113075980 | 113076480 | Crybb3 | Intron 4 | 4188 |
| 5 | 113075980 | 113076480 | Crybb3 | CDS 6 | 4631 |
| 5 | 113075980 | 113076480 | Crybb3 | Intron 5 | 4631 |
| 5 | 113075980 | 113076480 | Crybb3 | CDS 6 | 5105 |
| 5 | 113075980 | 113076480 | Crybb3 | Intron 5 | 5105 |
| 5 | 113075980 | 113076480 | Crybb3 | CDS 6 | 4631 |
| 5 | 113075980 | 113076480 | Crybb3 | Intron 5 | 4631 |
| 5 | 113075980 | 113076480 | Crybb3 | CDS 6 | 5104 |
| 5 | 113075980 | 113076480 | Crybb3 | Intron 5 | 5104 |
| 5 | 114040991 | 114041491 | Svop | Intron 10 | 50080 |
| 5 | 114040991 | 114041491 | Svop | Non-coding | 23728 |
| 5 | 114040991 | 114041491 | Svop | Promoter | 1391 |
| 5 | 114105991 | 114106491 | Usp30 | Intron 4 | 5676 |
| 5 | 114105991 | 114106491 | Usp30 | Intron 4 | 5631 |
| 5 | 114105991 | 114106491 | Usp30 | Promoter | 2279 |
| 5 | 114105991 | 114106491 | Gm17833 | Promoter | -2435 |
| 5 | 114105991 | 114106491 | Usp30 | Promoter | -4637 |
| 5 | 115172991 | 115173491 | Cabp1 | Intron 5 | 12818 |
| 5 | 115172991 | 115173491 | Cabp1 | CDS 5 | 12818 |
| 5 | 115172991 | 115173491 | Cabp1 | Intron 4 | 12818 |
| 5 | 115172991 | 115173491 | Cabp1 | Intron 4 | 20891 |
| 5 | 115172991 | 115173491 | Cabp1 | CDS 4 | 20891 |
| 5 | 115172991 | 115173491 | Cabp1 | Intron 3 | 20891 |
| 5 | 115172991 | 115173491 | Cabp1 | Non-coding | 7932 |
| 5 | 115172991 | 115173491 | Cabp1 | Intron 4 | 12631 |
| 5 | 115172991 | 115173491 | Cabp1 | CDS 4 | 12631 |
| 5 | 115172991 | 115173491 | Cabp1 | Intron 3 | 12631 |
| 5 | 115172991 | 115173491 | Cabp1 | 3' UTR | 12566 |
| 5 | 115172991 | 115173491 | Cabp1 | Non-coding | 7769 |
| 5 | 115541991 | 115542491 | Pxn | Intron 1 | 35315 |
| 5 | 115541991 | 115542491 | Pxn | Non-coding | 35280 |
| 5 | 115541991 | 115542491 | Pxn | Intron 1 | 35183 |
| 5 | 115541991 | 115542491 | Pxn | Intron 1 | 35135 |
| 5 | 115541991 | 115542491 | Pxn | Non-coding | 25636 |
| 5 | 115541991 | 115542491 | Pxn | Non-coding | 21686 |
| 5 | 115541991 | 115542491 | Pxn | Promoter | -423 |

| | | | | | |
|---|---|---|---|---|---|
| 5 | 115541991 | 115542491 | Pxn | Promoter | -459 |
| 5 | 115541991 | 115542491 | Pxn | Promoter | -1456 |
| 5 | 115541991 | 115542491 | Pxn | Promoter | -1594 |
| 5 | 115541991 | 115542491 | Pxn | Promoter | -1623 |
| 5 | 115541991 | 115542491 | Pxn | Promoter | -2305 |
| 5 | 115541991 | 115542491 | Pxn | Promoter | -2813 |
| 5 | 116194491 | 116194991 | Ccdc60 | Intron 2 | 93995 |
| 5 | 116158491 | 116158991 | Ccdc60 | Intron 6 | 129995 |
| 5 | 116194491 | 116194991 | Ccdc60 | Intron 2 | 93973 |
| 5 | 116317491 | 116317991 | B230112J18Rik | Non-coding | 5131 |
| 5 | 116491991 | 116492491 | Srrm4 | Intron 1 | 99327 |
| 5 | 118590991 | 118591491 | Med13l | Non-coding | 30312 |
| 5 | 118590991 | 118591491 | Med13l | Intron 1 | 30272 |
| 5 | 118590991 | 118591491 | Med13l | Intron 1 | 30252 |
| 5 | 118590991 | 118591491 | Med13l | Non-coding | 29416 |
| 5 | 120973491 | 120973991 | Rph3a | Intron 4 | 35548 |
| 5 | 120967491 | 120967991 | Rph3a | Intron 5 | 41548 |
| 5 | 120973491 | 120973991 | Rph3a | Intron 4 | 36090 |
| 5 | 120967491 | 120967991 | Rph3a | Intron 5 | 42090 |
| 5 | 120973491 | 120973991 | Rph3a | Intron 3 | 34552 |
| 5 | 120967491 | 120967991 | Rph3a | Intron 4 | 40552 |
| 5 | 120973491 | 120973991 | Gm42656 | Promoter | -1143 |
| 5 | 120973491 | 120973991 | Rph3a | Intron 5 | 36102 |
| 5 | 122038991 | 122039491 | Cux2 | 5' UTR | 10361 |
| 5 | 121950991 | 121951491 | Cux2 | 5' UTR | 98361 |
| 5 | 122038991 | 122039491 | Cux2 | 5' UTR | 8335 |
| 5 | 121950991 | 121951491 | Cux2 | 5' UTR | 96335 |
| 5 | 122038991 | 122039491 | Cux2 | 5' UTR | 10612 |
| 5 | 121950991 | 121951491 | Cux2 | 5' UTR | 98612 |
| 5 | 121950991 | 121951491 | Cux2 | 5' UTR | 50854 |
| 5 | 122038991 | 122039491 | Cux2 | Non-coding | 8401 |
| 5 | 121950991 | 121951491 | Cux2 | Non-coding | 96401 |
| 5 | 122453991 | 122454491 | Atp2a2 | 3' UTR | 47684 |
| 5 | 122453991 | 122454491 | Atp2a2 | 3' UTR | 47405 |
| 5 | 122453991 | 122454491 | Atp2a2 | CDS 21 | 47405 |
| 5 | 122453991 | 122454491 | Atp2a2 | Intron 20 | 47405 |
| 5 | 123657991 | 123658491 | Clip1 | 5' UTR | 25832 |
| 5 | 123657991 | 123658491 | Clip1 | 5' UTR | 26128 |
| 5 | 123657991 | 123658491 | Clip1 | 5' UTR | 25785 |
| 5 | 123657991 | 123658491 | Clip1 | 5' UTR | 25835 |
| 5 | 123657991 | 123658491 | Clip1 | Promoter | -1972 |
| 5 | 123657991 | 123658491 | Clip1 | Promoter | -2082 |
| 5 | 123657991 | 123658491 | Clip1 | 5' UTR | 8200 |
| 5 | 125156630 | 125157130 | Ncor2 | 5' UTR | 22090 |
| 5 | 125102630 | 125103130 | Ncor2 | Intron 8 | 76090 |
| 5 | 125102630 | 125103130 | Ncor2 | CDS 8 | 76090 |

| | | | | | |
|---|---|---|---|---|---|
| 5 | 125102630 | 125103130 | Ncor2 | Intron 7 | 76090 |
| 5 | 125156630 | 125157130 | Ncor2 | 5' UTR | 22085 |
| 5 | 125102630 | 125103130 | Ncor2 | Intron 7 | 76085 |
| 5 | 125102630 | 125103130 | Ncor2 | CDS 7 | 76085 |
| 5 | 125102630 | 125103130 | Ncor2 | Intron 6 | 76085 |
| 5 | 125156630 | 125157130 | Ncor2 | 5' UTR | 22051 |
| 5 | 125102630 | 125103130 | Ncor2 | Intron 7 | 76051 |
| 5 | 125102630 | 125103130 | Ncor2 | CDS 7 | 76051 |
| 5 | 125102630 | 125103130 | Ncor2 | Intron 6 | 76051 |
| 5 | 125156630 | 125157130 | Ncor2 | 5' UTR | 21924 |
| 5 | 125102630 | 125103130 | Ncor2 | Intron 7 | 75924 |
| 5 | 125102630 | 125103130 | Ncor2 | CDS 7 | 75924 |
| 5 | 125102630 | 125103130 | Ncor2 | Intron 6 | 75924 |
| 5 | 125102630 | 125103130 | Ncor2 | Intron 6 | 21771 |
| 5 | 125102630 | 125103130 | Ncor2 | CDS 6 | 21771 |
| 5 | 125102630 | 125103130 | Ncor2 | Intron 5 | 21771 |
| 5 | 125156630 | 125157130 | Ncor2 | 5' UTR | 22073 |
| 5 | 125102630 | 125103130 | Ncor2 | Intron 3 | 76073 |
| 5 | 125438130 | 125438630 | Dhx37 | Promoter | -4008 |
| 5 | 125438130 | 125438630 | Bri3bp | Promoter | -2938 |
| 5 | 125438130 | 125438630 | Bri3bp | Promoter | -3120 |
| 5 | 128206630 | 128207130 | Tmem132d | Intron 2 | 225948 |
| 5 | 128204630 | 128205130 | Tmem132d | Intron 2 | 227948 |
| 5 | 129541625 | 129542125 | Sfswap | Intron 9 | 40404 |
| 5 | 129541625 | 129542125 | Sfswap | CDS 10 | 40404 |
| 5 | 129541625 | 129542125 | Sfswap | 3' UTR | 40385 |
| 5 | 129541625 | 129542125 | Sfswap | Non-coding | 40374 |
| 5 | 129541625 | 129542125 | Sfswap | Promoter | 209 |
| 5 | 129541625 | 129542125 | Sfswap | Intron 1 | 209 |
| 5 | 129541625 | 129542125 | Sfswap | CDS 2 | 209 |
| 5 | 129541625 | 129542125 | Sfswap | Intron 2 | 209 |
| 5 | 134121130 | 134121630 | Castor2 | Intron 1 | 21419 |
| 5 | 134121130 | 134121630 | Castor2 | Non-coding | 21125 |
| 5 | 134544630 | 134545130 | Clip2 | 5' UTR | 7305 |
| 5 | 134544630 | 134545130 | Clip2 | 5' UTR | 7305 |
| 5 | 136197630 | 136198130 | Prkrip1 | Promoter | 834 |
| 5 | 136197630 | 136198130 | Prkrip1 | Intron 2 | 834 |
| 5 | 136197630 | 136198130 | Prkrip1 | CDS 2 | 834 |
| 5 | 136197630 | 136198130 | Prkrip1 | Intron 1 | 834 |
| 5 | 136197630 | 136198130 | Prkrip1 | Promoter | 781 |
| 5 | 136197630 | 136198130 | Prkrip1 | Promoter | 0 |
| 5 | 136197630 | 136198130 | Prkrip1 | Intron 1 | 0 |
| 5 | 136197630 | 136198130 | Prkrip1 | CDS 1 | 0 |
| 5 | 136197630 | 136198130 | Prkrip1 | Promoter | 779 |
| 5 | 136197630 | 136198130 | Prkrip1 | Intron 2 | 779 |
| 5 | 136197630 | 136198130 | Prkrip1 | CDS 2 | 779 |

| | | | | | |
|---|---|---|---|---|---|
| 5 | 136197630 | 136198130 | Prkrip1 | Intron 1 | 779 |
| 5 | 136746130 | 136746630 | Col26a1 | Intron 11 | 136492 |
| 5 | 136746130 | 136746630 | Col26a1 | Intron 11 | 136555 |
| 5 | 136746130 | 136746630 | Col26a1 | Promoter | -1580 |
| 5 | 136746130 | 136746630 | Col26a1 | Promoter | 1072 |
| 5 | 137366772 | 137367272 | Ephb4 | CDS 12 | 16663 |
| 5 | 137366772 | 137367272 | Ephb4 | Intron 12 | 16663 |
| 5 | 137354272 | 137354772 | Ephb4 | Intron 2 | 4163 |
| 5 | 137354272 | 137354772 | Ephb4 | CDS 3 | 4163 |
| 5 | 137354272 | 137354772 | Ephb4 | Intron 3 | 4163 |
| 5 | 137366772 | 137367272 | Ephb4 | CDS 12 | 16631 |
| 5 | 137366772 | 137367272 | Ephb4 | Intron 12 | 16631 |
| 5 | 137354272 | 137354772 | Ephb4 | Intron 2 | 4131 |
| 5 | 137354272 | 137354772 | Ephb4 | CDS 3 | 4131 |
| 5 | 137354272 | 137354772 | Ephb4 | Intron 3 | 4131 |
| 5 | 137366772 | 137367272 | Ephb4 | CDS 12 | 16600 |
| 5 | 137366772 | 137367272 | Ephb4 | Intron 12 | 16600 |
| 5 | 137354272 | 137354772 | Ephb4 | Intron 2 | 4100 |
| 5 | 137354272 | 137354772 | Ephb4 | CDS 3 | 4100 |
| 5 | 137354272 | 137354772 | Ephb4 | Intron 3 | 4100 |
| 5 | 137366772 | 137367272 | Ephb4 | CDS 13 | 16342 |
| 5 | 137366772 | 137367272 | Ephb4 | Intron 13 | 16342 |
| 5 | 137354272 | 137354772 | Ephb4 | Intron 3 | 3842 |
| 5 | 137354272 | 137354772 | Ephb4 | CDS 4 | 3842 |
| 5 | 137354272 | 137354772 | Ephb4 | Intron 4 | 3842 |
| 5 | 137366772 | 137367272 | Ephb4 | CDS 12 | 16259 |
| 5 | 137366772 | 137367272 | Ephb4 | Intron 12 | 16259 |
| 5 | 137354272 | 137354772 | Ephb4 | Intron 2 | 3759 |
| 5 | 137354272 | 137354772 | Ephb4 | CDS 3 | 3759 |
| 5 | 137354272 | 137354772 | Ephb4 | Intron 3 | 3759 |
| 5 | 139377546 | 139378046 | 3110082I17Rik | Intron 2 | 82457 |
| 5 | 139377546 | 139378046 | 3110082I17Rik | Intron 2 | 82482 |
| 5 | 139377546 | 139378046 | 3110082I17Rik | Non-coding | 82466 |
| 5 | 139377546 | 139378046 | Gpr146 | Promoter | 0 |
| 5 | 139377546 | 139378046 | Gpr146 | Promoter | 0 |
| 5 | 139377546 | 139378046 | Gpr146 | Promoter | -117 |
| 5 | 139377546 | 139378046 | Gpr146 | Promoter | -174 |
| 5 | 139377546 | 139378046 | Gpr146 | Promoter | -2535 |
| 5 | 139718046 | 139718546 | Micall2 | Intron 4 | 17791 |
| 5 | 139718046 | 139718546 | Micall2 | CDS 4 | 17791 |
| 5 | 139718046 | 139718546 | Micall2 | Intron 3 | 17791 |
| 5 | 139718046 | 139718546 | Micall2 | Intron 4 | 14289 |
| 5 | 139718046 | 139718546 | Micall2 | CDS 4 | 14289 |
| 5 | 139718046 | 139718546 | Micall2 | Intron 3 | 14289 |
| 5 | 139822546 | 139823046 | Gm26938 | Intron 1 | 3362 |
| 5 | 139822546 | 139823046 | Tmem184a | Promoter | -2628 |

| | | | | | |
|---|---|---|---|---|---|
| 5 | 141583546 | 141584046 | Sdk1 | Intron 2 | 342056 |
| 5 | 142139546 | 142140046 | Sdk1 | Intron 32 | 898056 |
| 5 | 141583546 | 141584046 | Sdk1 | Non-coding | 209892 |
| 5 | 142139546 | 142140046 | Sdk1 | Intron 28 | 282759 |
| 5 | 143764131 | 143764631 | D130017N08Rik | Non-coding | 5777 |
| 5 | 143764131 | 143764631 | D130017N08Rik | Non-coding | 4269 |
| 5 | 143764131 | 143764631 | D130017N08Rik | Promoter | 27 |
| 5 | 143764131 | 143764631 | D130017N08Rik | Promoter | -1680 |
| 5 | 143933131 | 143933631 | Pms2 | Non-coding | 23167 |
| 5 | 143933131 | 143933631 | Pms2 | 3' UTR | 23167 |
| 5 | 143933131 | 143933631 | Pms2 | Non-coding | 23167 |
| 5 | 143933131 | 143933631 | Gm17135 | Non-coding | 47361 |
| 5 | 143933131 | 143933631 | Gm42421 | Intron 3 | 4956 |
| 5 | 143933131 | 143933631 | Rsph10b | Promoter | 96 |
| 5 | 143933131 | 143933631 | Rsph10b | Promoter | 69 |
| 5 | 143933131 | 143933631 | Rsph10b | 5' UTR | 69 |
| 5 | 143933131 | 143933631 | Rsph10b | Promoter | -3452 |
| 5 | 144787631 | 144788131 | Trrap | Non-coding | 19539 |
| 5 | 144787631 | 144788131 | Trrap | Intron 12 | 18837 |
| 5 | 144787631 | 144788131 | Trrap | CDS 13 | 18837 |
| 5 | 144787631 | 144788131 | Trrap | Intron 13 | 18837 |
| 5 | 144787631 | 144788131 | Trrap | Intron 11 | 16542 |
| 5 | 144787631 | 144788131 | Trrap | CDS 12 | 16542 |
| 5 | 144787631 | 144788131 | Trrap | Intron 12 | 16542 |
| 5 | 144787631 | 144788131 | Trrap | Intron 11 | 16542 |
| 5 | 144787631 | 144788131 | Trrap | CDS 12 | 16542 |
| 5 | 144787631 | 144788131 | Trrap | Intron 12 | 16542 |
| 5 | 144787631 | 144788131 | Trrap | Intron 11 | 16481 |
| 5 | 144787631 | 144788131 | Trrap | CDS 12 | 16481 |
| 5 | 144787631 | 144788131 | Trrap | Intron 12 | 16481 |
| 5 | 144787631 | 144788131 | Trrap | Promoter | 2106 |
| 5 | 144787631 | 144788131 | Trrap | Intron 2 | 2106 |
| 5 | 144787631 | 144788131 | Trrap | CDS 3 | 2106 |
| 5 | 144787631 | 144788131 | Trrap | Intron 3 | 2106 |
| 5 | 144787631 | 144788131 | Trrap | Promoter | -3328 |
| 5 | 145074631 | 145075131 | Kpna7 | 5' UTR | 8900 |
| 5 | 147354424 | 147354924 | Flt3 | Intron 15 | 45566 |
| 5 | 147354424 | 147354924 | Flt3 | CDS 15 | 45566 |
| 5 | 147354424 | 147354924 | Flt3 | Intron 14 | 45566 |
| 5 | 147354424 | 147354924 | Flt3 | CDS 14 | 45566 |
| 5 | 147716924 | 147717424 | Flt1 | Intron 1 | 8588 |
| 5 | 147716924 | 147717424 | Flt1 | Intron 1 | 8565 |

| | | | | | |
|---|---|---|---|---|---|
| 5 | 147716924 | 147717424 | Flt1 | Non-coding | 8565 |
| 5 | 148015424 | 148015924 | Mtus2 | 5' UTR | 58104 |
| 5 | 148072924 | 148073424 | Mtus2 | 5' UTR | 115604 |
| 5 | 148015424 | 148015924 | Mtus2 | 5' UTR | 58082 |
| 5 | 148072924 | 148073424 | Mtus2 | 5' UTR | 115582 |
| 6 | 5413000 | 5413500 | Asb4 | Intron 3 | 29614 |
| 6 | 7675000 | 7675500 | Asns | 3' UTR | 17755 |
| 6 | 7675000 | 7675500 | Asns | CDS 12 | 17755 |
| 6 | 7675000 | 7675500 | Asns | 3' UTR | 17372 |
| 6 | 7675000 | 7675500 | Asns | CDS 12 | 17372 |
| 6 | 7675000 | 7675500 | Asns | Promoter | 1886 |
| 6 | 11998500 | 11999000 | Phf14 | Intron 14 | 72619 |
| 6 | 11998500 | 11999000 | Phf14 | Intron 14 | 72585 |
| 6 | 11998500 | 11999000 | Phf14 | Intron 14 | 72584 |
| 6 | 22205500 | 22206000 | Cped1 | Intron 15 | 219584 |
| 6 | 22010500 | 22011000 | Cped1 | Intron 1 | 24584 |
| 6 | 22010500 | 22011000 | Cped1 | Intron 1 | 24556 |
| 6 | 22010500 | 22011000 | Cped1 | Intron 1 | 23904 |
| 6 | 22010500 | 22011000 | Cped1 | Non-coding | 23608 |
| 6 | 22205500 | 22206000 | Cped1 | Intron 14 | 188434 |
| 6 | 27612500 | 27613000 | Grm8 | Intron 7 | 521546 |
| 6 | 27612500 | 27613000 | Grm8 | Intron 6 | 513126 |
| 6 | 27612500 | 27613000 | Grm8 | Intron 6 | 513196 |
| 6 | 31065500 | 31066000 | Mir29a | Promoter | -2752 |
| 6 | 31065500 | 31066000 | Mir29b-1 | Promoter | -2406 |
| 6 | 31065500 | 31066000 | Gm27019 | Promoter | -2004 |
| 6 | 31125000 | 31125500 | Lncpint | Non-coding | 41017 |
| 6 | 31125000 | 31125500 | Lncpint | Non-coding | 41017 |
| 6 | 31205500 | 31206000 | Lncpint | Non-coding | 12475 |
| 6 | 31125000 | 31125500 | Lncpint | Non-coding | 92975 |
| 6 | 31205500 | 31206000 | Lncpint | Non-coding | 12223 |
| 6 | 31205500 | 31206000 | Lncpint | Non-coding | 12434 |
| 6 | 31205500 | 31206000 | Lncpint | Non-coding | 12421 |
| 6 | 31205500 | 31206000 | Lncpint | Non-coding | 12347 |
| 6 | 31205500 | 31206000 | Lncpint | Non-coding | 10367 |
| 6 | 31624000 | 31624500 | Gm43154 | Non-coding | 18810 |
| 6 | 31624000 | 31624500 | Gm43154 | Non-coding | 18789 |
| 6 | 31624000 | 31624500 | Gm43154 | Non-coding | 18711 |
| 6 | 34932000 | 34932500 | Stra8 | Intron 5 | 11837 |
| 6 | 34932000 | 34932500 | Stra8 | Intron 4 | 11040 |
| 6 | 34932000 | 34932500 | Stra8 | Intron 4 | 10996 |
| 6 | 34932000 | 34932500 | Stra8 | Intron 3 | 10990 |
| 6 | 38066000 | 38066500 | Atp6v0a4 | Intron 14 | 58087 |
| 6 | 38123000 | 38123500 | Atp6v0a4 | Promoter | 1087 |
| 6 | 38123000 | 38123500 | Atp6v0a4 | 5' UTR | 1087 |
| 6 | 38066000 | 38066500 | Atp6v0a4 | Intron 15 | 43050 |

| | | | | | |
|---|---|---|---|---|---|
| 6 | 38123000 | 38123500 | Atp6v0a4 | Promoter | 990 |
| 6 | 38123000 | 38123500 | D630045J12Rik | 3' UTR | 130456 |
| 6 | 38123000 | 38123500 | D630045J12Rik | 3' UTR | 130510 |
| 6 | 40442501 | 40443001 | Wee2 | 5' UTR | 3413 |
| 6 | 40443501 | 40444001 | Wee2 | 5' UTR | 4413 |
| 6 | 40443501 | 40444001 | Wee2 | Intron 2 | 4413 |
| 6 | 40443501 | 40444001 | Wee2 | CDS 3 | 4413 |
| 6 | 40443001 | 40443501 | Wee2 | 5' UTR | 3913 |
| 6 | 40442501 | 40443001 | Wee2 | Promoter | 0 |
| 6 | 40442501 | 40443001 | Wee2 | 5' UTR | 0 |
| 6 | 40443501 | 40444001 | Wee2 | Promoter | 638 |
| 6 | 40443501 | 40444001 | Wee2 | 5' UTR | 638 |
| 6 | 40443501 | 40444001 | Wee2 | Intron 1 | 638 |
| 6 | 40443501 | 40444001 | Wee2 | CDS 2 | 638 |
| 6 | 40443001 | 40443501 | Wee2 | Promoter | 138 |
| 6 | 40443001 | 40443501 | Wee2 | 5' UTR | 138 |
| 6 | 40442501 | 40443001 | Wee2 | Promoter | 0 |
| 6 | 40442501 | 40443001 | Wee2 | 5' UTR | 0 |
| 6 | 40443501 | 40444001 | Wee2 | Promoter | 564 |
| 6 | 40443501 | 40444001 | Wee2 | 5' UTR | 564 |
| 6 | 40443501 | 40444001 | Wee2 | CDS 2 | 564 |
| 6 | 40443501 | 40444001 | Wee2 | Intron 2 | 564 |
| 6 | 40443501 | 40444001 | Wee2 | CDS 3 | 564 |
| 6 | 40443001 | 40443501 | Wee2 | Promoter | 64 |
| 6 | 40443001 | 40443501 | Wee2 | 5' UTR | 64 |
| 6 | 43309001 | 43309501 | Nobox | Promoter | 54 |
| 6 | 43309001 | 43309501 | Nobox | Intron 1 | 54 |
| 6 | 43309001 | 43309501 | Nobox | CDS 1 | 54 |
| 6 | 43309001 | 43309501 | Nobox | Promoter | -3865 |
| 6 | 43309001 | 43309501 | Nobox | Promoter | -1764 |
| 6 | 45220001 | 45220501 | Cntnap2 | Intron 2 | 160644 |
| 6 | 45220001 | 45220501 | Cntnap2 | Intron 1 | 159940 |
| 6 | 48741001 | 48741501 | Gm28053 | 5' UTR | 22394 |
| 6 | 48741001 | 48741501 | Gm28053 | Intron 2 | 22394 |
| 6 | 48741001 | 48741501 | Gm28053 | CDS 3 | 22394 |
| 6 | 48741001 | 48741501 | Gm28053 | Intron 3 | 22394 |
| 6 | 48741001 | 48741501 | Gimap1os | Promoter | -379 |
| 6 | 48741001 | 48741501 | Gimap1 | Promoter | 1947 |
| 6 | 48741001 | 48741501 | Gimap1 | 5' UTR | 1947 |
| 6 | 48741001 | 48741501 | Gimap1 | Intron 1 | 1947 |
| 6 | 48741001 | 48741501 | Gimap1 | CDS 2 | 1947 |
| 6 | 48741001 | 48741501 | Gimap1 | Intron 2 | 1947 |
| 6 | 48741001 | 48741501 | Gm28053 | Promoter | 1944 |
| 6 | 48741001 | 48741501 | Gm28053 | 5' UTR | 1944 |
| 6 | 48741001 | 48741501 | Gm28053 | Intron 1 | 1944 |
| 6 | 48741001 | 48741501 | Gm28053 | CDS 2 | 1944 |

| | | | | | |
|---|---|---|---|---|---|
| 6 | 48741001 | 48741501 | Gm28053 | Intron 2 | 1944 |
| 6 | 48741001 | 48741501 | Gimap1 | Promoter | 1928 |
| 6 | 48741001 | 48741501 | Gimap1 | 5' UTR | 1928 |
| 6 | 48741001 | 48741501 | Gimap1 | Intron 1 | 1928 |
| 6 | 48741001 | 48741501 | Gimap1 | CDS 2 | 1928 |
| 6 | 48741001 | 48741501 | Gimap1 | Intron 2 | 1928 |
| 6 | 48741001 | 48741501 | Gimap1 | Promoter | 1925 |
| 6 | 48741001 | 48741501 | Gimap1 | 5' UTR | 1925 |
| 6 | 48741001 | 48741501 | Gimap1 | Intron 1 | 1925 |
| 6 | 48741001 | 48741501 | Gimap1 | CDS 2 | 1925 |
| 6 | 48741001 | 48741501 | Gimap1 | Intron 2 | 1925 |
| 6 | 48741001 | 48741501 | Gimap1 | Promoter | 1914 |
| 6 | 48741001 | 48741501 | Gimap1 | Promoter | 1894 |
| 6 | 48741001 | 48741501 | Gimap1os | Promoter | 0 |
| 6 | 48741001 | 48741501 | Gimap1 | Promoter | -681 |
| 6 | 48741001 | 48741501 | Gimap5 | Promoter | -4696 |
| 6 | 48741001 | 48741501 | Gimap5 | Promoter | -4697 |
| 6 | 48741001 | 48741501 | Gimap5 | Promoter | -4698 |
| 6 | 48741001 | 48741501 | Gimap5 | Promoter | -4798 |
| 6 | 53394006 | 53394506 | Creb5 | Intron 4 | 106736 |
| 6 | 53394006 | 53394506 | Creb5 | Non-coding | 106720 |
| 6 | 53394006 | 53394506 | Creb5 | Non-coding | 106711 |
| 6 | 53394006 | 53394506 | Creb5 | Non-coding | 106688 |
| 6 | 53394006 | 53394506 | Creb5 | Non-coding | 106688 |
| 6 | 53394006 | 53394506 | Creb5 | Non-coding | 106685 |
| 6 | 53394006 | 53394506 | Creb5 | Non-coding | 106678 |
| 6 | 53394006 | 53394506 | Creb5 | Intron 4 | 103583 |
| 6 | 53394006 | 53394506 | Creb5 | Intron 4 | 83135 |
| 6 | 54689006 | 54689506 | Mturn | CDS 2 | 7382 |
| 6 | 54689006 | 54689506 | Mturn | Intron 2 | 7382 |
| 6 | 54689006 | 54689506 | Mturn | CDS 2 | 7252 |
| 6 | 54689006 | 54689506 | Mturn | Intron 2 | 7252 |
| 6 | 54689006 | 54689506 | Mturn | Non-coding | 7204 |
| 6 | 54689006 | 54689506 | Gm44008 | Promoter | -4140 |
| 6 | 55489006 | 55489506 | Adcyap1r1 | Intron 13 | 37028 |
| 6 | 55489006 | 55489506 | Adcyap1r1 | Intron 13 | 36978 |
| 6 | 55489006 | 55489506 | Adcyap1r1 | Intron 12 | 28770 |
| 6 | 55489006 | 55489506 | Adcyap1r1 | Intron 13 | 28770 |
| 6 | 55489006 | 55489506 | Adcyap1r1 | Intron 13 | 28770 |
| 6 | 55489006 | 55489506 | Adcyap1r1 | Intron 10 | 28770 |
| 6 | 56918006 | 56918506 | Nt5c3 | Intron 1 | 5427 |
| 6 | 56918006 | 56918506 | Nt5c3 | Intron 1 | 5314 |
| 6 | 56918006 | 56918506 | Nt5c3 | Intron 1 | 5371 |
| 6 | 56918006 | 56918506 | Nt5c3 | Intron 1 | 5219 |
| 6 | 56918006 | 56918506 | Nt5c3 | Non-coding | 5385 |
| 6 | 56918006 | 56918506 | Nt5c3 | Non-coding | 5352 |

| | | | | | |
|---|---|---|---|---|---|
| 6 | 72714506 | 72715006 | Tcf7l1 | Intron 3 | 74249 |
| 6 | 72714506 | 72715006 | Tcf7l1 | Intron 3 | 73947 |
| 6 | 72714506 | 72715006 | Tcf7l1 | Intron 2 | 73593 |
| 6 | 73070006 | 73070506 | Dnah6 | Intron 57 | 151126 |
| 6 | 73070006 | 73070506 | Dnah6 | CDS 57 | 151126 |
| 6 | 73070006 | 73070506 | Dnah6 | Intron 56 | 151146 |
| 6 | 73070006 | 73070506 | Dnah6 | CDS 56 | 151146 |
| 6 | 73070006 | 73070506 | Dnah6 | Intron 56 | 142158 |
| 6 | 73070006 | 73070506 | Dnah6 | CDS 56 | 142158 |
| 6 | 77232006 | 77232506 | Ctnna2 | Intron 8 | 747194 |
| 6 | 77232006 | 77232506 | Ctnna2 | Intron 8 | 747194 |
| 6 | 77232006 | 77232506 | Ctnna2 | Intron 7 | 747045 |
| 6 | 77232006 | 77232506 | Ctnna2 | Intron 7 | 747055 |
| 6 | 79539006 | 79539506 | Gm6261 | Promoter | -1702 |
| 6 | 87385006 | 87385506 | Gkn3 | Intron 4 | 3430 |
| 6 | 87385006 | 87385506 | Gkn3 | CDS 4 | 3430 |
| 6 | 87424006 | 87424506 | Bmp10 | Promoter | -4488 |
| 6 | 90477006 | 90477506 | Gm44421 | Promoter | -960 |
| 6 | 94200506 | 94201006 | Magi1 | Intron 1 | 82317 |
| 6 | 94200506 | 94201006 | Magi1 | Intron 1 | 82317 |
| 6 | 94200506 | 94201006 | Magi1 | Intron 1 | 82912 |
| 6 | 94200506 | 94201006 | Magi1 | Intron 1 | 82890 |
| 6 | 94200506 | 94201006 | Magi1 | Non-coding | 82020 |
| 6 | 94542006 | 94542506 | Slc25a26 | Intron 5 | 41675 |
| 6 | 94542006 | 94542506 | Slc25a26 | 3' UTR | 41673 |
| 6 | 94542006 | 94542506 | Slc25a26 | 3' UTR | 41637 |
| 6 | 94542006 | 94542506 | Slc25a26 | Intron 4 | 34428 |
| 6 | 94542006 | 94542506 | Slc25a26 | Non-coding | 7866 |
| 6 | 117994982 | 117995482 | Gm4875 | Promoter | -1725 |
| 6 | 126533482 | 126533982 | Kcna5 | Promoter | 1431 |
| 6 | 126533482 | 126533982 | Kcna5 | CDS 1 | 1431 |
| 6 | 126641982 | 126642482 | Kcna1 | Promoter | 2315 |
| 6 | 126641982 | 126642482 | Kcna1 | CDS 2 | 2315 |
| 6 | 126642482 | 126642982 | Kcna1 | Promoter | 1815 |
| 6 | 126642482 | 126642982 | Kcna1 | CDS 2 | 1815 |
| 6 | 126641982 | 126642482 | Kcna1 | Promoter | 2332 |
| 6 | 126641982 | 126642482 | Kcna1 | CDS 1 | 2332 |
| 6 | 126642482 | 126642982 | Kcna1 | Promoter | 1832 |
| 6 | 126642482 | 126642982 | Kcna1 | CDS 1 | 1832 |
| 6 | 126738482 | 126738982 | Kcna6 | Promoter | 1681 |
| 6 | 126738482 | 126738982 | Kcna6 | CDS 1 | 1681 |
| 6 | 126738482 | 126738982 | Kcna6 | Promoter | -1670 |
| 6 | 126738482 | 126738982 | Kcna6 | Promoter | 1693 |
| 6 | 126738482 | 126738982 | Kcna6 | CDS 1 | 1693 |
| 6 | 126738482 | 126738982 | Gm26673 | Non-coding | 19798 |
| 6 | 126738482 | 126738982 | Kcna6 | Promoter | 1170 |

| | | | | | |
|---|---|---|---|---|---|
| 6 | 126738482 | 126738982 | Kcna6 | CDS 1 | 1170 |
| 6 | 128077982 | 128078482 | Tspan9 | 5' UTR | 65082 |
| 6 | 128011482 | 128011982 | Tspan9 | Intron 2 | 131582 |
| 6 | 128077982 | 128078482 | Tspan9 | 5' UTR | 58228 |
| 6 | 128011482 | 128011982 | Tspan9 | Intron 3 | 124728 |
| 6 | 128077982 | 128078482 | Tspan9 | 5' UTR | 65079 |
| 6 | 128011482 | 128011982 | Tspan9 | Intron 3 | 131579 |
| 6 | 128011482 | 128011982 | Tspan9 | Intron 1 | 22611 |
| 6 | 128011482 | 128011982 | Tspan9 | Intron 2 | 23024 |
| 6 | 128077982 | 128078482 | Tspan9 | 5' UTR | 65079 |
| 6 | 128011482 | 128011982 | Tspan9 | Intron 4 | 131579 |
| 6 | 128077982 | 128078482 | Tspan9 | 5' UTR | 62265 |
| 6 | 128011482 | 128011982 | Tspan9 | Intron 3 | 128765 |
| 6 | 128077982 | 128078482 | Tspan9 | 5' UTR | 65072 |
| 6 | 128011482 | 128011982 | Tspan9 | 5' UTR | 131572 |
| 6 | 128077982 | 128078482 | Tspan9 | 5' UTR | 45835 |
| 6 | 128011482 | 128011982 | Tspan9 | Intron 3 | 112335 |
| 6 | 134144482 | 134144982 | Etv6 | Intron 2 | 108782 |
| 6 | 134182482 | 134182982 | Etv6 | Intron 2 | 146782 |
| 6 | 134144482 | 134144982 | Etv6 | 5' UTR | 108782 |
| 6 | 134182482 | 134182982 | Etv6 | 5' UTR | 146782 |
| 6 | 134144482 | 134144982 | Etv6 | Intron 2 | 108513 |
| 6 | 134182482 | 134182982 | Etv6 | Intron 2 | 146513 |
| 6 | 134981482 | 134981982 | Apold1 | Promoter | 0 |
| 6 | 134981482 | 134981982 | Apold1 | 5' UTR | 0 |
| 6 | 137401979 | 137402479 | Ptpro | Intron 12 | 149645 |
| 6 | 137401979 | 137402479 | Ptpro | Intron 12 | 149514 |
| 6 | 143153480 | 143153980 | Gm31108 | Promoter | 0 |
| 6 | 144742980 | 144743480 | Sox5 | Non-coding | 38498 |
| 6 | 144742980 | 144743480 | Sox5 | Non-coding | 38440 |
| 6 | 144742980 | 144743480 | Sox5os4 | Non-coding | 22069 |
| 6 | 145016480 | 145016980 | Bcat1 | Intron 6 | 31833 |
| 6 | 145016480 | 145016980 | Bcat1 | Intron 6 | 59205 |
| 6 | 145016480 | 145016980 | Bcat1 | Intron 6 | 59205 |
| 6 | 145016480 | 145016980 | Bcat1 | Non-coding | 18218 |
| 6 | 145016480 | 145016980 | Bcat1 | Intron 3 | 31631 |
| 6 | 145016480 | 145016980 | Bcat1 | Intron 2 | 15864 |
| 6 | 145754980 | 145755480 | Rassf8 | 5' UTR | 8232 |
| 6 | 145754980 | 145755480 | Rassf8 | 5' UTR | 4409 |
| 7 | 4062898 | 4063398 | Lair1 | Promoter | 0 |
| 7 | 4062898 | 4063398 | Lair1 | 5' UTR | 0 |
| 7 | 4062898 | 4063398 | Lair1 | Intron 1 | 0 |
| 7 | 4062898 | 4063398 | Lair1 | CDS 1 | 0 |
| 7 | 4062898 | 4063398 | Lair1 | Promoter | 0 |
| 7 | 4062898 | 4063398 | Lair1 | 5' UTR | 0 |
| 7 | 4062898 | 4063398 | Lair1 | Intron 1 | 0 |

| | | | | | |
|---|---|---|---|---|---|
| 7 | 4062898 | 4063398 | Lair1 | CDS 1 | 0 |
| 7 | 4062898 | 4063398 | Lair1 | Promoter | 0 |
| 7 | 4062898 | 4063398 | Lair1 | 5' UTR | 0 |
| 7 | 4062898 | 4063398 | Lair1 | Intron 1 | 0 |
| 7 | 4062898 | 4063398 | Lair1 | CDS 1 | 0 |
| 7 | 4062898 | 4063398 | Lair1 | Promoter | 0 |
| 7 | 4062898 | 4063398 | Lair1 | 5' UTR | 0 |
| 7 | 4062898 | 4063398 | Lair1 | Intron 1 | 0 |
| 7 | 4062898 | 4063398 | Lair1 | CDS 1 | 0 |
| 7 | 4062898 | 4063398 | Lair1 | Promoter | 0 |
| 7 | 4062898 | 4063398 | Lair1 | 5' UTR | 0 |
| 7 | 4062898 | 4063398 | Lair1 | Intron 1 | 0 |
| 7 | 4062898 | 4063398 | Lair1 | CDS 1 | 0 |
| 7 | 4062898 | 4063398 | Lair1 | Promoter | 0 |
| 7 | 4062898 | 4063398 | Lair1 | 5' UTR | 0 |
| 7 | 4062898 | 4063398 | Lair1 | Intron 1 | 0 |
| 7 | 4062898 | 4063398 | Lair1 | CDS 1 | 0 |
| 7 | 4062898 | 4063398 | Lair1 | Promoter | 0 |
| 7 | 4062898 | 4063398 | Lair1 | 5' UTR | 0 |
| 7 | 4062898 | 4063398 | Lair1 | Intron 1 | 0 |
| 7 | 4062898 | 4063398 | Lair1 | CDS 1 | 0 |
| 7 | 4062898 | 4063398 | Lair1 | Promoter | 0 |
| 7 | 4062898 | 4063398 | Lair1 | 5' UTR | 0 |
| 7 | 4062898 | 4063398 | Lair1 | Intron 1 | 0 |
| 7 | 4062898 | 4063398 | Lair1 | CDS 1 | 0 |
| 7 | 4062898 | 4063398 | Lair1 | Promoter | 0 |
| 7 | 4615898 | 4616398 | Gm18973 | Promoter | -2732 |
| 7 | 4950398 | 4950898 | Sbk2 | Intron 3 | 13509 |
| 7 | 12996151 | 12996651 | Slc27a5 | Promoter | 1500 |
| 7 | 12996151 | 12996651 | Slc27a5 | Intron 2 | 1500 |
| 7 | 12996151 | 12996651 | Slc27a5 | CDS 2 | 1500 |
| 7 | 12996151 | 12996651 | Slc27a5 | Intron 1 | 1500 |
| 7 | 12996151 | 12996651 | Slc27a5 | Promoter | 1542 |
| 7 | 12996151 | 12996651 | Slc27a5 | Intron 2 | 1542 |
| 7 | 12996151 | 12996651 | Slc27a5 | CDS 2 | 1542 |
| 7 | 12996151 | 12996651 | Slc27a5 | Intron 1 | 1542 |
| 7 | 12996151 | 12996651 | Slc27a5 | Promoter | 1027 |
| 7 | 12996151 | 12996651 | Slc27a5 | Intron 1 | 1027 |
| 7 | 12996151 | 12996651 | Slc27a5 | Promoter | 0 |
| 7 | 12996151 | 12996651 | Trim28 | Promoter | -2463 |
| 7 | 14596651 | 14597151 | Nlrp5-ps | Non-coding | 25329 |
| 7 | 14596651 | 14597151 | Nlrp5-ps | Non-coding | 12139 |
| 7 | 14596651 | 14597151 | Nlrp5-ps | Non-coding | 12189 |
| 7 | 16737651 | 16738151 | Ap2s1 | Promoter | -259 |
| 7 | 16737651 | 16738151 | Ap2s1 | Promoter | -293 |
| 7 | 16737651 | 16738151 | Ap2s1 | Promoter | -321 |

| | | | | | |
|---|---|---|---|---|---|
| 7 | 16737651 | 16738151 | Ap2s1 | Promoter | -387 |
| 7 | 18827651 | 18828151 | Gm18855 | Promoter | -3624 |
| 7 | 19086651 | 19087151 | Dmpk | Promoter | 2802 |
| 7 | 19086651 | 19087151 | Dmpk | Intron 4 | 2802 |
| 7 | 19086651 | 19087151 | Dmpk | CDS 5 | 2802 |
| 7 | 19086651 | 19087151 | Dmpk | Intron 5 | 2802 |
| 7 | 19086651 | 19087151 | Dmpk | Promoter | 2783 |
| 7 | 19086651 | 19087151 | Dmpk | Intron 3 | 2783 |
| 7 | 19086651 | 19087151 | Dmpk | CDS 4 | 2783 |
| 7 | 19086651 | 19087151 | Dmpk | Promoter | 2781 |
| 7 | 19086651 | 19087151 | Dmpk | Intron 4 | 2781 |
| 7 | 19086651 | 19087151 | Dmpk | CDS 5 | 2781 |
| 7 | 19086651 | 19087151 | Dmpk | Intron 5 | 2781 |
| 7 | 19086651 | 19087151 | Dmpk | Promoter | 2766 |
| 7 | 19086651 | 19087151 | Dmpk | Intron 4 | 2766 |
| 7 | 19086651 | 19087151 | Dmpk | CDS 5 | 2766 |
| 7 | 19086651 | 19087151 | Dmpk | Intron 5 | 2766 |
| 7 | 19086651 | 19087151 | Dmpk | Promoter | 2710 |
| 7 | 19086651 | 19087151 | Dmpk | Intron 4 | 2710 |
| 7 | 19086651 | 19087151 | Dmpk | CDS 5 | 2710 |
| 7 | 19086651 | 19087151 | Dmpk | Intron 5 | 2710 |
| 7 | 19086651 | 19087151 | Dmpk | Promoter | 2692 |
| 7 | 19086651 | 19087151 | Dmpk | Intron 4 | 2692 |
| 7 | 19086651 | 19087151 | Dmpk | CDS 5 | 2692 |
| 7 | 19086651 | 19087151 | Dmpk | Intron 5 | 2692 |
| 7 | 19086651 | 19087151 | Dmpk | Promoter | 1941 |
| 7 | 19086651 | 19087151 | Dmpk | Promoter | 151 |
| 7 | 19086651 | 19087151 | Dmpk | Promoter | 151 |
| 7 | 19086651 | 19087151 | Dmpk | Promoter | 151 |
| 7 | 19086651 | 19087151 | Mir3100 | Promoter | 0 |
| 7 | 19086651 | 19087151 | Dmpk | Promoter | -118 |
| 7 | 19086651 | 19087151 | Dmpk | Promoter | -342 |
| 7 | 19086651 | 19087151 | Dmpk | Promoter | -460 |
| 7 | 19086651 | 19087151 | Dmpk | Promoter | -595 |
| 7 | 19086651 | 19087151 | Dmpk | Promoter | -1034 |
| 7 | 19086651 | 19087151 | Dmpk | Promoter | -4332 |
| 7 | 19086651 | 19087151 | Dmpk | Promoter | -4465 |
| 7 | 19158651 | 19159151 | Gipr | Intron 12 | 6977 |
| 7 | 19159651 | 19160151 | Gipr | CDS 12 | 5977 |
| 7 | 19159651 | 19160151 | Gipr | Intron 11 | 5977 |
| 7 | 19159651 | 19160151 | Gipr | CDS 11 | 5977 |
| 7 | 19159651 | 19160151 | Gipr | Intron 10 | 5977 |
| 7 | 19159651 | 19160151 | Gipr | CDS 10 | 5977 |
| 7 | 19159651 | 19160151 | Gipr | Intron 9 | 5977 |
| 7 | 19158651 | 19159151 | Gipr | Non-coding | 6977 |
| 7 | 19159651 | 19160151 | Gipr | Non-coding | 5977 |

| | | | | | |
|---|---|---|---|---|---|
| 7 | 19465651 | 19466151 | Gm23099 | Promoter | -4715 |
| 7 | 19466151 | 19466651 | Exoc3l2 | Promoter | 2820 |
| 7 | 19466151 | 19466651 | Exoc3l2 | 5' UTR | 2820 |
| 7 | 19465651 | 19466151 | Exoc3l2 | Promoter | 2320 |
| 7 | 19465651 | 19466151 | Exoc3l2 | 5' UTR | 2320 |
| 7 | 19466151 | 19466651 | Gm45167 | Promoter | -1577 |
| 7 | 19465651 | 19466151 | Gm45167 | Promoter | -2077 |
| 7 | 23348981 | 23349481 | Nlrp4e | Intron 6 | 47789 |
| 7 | 24315981 | 24316481 | Zfp94 | Promoter | 125 |
| 7 | 24315981 | 24316481 | Zfp94 | 5' UTR | 125 |
| 7 | 24315981 | 24316481 | Zfp94 | Promoter | 186 |
| 7 | 24315981 | 24316481 | Zfp94 | 5' UTR | 186 |
| 7 | 24315981 | 24316481 | Zfp94 | Promoter | 144 |
| 7 | 24315981 | 24316481 | Zfp94 | Promoter | 128 |
| 7 | 24315981 | 24316481 | Zfp94 | Promoter | 102 |
| 7 | 24315981 | 24316481 | Gm44513 | Promoter | -1900 |
| 7 | 24327481 | 24327981 | Gm15465 | Promoter | -4696 |
| 7 | 24327481 | 24327981 | AC159140.1 | Promoter | 0 |
| 7 | 24547481 | 24547981 | Pinlyp | Promoter | -1362 |
| 7 | 24547481 | 24547981 | Xrcc1 | Promoter | 1192 |
| 7 | 24547481 | 24547981 | Xrcc1 | Intron 3 | 1192 |
| 7 | 24547481 | 24547981 | Xrcc1 | CDS 4 | 1192 |
| 7 | 24547481 | 24547981 | Xrcc1 | Intron 4 | 1192 |
| 7 | 24547481 | 24547981 | Xrcc1 | Promoter | 331 |
| 7 | 24547481 | 24547981 | Xrcc1 | Intron 1 | 331 |
| 7 | 24547481 | 24547981 | Xrcc1 | CDS 2 | 331 |
| 7 | 24547481 | 24547981 | Xrcc1 | Intron 2 | 331 |
| 7 | 24547481 | 24547981 | Xrcc1 | Promoter | 283 |
| 7 | 24547481 | 24547981 | Xrcc1 | Promoter | 280 |
| 7 | 24547481 | 24547981 | Xrcc1 | Promoter | 196 |
| 7 | 24547481 | 24547981 | Xrcc1 | Promoter | 148 |
| 7 | 27370481 | 27370981 | Sptbn4 | CDS 25 | 75619 |
| 7 | 27370481 | 27370981 | Sptbn4 | Intron 24 | 75619 |
| 7 | 27370481 | 27370981 | Sptbn4 | CDS 7 | 23526 |
| 7 | 27370481 | 27370981 | Sptbn4 | Intron 6 | 23526 |
| 7 | 27370481 | 27370981 | Sptbn4 | CDS 8 | 25573 |
| 7 | 27370481 | 27370981 | Sptbn4 | Intron 7 | 25573 |
| 7 | 27370481 | 27370981 | Sptbn4 | CDS 8 | 25218 |
| 7 | 27370481 | 27370981 | Sptbn4 | Intron 7 | 25218 |
| 7 | 27370481 | 27370981 | Sptbn4 | CDS 25 | 76706 |
| 7 | 27370481 | 27370981 | Sptbn4 | Intron 24 | 76706 |
| 7 | 30974481 | 30974981 | Lsr | Promoter | -1054 |
| 7 | 30974481 | 30974981 | Lsr | Promoter | -1016 |
| 7 | 30974481 | 30974981 | Lsr | Promoter | -1053 |
| 7 | 30974481 | 30974981 | Lsr | Promoter | -1158 |
| 7 | 30974481 | 30974981 | Lsr | Promoter | -1065 |

| | | | | | |
|---|---|---|---|---|---|
| 7 | 30974481 | 30974981 | Fam187b | Promoter | 691 |
| 7 | 30974481 | 30974981 | Fam187b | Promoter | 677 |
| 7 | 30974481 | 30974981 | Fam187b | 5' UTR | 677 |
| 7 | 30974481 | 30974981 | Fam187b | Promoter | -2062 |
| 7 | 34281481 | 34281981 | 4931406P16Rik | Intron 2 | 31571 |
| 7 | 34281481 | 34281981 | 4931406P16Rik | Intron 1 | 3619 |
| 7 | 34281481 | 34281981 | 4931406P16Rik | 5' UTR | 31542 |
| 7 | 37319981 | 37320481 | Zfp536 | 3' UTR | 247910 |
| 7 | 37599481 | 37599981 | Zfp536 | 5' UTR | 169644 |
| 7 | 37599481 | 37599981 | Zfp536 | 5' UTR | 169982 |
| 7 | 37599481 | 37599981 | Zfp536 | 5' UTR | 171000 |
| 7 | 37599481 | 37599981 | Zfp536 | 5' UTR | 172895 |
| 7 | 37599481 | 37599981 | Zfp536 | 5' UTR | 170802 |
| 7 | 37599481 | 37599981 | Zfp536 | 5' UTR | 173661 |
| 7 | 39450130 | 39450630 | Gm5590 | Promoter | -3955 |
| 7 | 39450130 | 39450630 | Gm5590 | Promoter | -908 |
| 7 | 39450130 | 39450630 | Zfp939 | Promoter | 612 |
| 7 | 44582130 | 44582630 | Napsa | Intron 4 | 9750 |
| 7 | 44582130 | 44582630 | Napsa | CDS 5 | 9750 |
| 7 | 44582130 | 44582630 | Napsa | Intron 5 | 9750 |
| 7 | 44582130 | 44582630 | Napsa | Non-coding | 9698 |
| 7 | 44582130 | 44582630 | Gm15396 | Non-coding | 7995 |
| 7 | 44582130 | 44582630 | Napsa | Promoter | 1192 |
| 7 | 44582130 | 44582630 | Napsa | Promoter | 0 |
| 7 | 44582130 | 44582630 | Napsa | 5' UTR | 0 |
| 7 | 44582130 | 44582630 | Napsa | CDS 1 | 0 |
| 7 | 44582130 | 44582630 | Napsa | Intron 1 | 0 |
| 7 | 44582130 | 44582630 | Napsa | Promoter | -2185 |
| 7 | 44746130 | 44746630 | Zfp473 | Promoter | -1899 |
| 7 | 44746130 | 44746630 | Zfp473 | Promoter | 1988 |
| 7 | 44746130 | 44746630 | Zfp473 | 5' UTR | 1988 |
| 7 | 44746130 | 44746630 | Zfp473 | Promoter | 1762 |
| 7 | 44746130 | 44746630 | Zfp473 | 5' UTR | 1762 |
| 7 | 44746130 | 44746630 | Zfp473 | Promoter | 1420 |
| 7 | 44746130 | 44746630 | Zfp473 | 5' UTR | 1420 |
| 7 | 44746130 | 44746630 | Zfp473 | Promoter | 1677 |
| 7 | 44746130 | 44746630 | Zfp473 | 5' UTR | 1677 |
| 7 | 44746130 | 44746630 | Zfp473 | Promoter | 1720 |
| 7 | 44746130 | 44746630 | Zfp473 | 5' UTR | 1720 |
| 7 | 44746130 | 44746630 | Zfp473 | 5' UTR | 4421 |
| 7 | 44746130 | 44746630 | Vrk3 | Promoter | -1783 |
| 7 | 44746130 | 44746630 | Vrk3 | Promoter | -1855 |
| 7 | 44746130 | 44746630 | Vrk3 | Promoter | -1954 |
| 7 | 44746130 | 44746630 | Vrk3 | Promoter | -2006 |
| 7 | 44746130 | 44746630 | Vrk3 | Promoter | -2016 |
| 7 | 44746130 | 44746630 | Vrk3 | Promoter | -2017 |

| | | | | | |
|---|---|---|---|---|---|
| 7 | 44746130 | 44746630 | Vrk3 | Promoter | -2313 |
| 7 | 44860130 | 44860630 | Pnkp | Promoter | 2991 |
| 7 | 44860130 | 44860630 | Pnkp | CDS 7 | 2991 |
| 7 | 44860130 | 44860630 | Pnkp | Intron 7 | 2991 |
| 7 | 44860130 | 44860630 | Pnkp | CDS 8 | 2991 |
| 7 | 44860130 | 44860630 | Pnkp | Intron 8 | 2991 |
| 7 | 44860130 | 44860630 | Pnkp | Promoter | 2979 |
| 7 | 44860130 | 44860630 | Pnkp | Promoter | 2971 |
| 7 | 44860130 | 44860630 | Pnkp | CDS 6 | 2971 |
| 7 | 44860130 | 44860630 | Pnkp | Intron 6 | 2971 |
| 7 | 44860130 | 44860630 | Pnkp | CDS 7 | 2971 |
| 7 | 44860130 | 44860630 | Pnkp | Intron 7 | 2971 |
| 7 | 44860130 | 44860630 | Pnkp | Promoter | 2963 |
| 7 | 44860130 | 44860630 | Pnkp | Promoter | 2957 |
| 7 | 44860130 | 44860630 | Pnkp | 3' UTR | 2957 |
| 7 | 44860130 | 44860630 | Pnkp | Promoter | 2790 |
| 7 | 44860130 | 44860630 | Pnkp | 3' UTR | 2790 |
| 7 | 44860130 | 44860630 | Pnkp | Promoter | 2788 |
| 7 | 44860130 | 44860630 | Pnkp | Intron 5 | 2788 |
| 7 | 44860130 | 44860630 | Pnkp | CDS 6 | 2788 |
| 7 | 44860130 | 44860630 | Pnkp | Intron 6 | 2788 |
| 7 | 44860130 | 44860630 | Pnkp | Promoter | 2782 |
| 7 | 44860130 | 44860630 | Pnkp | 3' UTR | 2782 |
| 7 | 44860130 | 44860630 | Pnkp | Promoter | 2751 |
| 7 | 44860130 | 44860630 | Pnkp | 3' UTR | 2751 |
| 7 | 44860130 | 44860630 | Pnkp | Promoter | 2736 |
| 7 | 44860130 | 44860630 | Pnkp | CDS 6 | 2736 |
| 7 | 44860130 | 44860630 | Pnkp | Intron 6 | 2736 |
| 7 | 44860130 | 44860630 | Pnkp | CDS 7 | 2736 |
| 7 | 44860130 | 44860630 | Pnkp | Intron 7 | 2736 |
| 7 | 44860130 | 44860630 | Pnkp | Promoter | 1634 |
| 7 | 44860130 | 44860630 | Pnkp | CDS 2 | 1634 |
| 7 | 44860130 | 44860630 | Pnkp | Intron 2 | 1634 |
| 7 | 44860130 | 44860630 | Pnkp | CDS 3 | 1634 |
| 7 | 44860130 | 44860630 | Pnkp | Intron 3 | 1634 |
| 7 | 44860130 | 44860630 | Pnkp | Promoter | 0 |
| 7 | 44860130 | 44860630 | Pnkp | Promoter | 0 |
| 7 | 44860130 | 44860630 | Pnkp | Promoter | -238 |
| 7 | 44860130 | 44860630 | Pnkp | Promoter | -745 |
| 7 | 44860130 | 44860630 | Pnkp | Promoter | -884 |
| 7 | 44860130 | 44860630 | Pnkp | Promoter | -1067 |
| 7 | 45093130 | 45093630 | Rcn3 | Promoter | -908 |
| 7 | 45093130 | 45093630 | Rcn3 | Promoter | -908 |
| 7 | 45093130 | 45093630 | Rcn3 | Promoter | -1675 |
| 7 | 45093130 | 45093630 | Rcn3 | Promoter | -4522 |
| 7 | 45093130 | 45093630 | Rcn3 | Promoter | -944 |

| | | | | | |
|---|---|---|---|---|---|
| 7 | 45093130 | 45093630 | Rcn3 | Promoter | -915 |
| 7 | 45093130 | 45093630 | Rcn3 | Promoter | -1404 |
| 7 | 45093130 | 45093630 | Rcn3 | Promoter | -1006 |
| 7 | 45093130 | 45093630 | Gm45669 | Promoter | 1721 |
| 7 | 45093130 | 45093630 | Fcgrt | 3' UTR | 10222 |
| 7 | 45093130 | 45093630 | Fcgrt | CDS 7 | 10222 |
| 7 | 45093130 | 45093630 | Fcgrt | Intron 6 | 10222 |
| 7 | 45093130 | 45093630 | Fcgrt | 3' UTR | 10116 |
| 7 | 45093130 | 45093630 | Fcgrt | CDS 7 | 10116 |
| 7 | 45093130 | 45093630 | Fcgrt | Intron 6 | 10116 |
| 7 | 45408130 | 45408630 | Kcna7 | Promoter | 2477 |
| 7 | 45408130 | 45408630 | Kcna7 | Intron 1 | 2477 |
| 7 | 45560630 | 45561130 | Hsd17b14 | Intron 5 | 5737 |
| 7 | 45560630 | 45561130 | Hsd17b14 | Intron 5 | 5693 |
| 7 | 45560630 | 45561130 | Hsd17b14 | 3' UTR | 4586 |
| 7 | 45560630 | 45561130 | Hsd17b14 | Promoter | 1424 |
| 7 | 45560630 | 45561130 | Hsd17b14 | Promoter | -966 |
| 7 | 45560630 | 45561130 | Hsd17b14 | Promoter | -977 |
| 7 | 45560630 | 45561130 | Hsd17b14 | Promoter | -1940 |
| 7 | 62420114 | 62420614 | Gm32061 | Promoter | 2632 |
| 7 | 62420114 | 62420614 | Mkrn3 | Promoter | 0 |
| 7 | 62420114 | 62420614 | Mkrn3 | 5' UTR | 0 |
| 7 | 64945614 | 64946114 | Fam189a1 | Intron 2 | 210457 |
| 7 | 65151114 | 65151614 | Fam189a1 | Intron 1 | 4957 |
| 7 | 64945614 | 64946114 | Fam189a1 | Intron 2 | 210303 |
| 7 | 65151114 | 65151614 | Fam189a1 | Intron 1 | 4803 |
| 7 | 66124114 | 66124614 | Chsy1 | Intron 1 | 14599 |
| 7 | 66124114 | 66124614 | Gm26827 | Non-coding | 48508 |
| 7 | 66124114 | 66124614 | Gm23042 | Promoter | -4710 |
| 7 | 66405114 | 66405614 | Aldh1a3 | Intron 9 | 21904 |
| 7 | 66405114 | 66405614 | Aldh1a3 | Promoter | 389 |
| 7 | 66405114 | 66405614 | Aldh1a3 | Intron 6 | 13622 |
| 7 | 68159614 | 68160114 | Igf1r | Intron 2 | 206787 |
| 7 | 68159614 | 68160114 | Igf1r | Non-coding | 11024 |
| 7 | 83598990 | 83599490 | Tmc3 | Non-coding | 14063 |
| 7 | 83598990 | 83599490 | Tmc3 | Intron 7 | 14059 |
| 7 | 83598990 | 83599490 | Tmc3 | Intron 7 | 13898 |
| 7 | 83598990 | 83599490 | Gm16638 | Non-coding | 32353 |
| 7 | 83598990 | 83599490 | Tmc3 | Promoter | 1463 |
| 7 | 91263490 | 91263990 | Dlg2 | Intron 6 | 786259 |
| 7 | 92181490 | 92181990 | Dlg2 | Intron 18 | 1704259 |
| 7 | 91263490 | 91263990 | Dlg2 | Intron 1 | 172784 |
| 7 | 92181490 | 92181990 | Dlg2 | Intron 13 | 1090784 |
| 7 | 91263490 | 91263990 | Dlg2 | Non-coding | 172764 |
| 7 | 92181490 | 92181990 | Dlg2 | Non-coding | 1090764 |
| 7 | 91263490 | 91263990 | Dlg2 | Intron 1 | 172762 |

| | | | | | |
|---|---|---|---|---|---|
| 7 | 92181490 | 92181990 | Dlg2 | Intron 13 | 1090762 |
| 7 | 92181490 | 92181990 | Dlg2 | Intron 12 | 509198 |
| 7 | 92181490 | 92181990 | Dlg2 | Intron 12 | 509116 |
| 7 | 92181490 | 92181990 | Dlg2 | Intron 12 | 508998 |
| 7 | 92181490 | 92181990 | Dlg2 | Intron 10 | 470070 |
| 7 | 92181490 | 92181990 | Dlg2 | Intron 4 | 119096 |
| 7 | 92181490 | 92181990 | Dlg2 | Promoter | 806 |
| 7 | 92181490 | 92181990 | Dlg2 | Promoter | 709 |
| 7 | 99075490 | 99075990 | Uvrag | Intron 5 | 65152 |
| 7 | 99075490 | 99075990 | Uvrag | Non-coding | 65139 |
| 7 | 99075490 | 99075990 | Uvrag | Non-coding | 65143 |
| 7 | 99075490 | 99075990 | Uvrag | 3' UTR | 65137 |
| 7 | 99075490 | 99075990 | Uvrag | Non-coding | 23438 |
| 7 | 100878486 | 100878986 | Arhgef17 | Intron 16 | 53122 |
| 7 | 100878486 | 100878986 | Arhgef17 | CDS 16 | 53122 |
| 7 | 100878486 | 100878986 | Arhgef17 | Intron 15 | 53122 |
| 7 | 100878486 | 100878986 | Arhgef17 | CDS 15 | 53122 |
| 7 | 100878486 | 100878986 | Arhgef17 | Intron 16 | 15705 |
| 7 | 100878486 | 100878986 | Arhgef17 | CDS 16 | 15705 |
| 7 | 100878486 | 100878986 | Arhgef17 | Intron 15 | 15705 |
| 7 | 100878486 | 100878986 | Arhgef17 | CDS 15 | 15705 |
| 7 | 100878486 | 100878986 | Arhgef17 | Promoter | 723 |
| 7 | 100878486 | 100878986 | Arhgef17 | Intron 3 | 723 |
| 7 | 100878486 | 100878986 | Arhgef17 | CDS 3 | 723 |
| 7 | 100878486 | 100878986 | Arhgef17 | Intron 2 | 723 |
| 7 | 100878486 | 100878986 | Arhgef17 | CDS 2 | 723 |
| 7 | 100878486 | 100878986 | Arhgef17 | Intron 1 | 723 |
| 7 | 101296986 | 101297486 | Atg16l2 | CDS 5 | 4603 |
| 7 | 101296986 | 101297486 | Atg16l2 | Intron 4 | 4603 |
| 7 | 101296986 | 101297486 | Atg16l2 | CDS 4 | 4603 |
| 7 | 101296986 | 101297486 | Atg16l2 | Intron 3 | 4603 |
| 7 | 101296986 | 101297486 | Atg16l2 | CDS 5 | 4542 |
| 7 | 101296986 | 101297486 | Atg16l2 | Intron 4 | 4542 |
| 7 | 101296986 | 101297486 | Atg16l2 | CDS 4 | 4542 |
| 7 | 101296986 | 101297486 | Atg16l2 | Intron 3 | 4542 |
| 7 | 101296986 | 101297486 | Atg16l2 | CDS 5 | 4586 |
| 7 | 101296986 | 101297486 | Atg16l2 | Intron 4 | 4586 |
| 7 | 101296986 | 101297486 | Atg16l2 | CDS 4 | 4586 |
| 7 | 101296986 | 101297486 | Atg16l2 | Intron 3 | 4586 |
| 7 | 101296986 | 101297486 | Atg16l2 | CDS 5 | 4766 |
| 7 | 101296986 | 101297486 | Atg16l2 | Intron 4 | 4766 |
| 7 | 101296986 | 101297486 | Atg16l2 | CDS 4 | 4766 |
| 7 | 101296986 | 101297486 | Atg16l2 | Intron 3 | 4766 |
| 7 | 101296986 | 101297486 | Atg16l2 | Promoter | -4842 |
| 7 | 101296986 | 101297486 | Atg16l2 | Promoter | -4839 |
| 7 | 101296986 | 101297486 | Atg16l2 | Promoter | -4597 |

| | | | | | |
|---|---|---|---|---|---|
| 7 | 101296986 | 101297486 | Atg16l2 | Promoter | -216 |
| 7 | 101296986 | 101297486 | Atg16l2 | Promoter | -203 |
| 7 | 101296986 | 101297486 | Atg16l2 | Promoter | -273 |
| 7 | 101296986 | 101297486 | Atg16l2 | Promoter | 0 |
| 7 | 101296986 | 101297486 | Atg16l2 | 3' UTR | 4586 |
| 7 | 101296986 | 101297486 | Atg16l2 | CDS 4 | 4586 |
| 7 | 101296986 | 101297486 | Atg16l2 | Intron 3 | 4586 |
| 7 | 101296986 | 101297486 | Atg16l2 | CDS 3 | 4586 |
| 7 | 101296986 | 101297486 | Atg16l2 | Intron 2 | 4586 |
| 7 | 104286986 | 104287486 | Trim5 | Promoter | 609 |
| 7 | 104286986 | 104287486 | Trim5 | 5' UTR | 609 |
| 7 | 106994486 | 106994986 | Olfr712-ps1 | Promoter | -4819 |
| 7 | 111030486 | 111030986 | Ctr9 | Promoter | 1535 |
| 7 | 111030486 | 111030986 | Ctr9 | Intron 2 | 1535 |
| 7 | 111030486 | 111030986 | Ctr9 | Promoter | 1491 |
| 7 | 111030486 | 111030986 | Ctr9 | Promoter | 1473 |
| 7 | 111030486 | 111030986 | Ctr9 | Promoter | 271 |
| 7 | 111030486 | 111030986 | Ctr9 | Promoter | -3050 |
| 7 | 111592486 | 111592986 | Galnt18 | Intron 3 | 186992 |
| 7 | 111592486 | 111592986 | Galnt18 | Intron 3 | 186992 |
| 7 | 112620486 | 112620986 | 2310014F06Rik | Non-coding | 59073 |
| 7 | 112620486 | 112620986 | Gm24154 | Promoter | -2260 |
| 7 | 119684986 | 119685486 | Acsm4 | Promoter | -4540 |
| 7 | 121393486 | 121393986 | Gm36736 | Promoter | -384 |
| 7 | 121393486 | 121393986 | Hs3st2 | Promoter | 1627 |
| 7 | 121393486 | 121393986 | Hs3st2 | Intron 2 | 1627 |
| 7 | 121393486 | 121393986 | Hs3st2 | Promoter | 1196 |
| 7 | 121393486 | 121393986 | Hs3st2 | Intron 1 | 1196 |
| 7 | 127903486 | 127903986 | Bckdk | Promoter | -96 |
| 7 | 127903486 | 127903986 | Bckdk | Promoter | -117 |
| 7 | 127903486 | 127903986 | Bckdk | Promoter | -654 |
| 7 | 127903486 | 127903986 | Bckdk | Promoter | -776 |
| 7 | 127903486 | 127903986 | Bckdk | Promoter | -776 |
| 7 | 127903486 | 127903986 | Bckdk | Promoter | -931 |
| 7 | 127903486 | 127903986 | Bckdk | Promoter | -3254 |
| 7 | 128004486 | 128004986 | Gm15533 | Promoter | 175 |
| 7 | 128004486 | 128004986 | Gm15533 | Promoter | 205 |
| 7 | 128004486 | 128004986 | Trim72 | Promoter | 537 |
| 7 | 128004486 | 128004986 | Trim72 | CDS 2 | 537 |
| 7 | 128004486 | 128004986 | Trim72 | Intron 2 | 537 |
| 7 | 128004486 | 128004986 | Trim72 | Promoter | 108 |
| 7 | 128004486 | 128004986 | Trim72 | CDS 1 | 108 |
| 7 | 128004486 | 128004986 | Trim72 | Intron 1 | 108 |
| 7 | 128273986 | 128274486 | BC017158 | Intron 10 | 23685 |
| 7 | 128273986 | 128274486 | BC017158 | Promoter | 363 |
| 7 | 128273986 | 128274486 | BC017158 | Non-coding | 23624 |

| | | | | | |
|---|---|---|---|---|---|
| 7 | 128273986 | 128274486 | BC017158 | Non-coding | 23623 |
| 7 | 128273986 | 128274486 | BC017158 | 3' UTR | 23610 |
| 7 | 131412486 | 131412986 | Fgfr2 | 5' UTR | 1710365 |
| 7 | 131412486 | 131412986 | Ikzf5 | Promoter | -1964 |
| 7 | 131412486 | 131412986 | Ikzf5 | Promoter | -1988 |
| 7 | 131412486 | 131412986 | Ikzf5 | Promoter | -2107 |
| 7 | 131412486 | 131412986 | Ikzf5 | Promoter | -2012 |
| 7 | 131412486 | 131412986 | Ikzf5 | Promoter | -1990 |
| 7 | 131412486 | 131412986 | Ikzf5 | Promoter | -2080 |
| 7 | 131412486 | 131412986 | Acadsb | Promoter | 1885 |
| 7 | 131412486 | 131412986 | Acadsb | Intron 1 | 1885 |
| 7 | 131412486 | 131412986 | Acadsb | Promoter | 1783 |
| 7 | 131412486 | 131412986 | Acadsb | Intron 1 | 1783 |
| 7 | 131412486 | 131412986 | Acadsb | Promoter | 1751 |
| 7 | 131412486 | 131412986 | Acadsb | Promoter | 1711 |
| 7 | 131412486 | 131412986 | Acadsb | Intron 1 | 1711 |
| 7 | 133916817 | 133917317 | Adam12 | Intron 16 | 307781 |
| 7 | 133916817 | 133917317 | Adam12 | Intron 6 | 43949 |
| 7 | 139020357 | 139020857 | Jakmip3 | Intron 5 | 79627 |
| 7 | 139020357 | 139020857 | Jakmip3 | CDS 6 | 79627 |
| 7 | 139020357 | 139020857 | Jakmip3 | Intron 6 | 79627 |
| 7 | 139020357 | 139020857 | Jakmip3 | CDS 7 | 79627 |
| 7 | 139020357 | 139020857 | Jakmip3 | Non-coding | 31037 |
| 7 | 139020357 | 139020857 | Jakmip3 | Non-coding | 31034 |
| 7 | 139020357 | 139020857 | Gm45613 | Promoter | 143 |
| 7 | 139301601 | 139302101 | Gm32486 | Promoter | -4735 |
| 7 | 139874601 | 139875101 | Adgra1 | Intron 7 | 40427 |
| 7 | 139874601 | 139875101 | Adgra1 | CDS 8 | 40427 |
| 7 | 140050101 | 140050601 | Msx3 | Promoter | -1011 |
| 7 | 140050101 | 140050601 | Msx3 | Promoter | -1012 |
| 7 | 140948601 | 140949101 | Ifitm5 | Promoter | 1191 |
| 7 | 140948601 | 140949101 | Ifitm5 | 3' UTR | 1191 |
| 7 | 144909095 | 144909595 | Gm39119 | Promoter | 491 |
| 7 | 144909095 | 144909595 | Bc1 | Promoter | -4875 |
| 8 | 8443000 | 8443500 | Gm31135 | Non-coding | 41222 |
| 8 | 8443000 | 8443500 | Gm31135 | Non-coding | 41231 |
| 8 | 9504500 | 9505000 | Fam155a | Intron 1 | 266162 |
| 8 | 9239000 | 9239500 | Fam155a | Intron 1 | 531662 |
| 8 | 9239000 | 9239500 | Fam155a | Non-coding | 84212 |
| 8 | 9504500 | 9505000 | Fam155a | Intron 1 | 266022 |
| 8 | 9239000 | 9239500 | Fam155a | Intron 1 | 531522 |
| 8 | 9239000 | 9239500 | Fam155a | Non-coding | 83789 |
| 8 | 9504500 | 9505000 | Fam155a | 3' UTR | 265105 |
| 8 | 12463000 | 12463500 | Gm33326 | Non-coding | 3465 |
| 8 | 14385000 | 14385500 | Dlgap2 | Intron 2 | 289135 |
| 8 | 14385000 | 14385500 | Dlgap2 | Intron 3 | 289125 |

| | | | | | |
|---|---|---|---|---|---|
| 8 | 14385000 | 14385500 | Dlgap2 | Intron 2 | 205333 |
| 8 | 14385000 | 14385500 | Dlgap2 | Intron 2 | 205333 |
| 8 | 14385000 | 14385500 | Dlgap2 | Non-coding | 3266 |
| 8 | 14385000 | 14385500 | Dlgap2 | Non-coding | 3052 |
| 8 | 14385000 | 14385500 | Dlgap2 | Non-coding | 3004 |
| 8 | 18741000 | 18741500 | Mcph1 | Intron 12 | 145869 |
| 8 | 18741000 | 18741500 | Angpt2 | Promoter | 63 |
| 8 | 18741000 | 18741500 | Angpt2 | 5' UTR | 63 |
| 8 | 18741000 | 18741500 | Angpt2 | CDS 1 | 63 |
| 8 | 23153528 | 23154028 | Gm15816 | Promoter | -3942 |
| 8 | 23153528 | 23154028 | Nkx6-3 | Promoter | 259 |
| 8 | 23153528 | 23154028 | Nkx6-3 | 5' UTR | 259 |
| 8 | 23153528 | 23154028 | Nkx6-3 | CDS 1 | 259 |
| 8 | 23153528 | 23154028 | Nkx6-3 | Intron 1 | 259 |
| 8 | 24948028 | 24948528 | Adam32 | Promoter | 277 |
| 8 | 24948028 | 24948528 | Adam32 | Intron 1 | 277 |
| 8 | 24948028 | 24948528 | Adam32 | Promoter | 268 |
| 8 | 24948028 | 24948528 | Adam32 | Intron 1 | 268 |
| 8 | 24948028 | 24948528 | Adam32 | Promoter | 254 |
| 8 | 24948028 | 24948528 | Adam32 | Intron 1 | 254 |
| 8 | 24948028 | 24948528 | Adam32 | Promoter | 277 |
| 8 | 26104028 | 26104528 | Hook3 | Intron 2 | 14697 |
| 8 | 26982028 | 26982528 | Gm45371 | Promoter | -2026 |
| 8 | 27202528 | 27203028 | Got1l1 | Promoter | 0 |
| 8 | 27202528 | 27203028 | Got1l1 | 5' UTR | 0 |
| 8 | 27202528 | 27203028 | Got1l1 | Promoter | -3810 |
| 8 | 27202528 | 27203028 | Got1l1 | Promoter | 0 |
| 8 | 27202528 | 27203028 | Got1l1 | Non-coding | 20801 |
| 8 | 27202528 | 27203028 | Got1l1 | 5' UTR | 20801 |
| 8 | 27553028 | 27553528 | Gm45861 | Intron 46 | 105928 |
| 8 | 27553028 | 27553528 | Gm45861 | Intron 45 | 105637 |
| 8 | 29090528 | 29091028 | Unc5d | Intron 1 | 128609 |
| 8 | 29090528 | 29091028 | Unc5d | Intron 1 | 128542 |
| 8 | 29090528 | 29091028 | Unc5d | Intron 1 | 128400 |
| 8 | 29090528 | 29091028 | Unc5d | Intron 1 | 128400 |
| 8 | 29090528 | 29091028 | Unc5d | Intron 1 | 128400 |
| 8 | 29090528 | 29091028 | Gm9911 | Promoter | -3041 |
| 8 | 29090528 | 29091028 | Gm45652 | Promoter | 876 |
| 8 | 31122028 | 31122528 | Rnf122 | Intron 2 | 10208 |
| 8 | 31122028 | 31122528 | Rnf122 | Non-coding | 10198 |
| 8 | 31122028 | 31122528 | Rnf122 | Intron 2 | 10198 |
| 8 | 31937528 | 31938028 | Nrg1 | Intron 5 | 71513 |
| 8 | 31937528 | 31938028 | Nrg1 | Intron 5 | 71514 |
| 8 | 31937528 | 31938028 | Nrg1 | Intron 5 | 946227 |
| 8 | 31937528 | 31938028 | Nrg1 | Intron 5 | 946770 |
| 8 | 31937528 | 31938028 | Nrg1 | Intron 5 | 71477 |

| | | | | | |
|---|---|---|---|---|---|
| 8 | 31937528 | 31938028 | Nrg1 | Intron 5 | 71444 |
| 8 | 31937528 | 31938028 | Nrg1 | Intron 5 | 71155 |
| 8 | 31937528 | 31938028 | Nrg1 | Intron 5 | 71424 |
| 8 | 31937528 | 31938028 | Nrg1 | Intron 3 | 71082 |
| 8 | 31937528 | 31938028 | Nrg1 | Intron 5 | 71094 |
| 8 | 31937528 | 31938028 | Nrg1 | Intron 3 | 71082 |
| 8 | 31937528 | 31938028 | Nrg1 | Intron 5 | 71126 |
| 8 | 31937528 | 31938028 | Nrg1 | Intron 2 | 12030 |
| 8 | 31937528 | 31938028 | Nrg1 | Non-coding | 12008 |
| 8 | 31937528 | 31938028 | Nrg1 | Intron 5 | 946227 |
| 8 | 31937528 | 31938028 | Nrg1 | Intron 5 | 71082 |
| 8 | 31937528 | 31938028 | Nrg1 | Intron 5 | 71514 |
| 8 | 31937528 | 31938028 | Nrg1 | Intron 3 | 946430 |
| 8 | 31937528 | 31938028 | Nrg1 | Intron 5 | 945835 |
| 8 | 31937528 | 31938028 | Nrg1 | Intron 3 | 71079 |
| 8 | 31937528 | 31938028 | Nrg1 | Non-coding | 71464 |
| 8 | 31937528 | 31938028 | Nrg1 | Non-coding | 354886 |
| 8 | 34053946 | 34054446 | Gm9951 | Non-coding | 4684 |
| 8 | 34053946 | 34054446 | Gm9951 | Promoter | -176 |
| 8 | 36600946 | 36601446 | Dlc1 | Intron 1 | 12498 |
| 8 | 36600946 | 36601446 | Dlc1 | Intron 1 | 131609 |
| 8 | 36600946 | 36601446 | Dlc1 | Intron 5 | 350997 |
| 8 | 36600946 | 36601446 | Dlc1 | Non-coding | 12429 |
| 8 | 40992646 | 40993146 | Mtus1 | 3' UTR | 140581 |
| 8 | 40992646 | 40993146 | Mtus1 | 3' UTR | 61649 |
| 8 | 40992646 | 40993146 | Mtus1 | 3' UTR | 121522 |
| 8 | 40992646 | 40993146 | Mtus1 | 3' UTR | 48721 |
| 8 | 47867146 | 47867646 | Wwc2 | Intron 11 | 122906 |
| 8 | 48341646 | 48342146 | Tenm3 | Intron 9 | 332545 |
| 8 | 48341646 | 48342146 | Tenm3 | Intron 9 | 332539 |
| 8 | 48341646 | 48342146 | Tenm3 | Intron 8 | 213168 |
| 8 | 49981146 | 49981646 | Gm45323 | Non-coding | 31739 |
| 8 | 54512146 | 54512646 | Gm45553 | Promoter | 1505 |
| 8 | 61517703 | 61518203 | Palld | Intron 8 | 72940 |
| 8 | 61517703 | 61518203 | Palld | Intron 16 | 241887 |
| 8 | 61517703 | 61518203 | Palld | Intron 18 | 384467 |
| 8 | 61517703 | 61518203 | Palld | Promoter | -1802 |
| 8 | 61517703 | 61518203 | Palld | Intron 16 | 384488 |
| 8 | 61517703 | 61518203 | Palld | Intron 8 | 72937 |
| 8 | 64041203 | 64041703 | Tll1 | Intron 17 | 164291 |
| 8 | 64041203 | 64041703 | Tll1 | CDS 17 | 164291 |
| 8 | 64041203 | 64041703 | Tll1 | Intron 16 | 164291 |
| 8 | 70114601 | 70115101 | Ncan | Intron 3 | 5773 |
| 8 | 70114601 | 70115101 | Ncan | CDS 3 | 5773 |
| 8 | 70452101 | 70452601 | Klhl26 | 3' UTR | 24350 |
| 8 | 70452101 | 70452601 | Klhl26 | CDS 3 | 24349 |

| | | | | | |
|---|---|---|---|---|---|
| 8 | 70452101 | 70452601 | Klhl26 | CDS 3 | 24305 |
| 8 | 70452101 | 70452601 | Klhl26 | CDS 2 | 24368 |
| 8 | 70452101 | 70452601 | Klhl26 | CDS 3 | 23298 |
| 8 | 83200101 | 83200601 | Tbc1d9 | Intron 1 | 34749 |
| 8 | 83200101 | 83200601 | Tbc1d9 | Intron 1 | 34241 |
| 8 | 83349601 | 83350101 | Mgat4d | Promoter | 1130 |
| 8 | 83349601 | 83350101 | Mgat4d | Intron 1 | 1130 |
| 8 | 83349601 | 83350101 | Gm15660 | Promoter | -2455 |
| 8 | 83654601 | 83655101 | Gipc1 | Promoter | 1924 |
| 8 | 83654601 | 83655101 | Gipc1 | 5' UTR | 1924 |
| 8 | 83654601 | 83655101 | Gipc1 | Promoter | 1915 |
| 8 | 83654601 | 83655101 | Gipc1 | 5' UTR | 1915 |
| 8 | 83654601 | 83655101 | Gipc1 | Promoter | 1903 |
| 8 | 83654601 | 83655101 | Gipc1 | 5' UTR | 1903 |
| 8 | 84128101 | 84128601 | Podnl1 | Promoter | 2112 |
| 8 | 84128101 | 84128601 | Podnl1 | Intron 5 | 2112 |
| 8 | 84128101 | 84128601 | Podnl1 | CDS 6 | 2112 |
| 8 | 84128101 | 84128601 | Podnl1 | Promoter | 283 |
| 8 | 84351101 | 84351601 | Cacna1a | 5' UTR | 12462 |
| 8 | 84611601 | 84612101 | Cacna1a | CDS 37 | 272962 |
| 8 | 84611601 | 84612101 | Cacna1a | Intron 37 | 272962 |
| 8 | 84556101 | 84556601 | Cacna1a | Intron 19 | 217462 |
| 8 | 84611601 | 84612101 | Cacna1a | CDS 35 | 196237 |
| 8 | 84611601 | 84612101 | Cacna1a | Intron 35 | 196237 |
| 8 | 84556101 | 84556601 | Cacna1a | Intron 17 | 140737 |
| 8 | 84611601 | 84612101 | Cacna1a | CDS 34 | 196224 |
| 8 | 84611601 | 84612101 | Cacna1a | Intron 34 | 196224 |
| 8 | 84556101 | 84556601 | Cacna1a | Intron 16 | 140724 |
| 8 | 84611601 | 84612101 | Cacna1a | CDS 35 | 195817 |
| 8 | 84611601 | 84612101 | Cacna1a | Intron 35 | 195817 |
| 8 | 84556101 | 84556601 | Cacna1a | Intron 17 | 140317 |
| 8 | 84611601 | 84612101 | Cacna1a | CDS 4 | 11107 |
| 8 | 84611601 | 84612101 | Cacna1a | Intron 4 | 11107 |
| 8 | 84611601 | 84612101 | Cacna1a | Non-coding | 10170 |
| 8 | 84611601 | 84612101 | Cacna1a | Non-coding | 10157 |
| 8 | 84611601 | 84612101 | Cacna1a | Non-coding | 10131 |
| 8 | 84611601 | 84612101 | Cacna1a | CDS 2 | 9628 |
| 8 | 84611601 | 84612101 | Cacna1a | Intron 2 | 9628 |
| 8 | 84611601 | 84612101 | Gm16183 | Non-coding | 5084 |
| 8 | 85789601 | 85790101 | Itfg1 | Intron 6 | 50821 |
| 8 | 86848601 | 86849101 | N4bp1 | Promoter | -4555 |
| 8 | 86848601 | 86849101 | N4bp1 | Intron 4 | 36158 |
| 8 | 86848601 | 86849101 | N4bp1 | Non-coding | 3320 |
| 8 | 87587601 | 87588101 | 4933402J07Rik | Intron 5 | 23748 |
| 8 | 91154101 | 91154601 | Aktip | Non-coding | 45376 |
| 8 | 91154101 | 91154601 | Aktip | 5' UTR | 45371 |

| | | | | | |
|---|---|---|---|---|---|
| 8 | 91154101 | 91154601 | Gm45665 | Promoter | 1894 |
| 8 | 91154101 | 91154601 | Gm8489 | Promoter | -3138 |
| 8 | 91505101 | 91505601 | Fto | Intron 7 | 191576 |
| 8 | 91505101 | 91505601 | Fto | Intron 7 | 191543 |
| 8 | 91505101 | 91505601 | Fto | Non-coding | 37416 |
| 8 | 94492100 | 94492600 | Nlrc5 | Intron 22 | 57709 |
| 8 | 94492100 | 94492600 | Nlrc5 | CDS 23 | 57709 |
| 8 | 94492100 | 94492600 | Nlrc5 | Intron 23 | 57709 |
| 8 | 94492100 | 94492600 | Nlrc5 | Intron 19 | 22090 |
| 8 | 94492100 | 94492600 | Nlrc5 | CDS 20 | 22090 |
| 8 | 94492100 | 94492600 | Nlrc5 | Intron 20 | 22090 |
| 8 | 94492100 | 94492600 | Nlrc5 | Intron 18 | 19337 |
| 8 | 94492100 | 94492600 | Nlrc5 | CDS 19 | 19337 |
| 8 | 94492100 | 94492600 | Nlrc5 | Intron 19 | 19337 |
| 8 | 103567600 | 103568100 | Gm45277 | Promoter | -3976 |
| 8 | 105285600 | 105286100 | 4931428F04Rik | 5' UTR | 3765 |
| 8 | 105285600 | 105286100 | 4931428F04Rik | CDS 2 | 3765 |
| 8 | 105285600 | 105286100 | 4931428F04Rik | Intron 1 | 3765 |
| 8 | 105285600 | 105286100 | 4931428F04Rik | Promoter | 2951 |
| 8 | 105285600 | 105286100 | 4931428F04Rik | 5' UTR | 2951 |
| 8 | 105285600 | 105286100 | 4931428F04Rik | CDS 2 | 2951 |
| 8 | 105285600 | 105286100 | 4931428F04Rik | Intron 1 | 2951 |
| 8 | 105285600 | 105286100 | 4931428F04Rik | 5' UTR | 3429 |
| 8 | 105285600 | 105286100 | 4931428F04Rik | CDS 3 | 3429 |
| 8 | 105285600 | 105286100 | 4931428F04Rik | Intron 2 | 3429 |
| 8 | 105285600 | 105286100 | 4931428F04Rik | Promoter | 2979 |
| 8 | 105285600 | 105286100 | 4931428F04Rik | 5' UTR | 2979 |
| 8 | 105285600 | 105286100 | 4931428F04Rik | CDS 2 | 2979 |
| 8 | 105285600 | 105286100 | 4931428F04Rik | Intron 1 | 2979 |
| 8 | 105285600 | 105286100 | Gm20163 | Promoter | -3578 |
| 8 | 105285600 | 105286100 | Gm20163 | Promoter | -3889 |
| 8 | 105336600 | 105337100 | Fhod1 | Intron 10 | 10854 |
| 8 | 105336600 | 105337100 | Fhod1 | CDS 10 | 10854 |
| 8 | 105336600 | 105337100 | Fhod1 | Intron 9 | 10854 |
| 8 | 105336600 | 105337100 | Fhod1 | CDS 9 | 10854 |
| 8 | 105336600 | 105337100 | Fhod1 | Intron 8 | 10854 |
| 8 | 105336600 | 105337100 | Fhod1 | Non-coding | 10810 |
| 8 | 105336600 | 105337100 | Fhod1 | Promoter | 808 |
| 8 | 105961100 | 105961600 | Slc12a4 | Intron 1 | 4445 |
| 8 | 105961100 | 105961600 | Slc12a4 | Intron 1 | 4498 |
| 8 | 105961100 | 105961600 | Slc12a4 | Promoter | 2821 |
| 8 | 105961100 | 105961600 | Slc12a4 | Intron 1 | 2821 |
| 8 | 105961100 | 105961600 | Slc12a4 | Non-coding | 4475 |
| 8 | 111741600 | 111742100 | Bcar1 | Promoter | 1710 |
| 8 | 111741600 | 111742100 | Bcar1 | Intron 1 | 1710 |
| 8 | 114641100 | 114641600 | Wwox | Intron 5 | 201445 |

| | | | | | |
|---|---|---|---|---|---|
| 8 | 115097600 | 115098100 | Wwox | Intron 8 | 657945 |
| 8 | 114641100 | 114641600 | Wwox | Intron 5 | 201405 |
| 8 | 114641100 | 114641600 | Wwox | Intron 5 | 201405 |
| 8 | 114641100 | 114641600 | Wwox | Non-coding | 32664 |
| 8 | 114641100 | 114641600 | Gm16117 | Promoter | 515 |
| 8 | 115097600 | 115098100 | Wwox | Non-coding | 385433 |
| 8 | 117293100 | 117293600 | Cmip | Intron 1 | 36036 |
| 8 | 117293100 | 117293600 | Cmip | Non-coding | 35983 |
| 8 | 118969100 | 118969600 | Cdh13 | Intron 6 | 685367 |
| 8 | 118933600 | 118934100 | Cdh13 | Intron 5 | 649867 |
| 8 | 118484100 | 118484600 | Cdh13 | Intron 1 | 200367 |
| 8 | 118484100 | 118484600 | Cdh13 | Non-coding | 200345 |
| 8 | 118933600 | 118934100 | Cdh13 | Non-coding | 649825 |
| 8 | 118484100 | 118484600 | Cdh13 | Non-coding | 200325 |
| 8 | 118484100 | 118484600 | Cdh13 | Non-coding | 200304 |
| 8 | 118969100 | 118969600 | Cdh13 | Non-coding | 685303 |
| 8 | 118933600 | 118934100 | Cdh13 | Non-coding | 649803 |
| 8 | 118484100 | 118484600 | Cdh13 | Non-coding | 200303 |
| 8 | 118484100 | 118484600 | Cdh13 | Non-coding | 169687 |
| 8 | 120867600 | 120868100 | Gm20388 | Intron 1 | 956759 |
| 8 | 122670100 | 122670600 | Gm20388 | Intron 1 | 2759259 |
| 8 | 122769100 | 122769600 | Gm20388 | Intron 1 | 2858259 |
| 8 | 122727600 | 122728100 | Gm20388 | Intron 1 | 2816759 |
| 8 | 122064600 | 122065100 | Gm20388 | Intron 1 | 2153759 |
| 8 | 122618100 | 122618600 | Gm20388 | Intron 1 | 2707259 |
| 8 | 122255100 | 122255600 | Gm20388 | Intron 1 | 2344259 |
| 8 | 121494100 | 121494600 | Gm20388 | Intron 1 | 1583259 |
| 8 | 121307600 | 121308100 | Gm20388 | Intron 1 | 1396759 |
| 8 | 120501100 | 120501600 | Gm20388 | Intron 1 | 590259 |
| 8 | 120103600 | 120104100 | Gm20388 | Intron 1 | 192759 |
| 8 | 120103600 | 120104100 | Zdhhc7 | Promoter | -2117 |
| 8 | 120103600 | 120104100 | Zdhhc7 | Promoter | -2527 |
| 8 | 120103600 | 120104100 | A130014A01Rik | Promoter | 2018 |
| 8 | 120501100 | 120501600 | Gse1 | Intron 2 | 272644 |
| 8 | 120501100 | 120501600 | Gse1 | Intron 1 | 12653 |
| 8 | 121307600 | 121308100 | Gm26815 | Non-coding | 48737 |
| 8 | 122255100 | 122255600 | Zfp469 | Promoter | -3020 |
| 8 | 122670100 | 122670600 | Cbfa2t3 | Intron 1 | 28510 |
| 8 | 122670100 | 122670600 | Cbfa2t3 | 5' UTR | 7473 |
| 8 | 122670100 | 122670600 | Cbfa2t3 | 5' UTR | 7576 |
| 8 | 122670100 | 122670600 | Cbfa2t3 | 5' UTR | 8095 |
| 8 | 122670100 | 122670600 | Cbfa2t3 | Non-coding | 7512 |
| 8 | 122670100 | 122670600 | Cbfa2t3 | Non-coding | 28303 |
| 8 | 122727600 | 122728100 | C230057M02Rik | Non-coding | 24226 |

| | | | | | |
|---|---|---|---|---|---|
| 8 | 122727600 | 122728100 | C230057M02Rik | Non-coding | 10412 |
| 8 | 127543600 | 127544100 | Pard3 | Intron 22 | 479707 |
| 8 | 127543600 | 127544100 | Pard3 | Intron 22 | 479544 |
| 8 | 127543600 | 127544100 | Pard3 | Intron 21 | 479355 |
| 8 | 127543600 | 127544100 | Pard3 | Intron 21 | 479305 |
| 8 | 127543600 | 127544100 | Pard3 | Intron 22 | 479305 |
| 8 | 127543600 | 127544100 | Pard3 | Intron 20 | 479255 |
| 9 | 9236500 | 9237000 | Arhgap42 | Promoter | 2102 |
| 9 | 9236500 | 9237000 | Arhgap42 | Intron 1 | 2102 |
| 9 | 9236500 | 9237000 | Arhgap42 | Promoter | 2102 |
| 9 | 9236500 | 9237000 | Arhgap42 | Promoter | 1901 |
| 9 | 9236500 | 9237000 | Arhgap42 | Intron 1 | 1901 |
| 9 | 9236500 | 9237000 | Arhgap42 | Promoter | 2027 |
| 9 | 9236500 | 9237000 | Arhgap42 | Intron 1 | 2027 |
| 9 | 9236500 | 9237000 | Gm16833 | Promoter | 212 |
| 9 | 14090056 | 14090556 | 1700019J19Rik | Promoter | 436 |
| 9 | 14090056 | 14090556 | 1700019J19Rik | Promoter | 0 |
| 9 | 14090056 | 14090556 | 1700019J19Rik | Promoter | 0 |
| 9 | 14274556 | 14275056 | Sesn3 | Promoter | -11 |
| 9 | 14274556 | 14275056 | Sesn3 | Promoter | -27 |
| 9 | 14274556 | 14275056 | Sesn3 | Promoter | -1245 |
| 9 | 14274556 | 14275056 | Sesn3 | Promoter | -1245 |
| 9 | 14602556 | 14603056 | Amotl1 | Intron 1 | 11951 |
| 9 | 14602556 | 14603056 | Amotl1 | Non-coding | 40474 |
| 9 | 14602556 | 14603056 | Amotl1 | Intron 2 | 42001 |
| 9 | 14602556 | 14603056 | Amotl1 | 5' UTR | 12428 |
| 9 | 14602556 | 14603056 | Amotl1 | Non-coding | 11531 |
| 9 | 15003556 | 15004056 | Panx1 | 3' UTR | 41423 |
| 9 | 15422056 | 15422556 | Gm2517 | Promoter | 2059 |
| 9 | 25555415 | 25555915 | Eepd1 | Intron 3 | 73868 |
| 9 | 35007915 | 35008415 | Kirrel3 | Intron 8 | 521789 |
| 9 | 35007915 | 35008415 | Kirrel3 | Intron 7 | 519182 |
| 9 | 35007915 | 35008415 | Kirrel3 | Intron 8 | 519120 |
| 9 | 35007915 | 35008415 | Kirrel3 | Intron 7 | 519120 |
| 9 | 35007915 | 35008415 | Kirrel3 | Intron 7 | 519087 |
| 9 | 35007915 | 35008415 | Kirrel3 | Intron 6 | 518979 |
| 9 | 37368415 | 37368915 | Hepacam | Promoter | 1064 |
| 9 | 37368415 | 37368915 | Hepacam | Intron 1 | 1064 |
| 9 | 37368415 | 37368915 | Hepacam | Promoter | 779 |
| 9 | 37368415 | 37368915 | Hepacam | Intron 1 | 779 |
| 9 | 37429915 | 37430415 | Robo3 | Promoter | 0 |
| 9 | 37429915 | 37430415 | Robo3 | 5' UTR | 0 |
| 9 | 37429915 | 37430415 | Robo3 | Promoter | 2832 |
| 9 | 37429915 | 37430415 | Robo3 | Intron 1 | 2832 |
| 9 | 37429915 | 37430415 | Robo3 | Promoter | 0 |

| | | | | | |
|---|---|---|---|---|---|
| 9 | 37429915 | 37430415 | Robo3 | 5' UTR | 0 |
| 9 | 40414415 | 40414915 | Gramd1b | Intron 1 | 40850 |
| 9 | 40414415 | 40414915 | Gramd1b | Intron 1 | 116469 |
| 9 | 40414415 | 40414915 | Gramd1b | Intron 1 | 40756 |
| 9 | 40414415 | 40414915 | Gramd1b | Non-coding | 50650 |
| 9 | 40414415 | 40414915 | Gramd1b | Non-coding | 50640 |
| 9 | 41612417 | 41612917 | Mir100hg | Non-coding | 285160 |
| 9 | 41612417 | 41612917 | Mir100hg | Non-coding | 236107 |
| 9 | 41612417 | 41612917 | Mir100hg | Non-coding | 236096 |
| 9 | 41612417 | 41612917 | Gm49317 | Non-coding | 137562 |
| 9 | 41612417 | 41612917 | Mir100hg | Non-coding | 137562 |
| 9 | 41612417 | 41612917 | Mir100hg | Non-coding | 137480 |
| 9 | 41612417 | 41612917 | Mir100hg | Non-coding | 31164 |
| 9 | 41612417 | 41612917 | Mir100hg | Non-coding | 31150 |
| 9 | 41612417 | 41612917 | Mir100hg | Non-coding | 31084 |
| 9 | 41612417 | 41612917 | Mir100hg | Non-coding | 31078 |
| 9 | 41612417 | 41612917 | Mir100hg | Non-coding | 31076 |
| 9 | 41612417 | 41612917 | Mir100hg | Non-coding | 31056 |
| 9 | 41612417 | 41612917 | Mir100hg | Non-coding | 12145 |
| 9 | 43222417 | 43222917 | Oaf | 3' UTR | 16995 |
| 9 | 43222417 | 43222917 | Oaf | CDS 4 | 16995 |
| 9 | 43222417 | 43222917 | Oaf | Intron 3 | 16995 |
| 9 | 43222417 | 43222917 | Oaf | Promoter | 1058 |
| 9 | 43222417 | 43222917 | Oaf | 3' UTR | 1058 |
| 9 | 43222417 | 43222917 | Oaf | CDS 2 | 1058 |
| 9 | 43222417 | 43222917 | Oaf | Intron 1 | 1058 |
| 9 | 43222417 | 43222917 | Oaf | Promoter | 2176 |
| 9 | 44537417 | 44537917 | Cxcr5 | 5' UTR | 23961 |
| 9 | 44537417 | 44537917 | Cxcr5 | 5' UTR | 18475 |
| 9 | 46098917 | 46099417 | Sik3 | Intron 1 | 86097 |
| 9 | 46198917 | 46199417 | Sik3 | Intron 13 | 186097 |
| 9 | 46098917 | 46099417 | Sik3 | Intron 1 | 86053 |
| 9 | 46198917 | 46199417 | Sik3 | Intron 13 | 186053 |
| 9 | 46098917 | 46099417 | Sik3 | Intron 1 | 85917 |
| 9 | 46198917 | 46199417 | Sik3 | Intron 12 | 185917 |
| 9 | 46198917 | 46199417 | Sik3 | Intron 12 | 75680 |
| 9 | 46198917 | 46199417 | Sik3 | Non-coding | 43538 |
| 9 | 46909417 | 46909917 | 2900052N01Rik | Promoter | -3576 |
| 9 | 46909417 | 46909917 | 2900052N01Rik | Promoter | -3686 |
| 9 | 46909417 | 46909917 | 2900052N01Rik | Promoter | -3710 |
| 9 | 47253917 | 47254417 | Gm31816 | Non-coding | 16327 |
| 9 | 48716395 | 48716895 | Zbtb16 | Intron 4 | 119051 |
| 9 | 48716395 | 48716895 | Zbtb16 | Intron 4 | 119328 |

| | | | | | |
|---|---|---|---|---|---|
| 9 | 49692895 | 49693395 | Ncam1 | Intron 1 | 105531 |
| 9 | 49692895 | 49693395 | Ncam1 | Intron 1 | 105478 |
| 9 | 49692895 | 49693395 | Ncam1 | Intron 1 | 105496 |
| 9 | 49692895 | 49693395 | Ncam1 | Intron 1 | 105364 |
| 9 | 49692895 | 49693395 | Ncam1 | Non-coding | 105528 |
| 9 | 50707895 | 50708395 | Dixdc1 | Intron 2 | 19581 |
| 9 | 50707895 | 50708395 | Dixdc1 | Intron 2 | 19673 |
| 9 | 50707895 | 50708395 | Dixdc1 | Intron 3 | 31123 |
| 9 | 51293395 | 51293895 | 1810046K07Rik | Intron 3 | 35023 |
| 9 | 51293395 | 51293895 | 1810046K07Rik | Intron 2 | 18821 |
| 9 | 51293395 | 51293895 | 1810046K07Rik | Intron 3 | 35030 |
| 9 | 52430395 | 52430895 | Gm33125 | Non-coding | 28299 |
| 9 | 58310193 | 58310693 | Loxl1 | Promoter | 2494 |
| 9 | 58310193 | 58310693 | Loxl1 | Intron 1 | 2494 |
| 9 | 63374693 | 63375193 | Map2k5 | Promoter | 2710 |
| 9 | 63374693 | 63375193 | Map2k5 | Intron 1 | 2710 |
| 9 | 63374693 | 63375193 | Map2k5 | Promoter | 2685 |
| 9 | 65128693 | 65129193 | Igdcc4 | CDS 11 | 27207 |
| 9 | 65128693 | 65129193 | Igdcc4 | Intron 11 | 27207 |
| 9 | 65128693 | 65129193 | Igdcc4 | CDS 12 | 27207 |
| 9 | 65128693 | 65129193 | Igdcc4 | Intron 12 | 27207 |
| 9 | 65128693 | 65129193 | Igdcc4 | CDS 14 | 27207 |
| 9 | 65128693 | 65129193 | Igdcc4 | Intron 14 | 27207 |
| 9 | 65128693 | 65129193 | Igdcc4 | CDS 15 | 27207 |
| 9 | 65128693 | 65129193 | Igdcc4 | Intron 15 | 27207 |
| 9 | 65128693 | 65129193 | Igdcc4 | CDS 11 | 27207 |
| 9 | 65128693 | 65129193 | Igdcc4 | Intron 11 | 27207 |
| 9 | 65128693 | 65129193 | Igdcc4 | CDS 12 | 27207 |
| 9 | 65128693 | 65129193 | Igdcc4 | Intron 12 | 27207 |
| 9 | 65128693 | 65129193 | Igdcc4 | Intron 2 | 27158 |
| 9 | 65128693 | 65129193 | Igdcc4 | Non-coding | 27158 |
| 9 | 66232693 | 66233193 | Dapk2 | Intron 4 | 74470 |
| 9 | 66232693 | 66233193 | Dapk2 | Intron 4 | 74467 |
| 9 | 66232693 | 66233193 | Dapk2 | Intron 2 | 11946 |
| 9 | 67182693 | 67183193 | Gm19299 | Non-coding | 49133 |
| 9 | 69101193 | 69101693 | Rora | Intron 1 | 447407 |
| 9 | 69101193 | 69101693 | Rora | Intron 1 | 447302 |
| 9 | 70930193 | 70930693 | Lipc | Intron 1 | 4116 |
| 9 | 70930193 | 70930693 | Lipc | 5' UTR | 21534 |
| 9 | 70930193 | 70930693 | Lipc | Intron 1 | 3921 |
| 9 | 70930193 | 70930693 | Lipc | Non-coding | 21529 |
| 9 | 70930193 | 70930693 | Gm32017 | Promoter | 0 |
| 9 | 71760693 | 71761193 | Cgnl1 | 5' UTR | 10410 |
| 9 | 71666193 | 71666693 | Cgnl1 | Intron 8 | 104910 |
| 9 | 71711193 | 71711693 | Cgnl1 | Intron 7 | 59910 |
| 9 | 71760693 | 71761193 | Cgnl1 | 5' UTR | 10410 |

| | | | | | |
|---|---|---|---|---|---|
| 9 | 71666193 | 71666693 | Cgnl1 | Intron 7 | 104910 |
| 9 | 71711193 | 71711693 | Cgnl1 | Intron 7 | 59910 |
| 9 | 71760693 | 71761193 | Cgnl1 | 5' UTR | 10368 |
| 9 | 71666193 | 71666693 | Cgnl1 | Intron 8 | 104868 |
| 9 | 71711193 | 71711693 | Cgnl1 | Intron 7 | 59868 |
| 9 | 74276693 | 74277193 | Wdr72 | Intron 19 | 166337 |
| 9 | 74276693 | 74277193 | Wdr72 | Intron 19 | 166337 |
| 9 | 74276693 | 74277193 | Wdr72 | Promoter | 2559 |
| 9 | 75774193 | 75774693 | Bmp5 | Promoter | -671 |
| 9 | 77496693 | 77497193 | Lrrc1 | Intron 2 | 47678 |
| 9 | 77435193 | 77435693 | Lrrc1 | CDS 12 | 109178 |
| 9 | 77435193 | 77435693 | Lrrc1 | Intron 11 | 109178 |
| 9 | 77496693 | 77497193 | Lrrc1 | Intron 2 | 46903 |
| 9 | 77435193 | 77435693 | Lrrc1 | CDS 12 | 108403 |
| 9 | 77435193 | 77435693 | Lrrc1 | Intron 11 | 108403 |
| 9 | 77435193 | 77435693 | Lrrc1 | CDS 2 | 6503 |
| 9 | 77435193 | 77435693 | Lrrc1 | Intron 1 | 6503 |
| 9 | 77435193 | 77435693 | Lrrc1 | Non-coding | 6799 |
| 9 | 77496693 | 77497193 | Lrrc1 | Intron 2 | 47658 |
| 9 | 77496693 | 77497193 | Lrrc1 | Non-coding | 35252 |
| 9 | 77496693 | 77497193 | Lrrc1 | Non-coding | 47656 |
| 9 | 78623193 | 78623693 | Cd109 | Intron 2 | 7647 |
| 9 | 79938193 | 79938693 | Filip1 | 5' UTR | 38967 |
| 9 | 79938193 | 79938693 | Filip1 | 5' UTR | 39112 |
| 9 | 79938193 | 79938693 | Filip1 | 5' UTR | 23341 |
| 9 | 79938193 | 79938693 | Filip1 | 5' UTR | 74159 |
| 9 | 80344693 | 80345193 | Impg1 | Intron 12 | 120246 |
| 9 | 80344693 | 80345193 | Impg1 | CDS 12 | 120246 |
| 9 | 80344693 | 80345193 | Impg1 | Intron 13 | 120289 |
| 9 | 80344693 | 80345193 | Impg1 | CDS 13 | 120289 |
| 9 | 83565393 | 83565893 | Sh3bgrl2 | Intron 1 | 17066 |
| 9 | 83565393 | 83565893 | Sh3bgrl2 | Intron 1 | 17065 |
| 9 | 83565393 | 83565893 | Sh3bgrl2 | Intron 1 | 17043 |
| 9 | 85823893 | 85824393 | 9330154J02Rik | Non-coding | 18893 |
| 9 | 85823893 | 85824393 | 9330154J02Rik | Non-coding | 19739 |
| 9 | 85823893 | 85824393 | 9330154J02Rik | Non-coding | 19931 |
| 9 | 85823893 | 85824393 | 9330154J02Rik | Non-coding | 18892 |
| 9 | 86549393 | 86549893 | Dop1a | Intron 35 | 82239 |
| 9 | 86549393 | 86549893 | Dop1a | 3' UTR | 82179 |
| 9 | 86549393 | 86549893 | Dop1a | Intron 34 | 63986 |
| 9 | 86549393 | 86549893 | Dop1a | Non-coding | 17816 |
| 9 | 86549393 | 86549893 | Dop1a | 3' UTR | 16390 |
| 9 | 86549393 | 86549893 | Gm17530 | Promoter | -2239 |
| 9 | 86549393 | 86549893 | Gm17530 | Promoter | -2597 |
| 9 | 86549393 | 86549893 | Dop1a | Promoter | 0 |
| 9 | 86594893 | 86595393 | Me1 | Intron 11 | 100561 |

| | | | | | |
|---|---|---|---|---|---|
| 9 | 86594893 | 86595393 | Me1 | Intron 11 | 91885 |
| 9 | 86594893 | 86595393 | Me1 | Non-coding | 23312 |
| 9 | 87145665 | 87146165 | Mrap2 | Promoter | 1359 |
| 9 | 87145665 | 87146165 | Mrap2 | 5' UTR | 1359 |
| 9 | 87145665 | 87146165 | Mrap2 | Promoter | 1359 |
| 9 | 87145665 | 87146165 | Mrap2 | 5' UTR | 1359 |
| 9 | 87145665 | 87146165 | Mrap2 | Promoter | 1322 |
| 9 | 87145665 | 87146165 | Mrap2 | 5' UTR | 1322 |
| 9 | 91538162 | 91538662 | Gm29478 | Non-coding | 133688 |
| 9 | 95073081 | 95073581 | Slc9a9 | Intron 12 | 403172 |
| 9 | 95564581 | 95565081 | Paqr9 | 3' UTR | 4924 |
| 9 | 97043081 | 97043581 | Gm18291 | Promoter | -218 |
| 9 | 98429581 | 98430081 | Rbp1 | Intron 2 | 6620 |
| 9 | 98496081 | 98496581 | Rbp2 | Intron 2 | 9966 |
| 9 | 98496081 | 98496581 | Rbp2 | Intron 2 | 9915 |
| 9 | 98496081 | 98496581 | Rbp2 | Intron 1 | 5544 |
| 9 | 98496081 | 98496581 | Rbp2 | Non-coding | 4384 |
| 9 | 98496081 | 98496581 | Gm37113 | Promoter | 2113 |
| 9 | 104377170 | 104377670 | Gm28548 | Non-coding | 15338 |
| 9 | 108106669 | 108107169 | Bsn | CDS 6 | 83216 |
| 9 | 108106669 | 108107169 | Bsn | CDS 2 | 3245 |
| 9 | 108106669 | 108107169 | Gm47303 | Promoter | -3422 |
| 9 | 108480169 | 108480669 | Lamb2 | Promoter | 433 |
| 9 | 108480169 | 108480669 | Lamb2 | Intron 3 | 433 |
| 9 | 108480169 | 108480669 | Lamb2 | CDS 4 | 433 |
| 9 | 108480169 | 108480669 | Lamb2 | Promoter | 312 |
| 9 | 108480169 | 108480669 | Lamb2 | Intron 2 | 312 |
| 9 | 108480169 | 108480669 | Lamb2 | CDS 3 | 312 |
| 9 | 108480169 | 108480669 | Lamb2 | Intron 3 | 312 |
| 9 | 108480169 | 108480669 | Lamb2 | CDS 4 | 312 |
| 9 | 108480169 | 108480669 | Lamb2 | Promoter | 297 |
| 9 | 108480169 | 108480669 | Lamb2 | Intron 2 | 297 |
| 9 | 108480169 | 108480669 | Lamb2 | CDS 3 | 297 |
| 9 | 108480169 | 108480669 | Lamb2 | Intron 3 | 297 |
| 9 | 108480169 | 108480669 | Lamb2 | CDS 4 | 297 |
| 9 | 108480169 | 108480669 | Lamb2 | Promoter | 291 |
| 9 | 108480169 | 108480669 | Lamb2 | Intron 2 | 291 |
| 9 | 108480169 | 108480669 | Lamb2 | CDS 3 | 291 |
| 9 | 108480169 | 108480669 | Lamb2 | Intron 3 | 291 |
| 9 | 108480169 | 108480669 | Lamb2 | CDS 4 | 291 |
| 9 | 108480169 | 108480669 | Lamb2 | Promoter | 288 |
| 9 | 110727496 | 110727996 | Pth1r | CDS 8 | 19127 |
| 9 | 110727496 | 110727996 | Pth1r | Intron 7 | 19127 |
| 9 | 110727496 | 110727996 | Pth1r | CDS 7 | 19127 |
| 9 | 110727496 | 110727996 | Pth1r | Intron 6 | 19127 |
| 9 | 110727496 | 110727996 | Pth1r | CDS 6 | 19127 |

| | | | | | |
|---|---|---|---|---|---|
| 9 | 110727496 | 110727996 | Pth1r | CDS 7 | 15691 |
| 9 | 110727496 | 110727996 | Pth1r | Intron 6 | 15691 |
| 9 | 110727496 | 110727996 | Pth1r | CDS 6 | 15691 |
| 9 | 110727496 | 110727996 | Pth1r | Intron 5 | 15691 |
| 9 | 110727496 | 110727996 | Pth1r | CDS 5 | 15691 |
| 9 | 110727496 | 110727996 | Pth1r | CDS 6 | 14724 |
| 9 | 110727496 | 110727996 | Pth1r | Intron 5 | 14724 |
| 9 | 110727496 | 110727996 | Pth1r | CDS 5 | 14724 |
| 9 | 110727496 | 110727996 | Pth1r | Intron 4 | 14724 |
| 9 | 110727496 | 110727996 | Pth1r | CDS 4 | 14724 |
| 9 | 110727496 | 110727996 | Pth1r | CDS 7 | 19061 |
| 9 | 110727496 | 110727996 | Pth1r | Intron 6 | 19061 |
| 9 | 110727496 | 110727996 | Pth1r | CDS 6 | 19061 |
| 9 | 110727496 | 110727996 | Pth1r | Intron 5 | 19061 |
| 9 | 110727496 | 110727996 | Pth1r | CDS 5 | 19061 |
| 9 | 110727496 | 110727996 | Pth1r | CDS 7 | 15653 |
| 9 | 110727496 | 110727996 | Pth1r | Intron 6 | 15653 |
| 9 | 110727496 | 110727996 | Pth1r | CDS 6 | 15653 |
| 9 | 110727496 | 110727996 | Pth1r | Intron 5 | 15653 |
| 9 | 110727496 | 110727996 | Pth1r | CDS 5 | 15653 |
| 9 | 112187496 | 112187996 | Arpp21 | Promoter | -1546 |
| 9 | 112187496 | 112187996 | Arpp21 | Promoter | 0 |
| 9 | 112187496 | 112187996 | Arpp21 | 5' UTR | 0 |
| 9 | 112187496 | 112187996 | Arpp21 | Promoter | 0 |
| 9 | 112187496 | 112187996 | Arpp21 | 5' UTR | 0 |
| 9 | 112187496 | 112187996 | Arpp21 | Promoter | -1726 |
| 9 | 112187496 | 112187996 | Arpp21 | Non-coding | 46428 |
| 9 | 112187496 | 112187996 | Arpp21 | 5' UTR | 47023 |
| 9 | 112187496 | 112187996 | Arpp21 | 5' UTR | 44506 |
| 9 | 112187496 | 112187996 | Arpp21 | Promoter | 0 |
| 9 | 112187496 | 112187996 | Arpp21 | 5' UTR | 0 |
| 9 | 112187496 | 112187996 | Arpp21 | 5' UTR | 29266 |
| 9 | 112187496 | 112187996 | Arpp21 | Non-coding | 46846 |
| 9 | 112187496 | 112187996 | Arpp21 | 5' UTR | 47023 |
| 9 | 112187496 | 112187996 | Arpp21 | 5' UTR | 29356 |
| 9 | 112187496 | 112187996 | Arpp21 | Non-coding | 45631 |
| 9 | 112187496 | 112187996 | Arpp21 | 5' UTR | 29331 |
| 9 | 112187496 | 112187996 | Arpp21 | 5' UTR | 29331 |
| 9 | 112187496 | 112187996 | Arpp21 | 5' UTR | 45820 |
| 9 | 112187496 | 112187996 | Arpp21 | Promoter | 0 |
| 9 | 112187496 | 112187996 | Arpp21 | 5' UTR | 0 |
| 9 | 112187496 | 112187996 | Arpp21 | Promoter | 0 |
| 9 | 112187496 | 112187996 | Arpp21 | 5' UTR | 0 |
| 9 | 112187496 | 112187996 | Arpp21 | Promoter | 0 |
| 9 | 112187496 | 112187996 | Arpp21 | Promoter | 0 |
| 9 | 112187496 | 112187996 | Arpp21 | Promoter | 0 |

| | | | | | |
|---|---|---|---|---|---|
| 9 | 112187496 | 112187996 | Arpp21 | Promoter | 0 |
| 9 | 112187496 | 112187996 | Arpp21 | Promoter | -1402 |
| 9 | 112187496 | 112187996 | Arpp21 | Promoter | -1546 |
| 9 | 112187496 | 112187996 | Arpp21 | Promoter | -1559 |
| 9 | 112187496 | 112187996 | Arpp21 | Non-coding | 43907 |
| 9 | 112187496 | 112187996 | Arpp21 | Non-coding | 45877 |
| 9 | 112187496 | 112187996 | Arpp21 | Non-coding | 47943 |
| 9 | 112187496 | 112187996 | Arpp21 | Promoter | 445 |
| 9 | 112187496 | 112187996 | Arpp21 | Promoter | 479 |
| 9 | 113928382 | 113928882 | Ubp1 | Promoter | -2052 |
| 9 | 113928382 | 113928882 | Ubp1 | Promoter | -2052 |
| 9 | 113928382 | 113928882 | Ubp1 | Promoter | -2553 |
| 9 | 113928382 | 113928882 | Ubp1 | Promoter | -2682 |
| 9 | 113928382 | 113928882 | Ubp1 | Promoter | -2747 |
| 9 | 113928382 | 113928882 | Ubp1 | Promoter | -2772 |
| 9 | 114530382 | 114530882 | Trim71 | Intron 1 | 33488 |
| 9 | 115301382 | 115301882 | Stt3b | Intron 1 | 8540 |
| 9 | 118921382 | 118921882 | Gm2415 | Non-coding | 4378 |
| 9 | 118921382 | 118921882 | Gm2415 | Non-coding | 4388 |
| 9 | 118921382 | 118921882 | Gm2415 | Non-coding | 4377 |
| 9 | 118921382 | 118921882 | Gm16295 | Promoter | -2991 |
| 9 | 118921382 | 118921882 | Ctdspl | Promoter | -4571 |
| 9 | 118921382 | 118921882 | Ctdspl | Promoter | -4980 |
| 9 | 119533882 | 119534382 | Scn5a | CDS 11 | 28297 |
| 9 | 119533882 | 119534382 | Scn5a | Intron 10 | 28297 |
| 9 | 119560882 | 119561382 | Scn5a | Promoter | 1297 |
| 9 | 119560882 | 119561382 | Scn5a | Intron 1 | 1297 |
| 9 | 119533882 | 119534382 | Scn5a | CDS 12 | 44635 |
| 9 | 119533882 | 119534382 | Scn5a | Intron 11 | 44635 |
| 9 | 119560882 | 119561382 | Scn5a | Intron 2 | 17635 |
| 9 | 119533882 | 119534382 | Scn5a | CDS 12 | 44635 |
| 9 | 119533882 | 119534382 | Scn5a | Intron 11 | 44635 |
| 9 | 119560882 | 119561382 | Scn5a | Intron 2 | 17635 |
| 9 | 119560882 | 119561382 | Scn5a | Promoter | 2122 |
| 9 | 119821882 | 119822382 | Scn11a | Promoter | 1479 |
| 9 | 119821882 | 119822382 | Scn11a | 5' UTR | 1479 |
| 9 | 119821882 | 119822382 | Scn11a | 5' UTR | 3075 |
| 9 | 121466882 | 121467382 | Trak1 | 3' UTR | 169380 |
| 9 | 121466882 | 121467382 | Trak1 | Intron 15 | 169380 |
| 9 | 121466882 | 121467382 | Trak1 | CDS 16 | 169380 |
| 9 | 121466882 | 121467382 | Trak1 | Intron 16 | 169380 |
| 9 | 121466882 | 121467382 | Trak1 | Intron 15 | 169370 |
| 9 | 121466882 | 121467382 | Trak1 | Intron 15 | 169370 |
| 9 | 121466882 | 121467382 | Trak1 | Intron 14 | 99866 |
| 9 | 121466882 | 121467382 | Trak1 | Intron 2 | 12586 |
| X | 53883823 | 53884323 | Atf1-ps | Promoter | 98 |

| | | | | | | |
|---|---|---|---|---|---|---|
| X | 71666825 | 71667325 | Gpr50 | CDS 2 | | 3158 |
| X | 71666825 | 71667325 | Gpr50 | Promoter | | 2859 |
| X | 71666825 | 71667325 | Gpr50 | Promoter | | 2859 |
| X | 74916161 | 74916661 | Xlr3e-ps | Non-coding | | 3744 |
| X | 84001161 | 84001661 | Dmd | Intron 43 | | 1052258 |
| X | 84001161 | 84001661 | Dmd | Intron 15 | | 145754 |
| X | 137203961 | 137204461 | Gm8036 | Promoter | | 2303 |
| 1 | 161442369 | 161442869 | Tnfsf4 | Intergenic | ? | |
| 1 | 60862656 | 60863156 | Gm38137 | Intergenic | ? | |
| 1 | 63784926 | 63785426 | 4933402D24Rik | Intergenic | ? | |
| 1 | 50386656 | 50387156 | Gm29665 | Intergenic | ? | |
| 1 | 40405155 | 40405655 | Il1rl1 | Intergenic | ? | |
| 1 | 91650423 | 91650923 | Gm28380 | Intergenic | ? | |
| 1 | 13104919 | 13105419 | Prdm14 | Intergenic | ? | |
| 1 | 92436923 | 92437423 | Ndufa10 | Intergenic | ? | |
| 1 | 128826423 | 128826923 | Gm26686 | Intergenic | ? | |
| 1 | 157803869 | 157804369 | Gm38256 | Intergenic | ? | |
| 1 | 177517869 | 177518369 | Gm37306 | Intergenic | ? | |
| 1 | 92754923 | 92755423 | Gm29482 | Intergenic | ? | |
| 1 | 177294869 | 177295369 | Gm38328 | Intergenic | ? | |
| 1 | 76251425 | 76251925 | Gm29068 | Intergenic | ? | |
| 1 | 152790370 | 152790870 | Ncf2 | Intergenic | ? | |
| 1 | 67097926 | 67098426 | Cps1 | Intergenic | ? | |
| 1 | 120963923 | 120964423 | Gm37043 | Intergenic | ? | |
| 1 | 128456423 | 128456923 | Gm23902 | Intergenic | ? | |
| 1 | 132292423 | 132292923 | Klhdc8a | Intergenic | ? | |
| 1 | 146106370 | 146106870 | Gm29514 | Intergenic | ? | |
| 1 | 133380423 | 133380923 | Sox13 | Intergenic | ? | |
| 1 | 161648369 | 161648869 | Gm37638 | Intergenic | ? | |
| 1 | 168879369 | 168879869 | Mir6354 | Intergenic | ? | |
| 1 | 193702806 | 193703306 | Gm21362 | Intergenic | ? | |
| 1 | 62647926 | 62648426 | Gm11600 | Intergenic | ? | |
| 1 | 78081425 | 78081925 | Pax3 | Intergenic | ? | |
| 10 | 24718194 | 24718694 | Enpp1 | Intergenic | ? | |
| 10 | 69306252 | 69306752 | Gm40685 | Intergenic | ? | |
| 10 | 8130702 | 8131202 | Gm30906 | Intergenic | ? | |
| 10 | 9302702 | 9303202 | Gm48748 | Intergenic | ? | |
| 10 | 42991194 | 42991694 | Sobp | Intergenic | ? | |
| 10 | 66618252 | 66618752 | Gm28139 | Intergenic | ? | |
| 10 | 85037755 | 85038255 | Ric8b | Intergenic | ? | |
| 10 | 126770944 | 126771444 | 4930484H19Rik | Intergenic | ? | |
| 10 | 61831252 | 61831752 | Col13a1 | Intergenic | ? | |
| 10 | 120140944 | 120141444 | Irak3 | Intergenic | ? | |
| 10 | 91714755 | 91715255 | Gm47091 | Intergenic | ? | |

| | | | | | |
|---|---|---|---|---|---|
| 10 | 33011694 | 33012194 | Trdn | Intergenic | ? |
| 10 | 67756252 | 67756752 | 4930543I11Rik | Intergenic | ? |
| 10 | 70060252 | 70060752 | Ccdc6 | Intergenic | ? |
| 10 | 74923255 | 74923755 | Rsph14 | Intergenic | ? |
| 10 | 104933944 | 104934444 | Gm48184 | Intergenic | ? |
| 10 | 117456444 | 117456944 | Gm32141 | Intergenic | ? |
| 10 | 61830752 | 61831252 | Col13a1 | Intergenic | ? |
| 11 | 63463498 | 63463998 | B130011K05Rik | Intergenic | ? |
| 11 | 45798998 | 45799498 | F630206G17Rik | Intergenic | ? |
| 11 | 63463998 | 63464498 | B130011K05Rik | Intergenic | ? |
| 11 | 114490186 | 114490686 | 4932435O22Rik | Intergenic | ? |
| 11 | 102335186 | 102335686 | Slc4a1 | Intergenic | ? |
| 11 | 100276686 | 100277186 | Gm14206 | Intergenic | ? |
| 11 | 82448498 | 82448998 | Tmem132e | Intergenic | ? |
| 11 | 102978686 | 102979186 | 2410004I01Rik | Intergenic | ? |
| 11 | 6538497 | 6538997 | Gm24000 | Intergenic | ? |
| 11 | 8107997 | 8108497 | Gm11988 | Intergenic | ? |
| 11 | 18181997 | 18182497 | Gm12019 | Intergenic | ? |
| 11 | 18182497 | 18182997 | Gm12019 | Intergenic | ? |
| 11 | 38913498 | 38913998 | Gm23520 | Intergenic | ? |
| 11 | 97590186 | 97590686 | 2410003L11Rik | Intergenic | ? |
| 11 | 26960000 | 26960500 | Gm12074 | Intergenic | ? |
| 11 | 92889686 | 92890186 | Gm25113 | Intergenic | ? |
| 11 | 51889998 | 51890498 | Gm39822 | Intergenic | ? |
| 11 | 59926498 | 59926998 | 1810063I02Rik | Intergenic | ? |
| 11 | 22376997 | 22377497 | Rpsa-ps3 | Intergenic | ? |
| 11 | 102725186 | 102725686 | Gm11637 | Intergenic | ? |
| 11 | 68911498 | 68911998 | Gm22608 | Intergenic | ? |
| 11 | 70075998 | 70076498 | Asgr2 | Intergenic | ? |
| 11 | 87749498 | 87749998 | Mir142hg | Intergenic | ? |
| 11 | 102347686 | 102348186 | Slc4a1 | Intergenic | ? |
| 11 | 105145186 | 105145686 | Gm24290 | Intergenic | ? |
| 11 | 109733186 | 109733686 | Gm11684 | Intergenic | ? |
| 11 | 109327686 | 109328186 | 1700096J18Rik | Intergenic | ? |
| 11 | 45746998 | 45747498 | F630206G17Rik | Intergenic | ? |
| 11 | 52724998 | 52725498 | Fstl4 | Intergenic | ? |
| 11 | 71801998 | 71802498 | Wscd1 | Intergenic | ? |
| 11 | 13040997 | 13041497 | Gm12004 | Intergenic | ? |
| 11 | 83788998 | 83789498 | Heatr6 | Intergenic | ? |
| 11 | 112528686 | 112529186 | BC006965 | Intergenic | ? |
| 12 | 28590635 | 28591135 | Colec11 | Intergenic | ? |

| | | | | | |
|---|---|---|---|---|---|
| 12 | 35895913 | 35896413 | Agr3 | Intergenic | ? |
| 12 | 50974513 | 50975013 | Gm40421 | Intergenic | ? |
| 12 | 28590135 | 28590635 | Colec11 | Intergenic | ? |
| 12 | 35186828 | 35187328 | Mir680-3 | Intergenic | ? |
| 12 | 53220513 | 53221013 | Npas3 | Intergenic | ? |
| 12 | 17184194 | 17184694 | Kcnf1 | Intergenic | ? |
| 12 | 35283828 | 35284328 | Gm44396 | Intergenic | ? |
| 12 | 106528790 | 106529290 | Gm17053 | Intergenic | ? |
| 12 | 25248635 | 25249135 | Gm19340 | Intergenic | ? |
| 12 | 51492513 | 51493013 | Gm18503 | Intergenic | ? |
| 12 | 51848013 | 51848513 | Gm19309 | Intergenic | ? |
| 12 | 28899135 | 28899635 | Gm31508 | Intergenic | ? |
| 12 | 32893135 | 32893635 | Gm47948 | Intergenic | ? |
| 12 | 20845139 | 20845639 | 3110053B16Rik | Intergenic | ? |
| 12 | 12571694 | 12572194 | AC107703.1 | Intergenic | ? |
| 12 | 118812027 | 118812527 | Gm6171 | Intergenic | ? |
| 12 | 17259194 | 17259694 | Pdia6 | Intergenic | ? |
| 12 | 100538290 | 100538790 | Rps6ka5 | Intergenic | ? |
| 12 | 106928290 | 106928790 | Gm16087 | Intergenic | ? |
| 12 | 29449635 | 29450135 | Gm20208 | Intergenic | ? |
| 12 | 77085513 | 77086013 | Gm22696 | Intergenic | ? |
| 12 | 16343694 | 16344194 | Gm48605 | Intergenic | ? |
| 12 | 17660694 | 17661194 | Gm25925 | Intergenic | ? |
| 12 | 24425135 | 24425635 | Gm9315 | Intergenic | ? |
| 12 | 44717013 | 44717513 | Gm6125 | Intergenic | ? |
| 12 | 105281790 | 105282290 | Tunar | Intergenic | ? |
| 12 | 105427790 | 105428290 | Gm46376 | Intergenic | ? |
| 12 | 18310194 | 18310694 | Gm7093 | Intergenic | ? |
| 12 | 21601639 | 21602139 | Gm4036 | Intergenic | ? |
| 12 | 22031048 | 22031548 | Gm22908 | Intergenic | ? |
| 12 | 106536790 | 106537290 | Gm17053 | Intergenic | ? |
| 12 | 19199145 | 19199645 | Gm48528 | Intergenic | ? |
| 12 | 28592135 | 28592635 | Colec11 | Intergenic | ? |
| 12 | 32910635 | 32911135 | 4933406C10Rik | Intergenic | ? |
| 12 | 52047013 | 52047513 | Gpr33 | Intergenic | ? |
| 12 | 68576513 | 68577013 | n-R5s61 | Intergenic | ? |
| 12 | 76517513 | 76518013 | Plekhg3 | Intergenic | ? |
| 12 | 23775135 | 23775635 | Gm10330 | Intergenic | ? |
| 12 | 51276513 | 51277013 | Rps11-ps4 | Intergenic | ? |
| 12 | 53221013 | 53221513 | Npas3 | Intergenic | ? |
| 12 | 86911550 | 86912050 | Irf2bpl | Intergenic | ? |
| 12 | 104678790 | 104679290 | Dicer1 | Intergenic | ? |
| 12 | 107046790 | 107047290 | Gm16086 | Intergenic | ? |
| 12 | 10572194 | 10572694 | Gm38407 | Intergenic | ? |
| 12 | 28872135 | 28872635 | Eipr1 | Intergenic | ? |
| 12 | 29453135 | 29453635 | Gm20208 | Intergenic | ? |

| | | | | | |
|---|---|---|---|---|---|
| 12 | 105256790 | 105257290 | Tcl1 | Intergenic | ? |
| 12 | 12995194 | 12995694 | Gm35208 | Intergenic | ? |
| 12 | 27670135 | 27670635 | Gm24326 | Intergenic | ? |
| 12 | 51065513 | 51066013 | 1700008C04Rik | Intergenic | ? |
| 12 | 51492013 | 51492513 | Gm18503 | Intergenic | ? |
| 12 | 27184135 | 27184635 | Gm9866 | Intergenic | ? |
| 12 | 111149789 | 111150289 | 4930595D18Rik | Intergenic | ? |
| 13 | 23499131 | 23499631 | Btn2a2 | Intergenic | ? |
| 13 | 98224045 | 98224545 | Utp15 | Intergenic | ? |
| 13 | 43592631 | 43593131 | Rnf182 | Intergenic | ? |
| 13 | 59856139 | 59856639 | Gm48384 | Intergenic | ? |
| 13 | 18456166 | 18456666 | Pou6f2 | Intergenic | ? |
| 13 | 9244254 | 9244754 | Gm8784 | Intergenic | ? |
| 13 | 115343792 | 115344292 | Gm47891 | Intergenic | ? |
| 13 | 54835139 | 54835639 | Gm16250 | Intergenic | ? |
| 13 | 56307139 | 56307639 | Gm17878 | Intergenic | ? |
| 13 | 70214122 | 70214622 | 4930520P13Rik | Intergenic | ? |
| 13 | 85921395 | 85921895 | Gm46396 | Intergenic | ? |
| 13 | 30629631 | 30630131 | 4930519D14Rik | Intergenic | ? |
| 13 | 53557631 | 53558131 | Gm5449 | Intergenic | ? |
| 13 | 55810139 | 55810639 | Catsper3 | Intergenic | ? |
| 13 | 42644131 | 42644631 | Gm23086 | Intergenic | ? |
| 13 | 110039292 | 110039792 | Rab3c | Intergenic | ? |
| 13 | 111665292 | 111665792 | Mier3 | Intergenic | ? |
| 13 | 53795131 | 53795631 | Gm48548 | Intergenic | ? |
| 13 | 102655416 | 102655916 | Cd180 | Intergenic | ? |
| 13 | 114734792 | 114735292 | 4930544M13Rik | Intergenic | ? |
| 13 | 52259631 | 52260131 | Gm48199 | Intergenic | ? |
| 13 | 56172639 | 56173139 | Tifab | Intergenic | ? |
| 13 | 84375895 | 84376395 | Gm26927 | Intergenic | ? |
| 13 | 94660045 | 94660545 | Gm48287 | Intergenic | ? |
| 14 | 89182193 | 89182693 | Gm36946 | Intergenic | ? |
| 14 | 99280281 | 99280781 | Gm4412 | Intergenic | ? |
| 14 | 50903825 | 50904325 | Osgep | Intergenic | ? |
| 14 | 66931163 | 66931663 | 4930578I07Rik | Intergenic | ? |
| 14 | 90987213 | 90987713 | Gm23225 | Intergenic | ? |
| 14 | 78224693 | 78225193 | Gm23652 | Intergenic | ? |
| 14 | 108892283 | 108892783 | Slitrk1 | Intergenic | ? |
| 14 | 25063014 | 25063514 | Gm38408 | Intergenic | ? |
| 14 | 62001163 | 62001663 | Gm47953 | Intergenic | ? |
| 14 | 118196778 | 118197278 | 4933431J24Rik | Intergenic | ? |
| 14 | 76295693 | 76296193 | 2900040C04Rik | Intergenic | ? |
| 14 | 93899281 | 93899781 | Pcdh9 | Intergenic | ? |

| | | | | | |
|---|---|---|---|---|---|
| 14 | 29863314 | 29863814 | Gm35281 | Intergenic | ? |
| 14 | 48957825 | 48958325 | Gm23470 | Intergenic | ? |
| 14 | 30699814 | 30700314 | Rft1 | Intergenic | ? |
| 15 | 38177245 | 38177745 | Gm35167 | Intergenic | ? |
| 15 | 97236569 | 97237069 | Amigo2 | Intergenic | ? |
| 15 | 86878570 | 86879070 | Gm49466 | Intergenic | ? |
| 15 | 94004569 | 94005069 | Gm30564 | Intergenic | ? |
| 15 | 18434245 | 18434745 | Gm49139 | Intergenic | ? |
| 15 | 34794745 | 34795245 | Gm48932 | Intergenic | ? |
| 15 | 28034245 | 28034745 | Gm19883 | Intergenic | ? |
| 15 | 51592454 | 51592954 | Gm34562 | Intergenic | ? |
| 15 | 56075445 | 56075945 | Gm49212 | Intergenic | ? |
| 15 | 78520570 | 78521070 | C1qtnf6 | Intergenic | ? |
| 15 | 99569069 | 99569569 | Aqp2 | Intergenic | ? |
| 15 | 71782070 | 71782570 | Gm47309 | Intergenic | ? |
| 15 | 78521070 | 78521570 | C1qtnf6 | Intergenic | ? |
| 15 | 91125069 | 91125569 | Gm7167 | Intergenic | ? |
| 15 | 11972245 | 11972745 | 1700047G03Rik | Intergenic | ? |
| 15 | 37114245 | 37114745 | Gm9509 | Intergenic | ? |
| 15 | 48835454 | 48835954 | Gm7517 | Intergenic | ? |
| 15 | 8748000 | 8748500 | Slc1a3 | Intergenic | ? |
| 15 | 73435070 | 73435570 | Gm10362 | Intergenic | ? |
| 15 | 89727569 | 89728069 | Gm18476 | Intergenic | ? |
| 15 | 5263000 | 5263500 | Gm49093 | Intergenic | ? |
| 15 | 8590000 | 8590500 | Slc1a3 | Intergenic | ? |
| 15 | 25282245 | 25282745 | 4930445E18Rik | Intergenic | ? |
| 15 | 84997570 | 84998070 | 5031439G07Rik | Intergenic | ? |
| 15 | 85715570 | 85716070 | Mirlet7b | Intergenic | ? |
| 15 | 84566570 | 84567070 | Gm34095 | Intergenic | ? |
| 16 | 84485755 | 84486255 | Gm6032 | Intergenic | ? |
| 16 | 21339427 | 21339927 | Magef1 | Intergenic | ? |
| 16 | 84569255 | 84569755 | Gm2477 | Intergenic | ? |
| 16 | 95026393 | 95026893 | Gm49641 | Intergenic | ? |
| 16 | 17051407 | 17051907 | 1700056N10Rik | Intergenic | ? |
| 16 | 39121387 | 39121887 | Gm36742 | Intergenic | ? |
| 17 | 9043135 | 9043635 | 1700010I14Rik | Intergenic | ? |
| 17 | 9047635 | 9048135 | 1700010I14Rik | Intergenic | ? |
| 17 | 79727660 | 79728160 | Gm33055 | Intergenic | ? |
| 17 | 65548660 | 65549160 | Gm49866 | Intergenic | ? |
| 17 | 67422160 | 67422660 | Gm22277 | Intergenic | ? |
| 17 | 48882175 | 48882675 | Lrfn2 | Intergenic | ? |
| 17 | 79472160 | 79472660 | Cdc42ep3 | Intergenic | ? |
| 17 | 79472660 | 79473160 | Cdc42ep3 | Intergenic | ? |

| | | | | | |
|---|---|---|---|---|---|
| 17 | 86909660 | 86910160 | Tmem247 | Intergenic | ? |
| 17 | 10329635 | 10330135 | 1700110C19Rik | Intergenic | ? |
| 17 | 14230993 | 14231493 | Gm34510 | Intergenic | ? |
| 17 | 62509151 | 62509651 | Gm25800 | Intergenic | ? |
| 17 | 8267635 | 8268135 | Ccr6 | Intergenic | ? |
| 17 | 87523660 | 87524160 | Gm30984 | Intergenic | ? |
| 17 | 28097555 | 28098055 | Tcp11 | Intergenic | ? |
| 17 | 84873160 | 84873660 | Gm9400 | Intergenic | ? |
| 17 | 81768160 | 81768660 | Rpl31-ps25 | Intergenic | ? |
| 17 | 85682160 | 85682660 | Six2 | Intergenic | ? |
| 17 | 9467135 | 9467635 | Gm17728 | Intergenic | ? |
| 17 | 10390135 | 10390635 | Pacrg | Intergenic | ? |
| 17 | 14094993 | 14095493 | 4930488N24Rik | Intergenic | ? |
| 17 | 35712055 | 35712555 | Gm4577 | Intergenic | ? |
| 18 | 83232108 | 83232608 | 4921531P14Rik | Intergenic | ? |
| 18 | 51081846 | 51082346 | Prr16 | Intergenic | ? |
| 18 | 69846846 | 69847346 | Gm50215 | Intergenic | ? |
| 18 | 72929846 | 72930346 | Gm31908 | Intergenic | ? |
| 18 | 77747261 | 77747761 | Haus1 | Intergenic | ? |
| 18 | 39605846 | 39606346 | Gm50397 | Intergenic | ? |
| 18 | 13106491 | 13106991 | Gm5240 | Intergenic | ? |
| 18 | 24043999 | 24044499 | Zfp24 | Intergenic | ? |
| 18 | 73363846 | 73364346 | n-R5-8s1 | Intergenic | ? |
| 18 | 82798108 | 82798608 | Gm50356 | Intergenic | ? |
| 18 | 42386346 | 42386846 | Pou4f3 | Intergenic | ? |
| 18 | 55121846 | 55122346 | AC163347.1 | Intergenic | ? |
| 18 | 82822108 | 82822608 | 4930445N18Rik | Intergenic | ? |
| 18 | 83943608 | 83944108 | Gm50420 | Intergenic | ? |
| 19 | 54049010 | 54049510 | Adra2a | Intergenic | ? |
| 19 | 26858510 | 26859010 | Gm815 | Intergenic | ? |
| 19 | 55697010 | 55697510 | Tcf7l2 | Intergenic | ? |
| 19 | 5937000 | 5937500 | Pola2 | Intergenic | ? |
| 19 | 20225510 | 20226010 | 1500015L24Rik | Intergenic | ? |
| 19 | 28045510 | 28046010 | Gm5517 | Intergenic | ? |
| 19 | 36477010 | 36477510 | F530104D19Rik | Intergenic | ? |
| 19 | 54889010 | 54889510 | Gm50191 | Intergenic | ? |
| 19 | 44717010 | 44717510 | Gm26644 | Intergenic | ? |
| 19 | 55062010 | 55062510 | Gpam | Intergenic | ? |
| 19 | 56195510 | 56196010 | Gm22271 | Intergenic | ? |
| 19 | 3429500 | 3430000 | Gal | Intergenic | ? |
| 2 | 172965499 | 172965999 | Spo11 | Intergenic | ? |
| 2 | 156805764 | 156806264 | Gm14231 | Intergenic | ? |
| 2 | 70938943 | 70939443 | Mettl8 | Intergenic | ? |

| | | | | | |
|---|---|---|---|---|---|
| 2 | 17899373 | 17899873 | H2al2a | Intergenic | ? |
| 2 | 21010873 | 21011373 | Gm13377 | Intergenic | ? |
| 2 | 74408943 | 74409443 | 4930441J16Rik | Intergenic | ? |
| 2 | 160406764 | 160407264 | Mafb | Intergenic | ? |
| 2 | 174627499 | 174627999 | Zfp831 | Intergenic | ? |
| 2 | 109848343 | 109848843 | Lin7c | Intergenic | ? |
| 2 | 131807264 | 131807764 | 5330413P13Rik | Intergenic | ? |
| 2 | 163776764 | 163777264 | Ccn5 | Intergenic | ? |
| 2 | 172963499 | 172963999 | Spo11 | Intergenic | ? |
| 2 | 174832499 | 174832999 | Gm14617 | Intergenic | ? |
| 2 | 32506980 | 32507480 | Gm37169 | Intergenic | ? |
| 2 | 52762966 | 52763466 | Gm13548 | Intergenic | ? |
| 2 | 72528943 | 72529443 | Rps13-ps8 | Intergenic | ? |
| 2 | 74063943 | 74064443 | A630050E04Rik | Intergenic | ? |
| 2 | 146546264 | 146546764 | 4933406D12Rik | Intergenic | ? |
| 2 | 118887764 | 118888264 | Ivd | Intergenic | ? |
| 2 | 178630795 | 178631295 | Cdh26 | Intergenic | ? |
| 2 | 51640480 | 51640980 | Tas2r134 | Intergenic | ? |
| 2 | 61344943 | 61345443 | Gm26422 | Intergenic | ? |
| 2 | 172058000 | 172058500 | Gm14271 | Intergenic | ? |
| 2 | 160921764 | 160922264 | Gm14228 | Intergenic | ? |
| 2 | 165106000 | 165106500 | Cdh22 | Intergenic | ? |
| 3 | 97309577 | 97310077 | Gm43073 | Intergenic | ? |
| 3 | 69912578 | 69913078 | Gm17895 | Intergenic | ? |
| 3 | 69655578 | 69656078 | Gm37380 | Intergenic | ? |
| 3 | 89511078 | 89511578 | Kcnn3 | Intergenic | ? |
| 3 | 96499077 | 96499577 | Gm22614 | Intergenic | ? |
| 3 | 132562036 | 132562536 | Gm25307 | Intergenic | ? |
| 3 | 146700036 | 146700536 | Gm9480 | Intergenic | ? |
| 3 | 96322077 | 96322577 | Gm24830 | Intergenic | ? |
| 3 | 110547582 | 110548082 | Gm43406 | Intergenic | ? |
| 3 | 145614036 | 145614536 | Znhit6 | Intergenic | ? |
| 3 | 52622078 | 52622578 | Gm10293 | Intergenic | ? |
| 3 | 53988578 | 53989078 | Gm8109 | Intergenic | ? |
| 3 | 101963577 | 101964077 | Slc22a15 | Intergenic | ? |
| 3 | 103711577 | 103712077 | Olfml3 | Intergenic | ? |
| 3 | 27294578 | 27295078 | Gm43345 | Intergenic | ? |
| 3 | 69712078 | 69712578 | Rpl32-ps | Intergenic | ? |
| 3 | 93779578 | 93780078 | Gm5773 | Intergenic | ? |
| 3 | 15582500 | 15583000 | Gm37468 | Intergenic | ? |
| 3 | 37595578 | 37596078 | Gm12565 | Intergenic | ? |
| 3 | 126590582 | 126591082 | Camk2d | Intergenic | ? |
| 3 | 52415078 | 52415578 | Gm38098 | Intergenic | ? |
| 3 | 32318078 | 32318578 | Zmat3 | Intergenic | ? |

| | | | | | |
|---|---|---|---|---|---|
| 3 | 107203582 | 107204082 | Cym | Intergenic | ? |
| 3 | 46299578 | 46300078 | Gm26081 | Intergenic | ? |
| 4 | 146486538 | 146487038 | Gm13245 | Intergenic | ? |
| 4 | 99625809 | 99626309 | Gm12687 | Intergenic | ? |
| 4 | 82769596 | 82770096 | Zdhhc21 | Intergenic | ? |
| 4 | 133488585 | 133489085 | Tent5b | Intergenic | ? |
| 4 | 148766891 | 148767391 | Casz1 | Intergenic | ? |
| 4 | 150808391 | 150808891 | Gm13049 | Intergenic | ? |
| 4 | 15314853 | 15315353 | Tmem64 | Intergenic | ? |
| 4 | 55515628 | 55516128 | Gm12507 | Intergenic | ? |
| 4 | 129094756 | 129095256 | Rnf19b | Intergenic | ? |
| 4 | 133940585 | 133941085 | Hmgn2 | Intergenic | ? |
| 4 | 138943085 | 138943585 | Rnf186 | Intergenic | ? |
| 4 | 38926467 | 38926967 | Gm12382 | Intergenic | ? |
| 4 | 128981256 | 128981756 | Ak2 | Intergenic | ? |
| 4 | 137072085 | 137072585 | 1700037C06Rik | Intergenic | ? |
| 4 | 138958585 | 138959085 | Rnf186 | Intergenic | ? |
| 4 | 153057891 | 153058391 | Gm25779 | Intergenic | ? |
| 4 | 24154853 | 24155353 | Gm11892 | Intergenic | ? |
| 4 | 61701966 | 61702466 | Mup-ps15 | Intergenic | ? |
| 4 | 88060096 | 88060596 | Gm12646 | Intergenic | ? |
| 4 | 94564309 | 94564809 | Plaa | Intergenic | ? |
| 4 | 138664585 | 138665085 | Ubxn10 | Intergenic | ? |
| 4 | 8085353 | 8085853 | Gm11800 | Intergenic | ? |
| 4 | 41064467 | 41064967 | Gm25637 | Intergenic | ? |
| 4 | 52317128 | 52317628 | Gm12475 | Intergenic | ? |
| 4 | 99560809 | 99561309 | Gm12690 | Intergenic | ? |
| 4 | 118296895 | 118297395 | Ptprf | Intergenic | ? |
| 4 | 125075256 | 125075756 | Snip1 | Intergenic | ? |
| 4 | 125203756 | 125204256 | 1700041M05Rik | Intergenic | ? |
| 4 | 130845085 | 130845585 | Gm22660 | Intergenic | ? |
| 4 | 131340585 | 131341085 | Gm25261 | Intergenic | ? |
| 4 | 45835628 | 45836128 | Stra6l | Intergenic | ? |
| 4 | 61715966 | 61716466 | Mup-ps15 | Intergenic | ? |
| 4 | 83064096 | 83064596 | Frem1 | Intergenic | ? |
| 4 | 84685596 | 84686096 | Bnc2 | Intergenic | ? |
| 4 | 113305395 | 113305895 | Gm12813 | Intergenic | ? |
| 4 | 128927756 | 128928256 | Azin2 | Intergenic | ? |
| 4 | 153738391 | 153738891 | Gm25880 | Intergenic | ? |
| 4 | 33526525 | 33527025 | Gm11935 | Intergenic | ? |
| 4 | 131021085 | 131021585 | A930031H19Rik | Intergenic | ? |
| 4 | 151906891 | 151907391 | Gm37141 | Intergenic | ? |
| 4 | 45742128 | 45742628 | Gm12409 | Intergenic | ? |
| 4 | 110021895 | 110022395 | Dmrta2 | Intergenic | ? |

| | | | | | |
|---|---|---|---|---|---|
| 4 | 127263756 | 127264256 | Smim12 | Intergenic | ? |
| 4 | 147968891 | 147969391 | Nppb | Intergenic | ? |
| 5 | 53678261 | 53678761 | Cckar | Intergenic | ? |
| 5 | 138956546 | 138957046 | Pdgfa | Intergenic | ? |
| 5 | 128527625 | 128528125 | Gm42499 | Intergenic | ? |
| 5 | 66735761 | 66736261 | Limch1 | Intergenic | ? |
| 5 | 67223761 | 67224261 | Gm42670 | Intergenic | ? |
| 5 | 117086991 | 117087491 | Suds3 | Intergenic | ? |
| 5 | 118317991 | 118318491 | Gm25076 | Intergenic | ? |
| 5 | 118841491 | 118841991 | Gm43782 | Intergenic | ? |
| 5 | 120393491 | 120393991 | Gm27199 | Intergenic | ? |
| 5 | 138875546 | 138876046 | Gm5294 | Intergenic | ? |
| 5 | 41556761 | 41557261 | Rab28 | Intergenic | ? |
| 5 | 106386481 | 106386981 | Gm42916 | Intergenic | ? |
| 5 | 111638480 | 111638980 | Gm42489 | Intergenic | ? |
| 5 | 143441632 | 143442132 | Gm4870 | Intergenic | ? |
| 5 | 102102981 | 102103481 | Gm29707 | Intergenic | ? |
| 5 | 106292981 | 106293481 | Gm38433 | Intergenic | ? |
| 5 | 140852546 | 140853046 | Card11 | Intergenic | ? |
| 5 | 3023500 | 3024000 | Gm3027 | Intergenic | ? |
| 5 | 111465980 | 111466480 | Gm43119 | Intergenic | ? |
| 5 | 33528351 | 33528851 | Fam53a | Intergenic | ? |
| 5 | 101530981 | 101531481 | Gm43103 | Intergenic | ? |
| 5 | 118361491 | 118361991 | Gm28563 | Intergenic | ? |
| 6 | 60892006 | 60892506 | Gm43867 | Intergenic | ? |
| 6 | 145343980 | 145344480 | Lmntd1 | Intergenic | ? |
| 6 | 51743501 | 51744001 | Gm38811 | Intergenic | ? |
| 6 | 55646506 | 55647006 | Neurod6 | Intergenic | ? |
| 6 | 73519006 | 73519506 | Gm4374 | Intergenic | ? |
| 6 | 4439000 | 4439500 | Gm37883 | Intergenic | ? |
| 6 | 50938501 | 50939001 | Gm31579 | Intergenic | ? |
| 6 | 126384482 | 126384982 | Gm44431 | Intergenic | ? |
| 6 | 14307500 | 14308000 | Gm8820 | Intergenic | ? |
| 6 | 127084482 | 127084982 | Tigar | Intergenic | ? |
| 6 | 127841982 | 127842482 | Gm23098 | Intergenic | ? |
| 6 | 49023001 | 49023501 | Gpnmb | Intergenic | ? |
| 6 | 50681501 | 50682001 | Npvf | Intergenic | ? |
| 6 | 53262506 | 53263006 | Creb5 | Intergenic | ? |
| 6 | 126700982 | 126701482 | Kcna6 | Intergenic | ? |
| 6 | 13684500 | 13685000 | Bmt2 | Intergenic | ? |
| 6 | 117114982 | 117115482 | Gm43930 | Intergenic | ? |
| 6 | 14620500 | 14621000 | Gm44446 | Intergenic | ? |
| 6 | 87119006 | 87119506 | D6Ertd527e | Intergenic | ? |
| 6 | 127399982 | 127400482 | Gm34091 | Intergenic | ? |
| 6 | 51822001 | 51822501 | Skap2 | Intergenic | ? |
| 6 | 140853479 | 140853979 | Gm43925 | Intergenic | ? |

| | | | | | |
|---|---|---|---|---|---|
| 6 | 4857000 | 4857500 | Gm26828 | Intergenic | ? |
| 6 | 31016500 | 31017000 | Mir29a | Intergenic | ? |
| 6 | 50955501 | 50956001 | Gm44402 | Intergenic | ? |
| 6 | 59185006 | 59185506 | Gm43905 | Intergenic | ? |
| 6 | 73369006 | 73369506 | Gm24370 | Intergenic | ? |
| 6 | 127337482 | 127337982 | Gm42458 | Intergenic | ? |
| 6 | 141876979 | 141877479 | Gm30784 | Intergenic | ? |
| 6 | 49652001 | 49652501 | Gm44219 | Intergenic | ? |
| 6 | 49978001 | 49978501 | Gm3455 | Intergenic | ? |
| 7 | 72949614 | 72950114 | Gm5335 | Intergenic | ? |
| 7 | 145110595 | 145111095 | Gm45181 | Intergenic | ? |
| 7 | 73921114 | 73921614 | St8sia2 | Intergenic | ? |
| 7 | 62719614 | 62720114 | Gm9445 | Intergenic | ? |
| 7 | 105723986 | 105724486 | Rrp8 | Intergenic | ? |
| 7 | 129516986 | 129517486 | Wdr11 | Intergenic | ? |
| 7 | 37253981 | 37254481 | Gm28077 | Intergenic | ? |
| 7 | 37845981 | 37846481 | Gm22302 | Intergenic | ? |
| 7 | 30338481 | 30338981 | Gm50483 | Intergenic | ? |
| 7 | 80160114 | 80160614 | Gm45202 | Intergenic | ? |
| 7 | 16191151 | 16191651 | Meis3 | Intergenic | ? |
| 7 | 34606481 | 34606981 | Gm12784 | Intergenic | ? |
| 7 | 70154114 | 70154614 | Gm29683 | Intergenic | ? |
| 7 | 128424486 | 128424986 | Gm15503 | Intergenic | ? |
| 7 | 24429481 | 24429981 | Irgc1 | Intergenic | ? |
| 7 | 55736130 | 55736630 | Siglech | Intergenic | ? |
| 8 | 113930600 | 113931100 | Adamts18 | Intergenic | ? |
| 8 | 117894600 | 117895100 | Gm10617 | Intergenic | ? |
| 8 | 47461646 | 47462146 | Gm22857 | Intergenic | ? |
| 8 | 94205100 | 94205600 | Nup93 | Intergenic | ? |
| 8 | 31101028 | 31101528 | Dusp26 | Intergenic | ? |
| 8 | 34679946 | 34680446 | Gm45627 | Intergenic | ? |
| 8 | 40187146 | 40187646 | Fgf20 | Intergenic | ? |
| 8 | 89730101 | 89730601 | Gm24212 | Intergenic | ? |
| 8 | 96151100 | 96151600 | Gm32005 | Intergenic | ? |
| 8 | 97935600 | 97936100 | Gm45246 | Intergenic | ? |
| 8 | 100788100 | 100788600 | Gm39232 | Intergenic | ? |
| 8 | 109984600 | 109985100 | Tat | Intergenic | ? |
| 8 | 112127100 | 112127600 | Gm22827 | Intergenic | ? |
| 8 | 26989028 | 26989528 | Gm45370 | Intergenic | ? |
| 8 | 35123446 | 35123946 | Rpl31-ps23 | Intergenic | ? |
| 8 | 35321446 | 35321946 | Gm45469 | Intergenic | ? |
| 8 | 40272646 | 40273146 | Fgf20 | Intergenic | ? |
| 8 | 109458100 | 109458600 | Gm23163 | Intergenic | ? |
| 8 | 57680203 | 57680703 | Gm45644 | Intergenic | ? |
| 8 | 10837500 | 10838000 | Gm32540 | Intergenic | ? |
| 8 | 18506500 | 18507000 | Gm24469 | Intergenic | ? |

| | | | | | |
|---|---|---|---|---|---|
| 8 | 24311028 | 24311528 | Gm45164 | Intergenic | ? |
| 8 | 48047646 | 48048146 | Gm2607 | Intergenic | ? |
| 8 | 91738101 | 91738601 | Gm36325 | Intergenic | ? |
| 8 | 104037100 | 104037600 | Gm8748 | Intergenic | ? |
| 8 | 125391600 | 125392100 | Sipa1l2 | Intergenic | ? |
| 8 | 103949600 | 103950100 | Gm8748 | Intergenic | ? |
| 8 | 62324203 | 62324703 | Gm2961 | Intergenic | ? |
| 8 | 10653500 | 10654000 | Myo16 | Intergenic | ? |
| 8 | 17739000 | 17739500 | Csmd1 | Intergenic | ? |
| 9 | 54527193 | 54527693 | Sh2d7 | Intergenic | ? |
| 9 | 69540193 | 69540693 | Anxa2 | Intergenic | ? |
| 9 | 95521081 | 95521581 | Gm28424 | Intergenic | ? |
| 9 | 32550415 | 32550915 | Gm37082 | Intergenic | ? |
| 9 | 70181193 | 70181693 | Myo1e | Intergenic | ? |
| 9 | 46653917 | 46654417 | Gm31557 | Intergenic | ? |
| 9 | 61490693 | 61491193 | Gm47240 | Intergenic | ? |
| 9 | 75740193 | 75740693 | Bmp5 | Intergenic | ? |
| 9 | 99445581 | 99446081 | Nme9 | Intergenic | ? |
| 9 | 107249669 | 107250169 | Mapkapk3 | Intergenic | ? |
| 9 | 69671693 | 69672193 | Gm47203 | Intergenic | ? |
| 9 | 83674393 | 83674893 | Gm36120 | Intergenic | ? |
| 9 | 98725081 | 98725581 | Pisrt1 | Intergenic | ? |
| 9 | 121514382 | 121514882 | Gm47095 | Intergenic | ? |
| 9 | 49975895 | 49976395 | Gm47543 | Intergenic | ? |
| 9 | 67799193 | 67799693 | C2cd4a | Intergenic | ? |
| 9 | 83726893 | 83727393 | Gm2109 | Intergenic | ? |
| 9 | 86902665 | 86903165 | Snap91 | Intergenic | ? |
| 9 | 114234882 | 114235382 | 4921528I07Rik | Intergenic | ? |

The gene annotation used Gencode.vM24. Location of H-sites may be related to more than one gene section. 3'-UTR is three prime untranslated regions; CDS is Coding Sequence.

**Table S3**- Differentially expressed genes in adult ALB-D2KO mouse liver as compared to littermate control samples from Cre-expressing mice.

| Gene ID | Total counts | P-value | FDR | Ratio | Fold change |
|---|---|---|---|---|---|
| Gm14434 | 66.13 | 0.02 | 0.61 | 0.08 | -12.38 |
| Gm31508 | 107.00 | 1.66E-04 | 0.18 | 0.13 | -7.76 |
| Acot3 | 1013.21 | 1.58E-03 | 0.34 | 0.15 | -6.73 |
| Cyp4a10 | 9119.81 | 7.47E-04 | 0.25 | 0.17 | -5.72 |
| Gm26232 | 268.41 | 0.04 | 0.64 | 0.18 | -5.67 |
| Gm21049 | 472.00 | 4.59E-05 | 0.11 | 0.19 | -5.39 |
| Cish | 5784.80 | 2.37E-05 | 0.08 | 0.20 | -5.04 |
| Cyp4a14 | 6246.99 | 0.01 | 0.53 | 0.20 | -4.89 |
| Gadd45g | 2990.00 | 0.03 | 0.62 | 0.21 | -4.75 |
| Gm28182 | 427.00 | 0.01 | 0.58 | 0.22 | -4.54 |
| Crybg2 | 273.00 | 2.44E-03 | 0.40 | 0.22 | -4.53 |
| Gm19963 | 32.00 | 0.01 | 0.58 | 0.23 | -4.32 |
| Gm13168 | 25.87 | 0.02 | 0.60 | 0.23 | -4.28 |
| Mt2 | 1861.78 | 0.02 | 0.61 | 0.24 | -4.17 |
| Gm49356 | 71.17 | 1.24E-03 | 0.31 | 0.24 | -4.12 |
| Gm18149 | 49.00 | 4.02E-03 | 0.46 | 0.24 | -4.10 |
| E2f8 | 2774.62 | 4.24E-04 | 0.21 | 0.25 | -4.06 |
| Csmd1 | 48.00 | 5.67E-04 | 0.24 | 0.25 | -4.00 |
| Socs2 | 3784.42 | 9.52E-05 | 0.16 | 0.25 | -3.96 |
| Serpinb1a | 148.00 | 6.39E-04 | 0.25 | 0.25 | -3.94 |
| Gm42796 | 466.00 | 9.11E-04 | 0.28 | 0.26 | -3.92 |
| Gbp11 | 1296.33 | 2.54E-04 | 0.19 | 0.26 | -3.90 |
| Gm20503 | 217.33 | 3.24E-04 | 0.19 | 0.26 | -3.79 |
| Gm48765 | 40.00 | 0.03 | 0.62 | 0.27 | -3.77 |
| Myc | 2294.00 | 4.40E-03 | 0.48 | 0.27 | -3.75 |
| CAAA01066123.1 | 55.17 | 0.01 | 0.58 | 0.28 | -3.54 |
| Gm49413 | 414.00 | 1.28E-03 | 0.32 | 0.29 | -3.50 |
| Gm45141 | 41.25 | 4.84E-03 | 0.48 | 0.29 | -3.49 |
| Gm49733 | 32.00 | 0.01 | 0.54 | 0.29 | -3.47 |
| Dio3os | 41.00 | 0.03 | 0.62 | 0.30 | -3.30 |
| Gm38228 | 46.00 | 3.28E-03 | 0.44 | 0.30 | -3.29 |
| Gm48111 | 46.00 | 3.73E-03 | 0.46 | 0.30 | -3.29 |
| Zfp125 | 1550.88 | 1.46E-03 | 0.33 | 0.31 | -3.24 |
| Gm45044 | 45.00 | 4.71E-03 | 0.48 | 0.31 | -3.20 |
| Ano5 | 35.00 | 0.01 | 0.58 | 0.31 | -3.20 |
| Gm8099 | 30.00 | 0.05 | 0.64 | 0.31 | -3.20 |
| Gm45244 | 5579.39 | 3.33E-05 | 0.09 | 0.31 | -3.19 |
| Gm13370 | 36.82 | 4.73E-03 | 0.48 | 0.31 | -3.18 |
| Gm10240 | 36.82 | 4.73E-03 | 0.48 | 0.31 | -3.18 |
| Ska3 | 29.00 | 0.03 | 0.63 | 0.33 | -3.07 |
| 5730521K06Rik | 175.15 | 0.04 | 0.64 | 0.33 | -3.02 |

| | | | | | |
|---|---|---|---|---|---|
| Clec2h | 1147.00 | 2.74E-04 | 0.19 | 0.33 | -3.01 |
| Olfr542-ps1 | 635.00 | 5.22E-05 | 0.11 | 0.33 | -2.99 |
| Gm44146 | 71.00 | 1.38E-03 | 0.32 | 0.33 | -2.99 |
| Ankmy1 | 30.22 | 0.03 | 0.62 | 0.34 | -2.97 |
| Gm16585 | 61.00 | 4.36E-03 | 0.48 | 0.34 | -2.95 |
| Gm2174 | 182.84 | 0.01 | 0.53 | 0.34 | -2.95 |
| Gm47341 | 56.00 | 4.67E-03 | 0.48 | 0.34 | -2.93 |
| Nkapl | 42.00 | 0.01 | 0.54 | 0.34 | -2.93 |
| Gm45553 | 56.00 | 0.01 | 0.58 | 0.34 | -2.93 |
| Spata17 | 28.00 | 0.03 | 0.61 | 0.34 | -2.93 |
| Gm12712 | 100.37 | 0.02 | 0.61 | 0.35 | -2.89 |
| Gm6155 | 57.52 | 2.58E-03 | 0.40 | 0.35 | -2.88 |
| Gm43111 | 32.00 | 0.02 | 0.61 | 0.35 | -2.86 |
| Gm26925 | 68.00 | 3.80E-03 | 0.46 | 0.35 | -2.83 |
| A230087F16Rik | 84.22 | 2.81E-03 | 0.42 | 0.36 | -2.80 |
| Olfr539 | 207.31 | 0.01 | 0.53 | 0.36 | -2.77 |
| Gm26646 | 93.83 | 2.13E-03 | 0.38 | 0.36 | -2.76 |
| Gm43909 | 220.62 | 0.03 | 0.61 | 0.36 | -2.76 |
| Nup62cl | 80.00 | 0.02 | 0.59 | 0.36 | -2.76 |
| Gm8261 | 53.00 | 0.02 | 0.60 | 0.37 | -2.73 |
| Gm43946 | 75.00 | 0.01 | 0.52 | 0.37 | -2.73 |
| Kcnk2 | 20.70 | 0.05 | 0.64 | 0.37 | -2.72 |
| Trib3 | 933.00 | 1.62E-03 | 0.34 | 0.37 | -2.72 |
| Gm7560 | 44.00 | 0.01 | 0.54 | 0.37 | -2.72 |
| Onecut1 | 45440.45 | 0.01 | 0.56 | 0.37 | -2.72 |
| Phlda1 | 33252.31 | 0.01 | 0.50 | 0.37 | -2.71 |
| Gm45112 | 101.00 | 0.01 | 0.50 | 0.37 | -2.71 |
| Egr1 | 5935.99 | 0.01 | 0.58 | 0.37 | -2.70 |
| Gm21982 | 97.67 | 0.03 | 0.62 | 0.37 | -2.69 |
| Gm36041 | 651.00 | 1.47E-03 | 0.33 | 0.37 | -2.67 |
| Gm13521 | 39.00 | 0.02 | 0.59 | 0.38 | -2.67 |
| Klrb1c | 33.83 | 0.01 | 0.58 | 0.38 | -2.66 |
| Ikzf4 | 497.00 | 4.34E-03 | 0.48 | 0.38 | -2.66 |
| Rab9b-ps1 | 30.00 | 0.01 | 0.56 | 0.38 | -2.63 |
| BC106179 | 121.27 | 4.02E-03 | 0.46 | 0.38 | -2.62 |
| Slc22a27 | 38.41 | 0.04 | 0.63 | 0.38 | -2.61 |
| Olfr45 | 298.45 | 4.75E-03 | 0.48 | 0.38 | -2.61 |
| Gm13536 | 34.00 | 0.04 | 0.64 | 0.38 | -2.60 |
| Gm2788 | 3137.39 | 3.02E-03 | 0.44 | 0.39 | -2.59 |
| Scd4 | 59.57 | 0.02 | 0.59 | 0.39 | -2.57 |
| Traip | 59.00 | 0.01 | 0.54 | 0.39 | -2.57 |
| B230220B15Rik | 34.42 | 0.02 | 0.59 | 0.39 | -2.56 |
| Irx3 | 200.00 | 2.18E-03 | 0.38 | 0.39 | -2.53 |
| 3110062G12Rik | 25.00 | 0.03 | 0.62 | 0.39 | -2.53 |
| H2-Oa | 29.00 | 0.02 | 0.61 | 0.40 | -2.51 |

| | | | | | |
|---|---|---|---|---|---|
| Npas1 | 29.00 | 0.03 | 0.63 | 0.40 | -2.51 |
| Fam229b | 29.00 | 0.04 | 0.63 | 0.40 | -2.51 |
| Gm44639 | 186.00 | 3.55E-03 | 0.45 | 0.40 | -2.51 |
| Mettl7a2 | 129.51 | 0.01 | 0.58 | 0.40 | -2.51 |
| Gm39465 | 33.00 | 0.03 | 0.62 | 0.40 | -2.50 |
| Gm37849 | 33.00 | 0.05 | 0.64 | 0.40 | -2.50 |
| Gm4045 | 69.24 | 0.03 | 0.62 | 0.40 | -2.50 |
| Gm10634 | 37.03 | 0.02 | 0.59 | 0.40 | -2.49 |
| D430020J02Rik | 37.00 | 0.03 | 0.62 | 0.40 | -2.49 |
| Lgals2 | 41.00 | 0.02 | 0.59 | 0.40 | -2.48 |
| A730085K08Rik | 110.57 | 0.04 | 0.64 | 0.40 | -2.47 |
| Gm43605 | 94.00 | 0.05 | 0.64 | 0.40 | -2.47 |
| Robo3 | 53.00 | 0.01 | 0.58 | 0.41 | -2.46 |
| Gm3896 | 53.00 | 0.02 | 0.60 | 0.41 | -2.46 |
| Fcho1 | 57.00 | 0.04 | 0.64 | 0.41 | -2.46 |
| C230071H17Rik | 58.42 | 0.04 | 0.63 | 0.41 | -2.44 |
| Gm47818 | 93.00 | 0.01 | 0.53 | 0.41 | -2.43 |
| Gm10599 | 122.04 | 0.01 | 0.58 | 0.41 | -2.41 |
| Fam221a | 52.00 | 0.02 | 0.59 | 0.42 | -2.40 |
| Gm48878 | 148.00 | 8.82E-04 | 0.28 | 0.42 | -2.40 |
| 2900093K20Rik | 104.00 | 3.05E-03 | 0.44 | 0.42 | -2.40 |
| Gm24233 | 132.00 | 0.01 | 0.53 | 0.42 | -2.40 |
| Gm44022 | 56.00 | 0.01 | 0.56 | 0.42 | -2.40 |
| Fam78b | 52.00 | 0.04 | 0.63 | 0.42 | -2.40 |
| 4930597O21Rik | 40.00 | 0.04 | 0.64 | 0.42 | -2.40 |
| Gm8344 | 20.00 | 0.04 | 0.64 | 0.42 | -2.40 |
| Gm48734 | 36.00 | 0.05 | 0.64 | 0.42 | -2.40 |
| Zfp941 | 168.00 | 0.02 | 0.61 | 0.42 | -2.40 |
| Cyp4a31 | 612.45 | 0.02 | 0.59 | 0.42 | -2.40 |
| Cxxc4 | 126.67 | 2.60E-03 | 0.40 | 0.42 | -2.40 |
| Gm17039 | 83.49 | 0.02 | 0.59 | 0.42 | -2.38 |
| Gm44017 | 75.00 | 0.01 | 0.54 | 0.42 | -2.36 |
| Gm20594 | 3276.00 | 1.09E-04 | 0.16 | 0.42 | -2.35 |
| Gm44562 | 122.00 | 0.01 | 0.54 | 0.43 | -2.35 |
| Gm45186 | 55.00 | 0.01 | 0.58 | 0.43 | -2.34 |
| Nxpe1-ps | 51.00 | 0.01 | 0.58 | 0.43 | -2.34 |
| Slc1a1 | 31.38 | 0.04 | 0.64 | 0.43 | -2.34 |
| Gm48010 | 47.00 | 0.01 | 0.58 | 0.43 | -2.33 |
| Slc25a30 | 9714.00 | 0.01 | 0.50 | 0.43 | -2.33 |
| Gm47204 | 41.70 | 0.03 | 0.62 | 0.43 | -2.32 |
| Snora78 | 39.00 | 0.02 | 0.59 | 0.43 | -2.32 |
| Cd69 | 39.00 | 0.03 | 0.62 | 0.43 | -2.32 |
| Kif14 | 35.00 | 0.02 | 0.61 | 0.43 | -2.31 |
| Gm6054 | 35.15 | 0.04 | 0.63 | 0.43 | -2.31 |
| Mns1 | 45.68 | 0.02 | 0.59 | 0.43 | -2.31 |

| | | | | | |
|---|---|---|---|---|---|
| Pcdhb17 | 136.00 | 0.01 | 0.50 | 0.43 | -2.31 |
| Gm44033 | 128.00 | 3.61E-03 | 0.45 | 0.43 | -2.30 |
| Hamp2 | 1972.02 | 0.01 | 0.57 | 0.44 | -2.29 |
| Gm8738 | 58.00 | 0.02 | 0.59 | 0.44 | -2.29 |
| Ccdc24 | 381.20 | 0.01 | 0.58 | 0.44 | -2.29 |
| Clstn2 | 139.00 | 0.01 | 0.50 | 0.44 | -2.29 |
| Siglecg | 108.00 | 0.03 | 0.62 | 0.44 | -2.29 |
| Car13 | 54.00 | 0.03 | 0.62 | 0.44 | -2.29 |
| Gm16552 | 89.49 | 0.01 | 0.57 | 0.44 | -2.28 |
| Gm37571 | 50.00 | 0.02 | 0.61 | 0.44 | -2.28 |
| Lars2 | 199871.71 | 6.95E-04 | 0.25 | 0.44 | -2.27 |
| AI504432 | 46.00 | 0.03 | 0.63 | 0.44 | -2.27 |
| Gm29383 | 88.00 | 0.04 | 0.63 | 0.44 | -2.26 |
| Zfp455 | 195.00 | 3.37E-03 | 0.44 | 0.44 | -2.26 |
| Gm20522 | 129.85 | 2.46E-03 | 0.40 | 0.44 | -2.25 |
| Lrrc49 | 61.00 | 0.02 | 0.61 | 0.44 | -2.25 |
| Gm35570 | 627.38 | 3.19E-03 | 0.44 | 0.45 | -2.24 |
| Gm7678 | 38.00 | 0.03 | 0.61 | 0.45 | -2.24 |
| Zfp57 | 91.00 | 0.01 | 0.50 | 0.45 | -2.23 |
| Anp32-ps | 87.10 | 0.04 | 0.63 | 0.45 | -2.23 |
| Gm15582 | 246.00 | 0.01 | 0.53 | 0.45 | -2.23 |
| Cacna1e | 121.00 | 0.01 | 0.49 | 0.45 | -2.22 |
| Rubcnl | 170.00 | 0.01 | 0.58 | 0.45 | -2.22 |
| Gm43916 | 83.00 | 0.01 | 0.58 | 0.45 | -2.22 |
| Gm26891 | 54.02 | 0.05 | 0.64 | 0.45 | -2.22 |
| Gm45807 | 146.57 | 0.02 | 0.59 | 0.45 | -2.22 |
| 1700007E05Rik | 49.00 | 0.02 | 0.59 | 0.45 | -2.22 |
| Gm10855 | 49.00 | 0.02 | 0.59 | 0.45 | -2.22 |
| Gm16217 | 113.00 | 0.01 | 0.58 | 0.45 | -2.21 |
| Gm1600 | 113.00 | 0.05 | 0.64 | 0.45 | -2.21 |
| Gm44787 | 109.00 | 0.02 | 0.61 | 0.45 | -2.21 |
| Prr18 | 109.00 | 0.02 | 0.59 | 0.45 | -2.21 |
| 2610307P16Rik | 45.00 | 0.02 | 0.60 | 0.45 | -2.20 |
| Tcp11l2 | 5144.01 | 6.09E-05 | 0.12 | 0.46 | -2.20 |
| Gm15746 | 1056.00 | 2.11E-03 | 0.38 | 0.46 | -2.20 |
| Gm37760 | 805.00 | 1.16E-03 | 0.31 | 0.46 | -2.20 |
| Acot2 | 804.65 | 4.35E-04 | 0.21 | 0.46 | -2.19 |
| Gm38227 | 56.00 | 0.01 | 0.58 | 0.46 | -2.19 |
| Gm50070 | 56.00 | 0.02 | 0.60 | 0.46 | -2.19 |
| Gm49101 | 41.00 | 0.04 | 0.64 | 0.46 | -2.18 |
| Gm48207 | 365.00 | 2.01E-03 | 0.38 | 0.46 | -2.18 |
| Gm6377 | 108.00 | 0.04 | 0.63 | 0.46 | -2.18 |
| Gm5417 | 69.09 | 0.03 | 0.61 | 0.46 | -2.17 |
| Ripply1 | 52.00 | 0.03 | 0.62 | 0.46 | -2.17 |
| Cmya5 | 47.05 | 0.02 | 0.60 | 0.46 | -2.17 |

| | | | | | |
|---|---|---|---|---|---|
| Unc93a2 | 700.12 | 4.53E-04 | 0.21 | 0.46 | -2.16 |
| Acot1 | 3545.70 | 3.47E-03 | 0.45 | 0.46 | -2.16 |
| Gm18284 | 48.08 | 0.03 | 0.61 | 0.46 | -2.16 |
| Trarg1 | 96.00 | 0.02 | 0.61 | 0.46 | -2.15 |
| Fam205a1 | 177.49 | 0.02 | 0.59 | 0.46 | -2.15 |
| Ckap2l | 59.00 | 0.02 | 0.59 | 0.47 | -2.15 |
| Gm7208 | 311.64 | 2.69E-03 | 0.41 | 0.47 | -2.14 |
| Gm16998 | 110.43 | 0.01 | 0.54 | 0.47 | -2.14 |
| Eif4a3l2 | 56.40 | 0.02 | 0.61 | 0.47 | -2.14 |
| D630039A03Rik | 3588.00 | 6.94E-04 | 0.25 | 0.47 | -2.14 |
| Acss3 | 4061.53 | 1.65E-05 | 0.07 | 0.47 | -2.14 |
| Stil | 55.00 | 0.03 | 0.61 | 0.47 | -2.13 |
| Gm37349 | 66.00 | 0.02 | 0.61 | 0.47 | -2.13 |
| Gm48269 | 44.00 | 0.04 | 0.63 | 0.47 | -2.13 |
| Gm31774 | 428.00 | 0.02 | 0.59 | 0.47 | -2.13 |
| Gm3436 | 232.34 | 0.01 | 0.58 | 0.47 | -2.13 |
| Ranbp3l | 234.00 | 3.35E-03 | 0.44 | 0.47 | -2.12 |
| Gm13351 | 106.00 | 0.01 | 0.52 | 0.47 | -2.12 |
| Gm48204 | 201.00 | 0.01 | 0.54 | 0.47 | -2.12 |
| Rnf186 | 1626.00 | 2.58E-03 | 0.40 | 0.47 | -2.12 |
| Kcnn4 | 73.00 | 0.04 | 0.64 | 0.47 | -2.12 |
| Gm16166 | 91.00 | 0.04 | 0.64 | 0.47 | -2.11 |
| Rpl17-ps4 | 156.72 | 0.01 | 0.50 | 0.47 | -2.11 |
| Gm48871 | 305.00 | 1.70E-03 | 0.35 | 0.48 | -2.10 |
| Rdh16 | 13472.37 | 8.59E-05 | 0.15 | 0.48 | -2.10 |
| Gm23804 | 1643.99 | 0.04 | 0.63 | 0.48 | -2.09 |
| Ttc16 | 55.07 | 0.03 | 0.62 | 0.48 | -2.09 |
| Gm48824 | 65.00 | 0.03 | 0.62 | 0.48 | -2.09 |
| Cpt1b | 272.90 | 4.55E-03 | 0.48 | 0.48 | -2.09 |
| Cyp2c38 | 3019.00 | 2.36E-03 | 0.40 | 0.48 | -2.09 |
| Gm38001 | 108.00 | 0.01 | 0.56 | 0.48 | -2.08 |
| Lmcd1 | 72.00 | 0.03 | 0.62 | 0.48 | -2.08 |
| Fermt1 | 72.00 | 0.04 | 0.64 | 0.48 | -2.08 |
| Gm11992 | 122.00 | 0.01 | 0.58 | 0.48 | -2.07 |
| Gm48501 | 122.00 | 0.03 | 0.62 | 0.48 | -2.07 |
| Gm15779 | 61.00 | 0.04 | 0.63 | 0.48 | -2.07 |
| Gm5131 | 104.00 | 0.01 | 0.57 | 0.48 | -2.07 |
| Hyal3 | 104.00 | 0.03 | 0.63 | 0.48 | -2.07 |
| Cyp2c55 | 999.00 | 1.32E-03 | 0.32 | 0.48 | -2.06 |
| Dnmt3l-ps1 | 111.00 | 0.03 | 0.61 | 0.48 | -2.06 |
| Gm18258 | 68.00 | 0.05 | 0.64 | 0.48 | -2.06 |
| Zyg11a | 3115.00 | 3.53E-03 | 0.45 | 0.49 | -2.06 |
| Gm20528 | 199.49 | 0.03 | 0.63 | 0.49 | -2.06 |
| Gm37332 | 50.00 | 0.04 | 0.63 | 0.49 | -2.06 |
| Gm12020 | 50.00 | 0.05 | 0.64 | 0.49 | -2.06 |

| | | | | | |
|---|---|---|---|---|---|
| Gm4952 | 6797.00 | 2.79E-04 | 0.19 | 0.49 | -2.06 |
| Gm17083 | 289.00 | 0.03 | 0.62 | 0.49 | -2.05 |
| Gm38218 | 260.00 | 4.78E-03 | 0.48 | 0.49 | -2.05 |
| D430041D05Rik | 96.00 | 0.01 | 0.58 | 0.49 | -2.04 |
| Cyp4a32 | 1769.77 | 3.08E-04 | 0.19 | 0.49 | -2.04 |
| Gm45515 | 96.95 | 0.01 | 0.58 | 0.49 | -2.04 |
| Gm43300 | 241.00 | 0.01 | 0.53 | 0.49 | -2.04 |
| Krt23 | 177.00 | 0.02 | 0.61 | 0.49 | -2.03 |
| Gm49942 | 46.00 | 0.02 | 0.61 | 0.49 | -2.03 |
| Gm46378 | 46.00 | 0.03 | 0.62 | 0.49 | -2.03 |
| Grhl3 | 46.00 | 0.03 | 0.63 | 0.49 | -2.03 |
| Ntf3 | 92.00 | 0.02 | 0.59 | 0.49 | -2.03 |
| Gm14414 | 313.02 | 0.01 | 0.50 | 0.49 | -2.03 |
| Gm26628 | 166.00 | 0.02 | 0.59 | 0.49 | -2.03 |
| Gm5112 | 60.00 | 0.03 | 0.63 | 0.49 | -2.02 |
| Prlr | 16821.56 | 1.99E-04 | 0.18 | 0.49 | -2.02 |
| Gm47451 | 67.00 | 0.03 | 0.62 | 0.49 | -2.02 |
| Gm44609 | 162.10 | 0.01 | 0.50 | 0.50 | -2.02 |
| Rapgef4os2 | 2370.50 | 1.20E-03 | 0.31 | 0.50 | -2.02 |
| Ssc5d | 176.00 | 0.02 | 0.59 | 0.50 | -2.02 |
| Gm28940 | 102.00 | 0.01 | 0.58 | 0.50 | -2.01 |
| Gm37818 | 61.35 | 0.04 | 0.63 | 0.50 | -2.01 |
| 1810064F22Rik | 596.00 | 0.01 | 0.58 | 0.50 | -2.00 |
| Smad9 | 5332.00 | 2.59E-04 | 0.19 | 0.50 | -2.00 |
| Gm3776 | 335.49 | 0.01 | 0.53 | 0.50 | -2.00 |
| Itgax | 161.00 | 0.01 | 0.54 | 0.50 | -2.00 |
| Gm48077 | 56.00 | 0.03 | 0.62 | 0.50 | -2.00 |
| Clec2i | 42.00 | 0.03 | 0.63 | 0.50 | -2.00 |
| Gm16588 | 28.00 | 0.04 | 0.63 | 0.50 | -2.00 |
| Tbc1d22bos | 84.00 | 0.05 | 0.64 | 0.50 | -2.00 |
| Gm12749 | 125.92 | 0.02 | 0.60 | 0.50 | -2.00 |
| Gm2996 | 168.43 | 0.01 | 0.56 | 0.50 | -2.00 |
| Rpl19-ps12 | 104.82 | 0.01 | 0.58 | 0.50 | -2.00 |
| Kif12 | 101.00 | 0.03 | 0.62 | 0.50 | -1.99 |
| Gm5826 | 66.74 | 0.02 | 0.61 | 0.50 | -1.98 |
| Gm49268 | 167.00 | 0.01 | 0.50 | 0.50 | -1.98 |
| Gm48708 | 73.00 | 0.03 | 0.62 | 0.50 | -1.98 |
| Hmgb1-rs16 | 73.00 | 0.04 | 0.64 | 0.50 | -1.98 |
| Slc52a3 | 73.00 | 0.03 | 0.63 | 0.50 | -1.98 |
| Gm10385 | 139.00 | 0.04 | 0.63 | 0.51 | -1.98 |
| Zfp114 | 38.46 | 0.05 | 0.64 | 0.51 | -1.98 |
| Fbxo32 | 1360.81 | 2.35E-03 | 0.40 | 0.51 | -1.98 |
| Kcnq5 | 59.00 | 0.03 | 0.63 | 0.51 | -1.98 |
| Gm30042 | 52.00 | 0.05 | 0.64 | 0.51 | -1.97 |
| Gm960 | 97.00 | 0.02 | 0.59 | 0.51 | -1.97 |

| | | | | | |
|---|---|---|---|---|---|
| 9530086O07Rik | 239.00 | 0.04 | 0.64 | 0.51 | -1.97 |
| Gm24265 | 142.00 | 0.03 | 0.63 | 0.51 | -1.97 |
| Gm13898 | 90.87 | 0.02 | 0.60 | 0.51 | -1.97 |
| 9430091E24Rik | 135.00 | 0.02 | 0.61 | 0.51 | -1.97 |
| Pde1c | 166.00 | 0.03 | 0.63 | 0.51 | -1.97 |
| Gm2568 | 152.00 | 0.03 | 0.62 | 0.51 | -1.96 |
| Gm48809 | 107.00 | 0.02 | 0.60 | 0.51 | -1.96 |
| CT010467.1 | 3331696.52 | 1.20E-03 | 0.31 | 0.51 | -1.96 |
| Dubr | 248.00 | 0.01 | 0.53 | 0.51 | -1.96 |
| Gm47357 | 93.00 | 0.03 | 0.62 | 0.51 | -1.96 |
| Gm45083 | 62.00 | 0.05 | 0.64 | 0.51 | -1.96 |
| Itk | 93.00 | 0.04 | 0.64 | 0.51 | -1.96 |
| Bhlhe41 | 2663.38 | 4.31E-03 | 0.48 | 0.51 | -1.95 |
| Tomt | 70.92 | 0.03 | 0.62 | 0.51 | -1.95 |
| Gm10513 | 439.00 | 2.64E-03 | 0.40 | 0.51 | -1.94 |
| 1700109K24Rik | 264.00 | 4.46E-03 | 0.48 | 0.51 | -1.94 |
| 4930455C13Rik | 72.51 | 0.05 | 0.64 | 0.51 | -1.94 |
| Hectd2os | 6788.69 | 5.68E-04 | 0.24 | 0.52 | -1.94 |
| Tagln3 | 65.00 | 0.02 | 0.60 | 0.52 | -1.94 |
| A530083M17Rik | 65.00 | 0.03 | 0.62 | 0.52 | -1.94 |
| Arhgef39 | 106.00 | 0.02 | 0.60 | 0.52 | -1.94 |
| Gm45776 | 997.00 | 0.01 | 0.53 | 0.52 | -1.93 |
| Gm48321 | 324.00 | 0.02 | 0.59 | 0.52 | -1.93 |
| Oaz3 | 68.30 | 0.03 | 0.62 | 0.52 | -1.93 |
| Baiap3 | 44.39 | 0.04 | 0.64 | 0.52 | -1.92 |
| Cd3e | 126.00 | 0.02 | 0.59 | 0.52 | -1.92 |
| Gm26931 | 68.00 | 0.03 | 0.61 | 0.52 | -1.92 |
| Gm6477 | 99.77 | 0.05 | 0.64 | 0.52 | -1.92 |
| 1700034J05Rik | 231.00 | 0.03 | 0.61 | 0.52 | -1.92 |
| Abcd2 | 1341.00 | 0.02 | 0.61 | 0.52 | -1.92 |
| 5330438I03Rik | 238.08 | 0.01 | 0.52 | 0.52 | -1.91 |
| Arid5a | 902.00 | 4.58E-03 | 0.48 | 0.52 | -1.91 |
| Gimap1os | 94.85 | 0.04 | 0.63 | 0.52 | -1.91 |
| Lratd1 | 369.00 | 4.99E-03 | 0.49 | 0.52 | -1.91 |
| Gm10804 | 132.00 | 0.03 | 0.63 | 0.52 | -1.91 |
| Gm29237 | 44.00 | 0.05 | 0.64 | 0.52 | -1.91 |
| Mrnip | 369.86 | 0.04 | 0.64 | 0.52 | -1.91 |
| 1700018L02Rik | 2415.34 | 0.01 | 0.50 | 0.53 | -1.90 |
| Zfa-ps | 46.88 | 0.03 | 0.62 | 0.53 | -1.90 |
| Rapgef4os3 | 4493.00 | 1.11E-03 | 0.31 | 0.53 | -1.90 |
| Gm15712 | 331.00 | 0.03 | 0.62 | 0.53 | -1.90 |
| Gm26813 | 233.00 | 0.03 | 0.62 | 0.53 | -1.90 |
| Arhgef9 | 746.00 | 1.32E-03 | 0.32 | 0.53 | -1.90 |
| Gm27043 | 54.00 | 0.02 | 0.60 | 0.53 | -1.90 |
| Ppfia2 | 191.47 | 0.01 | 0.58 | 0.53 | -1.90 |

| | | | | | |
|---|---|---|---|---|---|
| Gm7968 | 118.00 | 0.02 | 0.61 | 0.53 | -1.90 |
| Gm43059 | 111.00 | 0.02 | 0.61 | 0.53 | -1.89 |
| Gstm6 | 12415.89 | 3.47E-03 | 0.45 | 0.53 | -1.89 |
| Gm15616 | 96.54 | 0.03 | 0.62 | 0.53 | -1.89 |
| Gm2991 | 74.07 | 0.04 | 0.64 | 0.53 | -1.89 |
| Gm14111 | 107.38 | 0.03 | 0.62 | 0.53 | -1.89 |
| Wsb1 | 3514.00 | 1.85E-03 | 0.36 | 0.53 | -1.89 |
| H2bc15 | 130.58 | 0.03 | 0.62 | 0.53 | -1.88 |
| Cpxm1 | 171.00 | 0.01 | 0.58 | 0.53 | -1.88 |
| Gm44116 | 134.00 | 0.01 | 0.55 | 0.53 | -1.88 |
| Gm18949 | 77.00 | 0.05 | 0.64 | 0.53 | -1.88 |
| Trmt9b | 502.00 | 4.41E-03 | 0.48 | 0.53 | -1.88 |
| Lamc2 | 87.00 | 0.04 | 0.64 | 0.53 | -1.88 |
| Gm43178 | 127.00 | 0.02 | 0.59 | 0.53 | -1.87 |
| Ebf1 | 137.00 | 0.02 | 0.59 | 0.53 | -1.87 |
| Cyp2c39 | 147.00 | 0.02 | 0.59 | 0.53 | -1.87 |
| Gm12264 | 70.66 | 0.03 | 0.61 | 0.53 | -1.87 |
| Gm48872 | 247.00 | 0.01 | 0.58 | 0.53 | -1.87 |
| Slc22a2 | 120.10 | 0.02 | 0.59 | 0.54 | -1.87 |
| Mybl1 | 100.00 | 0.04 | 0.64 | 0.54 | -1.87 |
| Gm37879 | 210.00 | 0.01 | 0.54 | 0.54 | -1.87 |
| Gm37675 | 60.00 | 0.04 | 0.63 | 0.54 | -1.87 |
| Gm43958 | 50.00 | 0.05 | 0.64 | 0.54 | -1.87 |
| Dnah7b | 76.62 | 0.04 | 0.63 | 0.54 | -1.86 |
| Vnn1 | 3063.00 | 0.01 | 0.52 | 0.54 | -1.86 |
| Cnmd | 1102.00 | 1.66E-03 | 0.35 | 0.54 | -1.86 |
| Gm44207 | 103.00 | 0.04 | 0.63 | 0.54 | -1.86 |
| Gm50136 | 1892.48 | 0.02 | 0.59 | 0.54 | -1.86 |
| 2310001K24Rik | 154.03 | 0.01 | 0.58 | 0.54 | -1.86 |
| C230037L18Rik | 289.57 | 0.02 | 0.59 | 0.54 | -1.85 |
| Gm43365 | 189.45 | 0.01 | 0.55 | 0.54 | -1.85 |
| A430035B10Rik | 159.00 | 0.02 | 0.59 | 0.54 | -1.85 |
| Hsd17b6 | 20391.77 | 0.02 | 0.59 | 0.54 | -1.85 |
| Gm42892 | 1479.00 | 4.74E-04 | 0.21 | 0.54 | -1.85 |
| Gm16984 | 279.05 | 0.01 | 0.50 | 0.54 | -1.84 |
| Gm30970 | 175.79 | 0.01 | 0.58 | 0.54 | -1.84 |
| Gm47218 | 185.00 | 0.02 | 0.59 | 0.54 | -1.84 |
| Gm18303 | 109.00 | 0.03 | 0.62 | 0.54 | -1.84 |
| Gm48478 | 1169.00 | 0.04 | 0.63 | 0.54 | -1.84 |
| Gm37423 | 554.00 | 4.52E-03 | 0.48 | 0.54 | -1.84 |
| Gm43389 | 112.00 | 0.02 | 0.59 | 0.54 | -1.84 |
| Gpcpd1 | 24377.97 | 4.65E-03 | 0.48 | 0.55 | -1.83 |
| Lrrc15 | 217.00 | 0.04 | 0.63 | 0.55 | -1.83 |
| Gm15956 | 115.00 | 0.04 | 0.64 | 0.55 | -1.83 |
| Cbx3-ps6 | 128.23 | 0.03 | 0.63 | 0.55 | -1.83 |

| | | | | | |
|---|---|---|---|---|---|
| Relt | 82.00 | 0.04 | 0.64 | 0.55 | -1.82 |
| Acot4 | 11201.39 | 0.01 | 0.53 | 0.55 | -1.82 |
| Gm19667 | 344.00 | 0.01 | 0.58 | 0.55 | -1.82 |
| Gm47283 | 34415.39 | 0.02 | 0.61 | 0.55 | -1.82 |
| Nsl1 | 72.00 | 0.03 | 0.62 | 0.55 | -1.82 |
| Gm48045 | 92.20 | 0.02 | 0.61 | 0.55 | -1.82 |
| Gm15666 | 448.00 | 0.01 | 0.53 | 0.55 | -1.82 |
| G2e3 | 1268.00 | 4.41E-04 | 0.21 | 0.55 | -1.81 |
| E430024P14Rik | 131.81 | 0.02 | 0.60 | 0.55 | -1.81 |
| Gm13675 | 49.01 | 0.05 | 0.64 | 0.55 | -1.81 |
| Gm42941 | 480.00 | 0.01 | 0.58 | 0.55 | -1.81 |
| 1700112D23Rik | 124.00 | 0.02 | 0.61 | 0.55 | -1.81 |
| Ifi214 | 63.50 | 0.04 | 0.64 | 0.55 | -1.81 |
| Tchh | 2048.00 | 0.02 | 0.59 | 0.55 | -1.81 |
| Adamts6 | 375.00 | 0.04 | 0.64 | 0.55 | -1.81 |
| Bhlhb9 | 189.32 | 0.01 | 0.58 | 0.55 | -1.81 |
| Gm9984 | 101.00 | 0.02 | 0.61 | 0.55 | -1.81 |
| Gm10093 | 174.70 | 0.02 | 0.59 | 0.55 | -1.80 |
| Rps13-ps2 | 43.30 | 0.05 | 0.64 | 0.55 | -1.80 |
| Arhgap6 | 1618.11 | 3.30E-03 | 0.44 | 0.56 | -1.80 |
| Evi2 | 498.27 | 3.95E-03 | 0.46 | 0.56 | -1.80 |
| Raet1d | 954.10 | 0.03 | 0.62 | 0.56 | -1.79 |
| Gm22442 | 441.00 | 3.34E-03 | 0.44 | 0.56 | -1.79 |
| Gm6252 | 173.76 | 0.02 | 0.60 | 0.56 | -1.79 |
| Mpzl1 | 298.00 | 0.01 | 0.58 | 0.56 | -1.79 |
| Pcdhga3 | 240.50 | 0.03 | 0.63 | 0.56 | -1.79 |
| Gm28192 | 178.00 | 0.02 | 0.61 | 0.56 | -1.79 |
| Gm14388 | 55.13 | 0.04 | 0.64 | 0.56 | -1.79 |
| Gstt2 | 8522.99 | 3.12E-04 | 0.19 | 0.56 | -1.78 |
| Olfr648 | 91.26 | 0.04 | 0.63 | 0.56 | -1.78 |
| Gm17907 | 71.00 | 0.05 | 0.64 | 0.56 | -1.78 |
| Lypd1 | 109.09 | 0.03 | 0.63 | 0.56 | -1.78 |
| Dbndd1 | 229.00 | 0.02 | 0.59 | 0.56 | -1.78 |
| Ddah1 | 26394.18 | 4.25E-03 | 0.47 | 0.56 | -1.78 |
| Gm9144 | 156.03 | 0.02 | 0.61 | 0.56 | -1.78 |
| Aqp8 | 8337.03 | 3.11E-03 | 0.44 | 0.56 | -1.78 |
| Gm14049 | 155.10 | 0.02 | 0.59 | 0.56 | -1.78 |
| Gm37518 | 357.47 | 0.01 | 0.50 | 0.56 | -1.77 |
| H4c12 | 221.29 | 0.05 | 0.64 | 0.56 | -1.77 |
| Sgk2 | 9390.02 | 0.01 | 0.50 | 0.56 | -1.77 |
| Arhgap5 | 25730.83 | 7.28E-04 | 0.25 | 0.56 | -1.77 |
| Gm20517 | 339.66 | 0.01 | 0.58 | 0.57 | -1.77 |
| Gm42664 | 880.00 | 2.63E-03 | 0.40 | 0.57 | -1.77 |
| Tnfrsf23 | 125.95 | 0.02 | 0.61 | 0.57 | -1.77 |
| Gm12394 | 488.50 | 0.04 | 0.63 | 0.57 | -1.77 |

| | | | | | |
|---|---|---|---|---|---|
| Gm48795 | 144.50 | 0.03 | 0.62 | 0.57 | -1.76 |
| Gm5446 | 86.29 | 0.04 | 0.64 | 0.57 | -1.76 |
| Lrp2bp | 471.00 | 0.04 | 0.63 | 0.57 | -1.76 |
| Pde4d | 1187.00 | 4.05E-03 | 0.46 | 0.57 | -1.76 |
| Gnal | 385.91 | 0.02 | 0.59 | 0.57 | -1.76 |
| B230317F23Rik | 246.00 | 0.04 | 0.63 | 0.57 | -1.76 |
| Rpl39-ps | 115.00 | 0.04 | 0.63 | 0.57 | -1.76 |
| Gm4221 | 281.00 | 0.01 | 0.56 | 0.57 | -1.75 |
| Gm49713 | 134.00 | 0.02 | 0.61 | 0.57 | -1.75 |
| Fzd6 | 1024.00 | 2.59E-03 | 0.40 | 0.57 | -1.75 |
| Epb41l4aos | 960.00 | 1.58E-03 | 0.34 | 0.57 | -1.75 |
| Gm22988 | 6018.70 | 2.14E-03 | 0.38 | 0.57 | -1.75 |
| Gm37137 | 306.00 | 0.01 | 0.54 | 0.57 | -1.75 |
| Gm9920 | 842.40 | 0.01 | 0.53 | 0.57 | -1.75 |
| Kcnj13 | 303.10 | 0.01 | 0.58 | 0.57 | -1.75 |
| Gm2673 | 128.67 | 0.03 | 0.62 | 0.57 | -1.75 |
| Fgf12 | 172.00 | 0.02 | 0.59 | 0.57 | -1.75 |
| Hacl1 | 31482.79 | 8.06E-04 | 0.27 | 0.57 | -1.75 |
| Dclre1c | 2511.61 | 1.07E-03 | 0.31 | 0.57 | -1.74 |
| C230085N15Rik | 54.00 | 0.04 | 0.64 | 0.57 | -1.74 |
| Maff | 1099.00 | 0.01 | 0.54 | 0.57 | -1.74 |
| Ighm | 1746.00 | 0.01 | 0.55 | 0.57 | -1.74 |
| Dnal1 | 273.01 | 0.01 | 0.58 | 0.57 | -1.74 |
| Gm50358 | 117.24 | 0.03 | 0.62 | 0.58 | -1.73 |
| AC154200.1 | 212.00 | 0.02 | 0.61 | 0.58 | -1.73 |
| Arid5b | 37718.47 | 0.02 | 0.59 | 0.58 | -1.73 |
| Tceal1 | 98.00 | 0.03 | 0.62 | 0.58 | -1.73 |
| Spats1 | 79.00 | 0.05 | 0.64 | 0.58 | -1.73 |
| Mamstr | 139.00 | 0.04 | 0.64 | 0.58 | -1.73 |
| Rasl2-9 | 60.00 | 0.04 | 0.64 | 0.58 | -1.73 |
| Tc2n | 591.50 | 0.02 | 0.61 | 0.58 | -1.72 |
| Dnajb7 | 148.00 | 0.04 | 0.64 | 0.58 | -1.72 |
| St8sia4 | 318.00 | 0.01 | 0.58 | 0.58 | -1.72 |
| Tmem246 | 107.00 | 0.04 | 0.63 | 0.58 | -1.72 |
| 9830144P21Rik | 132.33 | 0.03 | 0.62 | 0.58 | -1.72 |
| Dsg1c | 1577.57 | 0.03 | 0.63 | 0.58 | -1.71 |
| Gm13684 | 200.07 | 0.04 | 0.63 | 0.58 | -1.71 |
| Gm47372 | 110.00 | 0.04 | 0.64 | 0.58 | -1.71 |
| C130023A14Rik | 355.00 | 0.01 | 0.53 | 0.58 | -1.71 |
| Pkn3 | 267.00 | 0.01 | 0.58 | 0.58 | -1.71 |
| Gm15163 | 534.35 | 0.01 | 0.53 | 0.58 | -1.71 |
| Gm10390 | 356.94 | 0.01 | 0.58 | 0.58 | -1.71 |
| Crybg3 | 8452.01 | 3.78E-03 | 0.46 | 0.58 | -1.71 |
| Adam12 | 204.00 | 0.03 | 0.61 | 0.58 | -1.71 |
| Gm37233 | 207.00 | 0.02 | 0.61 | 0.59 | -1.71 |

| | | | | | |
|---|---|---|---|---|---|
| Gm16291 | 1389.00 | 0.01 | 0.58 | 0.59 | -1.71 |
| Klf11 | 2787.00 | 0.01 | 0.54 | 0.59 | -1.71 |
| Cda | 790.00 | 0.01 | 0.55 | 0.59 | -1.71 |
| Gm5582 | 97.22 | 0.04 | 0.64 | 0.59 | -1.71 |
| A830035O19Rik | 522.07 | 0.01 | 0.54 | 0.59 | -1.70 |
| Gm45091 | 144.00 | 0.03 | 0.62 | 0.59 | -1.70 |
| Gm13543 | 194.00 | 0.02 | 0.60 | 0.59 | -1.70 |
| Cercam | 97.00 | 0.04 | 0.64 | 0.59 | -1.70 |
| Zdhhc23 | 244.00 | 0.04 | 0.64 | 0.59 | -1.70 |
| Abhd17b | 5673.00 | 0.02 | 0.61 | 0.59 | -1.70 |
| Gm49600 | 175.00 | 0.02 | 0.60 | 0.59 | -1.70 |
| Gm44443 | 375.00 | 0.02 | 0.61 | 0.59 | -1.70 |
| Ndrg1 | 2305.99 | 1.85E-04 | 0.18 | 0.59 | -1.70 |
| Ugt1a5 | 14439.47 | 3.31E-03 | 0.44 | 0.59 | -1.70 |
| Jph1 | 1143.00 | 0.01 | 0.56 | 0.59 | -1.70 |
| Srpk3 | 153.00 | 0.05 | 0.64 | 0.59 | -1.70 |
| Gm42937 | 128.00 | 0.05 | 0.64 | 0.59 | -1.70 |
| Gm6153 | 162.45 | 0.04 | 0.64 | 0.59 | -1.70 |
| Cldn34c1 | 142.21 | 0.05 | 0.64 | 0.59 | -1.70 |
| Gm17066 | 341.42 | 0.03 | 0.63 | 0.59 | -1.70 |
| Gm17300 | 2183.58 | 0.03 | 0.63 | 0.59 | -1.70 |
| Gm17056 | 184.00 | 0.02 | 0.59 | 0.59 | -1.69 |
| Gm19241 | 461.00 | 0.01 | 0.58 | 0.59 | -1.69 |
| Ptgds | 193.00 | 0.03 | 0.62 | 0.59 | -1.69 |
| Gm37215 | 308.00 | 0.01 | 0.53 | 0.59 | -1.69 |
| Slc25a18 | 140.00 | 0.03 | 0.62 | 0.59 | -1.69 |
| Gm47348 | 252.00 | 0.04 | 0.63 | 0.59 | -1.69 |
| Tox | 1879.00 | 0.05 | 0.64 | 0.59 | -1.69 |
| Cers6 | 4948.00 | 0.01 | 0.50 | 0.59 | -1.69 |
| Gm48230 | 1393.00 | 0.04 | 0.63 | 0.59 | -1.69 |
| Gm37788 | 171.00 | 0.04 | 0.63 | 0.59 | -1.69 |
| Gm26447 | 241.03 | 0.05 | 0.64 | 0.59 | -1.69 |
| 1110025M09Rik | 115.00 | 0.05 | 0.64 | 0.59 | -1.69 |
| Gm38057 | 376.00 | 0.01 | 0.53 | 0.59 | -1.69 |
| Rnf125 | 11636.57 | 6.39E-04 | 0.25 | 0.59 | -1.69 |
| Depp1 | 6896.02 | 0.02 | 0.58 | 0.59 | -1.69 |
| Gm29609 | 273.39 | 0.02 | 0.61 | 0.59 | -1.68 |
| Rgs3 | 3039.14 | 1.91E-03 | 0.37 | 0.59 | -1.68 |
| Serpina7 | 3306.00 | 0.01 | 0.55 | 0.59 | -1.68 |
| Gm9311 | 130.00 | 0.05 | 0.64 | 0.60 | -1.68 |
| B930036N10Rik | 238.86 | 0.05 | 0.64 | 0.60 | -1.68 |
| H3f3aos | 198.02 | 0.03 | 0.61 | 0.60 | -1.68 |
| Gm44005 | 1791.00 | 0.02 | 0.61 | 0.60 | -1.67 |
| Gm26684 | 2945.30 | 2.03E-03 | 0.38 | 0.60 | -1.67 |
| Gm32699 | 133.00 | 0.04 | 0.64 | 0.60 | -1.67 |

| | | | | | |
|---|---|---|---|---|---|
| B930059L03Rik | 269.00 | 0.02 | 0.61 | 0.60 | -1.67 |
| Gm43999 | 1459.00 | 0.02 | 0.59 | 0.60 | -1.67 |
| Gm20616 | 136.00 | 0.03 | 0.63 | 0.60 | -1.67 |
| Hhex | 35792.01 | 2.85E-03 | 0.43 | 0.60 | -1.67 |
| Sco2 | 4421.74 | 1.39E-04 | 0.17 | 0.60 | -1.67 |
| Aqp4 | 917.00 | 0.04 | 0.64 | 0.60 | -1.67 |
| Erbb4 | 2985.00 | 8.65E-04 | 0.28 | 0.60 | -1.67 |
| 4732463B04Rik | 646.39 | 0.04 | 0.64 | 0.60 | -1.67 |
| Spred1 | 4779.00 | 3.57E-04 | 0.20 | 0.60 | -1.66 |
| Rgs16 | 9242.29 | 0.04 | 0.64 | 0.60 | -1.66 |
| Klf9 | 40596.14 | 1.14E-03 | 0.31 | 0.60 | -1.66 |
| Gm37080 | 391.00 | 0.03 | 0.62 | 0.60 | -1.66 |
| Thbs1 | 369.00 | 0.02 | 0.61 | 0.60 | -1.66 |
| Gm16268 | 184.31 | 0.04 | 0.63 | 0.60 | -1.65 |
| Cdc45 | 227.00 | 0.02 | 0.61 | 0.60 | -1.65 |
| Abcc3 | 65316.53 | 5.98E-04 | 0.24 | 0.60 | -1.65 |
| Slc16a7 | 13719.64 | 0.02 | 0.59 | 0.60 | -1.65 |
| Chst7 | 411.00 | 0.02 | 0.60 | 0.60 | -1.65 |
| Esm1 | 147.00 | 0.05 | 0.64 | 0.61 | -1.65 |
| Acp1 | 8622.71 | 1.23E-04 | 0.16 | 0.61 | -1.65 |
| 1010001N08Rik | 260.00 | 0.03 | 0.62 | 0.61 | -1.65 |
| Cd36 | 4456.00 | 4.47E-03 | 0.48 | 0.61 | -1.65 |
| B430119L08Rik | 8828.95 | 0.01 | 0.56 | 0.61 | -1.64 |
| Ar | 5501.00 | 0.05 | 0.64 | 0.61 | -1.64 |
| Gm6425 | 176.88 | 0.04 | 0.64 | 0.61 | -1.64 |
| Resf1 | 11016.00 | 3.54E-04 | 0.20 | 0.61 | -1.64 |
| Gm20721 | 409.54 | 0.02 | 0.61 | 0.61 | -1.64 |
| Gsta2 | 2192.49 | 0.01 | 0.52 | 0.61 | -1.64 |
| Retreg1 | 7704.01 | 8.20E-04 | 0.27 | 0.61 | -1.64 |
| Retsat | 53840.98 | 3.59E-03 | 0.45 | 0.61 | -1.64 |
| Gm44981 | 618.00 | 0.04 | 0.63 | 0.61 | -1.64 |
| Shld3 | 425.35 | 0.03 | 0.62 | 0.61 | -1.64 |
| Tbc1d30 | 3052.03 | 0.01 | 0.58 | 0.61 | -1.64 |
| Gm48194 | 143.00 | 0.04 | 0.63 | 0.61 | -1.63 |
| Gm9887 | 269.48 | 0.02 | 0.61 | 0.61 | -1.63 |
| Card11 | 146.00 | 0.04 | 0.64 | 0.61 | -1.63 |
| Gstm2-ps1 | 1411.06 | 0.01 | 0.58 | 0.61 | -1.63 |
| Heyl | 653.00 | 0.01 | 0.53 | 0.61 | -1.63 |
| Adgrg2 | 267.00 | 0.02 | 0.61 | 0.61 | -1.63 |
| Gm28791 | 276.00 | 0.03 | 0.63 | 0.61 | -1.63 |
| Igf2bp3 | 219.90 | 0.04 | 0.64 | 0.61 | -1.63 |
| Gpr157 | 1395.00 | 1.53E-03 | 0.33 | 0.61 | -1.63 |
| Gm9042 | 282.00 | 0.02 | 0.59 | 0.62 | -1.63 |
| Gm4673 | 6069.01 | 1.33E-03 | 0.32 | 0.62 | -1.63 |
| C130051F05Rik | 285.00 | 0.04 | 0.63 | 0.62 | -1.63 |

| | | | | | |
|---|---|---|---|---|---|
| AC160336.1 | 1103.72 | 0.03 | 0.63 | 0.62 | -1.62 |
| Gm25291 | 1324.00 | 0.01 | 0.53 | 0.62 | -1.62 |
| Pnldc1 | 327.00 | 0.04 | 0.64 | 0.62 | -1.62 |
| Gm26083 | 896.60 | 0.02 | 0.59 | 0.62 | -1.62 |
| Gm28530 | 202.57 | 0.04 | 0.63 | 0.62 | -1.62 |
| Pkib | 121.00 | 0.05 | 0.64 | 0.62 | -1.62 |
| Gm18760 | 629.00 | 0.01 | 0.58 | 0.62 | -1.62 |
| Gm43201 | 290.00 | 0.02 | 0.61 | 0.62 | -1.62 |
| Lrrtm2 | 752.00 | 0.01 | 0.54 | 0.62 | -1.62 |
| Dcaf6 | 8374.99 | 2.85E-04 | 0.19 | 0.62 | -1.61 |
| Cdk5r1 | 211.00 | 0.04 | 0.64 | 0.62 | -1.61 |
| Cyp2c54 | 39507.63 | 0.01 | 0.52 | 0.62 | -1.61 |
| Camk1d | 14726.01 | 2.39E-03 | 0.40 | 0.62 | -1.61 |
| Gm5470 | 278.23 | 0.02 | 0.60 | 0.62 | -1.61 |
| Exd1 | 965.00 | 0.02 | 0.59 | 0.62 | -1.60 |
| Gm43713 | 171.56 | 0.04 | 0.63 | 0.62 | -1.60 |
| Lig1 | 474.03 | 0.01 | 0.58 | 0.62 | -1.60 |
| Tusc1 | 489.00 | 0.01 | 0.53 | 0.63 | -1.60 |
| Gm37873 | 177.00 | 0.03 | 0.62 | 0.63 | -1.60 |
| Cfhr3 | 135.00 | 0.05 | 0.64 | 0.63 | -1.60 |
| Cr2 | 204.00 | 0.04 | 0.63 | 0.63 | -1.60 |
| Cit | 488.00 | 0.01 | 0.58 | 0.63 | -1.60 |
| Pltp | 7430.85 | 0.02 | 0.60 | 0.63 | -1.59 |
| 9430038I01Rik | 1091.00 | 0.01 | 0.50 | 0.63 | -1.59 |
| Slc8b1 | 10800.00 | 3.99E-03 | 0.46 | 0.63 | -1.59 |
| Gm45853 | 457.00 | 0.04 | 0.63 | 0.63 | -1.59 |
| Gm49526 | 221.00 | 0.04 | 0.64 | 0.63 | -1.59 |
| Gm43323 | 3999.45 | 0.01 | 0.50 | 0.63 | -1.59 |
| Gjc2 | 200.00 | 0.03 | 0.62 | 0.63 | -1.59 |
| Vnn3 | 6859.00 | 3.74E-03 | 0.46 | 0.63 | -1.59 |
| Gm49759 | 270.00 | 0.03 | 0.61 | 0.63 | -1.59 |
| Atxn7l1 | 7117.72 | 0.01 | 0.58 | 0.63 | -1.59 |
| Mapk15 | 2586.00 | 1.91E-03 | 0.37 | 0.63 | -1.58 |
| 9930017N22Rik | 569.00 | 0.05 | 0.64 | 0.63 | -1.58 |
| Gm3734 | 5368.00 | 1.00E-03 | 0.29 | 0.63 | -1.58 |
| Gm4835 | 235.50 | 0.04 | 0.63 | 0.63 | -1.58 |
| Gm18290 | 390.00 | 0.02 | 0.60 | 0.63 | -1.58 |
| Pdgfd | 253.00 | 0.03 | 0.63 | 0.63 | -1.58 |
| Akr1b10 | 244.00 | 0.03 | 0.62 | 0.63 | -1.58 |
| Pard6g | 235.00 | 0.03 | 0.63 | 0.63 | -1.58 |
| 4930480G23Rik | 229.00 | 0.03 | 0.61 | 0.63 | -1.58 |
| Angpt2 | 386.00 | 0.04 | 0.64 | 0.63 | -1.58 |
| Zfp292 | 6238.00 | 1.86E-03 | 0.36 | 0.63 | -1.58 |
| Gm29253 | 794.84 | 0.02 | 0.58 | 0.63 | -1.58 |
| Pcdhgb7 | 359.39 | 0.02 | 0.59 | 0.63 | -1.57 |

| | | | | | |
|---|---|---|---|---|---|
| 4732416N19Rik | 1318.00 | 0.04 | 0.63 | 0.64 | -1.57 |
| 4930551O13Rik | 552.00 | 0.01 | 0.58 | 0.64 | -1.57 |
| Gm42866 | 1469.00 | 0.01 | 0.52 | 0.64 | -1.57 |
| Foxa2 | 9029.98 | 0.02 | 0.60 | 0.64 | -1.57 |
| A330009N23Rik | 181.00 | 0.05 | 0.64 | 0.64 | -1.57 |
| Gm13835 | 370.26 | 0.03 | 0.62 | 0.64 | -1.57 |
| Pknox2 | 356.00 | 0.03 | 0.61 | 0.64 | -1.57 |
| Gm26674 | 251.63 | 0.04 | 0.63 | 0.64 | -1.57 |
| Gm15285 | 510.26 | 0.04 | 0.63 | 0.64 | -1.57 |
| Tymp | 8714.55 | 0.03 | 0.63 | 0.64 | -1.57 |
| Mgst3 | 3027.00 | 3.99E-03 | 0.46 | 0.64 | -1.57 |
| Crebrf | 9545.00 | 0.01 | 0.58 | 0.64 | -1.57 |
| BC002059 | 634.00 | 0.03 | 0.62 | 0.64 | -1.57 |
| Slf1 | 666.00 | 0.02 | 0.59 | 0.64 | -1.57 |
| 3110082I17Rik | 1411.00 | 0.02 | 0.60 | 0.64 | -1.57 |
| Lratd2 | 1372.93 | 0.01 | 0.58 | 0.64 | -1.57 |
| Coq8a | 34771.67 | 0.01 | 0.58 | 0.64 | -1.57 |
| Rai2 | 520.00 | 0.02 | 0.59 | 0.64 | -1.56 |
| 1700047M11Rik | 254.00 | 0.04 | 0.64 | 0.64 | -1.56 |
| Pafah1b3 | 505.00 | 0.02 | 0.59 | 0.64 | -1.56 |
| Gm37589 | 2144.00 | 0.02 | 0.60 | 0.64 | -1.56 |
| Tspan13 | 183.00 | 0.05 | 0.64 | 0.64 | -1.56 |
| Cbx7 | 2788.00 | 4.91E-03 | 0.48 | 0.64 | -1.56 |
| Gm43775 | 4059.00 | 3.25E-03 | 0.44 | 0.64 | -1.56 |
| Gm37522 | 454.00 | 0.03 | 0.62 | 0.64 | -1.56 |
| Klb | 7177.00 | 1.47E-03 | 0.33 | 0.64 | -1.56 |
| Gm13375 | 442.00 | 0.01 | 0.58 | 0.64 | -1.56 |
| Bach2 | 3768.76 | 0.02 | 0.60 | 0.64 | -1.56 |
| Albfm1 | 5599.69 | 3.37E-03 | 0.44 | 0.64 | -1.56 |
| Slc13a2 | 500.61 | 0.05 | 0.64 | 0.64 | -1.56 |
| Gm14540 | 472.55 | 0.04 | 0.64 | 0.64 | -1.55 |
| Has3 | 812.01 | 0.03 | 0.62 | 0.64 | -1.55 |
| Gm31583 | 10427.00 | 0.01 | 0.58 | 0.64 | -1.55 |
| Pced1b | 738.00 | 0.02 | 0.59 | 0.64 | -1.55 |
| Slc25a27 | 335.00 | 0.03 | 0.62 | 0.64 | -1.55 |
| Inmt | 77205.92 | 0.01 | 0.50 | 0.65 | -1.55 |
| Btbd8 | 618.32 | 0.01 | 0.58 | 0.65 | -1.55 |
| Gm28875 | 434.00 | 0.02 | 0.58 | 0.65 | -1.55 |
| Tcaf2 | 214.00 | 0.05 | 0.64 | 0.65 | -1.55 |
| Gas7 | 2561.00 | 0.01 | 0.52 | 0.65 | -1.54 |
| Csnk2a3 | 520.45 | 0.02 | 0.61 | 0.65 | -1.54 |
| Grpel2 | 6794.00 | 2.23E-03 | 0.39 | 0.65 | -1.54 |
| B630019A10Rik | 7639.14 | 3.57E-03 | 0.45 | 0.65 | -1.54 |
| Gpr135 | 559.00 | 0.02 | 0.59 | 0.65 | -1.54 |
| Igsf10 | 755.36 | 0.01 | 0.57 | 0.65 | -1.54 |

| | | | | | |
|---|---|---|---|---|---|
| Pparg | 1850.00 | 0.04 | 0.63 | 0.65 | -1.54 |
| Gm15261 | 301.00 | 0.05 | 0.64 | 0.65 | -1.54 |
| 4933433G19Rik | 184.00 | 0.05 | 0.64 | 0.65 | -1.54 |
| Igfbp3 | 6691.00 | 0.01 | 0.57 | 0.65 | -1.54 |
| Mios | 2596.00 | 0.01 | 0.54 | 0.65 | -1.54 |
| Zfp36l2 | 26800.00 | 0.04 | 0.63 | 0.65 | -1.54 |
| Gm34667 | 680.00 | 0.03 | 0.62 | 0.65 | -1.53 |
| Dcaf12l1 | 607.00 | 0.03 | 0.62 | 0.65 | -1.53 |
| Gm11342 | 1027.00 | 0.02 | 0.59 | 0.65 | -1.53 |
| Gm13369 | 425.95 | 0.02 | 0.59 | 0.65 | -1.53 |
| D130043K22Rik | 945.00 | 0.04 | 0.64 | 0.65 | -1.53 |
| Ccdc141 | 3438.00 | 0.03 | 0.62 | 0.65 | -1.53 |
| Gm16559 | 2792.00 | 0.02 | 0.59 | 0.65 | -1.53 |
| 9130016M20Rik | 21202.00 | 0.01 | 0.58 | 0.65 | -1.53 |
| Mipol1 | 865.64 | 0.03 | 0.61 | 0.65 | -1.53 |
| Ehhadh | 39595.00 | 7.96E-04 | 0.27 | 0.65 | -1.53 |
| 1700012D14Rik | 549.57 | 0.02 | 0.60 | 0.65 | -1.53 |
| 9030622O22Rik | 2129.00 | 0.02 | 0.59 | 0.65 | -1.53 |
| Gm5898 | 564.08 | 0.04 | 0.64 | 0.65 | -1.53 |
| P2ry1 | 3407.93 | 0.04 | 0.64 | 0.65 | -1.53 |
| Rftn2 | 248.10 | 0.04 | 0.63 | 0.65 | -1.53 |
| Gm49979 | 1339.00 | 0.04 | 0.63 | 0.65 | -1.53 |
| Tlr5 | 3714.00 | 0.02 | 0.59 | 0.65 | -1.53 |
| Gm21451 | 488.78 | 0.03 | 0.62 | 0.66 | -1.53 |
| Palmd | 8681.27 | 2.65E-03 | 0.40 | 0.66 | -1.53 |
| Mir100hg | 828.13 | 0.02 | 0.59 | 0.66 | -1.53 |
| Flcn | 6699.99 | 0.02 | 0.59 | 0.66 | -1.52 |
| Mapkapk3 | 921.00 | 0.02 | 0.60 | 0.66 | -1.52 |
| Sgms2 | 16009.00 | 9.47E-04 | 0.28 | 0.66 | -1.52 |
| 9530085L11Rik | 528.00 | 0.03 | 0.61 | 0.66 | -1.52 |
| Klhl24 | 25004.00 | 2.36E-03 | 0.40 | 0.66 | -1.52 |
| Gm37529 | 1261.00 | 0.01 | 0.52 | 0.66 | -1.52 |
| Naaladl2 | 4307.00 | 0.02 | 0.59 | 0.66 | -1.52 |
| Gm37663 | 313.00 | 0.03 | 0.62 | 0.66 | -1.52 |
| Kdm5b | 8934.00 | 0.01 | 0.58 | 0.66 | -1.52 |
| Rgs7bp | 443.00 | 0.02 | 0.61 | 0.66 | -1.52 |
| Ppp1r3c | 6983.00 | 0.04 | 0.64 | 0.66 | -1.52 |
| Gstm2 | 7209.29 | 0.01 | 0.53 | 0.66 | -1.52 |
| Ces1e | 27616.55 | 1.84E-03 | 0.36 | 0.66 | -1.52 |
| Gm17477 | 272.00 | 0.05 | 0.64 | 0.66 | -1.51 |
| Ddb2 | 889.40 | 0.01 | 0.53 | 0.66 | -1.51 |
| Gm43857 | 456.20 | 0.04 | 0.63 | 0.66 | -1.51 |
| Pcmtd2 | 6452.01 | 1.12E-03 | 0.31 | 0.66 | -1.51 |
| Sytl2 | 214.00 | 0.04 | 0.64 | 0.66 | -1.51 |
| Micu3 | 853.00 | 0.01 | 0.58 | 0.66 | -1.51 |

| | | | | | |
|---|---|---|---|---|---|
| Pcdh17 | 1246.00 | 0.01 | 0.58 | 0.66 | -1.51 |
| Pctp | 12307.00 | 1.85E-03 | 0.36 | 0.66 | -1.51 |
| Arhgap29 | 15862.01 | 2.43E-03 | 0.40 | 0.66 | -1.51 |
| Gprin3 | 20866.00 | 3.23E-03 | 0.44 | 0.66 | -1.51 |
| Zfp704 | 4389.50 | 7.30E-04 | 0.25 | 0.66 | -1.51 |
| Ptbp2 | 2892.00 | 0.01 | 0.58 | 0.66 | -1.51 |
| Gmfg | 576.46 | 0.02 | 0.61 | 0.66 | -1.51 |
| Ocln | 3677.00 | 0.04 | 0.64 | 0.66 | -1.51 |
| Gm13339 | 379777.00 | 0.03 | 0.63 | 0.67 | -1.50 |
| Ces1g | 31715.50 | 2.37E-03 | 0.40 | 0.67 | -1.50 |
| Gm49331 | 262.00 | 0.05 | 0.64 | 0.67 | -1.50 |
| Ugt1a1 | 111540.59 | 3.27E-03 | 0.44 | 0.67 | -1.50 |
| Eri2 | 1799.86 | 0.01 | 0.53 | 0.67 | -1.50 |
| Gm15638 | 3405.00 | 0.03 | 0.63 | 0.67 | -1.50 |
| Acvr2b | 1760.00 | 0.01 | 0.58 | 0.67 | -1.50 |
| Tmem72 | 496.25 | 0.02 | 0.59 | 0.67 | -1.50 |
| Clec1a | 1018.00 | 0.02 | 0.59 | 0.67 | -1.50 |
| Gm26911 | 92.00 | 0.04 | 0.64 | 0.67 | -1.50 |
| Tmtc2 | 3547.97 | 0.02 | 0.61 | 0.67 | -1.50 |
| Mical1 | 337.91 | 0.05 | 0.64 | 0.67 | -1.50 |
| Gm6402 | 235.55 | 0.04 | 0.64 | 0.67 | -1.50 |
| Gm49971 | 645.00 | 0.03 | 0.63 | 0.67 | -1.49 |
| Gm26789 | 1049.00 | 0.02 | 0.61 | 0.67 | -1.49 |
| Ankrd44 | 2260.88 | 4.78E-03 | 0.48 | 0.67 | -1.49 |
| Mettl25 | 385.72 | 0.03 | 0.61 | 0.67 | -1.49 |
| Dsc2 | 11200.01 | 1.45E-03 | 0.33 | 0.67 | -1.49 |
| Serpina6 | 42919.00 | 0.01 | 0.58 | 0.67 | -1.49 |
| Gm10762 | 1301.11 | 0.01 | 0.53 | 0.67 | -1.49 |
| 5330406M23Rik | 2293.00 | 0.02 | 0.59 | 0.67 | -1.49 |
| Avpr1a | 4064.00 | 0.04 | 0.63 | 0.67 | -1.49 |
| Spry4 | 4755.00 | 0.01 | 0.50 | 0.67 | -1.49 |
| Rbm47 | 47628.47 | 0.01 | 0.53 | 0.67 | -1.49 |
| Ttc6 | 695.00 | 0.02 | 0.61 | 0.67 | -1.49 |
| Gm38125 | 286.00 | 0.04 | 0.64 | 0.67 | -1.49 |
| Gm48890 | 449.00 | 0.03 | 0.63 | 0.67 | -1.49 |
| Osmr | 1218.00 | 0.02 | 0.61 | 0.67 | -1.49 |
| Gm2225 | 303.39 | 0.04 | 0.64 | 0.67 | -1.49 |
| Chmp4c | 726.23 | 0.02 | 0.59 | 0.67 | -1.49 |
| Lipc | 20218.03 | 9.68E-04 | 0.29 | 0.67 | -1.49 |
| Rdh9 | 13255.69 | 0.02 | 0.59 | 0.67 | -1.48 |
| 9430034N14Rik | 945.00 | 0.03 | 0.62 | 0.67 | -1.48 |
| Plscr2 | 2651.00 | 0.01 | 0.53 | 0.67 | -1.48 |
| Zfpm2 | 916.00 | 0.04 | 0.63 | 0.67 | -1.48 |
| Zfp871 | 37065.75 | 0.01 | 0.50 | 0.67 | -1.48 |
| Sgce | 847.00 | 0.04 | 0.63 | 0.68 | -1.48 |

| | | | | | |
|---|---|---|---|---|---|
| Gm48225 | 4562.00 | 0.02 | 0.59 | 0.68 | -1.48 |
| Gm45191 | 248.00 | 0.05 | 0.64 | 0.68 | -1.48 |
| Stn1 | 473.00 | 0.03 | 0.63 | 0.68 | -1.48 |
| 4633401B06Rik | 809.00 | 0.03 | 0.61 | 0.68 | -1.48 |
| Elovl5 | 55069.08 | 4.07E-03 | 0.46 | 0.68 | -1.48 |
| Gm43800 | 729.00 | 0.03 | 0.63 | 0.68 | -1.48 |
| Pbx1 | 4180.00 | 0.04 | 0.64 | 0.68 | -1.48 |
| Fbxo21 | 27601.07 | 0.01 | 0.53 | 0.68 | -1.48 |
| Sox5 | 8744.24 | 4.29E-03 | 0.48 | 0.68 | -1.48 |
| Foxa1 | 9935.00 | 0.01 | 0.58 | 0.68 | -1.48 |
| Zc3h6 | 571.79 | 0.04 | 0.64 | 0.68 | -1.48 |
| Ica1 | 387.00 | 0.03 | 0.62 | 0.68 | -1.48 |
| Pdgfc | 856.47 | 0.03 | 0.62 | 0.68 | -1.48 |
| Prxl2b | 3861.00 | 0.05 | 0.64 | 0.68 | -1.48 |
| Dpysl3 | 734.00 | 0.01 | 0.58 | 0.68 | -1.48 |
| Gm37728 | 1499.00 | 0.02 | 0.59 | 0.68 | -1.48 |
| Dock7 | 4004.00 | 1.35E-03 | 0.32 | 0.68 | -1.47 |
| Zc2hc1c | 1109.00 | 0.05 | 0.64 | 0.68 | -1.47 |
| Zbed5 | 1112.18 | 0.01 | 0.58 | 0.68 | -1.47 |
| Gm10532 | 443.96 | 0.05 | 0.64 | 0.68 | -1.47 |
| Dnm3os | 321.00 | 0.05 | 0.64 | 0.68 | -1.47 |
| 1190005I06Rik | 1013.95 | 0.01 | 0.56 | 0.68 | -1.47 |
| Gm42763 | 460.00 | 0.05 | 0.64 | 0.68 | -1.47 |
| Pla1a | 5604.00 | 0.01 | 0.54 | 0.68 | -1.47 |
| D230025D16Rik | 14408.54 | 0.01 | 0.53 | 0.68 | -1.47 |
| Gm44187 | 658.00 | 0.03 | 0.62 | 0.68 | -1.47 |
| Bach1 | 15593.03 | 4.00E-03 | 0.46 | 0.68 | -1.47 |
| Hace1 | 1168.00 | 0.02 | 0.59 | 0.68 | -1.47 |
| Cyp4f17 | 2781.00 | 0.03 | 0.62 | 0.68 | -1.47 |
| Asap2 | 1858.29 | 0.02 | 0.59 | 0.68 | -1.47 |
| Calcoco1 | 8635.99 | 3.24E-03 | 0.44 | 0.68 | -1.47 |
| Zswim4 | 3648.00 | 0.02 | 0.59 | 0.68 | -1.47 |
| Gm10603 | 314.64 | 0.05 | 0.64 | 0.68 | -1.47 |
| Cpeb4 | 18055.95 | 0.01 | 0.58 | 0.68 | -1.46 |
| Chkb | 5374.17 | 2.97E-03 | 0.44 | 0.68 | -1.46 |
| Gm37101 | 560.00 | 0.03 | 0.63 | 0.68 | -1.46 |
| Nxt2 | 4265.00 | 0.01 | 0.58 | 0.68 | -1.46 |
| Id1 | 1052.00 | 0.03 | 0.62 | 0.68 | -1.46 |
| Keg1 | 35424.48 | 0.01 | 0.58 | 0.68 | -1.46 |
| Thbs2 | 639.00 | 0.03 | 0.63 | 0.68 | -1.46 |
| Scarna6 | 1455.00 | 0.05 | 0.64 | 0.68 | -1.46 |
| Mapk1ip1 | 1937.34 | 0.01 | 0.58 | 0.69 | -1.46 |
| Tmem267 | 2630.24 | 0.05 | 0.64 | 0.69 | -1.46 |
| Kdm3a | 4516.00 | 0.01 | 0.56 | 0.69 | -1.46 |
| Gm43593 | 700.00 | 0.03 | 0.62 | 0.69 | -1.46 |

| | | | | | |
|---|---|---|---|---|---|
| Impg2 | 1208.00 | 0.02 | 0.61 | 0.69 | -1.46 |
| Erbb3 | 39062.00 | 0.01 | 0.53 | 0.69 | -1.46 |
| Gm13341 | 45999.97 | 0.01 | 0.58 | 0.69 | -1.46 |
| Rnf144a | 736.00 | 0.03 | 0.62 | 0.69 | -1.46 |
| Gm26631 | 344.13 | 0.05 | 0.64 | 0.69 | -1.46 |
| Mcm10 | 18313.96 | 0.05 | 0.64 | 0.69 | -1.46 |
| Mug-ps1 | 25123.00 | 0.01 | 0.58 | 0.69 | -1.46 |
| Ppp2r2d | 4192.66 | 4.04E-03 | 0.46 | 0.69 | -1.45 |
| Arhgef37 | 3354.00 | 0.02 | 0.60 | 0.69 | -1.45 |
| Casp6 | 2585.00 | 0.01 | 0.50 | 0.69 | -1.45 |
| Spred2 | 5375.00 | 0.01 | 0.56 | 0.69 | -1.45 |
| Gm44888 | 695.00 | 0.05 | 0.64 | 0.69 | -1.45 |
| Gm44567 | 453.00 | 0.04 | 0.64 | 0.69 | -1.45 |
| Agpat4 | 332.00 | 0.04 | 0.64 | 0.69 | -1.45 |
| Crot | 50391.12 | 0.01 | 0.58 | 0.69 | -1.45 |
| Ctsc | 13689.98 | 0.05 | 0.64 | 0.69 | -1.45 |
| Mbtd1 | 5040.00 | 2.15E-03 | 0.38 | 0.69 | -1.45 |
| Tut4 | 7357.98 | 0.01 | 0.56 | 0.69 | -1.45 |
| Snhg1 | 1610.36 | 0.02 | 0.61 | 0.69 | -1.45 |
| Rtn4 | 6169.74 | 0.01 | 0.50 | 0.69 | -1.45 |
| Fam13a | 5433.00 | 0.02 | 0.61 | 0.69 | -1.45 |
| Rdh11 | 8297.00 | 0.01 | 0.52 | 0.69 | -1.45 |
| Ago4 | 1098.00 | 0.03 | 0.61 | 0.69 | -1.45 |
| Haus3 | 845.00 | 0.03 | 0.62 | 0.69 | -1.45 |
| Maoa | 1746.00 | 0.01 | 0.58 | 0.69 | -1.45 |
| Mnat1 | 1145.00 | 0.02 | 0.59 | 0.69 | -1.45 |
| Tm4sf4 | 8589.00 | 2.00E-03 | 0.38 | 0.69 | -1.44 |
| Amn1 | 5177.00 | 0.05 | 0.64 | 0.69 | -1.44 |
| Gm10268 | 438.44 | 0.03 | 0.62 | 0.69 | -1.44 |
| Fut8 | 1146.46 | 0.02 | 0.59 | 0.69 | -1.44 |
| Fam126b | 14220.36 | 0.05 | 0.64 | 0.69 | -1.44 |
| Commd10 | 843.00 | 0.02 | 0.59 | 0.69 | -1.44 |
| Gm43571 | 2497.51 | 0.02 | 0.60 | 0.69 | -1.44 |
| Abca8a | 27523.96 | 0.01 | 0.52 | 0.69 | -1.44 |
| Plscr1 | 1744.00 | 0.05 | 0.64 | 0.69 | -1.44 |
| Ptbp3 | 24264.91 | 0.01 | 0.57 | 0.70 | -1.44 |
| Gm38366 | 484.00 | 0.03 | 0.63 | 0.70 | -1.44 |
| Gstt3 | 9655.32 | 0.02 | 0.60 | 0.70 | -1.44 |
| Sms | 1056.59 | 0.02 | 0.59 | 0.70 | -1.44 |
| Gm8834 | 313.00 | 0.04 | 0.63 | 0.70 | -1.44 |
| Ap4m1 | 3373.98 | 0.02 | 0.59 | 0.70 | -1.44 |
| Clec14a | 4520.00 | 0.01 | 0.50 | 0.70 | -1.43 |
| Evi2a | 472.46 | 0.04 | 0.63 | 0.70 | -1.43 |
| Gstm1 | 287637.73 | 0.01 | 0.58 | 0.70 | -1.43 |
| Fnip1 | 12918.01 | 0.01 | 0.49 | 0.70 | -1.43 |

| | | | | | |
|---|---|---|---|---|---|
| Khnyn | 12128.96 | 0.03 | 0.63 | 0.70 | -1.43 |
| Nceh1 | 8256.00 | 0.03 | 0.62 | 0.70 | -1.43 |
| Pogk | 3512.00 | 0.01 | 0.58 | 0.70 | -1.43 |
| Tigd2 | 2285.00 | 0.01 | 0.53 | 0.70 | -1.43 |
| Gm16270 | 856.24 | 0.03 | 0.62 | 0.70 | -1.43 |
| Lpin2 | 81578.82 | 0.01 | 0.53 | 0.70 | -1.43 |
| Gm45630 | 463.00 | 0.04 | 0.63 | 0.70 | -1.43 |
| 7-Mar | 6189.00 | 0.01 | 0.53 | 0.70 | -1.43 |
| Rin2 | 4372.00 | 0.02 | 0.61 | 0.70 | -1.43 |
| Cd1d2 | 2577.77 | 0.03 | 0.63 | 0.70 | -1.43 |
| Tmem184a | 3806.96 | 0.01 | 0.55 | 0.70 | -1.43 |
| Mad2l2 | 1454.00 | 0.04 | 0.63 | 0.70 | -1.43 |
| 4930523C07Rik | 3860.09 | 0.01 | 0.55 | 0.70 | -1.43 |
| Adh7 | 6031.00 | 0.02 | 0.59 | 0.70 | -1.43 |
| Gm37653 | 707.00 | 0.05 | 0.64 | 0.70 | -1.43 |
| Gm48996 | 846.00 | 0.05 | 0.64 | 0.70 | -1.43 |
| Haus1 | 615.00 | 0.03 | 0.62 | 0.70 | -1.43 |
| Gabarapl1 | 54783.38 | 0.01 | 0.54 | 0.70 | -1.43 |
| Acsm5 | 30041.92 | 0.01 | 0.50 | 0.70 | -1.42 |
| Gm7265 | 1220.53 | 0.03 | 0.63 | 0.70 | -1.42 |
| Mtm1 | 2603.00 | 0.02 | 0.59 | 0.70 | -1.42 |
| Sash1 | 15712.37 | 0.01 | 0.50 | 0.70 | -1.42 |
| Gm49797 | 572.02 | 0.04 | 0.64 | 0.70 | -1.42 |
| Aldh3a2 | 70331.79 | 0.01 | 0.53 | 0.70 | -1.42 |
| Phc1 | 866.00 | 0.03 | 0.62 | 0.70 | -1.42 |
| Tasor | 13907.99 | 4.20E-03 | 0.47 | 0.70 | -1.42 |
| Sh2b2 | 1160.00 | 0.03 | 0.61 | 0.70 | -1.42 |
| Supt3 | 1132.00 | 0.02 | 0.59 | 0.70 | -1.42 |
| Gm37420 | 1351.00 | 0.04 | 0.63 | 0.70 | -1.42 |
| Eepd1 | 3988.00 | 0.03 | 0.62 | 0.70 | -1.42 |
| Slc16a9 | 615.51 | 0.04 | 0.64 | 0.71 | -1.42 |
| AU020206 | 1048.00 | 0.03 | 0.61 | 0.71 | -1.42 |
| Aplf | 596.00 | 0.04 | 0.64 | 0.71 | -1.42 |
| 2310001H17Rik | 3135.00 | 0.03 | 0.61 | 0.71 | -1.42 |
| Avl9 | 6680.00 | 0.01 | 0.58 | 0.71 | -1.42 |
| Pgap1 | 8740.00 | 1.19E-03 | 0.31 | 0.71 | -1.42 |
| S100pbp | 5196.00 | 0.01 | 0.54 | 0.71 | -1.42 |
| Atoh8 | 4905.00 | 0.01 | 0.58 | 0.71 | -1.42 |
| Wdr73 | 1938.00 | 0.01 | 0.58 | 0.71 | -1.41 |
| Slc6a9 | 15200.41 | 0.02 | 0.61 | 0.71 | -1.41 |
| Alms1 | 1350.87 | 0.02 | 0.61 | 0.71 | -1.41 |
| Gm10277 | 1146.00 | 0.04 | 0.64 | 0.71 | -1.41 |
| Gm19950 | 2585.00 | 0.02 | 0.59 | 0.71 | -1.41 |
| 6720427I07Rik | 3646.00 | 0.02 | 0.59 | 0.71 | -1.41 |
| Khdrbs3 | 2954.00 | 0.02 | 0.59 | 0.71 | -1.41 |

| | | | | | |
|---|---|---|---|---|---|
| Gm45413 | 522.00 | 0.04 | 0.64 | 0.71 | -1.41 |
| Klf16 | 1938.00 | 0.02 | 0.59 | 0.71 | -1.41 |
| Lncpint | 7754.00 | 0.01 | 0.58 | 0.71 | -1.41 |
| Arl4a | 4094.00 | 0.02 | 0.59 | 0.71 | -1.41 |
| 4833422C13Rik | 1476.86 | 0.02 | 0.61 | 0.71 | -1.41 |
| Mir22hg | 6347.39 | 0.01 | 0.58 | 0.71 | -1.41 |
| Irf6 | 8367.00 | 0.01 | 0.58 | 0.71 | -1.40 |
| Zup1 | 1914.00 | 0.02 | 0.59 | 0.71 | -1.40 |
| Rcl1 | 19214.14 | 4.96E-03 | 0.49 | 0.71 | -1.40 |
| Plscr4 | 911.00 | 0.03 | 0.63 | 0.71 | -1.40 |
| Hecw2 | 2124.00 | 0.01 | 0.58 | 0.71 | -1.40 |
| Cyp3a59 | 4189.00 | 0.03 | 0.62 | 0.71 | -1.40 |
| Hnmt | 3310.00 | 0.04 | 0.63 | 0.71 | -1.40 |
| Gm37033 | 1193.00 | 0.03 | 0.63 | 0.71 | -1.40 |
| Irs2 | 9652.99 | 0.05 | 0.64 | 0.71 | -1.40 |
| Acaa1b | 187937.39 | 0.01 | 0.58 | 0.71 | -1.40 |
| Ccdc163 | 839.17 | 0.03 | 0.62 | 0.71 | -1.40 |
| Cyp2e1 | 1048808.59 | 0.03 | 0.62 | 0.71 | -1.40 |
| Mfsd10 | 1124.00 | 0.03 | 0.61 | 0.72 | -1.40 |
| Gm15411 | 410.81 | 0.05 | 0.64 | 0.72 | -1.40 |
| Wdr45b | 9699.01 | 0.01 | 0.58 | 0.72 | -1.40 |
| Mospd2 | 4053.00 | 0.01 | 0.52 | 0.72 | -1.40 |
| Ttll5 | 4248.00 | 0.02 | 0.59 | 0.72 | -1.39 |
| Stx1b | 1324.00 | 0.04 | 0.64 | 0.72 | -1.39 |
| Zfp655 | 7033.00 | 0.02 | 0.59 | 0.72 | -1.39 |
| Ldb2 | 1063.00 | 0.04 | 0.64 | 0.72 | -1.39 |
| Syne3 | 2780.00 | 0.01 | 0.58 | 0.72 | -1.39 |
| Rnf152 | 9305.58 | 0.01 | 0.53 | 0.72 | -1.39 |
| Gm17251 | 1081.62 | 0.03 | 0.63 | 0.72 | -1.39 |
| Hykk | 32901.96 | 0.01 | 0.56 | 0.72 | -1.39 |
| Tada1 | 2604.40 | 0.01 | 0.56 | 0.72 | -1.39 |
| Gclc | 46748.09 | 0.02 | 0.60 | 0.72 | -1.39 |
| Tsc1 | 9262.99 | 0.01 | 0.54 | 0.72 | -1.39 |
| Ppargc1b | 5761.00 | 0.04 | 0.63 | 0.72 | -1.39 |
| Tab3 | 3020.00 | 0.02 | 0.60 | 0.72 | -1.39 |
| Sugct | 4250.06 | 0.01 | 0.54 | 0.72 | -1.39 |
| Rictor | 10993.00 | 0.01 | 0.58 | 0.72 | -1.39 |
| Steap2 | 1996.00 | 0.03 | 0.62 | 0.72 | -1.39 |
| Ugcg | 6000.00 | 0.01 | 0.53 | 0.72 | -1.39 |
| Gm17491 | 1984.00 | 0.03 | 0.63 | 0.72 | -1.39 |
| Ccng2 | 2055.00 | 0.03 | 0.62 | 0.72 | -1.39 |
| Cdo1 | 198155.64 | 0.01 | 0.53 | 0.72 | -1.39 |
| Gm21399 | 715.71 | 0.03 | 0.62 | 0.72 | -1.39 |
| Nfat5 | 21397.97 | 0.01 | 0.55 | 0.72 | -1.39 |
| Rnpc3 | 3686.87 | 0.02 | 0.61 | 0.72 | -1.39 |

| | | | | | |
|---|---|---|---|---|---|
| Sema4b | 2611.00 | 0.02 | 0.60 | 0.72 | -1.38 |
| Sp4 | 1767.00 | 0.03 | 0.62 | 0.72 | -1.38 |
| Mreg | 6863.00 | 0.01 | 0.58 | 0.72 | -1.38 |
| Acsl1 | 314038.69 | 0.03 | 0.61 | 0.72 | -1.38 |
| Camk2g | 2320.00 | 0.03 | 0.62 | 0.72 | -1.38 |
| Akap8l | 10720.01 | 0.01 | 0.58 | 0.72 | -1.38 |
| Pdk1 | 18683.99 | 0.03 | 0.63 | 0.72 | -1.38 |
| Xbp1 | 49934.16 | 0.03 | 0.62 | 0.72 | -1.38 |
| Cyp3a25 | 73436.38 | 0.01 | 0.58 | 0.72 | -1.38 |
| Zbtb2 | 2107.68 | 0.01 | 0.58 | 0.72 | -1.38 |
| Gm9752 | 786.08 | 0.04 | 0.64 | 0.72 | -1.38 |
| Zkscan17 | 4446.50 | 0.01 | 0.56 | 0.72 | -1.38 |
| Zbtb10 | 2466.00 | 0.05 | 0.64 | 0.72 | -1.38 |
| Nsmce2 | 1474.00 | 0.02 | 0.61 | 0.72 | -1.38 |
| Iigp1 | 41596.05 | 0.01 | 0.58 | 0.72 | -1.38 |
| Shroom2 | 9226.99 | 0.02 | 0.61 | 0.72 | -1.38 |
| Tfap4 | 1212.00 | 0.04 | 0.64 | 0.73 | -1.38 |
| Gm49599 | 3447.68 | 0.01 | 0.58 | 0.73 | -1.38 |
| Arhgap18 | 2050.00 | 0.02 | 0.59 | 0.73 | -1.38 |
| Cdc14b | 7502.00 | 0.01 | 0.53 | 0.73 | -1.38 |
| Haus4 | 2713.00 | 0.02 | 0.61 | 0.73 | -1.38 |
| Gm38372 | 2098.00 | 0.02 | 0.59 | 0.73 | -1.38 |
| Pon1 | 89849.06 | 0.01 | 0.54 | 0.73 | -1.38 |
| Ackr4 | 2964.70 | 0.02 | 0.59 | 0.73 | -1.38 |
| F830016B08Rik | 11933.00 | 0.04 | 0.64 | 0.73 | -1.38 |
| Trim21 | 1156.00 | 0.03 | 0.62 | 0.73 | -1.38 |
| Zfand5 | 10381.99 | 0.01 | 0.50 | 0.73 | -1.38 |
| Rnf217 | 10537.00 | 0.02 | 0.59 | 0.73 | -1.38 |
| Vim | 5253.00 | 0.03 | 0.62 | 0.73 | -1.37 |
| Zfp729b | 1805.85 | 0.02 | 0.59 | 0.73 | -1.37 |
| A1cf | 47402.24 | 0.01 | 0.58 | 0.73 | -1.37 |
| Eci2 | 28873.95 | 0.01 | 0.53 | 0.73 | -1.37 |
| Blvra | 1111.00 | 0.04 | 0.63 | 0.73 | -1.37 |
| Cavin2 | 2898.00 | 0.02 | 0.61 | 0.73 | -1.37 |
| Ypel3 | 10089.99 | 0.01 | 0.52 | 0.73 | -1.37 |
| Gm45769 | 1822.00 | 0.04 | 0.64 | 0.73 | -1.37 |
| Sfxn3 | 505.00 | 0.04 | 0.64 | 0.73 | -1.37 |
| Sqor | 26688.01 | 4.86E-03 | 0.48 | 0.73 | -1.37 |
| Bcl10 | 2152.00 | 0.02 | 0.61 | 0.73 | -1.37 |
| Map4k4 | 3023.00 | 0.02 | 0.61 | 0.73 | -1.37 |
| G6pdx | 1534.90 | 0.04 | 0.63 | 0.73 | -1.37 |
| Ppm1k | 26040.00 | 0.01 | 0.53 | 0.73 | -1.37 |
| Nox4 | 11362.03 | 0.02 | 0.59 | 0.73 | -1.37 |
| Ptpn9 | 4323.00 | 0.01 | 0.53 | 0.73 | -1.37 |
| Odf3b | 1062.00 | 0.04 | 0.63 | 0.73 | -1.37 |

| | | | | | |
|---|---|---|---|---|---|
| Cyp2c23 | 71558.56 | 0.02 | 0.59 | 0.73 | -1.37 |
| Syt3 | 1433.00 | 0.04 | 0.63 | 0.73 | -1.37 |
| Crtc3 | 6446.00 | 0.04 | 0.63 | 0.73 | -1.37 |
| Acox1 | 496840.98 | 0.03 | 0.62 | 0.73 | -1.37 |
| Gm37212 | 991.00 | 0.05 | 0.64 | 0.73 | -1.37 |
| Mospd1 | 1081.55 | 0.03 | 0.61 | 0.73 | -1.37 |
| Fzd8 | 9975.00 | 0.04 | 0.63 | 0.73 | -1.36 |
| Serpinb6a | 1908.59 | 0.01 | 0.58 | 0.73 | -1.36 |
| Zfand6 | 10474.00 | 0.01 | 0.58 | 0.73 | -1.36 |
| Rabep2 | 2515.26 | 0.02 | 0.61 | 0.73 | -1.36 |
| Pclo | 1342.00 | 0.03 | 0.62 | 0.73 | -1.36 |
| Mtx3 | 4521.00 | 0.01 | 0.58 | 0.73 | -1.36 |
| Nubpl | 1332.68 | 0.02 | 0.61 | 0.73 | -1.36 |
| Rida | 54422.48 | 0.01 | 0.55 | 0.73 | -1.36 |
| Cept1 | 7914.01 | 0.01 | 0.58 | 0.73 | -1.36 |
| 2410002F23Rik | 3164.91 | 0.04 | 0.64 | 0.73 | -1.36 |
| Tet2 | 7037.99 | 0.03 | 0.63 | 0.73 | -1.36 |
| Pnpla2 | 21333.96 | 0.03 | 0.62 | 0.73 | -1.36 |
| Zfp626 | 1305.00 | 0.04 | 0.63 | 0.73 | -1.36 |
| Tprkb | 19256.77 | 4.05E-03 | 0.46 | 0.73 | -1.36 |
| Dyrk1b | 6405.99 | 0.03 | 0.63 | 0.74 | -1.36 |
| Taf7 | 902.00 | 0.04 | 0.64 | 0.74 | -1.36 |
| Irf2bp2 | 22601.40 | 0.05 | 0.64 | 0.74 | -1.36 |
| 9330162G02Rik | 6587.53 | 0.04 | 0.63 | 0.74 | -1.36 |
| Htatip2 | 2546.00 | 0.02 | 0.61 | 0.74 | -1.36 |
| Nsd3 | 35057.10 | 0.04 | 0.63 | 0.74 | -1.36 |
| Nbn | 2709.00 | 0.01 | 0.56 | 0.74 | -1.36 |
| 1810008I18Rik | 12748.96 | 0.02 | 0.61 | 0.74 | -1.36 |
| Sgms1 | 4494.00 | 0.01 | 0.53 | 0.74 | -1.36 |
| Rbak | 981.00 | 0.04 | 0.64 | 0.74 | -1.36 |
| Ap4s1 | 1180.00 | 0.03 | 0.61 | 0.74 | -1.36 |
| Ces2a | 62680.11 | 0.01 | 0.53 | 0.74 | -1.35 |
| Rasa2 | 1080.00 | 0.05 | 0.64 | 0.74 | -1.35 |
| Lurap1l | 11650.00 | 0.03 | 0.62 | 0.74 | -1.35 |
| Rora | 27027.07 | 0.01 | 0.58 | 0.74 | -1.35 |
| Tgoln1 | 36996.00 | 0.02 | 0.61 | 0.74 | -1.35 |
| Fabp4 | 1069.00 | 0.04 | 0.64 | 0.74 | -1.35 |
| Tns2 | 30005.12 | 0.02 | 0.59 | 0.74 | -1.35 |
| Leprot | 2786.00 | 0.01 | 0.58 | 0.74 | -1.35 |
| Cnr2 | 1394.00 | 0.04 | 0.64 | 0.74 | -1.35 |
| Spg20 | 7802.00 | 0.04 | 0.63 | 0.74 | -1.35 |
| Gm5424 | 11630.52 | 0.03 | 0.62 | 0.74 | -1.35 |
| Myef2 | 3012.78 | 0.01 | 0.58 | 0.74 | -1.35 |
| Pld1 | 1936.00 | 0.05 | 0.64 | 0.74 | -1.35 |
| Cblb | 3866.00 | 0.02 | 0.61 | 0.74 | -1.35 |

| | | | | | |
|---|---|---|---|---|---|
| Nsun6 | 785.00 | 0.05 | 0.64 | 0.74 | -1.35 |
| Ccdc71l | 1371.00 | 0.03 | 0.63 | 0.74 | -1.35 |
| Rab9 | 5638.00 | 0.02 | 0.59 | 0.74 | -1.35 |
| Smc4 | 2088.00 | 0.04 | 0.63 | 0.74 | -1.35 |
| Cyp2c50 | 149349.11 | 0.05 | 0.64 | 0.74 | -1.35 |
| Rapgef4 | 39866.48 | 0.03 | 0.63 | 0.74 | -1.35 |
| 4632427E13Rik | 5444.01 | 0.03 | 0.62 | 0.74 | -1.35 |
| Zfp609 | 13041.00 | 0.03 | 0.62 | 0.74 | -1.35 |
| Manea | 3840.00 | 0.03 | 0.61 | 0.74 | -1.35 |
| Fbxl17 | 8353.01 | 0.01 | 0.54 | 0.74 | -1.35 |
| Gm47483 | 945.00 | 0.05 | 0.64 | 0.74 | -1.35 |
| Zfp60 | 2224.43 | 0.03 | 0.62 | 0.74 | -1.35 |
| Amdhd1 | 34899.00 | 0.03 | 0.62 | 0.74 | -1.35 |
| Pth1r | 2163.00 | 0.03 | 0.62 | 0.74 | -1.35 |
| Phf1 | 5167.76 | 0.01 | 0.54 | 0.74 | -1.35 |
| Jade1 | 13897.78 | 0.01 | 0.58 | 0.74 | -1.35 |
| Rsbn1 | 3175.42 | 0.03 | 0.62 | 0.74 | -1.35 |
| Dbp | 30655.63 | 0.02 | 0.59 | 0.74 | -1.35 |
| Sav1 | 3241.96 | 0.04 | 0.64 | 0.74 | -1.35 |
| Fbxl3 | 7592.00 | 0.01 | 0.54 | 0.74 | -1.34 |
| Ctdsp2 | 17738.96 | 0.01 | 0.56 | 0.74 | -1.34 |
| Dapk1 | 17837.06 | 0.02 | 0.58 | 0.74 | -1.34 |
| Rcor3 | 4571.93 | 0.03 | 0.63 | 0.74 | -1.34 |
| Cfap20 | 5938.16 | 0.01 | 0.58 | 0.74 | -1.34 |
| Gm22009 | 17966.00 | 0.04 | 0.64 | 0.74 | -1.34 |
| Tiparp | 1356.78 | 0.03 | 0.63 | 0.74 | -1.34 |
| Vps26c | 2599.00 | 0.02 | 0.60 | 0.74 | -1.34 |
| Cdk5rap2 | 1964.20 | 0.03 | 0.61 | 0.74 | -1.34 |
| Cd84 | 1256.00 | 0.03 | 0.63 | 0.74 | -1.34 |
| Acacb | 33824.01 | 0.01 | 0.58 | 0.75 | -1.34 |
| Slc23a2 | 6801.00 | 0.02 | 0.59 | 0.75 | -1.34 |
| Rnf146 | 3570.16 | 0.02 | 0.61 | 0.75 | -1.34 |
| Mcu | 5089.20 | 0.03 | 0.62 | 0.75 | -1.34 |
| Slc27a1 | 2050.00 | 0.04 | 0.63 | 0.75 | -1.34 |
| Lama4 | 2132.00 | 0.03 | 0.62 | 0.75 | -1.34 |
| Akr1c19 | 21257.97 | 0.05 | 0.64 | 0.75 | -1.34 |
| Dsp | 23405.68 | 0.02 | 0.60 | 0.75 | -1.34 |
| Fmo1 | 33597.35 | 0.04 | 0.64 | 0.75 | -1.34 |
| Prpf39 | 5952.15 | 0.04 | 0.63 | 0.75 | -1.34 |
| Foxk1 | 14920.00 | 0.02 | 0.59 | 0.75 | -1.34 |
| Usp28 | 1625.00 | 0.04 | 0.64 | 0.75 | -1.34 |
| Inca1 | 2103.00 | 0.04 | 0.64 | 0.75 | -1.34 |
| Aspg | 35400.07 | 0.03 | 0.62 | 0.75 | -1.34 |
| Prkn | 1726.00 | 0.03 | 0.63 | 0.75 | -1.34 |
| Maob | 27737.10 | 0.01 | 0.58 | 0.75 | -1.34 |

| | | | | | |
|---|---|---|---|---|---|
| Erg | 1918.00 | 0.03 | 0.62 | 0.75 | -1.34 |
| Slc35a5 | 4385.00 | 0.03 | 0.63 | 0.75 | -1.34 |
| Cdk19 | 2642.72 | 0.03 | 0.62 | 0.75 | -1.34 |
| Cnppd1 | 13171.98 | 0.02 | 0.59 | 0.75 | -1.34 |
| Tmem106b | 45196.04 | 0.02 | 0.59 | 0.75 | -1.34 |
| Nt5c3 | 2934.00 | 0.02 | 0.59 | 0.75 | -1.34 |
| Gk | 15283.96 | 0.02 | 0.61 | 0.75 | -1.34 |
| Acadl | 30711.00 | 0.02 | 0.60 | 0.75 | -1.34 |
| Zbtb39 | 5109.00 | 0.03 | 0.61 | 0.75 | -1.33 |
| Il15ra | 11564.00 | 0.02 | 0.59 | 0.75 | -1.33 |
| E030022I16Rik | 1299.00 | 0.04 | 0.64 | 0.75 | -1.33 |
| Dleu2 | 5590.00 | 0.04 | 0.64 | 0.75 | -1.33 |
| Reps1 | 3491.00 | 0.02 | 0.61 | 0.75 | -1.33 |
| Efnb2 | 1248.00 | 0.03 | 0.63 | 0.75 | -1.33 |
| Slc22a5 | 4612.14 | 0.02 | 0.61 | 0.75 | -1.33 |
| Nfib | 57465.90 | 0.01 | 0.58 | 0.75 | -1.33 |
| Mef2a | 12650.33 | 0.01 | 0.58 | 0.75 | -1.33 |
| Sclt1 | 1375.00 | 0.03 | 0.63 | 0.75 | -1.33 |
| Sdr42e1 | 20300.95 | 0.02 | 0.59 | 0.75 | -1.33 |
| Rgn | 139702.00 | 0.02 | 0.61 | 0.75 | -1.33 |
| Znrf3 | 5894.00 | 0.02 | 0.59 | 0.75 | -1.33 |
| Arhgap24 | 2438.00 | 0.05 | 0.64 | 0.75 | -1.33 |
| 9130221H12Rik | 1028.70 | 0.05 | 0.64 | 0.75 | -1.33 |
| D130020L05Rik | 4575.04 | 0.02 | 0.59 | 0.75 | -1.33 |
| Rbms1 | 10290.01 | 0.01 | 0.54 | 0.75 | -1.33 |
| Srsf5 | 38909.25 | 0.02 | 0.61 | 0.75 | -1.33 |
| Gm25395 | 3907.00 | 0.05 | 0.64 | 0.75 | -1.33 |
| Slc4a4 | 26473.01 | 0.02 | 0.59 | 0.75 | -1.33 |
| Nhsl2 | 1037.00 | 0.04 | 0.64 | 0.75 | -1.33 |
| Fam210a | 29205.46 | 0.01 | 0.52 | 0.75 | -1.33 |
| Cldn2 | 10762.99 | 0.05 | 0.64 | 0.75 | -1.33 |
| Epc2 | 3629.00 | 0.03 | 0.62 | 0.75 | -1.33 |
| Cobll1 | 19938.98 | 0.02 | 0.59 | 0.75 | -1.33 |
| Acsm3 | 17245.17 | 0.04 | 0.63 | 0.75 | -1.33 |
| Nr2c2 | 9838.14 | 0.03 | 0.62 | 0.75 | -1.33 |
| Ktn1 | 5430.00 | 0.02 | 0.59 | 0.75 | -1.33 |
| Prpf38a | 1281.00 | 0.05 | 0.64 | 0.75 | -1.33 |
| Zbtb44 | 23052.04 | 0.02 | 0.60 | 0.75 | -1.33 |
| Sesn1 | 2845.00 | 0.03 | 0.61 | 0.75 | -1.33 |
| Fbxl20 | 14445.99 | 0.02 | 0.59 | 0.76 | -1.32 |
| Bend7 | 2650.00 | 0.04 | 0.63 | 0.76 | -1.32 |
| Cnot6l | 13892.02 | 0.01 | 0.58 | 0.76 | -1.32 |
| Snx21 | 2477.56 | 0.03 | 0.61 | 0.76 | -1.32 |
| Gatm | 3132.00 | 0.04 | 0.63 | 0.76 | -1.32 |
| Agtr1a | 10206.00 | 0.02 | 0.59 | 0.76 | -1.32 |

| | | | | | |
|---|---|---|---|---|---|
| Asxl1 | 8324.75 | 0.02 | 0.61 | 0.76 | -1.32 |
| Snapc2 | 1788.00 | 0.04 | 0.63 | 0.76 | -1.32 |
| N4bp2 | 14023.51 | 0.02 | 0.59 | 0.76 | -1.32 |
| Robo4 | 3387.00 | 0.04 | 0.64 | 0.76 | -1.32 |
| Adgrl2 | 19506.19 | 0.01 | 0.58 | 0.76 | -1.32 |
| AL731706.1 | 1707.48 | 0.04 | 0.63 | 0.76 | -1.32 |
| Afmid | 26044.55 | 0.02 | 0.59 | 0.76 | -1.32 |
| Asf1a | 1593.64 | 0.05 | 0.64 | 0.76 | -1.32 |
| Pign | 3445.70 | 0.03 | 0.62 | 0.76 | -1.32 |
| Nmt2 | 5466.00 | 0.02 | 0.59 | 0.76 | -1.32 |
| Zfp532 | 1310.24 | 0.05 | 0.64 | 0.76 | -1.32 |
| Mms19 | 7896.45 | 0.04 | 0.64 | 0.76 | -1.32 |
| Utp11 | 2694.00 | 0.03 | 0.61 | 0.76 | -1.32 |
| E330009J07Rik | 3366.00 | 0.03 | 0.63 | 0.76 | -1.32 |
| Suox | 22942.00 | 0.01 | 0.58 | 0.76 | -1.32 |
| Cradd | 3047.08 | 0.04 | 0.64 | 0.76 | -1.32 |
| Slc40a1 | 25197.02 | 0.03 | 0.62 | 0.76 | -1.32 |
| Rabgap1l | 14104.59 | 0.01 | 0.58 | 0.76 | -1.32 |
| Ttc39b | 13784.98 | 0.02 | 0.59 | 0.76 | -1.32 |
| Dcaf17 | 3429.00 | 0.02 | 0.61 | 0.76 | -1.32 |
| Ces1c | 336150.62 | 0.03 | 0.62 | 0.76 | -1.31 |
| Eva1a | 13056.02 | 0.02 | 0.59 | 0.76 | -1.31 |
| Soat2 | 6229.00 | 0.03 | 0.62 | 0.76 | -1.31 |
| Klf12 | 17033.00 | 0.02 | 0.59 | 0.76 | -1.31 |
| Hmgcl | 27474.11 | 0.02 | 0.59 | 0.76 | -1.31 |
| Smim13 | 6312.00 | 0.03 | 0.62 | 0.76 | -1.31 |
| Lin7a | 4862.00 | 0.03 | 0.62 | 0.76 | -1.31 |
| Rad50 | 2725.00 | 0.03 | 0.62 | 0.76 | -1.31 |
| Fcho2 | 11728.29 | 0.05 | 0.64 | 0.76 | -1.31 |
| Pla2r1 | 1570.69 | 0.04 | 0.64 | 0.76 | -1.31 |
| Phka2 | 5907.00 | 0.02 | 0.61 | 0.76 | -1.31 |
| Angptl4 | 18722.97 | 0.05 | 0.64 | 0.76 | -1.31 |
| Smurf1 | 6890.99 | 0.04 | 0.63 | 0.76 | -1.31 |
| Gm38120 | 1995.00 | 0.05 | 0.64 | 0.76 | -1.31 |
| Wrn | 3602.00 | 0.03 | 0.61 | 0.76 | -1.31 |
| Zfp141 | 2088.34 | 0.04 | 0.63 | 0.76 | -1.31 |
| Eif2ak4 | 3036.00 | 0.04 | 0.64 | 0.76 | -1.31 |
| Stk24 | 5100.00 | 0.02 | 0.60 | 0.76 | -1.31 |
| Zcchc2 | 14457.02 | 0.04 | 0.64 | 0.76 | -1.31 |
| Add3 | 4242.00 | 0.02 | 0.59 | 0.76 | -1.31 |
| Cwf19l2 | 3593.00 | 0.04 | 0.63 | 0.76 | -1.31 |
| Ugt2a3 | 46534.70 | 0.02 | 0.59 | 0.76 | -1.31 |
| Rcn2 | 962.00 | 0.05 | 0.64 | 0.76 | -1.31 |
| 2810459M11Rik | 16139.04 | 0.04 | 0.64 | 0.76 | -1.31 |
| Ptpn12 | 4878.00 | 0.03 | 0.62 | 0.77 | -1.31 |

| | | | | | |
|---|---|---|---|---|---|
| Esco1 | 2719.78 | 0.03 | 0.62 | 0.77 | -1.31 |
| Dpys | 44400.45 | 0.02 | 0.59 | 0.77 | -1.31 |
| Rit1 | 1698.00 | 0.05 | 0.64 | 0.77 | -1.31 |
| Tm6sf2 | 4016.01 | 0.04 | 0.64 | 0.77 | -1.31 |
| Pdlim5 | 16267.99 | 0.02 | 0.61 | 0.77 | -1.30 |
| Ing2 | 1797.00 | 0.05 | 0.64 | 0.77 | -1.30 |
| Ireb2 | 21604.00 | 0.02 | 0.59 | 0.77 | -1.30 |
| Atp11b | 9892.00 | 0.02 | 0.61 | 0.77 | -1.30 |
| AW549877 | 8185.00 | 0.05 | 0.64 | 0.77 | -1.30 |
| Pdgfrb | 3056.00 | 0.03 | 0.63 | 0.77 | -1.30 |
| Dennd4a | 13704.02 | 0.03 | 0.62 | 0.77 | -1.30 |
| Stard9 | 3645.40 | 0.04 | 0.63 | 0.77 | -1.30 |
| Snx33 | 6446.00 | 0.02 | 0.59 | 0.77 | -1.30 |
| Ddx28 | 2394.00 | 0.03 | 0.63 | 0.77 | -1.30 |
| Sat1 | 6551.00 | 0.02 | 0.61 | 0.77 | -1.30 |
| Inpp4a | 10021.00 | 0.03 | 0.62 | 0.77 | -1.30 |
| Dnmbp | 13452.00 | 0.03 | 0.62 | 0.77 | -1.30 |
| Marcks | 5473.00 | 0.03 | 0.62 | 0.77 | -1.30 |
| Herc3 | 1930.00 | 0.04 | 0.64 | 0.77 | -1.30 |
| Appl2 | 4421.00 | 0.03 | 0.62 | 0.77 | -1.30 |
| Xpa | 2066.00 | 0.04 | 0.64 | 0.77 | -1.30 |
| Echdc3 | 11293.99 | 0.01 | 0.58 | 0.77 | -1.30 |
| Kdm3b | 10808.99 | 0.04 | 0.63 | 0.77 | -1.30 |
| Btg1 | 9430.73 | 0.03 | 0.61 | 0.77 | -1.30 |
| Zdhhc20 | 2972.00 | 0.02 | 0.61 | 0.77 | -1.30 |
| Ehbp1 | 3268.00 | 0.04 | 0.63 | 0.77 | -1.30 |
| 4732465J04Rik | 3504.00 | 0.04 | 0.64 | 0.77 | -1.30 |
| Kidins220 | 19498.78 | 0.01 | 0.58 | 0.77 | -1.30 |
| Rbm33 | 31584.90 | 0.03 | 0.62 | 0.77 | -1.30 |
| Myo1d | 5348.00 | 0.02 | 0.61 | 0.77 | -1.30 |
| Clec2d | 40351.00 | 0.02 | 0.59 | 0.77 | -1.30 |
| Dram2 | 6948.00 | 0.03 | 0.63 | 0.77 | -1.30 |
| Cirbp | 3187.00 | 0.05 | 0.64 | 0.77 | -1.30 |
| 1810026B05Rik | 4181.37 | 0.05 | 0.64 | 0.77 | -1.30 |
| Gpbp1l1 | 15689.50 | 0.02 | 0.59 | 0.77 | -1.30 |
| Mfap1a | 4384.01 | 0.04 | 0.64 | 0.77 | -1.30 |
| Mccc1 | 8614.84 | 0.02 | 0.60 | 0.77 | -1.30 |
| Malt1 | 2979.00 | 0.03 | 0.62 | 0.77 | -1.30 |
| Thoc2l | 7050.00 | 0.03 | 0.61 | 0.77 | -1.30 |
| Styx | 4791.41 | 0.04 | 0.64 | 0.77 | -1.30 |
| Akirin2 | 3685.00 | 0.04 | 0.63 | 0.77 | -1.30 |
| Stag2 | 15657.04 | 0.03 | 0.62 | 0.77 | -1.29 |
| Dyrk2 | 21663.29 | 0.03 | 0.63 | 0.77 | -1.29 |
| Chd6 | 7748.00 | 0.02 | 0.61 | 0.77 | -1.29 |
| Cryl1 | 2730.00 | 0.03 | 0.63 | 0.77 | -1.29 |

| | | | | | |
|---|---|---|---|---|---|
| Acss2 | 41473.84 | 0.03 | 0.62 | 0.77 | -1.29 |
| Ccnl1 | 7158.17 | 0.05 | 0.64 | 0.77 | -1.29 |
| Gm42715 | 4074.92 | 0.04 | 0.63 | 0.77 | -1.29 |
| Elk3 | 3309.35 | 0.03 | 0.63 | 0.77 | -1.29 |
| Rps7 | 18361.84 | 0.01 | 0.58 | 0.77 | -1.29 |
| Fcf1 | 2290.74 | 0.04 | 0.63 | 0.77 | -1.29 |
| Trim23 | 2079.00 | 0.05 | 0.64 | 0.77 | -1.29 |
| Dtd2 | 2296.00 | 0.04 | 0.63 | 0.77 | -1.29 |
| Gstt1 | 33148.97 | 0.01 | 0.58 | 0.77 | -1.29 |
| Tek | 3195.00 | 0.02 | 0.60 | 0.77 | -1.29 |
| Btbd7 | 9326.00 | 0.03 | 0.62 | 0.77 | -1.29 |
| Btd | 10562.01 | 0.03 | 0.62 | 0.77 | -1.29 |
| Ahnak | 17381.00 | 0.02 | 0.59 | 0.77 | -1.29 |
| Ces2e | 33783.03 | 0.02 | 0.60 | 0.77 | -1.29 |
| Bag4 | 11595.60 | 0.03 | 0.62 | 0.77 | -1.29 |
| Prxl2c | 15363.10 | 0.02 | 0.59 | 0.77 | -1.29 |
| Ubxn7 | 13534.98 | 0.02 | 0.59 | 0.78 | -1.29 |
| Pik3c2b | 3419.00 | 0.02 | 0.61 | 0.78 | -1.29 |
| Sec24b | 9015.99 | 0.03 | 0.62 | 0.78 | -1.29 |
| Erf | 9974.00 | 0.05 | 0.64 | 0.78 | -1.29 |
| Fam76b | 2107.00 | 0.05 | 0.64 | 0.78 | -1.29 |
| Gtpbp10 | 2951.77 | 0.04 | 0.64 | 0.78 | -1.29 |
| Zfp422 | 2471.11 | 0.04 | 0.63 | 0.78 | -1.29 |
| Papss2 | 41497.24 | 0.03 | 0.62 | 0.78 | -1.29 |
| Rnf167 | 8256.61 | 0.03 | 0.61 | 0.78 | -1.29 |
| Mga | 15477.01 | 0.04 | 0.64 | 0.78 | -1.29 |
| C77080 | 42744.42 | 0.03 | 0.62 | 0.78 | -1.29 |
| Odc1 | 11721.84 | 0.02 | 0.60 | 0.78 | -1.29 |
| Eif4a2 | 32168.10 | 0.02 | 0.59 | 0.78 | -1.29 |
| Cdc42bpg | 6081.00 | 0.04 | 0.64 | 0.78 | -1.29 |
| Eci1 | 14266.04 | 0.02 | 0.61 | 0.78 | -1.29 |
| 1700001C19Rik | 2054.74 | 0.05 | 0.64 | 0.78 | -1.29 |
| Samd8 | 13846.77 | 0.02 | 0.59 | 0.78 | -1.29 |
| Oit3 | 6966.00 | 0.03 | 0.63 | 0.78 | -1.29 |
| Upb1 | 50412.01 | 0.02 | 0.61 | 0.78 | -1.29 |
| Pdcd4 | 21211.00 | 0.03 | 0.62 | 0.78 | -1.29 |
| Irf2bpl | 13842.86 | 0.05 | 0.64 | 0.78 | -1.28 |
| Tmem56 | 56296.48 | 0.05 | 0.64 | 0.78 | -1.28 |
| Nqo2 | 15486.01 | 0.01 | 0.58 | 0.78 | -1.28 |
| Ngrn | 2067.00 | 0.04 | 0.64 | 0.78 | -1.28 |
| Hook2 | 6745.00 | 0.03 | 0.63 | 0.78 | -1.28 |
| Tbc1d20 | 9384.99 | 0.02 | 0.60 | 0.78 | -1.28 |
| Ilvbl | 4221.00 | 0.03 | 0.62 | 0.78 | -1.28 |
| Gucd1 | 32935.02 | 0.04 | 0.64 | 0.78 | -1.28 |
| Sypl | 10720.00 | 0.04 | 0.63 | 0.78 | -1.28 |

| | | | | | |
|---|---|---|---|---|---|
| Mtus1 | 61517.92 | 0.05 | 0.64 | 0.78 | -1.28 |
| Phlpp1 | 9952.00 | 0.03 | 0.62 | 0.78 | -1.28 |
| Gstm7 | 8667.00 | 0.04 | 0.64 | 0.78 | -1.28 |
| Strbp | 17664.97 | 0.03 | 0.63 | 0.78 | -1.28 |
| Clk1 | 11503.00 | 0.04 | 0.63 | 0.78 | -1.28 |
| Slc6a6 | 39422.00 | 0.03 | 0.61 | 0.78 | -1.28 |
| Zfp639 | 2662.00 | 0.05 | 0.64 | 0.78 | -1.28 |
| Lnpep | 22020.00 | 0.03 | 0.62 | 0.78 | -1.28 |
| Phip | 8323.00 | 0.04 | 0.63 | 0.78 | -1.28 |
| Rbpms | 10416.00 | 0.05 | 0.64 | 0.78 | -1.28 |
| Akip1 | 2256.00 | 0.05 | 0.64 | 0.78 | -1.28 |
| Bclaf1 | 16426.93 | 0.02 | 0.61 | 0.78 | -1.28 |
| Akr1d1 | 24691.00 | 0.03 | 0.62 | 0.78 | -1.28 |
| 9930021J03Rik | 8156.00 | 0.04 | 0.64 | 0.78 | -1.28 |
| Dixdc1 | 4380.00 | 0.05 | 0.64 | 0.78 | -1.28 |
| Acad8 | 9348.99 | 0.03 | 0.63 | 0.78 | -1.28 |
| Nr1d2 | 23965.04 | 0.02 | 0.61 | 0.78 | -1.28 |
| Kdm4c | 2553.00 | 0.05 | 0.64 | 0.78 | -1.28 |
| Chst15 | 6017.53 | 0.03 | 0.63 | 0.78 | -1.28 |
| Cpne1 | 4622.59 | 0.03 | 0.63 | 0.78 | -1.28 |
| Heca | 10806.01 | 0.02 | 0.61 | 0.78 | -1.28 |
| Brd8 | 3921.00 | 0.04 | 0.64 | 0.78 | -1.27 |
| Dyrk1a | 7948.00 | 0.03 | 0.62 | 0.78 | -1.27 |
| Rev3l | 8333.01 | 0.03 | 0.63 | 0.78 | -1.27 |
| Sp3 | 10676.00 | 0.04 | 0.63 | 0.79 | -1.27 |
| Abhd17c | 15630.99 | 0.04 | 0.63 | 0.79 | -1.27 |
| Rbbp4 | 25865.78 | 0.04 | 0.63 | 0.79 | -1.27 |
| Zwint | 9802.99 | 0.03 | 0.63 | 0.79 | -1.27 |
| Pcf11 | 9709.70 | 0.03 | 0.62 | 0.79 | -1.27 |
| Gstm4 | 7968.98 | 0.03 | 0.63 | 0.79 | -1.27 |
| Tet3 | 19732.00 | 0.04 | 0.64 | 0.79 | -1.27 |
| Calcrl | 4748.00 | 0.03 | 0.63 | 0.79 | -1.27 |
| Rap1a | 7113.23 | 0.03 | 0.62 | 0.79 | -1.27 |
| Acot12 | 25795.85 | 0.02 | 0.61 | 0.79 | -1.27 |
| Selenbp1 | 51085.03 | 0.02 | 0.59 | 0.79 | -1.27 |
| Ces1f | 70734.92 | 0.03 | 0.62 | 0.79 | -1.27 |
| Nicn1 | 2105.00 | 0.05 | 0.64 | 0.79 | -1.27 |
| Ppfibp1 | 7397.01 | 0.04 | 0.63 | 0.79 | -1.27 |
| Kdr | 18680.97 | 0.03 | 0.62 | 0.79 | -1.27 |
| Zbtb1 | 4558.00 | 0.04 | 0.63 | 0.79 | -1.27 |
| Decr1 | 20151.00 | 0.03 | 0.62 | 0.79 | -1.27 |
| Pnisr | 13597.99 | 0.04 | 0.63 | 0.79 | -1.27 |
| Tk2 | 3418.00 | 0.05 | 0.64 | 0.79 | -1.27 |
| Sin3a | 10167.00 | 0.04 | 0.63 | 0.79 | -1.27 |
| Usp6nl | 8045.53 | 0.04 | 0.63 | 0.79 | -1.27 |

| | | | | | |
|---|---|---|---|---|---|
| Acad10 | 6111.41 | 0.04 | 0.63 | 0.79 | -1.27 |
| Cbx3 | 6467.24 | 0.03 | 0.62 | 0.79 | -1.27 |
| Mier3 | 5051.00 | 0.04 | 0.63 | 0.79 | -1.26 |
| Ezh1 | 4454.00 | 0.05 | 0.64 | 0.79 | -1.26 |
| Smlr1 | 6518.00 | 0.04 | 0.64 | 0.79 | -1.26 |
| Wac | 17766.64 | 0.04 | 0.63 | 0.79 | -1.26 |
| Klf13 | 39853.03 | 0.03 | 0.62 | 0.79 | -1.26 |
| Ube2h | 30455.97 | 0.02 | 0.61 | 0.79 | -1.26 |
| Asah2 | 5898.00 | 0.04 | 0.64 | 0.79 | -1.26 |
| Shprh | 6945.00 | 0.04 | 0.64 | 0.79 | -1.26 |
| Kiz | 2460.00 | 0.05 | 0.64 | 0.79 | -1.26 |
| Rps3a3 | 6033.54 | 0.05 | 0.64 | 0.79 | -1.26 |
| Slc39a8 | 6625.00 | 0.05 | 0.64 | 0.79 | -1.26 |
| Csnk2a2 | 7106.00 | 0.04 | 0.63 | 0.79 | -1.26 |
| L3hypdh | 4324.00 | 0.04 | 0.64 | 0.79 | -1.26 |
| Zranb1 | 11111.00 | 0.04 | 0.64 | 0.79 | -1.26 |
| Fmr1 | 4432.98 | 0.04 | 0.63 | 0.79 | -1.26 |
| Large1 | 2063.00 | 0.05 | 0.64 | 0.79 | -1.26 |
| Acadm | 42972.08 | 0.02 | 0.61 | 0.79 | -1.26 |
| Fbxl5 | 5359.00 | 0.03 | 0.62 | 0.79 | -1.26 |
| BC024386 | 54620.32 | 0.05 | 0.64 | 0.79 | -1.26 |
| Tspan12 | 22937.02 | 0.02 | 0.59 | 0.79 | -1.26 |
| Sft2d1 | 4553.75 | 0.04 | 0.64 | 0.80 | -1.26 |
| Thnsl2 | 6376.00 | 0.04 | 0.63 | 0.80 | -1.26 |
| Sh3d19 | 35921.78 | 0.03 | 0.62 | 0.80 | -1.26 |
| Rilp | 4426.00 | 0.04 | 0.64 | 0.80 | -1.26 |
| Cd99l2 | 5402.99 | 0.03 | 0.62 | 0.80 | -1.26 |
| Dhtkd1 | 19822.97 | 0.05 | 0.64 | 0.80 | -1.26 |
| Kng2 | 34262.75 | 0.03 | 0.63 | 0.80 | -1.26 |
| Gramd3 | 7877.99 | 0.04 | 0.63 | 0.80 | -1.26 |
| Cryz | 10843.95 | 0.04 | 0.63 | 0.80 | -1.26 |
| Clk4 | 7436.00 | 0.04 | 0.64 | 0.80 | -1.26 |
| Acvr2a | 7488.53 | 0.05 | 0.64 | 0.80 | -1.26 |
| Ralgapa2 | 24621.04 | 0.03 | 0.62 | 0.80 | -1.25 |
| Abhd4 | 7532.99 | 0.04 | 0.64 | 0.80 | -1.25 |
| Rabgap1 | 6208.00 | 0.04 | 0.64 | 0.80 | -1.25 |
| Npc1 | 47525.76 | 0.03 | 0.62 | 0.80 | -1.25 |
| Hibadh | 50344.49 | 0.04 | 0.64 | 0.80 | -1.25 |
| Pum1 | 19803.99 | 0.04 | 0.64 | 0.80 | -1.25 |
| Prkaa1 | 3673.00 | 0.05 | 0.64 | 0.80 | -1.25 |
| Tlk2 | 5408.92 | 0.05 | 0.64 | 0.80 | -1.25 |
| Mtmr3 | 10842.99 | 0.05 | 0.64 | 0.80 | -1.25 |
| Syne2 | 33733.99 | 0.05 | 0.64 | 0.80 | -1.25 |
| Etfdh | 34269.66 | 0.04 | 0.63 | 0.80 | -1.25 |
| Mllt10 | 8614.36 | 0.05 | 0.64 | 0.80 | -1.25 |

| | | | | | |
|---|---|---|---|---|---|
| Nectin3 | 10349.00 | 0.04 | 0.64 | 0.80 | -1.25 |
| Kif1b | 73844.86 | 0.03 | 0.61 | 0.80 | -1.25 |
| Ripk1 | 8891.89 | 0.05 | 0.64 | 0.80 | -1.25 |
| Arih2 | 10471.36 | 0.04 | 0.63 | 0.80 | -1.25 |
| Tab2 | 15659.95 | 0.05 | 0.64 | 0.80 | -1.25 |
| Eif4b | 48135.14 | 0.04 | 0.63 | 0.80 | -1.25 |
| Pex7 | 11955.00 | 0.03 | 0.61 | 0.80 | -1.25 |
| Sult2a8 | 96944.10 | 0.05 | 0.64 | 0.80 | -1.25 |
| Tmem245 | 24565.02 | 0.05 | 0.64 | 0.80 | -1.25 |
| Smad2 | 6006.00 | 0.04 | 0.63 | 0.80 | -1.25 |
| Sbds | 8378.99 | 0.05 | 0.64 | 0.80 | -1.25 |
| Tfdp2 | 6355.00 | 0.05 | 0.64 | 0.80 | -1.25 |
| Uhrf1bp1l | 15850.00 | 0.03 | 0.62 | 0.80 | -1.25 |
| Cir1 | 3692.64 | 0.05 | 0.64 | 0.80 | -1.25 |
| Atn1 | 16150.22 | 0.04 | 0.64 | 0.80 | -1.25 |
| Chdh | 48575.54 | 0.04 | 0.64 | 0.80 | -1.25 |
| Ormdl1 | 4229.63 | 0.04 | 0.64 | 0.80 | -1.24 |
| Pan3 | 11267.00 | 0.05 | 0.64 | 0.80 | -1.24 |
| Aldh6a1 | 82426.05 | 0.05 | 0.64 | 0.80 | -1.24 |
| Ncor1 | 42004.87 | 0.05 | 0.64 | 0.80 | -1.24 |
| Eif5 | 47827.61 | 0.05 | 0.64 | 0.80 | -1.24 |
| Osbpl1a | 21667.21 | 0.05 | 0.64 | 0.80 | -1.24 |
| Ppp1r12a | 8956.00 | 0.04 | 0.63 | 0.80 | -1.24 |
| Stab2 | 27839.74 | 0.05 | 0.64 | 0.80 | -1.24 |
| Paics | 32391.74 | 0.04 | 0.64 | 0.80 | -1.24 |
| Rhou | 14992.00 | 0.04 | 0.63 | 0.80 | -1.24 |
| Cul3 | 12350.97 | 0.04 | 0.64 | 0.80 | -1.24 |
| Arl5a | 34550.00 | 0.05 | 0.64 | 0.81 | -1.24 |
| Trim33 | 6646.00 | 0.04 | 0.64 | 0.81 | -1.24 |
| Eea1 | 13716.01 | 0.04 | 0.64 | 0.81 | -1.24 |
| Nfx1 | 12773.99 | 0.04 | 0.63 | 0.81 | -1.24 |
| Gtf2h5 | 4203.00 | 0.04 | 0.64 | 0.81 | -1.24 |
| Suclg2 | 35856.80 | 0.03 | 0.63 | 0.81 | -1.24 |
| Rbbp6 | 17795.98 | 0.04 | 0.64 | 0.81 | -1.24 |
| Stard4 | 31769.00 | 0.04 | 0.64 | 0.81 | -1.24 |
| Ppm1a | 24069.02 | 0.03 | 0.62 | 0.81 | -1.24 |
| Adcy6 | 10518.96 | 0.04 | 0.64 | 0.81 | -1.24 |
| Lpcat3 | 23530.81 | 0.04 | 0.63 | 0.81 | -1.24 |
| Ugt2b36 | 76194.77 | 0.04 | 0.63 | 0.81 | -1.24 |
| Hsdl2 | 12460.95 | 0.05 | 0.64 | 0.81 | -1.24 |
| Aldh5a1 | 18766.89 | 0.03 | 0.62 | 0.81 | -1.24 |
| Lpp | 34895.01 | 0.04 | 0.63 | 0.81 | -1.24 |
| Abcc6 | 51691.00 | 0.04 | 0.64 | 0.81 | -1.24 |
| Cd82 | 15263.97 | 0.04 | 0.63 | 0.81 | -1.24 |
| Rc3h1 | 20128.00 | 0.05 | 0.64 | 0.81 | -1.23 |

| | | | | | |
|---|---|---|---|---|---|
| Creg1 | 55696.08 | 0.04 | 0.64 | 0.81 | -1.23 |
| Tjp1 | 17271.00 | 0.05 | 0.64 | 0.81 | -1.23 |
| Plxnd1 | 15614.00 | 0.05 | 0.64 | 0.81 | -1.23 |
| Hbp1 | 17093.16 | 0.04 | 0.64 | 0.81 | -1.23 |
| Kat2b | 22741.84 | 0.04 | 0.63 | 0.81 | -1.23 |
| Oxa1l | 10229.27 | 0.05 | 0.64 | 0.81 | -1.23 |
| Decr2 | 29282.02 | 0.04 | 0.64 | 0.81 | -1.23 |
| Tmem243 | 14723.30 | 0.04 | 0.63 | 0.81 | -1.23 |
| Adi1 | 24118.02 | 0.04 | 0.64 | 0.81 | -1.23 |
| Slc44a1 | 10992.22 | 0.04 | 0.63 | 0.81 | -1.23 |
| Cisd1 | 20167.00 | 0.05 | 0.64 | 0.82 | -1.23 |
| Smad4 | 9915.98 | 0.05 | 0.64 | 0.82 | -1.22 |
| Spp2 | 11308.00 | 0.04 | 0.64 | 0.82 | -1.22 |
| C2cd2 | 41758.96 | 0.05 | 0.64 | 0.82 | -1.22 |
| Atl3 | 37826.84 | 0.04 | 0.64 | 0.82 | -1.22 |
| Cds2 | 24579.07 | 0.05 | 0.64 | 0.82 | -1.22 |
| Cth | 38559.11 | 0.04 | 0.64 | 0.83 | -1.21 |
| Plxnb1 | 22015.93 | 0.05 | 0.64 | 1.25 | 1.25 |
| Slc35b1 | 9632.00 | 0.05 | 0.64 | 1.25 | 1.25 |
| Dnajb11 | 17204.03 | 0.04 | 0.64 | 1.26 | 1.26 |
| Ikbke | 5594.00 | 0.05 | 0.64 | 1.28 | 1.28 |
| Stip1 | 15053.84 | 0.05 | 0.64 | 1.29 | 1.29 |
| Pdia6 | 39593.78 | 0.03 | 0.62 | 1.29 | 1.29 |
| Shb | 14069.01 | 0.02 | 0.60 | 1.30 | 1.30 |
| Slc22a3 | 2002.51 | 0.02 | 0.61 | 1.33 | 1.33 |
| Mettl1 | 1490.00 | 0.05 | 0.64 | 1.33 | 1.33 |
| Ankrd33b | 14012.95 | 0.01 | 0.58 | 1.34 | 1.34 |
| Mid1ip1 | 17547.87 | 0.02 | 0.60 | 1.35 | 1.35 |
| Frmd8 | 5532.77 | 0.01 | 0.58 | 1.36 | 1.36 |
| Utp14b | 5306.98 | 0.01 | 0.58 | 1.37 | 1.37 |
| Hes6 | 18842.99 | 0.02 | 0.61 | 1.37 | 1.37 |
| Nnmt | 11472.01 | 0.01 | 0.58 | 1.38 | 1.38 |
| Tsc22d3 | 26082.97 | 0.02 | 0.59 | 1.38 | 1.38 |
| Ppl | 5246.00 | 0.03 | 0.62 | 1.39 | 1.39 |
| Ppp1r3b | 69026.90 | 0.04 | 0.64 | 1.39 | 1.39 |
| Srm | 7413.00 | 0.01 | 0.52 | 1.40 | 1.40 |
| Cad | 6068.00 | 0.02 | 0.60 | 1.40 | 1.40 |
| Dnaja4 | 1439.14 | 0.04 | 0.63 | 1.41 | 1.41 |
| Hyou1 | 81551.09 | 0.02 | 0.60 | 1.42 | 1.42 |
| Per2 | 18814.00 | 0.02 | 0.59 | 1.42 | 1.42 |
| Cyp2d40 | 13942.92 | 0.01 | 0.54 | 1.42 | 1.42 |
| Srebf1 | 125981.42 | 0.01 | 0.56 | 1.42 | 1.42 |
| Slc4a1 | 353.00 | 0.04 | 0.64 | 1.42 | 1.42 |
| Lrg1 | 96635.00 | 0.02 | 0.60 | 1.42 | 1.42 |
| Tkfc | 82094.48 | 0.02 | 0.59 | 1.43 | 1.43 |

| | | | | | |
|---|---|---|---|---|---|
| Mcm5 | 441.00 | 0.04 | 0.64 | 1.44 | 1.44 |
| Nfkbiz | 3211.00 | 0.01 | 0.50 | 1.44 | 1.44 |
| Tat | 310857.70 | 0.01 | 0.54 | 1.44 | 1.44 |
| Slc3a1 | 12321.15 | 0.01 | 0.50 | 1.44 | 1.44 |
| Serpina1e | 3234433.61 | 0.04 | 0.64 | 1.45 | 1.45 |
| Gm20634 | 3396.90 | 0.01 | 0.53 | 1.45 | 1.45 |
| Gm43611 | 4992.00 | 0.01 | 0.58 | 1.46 | 1.46 |
| H2ac4 | 660.24 | 0.02 | 0.61 | 1.47 | 1.47 |
| 2810408I11Rik | 330.47 | 0.05 | 0.64 | 1.47 | 1.47 |
| Wnt5b | 1123.00 | 0.03 | 0.63 | 1.48 | 1.48 |
| Ccnd1 | 5529.48 | 0.02 | 0.59 | 1.49 | 1.49 |
| Tsku | 75890.80 | 3.32E-03 | 0.44 | 1.49 | 1.49 |
| Gm5844 | 784.60 | 0.05 | 0.64 | 1.50 | 1.50 |
| Oasl1 | 1647.00 | 0.02 | 0.61 | 1.50 | 1.50 |
| Fkbp11 | 1919.00 | 0.01 | 0.53 | 1.51 | 1.51 |
| Csf1 | 3106.00 | 0.01 | 0.54 | 1.51 | 1.51 |
| Saa4 | 100155.00 | 0.01 | 0.50 | 1.51 | 1.51 |
| Sox9 | 1996.00 | 0.01 | 0.58 | 1.51 | 1.51 |
| Alpl | 3869.00 | 0.02 | 0.60 | 1.51 | 1.51 |
| Bdnf | 662.00 | 0.03 | 0.62 | 1.52 | 1.52 |
| Rcan1 | 5177.00 | 0.01 | 0.52 | 1.52 | 1.52 |
| Saa3 | 639.00 | 0.03 | 0.62 | 1.53 | 1.53 |
| Sik1 | 12142.99 | 0.04 | 0.64 | 1.54 | 1.54 |
| Acnat2 | 2363.52 | 0.02 | 0.60 | 1.54 | 1.54 |
| Tuba1c | 23471.96 | 1.73E-03 | 0.35 | 1.54 | 1.54 |
| Slc41a2 | 9189.00 | 0.03 | 0.63 | 1.54 | 1.54 |
| Col15a1 | 1875.00 | 0.02 | 0.61 | 1.55 | 1.55 |
| Fasn | 339850.78 | 0.01 | 0.53 | 1.56 | 1.56 |
| Agap2 | 2257.00 | 0.01 | 0.58 | 1.56 | 1.56 |
| Fgl1 | 21144.03 | 0.01 | 0.56 | 1.58 | 1.58 |
| Enho | 5276.00 | 0.01 | 0.58 | 1.58 | 1.58 |
| Gtpbp4-ps1 | 654.04 | 0.03 | 0.63 | 1.59 | 1.59 |
| C4a | 38703.21 | 0.01 | 0.58 | 1.59 | 1.59 |
| Gm44502 | 262.93 | 0.04 | 0.64 | 1.59 | 1.59 |
| Bag3 | 19867.00 | 0.02 | 0.61 | 1.59 | 1.59 |
| Insc | 2874.00 | 2.62E-03 | 0.40 | 1.60 | 1.60 |
| Cxcl11 | 647.33 | 0.01 | 0.58 | 1.60 | 1.60 |
| Hsf5 | 135.00 | 0.05 | 0.64 | 1.61 | 1.61 |
| Celsr1 | 6203.99 | 1.21E-03 | 0.31 | 1.61 | 1.61 |
| Thrsp | 128256.36 | 2.16E-03 | 0.38 | 1.62 | 1.62 |
| BC049987 | 2760.00 | 4.66E-04 | 0.21 | 1.63 | 1.63 |
| Tnfaip3 | 1052.00 | 0.04 | 0.63 | 1.63 | 1.63 |
| Gm43195 | 698.00 | 0.01 | 0.53 | 1.63 | 1.63 |
| Gm43702 | 691.00 | 0.04 | 0.63 | 1.64 | 1.64 |
| Slc17a9 | 1392.00 | 3.87E-03 | 0.46 | 1.64 | 1.64 |

| | | | | | |
|---|---|---|---|---|---|
| Fam47e | 3856.00 | 9.25E-04 | 0.28 | 1.65 | 1.65 |
| Gm7290 | 171.74 | 0.03 | 0.61 | 1.65 | 1.65 |
| S1pr5 | 1998.00 | 1.48E-03 | 0.33 | 1.65 | 1.65 |
| Col27a1 | 30165.02 | 1.17E-03 | 0.31 | 1.66 | 1.66 |
| Serpina12 | 10354.00 | 0.01 | 0.58 | 1.66 | 1.66 |
| Noct | 21520.70 | 0.02 | 0.59 | 1.67 | 1.67 |
| Fam186a | 292.00 | 0.05 | 0.64 | 1.67 | 1.67 |
| Sdf2l1 | 8334.00 | 0.01 | 0.52 | 1.68 | 1.68 |
| Hsp90aa1 | 38979.31 | 0.01 | 0.57 | 1.69 | 1.69 |
| Nmnat2 | 119.00 | 0.03 | 0.63 | 1.73 | 1.73 |
| Hsph1 | 13167.99 | 0.03 | 0.62 | 1.73 | 1.73 |
| Fam81a | 494.00 | 0.03 | 0.61 | 1.75 | 1.75 |
| B430219N15Rik | 768.66 | 0.01 | 0.58 | 1.78 | 1.78 |
| E030018B13Rik | 1150.00 | 0.03 | 0.62 | 1.78 | 1.78 |
| Tsc22d1 | 16315.95 | 5.99E-04 | 0.24 | 1.78 | 1.78 |
| Smpd3 | 803.34 | 4.55E-03 | 0.48 | 1.79 | 1.79 |
| Fam83f | 161.09 | 0.02 | 0.60 | 1.80 | 1.80 |
| Gnat2 | 179.00 | 0.02 | 0.59 | 1.82 | 1.82 |
| Rbp1 | 1999.00 | 6.51E-04 | 0.25 | 1.84 | 1.84 |
| Zfp454 | 188.00 | 0.02 | 0.59 | 1.84 | 1.84 |
| Cyp2d41-ps | 392.41 | 0.02 | 0.59 | 1.87 | 1.87 |
| Vmn2r20 | 175.82 | 0.01 | 0.53 | 1.89 | 1.89 |
| C130026I21Rik | 73.61 | 0.05 | 0.64 | 1.90 | 1.90 |
| Noxred1 | 66.00 | 0.04 | 0.64 | 1.92 | 1.92 |
| Ttbk1 | 1150.00 | 3.10E-03 | 0.44 | 1.92 | 1.92 |
| Gm20507 | 98.31 | 0.04 | 0.64 | 1.92 | 1.92 |
| Nat8f5 | 2745.67 | 1.88E-04 | 0.18 | 1.94 | 1.94 |
| Xist | 415.89 | 0.03 | 0.62 | 1.94 | 1.94 |
| Creld2 | 8788.00 | 0.01 | 0.58 | 1.96 | 1.96 |
| Gm44620 | 208.00 | 0.01 | 0.53 | 1.96 | 1.96 |
| Gm45416 | 114.00 | 0.01 | 0.54 | 1.99 | 1.99 |
| Sox5os1 | 153.76 | 0.01 | 0.54 | 2.04 | 2.04 |
| Gm28876 | 61.00 | 0.03 | 0.61 | 2.07 | 2.07 |
| Ntrk2 | 1157.00 | 0.01 | 0.58 | 2.07 | 2.07 |
| Pmaip1 | 88.00 | 0.01 | 0.53 | 2.08 | 2.08 |
| Zfp9 | 1049.00 | 0.03 | 0.62 | 2.09 | 2.09 |
| Sox5it | 35.00 | 0.05 | 0.64 | 2.12 | 2.12 |
| Atp2b2 | 447.00 | 0.02 | 0.59 | 2.12 | 2.12 |
| Cyp2d34 | 41.06 | 0.03 | 0.62 | 2.16 | 2.16 |
| Stambp-ps1 | 90.00 | 0.02 | 0.59 | 2.16 | 2.16 |
| Gm4211 | 44.20 | 0.04 | 0.63 | 2.16 | 2.16 |
| Gm16337 | 33.00 | 0.04 | 0.64 | 2.19 | 2.19 |
| Serpina5 | 64.00 | 0.04 | 0.63 | 2.23 | 2.23 |
| Nat8f6 | 410.63 | 4.53E-04 | 0.21 | 2.25 | 2.25 |
| Mmd2 | 781.00 | 0.02 | 0.58 | 2.25 | 2.25 |

| | | | | | |
|---|---|---|---|---|---|
| Pdxp | 176.99 | 0.01 | 0.58 | 2.27 | 2.27 |
| Tsix | 150.56 | 0.03 | 0.62 | 2.28 | 2.28 |
| Crybb3 | 113.00 | 0.05 | 0.64 | 2.28 | 2.28 |
| Slc8a1 | 7837.05 | 1.72E-04 | 0.18 | 2.29 | 2.29 |
| Lrp2 | 950.00 | 3.02E-04 | 0.19 | 2.31 | 2.31 |
| Derl3 | 1332.00 | 0.01 | 0.53 | 2.32 | 2.32 |
| Lcn2 | 1124.00 | 1.15E-04 | 0.16 | 2.40 | 2.40 |
| Chrm1 | 117.00 | 0.01 | 0.52 | 2.41 | 2.41 |
| Gm7993 | 295.52 | 0.01 | 0.50 | 2.43 | 2.43 |
| Ift57 | 189.00 | 0.01 | 0.53 | 2.44 | 2.44 |
| Mroh6 | 306.00 | 1.83E-04 | 0.18 | 2.50 | 2.50 |
| Gm47748 | 36.00 | 0.02 | 0.61 | 2.50 | 2.50 |
| Gm17396 | 36.00 | 0.03 | 0.62 | 2.50 | 2.50 |
| Gm29427 | 135.77 | 0.03 | 0.62 | 2.53 | 2.53 |
| Cyp21a1 | 707.25 | 0.03 | 0.61 | 2.53 | 2.53 |
| Slc35g3 | 34.00 | 0.01 | 0.58 | 2.61 | 2.61 |
| Gm50044 | 93.95 | 3.48E-03 | 0.45 | 2.68 | 2.68 |
| Avil | 57.09 | 0.01 | 0.54 | 2.69 | 2.69 |
| Apba2 | 130.00 | 0.01 | 0.54 | 2.71 | 2.71 |
| Gulp1 | 324.00 | 0.04 | 0.64 | 2.76 | 2.76 |
| Mup18 | 4410.39 | 0.04 | 0.64 | 2.86 | 2.86 |
| Myom3 | 96.00 | 0.04 | 0.63 | 2.89 | 2.89 |
| Capn8 | 2013.00 | 2.77E-04 | 0.19 | 2.89 | 2.89 |
| Gm19014 | 42.15 | 0.04 | 0.63 | 3.00 | 3.00 |
| Tff3 | 1050.00 | 2.07E-04 | 0.18 | 3.21 | 3.21 |
| Gm43802 | 45.94 | 0.01 | 0.58 | 3.22 | 3.22 |
| Cxcl2 | 47.00 | 0.01 | 0.54 | 3.27 | 3.27 |
| Hspa1b | 12716.67 | 0.03 | 0.63 | 3.30 | 3.30 |
| Gm12406 | 33.00 | 0.01 | 0.54 | 3.33 | 3.33 |
| Dkk4 | 107.00 | 2.73E-04 | 0.19 | 3.36 | 3.36 |
| Akap6 | 314.00 | 4.37E-04 | 0.21 | 3.48 | 3.48 |
| Gm49361 | 46.36 | 0.03 | 0.63 | 3.70 | 3.70 |
| Gm3953 | 112.00 | 6.90E-04 | 0.25 | 3.75 | 3.75 |
| Hspa1a | 972.21 | 0.02 | 0.59 | 3.91 | 3.91 |
| Gm7967 | 129.00 | 3.47E-03 | 0.45 | 4.12 | 4.12 |
| 5330439K02Rik | 39.00 | 3.88E-03 | 0.46 | 4.17 | 4.17 |
| Gm37158 | 60.23 | 4.28E-04 | 0.21 | 4.35 | 4.35 |
| Cxcl1 | 5470.98 | 0.04 | 0.64 | 4.85 | 4.85 |
| Gm6472 | 1976.74 | 1.55E-05 | 0.07 | 5.50 | 5.50 |
| Saa1 | 20786.37 | 1.72E-06 | 0.01 | 5.58 | 5.58 |
| Kcnk1 | 488.96 | 0.02 | 0.60 | 6.20 | 6.20 |
| Yy2 | 67.55 | 0.01 | 0.53 | 7.23 | 7.23 |
| Saa2 | 9234.35 | 5.56E-07 | 0.01 | 7.74 | 7.74 |
| Gm14438 | 57.34 | 3.60E-07 | 0.01 | 16.12 | 16.12 |

All genes differentially expressed in Alb-D2KO liver using Partek Flow statistical package; total counts is the number of reads overlapping a specific gene.

**Table S4**- Gene set enrichment analysis of differentially expressed genes in the ALB-D2KO mouse liver.

| Gene set | Description | Enrichment score | P-value |
|---|---|---|---|
| GO:0006631 | fatty acid metabolic process | 31.9 | 1.46E-14 |
| GO:0006629 | lipid metabolic process | 31.2 | 2.88E-14 |
| GO:0032787 | monocarboxylic acid metabolic process | 30.1 | 8.53E-14 |
| GO:0043436 | oxoacid metabolic process | 29.2 | 1.98E-13 |
| GO:0006082 | organic acid metabolic process | 29.2 | 2.04E-13 |
| GO:0019752 | carboxylic acid metabolic process | 28.6 | 3.79E-13 |
| GO:0044255 | cellular lipid metabolic process | 21.5 | 4.62E-10 |
| GO:0044281 | small molecule metabolic process | 18.2 | 1.29E-08 |
| GO:0016054 | organic acid catabolic process | 16 | 1.11E-07 |
| GO:0046395 | carboxylic acid catabolic process | 16 | 1.11E-07 |
| GO:0097164 | ammonium ion metabolic process | 16 | 1.12E-07 |
| GO:0072329 | monocarboxylic acid catabolic process | 14.6 | 4.67E-07 |
| GO:0055114 | oxidation-reduction process | 14.6 | 4.72E-07 |
| GO:0009062 | fatty acid catabolic process | 14.2 | 7.11E-07 |
| GO:0001676 | long-chain fatty acid metabolic process | 13.7 | 1.10E-06 |
| GO:0019395 | fatty acid oxidation | 13.5 | 1.37E-06 |
| GO:0006790 | sulfur compound metabolic process | 13.3 | 1.70E-06 |
| GO:0034440 | lipid oxidation | 13.3 | 1.73E-06 |
| GO:0006805 | xenobiotic metabolic process | 13 | 2.19E-06 |
| GO:0042178 | xenobiotic catabolic process | 12.9 | 2.47E-06 |
| GO:0044282 | small molecule catabolic process | 12.3 | 4.78E-06 |
| GO:0052689 | carboxylic ester hydrolase activity | 12 | 6.12E-06 |
| GO:0004364 | glutathione transferase activity | 11.9 | 6.68E-06 |
| GO:0008391 | arachidonic acid monooxygenase activity | 11.6 | 9.38E-06 |
| GO:0016491 | oxidoreductase activity | 11.4 | 1.17E-05 |
| GO:0062012 | regulation of small molecule metabolic process | 11.2 | 1.32E-05 |
| GO:0048037 | cofactor binding | 11.1 | 1.48E-05 |
| GO:0008392 | arachidonic acid epoxygenase activity | 11 | 1.71E-05 |
| GO:0016042 | lipid catabolic process | 11 | 1.73E-05 |
| GO:0006575 | cellular modified amino acid metabolic process | 10.8 | 2.11E-05 |
| GO:2000026 | regulation of multicellular organismal development | 10.3 | 3.30E-05 |
| GO:0006635 | fatty acid beta-oxidation | 10.2 | 3.66E-05 |
| GO:0006109 | regulation of carbohydrate metabolic process | 10.2 | 3.75E-05 |
| GO:0120161 | regulation of cold-induced thermogenesis | 9.9 | 5.13E-05 |
| GO:0019216 | regulation of lipid metabolic process | 9.8 | 5.46E-05 |
| GO:0030258 | lipid modification | 9.7 | 5.97E-05 |
| GO:0044242 | cellular lipid catabolic process | 9.5 | 7.60E-05 |
| GO:0033559 | unsaturated fatty acid metabolic process | 9.5 | 7.74E-05 |
| GO:0048519 | negative regulation of biological process | 9.2 | 9.70E-05 |

| | | | |
|---|---|---|---|
| GO:0033865 | nucleoside bisphosphate metabolic process | 9.2 | 1.05E-04 |
| GO:0033875 | ribonucleoside bisphosphate metabolic process | 9.2 | 1.05E-04 |
| GO:0034032 | purine nucleoside bisphosphate metabolic process | 9.2 | 1.05E-04 |
| GO:0006749 | glutathione metabolic process | 9.1 | 1.09E-04 |
| GO:0016790 | thiolester hydrolase activity | 9.1 | 1.15E-04 |
| GO:0009896 | positive regulation of catabolic process | 9 | 1.18E-04 |
| GO:0051186 | cofactor metabolic process | 9 | 1.22E-04 |
| GO:0006637 | acyl-CoA metabolic process | 9 | 1.24E-04 |
| GO:0035383 | thioester metabolic process | 9 | 1.24E-04 |
| GO:0006577 | amino-acid betaine metabolic process | 8.9 | 1.36E-04 |
| GO:0010033 | response to organic substance | 8.9 | 1.38E-04 |
| GO:0042221 | response to chemical | 8.8 | 1.58E-04 |
| GO:0031667 | response to nutrient levels | 8.6 | 1.81E-04 |
| GO:0046394 | carboxylic acid biosynthetic process | 8.6 | 1.83E-04 |
| GO:0016053 | organic acid biosynthetic process | 8.5 | 1.99E-04 |
| GO:0072529 | pyrimidine-containing compound catabolic process | 8.5 | 2.03E-04 |
| GO:0010675 | regulation of cellular carbohydrate metabolic process | 8.5 | 2.05E-04 |
| GO:0006357 | regulation of transcription by RNA polymerase II | 8.4 | 2.14E-04 |
| GO:0050793 | regulation of developmental process | 8.4 | 2.20E-04 |
| GO:0042133 | neurotransmitter metabolic process | 8.4 | 2.30E-04 |
| GO:0072330 | monocarboxylic acid biosynthetic process | 8.2 | 2.66E-04 |
| GO:0016765 | transferase activity, transferring alkyl or aryl (other than methyl) groups | 8.2 | 2.73E-04 |
| GO:0019222 | regulation of metabolic process | 8.1 | 3.10E-04 |
| GO:0048523 | negative regulation of cellular process | 8 | 3.30E-04 |
| GO:0015872 | dopamine transport | 8 | 3.30E-04 |
| GO:0018685 | alkane 1-monooxygenase activity | 8 | 3.30E-04 |
| GO:0042135 | neurotransmitter catabolic process | 8 | 3.51E-04 |
| GO:0047493 | ceramide cholinephosphotransferase activity | 8 | 3.51E-04 |
| GO:0002950 | ceramide phosphoethanolamine synthase activity | 8 | 3.51E-04 |
| GO:0004165 | dodecenoyl-CoA delta-isomerase activity | 8 | 3.51E-04 |
| GO:0032449 | CBM complex | 8 | 3.51E-04 |
| GO:0033188 | sphingomyelin synthase activity | 8 | 3.51E-04 |
| GO:0050662 | coenzyme binding | 7.9 | 3.53E-04 |
| GO:0004497 | monooxygenase activity | 7.9 | 3.75E-04 |
| GO:0017144 | drug metabolic process | 7.9 | 3.77E-04 |
| GO:0003712 | transcription coregulator activity | 7.8 | 3.91E-04 |
| GO:0140110 | transcription regulator activity | 7.8 | 4.00E-04 |
| GO:0009991 | response to extracellular stimulus | 7.8 | 4.07E-04 |
| GO:0120162 | positive regulation of cold-induced thermogenesis | 7.6 | 4.82E-04 |
| GO:0016323 | basolateral plasma membrane | 7.6 | 4.83E-04 |
| GO:0001505 | regulation of neurotransmitter levels | 7.6 | 5.15E-04 |
| GO:0060749 | mammary gland alveolus development | 7.5 | 5.28E-04 |

| | | | |
|---|---|---|---|
| GO:0019217 | regulation of fatty acid metabolic process | 7.5 | 5.53E-04 |
| GO:0044248 | cellular catabolic process | 7.5 | 5.56E-04 |
| GO:0009437 | carnitine metabolic process | 7.5 | 5.60E-04 |
| GO:0047617 | acyl-CoA hydrolase activity | 7.5 | 5.69E-04 |
| GO:0006732 | coenzyme metabolic process | 7.4 | 5.82E-04 |
| GO:0000981 | DNA-binding transcription factor activity, RNA polymerase II-specific | 7.4 | 5.86E-04 |
| GO:0001067 | regulatory region nucleic acid binding | 7.4 | 6.06E-04 |
| GO:0043651 | linoleic acid metabolic process | 7.2 | 7.27E-04 |
| GO:0050896 | response to stimulus | 7.2 | 7.64E-04 |
| GO:0019219 | regulation of nucleobase-containing compound metabolic process | 7.2 | 7.75E-04 |
| GO:0080090 | regulation of primary metabolic process | 7.2 | 7.75E-04 |
| GO:0009308 | amine metabolic process | 7.1 | 8.01E-04 |
| GO:0044212 | transcription regulatory region DNA binding | 7.1 | 8.18E-04 |
| GO:0099146 | intrinsic component of postsynaptic density membrane | 7.1 | 8.49E-04 |
| GO:0051969 | regulation of transmission of nerve impulse | 7 | 9.04E-04 |
| GO:0034663 | endoplasmic reticulum chaperone complex | 7 | 9.04E-04 |
| GO:0031331 | positive regulation of cellular catabolic process | 7 | 9.52E-04 |
| GO:0009894 | regulation of catabolic process | 7 | 9.53E-04 |
| GO:0016788 | hydrolase activity, acting on ester bonds | 7 | 9.57E-04 |
| GO:0016627 | oxidoreductase activity, acting on the CH-CH group of donors | 6.9 | 1.00E-03 |
| GO:0016289 | CoA hydrolase activity | 6.9 | 1.02E-03 |
| GO:0034364 | high-density lipoprotein particle | 6.9 | 1.02E-03 |
| GO:0006576 | cellular biogenic amine metabolic process | 6.8 | 1.06E-03 |
| GO:0006206 | pyrimidine nucleobase metabolic process | 6.8 | 1.08E-03 |
| GO:0050873 | brown fat cell differentiation | 6.8 | 1.10E-03 |
| GO:0055088 | lipid homeostasis | 6.8 | 1.14E-03 |
| GO:0006633 | fatty acid biosynthetic process | 6.8 | 1.16E-03 |
| GO:0031326 | regulation of cellular biosynthetic process | 6.8 | 1.17E-03 |
| GO:0061448 | connective tissue development | 6.7 | 1.18E-03 |
| GO:0051253 | negative regulation of RNA metabolic process | 6.7 | 1.20E-03 |
| GO:0010906 | regulation of glucose metabolic process | 6.7 | 1.25E-03 |
| GO:0044106 | cellular amine metabolic process | 6.7 | 1.29E-03 |
| GO:0018916 | nitrobenzene metabolic process | 6.6 | 1.33E-03 |
| GO:0019482 | beta-alanine metabolic process | 6.6 | 1.33E-03 |
| GO:0071614 | linoleic acid epoxygenase activity | 6.6 | 1.33E-03 |
| GO:0050051 | leukotriene-B4 20-monooxygenase activity | 6.6 | 1.33E-03 |
| GO:0050218 | propionate-CoA ligase activity | 6.6 | 1.33E-03 |
| GO:0090493 | catecholamine uptake | 6.6 | 1.33E-03 |
| GO:0090494 | dopamine uptake | 6.6 | 1.33E-03 |
| GO:0016878 | acid-thiol ligase activity | 6.6 | 1.33E-03 |
| GO:0007275 | multicellular organism development | 6.6 | 1.36E-03 |
| GO:0009409 | response to cold | 6.6 | 1.37E-03 |

| | | | |
|---|---|---|---|
| GO:0051937 | catecholamine transport | 6.6 | 1.37E-03 |
| GO:0033601 | positive regulation of mammary gland epithelial cell proliferation | 6.6 | 1.37E-03 |
| GO:0051239 | regulation of multicellular organismal process | 6.6 | 1.39E-03 |
| GO:1990837 | sequence-specific double-stranded DNA binding | 6.6 | 1.41E-03 |
| GO:0006793 | phosphorus metabolic process | 6.5 | 1.43E-03 |
| GO:0008610 | lipid biosynthetic process | 6.5 | 1.45E-03 |
| GO:0045732 | positive regulation of protein catabolic process | 6.5 | 1.45E-03 |
| GO:0005777 | peroxisome | 6.5 | 1.45E-03 |
| GO:0010565 | regulation of cellular ketone metabolic process | 6.5 | 1.53E-03 |
| GO:0051252 | regulation of RNA metabolic process | 6.5 | 1.55E-03 |
| GO:0015711 | organic anion transport | 6.5 | 1.55E-03 |
| GO:0008408 | 3'-5' exonuclease activity | 6.4 | 1.63E-03 |
| GO:1902679 | negative regulation of RNA biosynthetic process | 6.4 | 1.66E-03 |
| GO:0031669 | cellular response to nutrient levels | 6.4 | 1.67E-03 |
| GO:0072341 | modified amino acid binding | 6.4 | 1.68E-03 |
| GO:0006690 | icosanoid metabolic process | 6.4 | 1.68E-03 |
| GO:1901567 | fatty acid derivative binding | 6.4 | 1.70E-03 |
| GO:0009893 | positive regulation of metabolic process | 6.4 | 1.71E-03 |
| GO:0019373 | epoxygenase P450 pathway | 6.4 | 1.71E-03 |
| GO:0006796 | phosphate-containing compound metabolic process | 6.4 | 1.73E-03 |
| GO:0098936 | intrinsic component of postsynaptic membrane | 6.3 | 1.77E-03 |
| GO:0009889 | regulation of biosynthetic process | 6.3 | 1.86E-03 |
| GO:0000976 | transcription regulatory region sequence-specific DNA binding | 6.3 | 1.87E-03 |
| GO:0007589 | body fluid secretion | 6.3 | 1.88E-03 |
| GO:1901568 | fatty acid derivative metabolic process | 6.2 | 1.95E-03 |
| GO:0004806 | triglyceride lipase activity | 6.2 | 2.00E-03 |
| GO:2000112 | regulation of cellular macromolecule biosynthetic process | 6.2 | 2.01E-03 |
| GO:0043177 | organic acid binding | 6.2 | 2.04E-03 |
| GO:0065008 | regulation of biological quality | 6.2 | 2.12E-03 |
| GO:0043621 | protein self-association | 6.1 | 2.17E-03 |
| GO:0008395 | steroid hydroxylase activity | 6.1 | 2.17E-03 |
| GO:2001141 | regulation of RNA biosynthetic process | 6.1 | 2.17E-03 |
| GO:1905953 | negative regulation of lipid localization | 6.1 | 2.25E-03 |
| GO:0009755 | hormone-mediated signaling pathway | 6.1 | 2.31E-03 |
| GO:0016713 | oxidoreductase activity, acting on paired donors, with incorporation or reduction of molecular oxygen, reduced iron-sulfur protein as one donor, and incorporation of one atom of oxygen | 6.1 | 2.33E-03 |
| GO:0016812 | hydrolase activity, acting on carbon-nitrogen (but not peptide) bonds, in cyclic amides | 6.1 | 2.33E-03 |
| GO:1903507 | negative regulation of nucleic acid-templated transcription | 6.1 | 2.33E-03 |
| GO:0045934 | negative regulation of nucleobase-containing | 6.1 | 2.34E-03 |

| | | | |
|---|---|---|---|
| | compound metabolic process | | |
| GO:0003824 | catalytic activity | 6.1 | 2.35E-03 |
| GO:0010629 | negative regulation of gene expression | 6 | 2.37E-03 |
| GO:2000113 | negative regulation of cellular macromolecule biosynthetic process | 6 | 2.38E-03 |
| GO:0009749 | response to glucose | 6 | 2.38E-03 |
| GO:0010556 | regulation of macromolecule biosynthetic process | 6 | 2.40E-03 |
| GO:0009892 | negative regulation of metabolic process | 6 | 2.49E-03 |
| GO:0006766 | vitamin metabolic process | 5.9 | 2.63E-03 |
| GO:0005501 | retinoid binding | 5.9 | 2.63E-03 |
| GO:0003700 | DNA-binding transcription factor activity | 5.9 | 2.65E-03 |
| GO:1903506 | regulation of nucleic acid-templated transcription | 5.9 | 2.65E-03 |
| GO:0009056 | catabolic process | 5.9 | 2.66E-03 |
| GO:0000038 | very long-chain fatty acid metabolic process | 5.9 | 2.72E-03 |
| GO:0042579 | microbody | 5.9 | 2.73E-03 |
| GO:0031327 | negative regulation of cellular biosynthetic process | 5.9 | 2.76E-03 |
| GO:0010558 | negative regulation of macromolecule biosynthetic process | 5.9 | 2.78E-03 |
| GO:0009890 | negative regulation of biosynthetic process | 5.9 | 2.84E-03 |
| GO:0015850 | organic hydroxy compound transport | 5.9 | 2.88E-03 |
| GO:0009605 | response to external stimulus | 5.8 | 2.90E-03 |
| GO:0006355 | regulation of transcription, DNA-templated | 5.8 | 2.93E-03 |
| GO:0045892 | negative regulation of transcription, DNA-templated | 5.8 | 2.99E-03 |
| GO:0031406 | carboxylic acid binding | 5.8 | 2.99E-03 |
| GO:0071310 | cellular response to organic substance | 5.8 | 3.01E-03 |
| GO:0016705 | oxidoreductase activity, acting on paired donors, with incorporation or reduction of molecular oxygen | 5.8 | 3.07E-03 |
| GO:0030976 | thiamine pyrophosphate binding | 5.8 | 3.15E-03 |
| GO:1901681 | sulfur compound binding | 5.7 | 3.20E-03 |
| GO:0019904 | protein domain specific binding | 5.7 | 3.29E-03 |
| GO:0009746 | response to hexose | 5.7 | 3.38E-03 |
| GO:0070887 | cellular response to chemical stimulus | 5.7 | 3.39E-03 |
| GO:0000122 | negative regulation of transcription by RNA polymerase II | 5.7 | 3.41E-03 |
| GO:0016405 | CoA-ligase activity | 5.7 | 3.41E-03 |
| GO:0042537 | benzene-containing compound metabolic process | 5.7 | 3.41E-03 |
| GO:0051960 | regulation of nervous system development | 5.7 | 3.43E-03 |
| GO:0001012 | RNA polymerase II regulatory region DNA binding | 5.6 | 3.59E-03 |
| GO:0046703 | natural killer cell lectin-like receptor binding | 5.6 | 3.67E-03 |
| GO:0006213 | pyrimidine nucleoside metabolic process | 5.6 | 3.67E-03 |
| GO:0038180 | nerve growth factor signaling pathway | 5.6 | 3.67E-03 |
| GO:0090207 | regulation of triglyceride metabolic process | 5.6 | 3.69E-03 |
| GO:0051172 | negative regulation of nitrogen compound | 5.6 | 3.74E-03 |

| | | | |
|---|---|---|---|
| | metabolic process | | |
| GO:0034284 | response to monosaccharide | 5.6 | 3.78E-03 |
| GO:0042176 | regulation of protein catabolic process | 5.6 | 3.87E-03 |
| GO:0045109 | intermediate filament organization | 5.5 | 3.93E-03 |
| GO:0046337 | phosphatidylethanolamine metabolic process | 5.5 | 3.93E-03 |
| GO:0072527 | pyrimidine-containing compound metabolic process | 5.5 | 4.07E-03 |
| GO:0070330 | aromatase activity | 5.5 | 4.14E-03 |
| GO:0006767 | water-soluble vitamin metabolic process | 5.5 | 4.14E-03 |
| GO:0000977 | RNA polymerase II regulatory region sequence-specific DNA binding | 5.5 | 4.15E-03 |
| GO:0045935 | positive regulation of nucleobase-containing compound metabolic process | 5.5 | 4.27E-03 |
| GO:0031323 | regulation of cellular metabolic process | 5.5 | 4.29E-03 |
| GO:0099061 | integral component of postsynaptic density membrane | 5.4 | 4.31E-03 |
| GO:0045723 | positive regulation of fatty acid biosynthetic process | 5.4 | 4.34E-03 |
| GO:0000062 | fatty-acyl-CoA binding | 5.4 | 4.34E-03 |
| GO:1903747 | regulation of establishment of protein localization to mitochondrion | 5.4 | 4.34E-03 |
| GO:0043565 | sequence-specific DNA binding | 5.4 | 4.58E-03 |
| GO:0050848 | regulation of calcium-mediated signaling | 5.4 | 4.70E-03 |
| GO:0032501 | multicellular organismal process | 5.4 | 4.72E-03 |
| GO:0042594 | response to starvation | 5.3 | 4.75E-03 |
| GO:0031328 | positive regulation of cellular biosynthetic process | 5.3 | 4.76E-03 |
| GO:0016740 | transferase activity | 5.3 | 4.80E-03 |
| GO:0008406 | gonad development | 5.3 | 4.81E-03 |
| GO:0015939 | pantothenate metabolic process | 5.3 | 4.98E-03 |
| GO:0102209 | trans-permethrin hydrolase activity | 5.3 | 4.98E-03 |
| GO:0017159 | pantetheine hydrolase activity | 5.3 | 4.98E-03 |
| GO:1903381 | regulation of endoplasmic reticulum stress-induced neuron intrinsic apoptotic signaling pathway | 5.3 | 4.98E-03 |
| GO:1903382 | negative regulation of endoplasmic reticulum stress-induced neuron intrinsic apoptotic signaling pathway | 5.3 | 4.98E-03 |
| GO:0099540 | trans-synaptic signaling by neuropeptide | 5.3 | 4.98E-03 |
| GO:0099551 | trans-synaptic signaling by neuropeptide, modulating synaptic transmission | 5.3 | 4.98E-03 |
| GO:0002096 | polkadots | 5.3 | 4.98E-03 |
| GO:1905371 | ceramide phosphoethanolamine metabolic process | 5.3 | 4.98E-03 |
| GO:1905373 | ceramide phosphoethanolamine biosynthetic process | 5.3 | 4.98E-03 |
| GO:0004777 | succinate-semialdehyde dehydrogenase (NAD+) activity | 5.3 | 4.98E-03 |
| GO:0006574 | valine catabolic process | 5.3 | 4.98E-03 |

| | | | |
|---|---|---|---|
| GO:0032444 | activin responsive factor complex | 5.3 | 4.98E-03 |
| GO:0032788 | saturated monocarboxylic acid metabolic process | 5.3 | 4.98E-03 |
| GO:0032789 | unsaturated monocarboxylic acid metabolic process | 5.3 | 4.98E-03 |
| GO:0033189 | response to vitamin A | 5.3 | 4.98E-03 |
| GO:0008670 | 2,4-dienoyl-CoA reductase (NADPH) activity | 5.3 | 4.98E-03 |
| GO:0009013 | succinate-semialdehyde dehydrogenase [NAD(P)+] activity | 5.3 | 4.98E-03 |
| GO:0009631 | cold acclimation | 5.3 | 4.98E-03 |
| GO:0086038 | calcium:sodium antiporter activity involved in regulation of cardiac muscle cell membrane potential | 5.3 | 4.98E-03 |
| GO:1990444 | F-box domain binding | 5.3 | 4.98E-03 |
| GO:0014701 | junctional sarcoplasmic reticulum membrane | 5.3 | 4.98E-03 |
| GO:0090650 | cellular response to oxygen-glucose deprivation | 5.3 | 4.98E-03 |
| GO:0090649 | response to oxygen-glucose deprivation | 5.3 | 4.98E-03 |
| GO:0042304 | regulation of fatty acid biosynthetic process | 5.3 | 0.01 |
| GO:0099055 | integral component of postsynaptic membrane | 5.3 | 0.01 |
| GO:0043295 | glutathione binding | 5.2 | 0.01 |
| GO:1900750 | oligopeptide binding | 5.2 | 0.01 |
| GO:0006595 | polyamine metabolic process | 5.2 | 0.01 |
| GO:0007422 | peripheral nervous system development | 5.2 | 0.01 |
| GO:0042826 | histone deacetylase binding | 5.2 | 0.01 |
| GO:0016324 | apical plasma membrane | 5.2 | 0.01 |
| GO:0014068 | positive regulation of phosphatidylinositol 3-kinase signaling | 5.2 | 0.01 |
| GO:0045346 | regulation of MHC class II biosynthetic process | 5.2 | 0.01 |
| GO:0046459 | short-chain fatty acid metabolic process | 5.2 | 0.01 |
| GO:0031000 | response to caffeine | 5.2 | 0.01 |
| GO:0036270 | response to diuretic | 5.2 | 0.01 |
| GO:0000209 | protein polyubiquitination | 5.2 | 0.01 |
| GO:0043549 | regulation of kinase activity | 5.2 | 0.01 |
| GO:0009891 | positive regulation of biosynthetic process | 5.2 | 0.01 |
| GO:0016712 | oxidoreductase activity, acting on paired donors, with incorporation or reduction of molecular oxygen, reduced flavin or flavoprotein as one donor, and incorporation of one atom of oxygen | 5.2 | 0.01 |
| GO:0045944 | positive regulation of transcription by RNA polymerase II | 5.1 | 0.01 |
| GO:1901897 | regulation of relaxation of cardiac muscle | 5.1 | 0.01 |
| GO:0098908 | regulation of neuronal action potential | 5.1 | 0.01 |
| GO:0030870 | Mre11 complex | 5.1 | 0.01 |
| GO:0006686 | sphingomyelin biosynthetic process | 5.1 | 0.01 |
| GO:0007586 | digestion | 5.1 | 0.01 |
| GO:0070884 | regulation of calcineurin-NFAT signaling cascade | 5.1 | 0.01 |
| GO:0035265 | organ growth | 5.1 | 0.01 |

| | | | |
|---|---|---|---|
| GO:0009743 | response to carbohydrate | 5.1 | 0.01 |
| GO:0045923 | positive regulation of fatty acid metabolic process | 5.1 | 0.01 |
| GO:0005506 | iron ion binding | 5.1 | 0.01 |
| GO:0071496 | cellular response to external stimulus | 5.1 | 0.01 |
| GO:0097159 | organic cyclic compound binding | 5 | 0.01 |
| GO:0060255 | regulation of macromolecule metabolic process | 5 | 0.01 |
| GO:0048871 | multicellular organismal homeostasis | 5 | 0.01 |
| GO:0062013 | positive regulation of small molecule metabolic process | 5 | 0.01 |
| GO:0019840 | isoprenoid binding | 5 | 0.01 |
| GO:0007595 | lactation | 5 | 0.01 |
| GO:0015844 | monoamine transport | 5 | 0.01 |
| GO:0016780 | phosphotransferase activity, for other substituted phosphate groups | 5 | 0.01 |
| GO:0051338 | regulation of transferase activity | 5 | 0.01 |
| GO:0031668 | cellular response to extracellular stimulus | 5 | 0.01 |
| GO:0016747 | transferase activity, transferring acyl groups other than amino-acyl groups | 4.9 | 0.01 |
| GO:0106056 | regulation of calcineurin-mediated signaling | 4.9 | 0.01 |
| GO:0010468 | regulation of gene expression | 4.9 | 0.01 |
| GO:0001228 | DNA-binding transcription activator activity, RNA polymerase II-specific | 4.9 | 0.01 |
| GO:0033993 | response to lipid | 4.9 | 0.01 |
| GO:0098948 | intrinsic component of postsynaptic specialization membrane | 4.9 | 0.01 |
| GO:0004771 | sterol esterase activity | 4.9 | 0.01 |
| GO:0006596 | polyamine biosynthetic process | 4.9 | 0.01 |
| GO:0008584 | male gonad development | 4.9 | 0.01 |
| GO:0009888 | tissue development | 4.8 | 0.01 |
| GO:0010605 | negative regulation of macromolecule metabolic process | 4.8 | 0.01 |
| GO:0060487 | lung epithelial cell differentiation | 4.8 | 0.01 |
| GO:0001216 | DNA-binding transcription activator activity | 4.8 | 0.01 |
| GO:0045665 | negative regulation of neuron differentiation | 4.8 | 0.01 |
| GO:0097009 | energy homeostasis | 4.8 | 0.01 |
| GO:0034358 | plasma lipoprotein particle | 4.8 | 0.01 |
| GO:0060612 | adipose tissue development | 4.8 | 0.01 |
| GO:1990777 | lipoprotein particle | 4.8 | 0.01 |
| GO:0033293 | monocarboxylic acid binding | 4.8 | 0.01 |
| GO:0006820 | anion transport | 4.7 | 0.01 |
| GO:0046324 | regulation of glucose import | 4.7 | 0.01 |
| GO:0036041 | long-chain fatty acid binding | 4.7 | 0.01 |
| GO:0016810 | hydrolase activity, acting on carbon-nitrogen (but not peptide) bonds | 4.7 | 0.01 |
| GO:0043168 | anion binding | 4.7 | 0.01 |
| GO:0031324 | negative regulation of cellular metabolic process | 4.7 | 0.01 |

| | | | |
|---|---|---|---|
| GO:0010817 | regulation of hormone levels | 4.7 | 0.01 |
| GO:0050660 | flavin adenine dinucleotide binding | 4.7 | 0.01 |
| GO:0010604 | positive regulation of macromolecule metabolic process | 4.7 | 0.01 |
| GO:0042738 | exogenous drug catabolic process | 4.7 | 0.01 |
| GO:1901363 | heterocyclic compound binding | 4.6 | 0.01 |
| GO:0001954 | positive regulation of cell-matrix adhesion | 4.6 | 0.01 |
| GO:0006721 | terpenoid metabolic process | 4.6 | 0.01 |
| GO:0019852 | L-ascorbic acid metabolic process | 4.6 | 0.01 |
| GO:0047760 | butyrate-CoA ligase activity | 4.6 | 0.01 |
| GO:0002667 | regulation of T cell anergy | 4.6 | 0.01 |
| GO:0002911 | regulation of lymphocyte anergy | 4.6 | 0.01 |
| GO:0006208 | pyrimidine nucleobase catabolic process | 4.6 | 0.01 |
| GO:0033211 | adiponectin-activated signaling pathway | 4.6 | 0.01 |
| GO:0051240 | positive regulation of multicellular organismal process | 4.6 | 0.01 |
| GO:0060479 | lung cell differentiation | 4.6 | 0.01 |
| GO:0016877 | ligase activity, forming carbon-sulfur bonds | 4.6 | 0.01 |
| GO:0051254 | positive regulation of RNA metabolic process | 4.6 | 0.01 |
| GO:0016746 | transferase activity, transferring acyl groups | 4.6 | 0.01 |
| GO:0017128 | phospholipid scramblase activity | 4.6 | 0.01 |
| GO:1901722 | regulation of cell proliferation involved in kidney development | 4.6 | 0.01 |
| GO:0004535 | poly(A)-specific ribonuclease activity | 4.6 | 0.01 |
| GO:0006684 | sphingomyelin metabolic process | 4.6 | 0.01 |
| GO:0009636 | response to toxic substance | 4.5 | 0.01 |
| GO:0009267 | cellular response to starvation | 4.5 | 0.01 |
| GO:0015696 | ammonium transport | 4.5 | 0.01 |
| GO:0016709 | oxidoreductase activity, acting on paired donors, with incorporation or reduction of molecular oxygen, NAD(P)H as one donor, and incorporation of one atom of oxygen | 4.5 | 0.01 |
| GO:0009404 | toxin metabolic process | 4.5 | 0.01 |
| GO:0051173 | positive regulation of nitrogen compound metabolic process | 4.5 | 0.01 |
| GO:0030165 | PDZ domain binding | 4.5 | 0.01 |
| GO:1901605 | alpha-amino acid metabolic process | 4.5 | 0.01 |
| GO:0099240 | intrinsic component of synaptic membrane | 4.5 | 0.01 |
| GO:0016614 | oxidoreductase activity, acting on CH-OH group of donors | 4.5 | 0.01 |
| GO:0003714 | transcription corepressor activity | 4.4 | 0.01 |
| GO:0046914 | transition metal ion binding | 4.4 | 0.01 |
| GO:0031329 | regulation of cellular catabolic process | 4.4 | 0.01 |
| GO:0010557 | positive regulation of macromolecule biosynthetic process | 4.4 | 0.01 |
| GO:1903846 | positive regulation of cellular response to transforming growth factor beta stimulus | 4.4 | 0.01 |

| | | | |
|---|---|---|---|
| GO:0030511 | positive regulation of transforming growth factor beta receptor signaling pathway | 4.4 | 0.01 |
| GO:0003690 | double-stranded DNA binding | 4.4 | 0.01 |
| GO:0045833 | negative regulation of lipid metabolic process | 4.4 | 0.01 |
| GO:0009063 | cellular amino acid catabolic process | 4.4 | 0.01 |
| GO:0000978 | RNA polymerase II cis-regulatory region sequence-specific DNA binding | 4.4 | 0.01 |
| GO:0030097 | hemopoiesis | 4.4 | 0.01 |
| GO:0006869 | lipid transport | 4.4 | 0.01 |
| GO:0031519 | PcG protein complex | 4.4 | 0.01 |
| GO:1904029 | regulation of cyclin-dependent protein kinase activity | 4.3 | 0.01 |
| GO:0033613 | activating transcription factor binding | 4.3 | 0.01 |
| GO:0035326 | cis-regulatory region binding | 4.3 | 0.01 |
| GO:0045893 | positive regulation of transcription, DNA-templated | 4.3 | 0.01 |
| GO:0062014 | negative regulation of small molecule metabolic process | 4.3 | 0.01 |
| GO:0032994 | protein-lipid complex | 4.3 | 0.01 |
| GO:0042573 | retinoic acid metabolic process | 4.3 | 0.01 |
| GO:0045922 | negative regulation of fatty acid metabolic process | 4.3 | 0.01 |
| GO:0010866 | regulation of triglyceride biosynthetic process | 4.3 | 0.01 |
| GO:1901700 | response to oxygen-containing compound | 4.3 | 0.01 |
| GO:0051171 | regulation of nitrogen compound metabolic process | 4.3 | 0.01 |
| GO:0042401 | cellular biogenic amine biosynthetic process | 4.3 | 0.01 |
| GO:0043517 | positive regulation of DNA damage response, signal transduction by p53 class mediator | 4.3 | 0.01 |
| GO:0009081 | branched-chain amino acid metabolic process | 4.3 | 0.01 |
| GO:0009083 | branched-chain amino acid catabolic process | 4.3 | 0.01 |
| GO:0009309 | amine biosynthetic process | 4.3 | 0.01 |
| GO:0005887 | integral component of plasma membrane | 4.3 | 0.01 |
| GO:0009064 | glutamine family amino acid metabolic process | 4.3 | 0.01 |
| GO:1903508 | positive regulation of nucleic acid-templated transcription | 4.3 | 0.01 |
| GO:0015909 | long-chain fatty acid transport | 4.3 | 0.01 |
| GO:0016401 | palmitoyl-CoA oxidase activity | 4.3 | 0.01 |
| GO:0042713 | sperm ejaculation | 4.3 | 0.01 |
| GO:0043084 | penile erection | 4.3 | 0.01 |
| GO:0044346 | fibroblast apoptotic process | 4.3 | 0.01 |
| GO:0018992 | germ-line sex determination | 4.3 | 0.01 |
| GO:0019254 | carnitine metabolic process, CoA-linked | 4.3 | 0.01 |
| GO:0070458 | cellular detoxification of nitrogen compound | 4.3 | 0.01 |
| GO:0019857 | 5-methylcytosine metabolic process | 4.3 | 0.01 |
| GO:0019859 | thymine metabolic process | 4.3 | 0.01 |
| GO:0072110 | glomerular mesangial cell proliferation | 4.3 | 0.01 |
| GO:0046881 | positive regulation of follicle-stimulating hormone | 4.3 | 0.01 |

| | | | |
|---|---|---|---|
| | secretion | | |
| GO:0072576 | liver morphogenesis | 4.3 | 0.01 |
| GO:0097180 | serine protease inhibitor complex | 4.3 | 0.01 |
| GO:0048014 | Tie signaling pathway | 4.3 | 0.01 |
| GO:0047961 | glycine N-acyltransferase activity | 4.3 | 0.01 |
| GO:0048688 | negative regulation of sprouting of injured axon | 4.3 | 0.01 |
| GO:0050196 | [phosphorylase] phosphatase activity | 4.3 | 0.01 |
| GO:0099538 | synaptic signaling via neuropeptide | 4.3 | 0.01 |
| GO:1904048 | regulation of spontaneous neurotransmitter secretion | 4.3 | 0.01 |
| GO:1904071 | presynaptic active zone assembly | 4.3 | 0.01 |
| GO:0051410 | detoxification of nitrogen compound | 4.3 | 0.01 |
| GO:1904880 | response to hydrogen sulfide | 4.3 | 0.01 |
| GO:1904881 | cellular response to hydrogen sulfide | 4.3 | 0.01 |
| GO:0002581 | negative regulation of antigen processing and presentation of peptide or polysaccharide antigen via MHC class II | 4.3 | 0.01 |
| GO:0003987 | acetate-CoA ligase activity | 4.3 | 0.01 |
| GO:0004145 | diamine N-acetyltransferase activity | 4.3 | 0.01 |
| GO:0006210 | thymine catabolic process | 4.3 | 0.01 |
| GO:0006211 | 5-methylcytosine catabolic process | 4.3 | 0.01 |
| GO:0006573 | valine metabolic process | 4.3 | 0.01 |
| GO:0006578 | amino-acid betaine biosynthetic process | 4.3 | 0.01 |
| GO:0032278 | positive regulation of gonadotropin secretion | 4.3 | 0.01 |
| GO:0036100 | leukotriene catabolic process | 4.3 | 0.01 |
| GO:0036101 | leukotriene B4 catabolic process | 4.3 | 0.01 |
| GO:0036102 | leukotriene B4 metabolic process | 4.3 | 0.01 |
| GO:0031325 | positive regulation of cellular metabolic process | 4.2 | 0.01 |
| GO:1902680 | positive regulation of RNA biosynthetic process | 4.2 | 0.01 |
| GO:0003677 | DNA binding | 4.2 | 0.01 |
| GO:0042587 | glycogen granule | 4.2 | 0.02 |
| GO:0042758 | long-chain fatty acid catabolic process | 4.2 | 0.02 |
| GO:0070652 | HAUS complex | 4.2 | 0.02 |
| GO:0071415 | cellular response to purine-containing compound | 4.2 | 0.02 |
| GO:2000318 | positive regulation of T-helper 17 type immune response | 4.2 | 0.02 |
| GO:0072216 | positive regulation of metanephros development | 4.2 | 0.02 |
| GO:1901724 | positive regulation of cell proliferation involved in kidney development | 4.2 | 0.02 |
| GO:1903265 | positive regulation of tumor necrosis factor-mediated signaling pathway | 4.2 | 0.02 |
| GO:1903748 | negative regulation of establishment of protein localization to mitochondrion | 4.2 | 0.02 |
| GO:0005283 | amino acid:sodium symporter activity | 4.2 | 0.02 |
| GO:0008152 | metabolic process | 4.2 | 0.02 |
| GO:0048513 | animal organ development | 4.2 | 0.02 |

| | | | |
|---|---|---|---|
| GO:0071383 | cellular response to steroid hormone stimulus | 4.2 | 0.02 |
| GO:0004527 | exonuclease activity | 4.2 | 0.02 |
| GO:0008202 | steroid metabolic process | 4.2 | 0.02 |
| GO:0044237 | cellular metabolic process | 4.1 | 0.02 |
| GO:0000987 | cis-regulatory region sequence-specific DNA binding | 4.1 | 0.02 |
| GO:0016101 | diterpenoid metabolic process | 4.1 | 0.02 |
| GO:0016903 | oxidoreductase activity, acting on the aldehyde or oxo group of donors | 4.1 | 0.02 |
| GO:0043392 | negative regulation of DNA binding | 4.1 | 0.02 |
| GO:0019693 | ribose phosphate metabolic process | 4.1 | 0.02 |
| GO:0035634 | response to stilbenoid | 4.1 | 0.02 |
| GO:0042592 | homeostatic process | 4.1 | 0.02 |
| GO:0009259 | ribonucleotide metabolic process | 4.1 | 0.02 |
| GO:0005326 | neurotransmitter transmembrane transporter activity | 4.1 | 0.02 |
| GO:0015849 | organic acid transport | 4.1 | 0.02 |
| GO:0046942 | carboxylic acid transport | 4.1 | 0.02 |
| GO:0050768 | negative regulation of neurogenesis | 4.1 | 0.02 |
| GO:0030183 | B cell differentiation | 4.1 | 0.02 |
| GO:0010506 | regulation of autophagy | 4 | 0.02 |
| GO:0045859 | regulation of protein kinase activity | 4 | 0.02 |
| GO:0009150 | purine ribonucleotide metabolic process | 4 | 0.02 |
| GO:0016290 | palmitoyl-CoA hydrolase activity | 4 | 0.02 |
| GO:0017121 | plasma membrane phospholipid scrambling | 4 | 0.02 |
| GO:0044827 | modulation by host of viral genome replication | 4 | 0.02 |
| GO:0072520 | seminiferous tubule development | 4 | 0.02 |
| GO:0031091 | platelet alpha granule | 4 | 0.02 |
| GO:0032780 | negative regulation of ATPase activity | 4 | 0.02 |
| GO:0048008 | platelet-derived growth factor receptor signaling pathway | 4 | 0.02 |
| GO:0010628 | positive regulation of gene expression | 4 | 0.02 |
| GO:0055086 | nucleobase-containing small molecule metabolic process | 4 | 0.02 |
| GO:0019369 | arachidonic acid metabolic process | 4 | 0.02 |
| GO:0002526 | acute inflammatory response | 4 | 0.02 |
| GO:0051716 | cellular response to stimulus | 4 | 0.02 |
| GO:0019838 | growth factor binding | 4 | 0.02 |
| GO:0035690 | cellular response to drug | 4 | 0.02 |
| GO:1901342 | regulation of vasculature development | 4 | 0.02 |
| GO:0010827 | regulation of glucose transmembrane transport | 4 | 0.02 |
| GO:0018108 | peptidyl-tyrosine phosphorylation | 4 | 0.02 |
| GO:0042593 | glucose homeostasis | 4 | 0.02 |
| GO:0048511 | rhythmic process | 4 | 0.02 |
| GO:0036094 | small molecule binding | 3.9 | 0.02 |
| GO:0065007 | biological regulation | 3.9 | 0.02 |

| | | | |
|---|---|---|---|
| GO:0090184 | positive regulation of kidney development | 3.9 | 0.02 |
| GO:0042803 | protein homodimerization activity | 3.9 | 0.02 |
| GO:2001252 | positive regulation of chromosome organization | 3.9 | 0.02 |
| GO:0050767 | regulation of neurogenesis | 3.9 | 0.02 |
| GO:0005516 | calmodulin binding | 3.9 | 0.02 |
| GO:0033500 | carbohydrate homeostasis | 3.9 | 0.02 |
| GO:0016628 | oxidoreductase activity, acting on the CH-CH group of donors, NAD or NADP as acceptor | 3.9 | 0.02 |
| GO:0070412 | R-SMAD binding | 3.9 | 0.02 |
| GO:0045663 | positive regulation of myoblast differentiation | 3.9 | 0.02 |
| GO:0045736 | negative regulation of cyclin-dependent protein serine/threonine kinase activity | 3.9 | 0.02 |
| GO:0050849 | negative regulation of calcium-mediated signaling | 3.9 | 0.02 |
| GO:0060317 | cardiac epithelial to mesenchymal transition | 3.9 | 0.02 |
| GO:0044092 | negative regulation of molecular function | 3.9 | 0.02 |
| GO:1901616 | organic hydroxy compound catabolic process | 3.9 | 0.02 |
| GO:0090311 | regulation of protein deacetylation | 3.9 | 0.02 |
| GO:0009880 | embryonic pattern specification | 3.9 | 0.02 |
| GO:0019637 | organophosphate metabolic process | 3.9 | 0.02 |
| GO:1905269 | positive regulation of chromatin organization | 3.9 | 0.02 |
| GO:0099699 | integral component of synaptic membrane | 3.9 | 0.02 |
| GO:0010721 | negative regulation of cell development | 3.8 | 0.02 |
| GO:0042711 | maternal behavior | 3.8 | 0.02 |
| GO:0043568 | positive regulation of insulin-like growth factor receptor signaling pathway | 3.8 | 0.02 |
| GO:1900102 | negative regulation of endoplasmic reticulum unfolded protein response | 3.8 | 0.02 |
| GO:2000192 | negative regulation of fatty acid transport | 3.8 | 0.02 |
| GO:0000774 | adenyl-nucleotide exchange factor activity | 3.8 | 0.02 |
| GO:1905456 | regulation of lymphoid progenitor cell differentiation | 3.8 | 0.02 |
| GO:0005416 | amino acid:cation symporter activity | 3.8 | 0.02 |
| GO:0032094 | response to food | 3.8 | 0.02 |
| GO:0032927 | positive regulation of activin receptor signaling pathway | 3.8 | 0.02 |
| GO:0030098 | lymphocyte differentiation | 3.8 | 0.02 |
| GO:1905952 | regulation of lipid localization | 3.8 | 0.02 |
| GO:0014066 | regulation of phosphatidylinositol 3-kinase signaling | 3.8 | 0.02 |
| GO:0018107 | peptidyl-threonine phosphorylation | 3.8 | 0.02 |
| GO:0000079 | regulation of cyclin-dependent protein serine/threonine kinase activity | 3.8 | 0.02 |
| GO:2001057 | reactive nitrogen species metabolic process | 3.8 | 0.02 |
| GO:2001212 | regulation of vasculogenesis | 3.8 | 0.02 |
| GO:0046320 | regulation of fatty acid oxidation | 3.8 | 0.02 |
| GO:0045912 | negative regulation of carbohydrate metabolic process | 3.8 | 0.02 |

| | | | |
|---|---|---|---|
| GO:0031226 | intrinsic component of plasma membrane | 3.8 | 0.02 |
| GO:0016538 | cyclin-dependent protein serine/threonine kinase regulator activity | 3.8 | 0.02 |
| GO:0010883 | regulation of lipid storage | 3.8 | 0.02 |
| GO:0090183 | regulation of kidney development | 3.8 | 0.02 |
| GO:1901798 | positive regulation of signal transduction by p53 class mediator | 3.7 | 0.02 |
| GO:0004683 | calmodulin-dependent protein kinase activity | 3.7 | 0.02 |
| GO:0018212 | peptidyl-tyrosine modification | 3.7 | 0.02 |
| GO:0009791 | post-embryonic development | 3.7 | 0.02 |
| GO:1903828 | negative regulation of cellular protein localization | 3.7 | 0.02 |
| GO:0042802 | identical protein binding | 3.7 | 0.02 |
| GO:0034754 | cellular hormone metabolic process | 3.7 | 0.02 |
| GO:1901606 | alpha-amino acid catabolic process | 3.7 | 0.02 |
| GO:0055072 | iron ion homeostasis | 3.7 | 0.02 |
| GO:0009266 | response to temperature stimulus | 3.7 | 0.03 |
| GO:0043167 | ion binding | 3.7 | 0.03 |
| GO:0045834 | positive regulation of lipid metabolic process | 3.7 | 0.03 |
| GO:0051289 | protein homotetramerization | 3.7 | 0.03 |
| GO:0030173 | integral component of Golgi membrane | 3.7 | 0.03 |
| GO:1901575 | organic substance catabolic process | 3.7 | 0.03 |
| GO:0051961 | negative regulation of nervous system development | 3.7 | 0.03 |
| GO:0051094 | positive regulation of developmental process | 3.6 | 0.03 |
| GO:0042398 | cellular modified amino acid biosynthetic process | 3.6 | 0.03 |
| GO:0071466 | cellular response to xenobiotic stimulus | 3.6 | 0.03 |
| GO:0002931 | response to ischemia | 3.6 | 0.03 |
| GO:0098900 | regulation of action potential | 3.6 | 0.03 |
| GO:0120163 | negative regulation of cold-induced thermogenesis | 3.6 | 0.03 |
| GO:0033674 | positive regulation of kinase activity | 3.6 | 0.03 |
| GO:0016035 | zeta DNA polymerase complex | 3.6 | 0.03 |
| GO:0042424 | catecholamine catabolic process | 3.6 | 0.03 |
| GO:0018377 | protein myristoylation | 3.6 | 0.03 |
| GO:0044316 | cone cell pedicle | 3.6 | 0.03 |
| GO:0019107 | myristoyltransferase activity | 3.6 | 0.03 |
| GO:0019614 | catechol-containing compound catabolic process | 3.6 | 0.03 |
| GO:0070579 | methylcytosine dioxygenase activity | 3.6 | 0.03 |
| GO:0045347 | negative regulation of MHC class II biosynthetic process | 3.6 | 0.03 |
| GO:0071074 | eukaryotic initiation factor eIF2 binding | 3.6 | 0.03 |
| GO:0071141 | SMAD protein complex | 3.6 | 0.03 |
| GO:0071503 | response to heparin | 3.6 | 0.03 |
| GO:0071504 | cellular response to heparin | 3.6 | 0.03 |
| GO:2000195 | negative regulation of female gonad development | 3.6 | 0.03 |
| GO:2000301 | negative regulation of synaptic vesicle exocytosis | 3.6 | 0.03 |

| | | | |
|---|---|---|---|
| GO:0071947 | protein deubiquitination involved in ubiquitin-dependent protein catabolic process | 3.6 | 0.03 |
| GO:2000741 | positive regulation of mesenchymal stem cell differentiation | 3.6 | 0.03 |
| GO:0097179 | protease inhibitor complex | 3.6 | 0.03 |
| GO:0048104 | establishment of body hair or bristle planar orientation | 3.6 | 0.03 |
| GO:0048105 | establishment of body hair planar orientation | 3.6 | 0.03 |
| GO:0000320 | re-entry into mitotic cell cycle | 3.6 | 0.03 |
| GO:0000480 | endonucleolytic cleavage in 5'-ETS of tricistronic rRNA transcript (SSU-rRNA, 5.8S rRNA, LSU-rRNA) | 3.6 | 0.03 |
| GO:1904090 | peptidase inhibitor complex | 3.6 | 0.03 |
| GO:0051790 | short-chain fatty acid biosynthetic process | 3.6 | 0.03 |
| GO:0002134 | UTP binding | 3.6 | 0.03 |
| GO:0002669 | positive regulation of T cell anergy | 3.6 | 0.03 |
| GO:0052629 | phosphatidylinositol-3,5-bisphosphate 3-phosphatase activity | 3.6 | 0.03 |
| GO:1905605 | positive regulation of maintenance of permeability of blood-brain barrier | 3.6 | 0.03 |
| GO:0002904 | positive regulation of B cell apoptotic process | 3.6 | 0.03 |
| GO:0002913 | positive regulation of lymphocyte anergy | 3.6 | 0.03 |
| GO:0003158 | endothelium development | 3.6 | 0.03 |
| GO:0003433 | chondrocyte development involved in endochondral bone morphogenesis | 3.6 | 0.03 |
| GO:0003996 | acyl-CoA ligase activity | 3.6 | 0.03 |
| GO:0140213 | negative regulation of long-chain fatty acid import into cell | 3.6 | 0.03 |
| GO:0031860 | telomeric 3' overhang formation | 3.6 | 0.03 |
| GO:0006499 | N-terminal protein myristoylation | 3.6 | 0.03 |
| GO:0006659 | phosphatidylserine biosynthetic process | 3.6 | 0.03 |
| GO:0032229 | negative regulation of synaptic transmission, GABAergic | 3.6 | 0.03 |
| GO:0006848 | pyruvate transport | 3.6 | 0.03 |
| GO:0007171 | activation of transmembrane receptor protein tyrosine kinase activity | 3.6 | 0.03 |
| GO:0033695 | oxidoreductase activity, acting on CH or CH2 groups, quinone or similar compound as acceptor | 3.6 | 0.03 |
| GO:0008390 | testosterone 16-alpha-hydroxylase activity | 3.6 | 0.03 |
| GO:0008410 | CoA-transferase activity | 3.6 | 0.03 |
| GO:0060316 | positive regulation of ryanodine-sensitive calcium-release channel activity | 3.6 | 0.03 |
| GO:0034875 | caffeine oxidase activity | 3.6 | 0.03 |
| GO:0010748 | negative regulation of long-chain fatty acid import across plasma membrane | 3.6 | 0.03 |
| GO:0035441 | cell migration involved in vasculogenesis | 3.6 | 0.03 |
| GO:1990168 | protein K33-linked deubiquitination | 3.6 | 0.03 |
| GO:1990709 | presynaptic active zone organization | 3.6 | 0.03 |
| GO:1902115 | regulation of organelle assembly | 3.6 | 0.03 |

| | | | |
|---|---|---|---|
| GO:0043086 | negative regulation of catalytic activity | 3.6 | 0.03 |
| GO:0070402 | NADPH binding | 3.6 | 0.03 |
| GO:0070822 | Sin3-type complex | 3.6 | 0.03 |
| GO:0021799 | cerebral cortex radially oriented cell migration | 3.6 | 0.03 |
| GO:2000678 | negative regulation of transcription regulatory region DNA binding | 3.6 | 0.03 |
| GO:1903214 | regulation of protein targeting to mitochondrion | 3.6 | 0.03 |
| GO:0001504 | neurotransmitter uptake | 3.6 | 0.03 |
| GO:0030169 | low-density lipoprotein particle binding | 3.6 | 0.03 |
| GO:0030325 | adrenal gland development | 3.6 | 0.03 |
| GO:0033145 | positive regulation of intracellular steroid hormone receptor signaling pathway | 3.6 | 0.03 |
| GO:0010888 | negative regulation of lipid storage | 3.6 | 0.03 |
| GO:0070371 | ERK1 and ERK2 cascade | 3.6 | 0.03 |
| GO:1904030 | negative regulation of cyclin-dependent protein kinase activity | 3.6 | 0.03 |
| GO:0050794 | regulation of cellular process | 3.6 | 0.03 |
| GO:0016102 | diterpenoid biosynthetic process | 3.5 | 0.03 |
| GO:0044539 | long-chain fatty acid import into cell | 3.5 | 0.03 |
| GO:0046322 | negative regulation of fatty acid oxidation | 3.5 | 0.03 |
| GO:0046929 | negative regulation of neurotransmitter secretion | 3.5 | 0.03 |
| GO:1901077 | regulation of relaxation of muscle | 3.5 | 0.03 |
| GO:0000288 | nuclear-transcribed mRNA catabolic process, deadenylation-dependent decay | 3.5 | 0.03 |
| GO:0002138 | retinoic acid biosynthetic process | 3.5 | 0.03 |
| GO:0002664 | regulation of T cell tolerance induction | 3.5 | 0.03 |
| GO:0003995 | acyl-CoA dehydrogenase activity | 3.5 | 0.03 |
| GO:0140354 | lipid import into cell | 3.5 | 0.03 |
| GO:0006646 | phosphatidylethanolamine biosynthetic process | 3.5 | 0.03 |
| GO:0006809 | nitric oxide biosynthetic process | 3.5 | 0.03 |
| GO:0032525 | somite rostral/caudal axis specification | 3.5 | 0.03 |
| GO:0033539 | fatty acid beta-oxidation using acyl-CoA dehydrogenase | 3.5 | 0.03 |
| GO:0060746 | parental behavior | 3.5 | 0.03 |
| GO:1990226 | histone methyltransferase binding | 3.5 | 0.03 |
| GO:0099060 | integral component of postsynaptic specialization membrane | 3.5 | 0.03 |
| GO:0006140 | regulation of nucleotide metabolic process | 3.5 | 0.03 |
| GO:0015908 | fatty acid transport | 3.5 | 0.03 |
| GO:0046513 | ceramide biosynthetic process | 3.5 | 0.03 |
| GO:0001786 | phosphatidylserine binding | 3.5 | 0.03 |
| GO:0005504 | fatty acid binding | 3.5 | 0.03 |
| GO:0032007 | negative regulation of TOR signaling | 3.5 | 0.03 |
| GO:0046890 | regulation of lipid biosynthetic process | 3.5 | 0.03 |
| GO:0009719 | response to endogenous stimulus | 3.5 | 0.03 |
| GO:0016616 | oxidoreductase activity, acting on the CH-OH | 3.5 | 0.03 |

| | | | |
|---|---|---|---|
| | group of donors, NAD or NADP as acceptor | | |
| GO:0001217 | DNA-binding transcription repressor activity | 3.5 | 0.03 |
| GO:0001227 | DNA-binding transcription repressor activity, RNA polymerase II-specific | 3.5 | 0.03 |
| GO:0006520 | cellular amino acid metabolic process | 3.5 | 0.03 |
| GO:1903364 | positive regulation of cellular protein catabolic process | 3.5 | 0.03 |
| GO:1904018 | positive regulation of vasculature development | 3.5 | 0.03 |
| GO:0019932 | second-messenger-mediated signaling | 3.5 | 0.03 |
| GO:0016310 | phosphorylation | 3.5 | 0.03 |
| GO:0070988 | demethylation | 3.5 | 0.03 |
| GO:0031228 | intrinsic component of Golgi membrane | 3.5 | 0.03 |
| GO:0006468 | protein phosphorylation | 3.5 | 0.03 |
| GO:0071704 | organic substance metabolic process | 3.4 | 0.03 |
| GO:0071222 | cellular response to lipopolysaccharide | 3.4 | 0.03 |
| GO:1901861 | regulation of muscle tissue development | 3.4 | 0.03 |
| GO:0030154 | cell differentiation | 3.4 | 0.03 |
| GO:0051348 | negative regulation of transferase activity | 3.4 | 0.03 |
| GO:0048731 | system development | 3.4 | 0.03 |
| GO:0006836 | neurotransmitter transport | 3.4 | 0.03 |
| GO:0003713 | transcription coactivator activity | 3.4 | 0.03 |
| GO:0042056 | chemoattractant activity | 3.4 | 0.03 |
| GO:0000578 | embryonic axis specification | 3.4 | 0.03 |
| GO:0051354 | negative regulation of oxidoreductase activity | 3.4 | 0.03 |
| GO:0005496 | steroid binding | 3.4 | 0.03 |
| GO:0042737 | drug catabolic process | 3.4 | 0.03 |
| GO:0016874 | ligase activity | 3.4 | 0.03 |
| GO:0071333 | cellular response to glucose stimulus | 3.4 | 0.03 |
| GO:0098754 | detoxification | 3.4 | 0.03 |
| GO:0015645 | fatty acid ligase activity | 3.4 | 0.03 |
| GO:0042417 | dopamine metabolic process | 3.4 | 0.03 |
| GO:0070989 | oxidative demethylation | 3.4 | 0.03 |
| GO:0048745 | smooth muscle tissue development | 3.4 | 0.03 |
| GO:0002902 | regulation of B cell apoptotic process | 3.4 | 0.03 |
| GO:0004745 | retinol dehydrogenase activity | 3.4 | 0.03 |
| GO:0010743 | regulation of macrophage derived foam cell differentiation | 3.4 | 0.03 |
| GO:0015101 | organic cation transmembrane transporter activity | 3.4 | 0.03 |
| GO:0050789 | regulation of biological process | 3.4 | 0.03 |
| GO:0018210 | peptidyl-threonine modification | 3.4 | 0.03 |
| GO:0046983 | protein dimerization activity | 3.4 | 0.03 |
| GO:0017015 | regulation of transforming growth factor beta receptor signaling pathway | 3.4 | 0.03 |
| GO:0051087 | chaperone binding | 3.4 | 0.03 |
| GO:0007166 | cell surface receptor signaling pathway | 3.3 | 0.04 |
| GO:0050790 | regulation of catalytic activity | 3.3 | 0.04 |

| | | | |
|---|---|---|---|
| GO:0045595 | regulation of cell differentiation | 3.3 | 0.04 |
| GO:0050673 | epithelial cell proliferation | 3.3 | 0.04 |
| GO:0007584 | response to nutrient | 3.3 | 0.04 |
| GO:0048589 | developmental growth | 3.3 | 0.04 |
| GO:0001523 | retinoid metabolic process | 3.3 | 0.04 |
| GO:0010677 | negative regulation of cellular carbohydrate metabolic process | 3.3 | 0.04 |
| GO:0042493 | response to drug | 3.3 | 0.04 |
| GO:1903844 | regulation of cellular response to transforming growth factor beta stimulus | 3.3 | 0.04 |
| GO:0015651 | quaternary ammonium group transmembrane transporter activity | 3.3 | 0.04 |
| GO:0016233 | telomere capping | 3.3 | 0.04 |
| GO:0102991 | myristoyl-CoA hydrolase activity | 3.3 | 0.04 |
| GO:0043619 | regulation of transcription from RNA polymerase II promoter in response to oxidative stress | 3.3 | 0.04 |
| GO:0070411 | I-SMAD binding | 3.3 | 0.04 |
| GO:2000303 | regulation of ceramide biosynthetic process | 3.3 | 0.04 |
| GO:2000347 | positive regulation of hepatocyte proliferation | 3.3 | 0.04 |
| GO:2001214 | positive regulation of vasculogenesis | 3.3 | 0.04 |
| GO:0000289 | nuclear-transcribed mRNA poly(A) tail shortening | 3.3 | 0.04 |
| GO:0052695 | cellular glucuronidation | 3.3 | 0.04 |
| GO:0030322 | stabilization of membrane potential | 3.3 | 0.04 |
| GO:0005212 | structural constituent of eye lens | 3.3 | 0.04 |
| GO:0006534 | cysteine metabolic process | 3.3 | 0.04 |
| GO:0006658 | phosphatidylserine metabolic process | 3.3 | 0.04 |
| GO:0006968 | cellular defense response | 3.3 | 0.04 |
| GO:0032825 | positive regulation of natural killer cell differentiation | 3.3 | 0.04 |
| GO:0008401 | retinoic acid 4-hydroxylase activity | 3.3 | 0.04 |
| GO:0034755 | iron ion transmembrane transport | 3.3 | 0.04 |
| GO:0043620 | regulation of DNA-templated transcription in response to stress | 3.3 | 0.04 |
| GO:0030509 | BMP signaling pathway | 3.3 | 0.04 |
| GO:0006953 | acute-phase response | 3.3 | 0.04 |
| GO:0090208 | positive regulation of triglyceride metabolic process | 3.3 | 0.04 |
| GO:0090312 | positive regulation of protein deacetylation | 3.3 | 0.04 |
| GO:0010508 | positive regulation of autophagy | 3.3 | 0.04 |
| GO:0045104 | intermediate filament cytoskeleton organization | 3.3 | 0.04 |
| GO:0030518 | intracellular steroid hormone receptor signaling pathway | 3.3 | 0.04 |
| GO:0006163 | purine nucleotide metabolic process | 3.3 | 0.04 |
| GO:0045860 | positive regulation of protein kinase activity | 3.2 | 0.04 |
| GO:0007399 | nervous system development | 3.2 | 0.04 |
| GO:0008134 | transcription factor binding | 3.2 | 0.04 |
| GO:0048518 | positive regulation of biological process | 3.2 | 0.04 |

| | | | |
|---|---|---|---|
| GO:0042752 | regulation of circadian rhythm | 3.2 | 0.04 |
| GO:2000379 | positive regulation of reactive oxygen species metabolic process | 3.2 | 0.04 |
| GO:0009117 | nucleotide metabolic process | 3.2 | 0.04 |
| GO:0007623 | circadian rhythm | 3.2 | 0.04 |
| GO:0042572 | retinol metabolic process | 3.2 | 0.04 |
| GO:0045821 | positive regulation of glycolytic process | 3.2 | 0.04 |
| GO:0098688 | parallel fiber to Purkinje cell synapse | 3.2 | 0.04 |
| GO:0030057 | desmosome | 3.2 | 0.04 |
| GO:0005343 | organic acid:sodium symporter activity | 3.2 | 0.04 |
| GO:0031664 | regulation of lipopolysaccharide-mediated signaling pathway | 3.2 | 0.04 |
| GO:0071322 | cellular response to carbohydrate stimulus | 3.2 | 0.04 |
| GO:0050661 | NADP binding | 3.2 | 0.04 |
| GO:0045171 | intercellular bridge | 3.2 | 0.04 |
| GO:0015718 | monocarboxylic acid transport | 3.2 | 0.04 |
| GO:0071396 | cellular response to lipid | 3.2 | 0.04 |
| GO:0015727 | lactate transport | 3.1 | 0.04 |
| GO:0016554 | cytidine to uridine editing | 3.1 | 0.04 |
| GO:0019336 | phenol-containing compound catabolic process | 3.1 | 0.04 |
| GO:0070447 | positive regulation of oligodendrocyte progenitor proliferation | 3.1 | 0.04 |
| GO:0070813 | hydrogen sulfide metabolic process | 3.1 | 0.04 |
| GO:0071313 | cellular response to caffeine | 3.1 | 0.04 |
| GO:0106018 | phosphatidylinositol-3,5-bisphosphate phosphatase activity | 3.1 | 0.04 |
| GO:0046292 | formaldehyde metabolic process | 3.1 | 0.04 |
| GO:2000321 | positive regulation of T-helper 17 cell differentiation | 3.1 | 0.04 |
| GO:2000324 | positive regulation of glucocorticoid receptor signaling pathway | 3.1 | 0.04 |
| GO:0072012 | glomerulus vasculature development | 3.1 | 0.04 |
| GO:0021960 | anterior commissure morphogenesis | 3.1 | 0.04 |
| GO:0046668 | regulation of retinal cell programmed cell death | 3.1 | 0.04 |
| GO:1901523 | icosanoid catabolic process | 3.1 | 0.04 |
| GO:0098826 | endoplasmic reticulum tubular network membrane | 3.1 | 0.04 |
| GO:1902816 | regulation of protein localization to microtubule | 3.1 | 0.04 |
| GO:1902817 | negative regulation of protein localization to microtubule | 3.1 | 0.04 |
| GO:1902949 | positive regulation of tau-protein kinase activity | 3.1 | 0.04 |
| GO:1903215 | negative regulation of protein targeting to mitochondrion | 3.1 | 0.04 |
| GO:1903300 | negative regulation of hexokinase activity | 3.1 | 0.04 |
| GO:0001922 | B-1 B cell homeostasis | 3.1 | 0.04 |
| GO:0002580 | regulation of antigen processing and presentation of peptide or polysaccharide antigen via MHC class | 3.1 | 0.04 |

| | | | |
|---|---|---|---|
| | II | | |
| GO:1905451 | positive regulation of Fc-gamma receptor signaling pathway involved in phagocytosis | 3.1 | 0.04 |
| GO:0003847 | 1-alkyl-2-acetylglycerophosphocholine esterase activity | 3.1 | 0.04 |
| GO:0003997 | acyl-CoA oxidase activity | 3.1 | 0.04 |
| GO:0004157 | dihydropyrimidinase activity | 3.1 | 0.04 |
| GO:0004321 | fatty-acyl-CoA synthase activity | 3.1 | 0.04 |
| GO:0030124 | AP-4 adaptor complex | 3.1 | 0.04 |
| GO:0005173 | stem cell factor receptor binding | 3.1 | 0.04 |
| GO:0006104 | succinyl-CoA metabolic process | 3.1 | 0.04 |
| GO:0006498 | N-terminal protein lipidation | 3.1 | 0.04 |
| GO:0032557 | pyrimidine ribonucleotide binding | 3.1 | 0.04 |
| GO:0032792 | negative regulation of CREB transcription factor activity | 3.1 | 0.04 |
| GO:0033132 | negative regulation of glucokinase activity | 3.1 | 0.04 |
| GO:0008131 | primary amine oxidase activity | 3.1 | 0.04 |
| GO:0034184 | positive regulation of maintenance of mitotic sister chromatid cohesion | 3.1 | 0.04 |
| GO:0034433 | steroid esterification | 3.1 | 0.04 |
| GO:0034434 | sterol esterification | 3.1 | 0.04 |
| GO:0034435 | cholesterol esterification | 3.1 | 0.04 |
| GO:0008970 | phospholipase A1 activity | 3.1 | 0.04 |
| GO:0060406 | positive regulation of penile erection | 3.1 | 0.04 |
| GO:0010606 | positive regulation of cytoplasmic mRNA processing body assembly | 3.1 | 0.04 |
| GO:0010891 | negative regulation of sequestering of triglyceride | 3.1 | 0.04 |
| GO:0035523 | protein K29-linked deubiquitination | 3.1 | 0.04 |
| GO:0061156 | pulmonary artery morphogenesis | 3.1 | 0.04 |
| GO:0035873 | lactate transmembrane transport | 3.1 | 0.04 |
| GO:0035879 | plasma membrane lactate transport | 3.1 | 0.04 |
| GO:0061437 | renal system vasculature development | 3.1 | 0.04 |
| GO:0061440 | kidney vasculature development | 3.1 | 0.04 |
| GO:0061476 | response to anticoagulant | 3.1 | 0.04 |
| GO:0061642 | chemoattraction of axon | 3.1 | 0.04 |
| GO:0061734 | parkin-mediated stimulation of mitophagy in response to mitochondrial depolarization | 3.1 | 0.04 |
| GO:0150003 | regulation of spontaneous synaptic transmission | 3.1 | 0.04 |
| GO:0014889 | muscle atrophy | 3.1 | 0.04 |
| GO:0015220 | choline transmembrane transporter activity | 3.1 | 0.04 |
| GO:0045103 | intermediate filament-based process | 3.1 | 0.04 |
| GO:0071331 | cellular response to hexose stimulus | 3.1 | 0.04 |
| GO:0046326 | positive regulation of glucose import | 3.1 | 0.04 |
| GO:0004712 | protein serine/threonine/tyrosine kinase activity | 3.1 | 0.04 |
| GO:0008038 | neuron recognition | 3.1 | 0.04 |
| GO:0001932 | regulation of protein phosphorylation | 3.1 | 0.04 |

| | | | |
|---|---|---|---|
| GO:0071385 | cellular response to glucocorticoid stimulus | 3.1 | 0.04 |
| GO:0071813 | lipoprotein particle binding | 3.1 | 0.04 |
| GO:0071814 | protein-lipid complex binding | 3.1 | 0.04 |
| GO:0046470 | phosphatidylcholine metabolic process | 3.1 | 0.04 |
| GO:0035198 | miRNA binding | 3.1 | 0.04 |
| GO:0031461 | cullin-RING ubiquitin ligase complex | 3.1 | 0.04 |
| GO:0098590 | plasma membrane region | 3.1 | 0.04 |
| GO:0051347 | positive regulation of transferase activity | 3.1 | 0.04 |
| GO:2000351 | regulation of endothelial cell apoptotic process | 3.1 | 0.05 |
| GO:0051055 | negative regulation of lipid biosynthetic process | 3.1 | 0.05 |
| GO:0034308 | primary alcohol metabolic process | 3.1 | 0.05 |
| GO:0016597 | amino acid binding | 3.1 | 0.05 |
| GO:1901862 | negative regulation of muscle tissue development | 3.1 | 0.05 |
| GO:0001678 | cellular glucose homeostasis | 3.1 | 0.05 |
| GO:1901701 | cellular response to oxygen-containing compound | 3.1 | 0.05 |
| GO:0045913 | positive regulation of carbohydrate metabolic process | 3.1 | 0.05 |
| GO:0043524 | negative regulation of neuron apoptotic process | 3.1 | 0.05 |
| GO:0048534 | hematopoietic or lymphoid organ development | 3.1 | 0.05 |
| GO:0060284 | regulation of cell development | 3 | 0.05 |
| GO:2000027 | regulation of animal organ morphogenesis | 3 | 0.05 |
| GO:0042402 | cellular biogenic amine catabolic process | 3 | 0.05 |
| GO:0019585 | glucuronate metabolic process | 3 | 0.05 |
| GO:0045236 | CXCR chemokine receptor binding | 3 | 0.05 |
| GO:0046113 | nucleobase catabolic process | 3 | 0.05 |
| GO:0048019 | receptor antagonist activity | 3 | 0.05 |
| GO:1903131 | mononuclear cell differentiation | 3 | 0.05 |
| GO:0003188 | heart valve formation | 3 | 0.05 |
| GO:0003197 | endocardial cushion development | 3 | 0.05 |
| GO:0030224 | monocyte differentiation | 3 | 0.05 |
| GO:0006063 | uronic acid metabolic process | 3 | 0.05 |
| GO:0032454 | histone demethylase activity (H3-K9 specific) | 3 | 0.05 |
| GO:0008356 | asymmetric cell division | 3 | 0.05 |
| GO:0009310 | amine catabolic process | 3 | 0.05 |
| GO:0060368 | regulation of Fc receptor mediated stimulatory signaling pathway | 3 | 0.05 |
| GO:0010761 | fibroblast migration | 3 | 0.05 |
| GO:0048856 | anatomical structure development | 3 | 0.05 |
| GO:0042490 | mechanoreceptor differentiation | 3 | 0.05 |
| GO:1904666 | regulation of ubiquitin protein ligase activity | 3 | 0.05 |
| GO:0004602 | glutathione peroxidase activity | 3 | 0.05 |
| GO:0033599 | regulation of mammary gland epithelial cell proliferation | 3 | 0.05 |
| GO:0048525 | negative regulation of viral process | 3 | 0.05 |
| GO:0071345 | cellular response to cytokine stimulus | 3 | 0.05 |

| | | | |
|---|---|---|---|
| GO:0045332 | phospholipid translocation | 3 | 0.05 |
| GO:0071326 | cellular response to monosaccharide stimulus | 3 | 0.05 |
| GO:0050850 | positive regulation of calcium-mediated signaling | 3 | 0.05 |
| GO:0001102 | RNA polymerase II activating transcription factor binding | 3 | 0.05 |
| GO:0010464 | regulation of mesenchymal cell proliferation | 3 | 0.05 |
| GO:0009617 | response to bacterium | 3 | 0.05 |
| GO:0006879 | cellular iron ion homeostasis | 3 | 0.05 |
| GO:0000175 | 3'-5'-exoribonuclease activity | 3 | 0.05 |
| GO:1903959 | regulation of anion transmembrane transport | 3 | 0.05 |
| GO:0030291 | protein serine/threonine kinase inhibitor activity | 3 | 0.05 |
| GO:0032369 | negative regulation of lipid transport | 3 | 0.05 |
| GO:0090169 | regulation of spindle assembly | 3 | 0.05 |
| GO:0004842 | ubiquitin-protein transferase activity | 3 | 0.05 |
| GO:0110110 | positive regulation of animal organ morphogenesis | 3 | 0.05 |
| GO:0006497 | protein lipidation | 3 | 0.05 |
| GO:0016202 | regulation of striated muscle tissue development | 3 | 0.05 |
| GO:0071495 | cellular response to endogenous stimulus | 3 | 0.05 |
| GO:0017124 | SH3 domain binding | 3 | 0.05 |
| GO:0016811 | hydrolase activity, acting on carbon-nitrogen (but not peptide) bonds, in linear amides | 2.9 | 0.05 |
| GO:0061061 | muscle structure development | 2.9 | 0.05 |
| GO:0140297 | DNA-binding transcription factor binding | 2.9 | 0.05 |
| GO:0048878 | chemical homeostasis | 2.9 | 0.05 |
| GO:0016620 | oxidoreductase activity, acting on the aldehyde or oxo group of donors, NAD or NADP as acceptor | 2.9 | 0.05 |
| GO:0071364 | cellular response to epidermal growth factor stimulus | 2.9 | 0.05 |
| GO:1901570 | fatty acid derivative biosynthetic process | 2.9 | 0.05 |
| GO:0007098 | centrosome cycle | 2.9 | 0.05 |
| GO:0032270 | positive regulation of cellular protein metabolic process | 2.9 | 0.05 |
| GO:0043666 | regulation of phosphoprotein phosphatase activity | 2.9 | 0.05 |
| GO:0070936 | protein K48-linked ubiquitination | 2.9 | 0.05 |

Analysis done using Partek Flow package.

**Table S5**- Pathway enrichment analysis of differentially expressed genes in the ALB-D2KO mouse liver.

| Gene set | Description | Enrichment score | P-value |
|---|---|---|---|
| path:mmu00983 | Drug metabolism - other enzymes | 20.5 | 1.22E-09 |
| path:mmu05204 | Chemical carcinogenesis | 18.3 | 1.18E-08 |
| path:mmu00982 | Drug metabolism - cytochrome P450 | 17.4 | 2.68E-08 |
| path:mmu00830 | Retinol metabolism | 15.5 | 1.79E-07 |
| path:mmu00071 | Fatty acid degradation | 14.0 | 8.59E-07 |
| path:mmu03320 | PPAR signaling pathway | 12.5 | 3.61E-06 |
| path:mmu00980 | Metabolism of xenobiotics by cytochrome P450 | 11.8 | 7.28E-06 |
| path:mmu00140 | Steroid hormone biosynthesis | 11.5 | 9.92E-06 |
| path:mmu00480 | Glutathione metabolism | 10.9 | 1.78E-05 |
| path:mmu01100 | Metabolic pathways | 10.2 | 3.64E-05 |
| path:mmu01040 | Biosynthesis of unsaturated fatty acids | 7.9 | 3.79E-04 |
| path:mmu00590 | Arachidonic acid metabolism | 7.1 | 8.23E-04 |
| path:mmu00591 | Linoleic acid metabolism | 6.5 | 1.48E-03 |
| path:mmu01524 | Platinum drug resistance | 5.3 | 4.79E-03 |
| path:mmu01212 | Fatty acid metabolism | 5.2 | 0.01 |
| path:mmu00330 | Arginine and proline metabolism | 5.1 | 0.01 |
| path:mmu00920 | Sulfur metabolism | 4.9 | 0.01 |
| path:mmu00360 | Phenylalanine metabolism | 4.6 | 0.01 |
| path:mmu00053 | Ascorbate and aldarate metabolism | 4.6 | 0.01 |
| path:mmu00280 | Valine, leucine and isoleucine degradation | 4.5 | 0.01 |
| path:mmu00640 | Propanoate metabolism | 4.5 | 0.01 |
| path:mmu00040 | Pentose and glucuronate interconversions | 4.3 | 0.01 |
| path:mmu00340 | Histidine metabolism | 4.2 | 0.02 |
| path:mmu04979 | Cholesterol metabolism | 4.1 | 0.02 |
| path:mmu00410 | beta-Alanine metabolism | 3.9 | 0.02 |
| path:mmu00650 | Butanoate metabolism | 3.8 | 0.02 |
| path:mmu00561 | Glycerolipid metabolism | 3.7 | 0.03 |
| path:mmu04146 | Peroxisome | 3.6 | 0.03 |
| path:mmu00380 | Tryptophan metabolism | 3.4 | 0.03 |
| path:mmu00770 | Pantothenate and CoA biosynthesis | 3.3 | 0.04 |
| path:mmu05418 | Fluid shear stress and atherosclerosis | 3.1 | 0.05 |
| path:mmu05225 | Hepatocellular carcinoma | 3.1 | 0.05 |
| path:mmu00062 | Fatty acid elongation | 3.0 | 0.05 |

Analysis done using Partek Flow package.

**Table S6**- Genes downregulated in the Alb-D2KO mouse liver that are within 5,000 bp of a p-RCA area along with their coordinates. Also included are the function of the p-RCAs, the negative transcription factor footprints identified in these p-RCAs, the annotation of the nearest H-site, the RNA-seq Alb-D2KO/control ratio and relative drop in expression.

| | Function | Chr | Start | Stop | Motif | | | | | | | H-site annotation | Rna-seq data | |
|---|---|---|---|---|---|---|---|---|---|---|---|---|---|---|
| | | | | | **Nrf1** | **Sp1-2-3** | **Klf1** | **CTCF** | **Elk1** | **E2F3** | **EGR2** | | **ratio** | **drop** |
| Jph1 | Insulator | 1 | 16964560 | 17097890 | + | - | - | - | - | - | - | | 0.59 | -1.7 |
| Ankrd44 | Promoter | 1 | 54645340 | 54926388 | + | + | - | - | - | - | - | | 0.67 | -1.49 |
| Gm37423 | Promoter | 1 | 138968510 | 138971852 | - | + | - | - | + | - | - | | 0.54 | -1.84 |
| Rgs16 | Promoter | 1 | 153740349 | 153745469 | - | ++ | + | - | - | + | - | | 0.6 | -1.66 |
| Rc3h1 | Insulator | 1 | 160906418 | 160974979 | - | - | - | - | - | - | - | | 0.81 | -1.23 |
| Fmo1 | Insulator | 1 | 162829561 | 162866611 | - | - | - | - | - | - | - | | 0.75 | -1.34 |
| 1700047M11Rik* | Insulator | 1 | 182287990 | 182303326 | - | - | - | - | - | - | - | promoter | 0.64 | -1.56 |
| Tlr5 | Promoter | 1 | 182954788 | 182976045 | - | - | - | - | - | - | - | | 0.65 | -1.53 |
| Mcm10 | Promoter | 2 | 4989714 | 5012792 | - | - | - | - | - | - | - | | 0.69 | -1.46 |
| Echdc3 | Weak Enhancer | 2 | 6188465 | 6213034 | - | - | - | - | - | - | - | | 0.77 | -1.3 |
| Mllt10 | Promoter | 2 | 18055237 | 18212389 | - | - | + | - | - | - | - | | 0.8 | -1.25 |
| Cercam | Promoter | 2 | 29869164 | 29882841 | - | ++ | - | + | | - | - | | 0.59 | -1.7 |
| Acvr2a | Promoter | 2 | 48814109 | 48903270 | - | + | + | - | + | - | - | | 0.8 | -1.26 |
| Cobll1 | Promoter | 2 | 65088339 | 65239404 | + | + | - | - | + | - | - | | 0.75 | -1.33 |
| Sqor | Insulator | 2 | 122765237 | 122809570 | - | - | - | - | - | - | - | | 0.73 | -1.37 |
| Myef2 | Promoter | 2 | 125084628 | 125123662 | - | + | + | - | - | + | + | | 0.74 | -1.35 |
| 9030622O22Rik | Promoter | 2 | 147944740 | 148040204 | + | + | - | - | - | - | - | | 0.65 | -1.53 |
| Foxa2 | Promoter | 2 | 148042877 | 148046970 | + | + | - | - | - | - | - | | 0.64 | -1.57 |
| Slc52a3 | Insulator | 2 | 151996511 | 152009259 | + | - | + | - | - | + | - | | 0.51 | -1.98 |
| Zfp704 | Insulator | 3 | 9427020 | 9610086 | - | - | - | - | - | - | - | | 0.66 | -1.51 |
| 1700112D23Rik | Promoter | 3 | 28702667 | 28781109 | - | - | + | - | - | - | - | | 0.55 | -1.81 |
| Tm4sf4 | Promoter | 3 | 57425314 | 57441678 | - | - | - | - | - | - | - | | 0.69 | -1.44 |

| | | | | | | | | | | | | | | |
|---|---|---|---|---|---|---|---|---|---|---|---|---|---|---|
| Tiparp | Promoter | 3 | 65528410 | 65555519 | - | - | + | - | - | + | - | | 0.74 | -1.34 |
| Pdgfc | Promoter | 3 | 81036416 | 81214041 | - | + | + | - | - | - | - | | 0.68 | -1.48 |
| Gm38227 | Insulator | 3 | 96263785 | 96265387 | - | - | - | - | + | - | - | | 0.46 | -2.19 |
| Rap1a | Promoter | 3 | 105727267 | 105801337 | - | ++ | + | - | + | - | - | | 0.79 | -1.27 |
| Gstm6* | Weak Enhancer | 3 | 107938847 | 107943750 | - | - | + | - | - | - | - | promoter | 0.53 | -1.89 |
| Gstm1 | Enhancer | 3 | 108012255 | 108017974 | - | - | - | - | - | - | - | | 0.7 | -1.43 |
| Slc39a8 | Promoter | 3 | 135825279 | 135888573 | - | - | - | - | - | - | - | | 0.79 | -1.26 |
| Ddah1 | Insulator | 3 | 145758675 | 145894278 | + | - | - | - | - | + | - | | 0.56 | -1.78 |
| Adgrl2 | Promoter | 3 | 148815583 | 148990556 | + | + | - | - | + | + | - | | 0.76 | -1.32 |
| Bach2 | Weak Enhancer | 4 | 32238804 | 32586109 | - | - | - | - | - | - | - | | 0.64 | -1.56 |
| D630039A03Rik | Insulator | 4 | 57908483 | 57916298 | - | - | - | - | - | + | - | | 0.47 | -2.14 |
| Gm20503 | Insulator | 4 | 59003210 | 59041889 | + | + | + | - | + | + | - | | 0.26 | -3.79 |
| Lurap1l | Insulator | 4 | 80910646 | 80955629 | - | - | - | - | + | + | - | | 0.74 | -1.35 |
| Nfib | Promoter | 4 | 82290173 | 82705751 | - | ++ | + | - | - | + | - | | 0.75 | -1.33 |
| Dock7 | Promoter | 4 | 98936671 | 99120916 | + | - | - | - | - | - | - | | 0.68 | -1.47 |
| Tut4 | Promoter | 4 | 108459426 | 108559422 | - | - | - | - | + | - | + | | 0.69 | -1.45 |
| Pum1 | Promoter | 4 | 130663321 | 130781565 | - | + | - | - | - | - | - | | 0.8 | -1.25 |
| Rnf186 | Promoter | 4 | 138967112 | 138968361 | - | - | - | - | - | - | - | intergenic | 0.47 | -2.12 |
| N4bp2 | Promoter | 5 | 65763521 | 65830109 | - | - | - | - | - | - | - | | 0.76 | -1.32 |
| Rbm47 | Insulator | 5 | 66016549 | 66173119 | + | + | - | - | - | - | - | | 0.67 | -1.49 |
| Albfm1 | Insulator | 5 | 90561107 | 90597872 | - | - | - | - | - | - | - | | 0.64 | -1.56 |
| Arhgap24 | Promoter | 5 | 102481391 | 102897938 | - | + | - | - | - | - | + | | 0.75 | -1.33 |
| Gbp11 | Insulator | 5 | 105323042 | 105346473 | - | + | + | - | - | - | - | | 0.26 | -3.9 |
| 3110082I17Rik* | Weak Enhancer | 5 | 139359739 | 139460528 | - | + | + | - | - | - | - | | 0.64 | -1.57 |
| Smurf1 | Promoter | 5 | 144876495 | 144965848 | - | - | - | - | - | - | - | | 0.76 | -1.31 |
| Akr1d1 | Promoter | 6 | 37530173 | 37568816 | - | + | + | - | - | - | - | | 0.78 | -1.28 |
| E330009J07Rik | Promoter | 6 | 40401375 | 40436136 | + | + | + | - | - | - | - | intron | 0.76 | -1.32 |

| | | | | | | | | | | | | | | |
|---|---|---|---|---|---|---|---|---|---|---|---|---|---|---|
| Eva1a | Promoter | 6 | 82041043 | 82093100 | + | - | + | - | - | + | - | | 0.76 | -1.31 |
| Gm44005 | Insulator | 6 | 88738192 | 88741365 | - | - | - | - | - | + | - | | 0.6 | -1.67 |
| Gm43999 | Insulator | 6 | 88742727 | 88745616 | - | - | - | - | - | + | - | | 0.6 | -1.67 |
| Suclg2 | Promoter | 6 | 95473009 | 95718801 | + | - | - | - | - | - | - | | 0.81 | -1.24 |
| Atn1 | Promoter | 6 | 124742544 | 124756525 | - | - | - | - | - | - | - | | 0.8 | -1.25 |
| Dbp | Promoter | 7 | 45705088 | 45710204 | + | + | + | - | - | - | - | | 0.74 | -1.35 |
| 1810026B05Rik | Promoter | 7 | 73539801 | 73558396 | - | - | - | - | + | - | - | | 0.77 | -1.3 |
| Crtc3 | Promoter | 7 | 80586627 | 80688878 | - | - | - | - | - | + | - | | 0.73 | -1.37 |
| Acsm5 | Insulator | 7 | 119519463 | 119545552 | - | - | - | - | - | - | - | | 0.7 | -1.42 |
| Gm19950 | Insulator | 7 | 119607681 | 119643470 | - | - | - | - | - | - | - | | 0.71 | -1.41 |
| Olfr542-ps1 | Insulator | 7 | 140717473 | 140722113 | - | - | - | - | - | - | - | | 0.33 | -2.99 |
| BC024386 | Promoter | 7 | 140906014 | 140907984 | - | - | - | - | - | - | - | | 0.79 | -1.26 |
| Snapc2 | Insulator | 8 | 4253080 | 4256221 | - | + | - | + | - | - | - | | 0.76 | -1.32 |
| Gm17491 | Promoter | 8 | 22480291 | 22483333 | - | - | - | - | - | - | - | | 0.72 | -1.39 |
| Rbpms | Promoter | 8 | 33782643 | 33929864 | + | ++ | - | - | + | - | - | | 0.78 | -1.28 |
| Trmt9b | Promoter | 8 | 36457548 | 36516571 | - | - | - | - | - | + | - | | 0.53 | -1.88 |
| Mtus1* | Weak Promoter | 8 | 40990914 | 41133727 | - | - | - | - | - | - | - | 3’UTR | 0.78 | -1.28 |
| Large1 | Promoter | 8 | 72814599 | 73353541 | + | + | - | - | - | - | - | | 0.79 | -1.26 |
| Ces1c | Insulator | 8 | 93099015 | 93131284 | - | - | - | - | - | - | - | | 0.76 | -1.31 |
| Ces1e | Weak Enhancer | 8 | 93201218 | 93229620 | - | - | - | - | - | - | - | | 0.66 | -1.52 |
| Csnk2a2 | Weak Enhancer | 8 | 95446096 | 95490040 | + | + | - | - | - | + | - | | 0.79 | -1.26 |
| 1190005I06Rik | Promoter | 8 | 120608602 | 120647633 | + | ++ | - | - | - | + | - | | 0.68 | -1.47 |
| Pknox2 | Promoter | 9 | 36890982 | 37147408 | + | - | - | + | - | - | - | | 0.64 | -1.57 |
| Gm47204 | Weak Enhancer | 9 | 44477928 | 44479760 | - | - | - | + | - | - | - | | 0.43 | -2.32 |
| A830035O19Rik | Promoter | 9 | 45954101 | 45955999 | - | - | - | - | - | - | - | | 0.59 | -1.7 |
| Dixdc1* | - | 9 | 50662752 | 50739518 | - | - | - | - | - | - | - | intron | 0.78 | -1.28 |

| | | | | | | | | | | | | | | |
|---|---|---|---|---|---|---|---|---|---|---|---|---|---|---|
| Zfp609 | Insulator | 9 | 65692391 | 65827565 | - | - | - | - | - | + | - | | 0.74 | -1.35 |
| Gclc | Insulator | 9 | 77754535 | 77794486 | + | - | - | - | - | + | - | | 0.72 | -1.39 |
| Hyal3 | Enhancer | 9 | 107580818 | 107587360 | - | - | - | - | - | - | - | | 0.48 | -2.07 |
| Traip | Insulator | 9 | 107950136 | 107972271 | - | - | - | - | - | - | - | | 0.39 | -2.57 |
| Nicn1 | Promoter | 9 | 108290429 | 108296499 | - | ++ | - | - | - | - | - | | 0.79 | -1.27 |
| Acaa1b | Promoter | 9 | 119148023 | 119157094 | - | - | - | - | - | - | - | | 0.72 | -1.4 |
| Pex7 | Promoter | 10 | 19853900 | 19907690 | - | - | - | - | + | - | + | | 0.8 | -1.25 |
| Vnn3 | Insulator | 10 | 23851462 | 23869844 | - | - | - | - | - | - | - | | 0.63 | -1.59 |
| Gstt3 | Promoter | 10 | 75774115 | 75781415 | - | - | - | - | - | + | - | | 0.7 | -1.44 |
| Gstt1 | Promoter | 10 | 75783813 | 75798585 | - | - | - | - | - | + | - | | 0.77 | -1.29 |
| Appl2 | Weak Enhancer | 10 | 83600033 | 83648739 | - | - | - | - | - | - | - | | 0.77 | -1.3 |
| Stab2 | Insulator | 10 | 86841198 | 87008026 | - | - | - | - | - | - | - | | 0.81 | -1.24 |
| Cradd | Promoter | 10 | 95174746 | 95324134 | + | - | - | - | - | + | - | | 0.76 | -1.32 |
| Socs2 | Promoter | 10 | 95385362 | 95417181 | - | + | - | - | - | + | - | | 0.25 | -3.96 |
| Tbc1d30 | Promoter | 10 | 121263820 | 121351279 | + | ++ | - | - | - | + | - | | 0.61 | -1.64 |
| Ikzf4 | Promoter | 10 | 128630843 | 128669359 | - | - | - | - | - | + | - | | 0.38 | -2.66 |
| Flcn | Promoter | 11 | 59791408 | 59810017 | - | - | - | - | - | + | - | | 0.66 | -1.52 |
| Ncor1 | Promoter | 11 | 62316426 | 62458542 | + | - | - | - | + | - | - | | 0.8 | -1.24 |
| Rnf167 | Promoter | 11 | 70647235 | 70651422 | + | + | - | - | + | - | - | | 0.78 | -1.29 |
| Cdk5r1 | Promoter | 11 | 80477023 | 80481185 | - | - | - | - | - | - | - | | 0.62 | -1.61 |
| Abcc3 | Promoter | 11 | 94343295 | 94392998 | - | + | + | - | - | - | - | | 0.61 | -1.65 |
| Afmid | Promoter | 11 | 117825924 | 117839909 | - | + | - | - | + | - | - | | 0.76 | -1.32 |
| Atxn7l1 | Weak Enhancer | 12 | 33147693 | 33373186 | - | - | - | - | - | - | - | non-coding | 0.63 | -1.59 |
| Gm24233 | Insulator | 12 | 59134072 | 59134194 | - | - | + | - | - | - | - | | 0.42 | -2.4 |
| Sav1 | Promoter | 12 | 69965012 | 69987003 | + | - | - | - | - | + | - | | 0.74 | -1.35 |
| Zbtb1 | Promoter | 12 | 76370266 | 76396951 | + | ++ | - | - | - | - | + | | 0.79 | -1.27 |
| Acot2 | Promoter | 12 | 83987861 | 83993874 | - | - | + | - | - | - | | | 0.46 | -2.19 |

| | | | | | | | | | | | | | | |
|---|---|---|---|---|---|---|---|---|---|---|---|---|---|---|
| Dnal1 | Promoter | 12 | 84114366 | 84147499 | + | - | - | - | - | + | + | | 0.58 | -1.74 |
| Serpina6 | Insulator | 12 | 103646630 | 103657213 | - | - | - | - | - | + | - | | 0.67 | -1.49 |
| Gm34667 | Promoter | 12 | 109990882 | 110023874 | - | - | - | - | - | - | - | | 0.65 | -1.53 |
| Eif5* | Promoter | 12 | 111538016 | 111546753 | + | - | - | - | - | - | - | promoter | 0.8 | -1.24 |
| Ighm | Promoter | 12 | 113418558 | 113422731 | - | - | - | - | - | - | - | | 0.58 | -1.74 |
| Aldh5a1 | Promoter | 13 | 24907579 | 24937662 | - | + | - | + | - | + | - | | 0.81 | -1.24 |
| Dsp* | Promoter | 13 | 38151294 | 38198578 | - | + | + | - | - | - | - | Intron/Cds | 0.75 | -1.34 |
| Dapk1 | Insulator | 13 | 60601947 | 60763192 | - | - | - | - | - | - | - | | 0.74 | -1.34 |
| Pde4d* | Promoter | 13 | 108449948 | 109953462 | - | - | - | - | - | - | - | intron | 0.57 | -1.76 |
| Chdh | Promoter | 14 | 30009023 | 30040528 | - | ++ | - | - | - | + | - | | 0.8 | -1.25 |
| AW549877 | Promoter | 15 | 3982035 | 3995745 | - | - | + | - | - | - | + | | 0.77 | -1.3 |
| Prkaa1 | Promoter | 15 | 5143861 | 5181900 | - | ++ | - | - | - | + | - | | 0.8 | -1.25 |
| Dpys | Promoter | 15 | 39768487 | 39857471 | - | - | - | - | - | - | - | | 0.77 | -1.31 |
| Gm49413 | Weak Enhancer | 15 | 76577825 | 76580864 | - | - | - | - | - | - | - | | 0.29 | -3.5 |
| 1110025M09Rik | Promoter | 15 | 81584174 | 81585029 | - | + | - | - | - | - | - | | 0.59 | -1.69 |
| Sco2* | Promoter | 15 | 89371637 | 89373847 | - | + | - | + | - | - | - | promoter | 0.6 | -1.67 |
| Tymp* | Promoter | 15 | 89371931 | 89377040 | - | + | - | + | - | - | - | promoter | 0.64 | -1.57 |
| Odf3b* | Promoter | 15 | 89377449 | 89379588 | - | + | - | - | - | + | - | promoter | 0.73 | -1.37 |
| Adcy6 | Promoter | 15 | 98589973 | 98610077 | - | - | - | - | - | - | - | | 0.81 | -1.24 |
| Soat2 | Promoter | 15 | 102150526 | 102163470 | - | - | + | - | + | - | - | | 0.76 | -1.31 |
| Soat2 | Promoter | 15 | 102150526 | 102163470 | | | | | | | | | 0.76 | -1.31 |
| Kng2 | Insulator | 16 | 22985854 | 23029483 | - | - | - | - | - | - | - | | 0.8 | -1.26 |
| B630019A10Rik | Promoter | 16 | 23175280 | 23224447 | - | - | - | - | - | - | - | | 0.65 | -1.54 |
| Gm15746 | Promoter | 16 | 23197095 | 23198122 | - | - | - | - | - | - | - | | 0.46 | -2.2 |
| BC106179 | Promoter | 16 | 23220656 | 23225485 | + | - | - | - | - | - | - | | 0.38 | -2.62 |
| Gm31583 | Insulator | 16 | 24085083 | 24090090 | - | - | + | - | - | + | - | | 0.64 | -1.55 |
| Pla1a | Insulator | 16 | 38396117 | 38433146 | - | - | + | - | - | - | + | | 0.68 | -1.47 |
| Gm15638 | Promoter | 16 | 45945606 | 45971371 | - | - | - | - | - | - | - | | 0.67 | -1.5 |

| | | | | | | | | | | | | | |
|---|---|---|---|---|---|---|---|---|---|---|---|---|---|
| C2cd2 | Promoter | 16 | 97855209 | 97962599 | + | - | + | - | - | - | - | | 0.82 | -1.22 |
| Angptl4 | Promoter | 17 | 33773750 | 33781831 | - | ++ | + | - | - | - | - | | 0.76 | -1.31 |
| Gm20522 | Promoter | 17 | 35122738 | 35126441 | - | ++ | + | - | + | - | + | | 0.44 | -2.25 |
| 1700001C19Rik | Promoter | 17 | 47412734 | 47437377 | - | - | - | - | - | - | - | | 0.78 | -1.29 |
| Kat2b | Promoter | 17 | 53566861 | 53672721 | - | - | - | - | + | - | - | | 0.81 | -1.23 |
| Osbpl1a | Weak Enhancer | 18 | 12755314 | 12941842 | - | - | - | - | - | - | - | | 0.8 | -1.24 |
| Kdm3b | Insulator | 18 | 34777047 | 34838661 | - | + | - | - | | - | - | | 0.77 | -1.3 |
| Cdo1* | Promoter | 18 | 46713193 | 46728396 | - | + | - | - | + | - | - | Intron/3'UTR | 0.72 | -1.39 |
| Ahnak | Promoter | 19 | 8989284 | 9076915 | + | + | - | - | - | + | - | | 0.77 | -1.29 |
| 1700018L02Rik | Promoter | 19 | 29046245 | 29048730 | + | - | - | - | - | - | - | | 0.53 | -1.9 |
| Papss2 | Promoter | 19 | 32595790 | 32667188 | - | - | - | - | - | + | - | | 0.78 | -1.29 |
| Dnmbp | Promoter | 19 | 43846821 | 43940192 | + | + | - | + | + | - | - | | 0.77 | -1.3 |
| Stag2 | Insulator | X | 42149317 | 42277186 | - | - | - | - | + | - | - | | 0.77 | -1.29 |
| G6pdx | Promoter | X | 74409483 | 74429195 | - | + | + | - | - | + | - | | 0.73 | -1.37 |
| Gk | Promoter | X | 85701937 | 85776820 | - | - | - | - | - | - | - | | 0.75 | -1.34 |
| Rab9 | Promoter | X | 166457253 | 166479868 | - | + | + | - | - | - | - | | 0.74 | -1.35 |

The function for each p-RCA area was defined according to the H3K4me1 (enhancer) and H3K4me3 (promoter) signal ratio. Signals were summarized as transcripts per kilobase millions (TPM) values in each p-RCA region and grouped. The p-RCA regions with a TPM >4 for either H3K4me3 or H3K4me1 were placed into one of three categories: High ratios >1.5 were classified as a 'promoter'; ratios <0.67 were classified as 'enhancer', while intermediate values were classified as 'weak promoter'. p-RCA regions with TPM <=4 in both H3K4me3 and H3K4me1 were placed into one of two categories: if the ATAC-seq signal >H3K4me1 signal, the p-RCA was categorized as 'insulator'; if the ATAC-seq signal <H3K4me1 signal, the RCA was categorized as 'weak insulator'. The presence or absence of the motifs was defined using the nearest location of the p-RCA to each gene: "+" motif present and "-" motif not present. Genes containing (*) reflect presence of H-site in the promoter; Nrf1 is Nuclear respiratory factor 1; Sp is Specificity Protein; Klf1 is Kruppel Like Factor 1; CTCF is CCCTC-Binding Factor; Elk1 is ETS Transcription Factor; E2F3 is E2F Transcription Factor 3 and EGR2 is Early Growth Response 2.

**Table S7** – Presence of CTCF and/or Cohesin ChIP-seq peaks in the p-RCA areas of the genes that contain a p-RCA within 5 Kbp of a n-RNA-seq. Genes not shown do not contain either CTCF and/or a Cohesin ChIP-seq peak.

| | p-RCA coordinates | | | | | |
|---|---|---|---|---|---|---|
| | **chr** | **Start** | **End** | **gene location** | **CTCF** | **Cohesin** |
| Jph1 | 1 | 17097719 | 17098091 | 5' UTR | + | + |
| Rc3h1 | 1 | 160978442 | 160978644 | 3’ UTR | - | + |
| Fmo1 | 1 | 162866509 | 162866731 | 5' UTR | + | + |
| 1700047M11Rik | 1 | 182287826 | 182288094 | 5' UTR | + | + |
| Sqor | 2 | 122781608 | 122781964 | 5' UTR | + | + |
| Slc52a3 | 2 | 151996373 | 151996544 | 5' UTR | + | + |
| Zfp704 | 3 | 9610026 | 9610292 | 5' UTR | + | + |
| Gm38227 | 3 | 96268499 | 96268727 | 5' UTR | + | + |
| Ddah1 | 3 | 145758456 | 145758830 | 5' UTR | + | + |
| D630039A03Rik | 4 | 57916192 | 57916684 | 5' UTR | + | + |
| Gm20503 | 4 | 59003007 | 59003198 | 5' UTR | + | + |
| Lurap1l | 4 | 80910539 | 80910771 | 5' UTR | + | + |
| Rbm47 | 5 | 66098169 | 66098540 | intron | - | + |
| Albfm1 | 5 | 90560701 | 90561139 | 5' UTR | + | + |
| Gbp11 | 5 | 105346456 | 105346882 | 5' UTR | + | + |
| Gm44005/Gm43999 | 6 | 88742655 | 88743083 | intron | + | + |
| Acsm5 | 7 | 119525990 | 119526343 | 5' UTR | - | + |
| Gm19950 | 7 | 119620187 | 119620568 | intron | - | + |
| Olfr542-ps1 | 7 | 140717218 | 140717925 | - | + | + |
| Snapc2 | 8 | 4250688 | 4250867 | 5' UTR | + | + |
| Ces1c | 8 | 93131216 | 93131649 | 5' UTR | + | + |
| Zfp609 | 9 | 65827661 | 65827896 | 5' UTR | + | + |
| Gclc | 9 | 77754387 | 77754633 | 5' UTR | + | + |
| Traip | 9 | 107975388 | 107975651 | 3’ UTR | + | + |
| Vnn3 | 10 | 23851327 | 23851523 | 5' UTR | + | + |
| Stab2 | 10 | 86972034 | 86972224 | intron | + | + |
| Gm24233 | 12 | 59130264 | 59130503 | - | + | + |
| Serpina6 | 12 | 103657056 | 103657505 | 5' UTR | + | + |
| Dapk1 | 13 | 60601749 | 60601948 | 5' UTR | + | + |
| Kng2 | 16 | 23029015 | 23029479 | 5' UTR | + | + |
| Gm31583 | 16 | 24090027 | 24090422 | 5' UTR | + | + |
| Pla1a | 16 | 38432879 | 38433347 | 5' UTR | + | + |
| Kdm3b | 18 | 34776752 | 34777040 | 5' UTR | + | + |
| Stag2 | X | 42148924 | 42149305 | 5' UTR | + | + |
| | | | | **total** | **30** | **34** |

The presence “+” or absence “-“of chip-Seq peak was observed using the IGV browser. The majority of the CTCF or Cohesin peaks were located at the same position that P-RCA. Just 2 RCA sites have the CTCF/Cohesin peaks at ≤300bp of distance.  RCA is reduced chromatin accessibility; CTCF is CCCTC-binding factor; **-** 3'-UTR is three prime untranslated regions and 5’UTR is five prime untranslated regions.

**Table S8**- Number (upper panel) and density (lower panel) of H-sites, RCAs, p-RCAs and negative RNA-seq peaks in adult Alb-D2KO liver.

| Chr # | H-sites | RCAs | p-RCAs | i-RCA | Rna-seq |
|---|---|---|---|---|---|
| 1 | 82 | 416 | 61 | 140 | 104 |
| 2 | 104 | 551 | 96 | 182 | 102 |
| 3 | 73 | 379 | 72 | 135 | 76 |
| 4 | 120 | 608 | 98 | 179 | 67 |
| 5 | 94 | 517 | 108 | 133 | 74 |
| 6 | 78 | 463 | 78 | 155 | 99 |
| 7 | 74 | 463 | 93 | 141 | 103 |
| 8 | 92 | 453 | 72 | 137 | 56 |
| 9 | 83 | 454 | 98 | 140 | 74 |
| 10 | 74 | 440 | 82 | 133 | 93 |
| 11 | 105 | 687 | 166 | 205 | 58 |
| 12 | 151 | 319 | 59 | 113 | 78 |
| 13 | 54 | 315 | 51 | 115 | 65 |
| 14 | 54 | 307 | 50 | 84 | 33 |
| 15 | 65 | 413 | 96 | 112 | 58 |
| 16 | 45 | 253 | 50 | 58 | 38 |
| 17 | 60 | 383 | 87 | 108 | 49 |
| 18 | 57 | 237 | 35 | 72 | 48 |
| 19 | 38 | 321 | 66 | 77 | 45 |
| X | 5 | 40 | 33 | 6 | 42 |
| Y | 0 | 2 | 0 | 1 | 1 |

| Chr # | H-sites/Chr Mbp | RCAs/Chr Mbp | i-RCAs/Mbp | i-RCA | Rna-seq/Chr Mbp |
|---|---|---|---|---|---|
| 1 | 0.45 | 2.27 | 0.33 | 0.77 | 0.57 |
| 2 | 0.62 | 3.26 | 0.57 | 1.08 | 0.60 |
| 3 | 0.49 | 2.54 | 0.48 | 0.91 | 0.51 |
| 4 | 0.86 | 4.34 | 0.70 | 1.28 | 0.48 |
| 5 | 0.69 | 3.77 | 0.79 | 0.97 | 0.54 |
| 6 | 0.57 | 3.36 | 0.57 | 1.12 | 0.72 |
| 7 | 0.61 | 3.80 | 0.76 | 1.16 | 0.84 |
| 8 | 0.77 | 3.81 | 0.61 | 1.15 | 0.47 |
| 9 | 0.72 | 3.91 | 0.84 | 1.21 | 0.64 |
| 10 | 0.61 | 3.64 | 0.68 | 1.10 | 0.77 |
| 11 | 0.91 | 5.97 | 1.44 | 1.78 | 0.50 |
| 12 | 1.44 | 3.04 | 0.56 | 1.08 | 0.74 |
| 13 | 0.50 | 2.94 | 0.48 | 1.07 | 0.61 |
| 14 | 0.50 | 2.87 | 0.47 | 0.79 | 0.31 |
| 15 | 0.68 | 4.30 | 1.00 | 1.17 | 0.60 |
| 16 | 0.49 | 2.78 | 0.55 | 0.64 | 0.42 |
| 17 | 0.71 | 4.51 | 1.02 | 1.27 | 0.58 |
| 18 | 0.68 | 2.82 | 0.42 | 0.86 | 0.57 |
| 19 | 0.69 | 5.84 | 1.20 | 1.40 | 0.82 |
| X | 0.04 | 0.30 | 0.25 | 0.04 | 0.31 |
| Y | n.d. | n.d. | n.d | n.d. | n.d. |

n.d. not determined; Chr is chromosome; H-sites is site of DNA methylation; RCAs is reduced chromatin accessibility area; p-RCAs is RCA area in core promoters; i-RCA is RCA area intergenic; Mbp is mega base pair.

**Table S9** – Proximity of p-RCA, H-site and i-RCA sites to the *area-2* obtained from the Hi-C contact matrix.

| | **p-RCA Coordinates** | | **Hi-C *site 1* Coordinates** | | **Hi-C *site 2* Coordinates** | | **site-1 vs site-2 Distance** | | **H-site Coordinates** | | **Site-2 vs H-site Distance** | **i-RCA Coordinates** | | **Site-2 vs i-RCA Distance** |
|---|---|---|---|---|---|---|---|---|---|---|---|---|---|---|
| | **start** | **end** | **start** | **end** | **start** | **end** | **kb** | **logP** | **kb** | **end** | **kb** | **start** | **end** | **kb** |
| chr1 | 17097719 | 17098091 | 17089214 | 17099214 | 15888950 | 15898950 | 1200 | 5.6 | 13104919 | 13105419 | -2789 | 15997194 | 15997471 | 103 |
| chr1 | 17097719 | 17098091 | 17092099 | 17102099 | 15966954 | 15976954 | 1125 | 4.9 | 13104919 | 13105419 | -2867 | 15997194 | 15997471 | 25 |
| chr1 | 17097719 | 17098091 | 17087784 | 17097784 | 16111470 | 16121470 | 976 | 9.4 | 13104919 | 13105419 | -3011 | 15997194 | 15997471 | -119 |
| chr1 | 17097719 | 17098091 | 17098606 | 17108606 | 16297426 | 16307426 | 801 | 5.0 | 13104919 | 13105419 | -3197 | 15997194 | 15997471 | -305 |
| chr1 | 17097719 | 17098091 | 17088137 | 17098137 | 16454468 | 16464468 | 634 | 5.6 | 13104919 | 13105419 | -3354 | 15997194 | 15997471 | -462 |
| chr1 | 17097719 | 17098091 | 17094512 | 17104512 | 16512197 | 16522197 | 582 | 5.6 | 13104919 | 13105419 | -3412 | 15997194 | 15997471 | -520 |
| chr1 | 17097719 | 17098091 | 17088668 | 17098668 | 16585576 | 16595576 | 503 | 4.8 | 13104919 | 13105419 | -3485 | 15997194 | 15997471 | -593 |
| chr1 | 17097719 | 17098091 | 17087547 | 17097547 | 16820862 | 16830862 | 267 | 6.1 | 13104919 | 13105419 | -3721 | 15997194 | 15997471 | -829 |
| chr1 | 17097719 | 17098091 | 17088897 | 17098897 | 17048736 | 17058736 | 40 | 5.1 | 13104919 | 13105419 | -3949 | 15997194 | 15997471 | -1056 |
| chr1 | 54926248 | 54926723 | 54916257 | 54926257 | 54654473 | 54664473 | 262 | 10.9 | 53977156 | 53977656 | -682 | 54615018 | 54615302 | -44 |
| chr1 | 54926248 | 54926723 | 54925486 | 54935486 | 54827190 | 54837190 | 98 | 20.8 | 55362156 | 55362656 | 530 | 54615018 | 54615302 | -217 |
| chr1 | 138963183 | 138963655 | 138961480 | 138971480 | 138837959 | 138847959 | 124 | 7.3 | 135874423 | 135874923 | -2968 | 138912341 | 138912560 | 69 |
| chr1 | 138963183 | 138963655 | 138955863 | 138965863 | 138918414 | 138928414 | 37 | 7.1 | 135874423 | 135874923 | -3049 | 138912341 | 138912560 | -11 |
| chr1 | 138963183 | 138963655 | 138952541 | 138962541 | 138952541 | 138962541 | 0 | 5.1 | 135874423 | 135874923 | -3083 | 138912341 | 138912560 | -45 |
| chr1 | 153740184 | 153740436 | 153739881 | 153749881 | 153440107 | 153450107 | 300 | 9.4 | 153904870 | 153905370 | 460 | 153559366 | 153559537 | 114 |
| chr1 | 153749318 | 153749738 | 153739881 | 153749881 | 153440107 | 153450107 | 300 | 9.4 | 153904870 | 153905370 | 460 | 153559366 | 153559537 | 114 |
| chr1 | 153749318 | 153749738 | 153749361 | 153759361 | 153466914 | 153476914 | 282 | 4.6 | 153904870 | 153905370 | 433 | 153559366 | 153559537 | 88 |
| chr1 | 153749318 | 153749738 | 153741984 | 153751984 | 153543741 | 153553741 | 198 | 15.9 | 153904870 | 153905370 | 356 | 153559366 | 153559537 | 11 |
| chr1 | 153749318 | 153749738 | 153745439 | 153755439 | 153563213 | 153573213 | 182 | 34.5 | 153904870 | 153905370 | 337 | 153559366 | 153559537 | -9 |
| chr1 | 153749318 | 153749738 | 153743384 | 153753384 | 153611214 | 153621214 | 132 | 10.1 | 153904870 | 153905370 | 289 | 153559366 | 153559537 | -57 |
| chr1 | 153740184 | 153740436 | 153739448 | 153749448 | 153625809 | 153635809 | 114 | 6.8 | 153904870 | 153905370 | 274 | 153559366 | 153559537 | -71 |
| chr1 | 153749318 | 153749738 | 153739448 | 153749448 | 153625809 | 153635809 | 114 | 6.8 | 153904870 | 153905370 | 274 | 153559366 | 153559537 | -71 |
| chr1 | 153749318 | 153749738 | 153741555 | 153751555 | 153637870 | 153647870 | 104 | 7.9 | 153904870 | 153905370 | 262 | 153559366 | 153559537 | -83 |
| chr1 | 153740184 | 153740436 | 153730898 | 153740898 | 153648139 | 153658139 | 83 | 13.2 | 153904870 | 153905370 | 252 | 153559366 | 153559537 | -94 |
| chr1 | 153740184 | 153740436 | 153731882 | 153741882 | 153671125 | 153681125 | 61 | 10.7 | 153904870 | 153905370 | 229 | 153559366 | 153559537 | -117 |
| chr1 | 153740184 | 153740436 | 153739719 | 153749719 | 153692346 | 153702346 | 47 | 9.8 | 153904870 | 153905370 | 208 | 153559366 | 153559537 | -138 |
| chr1 | 153749318 | 153749738 | 153739719 | 153749719 | 153692346 | 153702346 | 47 | 9.8 | 153904870 | 153905370 | 208 | 153559366 | 153559537 | -138 |
| chr1 | 160978442 | 160978644 | 160974112 | 160984112 | 160401704 | 160411704 | 572 | 4.9 | 160988869 | 160989369 | 582 | 163338971 | 163339217 | 2932 |
| chr1 | 160978442 | 160978644 | 160974352 | 160984352 | 160519375 | 160529375 | 455 | 5.1 | 160988869 | 160989369 | 465 | 163338971 | 163339217 | 2815 |
| chr1 | 160978442 | 160978644 | 160971988 | 160981988 | 160608514 | 160618514 | 363 | 6.0 | 160988869 | 160989369 | 376 | 163338971 | 163339217 | 2726 |
| chr1 | 160978442 | 160978644 | 160973444 | 160983444 | 160738144 | 160748144 | 235 | 10.2 | 160988869 | 160989369 | 246 | 163338971 | 163339217 | 2596 |
| chr1 | 160978442 | 160978644 | 160976327 | 160986327 | 160782234 | 160792234 | 194 | 5.2 | 160988869 | 160989369 | 202 | 163338971 | 163339217 | 2552 |
| chr1 | 162866509 | 162866731 | 162867378 | 162877378 | 162846629 | 162856629 | 21 | 5.8 | 161648369 | 161648869 | -1203 | 163338971 | 163339217 | 487 |
| chr1 | 182287826 | 182288094 | 182284899 | 182294899 | 181896123 | 181906123 | 389 | 8.3 | 181510369 | 181510869 | -391 | 182268298 | 182268576 | 367 |
| chr1 | 182954688 | 182955252 | 182943795 | 182953795 | 182094297 | 182104297 | 849 | 5.6 | 182283369 | 182283869 | 184 | 182268298 | 182268576 | 169 |
| chr1 | 182287826 | 182288094 | 182280389 | 182290389 | 182110322 | 182120322 | 170 | 4.8 | 182283369 | 182283869 | 168 | 182268298 | 182268576 | 153 |
| chr1 | 182287826 | 182288094 | 182284696 | 182294696 | 182154515 | 182164515 | 130 | 38.0 | 182283369 | 182283869 | 124 | 182268298 | 182268576 | 109 |
| chr1 | 182287826 | 182288094 | 182285294 | 182295294 | 182178989 | 182188989 | 106 | 16.8 | 182283369 | 182283869 | 100 | 182268298 | 182268576 | 84 |
| chr1 | 182287826 | 182288094 | 182285788 | 182295788 | 182217365 | 182227365 | 68 | 13.6 | 182283369 | 182283869 | 61 | 182268298 | 182268576 | 46 |
| chr1 | 182287826 | 182288094 | 182278285 | 182288285 | 182244871 | 182254871 | 33 | 8.6 | 182283369 | 182283869 | 34 | 182268298 | 182268576 | 19 |
| chr1 | 182954688 | 182955252 | 182944361 | 182954361 | 182561825 | 182571825 | 383 | 5.3 | 182283369 | 182283869 | -283 | 182562757 | 182563574 | -4 |
| chr10 | 19907301 | 19907793 | 19907491 | 19917491 | 17759613 | 17769613 | 2148 | 5.6 | 17632194 | 17632694 | -132 | 17727481 | 17728179 | -37 |

| | | | | | | | | | | | | | | | |
|---|---|---|---|---|---|---|---|---|---|---|---|---|---|---|---|
| chr10 | 19907301 | 19907793 | 19902890 | 19912890 | 18544880 | 18554880 | 1358 | 7.4 | 18526194 | 18526694 | -23 | | 17727481 | 17728179 | -822 |
| chr10 | 19907301 | 19907793 | 19903299 | 19913299 | 18850680 | 18860680 | 1053 | 7.5 | 18526194 | 18526694 | -329 | | 17727481 | 17728179 | -1128 |
| chr10 | 19907301 | 19907793 | 19896879 | 19906879 | 18916753 | 18926753 | 980 | 5.3 | 18526194 | 18526694 | -395 | | 17727481 | 17728179 | -1194 |
| chr10 | 19907301 | 19907793 | 19902083 | 19912083 | 19236948 | 19246948 | 665 | 12.9 | 18526194 | 18526694 | -716 | | 17727481 | 17728179 | -1514 |
| chr10 | 19907301 | 19907793 | 19903096 | 19913096 | 19333107 | 19343107 | 570 | 5.3 | 18526194 | 18526694 | -812 | | 17727481 | 17728179 | -1610 |
| chr10 | 19907301 | 19907793 | 19898437 | 19908437 | 19345046 | 19355046 | 553 | 5.0 | 18526194 | 18526694 | -824 | | 17727481 | 17728179 | -1622 |
| chr10 | 19907301 | 19907793 | 19907003 | 19917003 | 19517094 | 19527094 | 390 | 7.9 | 18526194 | 18526694 | -996 | | 17727481 | 17728179 | -1794 |
| chr10 | 19907301 | 19907793 | 19902092 | 19912092 | 19612950 | 19622950 | 289 | 5.6 | 20147694 | 20148194 | 530 | | 21460081 | 21460369 | 1842 |
| chr10 | 19907301 | 19907793 | 19903927 | 19913927 | 19749714 | 19759714 | 154 | 6.4 | 20147694 | 20148194 | 393 | | 21460081 | 21460369 | 1706 |
| chr10 | 19907301 | 19907793 | 19906528 | 19916528 | 19839899 | 19849899 | 67 | 6.9 | 20147694 | 20148194 | 303 | | 21460081 | 21460369 | 1615 |
| chr10 | 19907301 | 19907793 | 19903812 | 19913812 | 19851211 | 19861211 | 53 | 7.5 | 20147694 | 20148194 | 292 | | 21460081 | 21460369 | 1604 |
| chr10 | 19907301 | 19907793 | 19897524 | 19907524 | 19894887 | 19904887 | 3 | 8.0 | 20147694 | 20148194 | 248 | | 21460081 | 21460369 | 1560 |
| chr10 | 23851327 | 23851523 | 23848526 | 23858526 | 23085611 | 23095611 | 763 | 7.2 | 24718194 | 24718694 | 1628 | | 23839016 | 23839261 | 749 |
| chr10 | 83637366 | 83637729 | 83629616 | 83639616 | 83549274 | 83559274 | 80 | 10.0 | 85037755 | 85038255 | 1484 | | 82899837 | 82900212 | -654 |
| chr10 | 83637366 | 83637729 | 83627441 | 83637441 | 83623127 | 83633127 | 4 | 10.2 | 85037755 | 85038255 | 1410 | | 82899837 | 82900212 | -728 |
| chr10 | 86972034 | 86972224 | 86964194 | 86974194 | 86754539 | 86764539 | 210 | 6.8 | 85037755 | 85038255 | -1722 | | 87051736 | 87052123 | 292 |
| chr10 | 86972034 | 86972224 | 86964141 | 86974141 | 86777699 | 86787699 | 186 | 7.3 | 85037755 | 85038255 | -1745 | | 87051736 | 87052123 | 269 |
| chr10 | 86972034 | 86972224 | 86965827 | 86975827 | 86832696 | 86842696 | 133 | 11.6 | 85037755 | 85038255 | -1800 | | 87051736 | 87052123 | 214 |
| chr10 | 86972034 | 86972224 | 86969132 | 86979132 | 86876820 | 86886820 | 92 | 6.8 | 85037755 | 85038255 | -1844 | | 87051736 | 87052123 | 170 |
| chr10 | 86972034 | 86972224 | 86963999 | 86973999 | 86921986 | 86931986 | 42 | 7.0 | 85037755 | 85038255 | -1889 | | 87051736 | 87052123 | 125 |
| chr10 | 86972034 | 86972224 | 86961525 | 86971525 | 86950430 | 86960430 | 11 | 4.7 | 85037755 | 85038255 | -1917 | | 87051736 | 87052123 | 96 |
| chr10 | 95416527 | 95417606 | 95418192 | 95428192 | 94164499 | 94174499 | 1254 | 5.6 | 94800755 | 94801255 | 632 | | 93425781 | 93425969 | -744 |
| chr10 | 95416527 | 95417606 | 95417856 | 95427856 | 94199086 | 94209086 | 1219 | 6.7 | 94800755 | 94801255 | 597 | | 93425781 | 93425969 | -778 |
| chr10 | 95323525 | 95324257 | 95323586 | 95333586 | 94488213 | 94498213 | 835 | 5.6 | 94800755 | 94801255 | 308 | | 94992236 | 94992405 | 499 |
| chr10 | 95416527 | 95417606 | 95416846 | 95426846 | 94801607 | 94811607 | 615 | 5.4 | 94800755 | 94801255 | -6 | | 94992236 | 94992405 | 186 |
| chr10 | 95323525 | 95324257 | 95324372 | 95334372 | 94964887 | 94974887 | 359 | 5.5 | 94800755 | 94801255 | -169 | | 94992236 | 94992405 | 22 |
| chr10 | 95323525 | 95324257 | 95321535 | 95331535 | 94984980 | 94994980 | 337 | 22.6 | 94800755 | 94801255 | -189 | | 94992236 | 94992405 | 2 |
| chr10 | 95416527 | 95417606 | 95416710 | 95426710 | 94986227 | 94996227 | 430 | 27.8 | 94800755 | 94801255 | -190 | | 94992236 | 94992405 | 1 |
| chr10 | 95416527 | 95417606 | 95415724 | 95425724 | 95020371 | 95030371 | 395 | 4.9 | 94800755 | 94801255 | -224 | | 94992236 | 94992405 | -33 |
| chr10 | 95323525 | 95324257 | 95319833 | 95329833 | 95043158 | 95053158 | 277 | 8.6 | 94800755 | 94801255 | -247 | | 94992236 | 94992405 | -56 |
| chr10 | 95416527 | 95417606 | 95411112 | 95421112 | 95059927 | 95069927 | 351 | 7.4 | 94800755 | 94801255 | -264 | | 94992236 | 94992405 | -73 |
| chr10 | 95416527 | 95417606 | 95417690 | 95427690 | 95111871 | 95121871 | 306 | 32.6 | 94800755 | 94801255 | -316 | | 95147335 | 95147885 | 31 |
| chr10 | 95323525 | 95324257 | 95321583 | 95331583 | 95128474 | 95138474 | 193 | 15.3 | 94800755 | 94801255 | -332 | | 95147335 | 95147885 | 14 |
| chr10 | 95416527 | 95417606 | 95411937 | 95421937 | 95164356 | 95174356 | 248 | 10.5 | 94800755 | 94801255 | -368 | | 95147335 | 95147885 | -22 |
| chr10 | 95323525 | 95324257 | 95322679 | 95332679 | 95176871 | 95186871 | 146 | 7.3 | 94800755 | 94801255 | -381 | | 95147335 | 95147885 | -34 |
| chr10 | 95416527 | 95417606 | 95412516 | 95422516 | 95182048 | 95192048 | 230 | 17.0 | 94800755 | 94801255 | -386 | | 95147335 | 95147885 | -39 |
| chr10 | 95323525 | 95324257 | 95322271 | 95332271 | 95226895 | 95236895 | 95 | 8.5 | 95605366 | 95605866 | 374 | | 95147335 | 95147885 | -84 |
| chr10 | 95416527 | 95417606 | 95411790 | 95421790 | 95227783 | 95237783 | 184 | 9.7 | 95605366 | 95605866 | 373 | | 95147335 | 95147885 | -85 |
| chr10 | 95323525 | 95324257 | 95323267 | 95333267 | 95242429 | 95252429 | 81 | 6.8 | 95605366 | 95605866 | 358 | | 95147335 | 95147885 | -100 |
| chr10 | 95323525 | 95324257 | 95321633 | 95331633 | 95274350 | 95284350 | 47 | 6.1 | 95605366 | 95605866 | 326 | | 95147335 | 95147885 | -132 |
| chr10 | 95416527 | 95417606 | 95417322 | 95427322 | 95304849 | 95314849 | 112 | 17.7 | 95605366 | 95605866 | 296 | | 95438907 | 95439296 | 129 |
| chr10 | 95323525 | 95324257 | 95325171 | 95335171 | 95323854 | 95333854 | 1 | 10.4 | 95605366 | 95605866 | 277 | | 95438907 | 95439296 | 110 |
| chr10 | 95416527 | 95417606 | 95412270 | 95422270 | 95338557 | 95348557 | 74 | 8.0 | 95605366 | 95605866 | 262 | | 95438907 | 95439296 | 96 |
| chr10 | 121310983 | 121311350 | 121307973 | 121317973 | 120909266 | 120919266 | 399 | 8.4 | 120876944 | 120877444 | -37 | | 122051930 | 122052126 | 1138 |
| chr10 | 121310983 | 121311350 | 121308477 | 121318477 | 120922654 | 120932654 | 386 | 4.9 | 120876944 | 120877444 | -50 | | 122051930 | 122052126 | 1124 |
| chr10 | 121351326 | 121351598 | 121351048 | 121361048 | 120928273 | 120938273 | 423 | 5.3 | 120876944 | 120877444 | -56 | | 122051930 | 122052126 | 1119 |
| chr10 | 121310983 | 121311350 | 121307865 | 121317865 | 120948393 | 120958393 | 359 | 4.6 | 120876944 | 120877444 | -76 | | 122051930 | 122052126 | 1099 |
| chr10 | 121310983 | 121311350 | 121307594 | 121317594 | 120972853 | 120982853 | 335 | 9.1 | 120876944 | 120877444 | -101 | | 122051930 | 122052126 | 1074 |

| | | | | | | | | | | | | | | |
|---|---|---|---|---|---|---|---|---|---|---|---|---|---|---|
| chr10 | 121310983 | 121311350 | 121304277 | 121314277 | 120996849 | 121006849 | 307 | 9.1 | 120876944 | 120877444 | -125 | 122051930 | 122052126 | 1050 |
| chr10 | 121310983 | 121311350 | 121304196 | 121314196 | 121027740 | 121037740 | 276 | 5.1 | 121918944 | 121919444 | 886 | 122051930 | 122052126 | 1019 |
| chr10 | 121310983 | 121311350 | 121307416 | 121317416 | 121107391 | 121117391 | 200 | 5.5 | 121918944 | 121919444 | 807 | 122051930 | 122052126 | 940 |
| chr10 | 128626258 | 128626577 | 128615259 | 128625259 | 128568626 | 128578626 | 47 | 15.7 | 126770944 | 126771444 | -1802 | 128590154 | 128590650 | 17 |
| chr10 | 128626258 | 128626577 | 128615508 | 128625508 | 128613056 | 128623056 | 2 | 8.3 | 126770944 | 126771444 | -1847 | 128590154 | 128590650 | -28 |
| chr11 | 59787286 | 59787779 | 59778044 | 59788044 | 59221492 | 59231492 | 557 | 5.6 | 59067498 | 59067998 | -159 | 58927776 | 58928598 | -298 |
| chr11 | 59787286 | 59787779 | 59786857 | 59796857 | 59319616 | 59329616 | 467 | 5.6 | 59067498 | 59067998 | -257 | 58927776 | 58928598 | -396 |
| chr11 | 59787286 | 59787779 | 59785963 | 59795963 | 59561483 | 59571483 | 224 | 5.0 | 59067498 | 59067998 | -499 | 60048716 | 60049133 | 482 |
| chr11 | 59787286 | 59787779 | 59788059 | 59798059 | 59674527 | 59684527 | 114 | 4.6 | 59067498 | 59067998 | -612 | 60048716 | 60049133 | 369 |
| chr11 | 59787286 | 59787779 | 59776552 | 59786552 | 59692314 | 59702314 | 84 | 4.8 | 59067498 | 59067998 | -630 | 60048716 | 60049133 | 352 |
| chr11 | 59787286 | 59787779 | 59786129 | 59796129 | 59769867 | 59779867 | 16 | 6.9 | 59067498 | 59067998 | -707 | 60048716 | 60049133 | 274 |
| chr11 | 94392549 | 94393110 | 94384219 | 94394219 | 94316158 | 94326158 | 68 | 7.7 | 95257686 | 95258186 | 937 | 94273324 | 94273803 | -48 |
| chr11 | 94392549 | 94393110 | 94382038 | 94392038 | 94336433 | 94346433 | 46 | 16.5 | 95257686 | 95258186 | 917 | 94273324 | 94273803 | -68 |
| chr11 | 94392549 | 94393110 | 94381648 | 94391648 | 94366261 | 94376261 | 15 | 18.1 | 95257686 | 95258186 | 887 | 94273324 | 94273803 | -98 |
| chr12 | 33315341 | 33315706 | 33307046 | 33317046 | 32630744 | 32640744 | 676 | 5.2 | 32893135 | 32893635 | 258 | 32681977 | 32682305 | 46 |
| chr12 | 33315341 | 33315706 | 33308686 | 33318686 | 32702092 | 32712092 | 607 | 8.6 | 32893135 | 32893635 | 186 | 32681977 | 32682305 | -25 |
| chr12 | 33315341 | 33315706 | 33312839 | 33322839 | 32935595 | 32945595 | 377 | 11.5 | 32893135 | 32893635 | -47 | 32910156 | 32910353 | -30 |
| chr12 | 33315341 | 33315706 | 33313773 | 33323773 | 32960574 | 32970574 | 353 | 10.1 | 32893135 | 32893635 | -72 | 32910156 | 32910353 | -55 |
| chr12 | 33315341 | 33315706 | 33305962 | 33315962 | 33025445 | 33035445 | 281 | 7.4 | 32893135 | 32893635 | -137 | 32910156 | 32910353 | -120 |
| chr12 | 33315341 | 33315706 | 33314151 | 33324151 | 33289557 | 33299557 | 25 | 5.5 | 33382635 | 33383135 | 88 | 32910156 | 32910353 | -384 |
| chr12 | 59130264 | 59130503 | 59128818 | 59138818 | 59052288 | 59062288 | 77 | 11.3 | 58925013 | 58925513 | -132 | 59195154 | 59195561 | 138 |
| chr12 | 59130264 | 59130503 | 59128797 | 59138797 | 59064059 | 59074059 | 65 | 21.9 | 58925013 | 58925513 | -144 | 59195154 | 59195561 | 126 |
| chr12 | 83987732 | 83988225 | 83987762 | 83997762 | 83638728 | 83648728 | 349 | 5.2 | 82179013 | 82179513 | -1464 | 83923227 | 83923460 | 280 |
| chr12 | 84114214 | 84114427 | 84112629 | 84122629 | 83683849 | 83693849 | 429 | 4.9 | 82179013 | 82179513 | -1510 | 83923227 | 83923460 | 234 |
| chr12 | 83987732 | 83988225 | 83988567 | 83998567 | 83688216 | 83698216 | 300 | 5.4 | 82179013 | 82179513 | -1514 | 83923227 | 83923460 | 230 |
| chr12 | 83987732 | 83988225 | 83979542 | 83989542 | 83724033 | 83734033 | 256 | 5.7 | 82179013 | 82179513 | -1550 | 83923227 | 83923460 | 194 |
| chr12 | 83987732 | 83988225 | 83980253 | 83990253 | 83818279 | 83828279 | 162 | 6.7 | 85336050 | 85336550 | 1513 | 83923227 | 83923460 | 100 |
| chr12 | 84114214 | 84114427 | 84110386 | 84120386 | 83965362 | 83975362 | 145 | 9.8 | 85336050 | 85336550 | 1366 | 83999362 | 83999585 | 29 |
| chr12 | 83987732 | 83988225 | 83980316 | 83990316 | 83979260 | 83989260 | 1 | 9.8 | 85336050 | 85336550 | 1352 | 83999362 | 83999585 | 15 |
| chr12 | 84114214 | 84114427 | 84110280 | 84120280 | 83982226 | 83992226 | 128 | 7.6 | 85336050 | 85336550 | 1349 | 83999362 | 83999585 | 12 |
| chr12 | 84114214 | 84114427 | 84111012 | 84121012 | 84092059 | 84102059 | 19 | 6.7 | 85336050 | 85336550 | 1239 | 83999362 | 83999585 | -98 |
| chr12 | 103657056 | 103657505 | 103646393 | 103656393 | 103528497 | 103538497 | 118 | 7.5 | 103316290 | 103316790 | -217 | 103659221 | 103659518 | 126 |
| chr12 | 103657056 | 103657505 | 103646984 | 103656984 | 103543626 | 103553626 | 103 | 7.5 | 103316290 | 103316790 | -232 | 103659221 | 103659518 | 111 |
| chr12 | 103657056 | 103657505 | 103647376 | 103657376 | 103573937 | 103583937 | 73 | 9.3 | 103316290 | 103316790 | -262 | 103659221 | 103659518 | 80 |
| chr12 | 103657056 | 103657505 | 103649912 | 103659912 | 103629570 | 103639570 | 20 | 7.9 | 103316290 | 103316790 | -318 | 103659221 | 103659518 | 25 |
| chr12 | 103657056 | 103657505 | 103649561 | 103659561 | 103647054 | 103657054 | 3 | 4.7 | 103316290 | 103316790 | -336 | 103659221 | 103659518 | 7 |
| chr12 | 111538597 | 111538916 | 111528789 | 111538789 | 110896642 | 110906642 | 632 | 7.2 | 111149789 | 111150289 | 248 | 110703033 | 110703403 | -198 |
| chr12 | 111538597 | 111538916 | 111530370 | 111540370 | 110917540 | 110927540 | 613 | 4.9 | 111149789 | 111150289 | 227 | 110703033 | 110703403 | -219 |
| chr12 | 111538597 | 111538916 | 111529410 | 111539410 | 110974508 | 110984508 | 555 | 5.6 | 111149789 | 111150289 | 171 | 110703033 | 110703403 | -276 |
| chr12 | 111538597 | 111538916 | 111536602 | 111546602 | 111313510 | 111323510 | 223 | 5.4 | 111357289 | 111357789 | 39 | 111505014 | 111505242 | 187 |
| chr12 | 111538597 | 111538916 | 111532043 | 111542043 | 111349206 | 111359206 | 183 | 5.9 | 111357289 | 111357789 | 3 | 111505014 | 111505242 | 151 |
| chr12 | 111538597 | 111538916 | 111531419 | 111541419 | 111373498 | 111383498 | 158 | 7.7 | 111357289 | 111357789 | -21 | 111505014 | 111505242 | 127 |
| chr12 | 111538597 | 111538916 | 111527710 | 111537710 | 111417551 | 111427551 | 110 | 4.7 | 111357289 | 111357789 | -65 | 111505014 | 111505242 | 83 |
| chr13 | 24937649 | 24938224 | 24927270 | 24937270 | 24482492 | 24492492 | 445 | 4.7 | 24296131 | 24296631 | -191 | 23730518 | 23730867 | -757 |
| chr13 | 24937649 | 24938224 | 24931679 | 24941679 | 24792755 | 24802755 | 139 | 5.8 | 24296131 | 24296631 | -501 | 23730518 | 23730867 | -1067 |
| chr13 | 24937649 | 24938224 | 24928872 | 24938872 | 24842660 | 24852660 | 86 | 6.7 | 24296131 | 24296631 | -551 | 23730518 | 23730867 | -1117 |
| chr13 | 38151862 | 38152358 | 38148959 | 38158959 | 38016517 | 38026517 | 132 | 4.6 | 38185131 | 38185631 | 164 | 37732302 | 37732574 | -289 |
| chr13 | 38151862 | 38152358 | 38142191 | 38152191 | 38053386 | 38063386 | 89 | 5.9 | 38185131 | 38185631 | 127 | 37732302 | 37732574 | -326 |

| | | | | | | | | | | | | | | |
|---|---|---|---|---|---|---|---|---|---|---|---|---|---|---|
| chr13 | 38151862 | 38152358 | 38146586 | 38156586 | 38101977 | 38111977 | 45 | 7.4 | 38185131 | 38185631 | 78 | 37732302 | 37732574 | -375 |
| chr13 | 38151862 | 38152358 | 38141495 | 38151495 | 38134835 | 38144835 | 7 | 9.1 | 38185131 | 38185631 | 46 | 37732302 | 37732574 | -407 |
| chr13 | 60601749 | 60601948 | 60592610 | 60602610 | 59824221 | 59834221 | 768 | 5.6 | 59856139 | 59856639 | 27 | 59840856 | 59841080 | 12 |
| chr13 | 60601749 | 60601948 | 60593932 | 60603932 | 60123339 | 60133339 | 471 | 5.8 | 60056139 | 60056639 | -72 | 59840856 | 59841080 | -287 |
| chr13 | 60601749 | 60601948 | 60594948 | 60604948 | 60412618 | 60422618 | 182 | 6.1 | 60427639 | 60428139 | 10 | 59840856 | 59841080 | -577 |
| chr13 | 60601749 | 60601948 | 60599075 | 60609075 | 60473331 | 60483331 | 126 | 23.9 | 60427639 | 60428139 | -50 | 59840856 | 59841080 | -637 |
| chr13 | 60601749 | 60601948 | 60598510 | 60608510 | 60519721 | 60529721 | 79 | 9.3 | 60427639 | 60428139 | -97 | 59840856 | 59841080 | -684 |
| chr13 | 109260331 | 109260761 | 109260752 | 109270752 | 108447356 | 108457356 | 813 | 12.0 | 108271806 | 108272306 | -180 | 107666771 | 107667124 | -785 |
| chr14 | 30008745 | 30009127 | 29998155 | 30008155 | 29968323 | 29978323 | 30 | 5.8 | 29863314 | 29863814 | -110 | 30907762 | 30908148 | 935 |
| chr14 | 30008745 | 30009127 | 29997781 | 30007781 | 29996601 | 30006601 | 1 | 19.6 | 29863314 | 29863814 | -138 | 30907762 | 30908148 | 906 |
| chr15 | 3979416 | 3979842 | 3969153 | 3979153 | 3182784 | 3192784 | 786 | 8.1 | 5263000 | 5263500 | 2075 | 3228960 | 3229299 | 41 |
| chr15 | 3979416 | 3979842 | 3970686 | 3980686 | 3233507 | 3243507 | 737 | 9.0 | 5263000 | 5263500 | 2025 | 3232534 | 3232810 | -6 |
| chr15 | 3979416 | 3979842 | 3974630 | 3984630 | 3267179 | 3277179 | 707 | 6.6 | 5263000 | 5263500 | 1991 | 3251000 | 3251286 | -21 |
| chr15 | 3979416 | 3979842 | 3975048 | 3985048 | 3288424 | 3298424 | 687 | 9.3 | 5263000 | 5263500 | 1970 | 3251000 | 3251286 | -42 |
| chr15 | 3979416 | 3979842 | 3970305 | 3980305 | 3307084 | 3317084 | 663 | 13.2 | 5263000 | 5263500 | 1951 | 3251000 | 3251286 | -61 |
| chr15 | 3979416 | 3979842 | 3971292 | 3981292 | 3332776 | 3342776 | 639 | 4.7 | 5263000 | 5263500 | 1925 | 3251000 | 3251286 | -87 |
| chr15 | 3979416 | 3979842 | 3970010 | 3980010 | 3351353 | 3361353 | 619 | 9.4 | 5263000 | 5263500 | 1907 | 3251000 | 3251286 | -105 |
| chr15 | 3979416 | 3979842 | 3971102 | 3981102 | 3394403 | 3404403 | 577 | 5.5 | 5263000 | 5263500 | 1864 | 3251000 | 3251286 | -148 |
| chr15 | 3979416 | 3979842 | 3970968 | 3980968 | 3447536 | 3457536 | 523 | 5.3 | 5263000 | 5263500 | 1811 | 3251000 | 3251286 | -201 |
| chr15 | 3979416 | 3979842 | 3975266 | 3985266 | 3567318 | 3577318 | 408 | 5.6 | 5263000 | 5263500 | 1691 | 3251000 | 3251286 | -321 |
| chr15 | 39857001 | 39857560 | 39848602 | 39858602 | 39288235 | 39298235 | 560 | 5.6 | 39937454 | 39937954 | 644 | 39859903 | 39860282 | 567 |
| chr15 | 39857001 | 39857560 | 39857250 | 39867250 | 39322779 | 39332779 | 534 | 8.6 | 39937454 | 39937954 | 610 | 39859903 | 39860282 | 532 |
| chr15 | 39857001 | 39857560 | 39851018 | 39861018 | 39361849 | 39371849 | 489 | 12.4 | 39937454 | 39937954 | 571 | 39859903 | 39860282 | 493 |
| chr15 | 39857001 | 39857560 | 39848736 | 39858736 | 39514793 | 39524793 | 334 | 5.7 | 39937454 | 39937954 | 418 | 39859903 | 39860282 | 340 |
| chr15 | 39857001 | 39857560 | 39856859 | 39866859 | 39809564 | 39819564 | 47 | 5.5 | 39937454 | 39937954 | 123 | 39859903 | 39860282 | 46 |
| chr15 | 98609420 | 98610224 | 98609101 | 98619101 | 98506757 | 98516757 | 102 | 8.8 | 98226069 | 98226569 | -285 | 98678044 | 98678269 | 166 |
| chr15 | 98609420 | 98610224 | 98610697 | 98620697 | 98540204 | 98550204 | 70 | 8.7 | 98226069 | 98226569 | -319 | 98678044 | 98678269 | 133 |
| chr15 | 98609420 | 98610224 | 98609964 | 98619964 | 98578288 | 98588288 | 32 | 60.9 | 98226069 | 98226569 | -357 | 98678044 | 98678269 | 95 |
| chr15 | 98609420 | 98610224 | 98599890 | 98609890 | 98599291 | 98609291 | 1 | 20.2 | 98226069 | 98226569 | -378 | 98678044 | 98678269 | 74 |
| chr15 | 102150305 | 102151117 | 102146594 | 102156594 | 102113536 | 102123536 | 33 | 8.5 | 101548569 | 101549069 | -570 | 102026505 | 102026834 | -92 |
| chr16 | 23029015 | 23029479 | 23024165 | 23034165 | 22242669 | 22252669 | 781 | 4.6 | 22621427 | 22621927 | 374 | 22899793 | 22900523 | 652 |
| chr16 | 23029015 | 23029479 | 23030032 | 23040032 | 22873874 | 22883874 | 156 | 17.7 | 22621427 | 22621927 | -257 | 22899793 | 22900523 | 21 |
| chr16 | 23029015 | 23029479 | 23018762 | 23028762 | 22899737 | 22909737 | 119 | 16.0 | 22621427 | 22621927 | -283 | 22899793 | 22900523 | -5 |
| chr16 | 23029015 | 23029479 | 23029135 | 23039135 | 23028318 | 23038318 | 1 | 14.5 | 22621427 | 22621927 | -412 | 23037745 | 23038307 | 5 |
| chr16 | 23225862 | 23226375 | 23222668 | 23232668 | 23074876 | 23084876 | 148 | 5.1 | 22621427 | 22621927 | -458 | 23039274 | 23039610 | -40 |
| chr16 | 23225862 | 23226375 | 23223025 | 23233025 | 23106509 | 23116509 | 117 | 6.3 | 22621427 | 22621927 | -490 | 23039274 | 23039610 | -72 |
| chr16 | 23225862 | 23226375 | 23222817 | 23232817 | 23125237 | 23135237 | 98 | 5.3 | 22621427 | 22621927 | -509 | 23039274 | 23039610 | -91 |
| chr16 | 23191668 | 23192220 | 23189789 | 23199789 | 23143231 | 23153231 | 47 | 8.9 | 22621427 | 22621927 | -527 | 23039274 | 23039610 | -109 |
| chr16 | 23191668 | 23192220 | 23180812 | 23190812 | 23160239 | 23170239 | 21 | 7.2 | 22621427 | 22621927 | -544 | 23039274 | 23039610 | -126 |
| chr16 | 24090027 | 24090422 | 24079470 | 24089470 | 23988648 | 23998648 | 91 | 32.6 | 22621427 | 22621927 | -1372 | 24061562 | 24061888 | 68 |
| chr16 | 24090027 | 24090422 | 24081275 | 24091275 | 24006509 | 24016509 | 75 | 10.2 | 22621427 | 22621927 | -1390 | 24061562 | 24061888 | 50 |
| chr16 | 24090027 | 24090422 | 24081407 | 24091407 | 24040750 | 24050750 | 41 | 7.8 | 22621427 | 22621927 | -1424 | 24061562 | 24061888 | 16 |
| chr16 | 24090027 | 24090422 | 24079086 | 24089086 | 24079086 | 24089086 | 0 | 8.6 | 22621427 | 22621927 | -1462 | 24061562 | 24061888 | -22 |
| chr16 | 38432879 | 38433347 | 38431463 | 38441463 | 37003631 | 37013631 | 1428 | 5.6 | 35183414 | 35183914 | -1825 | 37684790 | 37685150 | 676 |
| chr16 | 38432879 | 38433347 | 38429912 | 38439912 | 37989179 | 37999179 | 441 | 9.1 | 39121387 | 39121887 | 1127 | 37684790 | 37685150 | -309 |
| chr16 | 38432879 | 38433347 | 38423876 | 38433876 | 38219390 | 38229390 | 204 | 5.1 | 39121387 | 39121887 | 897 | 37684790 | 37685150 | -539 |
| chr16 | 38432879 | 38433347 | 38429044 | 38439044 | 38238247 | 38248247 | 191 | 4.6 | 39121387 | 39121887 | 878 | 37684790 | 37685150 | -558 |
| chr16 | 38432879 | 38433347 | 38428801 | 38438801 | 38283313 | 38293313 | 145 | 5.5 | 39121387 | 39121887 | 833 | 37684790 | 37685150 | -603 |

| | | | | | | | | | | | | | | |
|---|---|---|---|---|---|---|---|---|---|---|---|---|---|---|
| chr16 | 38432879 | 38433347 | 38432021 | 38442021 | 38347601 | 38357601 | 84 | 8.5 | 39121387 | 39121887 | 769 | 37684790 | 37685150 | -668 |
| chr16 | 38432879 | 38433347 | 38428673 | 38438673 | 38359720 | 38369720 | 69 | 8.2 | 39121387 | 39121887 | 757 | 37684790 | 37685150 | -680 |
| chr16 | 38432879 | 38433347 | 38422126 | 38432126 | 38376801 | 38386801 | 45 | 5.5 | 39121387 | 39121887 | 740 | 37684790 | 37685150 | -697 |
| chr16 | 38432879 | 38433347 | 38422298 | 38432298 | 38415146 | 38425146 | 7 | 7.6 | 39121387 | 39121887 | 701 | 37684790 | 37685150 | -735 |
| chr16 | 45945195 | 45945671 | 45939216 | 45949216 | 44378899 | 44388899 | 1560 | 5.3 | 44765387 | 44765887 | 382 | 44172273 | 44172469 | -212 |
| chr16 | 45945195 | 45945671 | 45945965 | 45955965 | 45769611 | 45779611 | 176 | 17.5 | 45579387 | 45579887 | -195 | 46010984 | 46011188 | 236 |
| chr16 | 45945195 | 45945671 | 45938286 | 45948286 | 45799391 | 45809391 | 139 | 7.6 | 45579387 | 45579887 | -225 | 45608512 | 45608960 | -196 |
| chr16 | 45945195 | 45945671 | 45945430 | 45955430 | 45835127 | 45845127 | 110 | 22.0 | 45579387 | 45579887 | -260 | 46010984 | 46011188 | 171 |
| chr16 | 45945195 | 45945671 | 45938280 | 45948280 | 45864773 | 45874773 | 74 | 11.6 | 45579387 | 45579887 | -290 | 46010984 | 46011188 | 141 |
| chr16 | 45945195 | 45945671 | 45938096 | 45948096 | 45933307 | 45943307 | 5 | 7.9 | 45579387 | 45579887 | -359 | 46010984 | 46011188 | 73 |
| chr16 | 97850958 | 97851358 | 97849238 | 97859238 | 97802050 | 97812050 | 47 | 6.2 | 96903393 | 96903893 | -903 | 97778817 | 97779000 | -28 |
| chr17 | 33781003 | 33781884 | 33780855 | 33790855 | 33719269 | 33729269 | 62 | 4.8 | 34130055 | 34130555 | 406 | 33897209 | 33897414 | 173 |
| chr17 | 35126241 | 35126660 | 35119081 | 35129081 | 34827420 | 34837420 | 292 | 7.0 | 34130055 | 34130555 | -702 | 34952688 | 34953036 | 120 |
| chr17 | 35126241 | 35126660 | 35120632 | 35130632 | 34927873 | 34937873 | 193 | 6.5 | 35554055 | 35554555 | 621 | 34952688 | 34953036 | 20 |
| chr17 | 35126241 | 35126660 | 35117503 | 35127503 | 35107870 | 35117870 | 10 | 12.7 | 35554055 | 35554555 | 441 | 34952688 | 34953036 | -160 |
| chr17 | 47437080 | 47437695 | 47436657 | 47446657 | 47382729 | 47392729 | 54 | 4.8 | 47512551 | 47513051 | 125 | 47472837 | 47473411 | 85 |
| chr17 | 53566237 | 53566760 | 53558843 | 53568843 | 52566546 | 52576546 | 992 | 5.2 | 48882175 | 48882675 | -3689 | 50508610 | 50508938 | -2063 |
| chr17 | 53566237 | 53566760 | 53555493 | 53565493 | 53488789 | 53498789 | 67 | 5.5 | 48882175 | 48882675 | -4611 | 55931674 | 55932329 | 2438 |
| chr17 | 53566237 | 53566760 | 53557835 | 53567835 | 53523072 | 53533072 | 35 | 6.0 | 48882175 | 48882675 | -4646 | 55931674 | 55932329 | 2404 |
| chr17 | 53566237 | 53566760 | 53560086 | 53570086 | 53556112 | 53566112 | 4 | 10.7 | 48882175 | 48882675 | -4679 | 55931674 | 55932329 | 2371 |
| chr18 | 12869110 | 12869537 | 12865807 | 12875807 | 12246085 | 12256085 | 620 | 5.0 | 12211991 | 12212491 | -39 | 10885882 | 10886060 | -1365 |
| chr18 | 12869110 | 12869537 | 12866286 | 12876286 | 12517233 | 12527233 | 349 | 5.4 | 12211991 | 12212491 | -310 | 10885882 | 10886060 | -1636 |
| chr18 | 12869110 | 12869537 | 12864760 | 12874760 | 12747403 | 12757403 | 117 | 18.5 | 12211991 | 12212491 | -540 | 10885882 | 10886060 | -1866 |
| chr18 | 12869110 | 12869537 | 12864923 | 12874923 | 12785263 | 12795263 | 80 | 20.3 | 12211991 | 12212491 | -578 | 10885882 | 10886060 | -1904 |
| chr18 | 12869110 | 12869537 | 12860475 | 12870475 | 12823953 | 12833953 | 37 | 13.3 | 12211991 | 12212491 | -617 | 10885882 | 10886060 | -1943 |
| chr18 | 34776752 | 34777040 | 34770552 | 34780552 | 32853492 | 32863492 | 1917 | 5.6 | 32406846 | 32407346 | -451 | 33378023 | 33378487 | 520 |
| chr18 | 34776752 | 34777040 | 34771485 | 34781485 | 34493310 | 34503310 | 278 | 5.2 | 35979846 | 35980346 | 1482 | 34850170 | 34850387 | 352 |
| chr18 | 34776752 | 34777040 | 34771723 | 34781723 | 34603063 | 34613063 | 169 | 6.6 | 35979846 | 35980346 | 1372 | 34850170 | 34850387 | 242 |
| chr18 | 34776752 | 34777040 | 34771846 | 34781846 | 34670630 | 34680630 | 101 | 5.1 | 35979846 | 35980346 | 1304 | 34850170 | 34850387 | 175 |
| chr18 | 34776752 | 34777040 | 34771591 | 34781591 | 34759196 | 34769196 | 12 | 13.0 | 35979846 | 35980346 | 1216 | 34850170 | 34850387 | 86 |
| chr18 | 46728256 | 46728682 | 46720201 | 46730201 | 45560698 | 45570698 | 1160 | 5.6 | 46273846 | 46274346 | 708 | 46335644 | 46335886 | 770 |
| chr18 | 46728256 | 46728682 | 46718680 | 46728680 | 46295896 | 46305896 | 423 | 6.1 | 46273846 | 46274346 | -27 | 46335644 | 46335886 | 35 |
| chr18 | 46728256 | 46728682 | 46724908 | 46734908 | 46333350 | 46343350 | 392 | 12.5 | 46273846 | 46274346 | -64 | 46335644 | 46335886 | -3 |
| chr18 | 46728256 | 46728682 | 46720239 | 46730239 | 46375760 | 46385760 | 344 | 8.5 | 46273846 | 46274346 | -107 | 46335644 | 46335886 | -45 |
| chr18 | 46728256 | 46728682 | 46728624 | 46738624 | 46484189 | 46494189 | 244 | 5.6 | 46273846 | 46274346 | -215 | 46629249 | 46629656 | 140 |
| chr18 | 46728256 | 46728682 | 46725206 | 46735206 | 46594576 | 46604576 | 131 | 64.1 | 46714846 | 46715346 | 116 | 46629249 | 46629656 | 30 |
| chr18 | 46728256 | 46728682 | 46721671 | 46731671 | 46621670 | 46631670 | 100 | 45.1 | 46714846 | 46715346 | 88 | 46629249 | 46629656 | 3 |
| chr18 | 46728256 | 46728682 | 46720460 | 46730460 | 46718804 | 46728804 | 2 | 25.9 | 46714846 | 46715346 | -9 | 46648659 | 46648961 | -75 |
| chr19 | 8989186 | 8989627 | 8980229 | 8990229 | 8837329 | 8847329 | 143 | 6.7 | 8712510 | 8713010 | -130 | 8630793 | 8631473 | -211 |
| chr19 | 8989186 | 8989627 | 8986245 | 8996245 | 8851769 | 8861769 | 134 | 8.4 | 8712510 | 8713010 | -144 | 8630793 | 8631473 | -226 |
| chr19 | 29047829 | 29048196 | 29047286 | 29057286 | 28962575 | 28972575 | 85 | 17.8 | 28045510 | 28046010 | -922 | 29049498 | 29049983 | 82 |
| chr19 | 29047829 | 29048196 | 29046485 | 29056485 | 28984040 | 28994040 | 62 | 10.2 | 28045510 | 28046010 | -943 | 29049498 | 29049983 | 61 |
| chr19 | 29047829 | 29048196 | 29045286 | 29055286 | 29013469 | 29023469 | 32 | 7.1 | 28045510 | 28046010 | -973 | 29049498 | 29049983 | 31 |
| chr19 | 32595581 | 32596045 | 32588737 | 32598737 | 32459638 | 32469638 | 129 | 4.6 | 36477010 | 36477510 | 4013 | 32532261 | 32532627 | 68 |
| chr19 | 32595581 | 32596045 | 32587107 | 32597107 | 32484909 | 32494909 | 102 | 6.1 | 36477010 | 36477510 | 3987 | 32532261 | 32532627 | 43 |
| chr19 | 32595581 | 32596045 | 32587452 | 32597452 | 32515532 | 32525532 | 72 | 8.1 | 36477010 | 36477510 | 3957 | 32532261 | 32532627 | 12 |
| chr19 | 32595581 | 32596045 | 32588093 | 32598093 | 32548939 | 32558939 | 39 | 14.2 | 36477010 | 36477510 | 3923 | 32573327 | 32573554 | 20 |
| chr19 | 32595581 | 32596045 | 32588289 | 32598289 | 32573555 | 32583555 | 15 | 8.5 | 36477010 | 36477510 | 3899 | 32573327 | 32573554 | -5 |

| | | | | | | | | | | | | | | |
|---|---|---|---|---|---|---|---|---|---|---|---|---|---|---|
| chr19 | 43940136 | 43940555 | 43940469 | 43950469 | 41552716 | 41562716 | 2388 | 7.4 | 40994010 | 40994510 | -563 | 41838940 | 41839215 | 281 |
| chr19 | 43940136 | 43940555 | 43939755 | 43949755 | 42579183 | 42589183 | 1361 | 7.4 | 42326010 | 42326510 | -258 | 42510326 | 42510775 | -74 |
| chr19 | 43940136 | 43940555 | 43938146 | 43948146 | 43709137 | 43719137 | 229 | 6.8 | 43016010 | 43016510 | -698 | 43999185 | 43999474 | 285 |
| chr19 | 43940136 | 43940555 | 43940697 | 43950697 | 43730397 | 43740397 | 210 | 5.8 | 43016010 | 43016510 | -719 | 43999185 | 43999474 | 264 |
| chr19 | 43940136 | 43940555 | 43935229 | 43945229 | 43755378 | 43765378 | 180 | 4.6 | 43016010 | 43016510 | -744 | 43999185 | 43999474 | 239 |
| chr19 | 43940136 | 43940555 | 43938211 | 43948211 | 43770961 | 43780961 | 167 | 6.3 | 43016010 | 43016510 | -760 | 43999185 | 43999474 | 223 |
| chr19 | 43940136 | 43940555 | 43940372 | 43950372 | 43840708 | 43850708 | 100 | 34.7 | 43016010 | 43016510 | -829 | 43999185 | 43999474 | 154 |
| chr19 | 43940136 | 43940555 | 43937617 | 43947617 | 43858183 | 43868183 | 79 | 14.5 | 43016010 | 43016510 | -847 | 43999185 | 43999474 | 136 |
| chr2 | 4995542 | 4996065 | 4994861 | 5004861 | 3623301 | 3633301 | 1372 | 5.6 | 4833954 | 4834454 | 1206 | 3799924 | 3800274 | 172 |
| chr2 | 4995542 | 4996065 | 4993615 | 5003615 | 4556580 | 4566580 | 437 | 5.0 | 4833954 | 4834454 | 273 | 4987626 | 4988100 | 426 |
| chr2 | 4995542 | 4996065 | 4989582 | 4999582 | 4583571 | 4593571 | 406 | 6.0 | 4833954 | 4834454 | 246 | 4987626 | 4988100 | 399 |
| chr2 | 4995542 | 4996065 | 4985231 | 4995231 | 4866184 | 4876184 | 119 | 7.6 | 4833954 | 4834454 | -37 | 4987626 | 4988100 | 117 |
| chr2 | 4995542 | 4996065 | 4992511 | 5002511 | 4912888 | 4922888 | 80 | 24.7 | 4833954 | 4834454 | -84 | 4987626 | 4988100 | 70 |
| chr2 | 4995542 | 4996065 | 4993951 | 5003951 | 4993020 | 5003020 | 1 | 5.1 | 4833954 | 4834454 | -164 | 4987626 | 4988100 | -10 |
| chr2 | 18063780 | 18064209 | 18059715 | 18069715 | 17420016 | 17430016 | 640 | 5.0 | 17899373 | 17899873 | 475 | 18766812 | 18767451 | 1342 |
| chr2 | 29869009 | 29869414 | 29865798 | 29875798 | 28754940 | 28764940 | 1111 | 5.6 | 27449980 | 27450480 | -1310 | 28629989 | 28630408 | -130 |
| chr2 | 29869009 | 29869414 | 29869239 | 29879239 | 29125888 | 29135888 | 743 | 7.4 | 27449980 | 27450480 | -1681 | 28629989 | 28630408 | -501 |
| chr2 | 29869009 | 29869414 | 29867358 | 29877358 | 29154023 | 29164023 | 713 | 5.1 | 27449980 | 27450480 | -1709 | 28629989 | 28630408 | -529 |
| chr2 | 29869009 | 29869414 | 29860264 | 29870264 | 29627531 | 29637531 | 233 | 4.7 | 31082480 | 31082980 | 1450 | 29817775 | 29817953 | 185 |
| chr2 | 29869009 | 29869414 | 29860558 | 29870558 | 29815414 | 29825414 | 45 | 5.9 | 31082480 | 31082980 | 1262 | 29817775 | 29817953 | -3 |
| chr2 | 29869009 | 29869414 | 29866272 | 29876272 | 29829966 | 29839966 | 36 | 9.8 | 31082480 | 31082980 | 1248 | 29817775 | 29817953 | -17 |
| chr2 | 29869009 | 29869414 | 29868745 | 29878745 | 29842461 | 29852461 | 26 | 10.5 | 31082480 | 31082980 | 1235 | 29817775 | 29817953 | -30 |
| chr2 | 29869009 | 29869414 | 29866808 | 29876808 | 29866074 | 29876074 | 1 | 6.7 | 31082480 | 31082980 | 1212 | 29817775 | 29817953 | -53 |
| chr2 | 48814020 | 48814435 | 48814256 | 48824256 | 47864442 | 47874442 | 950 | 6.7 | 51640480 | 51640980 | 3771 | 43470001 | 43470181 | -4399 |
| chr2 | 48814020 | 48814435 | 48814054 | 48824054 | 47927567 | 47937567 | 886 | 5.6 | 51640480 | 51640980 | 3708 | 43470001 | 43470181 | -4462 |
| chr2 | 48814020 | 48814435 | 48808531 | 48818531 | 47945325 | 47955325 | 863 | 5.6 | 51640480 | 51640980 | 3690 | 43470001 | 43470181 | -4480 |
| chr2 | 48814020 | 48814435 | 48813401 | 48823401 | 47993132 | 48003132 | 820 | 20.9 | 51640480 | 51640980 | 3643 | 43470001 | 43470181 | -4528 |
| chr2 | 48814020 | 48814435 | 48812510 | 48822510 | 48034209 | 48044209 | 778 | 7.4 | 51640480 | 51640980 | 3602 | 43470001 | 43470181 | -4569 |
| chr2 | 48814020 | 48814435 | 48814603 | 48824603 | 48127567 | 48137567 | 687 | 7.6 | 51640480 | 51640980 | 3508 | 43470001 | 43470181 | -4662 |
| chr2 | 48814020 | 48814435 | 48812406 | 48822406 | 48146633 | 48156633 | 666 | 5.1 | 51640480 | 51640980 | 3489 | 43470001 | 43470181 | -4682 |
| chr2 | 48814020 | 48814435 | 48810112 | 48820112 | 48182838 | 48192838 | 627 | 7.0 | 51640480 | 51640980 | 3453 | 43470001 | 43470181 | -4718 |
| chr2 | 48814020 | 48814435 | 48812880 | 48822880 | 48242000 | 48252000 | 571 | 6.9 | 51640480 | 51640980 | 3394 | 43470001 | 43470181 | -4777 |
| chr2 | 48814020 | 48814435 | 48813757 | 48823757 | 48310166 | 48320166 | 504 | 6.3 | 51640480 | 51640980 | 3326 | 43470001 | 43470181 | -4845 |
| chr2 | 48814020 | 48814435 | 48815220 | 48825220 | 48324006 | 48334006 | 491 | 29.0 | 51640480 | 51640980 | 3312 | 43470001 | 43470181 | -4859 |
| chr2 | 48814020 | 48814435 | 48812499 | 48822499 | 48394651 | 48404651 | 418 | 5.0 | 51640480 | 51640980 | 3241 | 43470001 | 43470181 | -4930 |
| chr2 | 48814020 | 48814435 | 48808486 | 48818486 | 48412182 | 48422182 | 396 | 5.6 | 51640480 | 51640980 | 3224 | 43470001 | 43470181 | -4947 |
| chr2 | 48814020 | 48814435 | 48814014 | 48824014 | 48462502 | 48472502 | 352 | 7.6 | 51640480 | 51640980 | 3173 | 43470001 | 43470181 | -4997 |
| chr2 | 65238470 | 65238972 | 65228758 | 65238758 | 64916063 | 64926063 | 313 | 5.9 | 66784943 | 66785443 | 1864 | 63861895 | 63862148 | -1059 |
| chr2 | 65239101 | 65239417 | 65228758 | 65238758 | 64916063 | 64926063 | 313 | 5.9 | 66784943 | 66785443 | 1864 | 63861895 | 63862148 | -1059 |
| chr2 | 65238470 | 65238972 | 65231829 | 65241829 | 64928528 | 64938528 | 303 | 5.6 | 66784943 | 66785443 | 1852 | 63861895 | 63862148 | -1072 |
| chr2 | 65239101 | 65239417 | 65231829 | 65241829 | 64928528 | 64938528 | 303 | 5.6 | 66784943 | 66785443 | 1852 | 63861895 | 63862148 | -1072 |
| chr2 | 65238470 | 65238972 | 65228213 | 65238213 | 64957110 | 64967110 | 271 | 10.6 | 66784943 | 66785443 | 1823 | 63861895 | 63862148 | -1100 |
| chr2 | 65239101 | 65239417 | 65228213 | 65238213 | 64957110 | 64967110 | 271 | 10.6 | 66784943 | 66785443 | 1823 | 63861895 | 63862148 | -1100 |
| chr2 | 65238470 | 65238972 | 65230795 | 65240795 | 64974492 | 64984492 | 256 | 5.2 | 66784943 | 66785443 | 1806 | 63861895 | 63862148 | -1117 |
| chr2 | 65239101 | 65239417 | 65230795 | 65240795 | 64974492 | 64984492 | 256 | 5.2 | 66784943 | 66785443 | 1806 | 63861895 | 63862148 | -1117 |
| chr2 | 65238470 | 65238972 | 65231808 | 65241808 | 65005751 | 65015751 | 226 | 6.8 | 66784943 | 66785443 | 1774 | 63861895 | 63862148 | -1149 |
| chr2 | 65239101 | 65239417 | 65231808 | 65241808 | 65005751 | 65015751 | 226 | 6.8 | 66784943 | 66785443 | 1774 | 63861895 | 63862148 | -1149 |
| chr2 | 65238470 | 65238972 | 65238693 | 65248693 | 65019320 | 65029320 | 219 | 16.6 | 66784943 | 66785443 | 1761 | 63861895 | 63862148 | -1162 |

| | | | | | | | | | | | | | | |
|---|---|---|---|---|---|---|---|---|---|---|---|---|---|---|
| chr2 | 65239101 | 65239417 | 65238693 | 65248693 | 65019320 | 65029320 | 219 | 16.6 | 66784943 | 66785443 | 1761 | 63861895 | 63862148 | -1162 |
| chr2 | 65238470 | 65238972 | 65229732 | 65239732 | 65115118 | 65125118 | 115 | 21.9 | 66784943 | 66785443 | 1665 | 63861895 | 63862148 | -1258 |
| chr2 | 65239101 | 65239417 | 65229732 | 65239732 | 65115118 | 65125118 | 115 | 21.9 | 66784943 | 66785443 | 1665 | 63861895 | 63862148 | -1258 |
| chr2 | 65238470 | 65238972 | 65229447 | 65239447 | 65133872 | 65143872 | 96 | 10.9 | 66784943 | 66785443 | 1646 | 63861895 | 63862148 | -1277 |
| chr2 | 65239101 | 65239417 | 65229447 | 65239447 | 65133872 | 65143872 | 96 | 10.9 | 66784943 | 66785443 | 1646 | 63861895 | 63862148 | -1277 |
| chr2 | 65238470 | 65238972 | 65228349 | 65238349 | 65144255 | 65154255 | 84 | 14.9 | 66784943 | 66785443 | 1636 | 63861895 | 63862148 | -1287 |
| chr2 | 65239101 | 65239417 | 65228349 | 65238349 | 65144255 | 65154255 | 84 | 14.9 | 66784943 | 66785443 | 1636 | 63861895 | 63862148 | -1287 |
| chr2 | 65238470 | 65238972 | 65229169 | 65239169 | 65166552 | 65176552 | 63 | 30.3 | 66784943 | 66785443 | 1614 | 63861895 | 63862148 | -1310 |
| chr2 | 65239101 | 65239417 | 65229169 | 65239169 | 65166552 | 65176552 | 63 | 30.3 | 66784943 | 66785443 | 1614 | 63861895 | 63862148 | -1310 |
| chr2 | 65238470 | 65238972 | 65229361 | 65239361 | 65193867 | 65203867 | 35 | 10.2 | 66784943 | 66785443 | 1586 | 63861895 | 63862148 | -1337 |
| chr2 | 65239101 | 65239417 | 65229361 | 65239361 | 65193867 | 65203867 | 35 | 10.2 | 66784943 | 66785443 | 1586 | 63861895 | 63862148 | -1337 |
| chr2 | 65238470 | 65238972 | 65229293 | 65239293 | 65207927 | 65217927 | 21 | 8.1 | 66784943 | 66785443 | 1572 | 63861895 | 63862148 | -1351 |
| chr2 | 65239101 | 65239417 | 65229293 | 65239293 | 65207927 | 65217927 | 21 | 8.1 | 66784943 | 66785443 | 1572 | 63861895 | 63862148 | -1351 |
| chr2 | 65238470 | 65238972 | 65228790 | 65238790 | 65228267 | 65238267 | 1 | 15.6 | 66784943 | 66785443 | 1552 | 63861895 | 63862148 | -1371 |
| chr2 | 65239101 | 65239417 | 65228790 | 65238790 | 65228267 | 65238267 | 1 | 15.6 | 66784943 | 66785443 | 1552 | 63861895 | 63862148 | -1371 |
| chr2 | 122781608 | 122781964 | 122775568 | 122785568 | 122166600 | 122176600 | 609 | 5.1 | 119126264 | 119126764 | -3045 | 122378630 | 122378837 | 207 |
| chr2 | 122781608 | 122781964 | 122782213 | 122792213 | 122368076 | 122378076 | 414 | 8.8 | 119126264 | 119126764 | -3247 | 122378630 | 122378837 | 6 |
| chr2 | 122781608 | 122781964 | 122772610 | 122782610 | 122771422 | 122781422 | 1 | 26.1 | 125660764 | 125661264 | 2885 | 122378630 | 122378837 | -398 |
| chr2 | 125123566 | 125123755 | 125123458 | 125133458 | 124111623 | 124121623 | 1012 | 6.9 | 125660764 | 125661264 | 1544 | 122378630 | 122378837 | -1738 |
| chr2 | 125123566 | 125123755 | 125122718 | 125132718 | 124237458 | 124247458 | 885 | 5.2 | 125660764 | 125661264 | 1419 | 122378630 | 122378837 | -1864 |
| chr2 | 125123566 | 125123755 | 125120518 | 125130518 | 124533570 | 124543570 | 587 | 5.1 | 125660764 | 125661264 | 1122 | 122378630 | 122378837 | -2160 |
| chr2 | 125123566 | 125123755 | 125123712 | 125133712 | 124926935 | 124936935 | 197 | 4.9 | 125660764 | 125661264 | 729 | 126951102 | 126951344 | 2019 |
| chr2 | 125123566 | 125123755 | 125122913 | 125132913 | 124943731 | 124953731 | 179 | 4.7 | 125660764 | 125661264 | 712 | 126951102 | 126951344 | 2002 |
| chr2 | 125123566 | 125123755 | 125122884 | 125132884 | 125021014 | 125031014 | 102 | 5.8 | 125660764 | 125661264 | 635 | 126951102 | 126951344 | 1925 |
| chr2 | 148040102 | 148040610 | 148029159 | 148039159 | 147938241 | 147948241 | 91 | 19.3 | 146546264 | 146546764 | -1397 | 147942625 | 147942972 | 0 |
| chr2 | 148040102 | 148040610 | 148030797 | 148040797 | 148029529 | 148039529 | 1 | 18.0 | 146546264 | 146546764 | -1488 | 147944182 | 147944694 | -90 |
| chr2 | 151996373 | 151996544 | 151991389 | 152001389 | 151852272 | 151862272 | 139 | 7.2 | 151865764 | 151866264 | 9 | 151994880 | 151995046 | 138 |
| chr2 | 151996373 | 151996544 | 151992389 | 152002389 | 151868260 | 151878260 | 124 | 22.3 | 151865764 | 151866264 | -7 | 151994880 | 151995046 | 122 |
| chr2 | 151996373 | 151996544 | 151991486 | 152001486 | 151911821 | 151921821 | 80 | 8.1 | 151865764 | 151866264 | -51 | 151994880 | 151995046 | 78 |
| chr2 | 151996373 | 151996544 | 151991833 | 152001833 | 151927450 | 151937450 | 64 | 18.2 | 151865764 | 151866264 | -66 | 151994880 | 151995046 | 63 |
| chr2 | 151996373 | 151996544 | 151991594 | 152001594 | 151946910 | 151956910 | 45 | 8.8 | 151865764 | 151866264 | -86 | 151994880 | 151995046 | 43 |
| chr3 | 9610026 | 9610292 | 9608270 | 9618270 | 8970445 | 8980445 | 638 | 5.0 | 5203000 | 5203500 | -3772 | 9947784 | 9947964 | 972 |
| chr3 | 9610026 | 9610292 | 9605301 | 9615301 | 9285553 | 9295553 | 320 | 5.6 | 5203000 | 5203500 | -4087 | 9947784 | 9947964 | 657 |
| chr3 | 9610026 | 9610292 | 9607053 | 9617053 | 9396635 | 9406635 | 210 | 11.8 | 5203000 | 5203500 | -4198 | 9947784 | 9947964 | 546 |
| chr3 | 9610026 | 9610292 | 9606413 | 9616413 | 9481034 | 9491034 | 125 | 12.2 | 5203000 | 5203500 | -4283 | 9947784 | 9947964 | 462 |
| chr3 | 9610026 | 9610292 | 9610928 | 9620928 | 9502720 | 9512720 | 108 | 8.3 | 5203000 | 5203500 | -4304 | 9947784 | 9947964 | 440 |
| chr3 | 9610026 | 9610292 | 9607325 | 9617325 | 9512922 | 9522922 | 94 | 7.6 | 5203000 | 5203500 | -4315 | 9947784 | 9947964 | 430 |
| chr3 | 28697751 | 28698115 | 28691118 | 28701118 | 28099722 | 28109722 | 591 | 5.6 | 27294578 | 27295078 | -810 | 27751707 | 27751920 | -353 |
| chr3 | 28697751 | 28698115 | 28689316 | 28699316 | 28151583 | 28161583 | 538 | 4.7 | 27294578 | 27295078 | -862 | 27899110 | 27899586 | -257 |
| chr3 | 28697751 | 28698115 | 28688711 | 28698711 | 28654138 | 28664138 | 35 | 8.3 | 27294578 | 27295078 | -1364 | 27899110 | 27899586 | -760 |
| chr3 | 28697751 | 28698115 | 28686778 | 28696778 | 28681998 | 28691998 | 5 | 12.4 | 27294578 | 27295078 | -1392 | 27899110 | 27899586 | -788 |
| chr3 | 57425183 | 57425592 | 57422463 | 57432463 | 55351858 | 55361858 | 2071 | 5.6 | 53988578 | 53989078 | -1368 | 54493643 | 54494037 | -863 |
| chr3 | 57425183 | 57425592 | 57423361 | 57433361 | 55814336 | 55824336 | 1609 | 5.6 | 53988578 | 53989078 | -1831 | 54493643 | 54494037 | -1325 |
| chr3 | 57425183 | 57425592 | 57418455 | 57428455 | 57298691 | 57308691 | 120 | 27.0 | 53988578 | 53989078 | -3315 | 58206974 | 58207362 | 903 |
| chr3 | 57425183 | 57425592 | 57421747 | 57431747 | 57344631 | 57354631 | 77 | 7.4 | 53988578 | 53989078 | -3361 | 58206974 | 58207362 | 858 |
| chr3 | 57425183 | 57425592 | 57417044 | 57427044 | 57386811 | 57396811 | 30 | 31.1 | 53988578 | 53989078 | -3403 | 58206974 | 58207362 | 815 |
| chr3 | 65529050 | 65529575 | 65528728 | 65538728 | 65311731 | 65321731 | 217 | 6.1 | 63297078 | 63297578 | -2019 | 64039820 | 64040282 | -1277 |
| chr3 | 65529050 | 65529575 | 65529760 | 65539760 | 65371605 | 65381605 | 158 | 19.7 | 63297078 | 63297578 | -2079 | 64039820 | 64040282 | -1337 |

| | | | | | | | | | | | | | | |
|---|---|---|---|---|---|---|---|---|---|---|---|---|---|---|
| chr3 | 65529050 | 65529575 | 65530348 | 65540348 | 65392107 | 65402107 | 138 | 7.4 | 63297078 | 63297578 | -2100 | 64039820 | 64040282 | -1357 |
| chr3 | 65529050 | 65529575 | 65524091 | 65534091 | 65406708 | 65416708 | 117 | 6.3 | 63297078 | 63297578 | -2114 | 64039820 | 64040282 | -1372 |
| chr3 | 81036684 | 81036990 | 81030070 | 81040070 | 79624102 | 79634102 | 1406 | 5.6 | 79519578 | 79520078 | -109 | 78919496 | 78919788 | -709 |
| chr3 | 81036684 | 81036990 | 81030159 | 81040159 | 79844172 | 79854172 | 1186 | 5.3 | 79519578 | 79520078 | -329 | 78919496 | 78919788 | -930 |
| chr3 | 81036684 | 81036990 | 81036838 | 81046838 | 80067087 | 80077087 | 970 | 7.0 | 79519578 | 79520078 | -552 | 78919496 | 78919788 | -1152 |
| chr3 | 81036684 | 81036990 | 81034491 | 81044491 | 80429426 | 80439426 | 605 | 4.7 | 79519578 | 79520078 | -915 | 78919496 | 78919788 | -1515 |
| chr3 | 81036684 | 81036990 | 81029377 | 81039377 | 80951515 | 80961515 | 78 | 17.2 | 79519578 | 79520078 | -1437 | 82022547 | 82022964 | 1066 |
| chr3 | 81036684 | 81036990 | 81037090 | 81047090 | 81037090 | 81047090 | 0 | 8.8 | 79519578 | 79520078 | -1522 | 82022547 | 82022964 | 981 |
| chr3 | 96268499 | 96268727 | 96257803 | 96267803 | 96085000 | 96095000 | 173 | 6.7 | 96322077 | 96322577 | 232 | 96094280 | 96094557 | 4 |
| chr3 | 96268499 | 96268727 | 96258718 | 96268718 | 96228675 | 96238675 | 30 | 5.1 | 96322077 | 96322577 | 89 | 96094280 | 96094557 | -139 |
| chr3 | 96268499 | 96268727 | 96259941 | 96269941 | 96259941 | 96269941 | 154 | 6.5 | 96322077 | 96322577 | 57 | 96094280 | 96094557 | -171 |
| chr3 | 105801297 | 105801647 | 105799031 | 105809031 | 104986846 | 104996846 | 812 | 5.6 | 104753082 | 104753582 | -239 | 104967870 | 104968267 | -24 |
| chr3 | 105801297 | 105801647 | 105798238 | 105808238 | 105682623 | 105692623 | 116 | 14.5 | 106042582 | 106043082 | 355 | 105058980 | 105059162 | -629 |
| chr3 | 108013289 | 108013838 | 108011242 | 108021242 | 107873784 | 107883784 | 137 | 7.4 | 106042582 | 106043082 | -1836 | 107867181 | 107867395 | -11 |
| chr3 | 108013289 | 108013838 | 108013762 | 108023762 | 107944353 | 107954353 | 69 | 48.4 | 106042582 | 106043082 | -1907 | 107867181 | 107867395 | -82 |
| chr3 | 108013289 | 108013838 | 108006655 | 108016655 | 107969192 | 107979192 | 37 | 34.9 | 106042582 | 106043082 | -1931 | 107867181 | 107867395 | -107 |
| chr3 | 135825071 | 135825300 | 135824934 | 135834934 | 135670461 | 135680461 | 154 | 5.7 | 135493036 | 135493536 | -182 | 135731383 | 135731600 | 56 |
| chr3 | 145758456 | 145758830 | 145749009 | 145759009 | 145200470 | 145210470 | 549 | 5.6 | 145614036 | 145614536 | 409 | 145624846 | 145625310 | 420 |
| chr3 | 145758456 | 145758830 | 145754493 | 145764493 | 145563132 | 145573132 | 191 | 6.7 | 145614036 | 145614536 | 46 | 145624846 | 145625310 | 57 |
| chr3 | 145758456 | 145758830 | 145754513 | 145764513 | 145576775 | 145586775 | 178 | 5.6 | 145614036 | 145614536 | 33 | 145624846 | 145625310 | 43 |
| chr4 | 32247529 | 32247819 | 32247607 | 32257607 | 31965953 | 31975953 | 282 | 7.8 | 33526525 | 33527025 | 1556 | 32153468 | 32153643 | 183 |
| chr4 | 32247529 | 32247819 | 32246489 | 32256489 | 32048114 | 32058114 | 198 | 13.2 | 33526525 | 33527025 | 1474 | 32153468 | 32153643 | 100 |
| chr4 | 32247529 | 32247819 | 32243060 | 32253060 | 32062363 | 32072363 | 181 | 6.9 | 33526525 | 33527025 | 1459 | 32153468 | 32153643 | 86 |
| chr4 | 32247529 | 32247819 | 32248323 | 32258323 | 32080418 | 32090418 | 168 | 10.6 | 33526525 | 33527025 | 1441 | 32153468 | 32153643 | 68 |
| chr4 | 32247529 | 32247819 | 32248626 | 32258626 | 32107433 | 32117433 | 141 | 5.8 | 33526525 | 33527025 | 1414 | 32153468 | 32153643 | 41 |
| chr4 | 32247529 | 32247819 | 32242910 | 32252910 | 32167815 | 32177815 | 75 | 13.9 | 33526525 | 33527025 | 1354 | 32153468 | 32153643 | -19 |
| chr4 | 57916192 | 57916684 | 57909614 | 57919614 | 57504734 | 57514734 | 405 | 9.1 | 55515628 | 55516128 | -1994 | 57370066 | 57370246 | -140 |
| chr4 | 59003007 | 59003198 | 59003420 | 59013420 | 57956207 | 57966207 | 1047 | 5.6 | 55515628 | 55516128 | -2445 | 57370066 | 57370246 | -591 |
| chr4 | 59003007 | 59003198 | 58996639 | 59006639 | 58824332 | 58834332 | 172 | 4.9 | 55515628 | 55516128 | -3313 | 58913583 | 58913874 | 84 |
| chr4 | 59003007 | 59003198 | 58993319 | 59003319 | 58835524 | 58845524 | 158 | 5.8 | 55515628 | 55516128 | -3325 | 58913583 | 58913874 | 73 |
| chr4 | 59003007 | 59003198 | 59003395 | 59013395 | 58838422 | 58848422 | 165 | 8.3 | 55515628 | 55516128 | -3328 | 58913583 | 58913874 | 70 |
| chr4 | 59003007 | 59003198 | 58993790 | 59003790 | 58905917 | 58915917 | 88 | 6.3 | 55515628 | 55516128 | -3395 | 58913583 | 58913874 | 3 |
| chr4 | 82505623 | 82505970 | 82498706 | 82508706 | 80922823 | 80932823 | 1576 | 7.4 | 82769596 | 82770096 | 1842 | 80874185 | 80874441 | -54 |
| chr4 | 82507856 | 82508070 | 82498706 | 82508706 | 80922823 | 80932823 | 1576 | 7.4 | 82769596 | 82770096 | 1842 | 80874185 | 80874441 | -54 |
| chr4 | 82505623 | 82505970 | 82500142 | 82510142 | 80935878 | 80945878 | 1564 | 5.6 | 82769596 | 82770096 | 1829 | 80874185 | 80874441 | -67 |
| chr4 | 82507856 | 82508070 | 82500142 | 82510142 | 80935878 | 80945878 | 1564 | 5.6 | 82769596 | 82770096 | 1829 | 80874185 | 80874441 | -67 |
| chr4 | 82505623 | 82505970 | 82501717 | 82511717 | 81346349 | 81356349 | 1155 | 7.4 | 82769596 | 82770096 | 1418 | 81276365 | 81276584 | -75 |
| chr4 | 82507856 | 82508070 | 82501717 | 82511717 | 81346349 | 81356349 | 1155 | 7.4 | 82769596 | 82770096 | 1418 | 81276365 | 81276584 | -75 |
| chr4 | 82505623 | 82505970 | 82506066 | 82516066 | 81364583 | 81374583 | 1141 | 5.6 | 82769596 | 82770096 | 1400 | 81276365 | 81276584 | -93 |
| chr4 | 82507856 | 82508070 | 82506066 | 82516066 | 81364583 | 81374583 | 1141 | 5.6 | 82769596 | 82770096 | 1400 | 81276365 | 81276584 | -93 |
| chr4 | 82505623 | 82505970 | 82494658 | 82504658 | 81377103 | 81387103 | 1118 | 6.6 | 82769596 | 82770096 | 1388 | 81519221 | 81519504 | 137 |
| chr4 | 82505623 | 82505970 | 82499017 | 82509017 | 81437236 | 81447236 | 1062 | 5.6 | 82769596 | 82770096 | 1328 | 81519221 | 81519504 | 77 |
| chr4 | 82507856 | 82508070 | 82499017 | 82509017 | 81437236 | 81447236 | 1062 | 5.6 | 82769596 | 82770096 | 1328 | 81519221 | 81519504 | 77 |
| chr4 | 82505623 | 82505970 | 82500380 | 82510380 | 81499945 | 81509945 | 1000 | 13.7 | 82769596 | 82770096 | 1265 | 81519221 | 81519504 | 14 |
| chr4 | 82507856 | 82508070 | 82500380 | 82510380 | 81499945 | 81509945 | 1000 | 13.7 | 82769596 | 82770096 | 1265 | 81519221 | 81519504 | 14 |
| chr4 | 82505623 | 82505970 | 82501315 | 82511315 | 81574013 | 81584013 | 927 | 5.6 | 82769596 | 82770096 | 1191 | 81519221 | 81519504 | -60 |
| chr4 | 82507856 | 82508070 | 82501315 | 82511315 | 81574013 | 81584013 | 927 | 5.6 | 82769596 | 82770096 | 1191 | 81519221 | 81519504 | -60 |
| chr4 | 82505623 | 82505970 | 82500070 | 82510070 | 81587276 | 81597276 | 913 | 5.6 | 82769596 | 82770096 | 1178 | 81519221 | 81519504 | -73 |

| | | | | | | | | | | | | | | |
|---|---|---|---|---|---|---|---|---|---|---|---|---|---|---|
| chr4 | 82507856 | 82508070 | 82500070 | 82510070 | 81587276 | 81597276 | 913 | 5.6 | 82769596 | 82770096 | 1178 | 81519221 | 81519504 | -73 |
| chr4 | 82505623 | 82505970 | 82494622 | 82504622 | 81670420 | 81680420 | 824 | 14.8 | 82769596 | 82770096 | 1094 | 81789319 | 81789561 | 114 |
| chr4 | 82507856 | 82508070 | 82507531 | 82517531 | 81722458 | 81732458 | 785 | 9.4 | 82769596 | 82770096 | 1042 | 81789319 | 81789561 | 62 |
| chr4 | 82507856 | 82508070 | 82507323 | 82517323 | 81925668 | 81935668 | 582 | 10.8 | 82769596 | 82770096 | 839 | 81789319 | 81789561 | -141 |
| chr4 | 82505623 | 82505970 | 82495320 | 82505320 | 81993791 | 82003791 | 502 | 8.5 | 82769596 | 82770096 | 771 | 81789319 | 81789561 | -209 |
| chr4 | 82505623 | 82505970 | 82506058 | 82516058 | 82067926 | 82077926 | 438 | 7.0 | 82769596 | 82770096 | 697 | 81789319 | 81789561 | -283 |
| chr4 | 82507856 | 82508070 | 82506058 | 82516058 | 82067926 | 82077926 | 438 | 7.0 | 82769596 | 82770096 | 697 | 81789319 | 81789561 | -283 |
| chr4 | 82505623 | 82505970 | 82503632 | 82513632 | 82092983 | 82102983 | 411 | 5.6 | 82769596 | 82770096 | 672 | 81789319 | 81789561 | -309 |
| chr4 | 82507856 | 82508070 | 82503632 | 82513632 | 82092983 | 82102983 | 411 | 5.6 | 82769596 | 82770096 | 672 | 81789319 | 81789561 | -309 |
| chr4 | 82505623 | 82505970 | 82505758 | 82515758 | 82256205 | 82266205 | 250 | 10.0 | 82769596 | 82770096 | 509 | 81789319 | 81789561 | -472 |
| chr4 | 82507856 | 82508070 | 82505758 | 82515758 | 82256205 | 82266205 | 250 | 10.0 | 82769596 | 82770096 | 509 | 81789319 | 81789561 | -472 |
| chr4 | 82505623 | 82505970 | 82506317 | 82516317 | 82310831 | 82320831 | 195 | 17.9 | 82769596 | 82770096 | 454 | 81789319 | 81789561 | -526 |
| chr4 | 82507856 | 82508070 | 82506317 | 82516317 | 82310831 | 82320831 | 195 | 17.9 | 82769596 | 82770096 | 454 | 81789319 | 81789561 | -526 |
| chr4 | 82505623 | 82505970 | 82500488 | 82510488 | 82333666 | 82343666 | 167 | 20.3 | 82769596 | 82770096 | 431 | 81789319 | 81789561 | -549 |
| chr4 | 82507856 | 82508070 | 82500488 | 82510488 | 82333666 | 82343666 | 167 | 20.3 | 82769596 | 82770096 | 431 | 81789319 | 81789561 | -549 |
| chr4 | 82505623 | 82505970 | 82506786 | 82516786 | 82378128 | 82388128 | 129 | 13.0 | 82769596 | 82770096 | 387 | 81789319 | 81789561 | -594 |
| chr4 | 82507856 | 82508070 | 82506786 | 82516786 | 82378128 | 82388128 | 129 | 13.0 | 82769596 | 82770096 | 387 | 81789319 | 81789561 | -594 |
| chr4 | 82505623 | 82505970 | 82500645 | 82510645 | 82406176 | 82416176 | 94 | 25.1 | 82769596 | 82770096 | 359 | 81789319 | 81789561 | -622 |
| chr4 | 82507856 | 82508070 | 82500645 | 82510645 | 82406176 | 82416176 | 94 | 25.1 | 82769596 | 82770096 | 359 | 81789319 | 81789561 | -622 |
| chr4 | 99120860 | 99121120 | 99116090 | 99126090 | 98467130 | 98477130 | 649 | 4.7 | 97900309 | 97900809 | -572 | 101400076 | 101400455 | 2928 |
| chr4 | 99120860 | 99121120 | 99110611 | 99120611 | 98924398 | 98934398 | 186 | 5.4 | 99560809 | 99561309 | 632 | 101400076 | 101400455 | 2471 |
| chr4 | 99120860 | 99121120 | 99110750 | 99120750 | 98938440 | 98948440 | 172 | 6.5 | 99560809 | 99561309 | 618 | 101400076 | 101400455 | 2457 |
| chr4 | 99120860 | 99121120 | 99117494 | 99127494 | 98995101 | 99005101 | 122 | 5.2 | 99560809 | 99561309 | 561 | 101400076 | 101400455 | 2400 |
| chr4 | 99120860 | 99121120 | 99112049 | 99122049 | 99014867 | 99024867 | 97 | 5.5 | 99560809 | 99561309 | 541 | 101400076 | 101400455 | 2380 |
| chr4 | 99120860 | 99121120 | 99115830 | 99125830 | 99077382 | 99087382 | 38 | 5.8 | 99560809 | 99561309 | 479 | 101400076 | 101400455 | 2318 |
| chr4 | 108459573 | 108460023 | 108457173 | 108467173 | 107909467 | 107919467 | 548 | 5.6 | 107442895 | 107443395 | -471 | 107900290 | 107900506 | -14 |
| chr4 | 108459573 | 108460023 | 108456779 | 108466779 | 107937051 | 107947051 | 520 | 5.6 | 107442895 | 107443395 | -499 | 107900290 | 107900506 | -42 |
| chr4 | 108459573 | 108460023 | 108454196 | 108464196 | 108153379 | 108163379 | 301 | 7.0 | 108890395 | 108890895 | 732 | 108225312 | 108225787 | 67 |
| chr4 | 108459573 | 108460023 | 108457607 | 108467607 | 108291118 | 108301118 | 166 | 6.5 | 108890395 | 108890895 | 595 | 108349144 | 108349382 | 53 |
| chr4 | 108459573 | 108460023 | 108455185 | 108465185 | 108322762 | 108332762 | 132 | 15.8 | 108890395 | 108890895 | 563 | 108349144 | 108349382 | 22 |
| chr4 | 108459573 | 108460023 | 108455512 | 108465512 | 108341455 | 108351455 | 114 | 20.7 | 108890395 | 108890895 | 544 | 108349144 | 108349382 | 3 |
| chr4 | 108459573 | 108460023 | 108459022 | 108469022 | 108395353 | 108405353 | 64 | 27.9 | 108890395 | 108890895 | 490 | 108362579 | 108362962 | -38 |
| chr4 | 108459573 | 108460023 | 108454572 | 108464572 | 108420407 | 108430407 | 34 | 8.2 | 108890395 | 108890895 | 465 | 108456125 | 108456424 | 31 |
| chr4 | 130663132 | 130663670 | 130653553 | 130663553 | 130309535 | 130319535 | 344 | 5.4 | 130845085 | 130845585 | 531 | 130198999 | 130199187 | -115 |
| chr4 | 138966900 | 138968000 | 138959299 | 138969299 | 136548619 | 136558619 | 2411 | 5.6 | 136981585 | 136982085 | 428 | 136160215 | 136160417 | -393 |
| chr4 | 138966900 | 138968000 | 138958228 | 138968228 | 138444181 | 138454181 | 514 | 5.2 | 138664585 | 138665085 | 216 | 138379198 | 138379425 | -70 |
| chr4 | 138966900 | 138968000 | 138957911 | 138967911 | 138919329 | 138929329 | 39 | 7.2 | 138958585 | 138959085 | 35 | 138922141 | 138922365 | -2 |
| chr4 | 138966900 | 138968000 | 138957866 | 138967866 | 138957557 | 138967557 | 0 | 31.6 | 138958585 | 138959085 | -4 | 138969205 | 138969637 | 7 |
| chr5 | 65763083 | 65763436 | 65758232 | 65768232 | 65383266 | 65393266 | 375 | 5.6 | 66527261 | 66527761 | 1139 | 65724355 | 65725015 | 336 |
| chr5 | 65763083 | 65763436 | 65757579 | 65767579 | 65518033 | 65528033 | 240 | 9.4 | 66527261 | 66527761 | 1004 | 65724355 | 65725015 | 202 |
| chr5 | 65763083 | 65763436 | 65757010 | 65767010 | 65535365 | 65545365 | 222 | 6.6 | 66527261 | 66527761 | 987 | 65724355 | 65725015 | 184 |
| chr5 | 65763083 | 65763436 | 65752533 | 65762533 | 65715385 | 65725385 | 37 | 11.7 | 66527261 | 66527761 | 807 | 65724355 | 65725015 | 4 |
| chr5 | 66151323 | 66151672 | 66146416 | 66156416 | 66028287 | 66038287 | 118 | 15.6 | 66527261 | 66527761 | 494 | 66239366 | 66239665 | 206 |
| chr5 | 66098169 | 66098540 | 66097779 | 66107779 | 66030235 | 66040235 | 68 | 8.5 | 66527261 | 66527761 | 492 | 66239366 | 66239665 | 204 |
| chr5 | 66098169 | 66098540 | 66097367 | 66107367 | 66049512 | 66059512 | 48 | 8.8 | 66527261 | 66527761 | 473 | 66239366 | 66239665 | 185 |
| chr5 | 66151323 | 66151672 | 66151961 | 66161961 | 66052763 | 66062763 | 99 | 11.5 | 66527261 | 66527761 | 470 | 66239366 | 66239665 | 182 |
| chr5 | 66098169 | 66098540 | 66088802 | 66098802 | 66072294 | 66082294 | 17 | 7.9 | 66527261 | 66527761 | 450 | 66239366 | 66239665 | 162 |
| chr5 | 66151323 | 66151672 | 66147520 | 66157520 | 66074466 | 66084466 | 73 | 14.1 | 66527261 | 66527761 | 448 | 66239366 | 66239665 | 160 |

| | | | | | | | | | | | | | | |
|---|---|---|---|---|---|---|---|---|---|---|---|---|---|---|
| chr5 | 90560701 | 90561139 | 90554115 | 90564115 | 90322877 | 90332877 | 231 | 5.7 | 92320974 | 92321474 | 1993 | 90439171 | 90439486 | 111 |
| chr5 | 90560701 | 90561139 | 90554725 | 90564725 | 90401417 | 90411417 | 153 | 8.1 | 92320974 | 92321474 | 1915 | 90439171 | 90439486 | 33 |
| chr5 | 90560701 | 90561139 | 90560003 | 90570003 | 90414897 | 90424897 | 145 | 12.1 | 92320974 | 92321474 | 1901 | 90439171 | 90439486 | 19 |
| chr5 | 90560701 | 90561139 | 90553772 | 90563772 | 90479341 | 90489341 | 74 | 5.9 | 92320974 | 92321474 | 1837 | 90476631 | 90478042 | -7 |
| chr5 | 102481265 | 102481627 | 102472945 | 102482945 | 102255388 | 102265388 | 218 | 5.5 | 102102981 | 102103481 | -157 | 103927511 | 103927872 | 1667 |
| chr5 | 102481265 | 102481627 | 102479536 | 102489536 | 102319408 | 102329408 | 160 | 4.8 | 102102981 | 102103481 | -221 | 103927511 | 103927872 | 1603 |
| chr5 | 102481265 | 102481627 | 102479699 | 102489699 | 102421674 | 102431674 | 58 | 8.1 | 102102981 | 102103481 | -323 | 103927511 | 103927872 | 1501 |
| chr5 | 102481265 | 102481627 | 102471327 | 102481327 | 102432976 | 102442976 | 38 | 5.6 | 102102981 | 102103481 | -335 | 103927511 | 103927872 | 1490 |
| chr5 | 102481265 | 102481627 | 102471064 | 102481064 | 102469186 | 102479186 | 2 | 8.7 | 102102981 | 102103481 | -371 | 103927511 | 103927872 | 1454 |
| chr5 | 139389705 | 139390033 | 139389778 | 139399778 | 139337605 | 139347605 | 52 | 5.4 | 139377546 | 139378046 | 35 | 139331634 | 139331978 | -11 |
| chr5 | 139389705 | 139390033 | 139385929 | 139395929 | 139365111 | 139375111 | 21 | 10.5 | 139377546 | 139378046 | 8 | 139358688 | 139359076 | -11 |
| chr5 | 139389705 | 139390033 | 139384840 | 139394840 | 139382371 | 139392371 | 2 | 5.0 | 139377546 | 139378046 | -10 | 139358688 | 139359076 | -28 |
| chr6 | 37530036 | 37530381 | 37527803 | 37537803 | 37076848 | 37086848 | 451 | 5.8 | 38066000 | 38066500 | 984 | 37448049 | 37448230 | 366 |
| chr6 | 37530036 | 37530381 | 37522922 | 37532922 | 37468135 | 37478135 | 55 | 15.0 | 38066000 | 38066500 | 593 | 37448049 | 37448230 | -25 |
| chr6 | 37530036 | 37530381 | 37527571 | 37537571 | 37490420 | 37500420 | 37 | 8.4 | 38066000 | 38066500 | 571 | 37448049 | 37448230 | -47 |
| chr6 | 37530036 | 37530381 | 37528198 | 37538198 | 37504350 | 37514350 | 24 | 14.8 | 38066000 | 38066500 | 557 | 37448049 | 37448230 | -61 |
| chr6 | 37530036 | 37530381 | 37528389 | 37538389 | 37525425 | 37535425 | 3 | 7.3 | 38066000 | 38066500 | 536 | 37592486 | 37592889 | 62 |
| chr6 | 82041470 | 82041995 | 82034924 | 82044924 | 81973009 | 81983009 | 62 | 24.2 | 79539006 | 79539506 | -2439 | 82012493 | 82012687 | 35 |
| chr6 | 82041470 | 82041995 | 82033647 | 82043647 | 81985557 | 81995557 | 48 | 35.4 | 79539006 | 79539506 | -2451 | 82012493 | 82012687 | 22 |
| chr6 | 82052145 | 82052406 | 82046252 | 82056252 | 82012779 | 82022779 | 33 | 5.0 | 79539006 | 79539506 | -2479 | 82012493 | 82012687 | -5 |
| chr6 | 82041470 | 82041995 | 82040032 | 82050032 | 82031523 | 82041523 | 9 | 5.0 | 79539006 | 79539506 | -2497 | 82039045 | 82039630 | 3 |
| chr6 | 88742655 | 88743083 | 88738594 | 88748594 | 88523628 | 88533628 | 215 | 5.3 | 87424006 | 87424506 | -1104 | 88831573 | 88831756 | 303 |
| chr6 | 88742655 | 88743083 | 88743237 | 88753237 | 88574455 | 88584455 | 169 | 4.8 | 87424006 | 87424506 | -1155 | 88831573 | 88831756 | 252 |
| chr6 | 88742655 | 88743083 | 88739074 | 88749074 | 88602461 | 88612461 | 137 | 9.8 | 87424006 | 87424506 | -1183 | 88831573 | 88831756 | 224 |
| chr6 | 88742655 | 88743083 | 88743060 | 88753060 | 88616842 | 88626842 | 126 | 18.0 | 87424006 | 87424506 | -1198 | 88831573 | 88831756 | 210 |
| chr6 | 88742655 | 88743083 | 88737911 | 88747911 | 88633029 | 88643029 | 105 | 11.6 | 87424006 | 87424506 | -1214 | 88831573 | 88831756 | 194 |
| chr6 | 88742655 | 88743083 | 88742743 | 88752743 | 88677961 | 88687961 | 65 | 5.6 | 87424006 | 87424506 | -1259 | 88831573 | 88831756 | 149 |
| chr6 | 95718723 | 95718972 | 95711008 | 95721008 | 94645184 | 94655184 | 1066 | 5.6 | 94542006 | 94542506 | -108 | 94701087 | 94701330 | 51 |
| chr6 | 95718723 | 95718972 | 95717902 | 95727902 | 94687433 | 94697433 | 1030 | 5.2 | 94542006 | 94542506 | -150 | 94701087 | 94701330 | 9 |
| chr6 | 95718723 | 95718972 | 95717607 | 95727607 | 95114750 | 95124750 | 603 | 9.7 | 94542006 | 94542506 | -577 | 94999088 | 94999352 | -121 |
| chr6 | 95718723 | 95718972 | 95714191 | 95724191 | 95241900 | 95251900 | 472 | 5.9 | 94542006 | 94542506 | -705 | 94999088 | 94999352 | -248 |
| chr6 | 95718723 | 95718972 | 95714955 | 95724955 | 95382255 | 95392255 | 333 | 8.0 | 94542006 | 94542506 | -845 | 94999088 | 94999352 | -388 |
| chr6 | 95718723 | 95718972 | 95716679 | 95726679 | 95455484 | 95465484 | 261 | 5.5 | 94542006 | 94542506 | -918 | 94999088 | 94999352 | -461 |
| chr6 | 95718723 | 95718972 | 95711626 | 95721626 | 95507621 | 95517621 | 204 | 14.9 | 94542006 | 94542506 | -970 | 95936422 | 95936903 | 424 |
| chr6 | 95718723 | 95718972 | 95715376 | 95725376 | 95542017 | 95552017 | 173 | 8.5 | 94542006 | 94542506 | -1005 | 95936422 | 95936903 | 390 |
| chr6 | 95718723 | 95718972 | 95713932 | 95723932 | 95592209 | 95602209 | 122 | 17.9 | 94542006 | 94542506 | -1055 | 95936422 | 95936903 | 339 |
| chr6 | 95718723 | 95718972 | 95711580 | 95721580 | 95609238 | 95619238 | 102 | 10.6 | 94542006 | 94542506 | -1072 | 95936422 | 95936903 | 322 |
| chr6 | 124738514 | 124738987 | 124729105 | 124739105 | 124423049 | 124433049 | 306 | 8.6 | 126384482 | 126384982 | 1957 | 124425013 | 124425744 | -3 |
| chr6 | 124738514 | 124738987 | 124731572 | 124741572 | 124637142 | 124647142 | 94 | 15.1 | 126384482 | 126384982 | 1743 | 124719552 | 124719929 | 78 |
| chr6 | 124755912 | 124756152 | 124756991 | 124766991 | 124643411 | 124653411 | 114 | 16.0 | 126384482 | 126384982 | 1736 | 124719552 | 124719929 | 71 |
| chr6 | 124738514 | 124738987 | 124731447 | 124741447 | 124667037 | 124677037 | 64 | 15.6 | 126384482 | 126384982 | 1713 | 124719552 | 124719929 | 48 |
| chr7 | 45706878 | 45707151 | 45704884 | 45714884 | 45222793 | 45232793 | 482 | 5.2 | 45093130 | 45093630 | -134 | 45299859 | 45300028 | 72 |
| chr7 | 45700841 | 45701040 | 45692493 | 45702493 | 45429287 | 45439287 | 263 | 7.8 | 45408130 | 45408630 | -26 | 45299859 | 45300028 | -134 |
| chr7 | 45700841 | 45701040 | 45701067 | 45711067 | 45452753 | 45462753 | 248 | 5.6 | 45408130 | 45408630 | -49 | 45610798 | 45611287 | 153 |
| chr7 | 45706878 | 45707151 | 45701067 | 45711067 | 45452753 | 45462753 | 248 | 5.6 | 45408130 | 45408630 | -49 | 45610798 | 45611287 | 153 |
| chr7 | 45706878 | 45707151 | 45702233 | 45712233 | 45480056 | 45490056 | 222 | 5.5 | 45560630 | 45561130 | 76 | 45610798 | 45611287 | 126 |
| chr7 | 45706878 | 45707151 | 45705739 | 45715739 | 45499837 | 45509837 | 206 | 4.9 | 45560630 | 45561130 | 56 | 45610798 | 45611287 | 106 |
| chr7 | 45700841 | 45701040 | 45691457 | 45701457 | 45604094 | 45614094 | 87 | 5.9 | 45560630 | 45561130 | -48 | 45610798 | 45611287 | 2 |

| | | | | | | | | | | | | | | |
|---|---|---|---|---|---|---|---|---|---|---|---|---|---|---|
| chr7 | 45706878 | 45707151 | 45706495 | 45716495 | 45604867 | 45614867 | 102 | 9.3 | 45560630 | 45561130 | -49 | 45610798 | 45611287 | 1 |
| chr7 | 45700841 | 45701040 | 45691222 | 45701222 | 45639615 | 45649615 | 52 | 6.3 | 45560630 | 45561130 | -84 | 45610798 | 45611287 | -34 |
| chr7 | 73541414 | 73541797 | 73534673 | 73544673 | 73534673 | 73544673 | 0 | 14.5 | 73921114 | 73921614 | 382 | 73577812 | 73578260 | 38 |
| chr7 | 80688909 | 80689256 | 80678083 | 80688083 | 80599008 | 80609008 | 79 | 13.6 | 80160114 | 80160614 | -444 | | | 0 |
| chr7 | 80688909 | 80689256 | 80677957 | 80687957 | 80649091 | 80659091 | 29 | 8.6 | 80160114 | 80160614 | -494 | 80857140 | 80857477 | 203 |
| chr7 | 119525990 | 119526343 | 119516109 | 119526109 | 118786193 | 118796193 | 730 | 7.4 | 119684986 | 119685486 | 894 | 118448577 | 118448960 | -342 |
| chr7 | 119620187 | 119620568 | 119611100 | 119621100 | 119524510 | 119534510 | 87 | 14.7 | 119684986 | 119685486 | 156 | 119748913 | 119749179 | 220 |
| chr7 | 140717218 | 140717925 | 140713948 | 140723948 | 140092877 | 140102877 | 621 | 5.6 | 140050101 | 140050601 | -48 | 140707534 | 140708277 | 610 |
| chr7 | 140720983 | 140721476 | 140713948 | 140723948 | 140092877 | 140102877 | 621 | 5.6 | 140050101 | 140050601 | -48 | 140707534 | 140708277 | 610 |
| chr7 | 140717218 | 140717925 | 140712480 | 140722480 | 140252306 | 140262306 | 460 | 5.0 | 140050101 | 140050601 | -207 | 140707534 | 140708277 | 451 |
| chr7 | 140720983 | 140721476 | 140712480 | 140722480 | 140252306 | 140262306 | 460 | 5.0 | 140050101 | 140050601 | -207 | 140707534 | 140708277 | 451 |
| chr7 | 140717218 | 140717925 | 140712343 | 140722343 | 140305002 | 140315002 | 407 | 6.1 | 140050101 | 140050601 | -260 | 140707534 | 140708277 | 398 |
| chr7 | 140720983 | 140721476 | 140712343 | 140722343 | 140305002 | 140315002 | 407 | 6.1 | 140050101 | 140050601 | -260 | 140707534 | 140708277 | 398 |
| chr7 | 140717218 | 140717925 | 140712544 | 140722544 | 140354647 | 140364647 | 358 | 13.6 | 140050101 | 140050601 | -309 | 140707534 | 140708277 | 348 |
| chr7 | 140720983 | 140721476 | 140712544 | 140722544 | 140354647 | 140364647 | 358 | 13.6 | 140050101 | 140050601 | -309 | 140707534 | 140708277 | 348 |
| chr7 | 140717218 | 140717925 | 140712710 | 140722710 | 140380989 | 140390989 | 332 | 5.0 | 140050101 | 140050601 | -336 | 140707534 | 140708277 | 322 |
| chr7 | 140720983 | 140721476 | 140712710 | 140722710 | 140380989 | 140390989 | 332 | 5.0 | 140050101 | 140050601 | -336 | 140707534 | 140708277 | 322 |
| chr7 | 140717218 | 140717925 | 140712566 | 140722566 | 140554267 | 140564267 | 158 | 5.5 | 140948601 | 140949101 | 390 | 140707534 | 140708277 | 149 |
| chr7 | 140720983 | 140721476 | 140712566 | 140722566 | 140554267 | 140564267 | 158 | 5.5 | 140948601 | 140949101 | 390 | 140707534 | 140708277 | 149 |
| chr8 | 4250688 | 4250867 | 4244454 | 4254454 | 3648330 | 3658330 | 596 | 5.6 | 8443000 | 8443500 | 4790 | 5106857 | 5107148 | 1454 |
| chr8 | 4250688 | 4250867 | 4242587 | 4252587 | 4132043 | 4142043 | 111 | 7.2 | 8443000 | 8443500 | 4306 | 5106857 | 5107148 | 970 |
| chr8 | 22476197 | 22476899 | 22472821 | 22482821 | 22163379 | 22173379 | 309 | 9.5 | 23153528 | 23154028 | 985 | 22400541 | 22400897 | 232 |
| chr8 | 22476197 | 22476899 | 22473032 | 22483032 | 22431494 | 22441494 | 42 | 12.0 | 23153528 | 23154028 | 717 | 22400541 | 22400897 | -36 |
| chr8 | 33929759 | 33930577 | 33931034 | 33941034 | 33743119 | 33753119 | 188 | 5.8 | 34053946 | 34054446 | 306 | 33931553 | 33931831 | 184 |
| chr8 | 33929759 | 33930577 | 33929244 | 33939244 | 33761831 | 33771831 | 167 | 14.4 | 34053946 | 34054446 | 287 | 33931553 | 33931831 | 165 |
| chr8 | 33929759 | 33930577 | 33929370 | 33939370 | 33805859 | 33815859 | 124 | 5.4 | 34053946 | 34054446 | 243 | 33931553 | 33931831 | 121 |
| chr8 | 33929759 | 33930577 | 33929060 | 33939060 | 33822269 | 33832269 | 107 | 8.5 | 34053946 | 34054446 | 227 | 33931553 | 33931831 | 104 |
| chr8 | 33929759 | 33930577 | 33919043 | 33929043 | 33854437 | 33864437 | 65 | 13.7 | 34053946 | 34054446 | 195 | 33931553 | 33931831 | 72 |
| chr8 | 33929759 | 33930577 | 33928289 | 33938289 | 33871619 | 33881619 | 57 | 5.2 | 34053946 | 34054446 | 178 | 33931553 | 33931831 | 55 |
| chr8 | 36457439 | 36458396 | 36458293 | 36468293 | 36153614 | 36163614 | 305 | 5.0 | 36600946 | 36601446 | 443 | 36187081 | 36187339 | 29 |
| chr8 | 41133523 | 41134266 | 41128721 | 41138721 | 40987671 | 40997671 | 141 | 5.4 | 40992646 | 40993146 | 0 | 41135507 | 41135821 | 143 |
| chr8 | 41133523 | 41134266 | 41126457 | 41136457 | 41038075 | 41048075 | 88 | 6.2 | 40992646 | 40993146 | -50 | 41135507 | 41135821 | 93 |
| chr8 | 41133523 | 41134266 | 41126234 | 41136234 | 41094112 | 41104112 | 32 | 5.0 | 40992646 | 40993146 | -106 | 41135507 | 41135821 | 37 |
| chr8 | 41133523 | 41134266 | 41126415 | 41136415 | 41112571 | 41122571 | 14 | 11.9 | 40992646 | 40993146 | -125 | 41135507 | 41135821 | 18 |
| chr8 | 73352394 | 73352992 | 73349084 | 73359084 | 72824513 | 72834513 | 525 | 5.2 | 70452101 | 70452601 | -2377 | 72549608 | 72549863 | -280 |
| chr8 | 73352394 | 73352992 | 73341588 | 73351588 | 72959353 | 72969353 | 382 | 8.7 | 70452101 | 70452601 | -2512 | 72549608 | 72549863 | -415 |
| chr8 | 73352394 | 73352992 | 73353650 | 73363650 | 72989127 | 72999127 | 365 | 9.2 | 70452101 | 70452601 | -2542 | 72549608 | 72549863 | -444 |
| chr8 | 73352394 | 73352992 | 73348003 | 73358003 | 73056199 | 73066199 | 292 | 6.2 | 70452101 | 70452601 | -2609 | 72549608 | 72549863 | -511 |
| chr8 | 73352394 | 73352992 | 73347838 | 73357838 | 73228623 | 73238623 | 119 | 11.6 | 70452101 | 70452601 | -2781 | 72549608 | 72549863 | -684 |
| chr8 | 93131216 | 93131649 | 93124512 | 93134512 | 93054760 | 93064760 | 70 | 6.5 | 94205100 | 94205600 | 1146 | 93233584 | 93234345 | 174 |
| chr8 | 93197926 | 93198455 | 93191768 | 93201768 | 93068500 | 93078500 | 123 | 9.8 | 94205100 | 94205600 | 1132 | 93233584 | 93234345 | 160 |
| chr8 | 93131216 | 93131649 | 93126440 | 93136440 | 93069572 | 93079572 | 57 | 12.5 | 94205100 | 94205600 | 1131 | 93233584 | 93234345 | 159 |
| chr8 | 93197926 | 93198455 | 93189615 | 93199615 | 93120846 | 93130846 | 69 | 7.7 | 94205100 | 94205600 | 1080 | 93233584 | 93234345 | 108 |
| chr8 | 93197926 | 93198455 | 93194637 | 93204637 | 93151060 | 93161060 | 44 | 9.6 | 94205100 | 94205600 | 1049 | 93233584 | 93234345 | 78 |
| chr8 | 93197926 | 93198455 | 93189121 | 93199121 | 93184685 | 93194685 | 4 | 11.5 | 94205100 | 94205600 | 1016 | 93233584 | 93234345 | 44 |
| chr8 | 95488829 | 95489152 | 95481391 | 95491391 | 93998425 | 94008425 | 1483 | 5.6 | 94205100 | 94205600 | 202 | 94013134 | 94013362 | 10 |
| chr8 | 95488829 | 95489152 | 95479521 | 95489521 | 94068298 | 94078298 | 1411 | 5.5 | 94492100 | 94492600 | 419 | 94013134 | 94013362 | -60 |
| chr8 | 95488829 | 95489152 | 95479826 | 95489826 | 95304217 | 95314217 | 176 | 5.4 | 94492100 | 94492600 | -817 | 95377858 | 95378532 | 69 |

| | | | | | | | | | | | | | | |
|---|---|---|---|---|---|---|---|---|---|---|---|---|---|---|
| chr8 | 95488829 | 95489152 | 95479726 | 95489726 | 95328380 | 95338380 | 151 | 18.0 | 96151100 | 96151600 | 818 | 95377858 | 95378532 | 45 |
| chr8 | 95488829 | 95489152 | 95480186 | 95490186 | 95343800 | 95353800 | 136 | 22.1 | 96151100 | 96151600 | 803 | 95377858 | 95378532 | 29 |
| chr8 | 95488829 | 95489152 | 95484721 | 95494721 | 95362083 | 95372083 | 123 | 8.3 | 96151100 | 96151600 | 784 | 95377858 | 95378532 | 11 |
| chr8 | 95488829 | 95489152 | 95481092 | 95491092 | 95379522 | 95389522 | 102 | 5.8 | 96151100 | 96151600 | 767 | 95377858 | 95378532 | -6 |
| chr8 | 95488829 | 95489152 | 95481098 | 95491098 | 95457699 | 95467699 | 23 | 4.9 | 96151100 | 96151600 | 689 | 95377858 | 95378532 | -85 |
| chr8 | 95488829 | 95489152 | 95483159 | 95493159 | 95473389 | 95483389 | 10 | 9.1 | 96151100 | 96151600 | 673 | 95377858 | 95378532 | -100 |
| chr8 | 120634241 | 120634798 | 120633929 | 120643929 | 120166512 | 120176512 | 467 | 8.5 | 120103600 | 120104100 | -68 | 119429595 | 119430005 | -742 |
| chr8 | 120634241 | 120634798 | 120631319 | 120641319 | 120297620 | 120307620 | 334 | 6.0 | 120103600 | 120104100 | -199 | 119429595 | 119430005 | -873 |
| chr8 | 120634241 | 120634798 | 120629597 | 120639597 | 120550592 | 120560592 | 79 | 5.8 | 120103600 | 120104100 | -452 | 119429595 | 119430005 | -1126 |
| chr9 | 37146962 | 37147554 | 37139605 | 37149605 | 35612852 | 35622852 | 1527 | 5.1 | 35007915 | 35008415 | -610 | 35038938 | 35039648 | -579 |
| chr9 | 37146962 | 37147554 | 37138403 | 37148403 | 36910252 | 36920252 | 228 | 6.5 | 37368415 | 37368915 | 453 | 35038938 | 35039648 | -1876 |
| chr9 | 37146962 | 37147554 | 37139227 | 37149227 | 36955426 | 36965426 | 184 | 8.1 | 37368415 | 37368915 | 408 | 35038938 | 35039648 | -1921 |
| chr9 | 37146962 | 37147554 | 37138144 | 37148144 | 36973043 | 36983043 | 165 | 13.4 | 37368415 | 37368915 | 391 | 35038938 | 35039648 | -1939 |
| chr9 | 37146962 | 37147554 | 37137528 | 37147528 | 37012815 | 37022815 | 125 | 10.6 | 37368415 | 37368915 | 351 | 35038938 | 35039648 | -1979 |
| chr9 | 37146962 | 37147554 | 37137577 | 37147577 | 37042863 | 37052863 | 95 | 9.1 | 37368415 | 37368915 | 321 | 35038938 | 35039648 | -2009 |
| chr9 | 37146962 | 37147554 | 37139213 | 37149213 | 37138621 | 37148621 | 1 | 10.8 | 37368415 | 37368915 | 225 | 35038938 | 35039648 | -2104 |
| chr9 | 44477755 | 44478071 | 44477377 | 44487377 | 44103171 | 44113171 | 374 | 5.2 | 44537417 | 44537917 | 429 | 44604000 | 44604656 | 496 |
| chr9 | 44482427 | 44482800 | 44477377 | 44487377 | 44103171 | 44113171 | 374 | 5.2 | 44537417 | 44537917 | 429 | 44604000 | 44604656 | 496 |
| chr9 | 44483685 | 44484190 | 44477377 | 44487377 | 44103171 | 44113171 | 374 | 5.2 | 44537417 | 44537917 | 429 | 44604000 | 44604656 | 496 |
| chr9 | 44482427 | 44482800 | 44482407 | 44492407 | 44319785 | 44329785 | 163 | 5.4 | 44537417 | 44537917 | 213 | 44604000 | 44604656 | 280 |
| chr9 | 44483685 | 44484190 | 44482407 | 44492407 | 44319785 | 44329785 | 163 | 5.4 | 44537417 | 44537917 | 213 | 44604000 | 44604656 | 280 |
| chr9 | 44477755 | 44478071 | 44471922 | 44481922 | 44409856 | 44419856 | 62 | 6.0 | 44537417 | 44537917 | 123 | 44604000 | 44604656 | 189 |
| chr9 | 44482427 | 44482800 | 44471922 | 44481922 | 44409856 | 44419856 | 62 | 6.0 | 44537417 | 44537917 | 123 | 44604000 | 44604656 | 189 |
| chr9 | 44477755 | 44478071 | 44478638 | 44488638 | 44477633 | 44487633 | 1 | 10.0 | 44537417 | 44537917 | 55 | 44604000 | 44604656 | 122 |
| chr9 | 44482427 | 44482800 | 44478638 | 44488638 | 44477633 | 44487633 | 1 | 10.0 | 44537417 | 44537917 | 55 | 44604000 | 44604656 | 122 |
| chr9 | 44483685 | 44484190 | 44478638 | 44488638 | 44477633 | 44487633 | 1 | 10.0 | 44537417 | 44537917 | 55 | 44604000 | 44604656 | 122 |
| chr9 | 65827661 | 65827896 | 65820114 | 65830114 | 65330908 | 65340908 | 489 | 4.7 | 65128693 | 65129193 | -207 | 65290061 | 65290329 | -46 |
| chr9 | 65827661 | 65827896 | 65820189 | 65830189 | 65351724 | 65361724 | 468 | 5.0 | 65128693 | 65129193 | -228 | 65290061 | 65290329 | -67 |
| chr9 | 65827661 | 65827896 | 65817174 | 65827174 | 65474046 | 65484046 | 343 | 5.8 | 65128693 | 65129193 | -350 | 65290061 | 65290329 | -189 |
| chr9 | 65827661 | 65827896 | 65827015 | 65837015 | 65584978 | 65594978 | 242 | 10.1 | 65128693 | 65129193 | -461 | 65290061 | 65290329 | -300 |
| chr9 | 65827661 | 65827896 | 65823018 | 65833018 | 65608044 | 65618044 | 215 | 15.0 | 65128693 | 65129193 | -484 | 65290061 | 65290329 | -323 |
| chr9 | 65827661 | 65827896 | 65828046 | 65838046 | 65630267 | 65640267 | 198 | 17.6 | 65128693 | 65129193 | -506 | 65290061 | 65290329 | -345 |
| chr9 | 65827661 | 65827896 | 65823149 | 65833149 | 65691470 | 65701470 | 132 | 11.3 | 65128693 | 65129193 | -568 | 66084188 | 66084480 | 388 |
| chr9 | 65827661 | 65827896 | 65819855 | 65829855 | 65756951 | 65766951 | 63 | 40.0 | 65128693 | 65129193 | -633 | 66084188 | 66084480 | 322 |
| chr9 | 65827661 | 65827896 | 65826511 | 65836511 | 65785665 | 65795665 | 41 | 13.1 | 65128693 | 65129193 | -662 | 66084188 | 66084480 | 294 |
| chr9 | 65827661 | 65827896 | 65818884 | 65828884 | 65810404 | 65820404 | 8 | 5.9 | 65128693 | 65129193 | -686 | 66084188 | 66084480 | 269 |
| chr9 | 77754387 | 77754633 | 77744037 | 77754037 | 77600961 | 77610961 | 143 | 7.1 | 77496693 | 77497193 | -109 | 78006586 | 78006948 | 401 |
| chr9 | 77754387 | 77754633 | 77743849 | 77753849 | 77684198 | 77694198 | 60 | 32.6 | 77496693 | 77497193 | -192 | 78006586 | 78006948 | 318 |
| chr9 | 107590305 | 107590957 | 107582069 | 107592069 | 107579341 | 107589341 | 3 | 12.1 | 107249669 | 107250169 | -334 | 107644668 | 107645057 | 61 |
| chr9 | 108296405 | 108296953 | 108286844 | 108296844 | 107732769 | 107742769 | 554 | 6.0 | 107249669 | 107250169 | -488 | 107738141 | 107738507 | 1 |
| chr9 | 108296405 | 108296953 | 108297761 | 108307761 | 107850207 | 107860207 | 448 | 5.1 | 107249669 | 107250169 | -605 | 107738141 | 107738507 | -117 |
| chr9 | 108296405 | 108296953 | 108285668 | 108295668 | 107889182 | 107899182 | 396 | 6.2 | 107249669 | 107250169 | -644 | 107738141 | 107738507 | -156 |
| chr9 | 108296405 | 108296953 | 108287112 | 108297112 | 107999072 | 108009072 | 288 | 12.4 | 107249669 | 107250169 | -754 | 108095683 | 108095889 | 92 |
| chr9 | 108296405 | 108296953 | 108293247 | 108303247 | 108014217 | 108024217 | 279 | 5.3 | 108106669 | 108107169 | 88 | 108095683 | 108095889 | 77 |
| chr9 | 108296405 | 108296953 | 108287126 | 108297126 | 108070853 | 108080853 | 216 | 12.7 | 108106669 | 108107169 | 31 | 108095683 | 108095889 | 20 |
| chr9 | 108296405 | 108296953 | 108288583 | 108298583 | 108116552 | 108126552 | 172 | 7.6 | 108106669 | 108107169 | -15 | 108095683 | 108095889 | -26 |
| chr9 | 108296405 | 108296953 | 108286199 | 108296199 | 108133188 | 108143188 | 153 | 8.3 | 108106669 | 108107169 | -31 | 108095683 | 108095889 | -42 |
| chr9 | 108296405 | 108296953 | 108287327 | 108297327 | 108159665 | 108169665 | 128 | 9.4 | 108106669 | 108107169 | -58 | 108095683 | 108095889 | -69 |

| | | | | | | | | | | | | | | |
|---|---|---|---|---|---|---|---|---|---|---|---|---|---|---|
| chr9 | 108296405 | 108296953 | 108296758 | 108306758 | 108215279 | 108225279 | 81 | 8.5 | 108106669 | 108107169 | -113 | 108284505 | 108284704 | 64 |
| chr9 | 108296405 | 108296953 | 108288550 | 108298550 | 108287596 | 108297596 | 1 | 43.2 | 108106669 | 108107169 | -186 | 108284505 | 108284704 | -8 |

only p-RCA within 1,000 bp of area-1 were included; site-1 vs site-2 log-p.
